# Targeting de novo cholesterol synthesis in rhabdomyosarcoma induces cell cycle arrest and triggers apoptosis through ER stress-mediated pathways

**DOI:** 10.1101/2024.12.18.628900

**Authors:** Nebeyu Yosef Gizaw, Kalle Kolari, Kari Alitalo, Riikka Kivelä

## Abstract

Rhabdomyosarcoma (RMS) is the most common soft tissue sarcoma in children, but the outcomes of high-grade RMS patients remain poor, underscoring the critical need for novel therapeutic strategies. Although metabolic pathways in RMS are incompletely characterized, emerging evidence suggests that metabolic adaptations in RMS resemble those in other malignancies. Here, we identify elevated cholesterol biosynthesis driven by the PROX1 transcription factor as a defining feature of RMS. Our findings demonstrate that the cholesterol biosynthesis pathway is essential for RMS cell growth, proliferation, and survival. Blocking this pathway through genetic or pharmacological inhibition of the key cholesterol biosynthesis enzymes significantly impairs RMS cell proliferation, halts cell cycle progression, and triggers apoptosis through activation of endoplasmic reticulum stress pathways. We furthermore validate the critical role of cholesterol biosynthesis in RMS progression in tumor xenograft models, demonstrating that silencing of the DHCR7 gene significantly suppresses tumor growth. Transcriptomic analysis revealed widespread downregulation of cell cycle-related genes following DHCR7 silencing, further supporting the role of cholesterol metabolism in cell cycle regulation. These results highlight the vulnerability of RMS cells to cholesterol biosynthesis inhibition and suggest that targeting this metabolic pathway as a promising therapeutic approach for improving RMS outcomes. Our findings provide a rationale for the development of novel therapies targeted to cholesterol biosynthesis in this aggressive cancer.

**Significance:** This study reveals that targeting cholesterol biosynthesis in rhabdomyosarcoma induces ER-stress, apoptosis and cell cycle arrest, highlighting a potential therapeutic strategy for treating this aggressive pediatric cancer.

## Introduction

Rhabdomyosarcoma (RMS), an aggressive childhood cancer with poor prognosis, accounts for approximately 7% of all pediatric cancers and about half of all soft tissue sarcomas in children (1). RMS tumors exhibit features of skeletal muscle and are believed to originate from mesenchymal cell precursors that have failed to differentiate or have developed abnormally along the skeletal muscle lineage (2). The two major histological subtypes of RMS are embryonal RMS (ERMS) and alveolar RMS (ARMS), each driven by distinct mechanisms (3). The ARMS variant, which generally has a worse prognosis, harbors pathogenic chromosomal translocations in about 80% of cases, resulting in the expression of PAX3-FOXO1 or PAX7-FOXO1 fusion proteins (4). The remaining 20% of ARMS tumors lack these translocations and are classified as fusion-negative, with outcomes similar to those of ERMS. Consequently, RMS is often categorized in clinical settings as either fusion-positive (FP-RMS) or fusion-negative (FN-RMS) (5). Treatment for RMS involves a multidisciplinary approach, including chemotherapy combined with local control through surgical resection and/or radiation therapy (6). However, the five-year event-free survival rate for patients with metastatic disease at diagnosis remains under 30%, and those with relapsed disease face similarly poor outcomes. Over the past three decades, neither the survival rates nor the treatment side effects for high-risk RMS have significantly improved (1). Thus, improving these survival rates hinges on identifying clinically effective agents targeting RMS vulnerabilities.

We aimed to investigate the molecular mechanisms driving RMS tumorigenesis. Our previous work identified that the transcription factor Prospero homeobox 1 (PROX1) is highly expressed in both FN-RMS and FP-RMS. PROX1 is a transcriptional regulator essential for the development of various organs during embryogenesis and also plays a role in oncogenesis (7). PROX1 depletion in RMS cells decreased tumor growth and transformed the RD RMS cells to resemble that of benign mesenchymal stem cells (8).

Through immunohistochemistry and RNA sequencing (RNA-Seq) analysis, we found that PROX1 regulates genes associated with cholesterol biosynthesis in both RMS subtypes. Deregulated lipid and cholesterol homeostasis often drive tumorigenesis and cancer progression (9). Elevated expression of the cholesterol-biosynthesis pathway strongly correlates with poor prognosis in many solid tumors. This link is unsurprising, as cholesterol is crucial for cell membrane structure, and rapidly proliferating cancer cells require high cholesterol levels for membrane biogenesis and other functions (10, 11). The mevalonate-cholesterol biosynthesis pathway is a complex biochemical route that produces several critical end-products, including cholesterol, isoprenoids, dolichol, ubiquinone, and isopentenyl adenine (12). Additionally, reprogramming of lipid metabolism can influence other crucial processes involved in cancer progression, including endoplasmic reticulum (ER) stress response and apoptosis (9). Statins, which inhibit HMGCR, the first rate-limiting enzyme of the MVA pathway, have been demonstrated to reduce growth and increase apoptosis in various cancers both in vitro and in vivo (13, 14). Despite these insights, metabolic reprogramming in RMS, especially that of lipid and cholesterol pathways, is poorly understood and remains an untapped therapeutic target.

In the current study, we show that cholesterol biosynthesis is highly induced in the majority of human RMS cases. Inhibition of cholesterol biosynthesis resulted in decreased cell proliferation and colony formation in vitro. Additionally, cholesterol biosynthesis was found to be essential for the growth of tumor xenografts following the engraftment of human RMS into immune-compromized mice. Silencing of DHCR7, the last enzyme regulating cholesterol biosynthesis, induced ER stress and ER stress-mediated intrinsic apoptosis in RMS cells and tumors. These findings suggest that targeting cholesterol biosynthesis could be a promising therapeutic strategy for RMS.

## Materials and Methods

### Human RMS cell lines and constructs

Human RD cells (CCL-136) were acquired from the American Type Culture Collection (ATCC), while KLHEL1 cells were derived from a biopsy of a 27-year-old male diagnosed with high-grade alveolar rhabdomyosarcoma (ARMS). The biopsy was confirmed positive for FKHR (FOXO1) 13q14 gene fusion by fluorescence in situ hybridization (FISH), and the cells expressed vimentin, MYF4, myosin, and desmin. Ethical approval was obtained from the Helsinki University Hospital’s ethical committee, and written consent was provided by the patient. Before use, all cell lines underwent authentication via short tandem repeat (STR) and single nucleotide polymorphism (SNP) profiling using the ForenSeq DNA Signature Kit (Verogen), and regular testing for mycoplasma contamination was conducted. Cells were cultured in Dulbecco’s Modified Eagle Medium (DMEM) high glucose supplemented with 10% fetal bovine serum (FBS), 2 mM L-glutamine, penicillin (100 U/ml), and streptomycin (100 U/ml), and maintained at 37°C in a humidified 5% CO2 incubator. Lentiviral constructs targeting HMGCR (TRCN0000046448 and TRCN0000046448), DHCR7 (TRCN0000046598, TRCN0000046600, and TRCN0000046602), and a scramble control (SH00200) were sourced from the TRC library. For viral packaging, 4.5 μg of CMVg, 6.5 μg of CMVd8.9, and 9 μg of shRNA plasmid were transfected into 293FT cells (RRID:CVCL_6911) using jetPEI® transfection reagent. RMS cells were transduced with 293FT supernatant containing 4 μg/ml lentivirus for two to three days. Following transduction, cells were subjected to selection with medium containing 3 μg/ml puromycin for 48 hours to establish stable infection.

### RMS cell growth and apoptosis

The growth dynamics and apoptotic responses of HMGCR or DHCR7-silenced cells, alongside their respective controls, were investigated using the IncuCyte live-cell analysis system (Essen Bioscience, Sartorius). Initially, 1000 RD and KLHEL1 cells transduced with lentiviral shHMGCR, shDHCR7, or shSCR were seeded into individual wells of 96-well plates (Corning), with five replicates per construct. Images were acquired every 2 hours throughout the experiment.

To explore the impact of HMGCR and DHCR7 chemical inhibition on cell proliferation, 1000 - 2000 RD and KLHEL1 cells were plated in 96-well plates (Corning) with five replicates per condition. After 24 hours, cells were exposed to varying concentrations of Lovastatin (# HY- N0504, MCE) or AY9944 (DHCR7 inhibitor) (# HY-107420, MCE), and images were captured every 2 hours to monitor growth dynamics using IncuCyte software. Apoptotic events were monitored by assessing Caspase-3/7 activity. For this, RD and KLHEL1 cells were plated at densities of 4000 to 5000 cells/well in 96-well plates and allowed to adhere for 24 hours. Following this, cells were treated with either DMSO or DHCR7 inhibitor, along with IncuCyte Caspase-3/7 Green Apoptosis Assay Reagent (at a 1:1000 dilution). Images were captured every 2 hours over a 3-day period to observe apoptotic changes. For cells transduced with shDHCR7 or shSCR, 2000 to 3000 cells were seeded into wells containing media supplemented with IncuCyte Caspase-3/7 Green Apoptosis Assay Reagent (at a 1:1000 dilution), and images were acquired every 2 hours for 3 days to assess apoptotic responses.

### Mouse xenograft models

Female NOD scid gamma (NSG) mice (NOD.Cg-Prkdc scid Il2rg tm1Wjl/SzJ, RRID:IMSR_JAX:005557) from Jackson Laboratory were utilized for tumor xenograft experiments. NSG mice were anesthetized with isoflurane, and 2 x 10^6 shDHCR7 or shSCR RD cells suspended in 100 μl Matrigel were subcutaneously injected into the right and left scapular areas. Similarly, 2 x 10^6 shDHCR7 or shSCR KLHEL1 cells in 100 μl Matrigel were injected into the same regions. Mice were regularly monitored for palpable tumor formation, and tumor growth was assessed weekly using calipers to measure height (H), width (W), and depth (D), which were then converted into relative tumor volume. At predefined time points, mice were euthanized, and tumors were excised. The volume and mass of the tumors were measured, and the tumors were subjected to histological and gene expression analysis. All animal procedures were approved by the National Animal Experiment Board in Finland (ESAVI/5365/2022).

### Gene expression analysis

RNA isolation and cDNA preparation procedures were conducted as described previously(8). The sequencing data is deposited in Gene Expression Omnibus (RRID:SCR_005012) under accession number GSE279213. Quantitative real-time PCR was performed using a CFX96 Touch Real-Time PCR system. Primer sequences for PCR are provided in Supplementary Table 3.

### Histology and immunohistochemistry

Harvested tumors were fixed in 4% paraformaldehyde (PFA), processed, and embedded in paraffin. Sections of 5 μm thickness were then deparaffinized and subjected to staining with either hematoxylin and eosin or specific antibodies. Heat-induced epitope retrieval using High pH Retrieval solution (DAKO) was performed prior to immunostaining for Ki67 (Dako, clone MIB-1, M7240; 1:200) and CC3 (R and D Systems Cat# AF835, RRID:AB_2243952, 1:200). Signal detection was facilitated by employing Alexa Fluor 488 and 594-conjugated secondary antibodies (Molecular Probes) at a dilution of 1:500. For patient-derived RMS samples, PROX1 (R and D Systems Cat# AF2727, RRID:AB_2170716) staining was carried out utilizing the BenchMark XT automated system on 5 μm thick sections that had undergone antigen retrieval. Imaging was performed using Zeiss Axioplan fluorescent microscope and 3DHISTECH Pannoramic 250 FLASH III digital slide scanner at Genome Biology Unit supported by HiLIFE and the Faculty of Medicine, University of Helsinki, and Biocenter Finland. Image processing and analysis were carried out using ImageJ software (RRID:SCR_002285).

### Immunoblots

Total protein from human RMS cells was extracted using RIPA buffer (50 mM Tris-HCl at pH 7.6, 150 mM NaCl, 1% NP-40, 0.5% DOC, 0.1% SDS) supplemented with a protease and phosphatase inhibitor cocktail (Pierce). Protein concentrations from the samples were measured using the Bicinchoninic Acid (BCA) Protein Assay Kit (Pierce Biotechnology, Rockford, IL, USA). A total of 8-10 µg of protein was denatured at 95°C and loaded onto a 4–20% Criterion TGX Stain-Free protein gel (Bio-Rad Laboratories, catalog no. 5678094). Protein separation was achieved via SDS-PAGE at 270V for 35 minutes. After gel activation, proteins were transferred to low-fluorescence PVDF membranes. Primary antibodies were incubated with the membranes overnight at 4°C. Detection was performed using chemiluminescence with SuperSignal West Femto Maximum Sensitivity Substrate (Pierce Biotechnology), and images were captured using a ChemiDoc MP system (Bio-Rad Laboratories, RRID:SCR_019037). Normalization was done by quantifying total protein in each lane using Stain-Free technology with ImageLab 6.0.1 software (Bio-Rad). The primary antibodies used included: phospho-eIF2α (Ser51) (Cell Signaling Technology Cat# 3398, RRID:AB_2096481, 1:1000), total eIF2α (Cell Signaling Technology Cat# 5324, RRID:AB_10692650, 1:1000), PERK (Cell Signaling Technology Cat# 3192, RRID:AB_2095847, 1:1000), ATF-4 (Cell Signaling Technology Cat# 11815, RRID:AB_2616025, 1:1000), and CHOP (Cell Signaling Technology Cat# 2895, RRID:AB_2089254, 1:1000). HRP-conjugated secondary antibodies (Jackson ImmunoResearch Laboratories, 1:10000) were used.

### RNA sequencing and gene set enrichment analysis

RNA-seq and Gene Set Enrichment Analysis (GSEA) were conducted for shPROX1 KLHEL1 cells and human myoblasts, along with their respective controls, as described previously (8). For shDHCR7 and control RNA-seq, RNA samples were obtained from RD cells, and messenger RNA (mRNA) was purified from total RNA using poly-T oligo-attached magnetic beads. Subsequently, the mRNA was fragmented, and the first strand cDNA was synthesized using random hexamer primers, followed by the synthesis of the second strand cDNA. The library preparation involved end repair, A-tailing, adapter ligation, size selection, amplification, and purification steps. The quality and quantity of the library were assessed using Qubit and real-time PCR for quantification, as well as a bioanalyzer for size distribution detection. The quantified libraries were pooled and sequenced on Illumina platforms based on effective library concentration and data amount criteria. The analysis of the generated FASTQ data was performed using Chipster software (www.chipster.csc.fi),(15) as previously described. Gene Set Enrichment Analysis (GSEA) (16) was also conducted using the GSEA software (RRID:SCR_003199) (http://www.broadinstitute.org/gsea), as described previously.

### Cell Cycle Assay

For cell cycle analysis, RD cells were harvested and washed with cold PBS. The cells were then fixed in 70% cold ethanol andstored at −20 °C overnight. Following fixation, the cells were stained with propidium iodide and subjected to flow cytometry using the NovoCyte Quanteon 4025. Flow cytometry data were analyzed using FlowJo software (RRID:SCR_008520).

### Statistics

Excluding RNA sequencing, statistical analyses were conducted using GraphPad Prism (v8.0) (RRID:SCR_002798). For comparisons between two groups, an unpaired two-tailed t-test was applied, with values expressed as means ± SE. For experiments with more than two groups, one-way ANOVA followed by Tukey’s multiple comparisons test was performed. Statistical significance levels were set at *P < 0.05; **P < 0.01; and ***P < 0.001.

## Results

### Cholesterol biosynthesis is elevated in RMS and regulated by PROX1

Our previous studies have demonstrated that PROX1 plays a regulatory role in the myogenic phenotype in both skeletal myoblasts and rhabdomyosarcoma (RMS) cells (8, 17). Strikingly, increased PROX1 expression induces terminal differentiation in myoblasts, whereas in RMS, it promotes proliferation.

To identify the core transcriptional program induced by PROX1 in RMS cells but not in healthy myoblasts, we conducted RNA-seq analysis on PROX1-silenced RD cells (FN- RMS), KLHEL1 cells (FP-RMS), and human myoblasts (**Fig.1A**). Gene Ontology (GO) analysis of the 436 transcripts downregulated in both RMS cell lines, but not in the healthy myoblasts, revealed that cholesterol biosynthesis was among the most highly enriched pathways **(Fig. 1B**). Also gene-set enrichment analysis (GSEA) clearly indicated that the hallmarks of the cholesterol biosynthesis pathway were significantly affected by PROX1 silencing in both RD and KLHEL1 cells (**Fig.1C**). Importantly, in agreement with the observed effects on gene expression, PROX1 silencing significantly reduced the cellular cholesterol content of the RMS cells (**Fig. 1D**). Further analysis using previously published gene expression data from primary RMS patient samples (GEO: GSE108022) (18) revealed a significant upregulation of genes associated with cholesterol biosynthesis in both fusion-negative and fusion-positive RMS compared to healthy skeletal muscle (**Fig.1E**). Next, we analyzed human RMS tumor samples obtained from the Helsinki Biobank. Immunohistochemistry revealed that elevated expression of PROX1 was associated with increased expression of the cholesterol synthesis enzyme DHCR7; both were more strongly expressed in tumors than the adjacent healthy muscle tissue (**Fig. 1F**).

**Figure 1.**
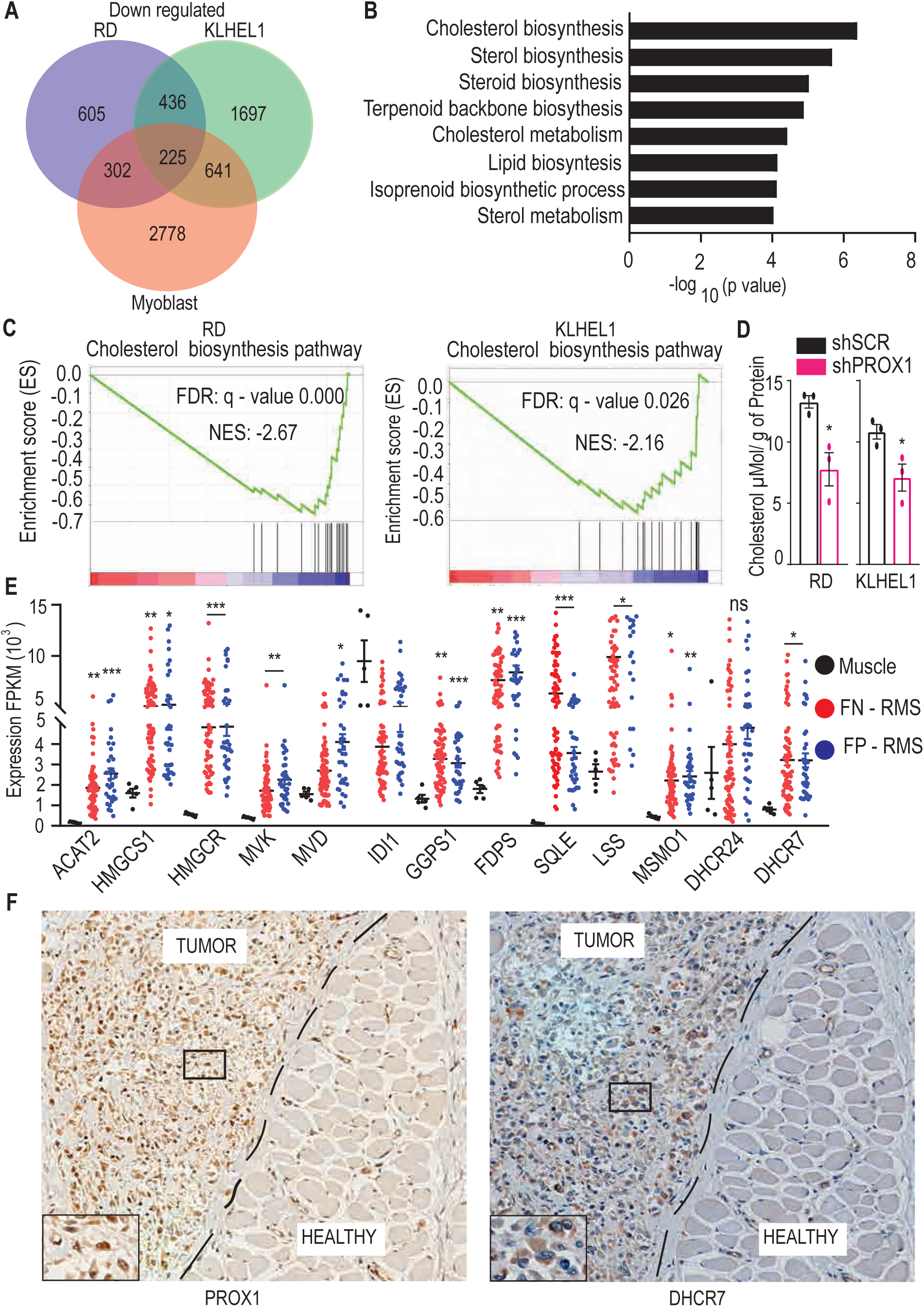
Cholesterol biosynthesis is elevated in rhabdomyosarcoma (RMS). (A) Venn diagram illustrating the number of significantly downregulated genes identified through RNA-seq analysis following PROX1 silencing in RMS cell lines RD and KLHEL1 and compared to healthy human myoblasts. (B) Gene ontology analysis of the 436 common genes downregulated upon PROX1 silencing in both RD and KLHEL1 cells but not in human myoblasts. (C) Gene Set Enrichment Analysis (GSEA) demonstrates the consistent repression of genes involved in cholesterol biosynthesis pathway upon PROX1 silencing in RD cells (left) and in KLHEL1 cells (right). NES denotes normalized enrichment score, and FDR indicates false-discovery rate. (D) Quantification of total cellular cholesterol content in shSCR and shPROX1 treated RD cells (left) and KLHEL1 cells (right) at 3 days post-transduction. The data were analyzed following organic extraction and normalization to protein levels. Data presented as mean ± standard error of the mean (SEM) (*p < 0.05). (E) Normalized expression (FPKM) of genes involved in cholesterol biosynthesis in normal muscles, fusion-negative RMS (FN-RMS), and fusion-positive RMS (FP-RMS) tumor samples in analyzed using a publicly available RNA-seq data set (GSE108022, FDR < 0.05). Each data point represents an individual sample. The horizontal line indicates the mean expression and asterisks denoting significance (*p < 0.05, **p < 0.01, and ***p < 0.001; n.s., not significant, Student’s t-test). (F) Immunohistochemical staining of PROX1 (left) and DHCR7 (right) in clinical RMS tumor samples obtained from the Helsinki Biobank. The tumors exhibit robust nuclear PROX1 expression and cytoplasmic DHCR7 expression compared to adjacent healthy muscle tissue (tumor border marked with a dashed line).

### Inhibition of Cholesterol Biosynthesis Suppresses RMS Cell Growth and Survival

To evaluate the significance of cholesterol biosynthesis in RMS growth, we first silenced HMGCR, a rate-limiting enzyme in the mevalonate–cholesterol biosynthesis pathway, in both the RD (FN-RMS) and KLHEL1 (FP-RMS) cell lines by using two independent lentiviral short hairpin RNAs (shRNAs) (**Supplementary Fig.1A and 1E**). Live cell imaging and analysis revealed that the growth of HMGCR-silenced RD and KLHEL1 cells was markedly inhibited vs shSCR transfected control cells (**Supplementary Fig.1B and 1F**). To assess whether HMGCR knockdown affected the clonogenic potential of RD and KLHEL1 cells, we conducted colony formation assays, which revealed a remarkable reduction in both the number and size of colonies formed upon HMGCR knockdown (**Supplementary Fig.1C, D, and G, H**). In line with the genetic experiments, treatment with the HMG CoA reductase inhibitor lovastatin that slows down cholesterol biosynthesis also significantly reduced RMS cell growth (**Supplementary Fig.1G and 1H**).

The mevalonate–cholesterol biosynthesis pathway is critical not only for cholesterol production but also for the synthesis of other metabolites such as isoprenoids, which have been implicated in tumor progression(12). To elucidate whether it is indeed cholesterol or other upstream metabolites in the pathway that are essential for RMS growth, we specifically targeted the last enzyme in the pathway, DHCR7, using three independent shRNAs (**Fig. 2A and H**). DHCR7 silencing completely halted the growth of RD and KLHEL1 cells (**Fig. 2B and I),** and significantly reduced their colony formation capacity (**Fig. 2C, D, and J, K**). Interestingly, the growth inhibition was not reversed by supplementing the culture media with LDL cholesterol, indicating that RMS cells are highly dependent on the de novo cholesterol synthesis (**Supplementary Fig.1K and 1L**). Additionally, DHCR7-silenced RD and KLHEL1 cells exhibited more apoptotic cells than their shSCR-transduced controls (**Fig. 2E, and L**). Consistent with the effects of DHCR7 gene silencing on tumor cell growth, the DHCR7 specific inhibitor AY9944 decreased the growth of RD and KLHEL1 cells in a dose-dependent manner (**Fig. 2F and M**). Furthermore, treatment with AY9944 compromised the survival of RD and KLHEL1 cells, as evidenced by pronounced apoptosis, measured by the activation of caspase 3/7 (**Fig. 2G and N**). These data indicate that cholesterol biosynthesis is essential for the proliferation, viability, and clonogenic capacity of RMS cells, irrespective of the RMS tumor subtype.

**Figure 2.**
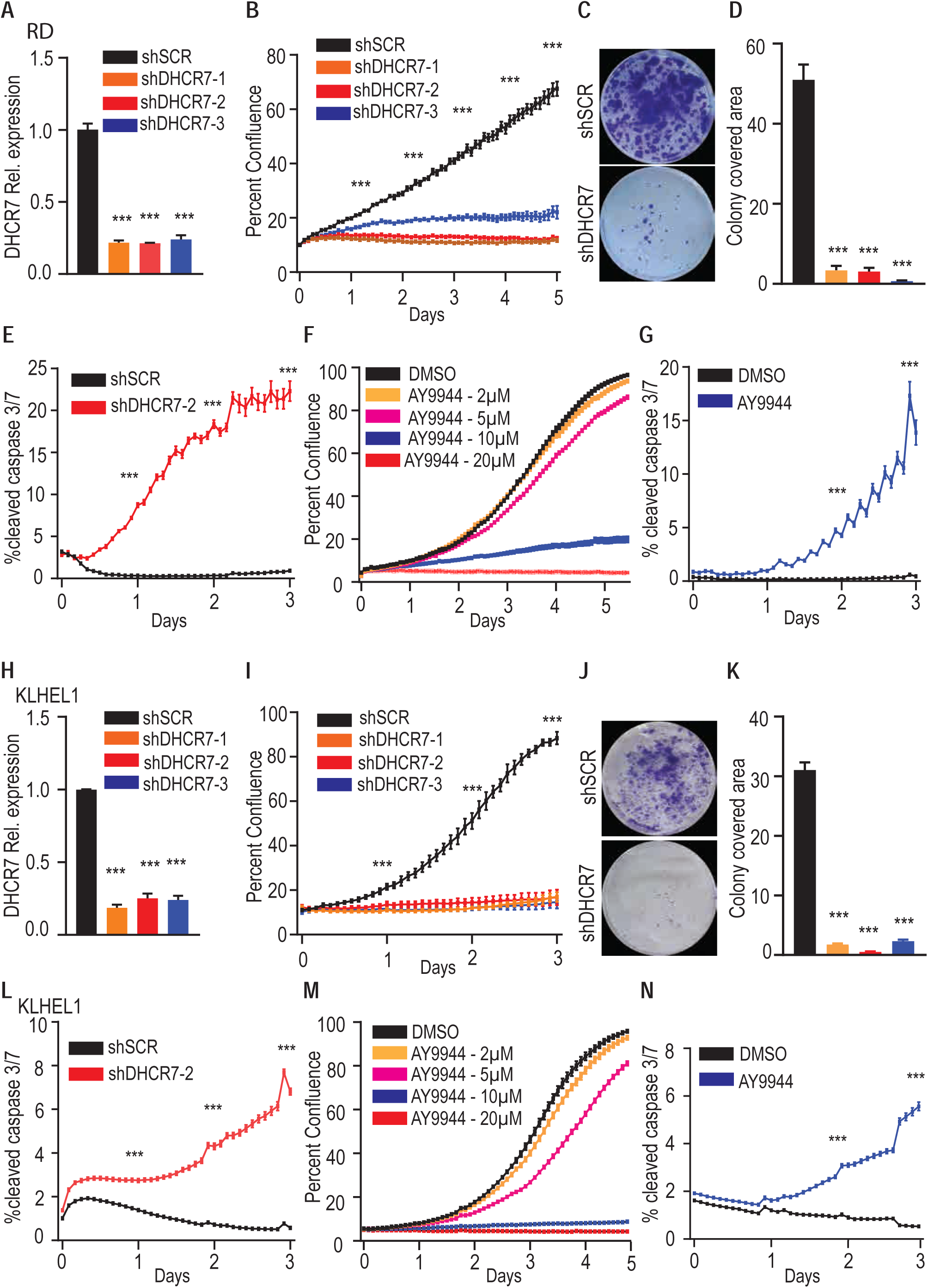
Inhibition of cholesterol biosynthesis suppresses RMS cell growth and survival. (A) Quantitative PCR analysis of DHCR7 mRNA expression in RD cells following shSCR (control) and shDHCR7 transduction with three independent shRNA constructs. (B) Cell growth assessed by IncuCyte live cell imaging for stably expressing shSCR or shDHCR7 RD cells utilizing three different silencing constructs. (C, D) Colony formation assay and corresponding quantification in RD cells. (E) Quantification of cleaved caspase 3/7 activity in control and DHCR7-silenced RD cells using a fluorescent reporter and live imaging. (F) Growth curves of RD cells treated with different concentrations of DHCR7 inhibitor (AY9944) and analyzed by live cell imaging. (G) Caspase 3/7 activity in dimethyl sulfoxide (DMSO) and DHCR7 inhibitor-treated RD cells. (H) qPCR analysis of DHCR7 mRNA expression in KLHEL1 cells following shSCR (control) and shDHCR7 transduction with three independent shRNA constructs. (I) Cell growth based on IncuCyte live cell imaging for stably expressing shSCR or shDHCR7 KLHEL1 cells. (J, K) Colony formation assay and corresponding quantification in KLHEL1 cells. (I) Caspase 3/7 activity in control and PROX1-silenced KLHEL1 cells utilizing a fluorescent reporter. (M) Growth curves of KLHEL1 cells treated with indicated concentrations of DHCR7 inhibitor (AY9944) and analyzed by live cell imaging. (N) Caspase 3/7 activity in DMSO and DHCR7 inhibitor-treated KLHEL1 cells assessed using a fluorescent reporter. *p < 0.05, **p < 0.01, and ***p < 0.001. Data presented as mean ± standard error of the mean (SEM).

### Cholesterol biosynthesis is essential for the growth of RMS tumor xenografts

We next analysed the tumor-propagating potential of cholesterol biosynthesis *in vivo* by engrafting control and DHCR7-silenced RD (FN-RMS) and KLHEL1 (FP-RMS) cells into the left and right flanks of female NOD/SCID/IL2rg mice. Serial tumor volume measurements demonstrated significant growth inhibition in the DHCR7-silenced tumors compared to the shSCR-transduced control tumors (**Fig. 3A**). Tumors derived from the DHCR7-silenced cells were notably smaller and lighter than their respective controls (**Fig. 3 B and C**). Hematoxylin and eosin (H&E)-stained tumor sections from DHCR7-silenced RD and KLHEL1 cells exhibited also reduced cell density compared to control tumors (**Fig. 3 D, H, and L and Supplementary, Fig. S2 C, G, and K**). Immunofluorescence staining for Ki67 and cleaved caspase 3 (CC3) revealed decreased cell proliferation and increased apoptosis in the DHCR7-silenced tumors (**Fig. 3 E, M, and I and, Supplementary Fig. S2 D, H, and L**). These findings provided compelling evidence that cholesterol biosynthesis is indispensable for RMS tumor growth in vivo.

**Figure 3.**
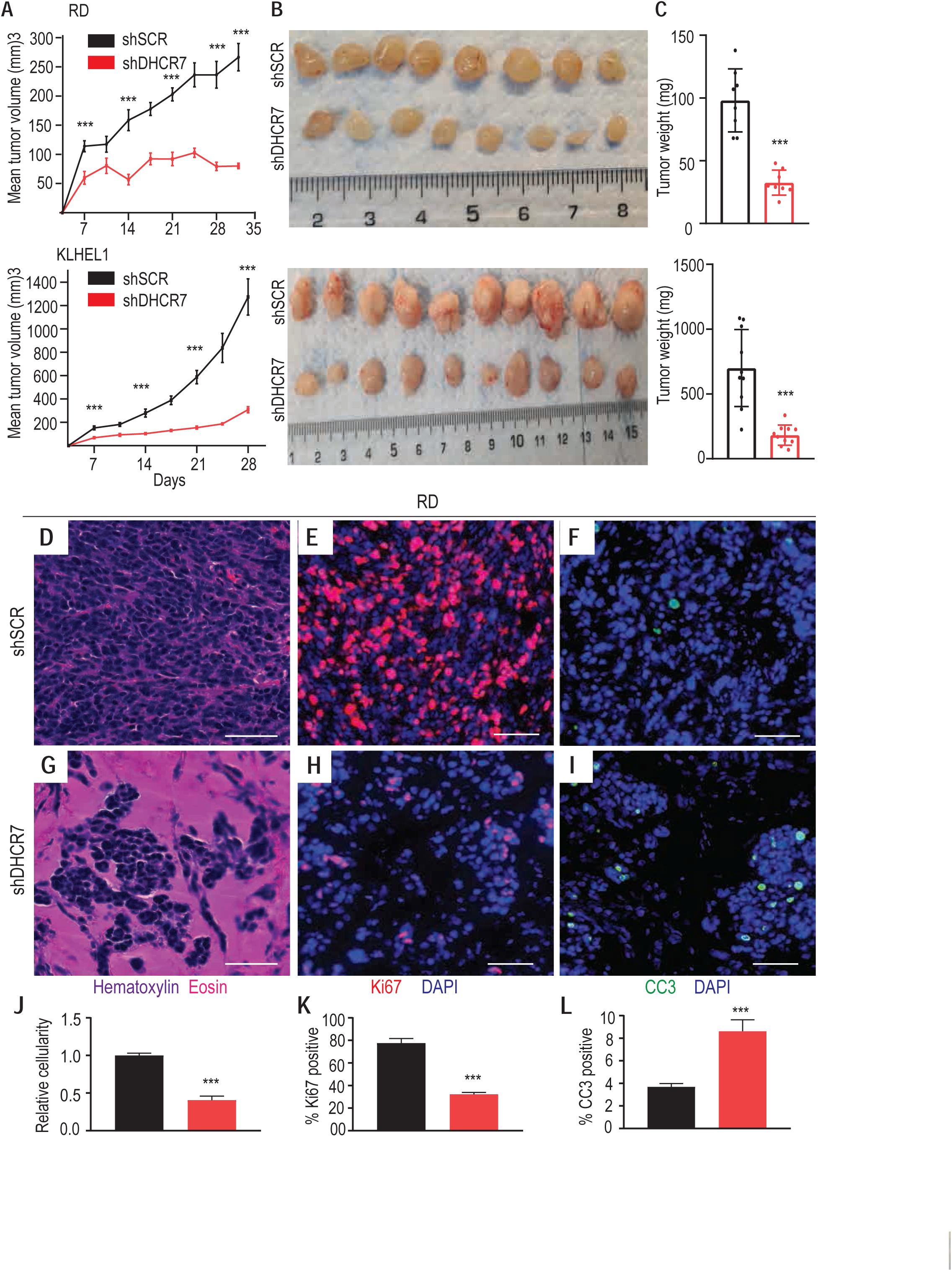
Cholesterol biosynthesis is essential for RMS tumor xenograft growth (A) Tumor growth was quantified by measuring the tumor volumes (mm^3) in RD (top) and KLHEL1 (bottom) xenografts, with 8 replicates for RD and 10 for KLHEL1 per group. (B) Representative images depicting shSCR and shDHCR xenograft tumors derived from RD (Top) and KLHEL1 (Bottom) cells at the end of the experiment. (C) Tumor weights of shSCR and shDHCR7 tumors at the end of the experiment for RD (top) and KLHEL1 (bottom) xenografts. (D–K) Histological analysis of RD xenograft tumors with shSCR (D–F) and shDHCR7 treatment (G–I). Representative H&E-stained sections of the tumors are shown in (D) and (G). Immunofluorescent staining for Ki67 (red) and DAPI (blue) is displayed in (E) and (H), while cleaved caspase 3 (CC3) and DAPI (blue) staining is shown in (F) and (I). (J) Quantification of nuclei per tumor area, (K) percentage of Ki67-positive cells, and (L) percentage of cleaved caspase 3 (CC3)-positive cells. Statistical significance is indicated as follows: *P < 0.05, **P < 0.01, ***P < 0.001. Data are presented as mean ± SEM, with a scale bar of 100 µm.

### Blocking cholesterol biosynthesis in RMS inhibits cell cycle progression and activates ER stress-triggered apoptosis

To elucidate global changes in gene expression in response to cholesterol inhibition in RMS tumors, we isolated tumor RNA and conducted whole transcriptome analysis on DHCR7-silenced and control RD cells. Principal component analysis (PCA) and hierarchical clustering of the RNA-Seq data revealed that the DHCR7-silenced and control cells formed clearly distinct groups (**Fig. 4A and B**). Analysis of the differentially expressed genes (DEGs), using the statistical significance cutoff at false discovery rate (FDR) of < 0.05 and a biological significance cutoff of ≥ 1.5-fold, identified 1961 upregulated and 2698 downregulated genes in the shDHCR7 cells (**Fig. 4C**). Among the top 10 highly downregulated genes were the proliferation marker MKI67 and the transcription factor E2F2 (Supplementary table 1). The family of E2F family transcription factors (E2F1, E2F2, and E2F3), are known activators of cell cycle progression; many crucial cell cycle genes are bona fide E2F targets (19–22). We found that all members of the E2F family were downregulated in the DHCR7 silenced tumor cells **(Supplementary Table 1)**, and this was confirmed by Gene Set Enrichment Analysis (GSEA) GSEA analysis further indicated that several genes involved in various phases of cell cycle progression were significantly downregulated (**Fig. 4D - 4I and Supplementary Fig. 4A and 4E**). Analysis of the effects of DHCR7 knockdown on RD cell cycle distribution demonstrated that DHCR7 depletion caused a progressive increase in the proportion of cells in the G2/M phase, accompanied by a corresponding decrease in the G0/G1 and S phases (**Fig. 4D - 4I**). These findings suggest that inhibition of cholesterol biosynthesis abrogates cell cycle progression and induces cell cycle arrest predominantly at the G2/M phase, by modulating the expression of key cell cycle-related genes.

**Figure 4.**
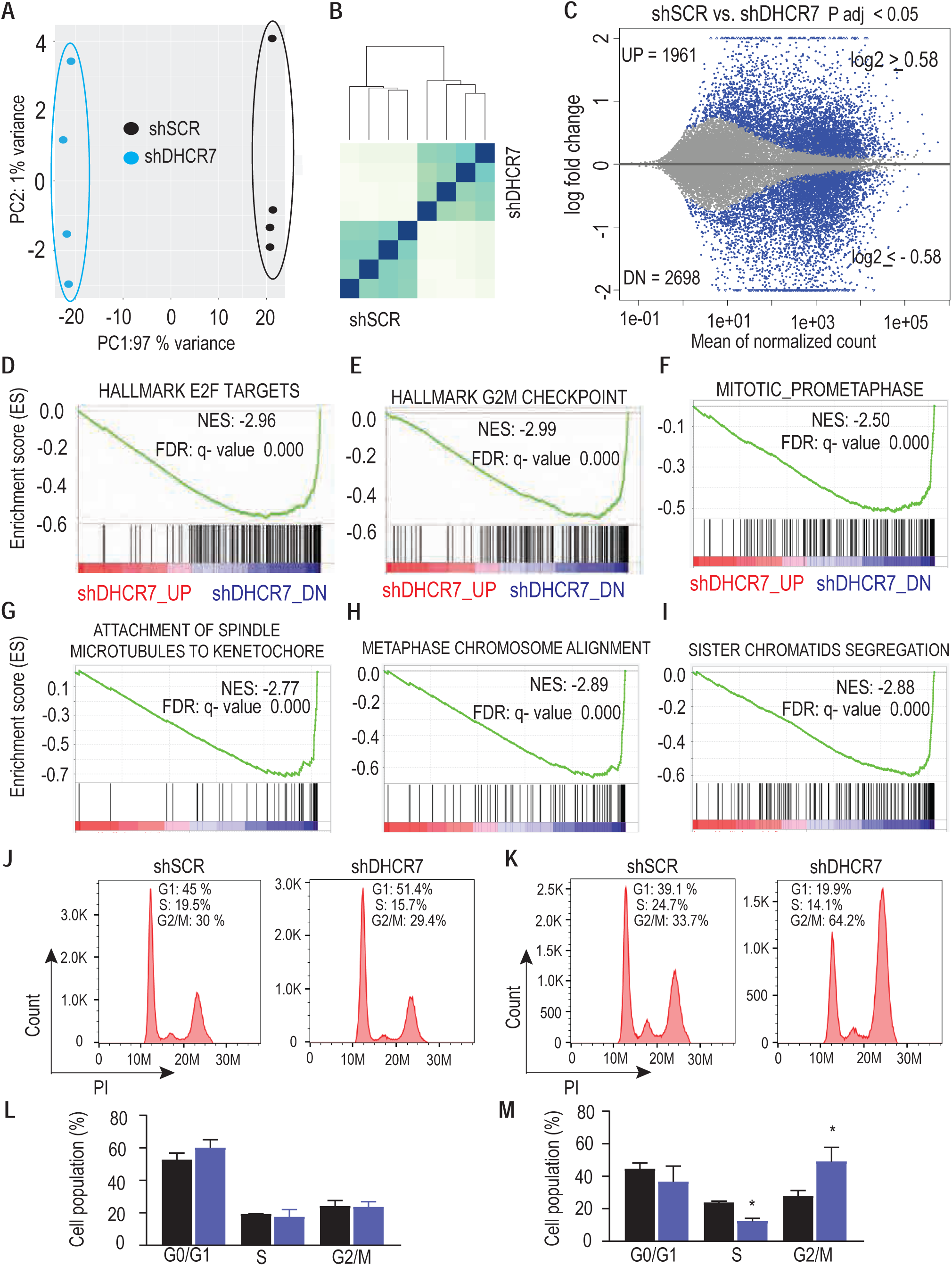
Inhibition of cholesterol synthesis impedes cell cycle progression. (A) Principal Component Analysis (PCA) illustrating the variance between the samples of DHCR7-silenced RD cells compared to control RD cells. (B) Sample distance analysis of RNA-seq samples (n = 4 + 4) depicting the relationship between DHCR7-silenced and control RD cells. (C) MA plot displaying Differentially Expressed Genes (DEGs) in red (FDR < 0.05, log2 fold change (FC) cutoff 0.58) in DHCR7-silenced RD cells compared to controls, encompassing 1,961 upregulated and 2,698 downregulated genes. (D-I) Gene Set Enrichment Analysis (GSEA) plots highlighting the most significantly affected functional categories among downregulated genes in DHCR7-silenced RD cells. The Normalized Enrichment Score (NES) and False Discovery Rate (FDR) values are provided for each category. (J - M) Propidium Iodide (PI) flow cytometry analysis of cell cycle distribution and corresponding statistical analysis in RD cells at first day (J and L) and second day (K and M) post-infection with DHCR7 shRNA. *P < 0.05. Data are presented as mean ± SEM.

Similarly to the response observed in breast cancer cell lines sensitive to atorvastatin-mediated cholesterol inhibition (23), DHCR7 knockdown in RMS cells also increased several key transcripts of the cholesterol biosynthesis pathway (**Fig. 5A),** likely as a compensatory mechanism. Further GSEA analysis of the upregulated genes revealed strong enrichment of genes involved in the endoplasmic reticulum (ER) unfolded protein response (UPR) (Fig. 5B). UPR is typically initiated and regulated by three ER sensors: inositol-requiring enzyme 1 (IRE1), PKR-like ER kinase (PERK), and activating transcription factor 6 (ATF6) (24, 25). Our data demonstrated that transcripts linked to these pathways were increased following DHCR7 silencing (**Supplementary Table 2**). Notably, GSEA analysis also revealed activation of the intrinsic apoptosis signaling pathway activated during prolonged ER stress, (**Fig. 5C**). Consistently with this, our data showed significant upregulation of transcripts of the PERK branch of the UPR, which is known to elicit pro-apoptotic effects (26), and **(Fig. 5D)**. Among these we found the transcription factor activating transcription factor-4 (ATF4), which induces the expression of C/EBP-homologous protein (CHOP), a regulator of pro-apoptotic genes during ER stress. Also ATF4 and its target genes, including CHOP and the CHOP-induced pro-apoptotic gene GADD34, were significantly upregulated by DHCR7 silencing **(Fig. 5E and Supplementary Table 2**). Upregulation of these PERK-related transcipts was also observed upon silencing DHCR7 in KLHEL1 cells (**Fig. 5F**). Since PERK enhances ATF4 expression by phosphorylating its only known target, eukaryotic translation initiation factor 2α (eIF2α), we evaluated the phosphorylation levels of eIF2α. Our results showed increased phosphorylation of eIF2α in both RD and KLHEL1 cells following DHCR7 knockdown. This was accompanied by elevated ATF4 expression and increased CHOP transcription (**Fig. 5G and H**), indicating activation of the PERK-eIF2α-ATF4-CHOP axis. Collectively, our findings demonstrate that inhibition of cholesterol biosynthesis in RMS activates UPR pathways and induces the PERK-ATF4-CHOP axis, leading to ER stress-induced intrinsic apoptosis.

**Figure 5.**
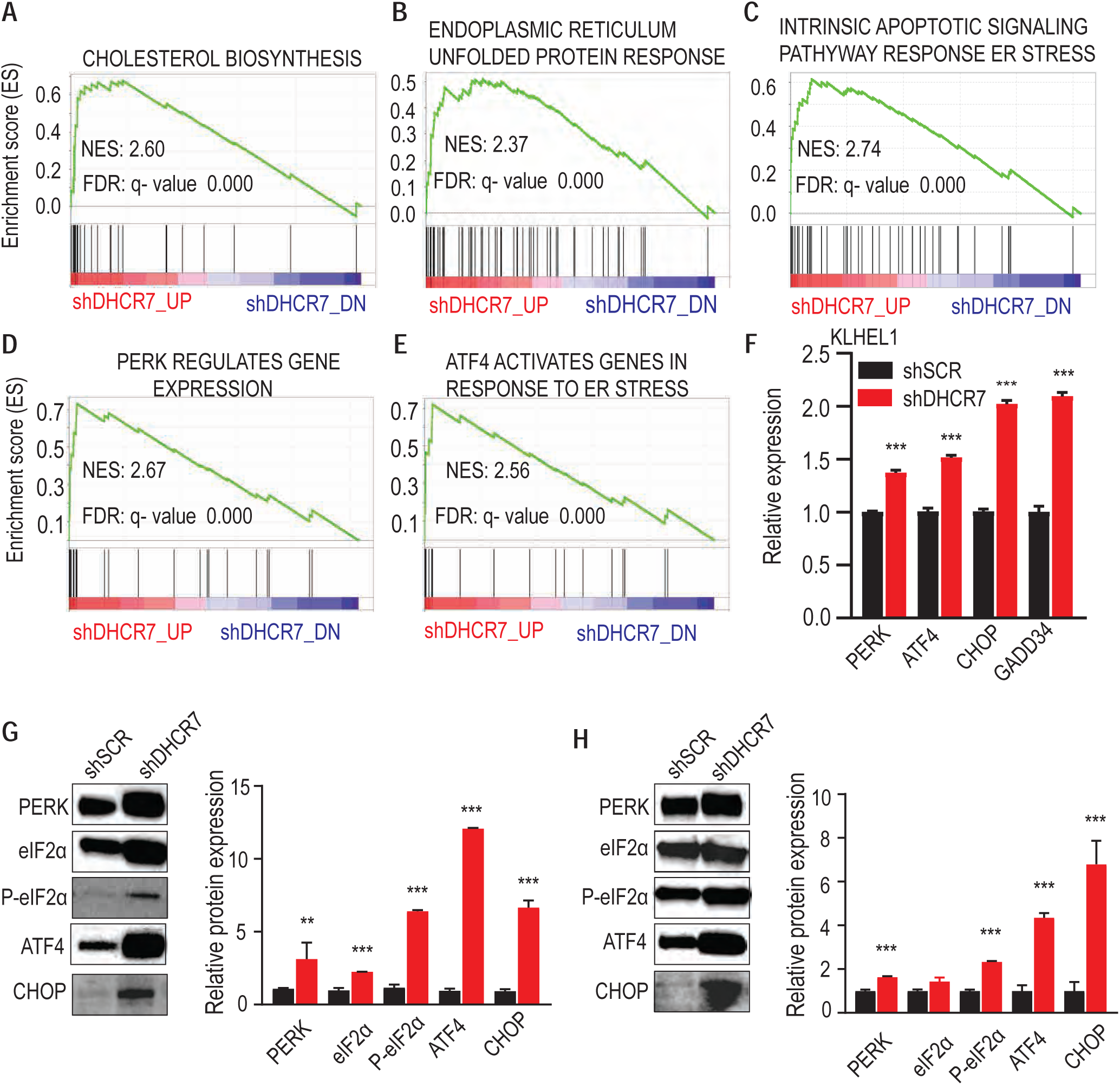
Inhibition of cholesterol biosynthesis induces ER stress-mediated apoptosis. (A-E) Gene Set Enrichment Analysis (GSEA) plots illustrating the most significantly enriched functional categories among the upregulated genes in DHCR7-silenced RD cells. Normalized Enrichment Score (NES) and False Discovery Rate (FDR) are shown for each category. (F) Real-time qPCR analysis of genes’ expression levels involved in the ER stress-induced apoptotic pathway following DHCR7 knockdown in KLHEL cells. Data are presented as mean ± SEM. (G and H) Western blot analysis demonstrating the phosphorylation of eIF2α and the expression of proteins associated with the ER stress-induced apoptotic pathway following DHCR7 knockdown in RD cells (G) and KLHEL1 cells (H).*p < 0.05, **p < 0.01, and ***p < 0.001. Data are presented as mean ± SEM.

## Discussion

Our study identifies cholesterol biosynthesis as a critical metabolic pathway driving RMS progression, offering a novel therapeutic target in both fusion-positive and fusion-negative RMS subtypes. Our findings demonstrate that RMS cells are highly dependent on de novo cholesterol biosynthesis for their growth and survival, and that targeting this pathway, either through genetic or pharmacological inhibition, leads to significant reduction in RMS cell proliferation and increased ER stress-mediated apoptosis, resulting in inhibition of RMS tumor growth. In particular, the central role of the DHCR7 enzyme that carries out the final step in cholesterol biosynthesis, became evident as its silencing halted cell proliferation and significantly suppressed tumor progression in xenograft models. These results not only highlight a previously underexplored metabolic vulnerability in RMS but also suggest that cholesterol biosynthesis inhibition could offer a promising avenue for therapeutic development in high-risk RMS.

Previous studies have shown that cholesterol and lipid metabolism are frequently dysregulated in various cancers (9, 27). Our current findings demonstrate that in RMS, the PROX1 transcription factor, which was recently implicated in RMS growth (8), plays a critical role in regulating cholesterol biosynthesis. Elevated expression of cholesterol biosynthesis genes was observed in both patient-derived RMS tumor samples and RMS cell lines. The regulatory function of PROX1 in RMS metabolism aligns with its role in other cancers, in which it has been shown to promote tumorigenesis via alteration of tumor cell metabolism (28). RNA-seq analysis of the PROX1-silenced RMS cells (FN-RMS and FP-RMS) revealed a core transcriptional program including downregulation of transcripts involved in cholesterol biosynthesis, an effect not observed in-healthy myoblasts. This suggests that PROX1 selectively drives cholesterol metabolism in RMS cells, thereby contributing to their proliferative advantage.

Our experiments demonstrate that the inhibition of cholesterol biosynthesis via genetic silencing of DHCR7 and HMGCR, or pharmacological treatment with specific inhibitors, has profound effects on RMS cell survival and proliferation. Interestingly, the silencing of both HMGCR (at the origin of the mevalonate pathway) and DHCR7 resulted in RMS growth inhibition, suggesting that the critical product is cholesterol rather than its upstream intermediates. RNA sequencing analysis confirmed this by showing that DHCR7 silencing increased the expression of almost all upstream cholesterol biosynthesis genes, which, however, failed to rescue RMS growth inhibition. This indicates that prenylation products derived from the upstream mevalonate pathway, which is important for modification of oncogenic proteins, such as Rho or Ras, are not key regulators of RMS growth. Thus, the suppression of RMS proliferation is primarily linked to the lack of cholesterol production by the tumor cells.

Unlike most normal human cells that rely on exogenous fatty acids, tumors tend to shift towards de novo lipid biosynthesis despite the availability of extracellular lipids (29–31). However, upon cholesterol biosynthesis inhibition, many types of tumors can increase exogenous cholesterol uptake via LDL receptors (LDLR). Thus, sensitizing these cancers to cholesterol biosynthesis inhibitors may require additional restriction of the availability of exogenous cholesterol or a combination with cholesterol uptake blockers (10, 11). However, our results indicate that even in the presence of high levels of external cholesterol, RMS cells and tumors remain dependent on the de novo cholesterol synthesis for growth, underscoring the essential nature of the cholesterol biosynthesis pathway.

The induction of apoptosis upon disruption of cholesterol synthesis pathway was mechanistically linked to the activation of the unfolded protein response (UPR) and all three ER stress pathways—IRE1, PERK, and ATF6. Specifically, the activation of the PERK- eIF2α-ATF4-CHOP axis pointed to an ER stress-triggered apoptotic pathway, linking cholesterol biosynthesis inhibition to apoptosis in RMS cells. This finding suggests that RMS cells rely on an unperturbed cholesterol biosynthesis pathway to maintain ER homeostasis. Blocking of this pathway in RMS cells triggers lethal levels of ER stress that ultimately leads to cell death.

Arrest of the cell cycle at the G2/M phase and downregulation of key cell cycle genes, including members of the E2F transcription factor family and genes involved in various stages of the cell cycle (G1, S, G2, and M) upon DHCR7 silencing indicated that cholesterol biosynthesis is essential not only for maintaining ER stability, but also for the normal progression of the cell cycle in RMS cells. Our RNA sequencing analysis elucidated further molecular consequences of cholesterol biosynthesis disruption in RMS. When we silenced DHCR7, there was a compensatory upregulation of nearly all genes involved in the cholesterol biosynthesis pathway, suggesting that RMS cells possess a hard-wired mechanism to preserve cholesterol production under metabolic stress. Interestingly, similar patterns of transcriptional changes in cholesterol biosynthesis, cell cycle progression, and apoptosis were reported in a global transcriptomic study analysing the effects of statin treatment in breast cancer (23).

The combination of apoptosis, cell cycle arrest, and compensatory upregulation of cholesterol biosynthesis genes underscores the critical dependency of RMS cells on this metabolic pathway and highlights the therapeutic potential of targeting cholesterol metabolism as a strategy to halt RMS progression. Repurposing cholesterol-lowering drugs, such as statins, which inhibit the rate-limiting enzyme HMGCR in the cholesterol biosynthesis pathway, should be tested in combination with current RMS treatment protocols. While the use of statins has resulted in promising treatment results in various preclinical cancer models, our findings suggest that more selective inhibitors targeting the terminal stages of cholesterol biosynthesis, such as DHCR7, may provide a more effective strategy by directly impairing the cholesterol supply required for RMS cell survival.

In conclusion, our study identifies de novo cholesterol biosynthesis as a key metabolic pathway that supports RMS growth and survival. The dependency of RMS cells on this pathway for cell cycle progression, ER homeostasis, and overall survival makes cholesterol biosynthesis an attractive therapeutic target. By inhibiting cholesterol biosynthesis, we could effectively suppress tumor growth and induce apoptosis via ER stress mechanisms, which provides a strong rationale for the development of novel therapies that target a key metabolic vulnerability in RMS cells. Future research should focus on optimising cholesterol biosynthesis inhibition and delivery in preclinical RMS models, and on exploration of combination therapies that may enhance the therapeutic efficacy. Ultimately, targeting cholesterol biosynthesis could offer a novel strategy to overcome drug resistance in RMS patients.

## Acknowledgment

We would like to thank Mari Jokinen, Maria Arrano, Tanja Laakkonen, and Tapio Tainola for their excellent technical help. Elina Ikonen is acknowledged for discussions and comments during the project. We also thank the Laboratory Animal Center at the University of Helsinki for expert animal care, the Biomedicum Imaging Unit for microscope support and the Biomedicum Virus Core and Genome Biology Unit for TRC library construct and virus preparation. The work was funded by the Cancer Foundation Finland sr, Barncancerfonden, Children’s Cancer Foundation Väre, Children’s Cancer Foundation Aamu and Orion Research Foundation sr.

## Authors’ Contributions

N.Y.G: Conceptualization, performed experiments and analyzed data, generated and analyzed transcriptomic data, visualization, writing– original draft, funding acquisition, writing–review and editing. K.K: performed immunoblotting, methodology, and visualization. K.A.: resources, analyzed data, validation, provided expertise and writing–review and editing. R.K.: Conceptualization, supervision, analyzed data, resources, funding acquisition, project administration, writing–review and editing.

