## Supplemental Files for "Targeting de novo cholesterol synthesis in rhabdomyosarcoma induces cell cycle arrest and triggers apoptosis through ER stress-mediated pathways"

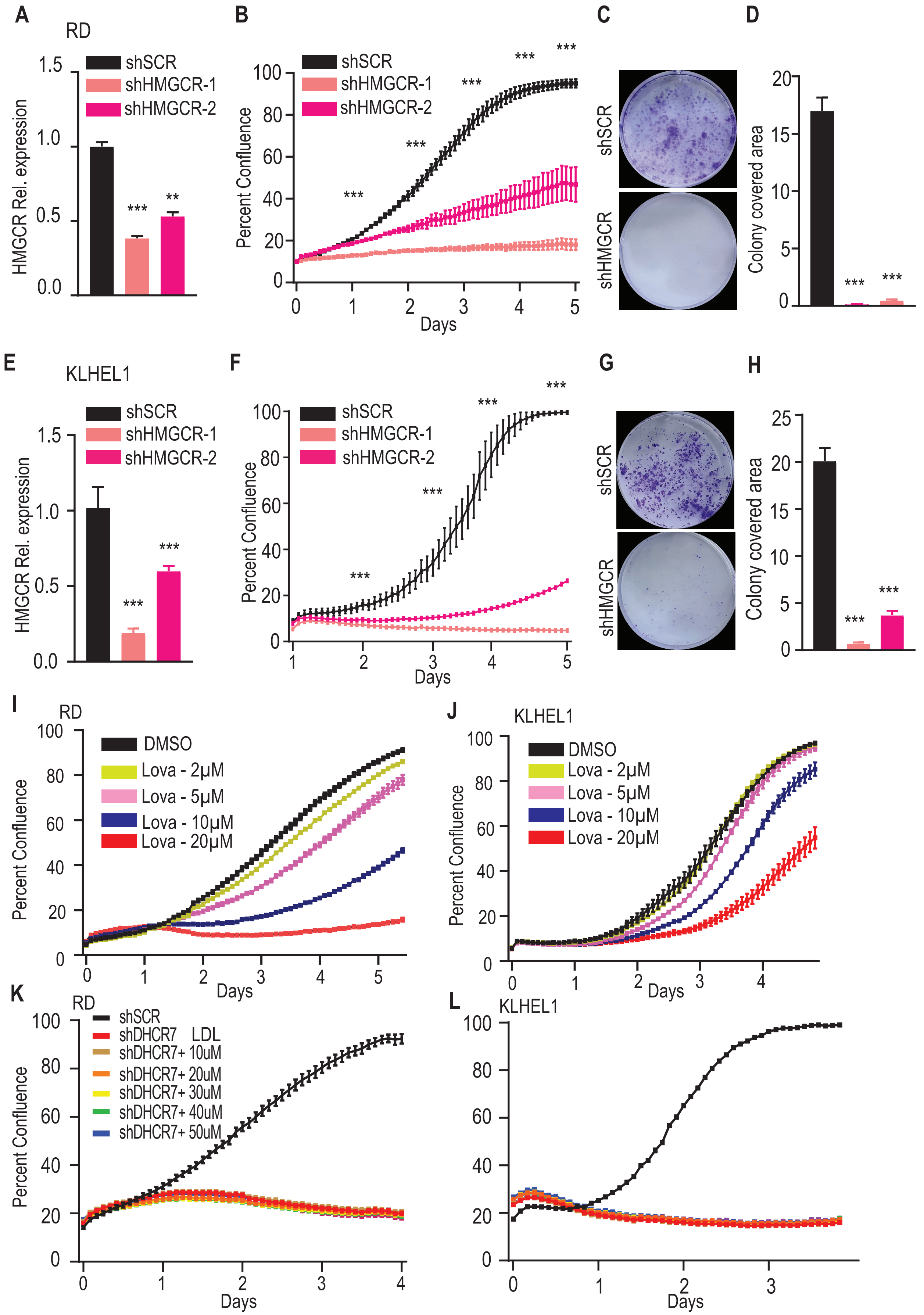

A
RD
HMGCR Rel. expression
shSCR
shHMGCR-1
shHMGCR-2
B
Percent Confluence
Days
C
shSCR
shHMGCR
D
Colony covered area
E
KLHEL1
F
G
H
I
DMSO
Lova - 2μM
Lova - 5μM
Lova - 10μM
Lova - 20μM
J
K
shDHCR7 LDL
shDHCR7+ 10uM
shDHCR7+ 20uM
shDHCR7+ 30uM
shDHCR7+ 40uM
shDHCR7+ 50uM
L

**Supplementary Figure S1.**

Inhibition of cholesterol biosynthesis impairs rhabdomyosarcoma (RMS) cell growth. (A) Quantitative PCR analysis showing HMGCR mRNA expression levels in RD cells transduced with shSCR (control) or two independent shHMGCR shRNA constructs. (B) Cell growth was monitored via IncuCyte live cell imaging for RD cells stably expressing shSCR or shHMGCR across two silencing constructs. (C, D) Colony formation assay in RD cells, along with its quantification. (E) qPCR analysis of HMGCR mRNA levels in KLHEL1 cells following transduction with shSCR (control) or two independent shHMGCR constructs. (F) Cell growth in KLHEL1 cells expressing shSCR or shHMGCR was tracked using IncuCyte live imaging. (G, H) Colony formation assay and its corresponding quantification in KLHEL1 cells. (I, J) Growth curves of RD (I) and KLHEL1 (J) cells treated with different concentrations of lovastatin, analyzed through live cell imaging. (K, L) Supplementation with varying concentrations of LDL cholesterol (LDL-c) could not reverse the antiproliferative effects of cholesterol biosynthesis inhibition with shDHCR7 in RD (K) and KLHEL1 (L) cells. *$p < 0.05$, **$p < 0.01$, and ***$p < 0.001$. Data are shown as mean ± standard error of the mean (SEM).

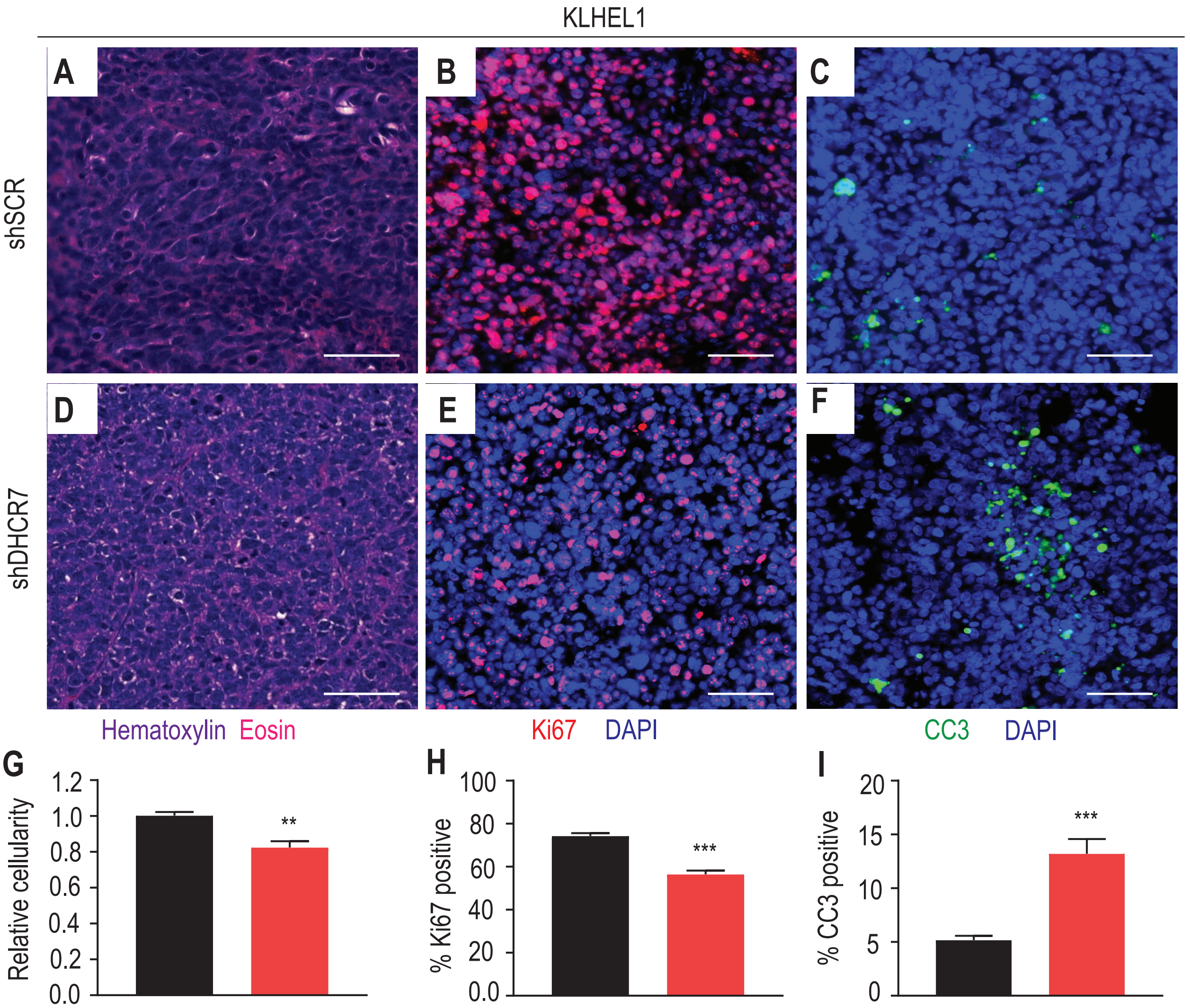


**Supplementary Figure S2.**

Cholesterol biosynthesis is essential for RMS tumor xenograft growth. (A–F) Histological analysis of KLHEL1 xenograft tumors derived from cells transduced with shSCR (A–C) or shDHCR7 (D–F). Representative images of H&E-stained tumor sections (A and D), Ki-67 immunostaining (red) and DAPI (blue) for nuclear visualization (B and E), and cleaved caspase 3 (CC3) immunostaining (green) and DAPI (blue) for apoptosis detection (C and F). (G) Quantification of nuclei count per tumor area. (H) Percentage of Ki67-positive proliferating cells within the tumors. (I) Percentage of activated caspase 3 (CC3)-positive cells, indicating apoptosis. *p < 0.05, **p < 0.01, and ***p < 0.001. Data are presented as mean ± SEM. Scale bar: 100 μm.

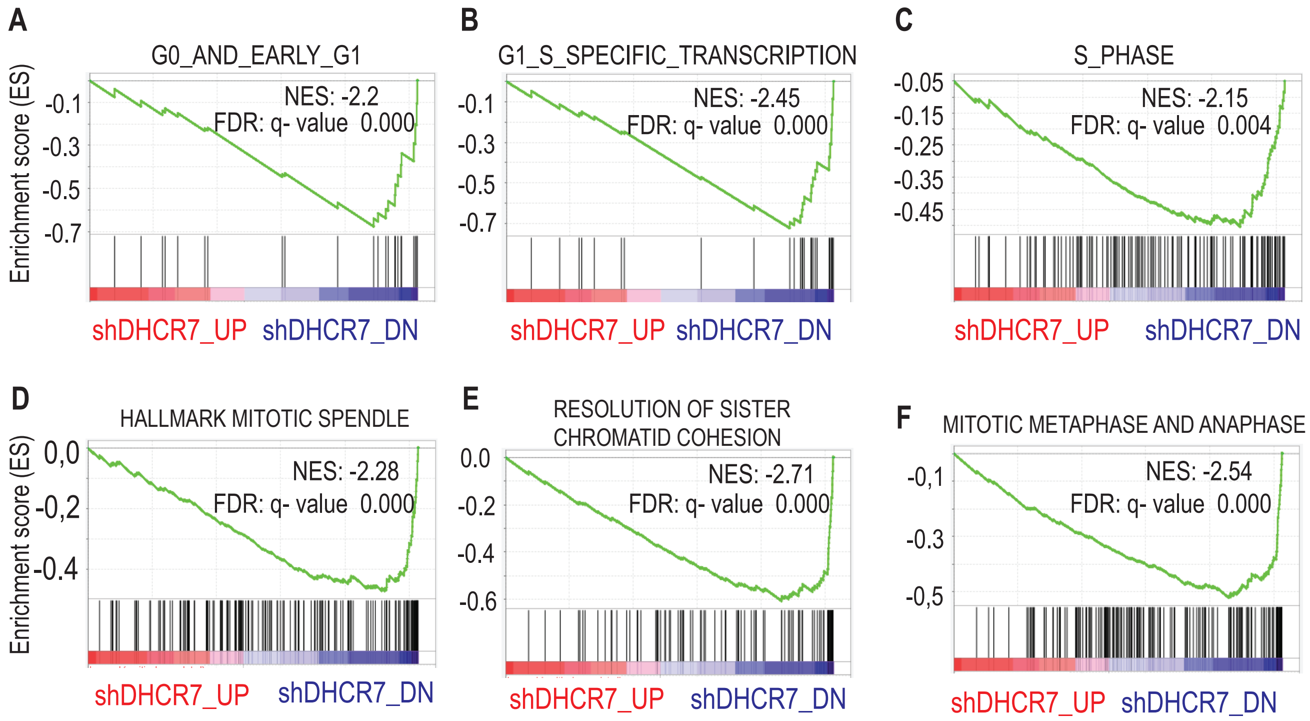


**Supplementary Figure S3.**

Cholesterol synthesis inhibition disrupts cell cycle progression. (A–F) Gene Set Enrichment Analysis (GSEA) plots showing the top functional categories significantly impacted by the downregulation of genes in DHCR7-silenced RD cells. Each plot includes the respective pathways' Normalized Enrichment Score (NES) and False Discovery Rate (FDR).

**Table S1. Downregulated genes in DHCR7 silenced RD cells**

| | symbol | log2FoldChange | lfcSE | pvalue | padj |
|---|---|---|---|---|---|
| ENSG00000196090 | PTPRT | -3.69 | 0.15 | 4.16E-142 | 4.09E-140 |
| ENSG00000213981 | | -3.36 | 1.39 | 4.66E-08 | 2.22E-07 |
| ENSG00000189057 | FAM111B | -3.35 | 0.16 | 4.26E-105 | 2.64E-103 |
| ENSG00000133055 | MYBPH | -3.34 | 0.1 | 3.41E-226 | 8.78E-224 |
| ENSG00000171320 | ESCO2 | -3.28 | 0.11 | 2.90E-213 | 6.72E-211 |
| ENSG00000148773 | MKI67 | -3.25 | 0.05 | 0.00E+00 | 0.00E+00 |
| ENSG00000166803 | PCLAF | -3.25 | 0.17 | 6.73E-88 | 3.17E-86 |
| ENSG00000162975 | KCNF1 | -3.21 | 0.32 | 2.92E-26 | 3.45E-25 |
| ENSG00000007968 | E2F2 | -3.13 | 0.13 | 3.12E-130 | 2.66E-128 |
| ENSG00000186185 | KIF18B | -3.12 | 0.11 | 2.10E-183 | 3.44E-181 |
| ENSG00000005022 | SLC25A5 | -3.06 | 0.06 | 0.00E+00 | 0.00E+00 |
| ENSG00000095139 | ARCN1 | -3.05 | 0.05 | 0.00E+00 | 0.00E+00 |
| ENSG00000166317 | SYNPO2L | -2.98 | 0.11 | 1.06E-158 | 1.23E-156 |
| ENSG00000168418 | KCNG4 | -2.97 | 0.39 | 4.23E-17 | 3.46E-16 |
| ENSG00000262179 | MYMX | -2.87 | 0.2 | 7.76E-48 | 1.66E-46 |
| ENSG00000167767 | KRT80 | -2.84 | 0.1 | 1.94E-196 | 3.65E-194 |
| ENSG00000171848 | RRM2 | -2.83 | 0.06 | 0.00E+00 | 0.00E+00 |
| ENSG00000065328 | MCM10 | -2.83 | 0.12 | 6.05E-132 | 5.24E-130 |
| ENSG00000185585 | OLFML2A | -2.79 | 0.1 | 5.00E-190 | 8.66E-188 |
| ENSG00000131747 | TOP2A | -2.78 | 0.06 | 0.00E+00 | 0.00E+00 |
| ENSG00000122180 | MYOG | -2.77 | 0.07 | 0.00E+00 | 0.00E+00 |
| ENSG00000135451 | TROAP | -2.77 | 0.13 | 2.44E-104 | 1.49E-102 |
| ENSG00000146950 | SHROOM2 | -2.75 | 0.13 | 7.24E-105 | 4.46E-103 |
| ENSG00000133048 | CHI3L1 | -2.73 | 0.23 | 9.75E-36 | 1.57E-34 |
| ENSG00000265763 | ZNF488 | -2.72 | 0.12 | 3.74E-117 | 2.65E-115 |
| ENSG00000104415 | CCN4 | -2.71 | 0.17 | 8.87E-61 | 2.52E-59 |
| ENSG00000178999 | AURKB | -2.7 | 0.08 | 1.22E-227 | 3.20E-225 |
| ENSG00000165272 | AQP3 | -2.7 | 0.1 | 4.71E-179 | 7.08E-177 |
| ENSG00000152253 | SPC25 | -2.67 | 0.15 | 2.91E-76 | 1.13E-74 |
| ENSG00000122966 | CIT | -2.64 | 0.08 | 7.73E-267 | 2.85E-264 |
| ENSG00000114115 | RBP1 | -2.63 | 0.13 | 2.58E-87 | 1.19E-85 |
| ENSG00000109674 | NEIL3 | -2.63 | 0.19 | 2.11E-43 | 4.14E-42 |
| ENSG00000123485 | HJURP | -2.61 | 0.07 | 0.00E+00 | 0.00E+00 |
| ENSG00000024526 | DEPDC1 | -2.6 | 0.08 | 1.94E-236 | 5.63E-234 |
| ENSG00000228340 | MIR646HG | -2.6 | 0.23 | 1.71E-31 | 2.41E-30 |
| ENSG00000164318 | EGFLAM | -2.59 | 0.07 | 6.88E-269 | 2.61E-266 |
| ENSG00000145386 | CCNA2 | -2.58 | 0.07 | 2.35E-287 | 1.06E-284 |
| ENSG00000128284 | APOL3 | -2.58 | 0.29 | 1.45E-21 | 1.43E-20 |
| ENSG00000142945 | KIF2C | -2.57 | 0.07 | 0.00E+00 | 0.00E+00 |
| ENSG00000184113 | CLDN5 | -2.57 | 0.44 | 1.62E-11 | 9.84E-11 |
| ENSG00000090889 | KIF4A | -2.56 | 0.07 | 2.54E-275 | 1.01E-272 |
| ENSG00000126787 | DLGAP5 | -2.53 | 0.08 | 5.88E-218 | 1.41E-215 |
| ENSG00000168509 | HJV | -2.53 | 0.3 | 1.18E-19 | 1.08E-18 |
| ENSG00000163380 | LMOD3 | -2.52 | 0.24 | 9.83E-27 | 1.18E-25 |
| ENSG00000167900 | TK1 | -2.51 | 0.09 | 1.25E-158 | 1.45E-156 |
| ENSG00000198842 | STYXL2 | -2.51 | 0.13 | 5.40E-86 | 2.45E-84 |
| ENSG00000135677 | GNS | -2.5 | 0.04 | 0.00E+00 | 0.00E+00 |
| ENSG00000138180 | CEP55 | -2.5 | 0.08 | 1.22E-212 | 2.79E-210 |

| | | | | | |
|---|---|---|---|---|---|
| ENSG00000168078 | PBK | -2.5 | 0.1 | 1.10E-140 | 1.05E-138 |
| ENSG00000134690 | CDCA8 | -2.49 | 0.07 | 3.43E-283 | 1.52E-280 |
| ENSG00000161888 | SPC24 | -2.49 | 0.13 | 1.40E-82 | 6.07E-81 |
| ENSG00000183044 | ABAT | -2.48 | 0.09 | 4.32E-154 | 4.72E-152 |
| ENSG00000184787 | UBE2G2 | -2.46 | 0.05 | 0.00E+00 | 0.00E+00 |
| ENSG00000280079 | | -2.46 | 0.29 | 4.22E-19 | 3.76E-18 |
| ENSG00000134602 | STK26 | -2.45 | 0.06 | 0.00E+00 | 0.00E+00 |
| ENSG00000115163 | CENPA | -2.45 | 0.14 | 9.60E-66 | 3.03E-64 |
| ENSG00000165480 | SKA3 | -2.45 | 0.15 | 6.47E-59 | 1.78E-57 |
| ENSG00000181826 | RELL1 | -2.44 | 0.09 | 3.87E-169 | 5.05E-167 |
| ENSG00000092470 | WDR76 | -2.44 | 0.14 | 8.07E-68 | 2.63E-66 |
| ENSG00000086967 | MYBPC2 | -2.44 | 0.21 | 1.75E-32 | 2.54E-31 |
| ENSG00000163171 | CDC42EP3 | -2.43 | 0.05 | 0.00E+00 | 0.00E+00 |
| ENSG00000109805 | NCAPG | -2.43 | 0.07 | 9.14E-251 | 2.99E-248 |
| ENSG00000051341 | POLQ | -2.43 | 0.1 | 1.05E-144 | 1.05E-142 |
| ENSG00000129195 | PIMREG | -2.43 | 0.12 | 1.28E-91 | 6.48E-90 |
| ENSG00000055070 | SZRD1 | -2.41 | 0.05 | 0.00E+00 | 0.00E+00 |
| ENSG00000103966 | EHD4 | -2.41 | 0.06 | 0.00E+00 | 0.00E+00 |
| ENSG00000093009 | CDC45 | -2.4 | 0.09 | 3.60E-145 | 3.64E-143 |
| ENSG00000183856 | IQGAP3 | -2.4 | 0.1 | 6.08E-138 | 5.71E-136 |
| ENSG00000171388 | APLN | -2.39 | 0.09 | 4.35E-161 | 5.17E-159 |
| ENSG00000075218 | GTSE1 | -2.38 | 0.07 | 2.33E-233 | 6.55E-231 |
| ENSG00000112984 | KIF20A | -2.38 | 0.08 | 8.18E-189 | 1.41E-186 |
| ENSG00000129173 | E2F8 | -2.38 | 0.16 | 1.40E-54 | 3.55E-53 |
| ENSG00000163808 | KIF15 | -2.37 | 0.08 | 2.10E-175 | 3.02E-173 |
| ENSG00000118194 | TNNT2 | -2.37 | 0.17 | 3.26E-46 | 6.72E-45 |
| ENSG00000124491 | F13A1 | -2.37 | 0.21 | 7.79E-31 | 1.08E-29 |
| ENSG00000276672 | | -2.37 | 0.24 | 9.42E-25 | 1.06E-23 |
| ENSG00000183778 | B3GALT5 | -2.36 | 0.14 | 2.39E-64 | 7.35E-63 |
| ENSG00000270587 | FAM136DP | -2.36 | 0.34 | 2.62E-14 | 1.88E-13 |
| ENSG00000170312 | CDK1 | -2.34 | 0.07 | 6.26E-254 | 2.10E-251 |
| ENSG00000237649 | KIFC1 | -2.34 | 0.09 | 3.39E-153 | 3.65E-151 |
| ENSG00000087589 | CASS4 | -2.34 | 0.12 | 6.34E-92 | 3.25E-90 |
| ENSG00000197299 | BLM | -2.34 | 0.12 | 2.52E-90 | 1.23E-88 |
| ENSG00000101144 | BMP7 | -2.34 | 0.5 | 1.17E-08 | 5.81E-08 |
| ENSG00000147100 | SLC16A2 | -2.33 | 0.09 | 8.56E-153 | 9.18E-151 |
| ENSG00000169607 | CKAP2L | -2.31 | 0.09 | 1.79E-134 | 1.60E-132 |
| ENSG00000146410 | MTFR2 | -2.31 | 0.15 | 8.31E-55 | 2.11E-53 |
| ENSG00000113749 | HRH2 | -2.3 | 0.06 | 0.00E+00 | 0.00E+00 |
| ENSG00000089685 | BIRC5 | -2.3 | 0.08 | 8.08E-208 | 1.76E-205 |
| ENSG00000133063 | CHIT1 | -2.3 | 0.48 | 9.58E-09 | 4.81E-08 |
| ENSG00000159251 | ACTC1 | -2.29 | 0.05 | 0.00E+00 | 0.00E+00 |
| ENSG00000087495 | PHACTR3 | -2.29 | 0.1 | 3.04E-128 | 2.54E-126 |
| ENSG00000105011 | ASF1B | -2.29 | 0.11 | 3.31E-105 | 2.06E-103 |
| ENSG00000198826 | ARHGAP11A | -2.28 | 0.08 | 2.96E-186 | 4.94E-184 |
| ENSG00000179750 | APOBEC3B | -2.28 | 0.14 | 9.75E-63 | 2.87E-61 |
| ENSG00000104879 | CKM | -2.27 | 0.12 | 1.07E-78 | 4.31E-77 |
| ENSG00000171241 | SHCBP1 | -2.24 | 0.07 | 2.77E-204 | 5.77E-202 |
| ENSG00000166851 | PLK1 | -2.24 | 0.08 | 1.45E-181 | 2.28E-179 |
| ENSG00000019186 | CYP24A1 | -2.24 | 0.31 | 8.85E-15 | 6.50E-14 |

| | | | | | |
|---|---|---|---|---|---|
| ENSG00000134222 | PSRC1 | -2.23 | 0.09 | 7.22E-141 | 6.93E-139 |
| ENSG00000180730 | SHISA2 | -2.23 | 0.09 | 9.15E-127 | 7.37E-125 |
| ENSG00000159173 | TNNI1 | -2.22 | 0.06 | 1.90E-282 | 8.16E-280 |
| ENSG00000165244 | ZNF367 | -2.21 | 0.12 | 4.88E-73 | 1.79E-71 |
| ENSG00000163762 | TM4SF18 | -2.21 | 0.17 | 1.50E-41 | 2.82E-40 |
| ENSG00000241399 | CD302 | -2.21 | 0.23 | 6.71E-23 | 7.01E-22 |
| ENSG00000122952 | ZWINT | -2.2 | 0.06 | 0.00E+00 | 0.00E+00 |
| ENSG00000172156 | CCL11 | -2.2 | 0.32 | 5.64E-14 | 3.98E-13 |
| ENSG00000066279 | ASPM | -2.19 | 0.07 | 2.78E-194 | 5.17E-192 |
| ENSG00000144868 | TMEM108 | -2.19 | 0.17 | 1.35E-41 | 2.54E-40 |
| ENSG00000231826 | LINC01819 | -2.19 | 0.71 | 3.67E-06 | 1.48E-05 |
| ENSG00000167325 | RRM1 | -2.18 | 0.06 | 0.00E+00 | 0.00E+00 |
| ENSG00000084070 | SMAP2 | -2.18 | 0.06 | 1.01E-260 | 3.66E-258 |
| ENSG00000260439 | LMF1-AS1 | -2.18 | 0.2 | 1.10E-30 | 1.51E-29 |
| ENSG00000170961 | HAS2 | -2.17 | 0.13 | 6.56E-66 | 2.08E-64 |
| ENSG00000129810 | SGO1 | -2.17 | 0.14 | 5.46E-55 | 1.39E-53 |
| ENSG00000137310 | TCF19 | -2.17 | 0.16 | 2.25E-43 | 4.41E-42 |
| ENSG00000288076 | | -2.17 | 0.22 | 1.15E-25 | 1.33E-24 |
| ENSG00000087586 | AURKA | -2.16 | 0.05 | 0.00E+00 | 0.00E+00 |
| ENSG00000144668 | ITGA9 | -2.16 | 0.06 | 1.52E-275 | 6.22E-273 |
| ENSG00000174371 | EXO1 | -2.16 | 0.11 | 3.12E-82 | 1.34E-80 |
| ENSG00000261305 | | -2.16 | 0.46 | 1.74E-08 | 8.55E-08 |
| ENSG00000100410 | PHF5A | -2.15 | 0.07 | 2.22E-204 | 4.65E-202 |
| ENSG00000175985 | PLEKHD1 | -2.15 | 0.17 | 1.59E-37 | 2.68E-36 |
| ENSG00000113578 | FGF1 | -2.15 | 0.18 | 4.02E-33 | 5.94E-32 |
| ENSG00000172000 | ZNF556 | -2.14 | 0.07 | 4.78E-191 | 8.39E-189 |
| ENSG00000092853 | CLSPN | -2.14 | 0.08 | 4.65E-143 | 4.61E-141 |
| ENSG00000117650 | NEK2 | -2.14 | 0.13 | 4.50E-65 | 1.40E-63 |
| ENSG00000158220 | ESYT3 | -2.14 | 0.17 | 3.44E-38 | 5.94E-37 |
| ENSG00000176890 | TYMS | -2.14 | 0.17 | 2.25E-36 | 3.67E-35 |
| ENSG00000185739 | SRL | -2.14 | 0.25 | 3.40E-19 | 3.06E-18 |
| ENSG00000106554 | CHCHD3 | -2.13 | 0.07 | 2.93E-228 | 7.75E-226 |
| ENSG00000168077 | SCARA3 | -2.13 | 0.08 | 1.45E-172 | 2.00E-170 |
| ENSG00000101057 | MYBL2 | -2.13 | 0.08 | 8.52E-168 | 1.09E-165 |
| ENSG00000115593 | SMYD1 | -2.13 | 0.12 | 6.02E-73 | 2.20E-71 |
| ENSG00000042980 | ADAM28 | -2.13 | 0.14 | 1.92E-56 | 5.00E-55 |
| ENSG00000091704 | CPA1 | -2.13 | 0.23 | 4.51E-22 | 4.55E-21 |
| ENSG00000172482 | AGXT | -2.13 | 0.24 | 1.44E-20 | 1.37E-19 |
| ENSG00000255561 | FDXACB1 | -2.12 | 0.2 | 1.29E-28 | 1.65E-27 |
| ENSG00000164530 | PI16 | -2.12 | 0.29 | 8.18E-15 | 6.02E-14 |
| ENSG00000233290 | | -2.12 | 0.48 | 7.52E-08 | 3.52E-07 |
| ENSG00000156970 | BUB1B | -2.11 | 0.07 | 5.30E-221 | 1.32E-218 |
| ENSG00000122367 | LDB3 | -2.11 | 0.15 | 3.50E-48 | 7.52E-47 |
| ENSG00000107562 | CXCL12 | -2.11 | 0.24 | 4.10E-21 | 4.00E-20 |
| ENSG00000143816 | WNT9A | -2.1 | 0.08 | 7.50E-142 | 7.35E-140 |
| ENSG00000143228 | NUF2 | -2.1 | 0.1 | 5.75E-110 | 3.75E-108 |
| ENSG00000161896 | IP6K3 | -2.1 | 0.11 | 9.87E-85 | 4.44E-83 |
| ENSG00000169116 | PARM1 | -2.1 | 0.17 | 1.82E-35 | 2.91E-34 |
| ENSG00000101198 | NKAIN4 | -2.09 | 0.09 | 2.40E-116 | 1.67E-114 |
| ENSG00000111206 | FOXM1 | -2.08 | 0.09 | 1.99E-130 | 1.71E-128 |

| | | | | | |
|---|---|---|---|---|---|
| ENSG00000127951 | FGL2 | -2.08 | 0.23 | 7.63E-22 | 7.63E-21 |
| ENSG00000177613 | CSTF2T | -2.07 | 0.06 | 7.40E-294 | 3.52E-291 |
| ENSG00000138160 | KIF11 | -2.07 | 0.06 | 4.42E-230 | 1.18E-227 |
| ENSG00000154328 | NEIL2 | -2.07 | 0.08 | 1.68E-152 | 1.79E-150 |
| ENSG00000187908 | DMBT1 | -2.07 | 0.55 | 9.53E-07 | 4.05E-06 |
| ENSG00000169504 | CLIC4 | -2.06 | 0.04 | 0.00E+00 | 0.00E+00 |
| ENSG00000085840 | ORC1 | -2.06 | 0.08 | 2.62E-143 | 2.61E-141 |
| ENSG00000166033 | HTRA1 | -2.06 | 0.11 | 2.79E-78 | 1.12E-76 |
| ENSG00000127564 | PKMYT1 | -2.06 | 0.14 | 3.76E-51 | 8.68E-50 |
| ENSG00000116985 | BMP8B | -2.06 | 0.3 | 5.84E-14 | 4.12E-13 |
| ENSG00000222022 | | -2.06 | 0.43 | 1.27E-08 | 6.33E-08 |
| ENSG00000085999 | RAD54L | -2.05 | 0.14 | 4.13E-49 | 9.09E-48 |
| ENSG00000205097 | FRG2 | -2.05 | 0.24 | 3.31E-19 | 2.98E-18 |
| ENSG00000136205 | TNS3 | -2.04 | 0.06 | 6.25E-283 | 2.73E-280 |
| ENSG00000184661 | CDCA2 | -2.04 | 0.08 | 2.94E-163 | 3.55E-161 |
| ENSG00000175063 | UBE2C | -2.03 | 0.06 | 5.89E-239 | 1.77E-236 |
| ENSG00000137812 | KNL1 | -2.03 | 0.11 | 4.10E-81 | 1.72E-79 |
| ENSG00000068903 | SIRT2 | -2.02 | 0.08 | 2.40E-157 | 2.76E-155 |
| ENSG00000131153 | GINS2 | -2.01 | 0.1 | 1.40E-97 | 7.77E-96 |
| ENSG00000154839 | SKA1 | -2.01 | 0.12 | 6.24E-69 | 2.09E-67 |
| ENSG00000175841 | FAM172BP | -2.01 | 0.44 | 5.08E-08 | 2.41E-07 |
| ENSG00000136231 | IGF2BP3 | -2 | 0.05 | 0.00E+00 | 0.00E+00 |
| ENSG00000205221 | VIT | -2 | 0.09 | 2.14E-121 | 1.61E-119 |
| ENSG00000104419 | NDRG1 | -1.99 | 0.07 | 3.67E-194 | 6.62E-192 |
| ENSG00000170439 | METTL7B | -1.99 | 0.09 | 8.32E-107 | 5.30E-105 |
| ENSG00000167306 | MYO5B | -1.99 | 0.1 | 6.85E-88 | 3.22E-86 |
| ENSG00000215784 | FAM72D | -1.99 | 0.21 | 4.01E-23 | 4.21E-22 |
| ENSG00000117724 | CENPF | -1.98 | 0.06 | 3.63E-220 | 8.80E-218 |
| ENSG00000172893 | DHCR7 | -1.98 | 0.07 | 5.02E-171 | 6.74E-169 |
| ENSG00000159337 | PLA2G4D | -1.98 | 0.53 | 1.36E-06 | 5.72E-06 |
| ENSG00000138778 | CENPE | -1.97 | 0.08 | 2.20E-151 | 2.32E-149 |
| ENSG00000165304 | MELK | -1.97 | 0.09 | 5.19E-112 | 3.46E-110 |
| ENSG00000266709 | | -1.97 | 0.1 | 2.26E-92 | 1.16E-90 |
| ENSG00000197943 | PLCG2 | -1.97 | 0.13 | 4.49E-53 | 1.09E-51 |
| ENSG00000164124 | TMEM144 | -1.97 | 0.15 | 1.68E-40 | 3.06E-39 |
| ENSG00000259182 | | -1.97 | 0.33 | 3.66E-11 | 2.18E-10 |
| ENSG00000115461 | IGFBP5 | -1.96 | 0.08 | 5.80E-149 | 6.00E-147 |
| ENSG00000080986 | NDC80 | -1.96 | 0.11 | 1.27E-79 | 5.21E-78 |
| ENSG00000183580 | FBXL7 | -1.96 | 0.11 | 1.80E-71 | 6.36E-70 |
| ENSG00000092036 | HAUS4 | -1.96 | 0.29 | 2.27E-13 | 1.55E-12 |
| ENSG00000151224 | MAT1A | -1.96 | 0.97 | 2.29E-05 | 8.54E-05 |
| ENSG00000117266 | CDK18 | -1.95 | 0.07 | 1.18E-171 | 1.60E-169 |
| ENSG00000225826 | LINC00626 | -1.95 | 0.07 | 2.95E-169 | 3.86E-167 |
| ENSG00000173281 | PPP1R3B | -1.95 | 0.07 | 7.08E-155 | 7.92E-153 |
| ENSG00000169679 | BUB1 | -1.95 | 0.08 | 8.31E-147 | 8.50E-145 |
| ENSG00000256663 | | -1.95 | 0.23 | 1.03E-19 | 9.42E-19 |
| ENSG00000118193 | KIF14 | -1.94 | 0.07 | 2.75E-181 | 4.29E-179 |
| ENSG00000187616 | MYMK | -1.94 | 0.2 | 9.99E-25 | 1.12E-23 |
| ENSG00000196754 | S100A2 | -1.93 | 0.06 | 5.02E-239 | 1.53E-236 |
| ENSG00000164104 | HMGB2 | -1.93 | 0.08 | 5.43E-132 | 4.74E-130 |

| | | | | | |
|---|---|---|---|---|---|
| ENSG00000112964 | GHR | -1.93 | 0.15 | 1.21E-41 | 2.27E-40 |
| ENSG00000150938 | CRIM1 | -1.92 | 0.04 | 0.00E+00 | 0.00E+00 |
| ENSG00000196562 | SULF2 | -1.92 | 0.06 | 1.70E-250 | 5.50E-248 |
| ENSG00000136492 | BRIP1 | -1.92 | 0.12 | 7.37E-57 | 1.93E-55 |
| ENSG00000069431 | ABCC9 | -1.92 | 0.21 | 5.56E-21 | 5.39E-20 |
| ENSG00000164440 | TXLNB | -1.92 | 0.22 | 1.31E-20 | 1.25E-19 |
| ENSG00000134121 | CHL1 | -1.92 | 1.02 | 2.45E-05 | 9.11E-05 |
| ENSG00000248774 | | -1.92 | 0.96 | 2.89E-05 | 1.07E-04 |
| ENSG00000143476 | DTL | -1.91 | 0.07 | 3.99E-177 | 5.87E-175 |
| ENSG00000142731 | PLK4 | -1.91 | 0.1 | 3.55E-76 | 1.38E-74 |
| ENSG00000186871 | ERCC6L | -1.91 | 0.11 | 1.05E-71 | 3.71E-70 |
| ENSG00000276043 | UHRF1 | -1.9 | 0.07 | 1.49E-179 | 2.26E-177 |
| ENSG00000072571 | HMMR | -1.9 | 0.1 | 1.36E-82 | 5.89E-81 |
| ENSG00000079385 | CEACAM1 | -1.9 | 0.14 | 5.84E-41 | 1.08E-39 |
| ENSG00000188167 | TMPPE | -1.9 | 0.15 | 1.68E-40 | 3.06E-39 |
| ENSG00000104267 | CA2 | -1.9 | 0.64 | 1.43E-05 | 5.44E-05 |
| ENSG00000094880 | CDC23 | -1.89 | 0.08 | 1.98E-135 | 1.79E-133 |
| ENSG00000183775 | KCTD16 | -1.89 | 0.19 | 5.17E-24 | 5.62E-23 |
| ENSG00000117069 | ST6GALNAC5 | -1.89 | 0.39 | 1.21E-08 | 6.04E-08 |
| ENSG00000153904 | DDAH1 | -1.88 | 0.04 | 0.00E+00 | 0.00E+00 |
| ENSG00000101447 | FAM83D | -1.88 | 0.06 | 5.53E-206 | 1.19E-203 |
| ENSG00000048740 | CELF2 | -1.88 | 0.06 | 2.78E-201 | 5.55E-199 |
| ENSG00000021826 | CPS1 | -1.88 | 0.07 | 2.04E-142 | 2.01E-140 |
| ENSG00000163909 | HEYL | -1.87 | 0.06 | 3.20E-230 | 8.64E-228 |
| ENSG00000140451 | PIF1 | -1.87 | 0.13 | 9.02E-47 | 1.89E-45 |
| ENSG00000102445 | RUBCNL | -1.87 | 0.2 | 1.40E-22 | 1.44E-21 |
| ENSG00000174238 | PITPNA | -1.86 | 0.05 | 9.84E-298 | 4.77E-295 |
| ENSG00000197296 | FITM2 | -1.86 | 0.07 | 1.19E-168 | 1.54E-166 |
| ENSG00000112742 | TTK | -1.86 | 0.09 | 3.39E-89 | 1.63E-87 |
| ENSG00000082497 | SERTAD4 | -1.86 | 0.14 | 7.46E-41 | 1.37E-39 |
| ENSG00000095627 | TDRD1 | -1.86 | 0.48 | 8.65E-07 | 3.69E-06 |
| ENSG00000113048 | MRPS27 | -1.85 | 0.06 | 2.43E-236 | 6.99E-234 |
| ENSG00000151726 | ACSL1 | -1.85 | 0.06 | 8.17E-201 | 1.62E-198 |
| ENSG00000129103 | SUMF2 | -1.85 | 0.06 | 3.66E-181 | 5.67E-179 |
| ENSG00000109063 | MYH3 | -1.85 | 0.08 | 1.72E-130 | 1.48E-128 |
| ENSG00000115756 | HPCAL1 | -1.85 | 0.08 | 5.16E-121 | 3.86E-119 |
| ENSG00000137507 | LRRC32 | -1.85 | 0.12 | 2.85E-53 | 6.95E-52 |
| ENSG00000226237 | GAS1RR | -1.85 | 0.3 | 1.39E-11 | 8.48E-11 |
| ENSG00000146386 | ABRACL | -1.84 | 0.1 | 2.88E-82 | 1.23E-80 |
| ENSG00000111665 | CDCA3 | -1.84 | 0.1 | 1.79E-79 | 7.30E-78 |
| ENSG00000105251 | SHD | -1.84 | 0.1 | 4.31E-78 | 1.72E-76 |
| ENSG00000158246 | TENT5B | -1.84 | 0.15 | 1.55E-35 | 2.48E-34 |
| ENSG00000273893 | | -1.84 | 0.25 | 6.87E-15 | 5.07E-14 |
| ENSG00000186198 | SLC51B | -1.84 | 0.32 | 1.59E-10 | 9.09E-10 |
| ENSG00000116774 | OLFML3 | -1.83 | 0.05 | 1.30E-302 | 6.69E-300 |
| ENSG00000144554 | FANCD2 | -1.83 | 0.08 | 9.28E-117 | 6.48E-115 |
| ENSG00000108924 | HLF | -1.83 | 0.17 | 2.62E-30 | 3.56E-29 |
| ENSG00000254978 | ALG1L9P | -1.83 | 0.89 | 5.50E-05 | 1.97E-04 |
| ENSG00000127337 | YEATS4 | -1.82 | 0.08 | 5.20E-131 | 4.48E-129 |
| ENSG00000117308 | GALE | -1.82 | 0.08 | 2.40E-128 | 2.01E-126 |

| | | | | | |
|---|---|---|---|---|---|
| ENSG00000134539 | KLRD1 | -1.82 | 0.19 | 1.76E-23 | 1.88E-22 |
| ENSG00000117586 | TNFSF4 | -1.82 | 0.2 | 2.31E-22 | 2.36E-21 |
| ENSG00000278195 | SSTR3 | -1.82 | 0.32 | 3.02E-10 | 1.69E-09 |
| ENSG00000272989 | LINC02012 | -1.82 | 0.33 | 7.55E-10 | 4.11E-09 |
| ENSG00000108654 | DDX5 | -1.81 | 0.05 | 0.00E+00 | 0.00E+00 |
| ENSG00000162063 | CCNF | -1.81 | 0.08 | 1.92E-117 | 1.37E-115 |
| ENSG00000111247 | RAD51AP1 | -1.81 | 0.09 | 1.07E-97 | 5.95E-96 |
| ENSG00000166106 | ADAMTS15 | -1.81 | 0.21 | 5.33E-19 | 4.73E-18 |
| ENSG00000267655 | | -1.81 | 0.42 | 1.76E-07 | 8.00E-07 |
| ENSG00000017483 | SLC38A5 | -1.81 | 0.61 | 1.93E-05 | 7.25E-05 |
| ENSG00000273024 | INTS4P2 | -1.81 | 0.77 | 5.91E-05 | 2.11E-04 |
| ENSG00000110092 | CCND1 | -1.8 | 0.03 | 0.00E+00 | 0.00E+00 |
| ENSG00000121152 | NCAPH | -1.8 | 0.06 | 1.31E-181 | 2.08E-179 |
| ENSG00000128849 | CGNL1 | -1.8 | 0.12 | 1.39E-51 | 3.24E-50 |
| ENSG00000183287 | CCBE1 | -1.8 | 0.38 | 2.80E-08 | 1.36E-07 |
| ENSG00000253311 | LINC01847 | -1.8 | 0.43 | 3.21E-07 | 1.43E-06 |
| ENSG00000124098 | FAM210B | -1.79 | 0.06 | 1.27E-170 | 1.69E-168 |
| ENSG00000100526 | CDKN3 | -1.79 | 0.15 | 1.96E-35 | 3.12E-34 |
| ENSG00000196550 | FAM72A | -1.79 | 0.21 | 5.36E-20 | 4.99E-19 |
| ENSG00000285867 | | -1.79 | 0.4 | 1.21E-07 | 5.57E-07 |
| ENSG00000287446 | | -1.79 | 0.6 | 1.96E-05 | 7.35E-05 |
| ENSG00000029534 | ANK1 | -1.78 | 0.04 | 0.00E+00 | 0.00E+00 |
| ENSG00000100167 | SEPTIN3 | -1.78 | 0.1 | 7.68E-68 | 2.51E-66 |
| ENSG00000086848 | ALG9 | -1.78 | 0.14 | 8.95E-37 | 1.48E-35 |
| ENSG00000164764 | SBSPON | -1.78 | 0.19 | 3.45E-22 | 3.50E-21 |
| ENSG00000224043 | CCNT2-AS1 | -1.78 | 0.21 | 1.47E-19 | 1.34E-18 |
| ENSG00000137845 | ADAM10 | -1.77 | 0.06 | 2.91E-225 | 7.39E-223 |
| ENSG00000065361 | ERBB3 | -1.77 | 0.08 | 9.45E-105 | 5.81E-103 |
| ENSG00000076382 | SPAG5 | -1.77 | 0.09 | 9.72E-95 | 5.23E-93 |
| ENSG00000128285 | MCHR1 | -1.77 | 0.09 | 2.72E-89 | 1.32E-87 |
| ENSG00000281406 | BLACAT1 | -1.77 | 0.26 | 8.08E-14 | 5.64E-13 |
| ENSG00000289357 | | -1.77 | 0.31 | 1.96E-10 | 1.12E-09 |
| ENSG00000166503 | HDGFL3 | -1.76 | 0.06 | 2.98E-175 | 4.26E-173 |
| ENSG00000163485 | ADORA1 | -1.76 | 0.08 | 2.38E-114 | 1.62E-112 |
| ENSG00000144810 | COL8A1 | -1.76 | 0.26 | 1.53E-13 | 1.05E-12 |
| ENSG00000170927 | PKHD1 | -1.76 | 0.68 | 5.21E-05 | 1.87E-04 |
| ENSG00000140937 | CDH11 | -1.75 | 0.05 | 1.80E-268 | 6.74E-266 |
| ENSG00000196296 | ATP2A1 | -1.75 | 0.07 | 1.23E-125 | 9.74E-124 |
| ENSG00000162630 | B3GALT2 | -1.75 | 0.07 | 3.32E-125 | 2.60E-123 |
| ENSG00000013810 | TACC3 | -1.75 | 0.07 | 5.00E-125 | 3.90E-123 |
| ENSG00000198894 | CIPC | -1.75 | 0.09 | 4.88E-87 | 2.23E-85 |
| ENSG00000130052 | STARD8 | -1.75 | 0.1 | 1.19E-74 | 4.43E-73 |
| ENSG00000261167 | | -1.75 | 0.18 | 3.55E-25 | 4.02E-24 |
| ENSG00000182533 | CAV3 | -1.75 | 0.22 | 1.26E-16 | 1.01E-15 |
| ENSG00000137142 | IGFBPL1 | -1.74 | 0.05 | 3.60E-271 | 1.39E-268 |
| ENSG00000107738 | VSIR | -1.74 | 0.08 | 8.03E-114 | 5.45E-112 |
| ENSG00000138131 | LOXL4 | -1.74 | 0.1 | 2.56E-70 | 8.86E-69 |
| ENSG00000141161 | UNC45B | -1.74 | 0.12 | 6.05E-52 | 1.43E-50 |
| ENSG00000120162 | MOB3B | -1.74 | 0.18 | 7.08E-24 | 7.67E-23 |
| ENSG00000113368 | LMNB1 | -1.73 | 0.05 | 7.80E-233 | 2.17E-230 |

| | | | | | |
|---|---|---|---|---|---|
| ENSG00000179151 | EDC3 | -1.73 | 0.07 | 2.05E-122 | 1.57E-120 |
| ENSG00000129038 | LOXL1 | -1.73 | 0.1 | 2.96E-67 | 9.55E-66 |
| ENSG00000171992 | SYNPO | -1.73 | 0.11 | 5.05E-54 | 1.26E-52 |
| ENSG00000123454 | DBH | -1.73 | 0.13 | 5.47E-41 | 1.01E-39 |
| ENSG00000229267 | SNHG31 | -1.73 | 0.36 | 2.63E-08 | 1.28E-07 |
| ENSG00000100297 | MCM5 | -1.72 | 0.06 | 4.82E-169 | 6.24E-167 |
| ENSG00000112992 | NNT | -1.72 | 0.07 | 8.30E-144 | 8.33E-142 |
| ENSG00000134198 | TSPAN2 | -1.72 | 0.41 | 3.57E-07 | 1.58E-06 |
| ENSG00000231811 | | -1.72 | 0.68 | 6.69E-05 | 2.38E-04 |
| ENSG00000143333 | RGS16 | -1.71 | 0.04 | 0.00E+00 | 0.00E+00 |
| ENSG00000183873 | SCN5A | -1.71 | 0.06 | 2.30E-167 | 2.90E-165 |
| ENSG00000129910 | CDH15 | -1.71 | 0.08 | 1.40E-99 | 8.12E-98 |
| ENSG00000134207 | SYT6 | -1.71 | 0.19 | 6.85E-22 | 6.87E-21 |
| ENSG00000070808 | CAMK2A | -1.71 | 0.24 | 1.57E-14 | 1.14E-13 |
| ENSG00000185028 | LRRC14B | -1.71 | 0.77 | 1.07E-04 | 3.70E-04 |
| ENSG00000263513 | FAM72C | -1.7 | 0.25 | 2.45E-13 | 1.67E-12 |
| ENSG00000185608 | MRPL40 | -1.69 | 0.09 | 1.18E-80 | 4.90E-79 |
| ENSG00000260804 | LINC01963 | -1.69 | 0.09 | 1.17E-72 | 4.23E-71 |
| ENSG00000279348 | | -1.69 | 0.11 | 3.07E-59 | 8.52E-58 |
| ENSG00000136943 | CTSV | -1.69 | 0.12 | 1.06E-48 | 2.30E-47 |
| ENSG00000183196 | CHST6 | -1.69 | 0.12 | 3.77E-44 | 7.49E-43 |
| ENSG00000272711 | HK2-DT | -1.69 | 0.15 | 2.78E-31 | 3.91E-30 |
| ENSG00000283329 | FAM240B | -1.69 | 0.19 | 3.26E-20 | 3.06E-19 |
| ENSG00000214357 | NEURL1B | -1.69 | 0.21 | 8.74E-18 | 7.36E-17 |
| ENSG00000105642 | KCNN1 | -1.69 | 0.28 | 3.25E-11 | 1.94E-10 |
| ENSG00000176659 | LINC02910 | -1.69 | 0.38 | 1.27E-07 | 5.84E-07 |
| ENSG00000169813 | HNRNPF | -1.68 | 0.04 | 0.00E+00 | 0.00E+00 |
| ENSG00000133985 | TTC9 | -1.68 | 0.05 | 1.08E-211 | 2.43E-209 |
| ENSG00000146670 | CDCA5 | -1.68 | 0.08 | 4.24E-109 | 2.75E-107 |
| ENSG00000132640 | BTBD3 | -1.68 | 0.08 | 1.42E-93 | 7.50E-92 |
| ENSG00000105889 | STEAP1B | -1.68 | 0.09 | 4.77E-74 | 1.78E-72 |
| ENSG00000102678 | FGF9 | -1.68 | 0.14 | 1.73E-33 | 2.60E-32 |
| ENSG00000173918 | C1QTNF1 | -1.68 | 0.19 | 1.34E-19 | 1.22E-18 |
| ENSG00000157119 | KLHL40 | -1.68 | 0.22 | 9.97E-16 | 7.62E-15 |
| ENSG00000174403 | CRMA | -1.68 | 0.28 | 5.22E-11 | 3.08E-10 |
| ENSG00000250360 | | -1.68 | 0.53 | 1.48E-05 | 5.63E-05 |
| ENSG00000165097 | KDM1B | -1.67 | 0.08 | 1.07E-99 | 6.20E-98 |
| ENSG00000166801 | FAM111A | -1.67 | 0.09 | 2.36E-82 | 1.02E-80 |
| ENSG00000011347 | SYT7 | -1.67 | 0.1 | 1.13E-67 | 3.69E-66 |
| ENSG00000197020 | ZNF100 | -1.67 | 0.1 | 2.35E-62 | 6.85E-61 |
| ENSG00000290555 | LINC00680 | -1.67 | 0.11 | 1.86E-51 | 4.32E-50 |
| ENSG00000100629 | CEP128 | -1.67 | 0.14 | 1.38E-32 | 2.01E-31 |
| ENSG00000188610 | FAM72B | -1.67 | 0.18 | 4.85E-21 | 4.71E-20 |
| ENSG00000229619 | MBNL1-AS1 | -1.67 | 0.33 | 6.81E-09 | 3.45E-08 |
| ENSG00000163072 | NOSTRIN | -1.67 | 0.4 | 4.14E-07 | 1.83E-06 |
| ENSG00000249738 | | -1.67 | 0.41 | 7.50E-07 | 3.22E-06 |
| ENSG00000274600 | RIMBP3B | -1.67 | 0.45 | 2.67E-06 | 1.09E-05 |
| ENSG00000116830 | TTF2 | -1.66 | 0.05 | 5.83E-255 | 1.98E-252 |
| ENSG00000165806 | CASP7 | -1.66 | 0.07 | 5.45E-141 | 5.27E-139 |
| ENSG00000158402 | CDC25C | -1.66 | 0.17 | 1.42E-23 | 1.52E-22 |

| | | | | | |
|---|---|---|---|---|---|
| ENSG00000169067 | ACTBL2 | -1.66 | 0.25 | 7.67E-13 | 5.07E-12 |
| ENSG00000230499 | | -1.66 | 0.29 | 1.45E-10 | 8.30E-10 |
| ENSG00000243978 | RTL9 | -1.66 | 0.44 | 1.99E-06 | 8.22E-06 |
| ENSG00000143324 | XPR1 | -1.65 | 0.05 | 3.35E-200 | 6.55E-198 |
| ENSG00000149503 | INCENP | -1.65 | 0.07 | 7.79E-136 | 7.10E-134 |
| ENSG00000156172 | CFAP418 | -1.65 | 0.18 | 8.03E-22 | 8.03E-21 |
| ENSG00000115221 | ITGB6 | -1.65 | 0.22 | 3.98E-16 | 3.11E-15 |
| ENSG00000274719 | | -1.65 | 0.78 | 1.48E-04 | 5.04E-04 |
| ENSG00000137124 | ALDH1B1 | -1.64 | 0.04 | 1.16E-304 | 6.08E-302 |
| ENSG00000064300 | NGFR | -1.64 | 0.06 | 8.96E-161 | 1.06E-158 |
| ENSG00000101335 | MYL9 | -1.64 | 0.06 | 2.29E-157 | 2.64E-155 |
| ENSG00000138182 | KIF20B | -1.64 | 0.08 | 8.09E-98 | 4.51E-96 |
| ENSG00000102743 | SLC25A15 | -1.64 | 0.1 | 5.11E-64 | 1.55E-62 |
| ENSG00000183798 | EMILIN3 | -1.64 | 0.16 | 2.84E-25 | 3.24E-24 |
| ENSG00000088325 | TPX2 | -1.63 | 0.05 | 1.96E-250 | 6.25E-248 |
| ENSG00000116171 | SCP2 | -1.63 | 0.05 | 2.70E-220 | 6.61E-218 |
| ENSG00000011426 | ANLN | -1.63 | 0.05 | 1.65E-209 | 3.64E-207 |
| ENSG00000053371 | AKR7A2 | -1.63 | 0.07 | 1.61E-119 | 1.18E-117 |
| ENSG00000095383 | TBC1D2 | -1.63 | 0.13 | 7.90E-38 | 1.35E-36 |
| ENSG00000119986 | AVPI1 | -1.63 | 0.16 | 4.56E-25 | 5.15E-24 |
| ENSG00000140534 | TICRR | -1.62 | 0.07 | 2.12E-118 | 1.52E-116 |
| ENSG00000173546 | CSPG4 | -1.62 | 0.08 | 1.28E-98 | 7.31E-97 |
| ENSG00000151116 | UEVLD | -1.62 | 0.09 | 1.69E-79 | 6.92E-78 |
| ENSG00000166377 | ATP9B | -1.62 | 0.1 | 1.86E-56 | 4.86E-55 |
| ENSG00000036448 | MYOM2 | -1.62 | 0.12 | 1.00E-41 | 1.89E-40 |
| ENSG00000180999 | C1orf105 | -1.62 | 0.18 | 1.66E-21 | 1.64E-20 |
| ENSG00000188015 | S100A3 | -1.62 | 0.2 | 3.81E-17 | 3.12E-16 |
| ENSG00000145861 | C1QTNF2 | -1.62 | 0.28 | 3.00E-10 | 1.68E-09 |
| ENSG00000018625 | ATP1A2 | -1.62 | 0.41 | 1.18E-06 | 4.99E-06 |
| ENSG00000125844 | RRBP1 | -1.61 | 0.05 | 2.28E-202 | 4.63E-200 |
| ENSG00000140525 | FANCI | -1.61 | 0.06 | 2.55E-154 | 2.80E-152 |
| ENSG00000112144 | CILK1 | -1.61 | 0.09 | 8.93E-74 | 3.32E-72 |
| ENSG00000182010 | RTKN2 | -1.61 | 0.15 | 3.24E-29 | 4.24E-28 |
| ENSG00000186994 | KANK3 | -1.61 | 0.27 | 6.77E-11 | 3.96E-10 |
| ENSG00000137809 | ITGA11 | -1.61 | 0.34 | 3.03E-08 | 1.47E-07 |
| ENSG00000167850 | CD300C | -1.61 | 0.43 | 2.67E-06 | 1.09E-05 |
| ENSG00000169258 | GPRIN1 | -1.6 | 0.1 | 2.25E-63 | 6.71E-62 |
| ENSG00000170779 | CDCA4 | -1.6 | 0.11 | 1.50E-47 | 3.20E-46 |
| ENSG00000101470 | TNNC2 | -1.6 | 0.11 | 1.71E-47 | 3.63E-46 |
| ENSG00000198937 | CCDC167 | -1.6 | 0.14 | 9.92E-34 | 1.50E-32 |
| ENSG00000102575 | ACP5 | -1.6 | 0.15 | 1.69E-28 | 2.15E-27 |
| ENSG00000156466 | GDF6 | -1.6 | 0.21 | 1.06E-15 | 8.12E-15 |
| ENSG00000198739 | LRRTM3 | -1.59 | 0.05 | 5.09E-260 | 1.78E-257 |
| ENSG00000105323 | HNRNPUL1 | -1.59 | 0.05 | 1.50E-193 | 2.69E-191 |
| ENSG00000107551 | RASSF4 | -1.59 | 0.07 | 8.64E-107 | 5.48E-105 |
| ENSG00000007129 | CEACAM21 | -1.59 | 0.07 | 3.02E-106 | 1.90E-104 |
| ENSG00000196584 | XRCC2 | -1.59 | 0.09 | 1.59E-66 | 5.08E-65 |
| ENSG00000130844 | ZNF331 | -1.59 | 0.12 | 2.75E-43 | 5.36E-42 |
| ENSG00000164978 | NUDT2 | -1.59 | 0.16 | 3.36E-25 | 3.81E-24 |
| ENSG00000166897 | ELFN2 | -1.59 | 0.6 | 8.01E-05 | 2.82E-04 |

| | | | | | |
|---|---|---|---|---|---|
| ENSG00000151388 | ADAMTS12 | -1.58 | 0.06 | 2.99E-140 | 2.84E-138 |
| ENSG00000205056 | LINC02397 | -1.58 | 0.24 | 2.45E-12 | 1.57E-11 |
| ENSG00000120457 | KCNJ5 | -1.58 | 0.61 | 1.01E-04 | 3.52E-04 |
| ENSG00000144824 | PHLDB2 | -1.57 | 0.05 | 1.55E-238 | 4.61E-236 |
| ENSG00000112183 | RBM24 | -1.57 | 0.06 | 3.74E-167 | 4.70E-165 |
| ENSG00000137807 | KIF23 | -1.57 | 0.06 | 7.27E-162 | 8.70E-160 |
| ENSG00000136982 | DSCC1 | -1.57 | 0.08 | 7.48E-91 | 3.72E-89 |
| ENSG00000130313 | PGLS | -1.57 | 0.08 | 3.29E-81 | 1.38E-79 |
| ENSG00000132613 | MTSS2 | -1.57 | 0.09 | 2.62E-75 | 9.98E-74 |
| ENSG00000249115 | HAUS5 | -1.57 | 0.12 | 1.07E-40 | 1.96E-39 |
| ENSG00000232334 | | -1.57 | 0.53 | 3.94E-05 | 1.43E-04 |
| ENSG00000102172 | SMS | -1.56 | 0.06 | 3.73E-141 | 3.62E-139 |
| ENSG00000101003 | GINS1 | -1.56 | 0.07 | 5.12E-116 | 3.53E-114 |
| ENSG00000110104 | CCDC86 | -1.56 | 0.07 | 3.47E-98 | 1.95E-96 |
| ENSG00000172159 | FRMD3 | -1.56 | 0.08 | 2.72E-81 | 1.14E-79 |
| ENSG00000125650 | PSPN | -1.56 | 0.08 | 5.91E-80 | 2.43E-78 |
| ENSG00000157045 | NTAN1 | -1.56 | 0.12 | 5.63E-43 | 1.09E-41 |
| ENSG00000185269 | NOTUM | -1.56 | 0.12 | 1.35E-39 | 2.41E-38 |
| ENSG00000091137 | SLC26A4 | -1.56 | 0.12 | 1.83E-39 | 3.24E-38 |
| ENSG00000188662 | H1-9P | -1.56 | 0.35 | 1.50E-07 | 6.88E-07 |
| ENSG00000163535 | SGO2 | -1.55 | 0.08 | 8.84E-78 | 3.52E-76 |
| ENSG00000149596 | JPH2 | -1.55 | 0.1 | 3.48E-58 | 9.39E-57 |
| ENSG00000158164 | TMSB15A | -1.55 | 0.13 | 7.85E-36 | 1.27E-34 |
| ENSG00000253636 | | -1.55 | 0.17 | 1.49E-20 | 1.42E-19 |
| ENSG00000107859 | PITX3 | -1.55 | 0.18 | 6.77E-20 | 6.26E-19 |
| ENSG00000106804 | C5 | -1.55 | 0.22 | 3.41E-14 | 2.43E-13 |
| ENSG00000146276 | GABRR1 | -1.55 | 0.25 | 2.10E-11 | 1.27E-10 |
| ENSG00000130943 | PKDREJ | -1.55 | 0.28 | 8.69E-10 | 4.71E-09 |
| ENSG00000147408 | CSGALNACT1 | -1.55 | 0.35 | 2.14E-07 | 9.66E-07 |
| ENSG00000127870 | RNF6 | -1.54 | 0.09 | 7.91E-68 | 2.58E-66 |
| ENSG00000175928 | LRRN1 | -1.54 | 0.11 | 1.95E-49 | 4.31E-48 |
| ENSG00000280202 | | -1.54 | 0.17 | 2.14E-21 | 2.11E-20 |
| ENSG00000143416 | SELENBP1 | -1.54 | 0.28 | 5.22E-10 | 2.88E-09 |
| ENSG00000277534 | | -1.54 | 0.28 | 5.97E-10 | 3.28E-09 |
| ENSG00000228998 | PERPP2 | -1.54 | 0.35 | 1.99E-07 | 9.04E-07 |
| ENSG00000130309 | COLGALT1 | -1.53 | 0.06 | 1.29E-133 | 1.15E-131 |
| ENSG00000117643 | MAN1C1 | -1.53 | 0.06 | 2.88E-133 | 2.54E-131 |
| ENSG00000163840 | DTX3L | -1.53 | 0.07 | 1.17E-115 | 8.05E-114 |
| ENSG00000086289 | EPDR1 | -1.53 | 0.07 | 1.50E-107 | 9.61E-106 |
| ENSG00000151422 | FER | -1.53 | 0.09 | 7.17E-73 | 2.60E-71 |
| ENSG00000175920 | DOK7 | -1.53 | 0.13 | 9.16E-34 | 1.38E-32 |
| ENSG00000142149 | HUNK | -1.52 | 0.05 | 4.92E-207 | 1.06E-204 |
| ENSG00000185900 | POMK | -1.52 | 0.07 | 3.26E-98 | 1.83E-96 |
| ENSG00000077152 | UBE2T | -1.52 | 0.07 | 1.02E-94 | 5.45E-93 |
| ENSG00000135643 | KCNMB4 | -1.52 | 0.08 | 1.78E-85 | 8.07E-84 |
| ENSG00000165194 | PCDH19 | -1.52 | 0.08 | 2.53E-78 | 1.02E-76 |
| ENSG00000170214 | ADRA1B | -1.52 | 0.1 | 1.35E-50 | 3.07E-49 |
| ENSG00000225964 | NRIR | -1.52 | 0.37 | 6.64E-07 | 2.87E-06 |
| ENSG00000259236 | GOLGA8VP | -1.52 | 0.58 | 1.08E-04 | 3.75E-04 |
| ENSG00000204889 | KRT40 | -1.52 | 0.94 | 2.01E-04 | 6.75E-04 |

| | | | | | |
|---|---|---|---|---|---|
| ENSG00000131724 | IL13RA1 | -1.51 | 0.06 | 2.19E-143 | 2.19E-141 |
| ENSG00000089693 | MLF2 | -1.51 | 0.07 | 3.86E-108 | 2.49E-106 |
| ENSG00000085117 | CD82 | -1.51 | 0.08 | 2.46E-82 | 1.06E-80 |
| ENSG00000152217 | SETBP1 | -1.51 | 0.11 | 4.71E-47 | 9.92E-46 |
| ENSG00000177602 | HASPIN | -1.51 | 0.11 | 5.23E-42 | 9.95E-41 |
| ENSG00000196074 | SYCP2 | -1.51 | 0.17 | 1.04E-20 | 9.96E-20 |
| ENSG00000138356 | AOX1 | -1.51 | 0.25 | 2.67E-11 | 1.61E-10 |
| ENSG00000248964 | | -1.51 | 0.32 | 6.49E-08 | 3.05E-07 |
| ENSG00000157570 | TSPAN18 | -1.51 | 0.58 | 1.21E-04 | 4.16E-04 |
| ENSG00000164794 | KCNV1 | -1.51 | 0.63 | 1.82E-04 | 6.14E-04 |
| ENSG00000135476 | ESPL1 | -1.5 | 0.08 | 1.68E-73 | 6.20E-72 |
| ENSG00000129480 | DTD2 | -1.5 | 0.1 | 5.81E-51 | 1.33E-49 |
| ENSG00000104313 | EYA1 | -1.5 | 0.11 | 6.64E-47 | 1.40E-45 |
| ENSG00000289194 | | -1.5 | 0.19 | 6.82E-17 | 5.54E-16 |
| ENSG00000231607 | DLEU2 | -1.5 | 0.21 | 3.44E-14 | 2.45E-13 |
| ENSG00000242689 | CNTF | -1.5 | 0.23 | 3.66E-12 | 2.32E-11 |
| ENSG00000267060 | PTGES3L | -1.5 | 0.3 | 1.56E-08 | 7.71E-08 |
| ENSG00000183134 | PTGDR2 | -1.5 | 0.33 | 1.52E-07 | 6.93E-07 |
| ENSG00000064655 | EYA2 | -1.5 | 0.66 | 2.40E-04 | 7.96E-04 |
| ENSG00000230834 | | -1.5 | 0.8 | 3.40E-04 | 1.10E-03 |
| ENSG00000117114 | ADGRL2 | -1.49 | 0.05 | 4.86E-175 | 6.88E-173 |
| ENSG00000145819 | ARHGAP26 | -1.49 | 0.07 | 5.27E-98 | 2.95E-96 |
| ENSG00000012048 | BRCA1 | -1.49 | 0.08 | 2.43E-75 | 9.28E-74 |
| ENSG00000157456 | CCNB2 | -1.49 | 0.09 | 1.19E-67 | 3.85E-66 |
| ENSG00000168530 | MYL1 | -1.49 | 0.09 | 4.11E-60 | 1.15E-58 |
| ENSG00000107960 | STN1 | -1.49 | 0.11 | 1.46E-45 | 2.97E-44 |
| ENSG00000198056 | PRIM1 | -1.49 | 0.14 | 2.85E-28 | 3.59E-27 |
| ENSG00000100867 | DHRS2 | -1.49 | 0.21 | 3.11E-14 | 2.23E-13 |
| ENSG00000158008 | EXTL1 | -1.49 | 0.21 | 3.40E-14 | 2.42E-13 |
| ENSG00000081248 | CACNA1S | -1.49 | 0.23 | 3.31E-12 | 2.11E-11 |
| ENSG00000128408 | RIBC2 | -1.49 | 0.3 | 2.26E-08 | 1.10E-07 |
| ENSG00000100628 | ASB2 | -1.49 | 0.7 | 3.11E-04 | 1.01E-03 |
| ENSG00000124942 | AHNAK | -1.48 | 0.04 | 2.97E-251 | 9.84E-249 |
| ENSG00000139433 | GLTP | -1.48 | 0.05 | 2.57E-182 | 4.13E-180 |
| ENSG00000111913 | RIPOR2 | -1.48 | 0.06 | 1.01E-154 | 1.12E-152 |
| ENSG00000160867 | FGFR4 | -1.48 | 0.07 | 1.04E-97 | 5.82E-96 |
| ENSG00000180209 | MYL11 | -1.48 | 0.07 | 1.44E-94 | 7.69E-93 |
| ENSG00000111077 | TNS2 | -1.48 | 0.08 | 1.23E-80 | 5.11E-79 |
| ENSG00000109265 | CRACD | -1.48 | 0.08 | 1.05E-78 | 4.23E-77 |
| ENSG00000183763 | TRAIP | -1.48 | 0.12 | 9.77E-38 | 1.66E-36 |
| ENSG00000165606 | DRGX | -1.48 | 0.17 | 2.71E-19 | 2.45E-18 |
| ENSG00000178700 | DHFR2 | -1.48 | 0.19 | 1.33E-16 | 1.06E-15 |
| ENSG00000287559 | | -1.48 | 0.19 | 5.52E-16 | 4.29E-15 |
| ENSG00000235387 | SPAAR | -1.48 | 0.44 | 1.43E-05 | 5.45E-05 |
| ENSG00000239455 | PTPN11P1 | -1.48 | 0.45 | 1.83E-05 | 6.90E-05 |
| ENSG00000172927 | MYEOV | -1.48 | 0.52 | 7.26E-05 | 2.57E-04 |
| ENSG00000231419 | LINC00689 | -1.48 | 0.53 | 7.45E-05 | 2.63E-04 |
| ENSG00000179104 | TMTC2 | -1.47 | 0.06 | 1.06E-141 | 1.03E-139 |
| ENSG00000171792 | RHNO1 | -1.47 | 0.07 | 6.66E-93 | 3.50E-91 |
| ENSG00000137726 | FXYD6 | -1.47 | 0.1 | 2.45E-53 | 5.98E-52 |

| | | | | | |
|---|---|---|---|---|---|
| ENSG00000051180 | RAD51 | -1.47 | 0.12 | 5.11E-36 | 8.30E-35 |
| ENSG00000162946 | DISC1 | -1.47 | 0.12 | 6.93E-35 | 1.08E-33 |
| ENSG00000163517 | HDAC11 | -1.47 | 0.13 | 1.92E-30 | 2.62E-29 |
| ENSG00000131969 | ABHD12B | -1.47 | 0.2 | 2.00E-14 | 1.44E-13 |
| ENSG00000153064 | BANK1 | -1.47 | 0.22 | 1.34E-12 | 8.71E-12 |
| ENSG00000174059 | CD34 | -1.47 | 0.26 | 3.31E-10 | 1.85E-09 |
| ENSG00000166159 | LRTM2 | -1.47 | 0.46 | 2.41E-05 | 8.96E-05 |
| ENSG00000279622 | | -1.47 | 0.92 | 2.72E-04 | 8.98E-04 |
| ENSG00000131475 | VPS25 | -1.46 | 0.06 | 1.42E-150 | 1.50E-148 |
| ENSG00000167693 | NXN | -1.46 | 0.06 | 4.56E-113 | 3.08E-111 |
| ENSG00000137868 | STRA6 | -1.46 | 0.07 | 6.22E-103 | 3.74E-101 |
| ENSG00000184985 | SORCS2 | -1.46 | 0.07 | 4.22E-102 | 2.51E-100 |
| ENSG00000138346 | DNA2 | -1.46 | 0.07 | 2.12E-96 | 1.16E-94 |
| ENSG00000161381 | PLXDC1 | -1.46 | 0.14 | 8.08E-26 | 9.38E-25 |
| ENSG00000260877 | | -1.46 | 0.25 | 2.52E-10 | 1.42E-09 |
| ENSG00000135052 | GOLM1 | -1.45 | 0.05 | 3.01E-223 | 7.57E-221 |
| ENSG00000119906 | SLF2 | -1.45 | 0.06 | 1.62E-112 | 1.09E-110 |
| ENSG00000088298 | EDEM2 | -1.45 | 0.08 | 6.76E-78 | 2.69E-76 |
| ENSG00000157214 | STEAP2 | -1.45 | 0.1 | 6.40E-52 | 1.51E-50 |
| ENSG00000275793 | RIMBP3 | -1.45 | 0.19 | 1.73E-15 | 1.31E-14 |
| ENSG00000174370 | KCNJ5-AS1 | -1.45 | 0.21 | 1.27E-13 | 8.77E-13 |
| ENSG00000134668 | SPOCD1 | -1.45 | 0.25 | 2.91E-10 | 1.63E-09 |
| ENSG00000147649 | MTDH | -1.44 | 0.04 | 1.33E-275 | 5.51E-273 |
| ENSG00000008311 | AASS | -1.44 | 0.06 | 1.20E-122 | 9.21E-121 |
| ENSG00000106785 | TRIM14 | -1.44 | 0.08 | 2.01E-77 | 7.98E-76 |
| ENSG00000143603 | KCNN3 | -1.44 | 0.09 | 4.23E-62 | 1.23E-60 |
| ENSG00000151413 | NUBPL | -1.44 | 0.1 | 6.99E-45 | 1.40E-43 |
| ENSG00000172731 | LRRC20 | -1.44 | 0.11 | 1.31E-37 | 2.22E-36 |
| ENSG00000141401 | IMPA2 | -1.44 | 0.11 | 1.42E-37 | 2.39E-36 |
| ENSG00000163634 | THOC7 | -1.44 | 0.12 | 1.28E-36 | 2.11E-35 |
| ENSG00000203867 | RBM20 | -1.44 | 0.15 | 5.13E-22 | 5.17E-21 |
| ENSG00000227992 | | -1.44 | 0.19 | 2.18E-15 | 1.65E-14 |
| ENSG00000227877 | MRLN | -1.44 | 0.22 | 1.23E-12 | 8.01E-12 |
| ENSG00000135604 | STX11 | -1.44 | 0.22 | 3.24E-12 | 2.06E-11 |
| ENSG00000162391 | FAM151A | -1.44 | 0.35 | 7.57E-07 | 3.25E-06 |
| ENSG00000288071 | | -1.44 | 0.99 | 1.87E-04 | 6.28E-04 |
| ENSG00000112584 | FAM120B | -1.43 | 0.05 | 4.69E-210 | 1.04E-207 |
| ENSG00000197712 | FAM114A1 | -1.43 | 0.06 | 7.28E-125 | 5.65E-123 |
| ENSG00000068489 | PRR11 | -1.43 | 0.07 | 1.18E-100 | 6.85E-99 |
| ENSG00000145332 | KLHL8 | -1.43 | 0.08 | 2.58E-79 | 1.05E-77 |
| ENSG00000213853 | EMP2 | -1.43 | 0.1 | 5.61E-51 | 1.29E-49 |
| ENSG00000142661 | MYOM3 | -1.43 | 0.22 | 4.40E-12 | 2.77E-11 |
| ENSG00000109667 | SLC2A9 | -1.43 | 0.44 | 2.61E-05 | 9.66E-05 |
| ENSG00000162490 | DRAXIN | -1.43 | 0.56 | 1.68E-04 | 5.70E-04 |
| ENSG00000137693 | YAP1 | -1.42 | 0.05 | 1.31E-182 | 2.11E-180 |
| ENSG00000137710 | RDX | -1.42 | 0.05 | 1.04E-151 | 1.11E-149 |
| ENSG00000278259 | MYO19 | -1.42 | 0.06 | 4.90E-129 | 4.13E-127 |
| ENSG00000117399 | CDC20 | -1.42 | 0.07 | 3.28E-103 | 1.98E-101 |
| ENSG00000075234 | TTC38 | -1.42 | 0.09 | 2.37E-60 | 6.70E-59 |
| ENSG00000239789 | MRPS17 | -1.42 | 0.09 | 1.25E-53 | 3.09E-52 |

| | | | | | |
|---|---|---|---|---|---|
| ENSG00000126822 | PLEKHG3 | -1.42 | 0.1 | 1.51E-50 | 3.45E-49 |
| ENSG00000128274 | A4GALT | -1.42 | 0.11 | 1.01E-41 | 1.91E-40 |
| ENSG00000186638 | KIF24 | -1.42 | 0.12 | 2.15E-35 | 3.42E-34 |
| ENSG00000175305 | CCNE2 | -1.42 | 0.19 | 9.09E-15 | 6.68E-14 |
| ENSG00000265485 | LINC01915 | -1.42 | 0.27 | 4.72E-09 | 2.41E-08 |
| ENSG00000147166 | ITGB1BP2 | -1.42 | 0.39 | 6.18E-06 | 2.44E-05 |
| ENSG00000184363 | PKP3 | -1.42 | 0.65 | 3.94E-04 | 1.27E-03 |
| ENSG00000111252 | SH2B3 | -1.41 | 0.04 | 1.07E-241 | 3.31E-239 |
| ENSG00000154127 | UBASH3B | -1.41 | 0.05 | 3.69E-173 | 5.14E-171 |
| ENSG00000135048 | CEMIP2 | -1.41 | 0.05 | 3.30E-171 | 4.46E-169 |
| ENSG00000167670 | CHAF1A | -1.41 | 0.06 | 5.63E-110 | 3.68E-108 |
| ENSG00000138594 | TMOD3 | -1.41 | 0.07 | 2.13E-92 | 1.10E-90 |
| ENSG00000121766 | ZCCHC17 | -1.41 | 0.07 | 4.75E-83 | 2.09E-81 |
| ENSG00000105173 | CCNE1 | -1.41 | 0.08 | 4.07E-77 | 1.61E-75 |
| ENSG00000148082 | SHC3 | -1.41 | 0.09 | 6.84E-63 | 2.02E-61 |
| ENSG00000275993 | | -1.41 | 0.11 | 2.46E-42 | 4.71E-41 |
| ENSG00000168874 | ATOH8 | -1.41 | 0.11 | 9.65E-42 | 1.82E-40 |
| ENSG00000253669 | GASAL1 | -1.41 | 0.13 | 6.73E-28 | 8.38E-27 |
| ENSG00000213062 | | -1.41 | 0.14 | 2.94E-26 | 3.46E-25 |
| ENSG00000253377 | | -1.41 | 0.16 | 4.62E-20 | 4.30E-19 |
| ENSG00000100479 | POLE2 | -1.41 | 0.16 | 1.09E-19 | 9.99E-19 |
| ENSG00000137135 | ARHGEF39 | -1.41 | 0.21 | 2.49E-13 | 1.69E-12 |
| ENSG00000152078 | TLCD4 | -1.41 | 0.24 | 1.74E-10 | 9.93E-10 |
| ENSG00000170122 | FOXD4 | -1.41 | 0.31 | 1.07E-07 | 4.97E-07 |
| ENSG00000258502 | LINC02290 | -1.41 | 0.32 | 2.23E-07 | 1.01E-06 |
| ENSG00000038427 | VCAN | -1.4 | 0.05 | 2.10E-152 | 2.23E-150 |
| ENSG00000147601 | TERF1 | -1.4 | 0.05 | 9.88E-148 | 1.01E-145 |
| ENSG00000168496 | FEN1 | -1.4 | 0.06 | 1.42E-132 | 1.25E-130 |
| ENSG00000150687 | PRSS23 | -1.4 | 0.06 | 2.00E-119 | 1.46E-117 |
| ENSG00000280219 | | -1.4 | 0.08 | 1.45E-69 | 4.95E-68 |
| ENSG00000187672 | ERC2 | -1.4 | 0.11 | 1.76E-36 | 2.88E-35 |
| ENSG00000171812 | COL8A2 | -1.4 | 0.2 | 3.59E-14 | 2.56E-13 |
| ENSG00000264345 | LINC01894 | -1.4 | 0.21 | 2.44E-12 | 1.57E-11 |
| ENSG00000196196 | HRCT1 | -1.4 | 0.33 | 5.35E-07 | 2.34E-06 |
| ENSG00000108798 | ABI3 | -1.4 | 0.4 | 1.29E-05 | 4.94E-05 |
| ENSG00000168079 | SCARA5 | -1.4 | 0.42 | 2.19E-05 | 8.20E-05 |
| ENSG00000204876 | | -1.4 | 0.52 | 1.21E-04 | 4.17E-04 |
| ENSG00000149633 | KIAA1755 | -1.4 | 0.52 | 1.39E-04 | 4.75E-04 |
| ENSG00000149136 | SSRP1 | -1.39 | 0.04 | 9.92E-221 | 2.45E-218 |
| ENSG00000137486 | ARRB1 | -1.39 | 0.05 | 9.23E-183 | 1.50E-180 |
| ENSG00000119408 | NEK6 | -1.39 | 0.06 | 5.63E-115 | 3.86E-113 |
| ENSG00000172331 | BPGM | -1.39 | 0.1 | 6.72E-46 | 1.38E-44 |
| ENSG00000254685 | FPGT | -1.39 | 0.12 | 5.80E-31 | 8.07E-30 |
| ENSG00000166343 | MSS51 | -1.39 | 0.14 | 1.03E-23 | 1.11E-22 |
| ENSG00000135537 | AFG1L | -1.39 | 0.15 | 3.42E-22 | 3.47E-21 |
| ENSG00000241322 | FBXW10B | -1.39 | 0.17 | 1.45E-17 | 1.21E-16 |
| ENSG00000132016 | BRME1 | -1.39 | 0.19 | 7.32E-15 | 5.40E-14 |
| ENSG00000141574 | SECTM1 | -1.39 | 0.21 | 3.89E-13 | 2.62E-12 |
| ENSG00000181350 | LRRC75A | -1.39 | 0.21 | 1.97E-12 | 1.27E-11 |
| ENSG00000232774 | LINC03033 | -1.39 | 0.28 | 3.09E-08 | 1.49E-07 |

| | | | | | |
|---|---|---|---|---|---|
| ENSG00000174885 | NLRP6 | -1.39 | 0.47 | 6.88E-05 | 2.44E-04 |
| ENSG00000223511 | LINC02968 | -1.39 | 0.62 | 3.90E-04 | 1.26E-03 |
| ENSG00000119689 | DLST | -1.38 | 0.04 | 8.69E-217 | 2.07E-214 |
| ENSG00000124126 | PREX1 | -1.38 | 0.05 | 1.40E-199 | 2.69E-197 |
| ENSG00000172403 | SYNPO2 | -1.38 | 0.06 | 4.73E-126 | 3.75E-124 |
| ENSG00000138496 | PARP9 | -1.38 | 0.08 | 2.26E-66 | 7.19E-65 |
| ENSG00000146263 | MMS22L | -1.38 | 0.08 | 4.17E-64 | 1.27E-62 |
| ENSG00000185567 | AHNAK2 | -1.38 | 0.09 | 2.96E-60 | 8.34E-59 |
| ENSG00000130528 | HRC | -1.38 | 0.1 | 1.01E-44 | 2.03E-43 |
| ENSG00000149929 | HIRIP3 | -1.38 | 0.12 | 1.46E-34 | 2.26E-33 |
| ENSG00000103995 | CEP152 | -1.38 | 0.12 | 8.29E-30 | 1.11E-28 |
| ENSG00000290994 | FAM86C2P | -1.38 | 0.26 | 4.34E-09 | 2.23E-08 |
| ENSG00000286878 | | -1.38 | 0.4 | 1.40E-05 | 5.33E-05 |
| ENSG00000271524 | BNIP3P17 | -1.38 | 0.63 | 4.23E-04 | 1.36E-03 |
| ENSG00000198883 | PNMA5 | -1.37 | 0.06 | 8.26E-137 | 7.70E-135 |
| ENSG00000110063 | DCPS | -1.37 | 0.06 | 3.52E-110 | 2.31E-108 |
| ENSG00000251381 | LINC00958 | -1.37 | 0.06 | 5.42E-108 | 3.48E-106 |
| ENSG00000180198 | RCC1 | -1.37 | 0.07 | 8.30E-95 | 4.47E-93 |
| ENSG00000005513 | SOX8 | -1.37 | 0.09 | 8.60E-55 | 2.18E-53 |
| ENSG00000103175 | WFDC1 | -1.37 | 0.21 | 5.00E-12 | 3.14E-11 |
| ENSG00000250132 | | -1.37 | 0.21 | 5.06E-12 | 3.18E-11 |
| ENSG00000134955 | SLC37A2 | -1.37 | 0.28 | 2.84E-08 | 1.38E-07 |
| ENSG00000168214 | RBPJ | -1.36 | 0.05 | 2.05E-138 | 1.94E-136 |
| ENSG00000082153 | BZW1 | -1.36 | 0.06 | 7.41E-118 | 5.29E-116 |
| ENSG00000165934 | CPSF2 | -1.36 | 0.06 | 8.95E-111 | 5.92E-109 |
| ENSG00000228716 | DHFR | -1.36 | 0.06 | 1.09E-103 | 6.64E-102 |
| ENSG00000196810 | CTBP1-DT | -1.36 | 0.08 | 6.67E-75 | 2.51E-73 |
| ENSG00000142733 | MAP3K6 | -1.36 | 0.09 | 2.00E-50 | 4.54E-49 |
| ENSG00000158715 | SLC45A3 | -1.36 | 0.13 | 3.10E-28 | 3.90E-27 |
| ENSG00000006534 | ALDH3B1 | -1.36 | 0.19 | 1.73E-14 | 1.25E-13 |
| ENSG00000273226 | | -1.36 | 0.34 | 1.88E-06 | 7.76E-06 |
| ENSG00000151702 | FLI1 | -1.36 | 0.43 | 3.33E-05 | 1.22E-04 |
| ENSG00000229459 | | -1.36 | 0.79 | 7.28E-04 | 2.26E-03 |
| ENSG00000223949 | ROR1-AS1 | -1.36 | 0.77 | 7.34E-04 | 2.28E-03 |
| ENSG00000187979 | FAM230D | -1.36 | 0.74 | 7.38E-04 | 2.29E-03 |
| ENSG00000105341 | DMAC2 | -1.35 | 0.06 | 1.20E-118 | 8.62E-117 |
| ENSG00000140564 | FURIN | -1.35 | 0.09 | 1.46E-51 | 3.40E-50 |
| ENSG00000114854 | TNNC1 | -1.35 | 0.13 | 5.87E-27 | 7.10E-26 |
| ENSG00000183801 | OLFML1 | -1.35 | 0.16 | 6.78E-19 | 5.98E-18 |
| ENSG00000167680 | SEMA6B | -1.35 | 0.16 | 7.49E-19 | 6.60E-18 |
| ENSG00000255129 | TTC12-DT | -1.35 | 0.52 | 2.03E-04 | 6.81E-04 |
| ENSG00000060138 | YBX3 | -1.34 | 0.05 | 1.49E-185 | 2.47E-183 |
| ENSG00000161800 | RACGAP1 | -1.34 | 0.07 | 6.68E-78 | 2.67E-76 |
| ENSG00000147536 | GINS4 | -1.34 | 0.08 | 3.52E-69 | 1.18E-67 |
| ENSG00000111684 | LPCAT3 | -1.34 | 0.08 | 1.72E-58 | 4.66E-57 |
| ENSG00000223459 | TCAF1P1 | -1.34 | 0.1 | 1.52E-41 | 2.85E-40 |
| ENSG00000102384 | CENPI | -1.34 | 0.11 | 2.41E-36 | 3.94E-35 |
| ENSG00000149548 | CCDC15 | -1.34 | 0.11 | 2.75E-34 | 4.21E-33 |
| ENSG00000113356 | POLR3G | -1.34 | 0.11 | 1.27E-33 | 1.91E-32 |
| ENSG00000115884 | SDC1 | -1.34 | 0.12 | 1.31E-30 | 1.81E-29 |

| | | | | | |
|---|---|---|---|---|---|
| ENSG00000198892 | SHISA4 | -1.34 | 0.12 | 7.21E-29 | 9.34E-28 |
| ENSG00000196811 | CHRNG | -1.34 | 0.13 | 7.90E-28 | 9.83E-27 |
| ENSG00000123453 | SARDH | -1.34 | 0.15 | 2.59E-21 | 2.55E-20 |
| ENSG00000121440 | PDZRN3 | -1.34 | 0.18 | 1.28E-15 | 9.71E-15 |
| ENSG00000250007 | GJD2-DT | -1.34 | 0.23 | 2.56E-10 | 1.44E-09 |
| ENSG00000249863 | | -1.34 | 0.3 | 2.94E-07 | 1.31E-06 |
| ENSG00000222032 | | -1.34 | 0.32 | 9.63E-07 | 4.09E-06 |
| ENSG00000104059 | ENTREP2 | -1.34 | 0.33 | 1.19E-06 | 5.04E-06 |
| ENSG00000279386 | | -1.34 | 0.47 | 1.09E-04 | 3.78E-04 |
| ENSG00000079112 | CDH17 | -1.34 | 0.5 | 1.65E-04 | 5.57E-04 |
| ENSG00000206754 | SNORD101 | -1.34 | 0.5 | 1.71E-04 | 5.76E-04 |
| ENSG00000177354 | C10orf71 | -1.34 | 0.51 | 1.78E-04 | 5.99E-04 |
| ENSG00000120645 | IQSEC3 | -1.34 | 0.53 | 2.42E-04 | 8.04E-04 |
| ENSG00000162511 | LAPTM5 | -1.34 | 0.59 | 4.15E-04 | 1.33E-03 |
| ENSG00000224982 | TMEM233 | -1.33 | 0.05 | 3.63E-154 | 3.98E-152 |
| ENSG00000025293 | PHF20 | -1.33 | 0.05 | 3.55E-146 | 3.62E-144 |
| ENSG00000166444 | DENND2B | -1.33 | 0.06 | 4.14E-128 | 3.43E-126 |
| ENSG00000196924 | FLNA | -1.33 | 0.06 | 5.09E-124 | 3.92E-122 |
| ENSG00000168228 | ZCCHC4 | -1.33 | 0.08 | 2.36E-60 | 6.68E-59 |
| ENSG00000157823 | AP3S2 | -1.33 | 0.08 | 1.82E-58 | 4.92E-57 |
| ENSG00000262919 | CCNQ | -1.33 | 0.09 | 3.34E-53 | 8.14E-52 |
| ENSG00000123374 | CDK2 | -1.33 | 0.1 | 1.67E-40 | 3.05E-39 |
| ENSG00000168334 | XIRP1 | -1.33 | 0.1 | 3.57E-39 | 6.30E-38 |
| ENSG00000197046 | SIGLEC15 | -1.33 | 0.12 | 1.90E-30 | 2.60E-29 |
| ENSG00000235859 | | -1.33 | 0.16 | 3.61E-19 | 3.23E-18 |
| ENSG00000260398 | | -1.33 | 0.16 | 9.57E-18 | 8.05E-17 |
| ENSG00000133256 | PDE6B | -1.33 | 0.2 | 2.33E-12 | 1.50E-11 |
| ENSG00000124701 | APOBEC2 | -1.33 | 0.3 | 3.30E-07 | 1.47E-06 |
| ENSG00000110195 | FOLR1 | -1.33 | 0.4 | 2.79E-05 | 1.03E-04 |
| ENSG00000278214 | LINC02139 | -1.33 | 0.44 | 7.14E-05 | 2.53E-04 |
| ENSG00000180712 | LINC02363 | -1.33 | 0.55 | 3.21E-04 | 1.05E-03 |
| ENSG00000010030 | ETV7 | -1.33 | 0.78 | 8.44E-04 | 2.60E-03 |
| ENSG00000135902 | CHRND | -1.32 | 0.04 | 1.29E-289 | 5.94E-287 |
| ENSG00000130707 | ASS1 | -1.32 | 0.04 | 9.28E-216 | 2.17E-213 |
| ENSG00000153179 | RASSF3 | -1.32 | 0.05 | 2.62E-153 | 2.83E-151 |
| ENSG00000158623 | COPG2 | -1.32 | 0.06 | 6.97E-99 | 3.98E-97 |
| ENSG00000047644 | WWC3 | -1.32 | 0.07 | 7.16E-92 | 3.66E-90 |
| ENSG00000170379 | TCAF2 | -1.32 | 0.07 | 4.66E-81 | 1.95E-79 |
| ENSG00000151692 | RNF144A | -1.32 | 0.07 | 1.14E-78 | 4.61E-77 |
| ENSG00000055118 | KCNH2 | -1.32 | 0.1 | 6.36E-41 | 1.18E-39 |
| ENSG00000170681 | CAVIN4 | -1.32 | 0.12 | 3.57E-30 | 4.83E-29 |
| ENSG00000005189 | REXO5 | -1.32 | 0.21 | 8.03E-12 | 4.98E-11 |
| ENSG00000287707 | | -1.32 | 0.28 | 5.67E-08 | 2.68E-07 |
| ENSG00000184613 | NELL2 | -1.32 | 0.3 | 4.81E-07 | 2.11E-06 |
| ENSG00000103489 | XYLT1 | -1.32 | 0.35 | 4.46E-06 | 1.78E-05 |
| ENSG00000241634 | RPL13P8 | -1.32 | 0.39 | 1.68E-05 | 6.36E-05 |
| ENSG00000269404 | SPIB | -1.32 | 0.52 | 2.45E-04 | 8.11E-04 |
| ENSG00000074590 | NUAK1 | -1.31 | 0.04 | 2.43E-260 | 8.71E-258 |
| ENSG00000174444 | RPL4 | -1.31 | 0.04 | 7.73E-211 | 1.73E-208 |
| ENSG00000113580 | NR3C1 | -1.31 | 0.06 | 1.34E-125 | 1.06E-123 |

| | | | | | |
|---|---|---|---|---|---|
| ENSG00000158457 | TSPAN33 | -1.31 | 0.06 | 3.04E-115 | 2.09E-113 |
| ENSG00000162430 | SELENON | -1.31 | 0.07 | 1.11E-76 | 4.36E-75 |
| ENSG00000213516 | RBMXL1 | -1.31 | 0.08 | 1.55E-66 | 4.94E-65 |
| ENSG00000138376 | BARD1 | -1.31 | 0.08 | 8.11E-63 | 2.39E-61 |
| ENSG00000103196 | CRISPLD2 | -1.31 | 0.14 | 6.18E-22 | 6.21E-21 |
| ENSG00000279443 | | -1.31 | 0.22 | 4.08E-11 | 2.42E-10 |
| ENSG00000203999 | LINC01270 | -1.31 | 0.26 | 1.62E-08 | 7.96E-08 |
| ENSG00000289504 | | -1.31 | 0.3 | 5.28E-07 | 2.31E-06 |
| ENSG00000162520 | SYNC | -1.31 | 0.32 | 1.33E-06 | 5.57E-06 |
| ENSG00000158042 | MRPL17 | -1.3 | 0.06 | 1.80E-125 | 1.41E-123 |
| ENSG00000164109 | MAD2L1 | -1.3 | 0.06 | 1.89E-100 | 1.10E-98 |
| ENSG00000132128 | LRRC41 | -1.3 | 0.06 | 1.67E-98 | 9.47E-97 |
| ENSG00000157216 | SSBP3 | -1.3 | 0.07 | 3.37E-90 | 1.65E-88 |
| ENSG00000134452 | FBH1 | -1.3 | 0.07 | 8.63E-82 | 3.66E-80 |
| ENSG00000160712 | IL6R | -1.3 | 0.07 | 5.83E-73 | 2.13E-71 |
| ENSG00000165495 | PKNOX2 | -1.3 | 0.07 | 5.11E-72 | 1.83E-70 |
| ENSG00000164976 | MYORG | -1.3 | 0.07 | 1.49E-68 | 4.93E-67 |
| ENSG00000138316 | ADAMTS14 | -1.3 | 0.08 | 2.16E-66 | 6.87E-65 |
| ENSG00000151502 | VPS26B | -1.3 | 0.08 | 1.89E-63 | 5.67E-62 |
| ENSG00000169689 | CENPX | -1.3 | 0.1 | 1.49E-43 | 2.92E-42 |
| ENSG00000167513 | CDT1 | -1.3 | 0.12 | 8.58E-30 | 1.15E-28 |
| ENSG00000065534 | MYLK | -1.3 | 0.2 | 5.09E-12 | 3.20E-11 |
| ENSG00000171094 | ALK | -1.3 | 0.31 | 8.71E-07 | 3.72E-06 |
| ENSG00000224609 | FGGY-DT | -1.3 | 0.45 | 1.13E-04 | 3.91E-04 |
| ENSG00000167641 | PPP1R14A | -1.3 | 0.48 | 1.67E-04 | 5.65E-04 |
| ENSG00000075702 | WDR62 | -1.29 | 0.08 | 1.43E-58 | 3.89E-57 |
| ENSG00000076770 | MBNL3 | -1.29 | 0.11 | 1.19E-32 | 1.73E-31 |
| ENSG00000113555 | PCDH12 | -1.29 | 0.12 | 1.10E-29 | 1.47E-28 |
| ENSG00000224273 | RABGEF1P3 | -1.29 | 0.35 | 6.94E-06 | 2.73E-05 |
| ENSG00000183150 | GPR19 | -1.29 | 0.44 | 1.02E-04 | 3.54E-04 |
| ENSG00000227482 | | -1.29 | 0.51 | 2.73E-04 | 8.99E-04 |
| ENSG00000273056 | | -1.29 | 0.93 | 5.11E-04 | 1.63E-03 |
| ENSG00000198420 | TCAF1 | -1.28 | 0.04 | 3.69E-188 | 6.27E-186 |
| ENSG00000136271 | DDX56 | -1.28 | 0.04 | 3.18E-184 | 5.23E-182 |
| ENSG00000165661 | QSOX2 | -1.28 | 0.06 | 2.22E-119 | 1.62E-117 |
| ENSG00000133065 | SLC41A1 | -1.28 | 0.06 | 1.74E-110 | 1.15E-108 |
| ENSG00000113273 | ARSB | -1.28 | 0.06 | 3.71E-107 | 2.37E-105 |
| ENSG00000184992 | BRI3BP | -1.28 | 0.07 | 6.01E-67 | 1.93E-65 |
| ENSG00000187736 | NHEJ1 | -1.28 | 0.08 | 1.07E-53 | 2.64E-52 |
| ENSG00000139364 | TMEM132B | -1.28 | 0.1 | 2.35E-42 | 4.50E-41 |
| ENSG00000129521 | EGLN3 | -1.28 | 0.1 | 1.70E-41 | 3.18E-40 |
| ENSG00000182858 | ALG12 | -1.28 | 0.11 | 1.59E-34 | 2.44E-33 |
| ENSG00000134574 | DDB2 | -1.28 | 0.11 | 3.58E-30 | 4.84E-29 |
| ENSG00000142178 | SIK1 | -1.28 | 0.12 | 6.81E-27 | 8.22E-26 |
| ENSG00000171860 | C3AR1 | -1.28 | 0.19 | 9.54E-13 | 6.27E-12 |
| ENSG00000179841 | AKAP5 | -1.28 | 0.25 | 1.11E-08 | 5.57E-08 |
| ENSG00000147082 | CCNB3 | -1.28 | 0.3 | 8.25E-07 | 3.53E-06 |
| ENSG00000124839 | RAB17 | -1.28 | 0.5 | 2.57E-04 | 8.49E-04 |
| ENSG00000261371 | PECAM1 | -1.28 | 0.5 | 2.95E-04 | 9.65E-04 |
| ENSG00000106078 | COBL | -1.27 | 0.05 | 8.97E-129 | 7.53E-127 |

| | | | | | |
|---|---|---|---|---|---|
| ENSG00000135679 | MDM2 | -1.27 | 0.05 | 7.82E-120 | 5.75E-118 |
| ENSG00000108064 | TFAM | -1.27 | 0.06 | 7.47E-117 | 5.26E-115 |
| ENSG00000123643 | SLC36A1 | -1.27 | 0.06 | 4.75E-106 | 2.98E-104 |
| ENSG00000146122 | DAAM2 | -1.27 | 0.06 | 2.80E-99 | 1.61E-97 |
| ENSG00000114450 | GNB4 | -1.27 | 0.06 | 1.45E-98 | 8.26E-97 |
| ENSG00000114346 | ECT2 | -1.27 | 0.06 | 1.54E-98 | 8.72E-97 |
| ENSG00000094804 | CDC6 | -1.27 | 0.06 | 7.30E-96 | 3.97E-94 |
| ENSG00000144354 | CDCA7 | -1.27 | 0.06 | 6.74E-87 | 3.07E-85 |
| ENSG00000107105 | ELAVL2 | -1.27 | 0.1 | 2.60E-42 | 4.98E-41 |
| ENSG00000149177 | PTPRJ | -1.27 | 0.11 | 1.57E-34 | 2.42E-33 |
| ENSG00000204060 | FOXO6 | -1.27 | 0.13 | 1.90E-24 | 2.10E-23 |
| ENSG00000169683 | LRRC45 | -1.27 | 0.15 | 8.70E-20 | 8.02E-19 |
| ENSG00000113790 | EHHADH | -1.27 | 0.19 | 1.22E-12 | 7.94E-12 |
| ENSG00000139508 | SLC46A3 | -1.27 | 0.26 | 3.10E-08 | 1.50E-07 |
| ENSG00000138100 | TRIM54 | -1.27 | 0.27 | 8.62E-08 | 4.02E-07 |
| ENSG00000182873 | PRKCZ-AS1 | -1.27 | 0.3 | 6.16E-07 | 2.67E-06 |
| ENSG00000261780 | LINC02582 | -1.27 | 0.81 | 1.07E-03 | 3.24E-03 |
| ENSG00000142230 | SAE1 | -1.26 | 0.04 | 7.28E-179 | 1.09E-176 |
| ENSG00000154380 | ENAH | -1.26 | 0.05 | 4.66E-170 | 6.13E-168 |
| ENSG00000144320 | LNPK | -1.26 | 0.06 | 3.48E-89 | 1.68E-87 |
| ENSG00000189308 | LIN54 | -1.26 | 0.08 | 9.79E-63 | 2.88E-61 |
| ENSG00000188266 | HYKK | -1.26 | 0.11 | 5.10E-32 | 7.31E-31 |
| ENSG00000198934 | MAGEE1 | -1.26 | 0.13 | 2.89E-24 | 3.17E-23 |
| ENSG00000135824 | RGS8 | -1.26 | 0.19 | 1.81E-12 | 1.17E-11 |
| ENSG00000232077 | LINC01031 | -1.26 | 0.21 | 5.44E-11 | 3.21E-10 |
| ENSG00000204516 | MICB | -1.26 | 0.21 | 1.08E-10 | 6.22E-10 |
| ENSG00000130270 | ATP8B3 | -1.26 | 0.24 | 7.90E-09 | 3.98E-08 |
| ENSG00000273476 | | -1.26 | 0.42 | 7.80E-05 | 2.75E-04 |
| ENSG00000287016 | | -1.26 | 0.56 | 5.82E-04 | 1.84E-03 |
| ENSG00000174600 | CMKLR1 | -1.26 | 0.59 | 7.61E-04 | 2.36E-03 |
| ENSG00000106236 | NPTX2 | -1.25 | 0.04 | 4.32E-260 | 1.53E-257 |
| ENSG00000203727 | SAMD5 | -1.25 | 0.04 | 1.31E-180 | 2.01E-178 |
| ENSG00000158773 | USF1 | -1.25 | 0.06 | 8.14E-103 | 4.88E-101 |
| ENSG00000105976 | MET | -1.25 | 0.07 | 2.23E-69 | 7.55E-68 |
| ENSG00000183160 | TMEM119 | -1.25 | 0.09 | 3.50E-44 | 6.97E-43 |
| ENSG00000171385 | KCND3 | -1.25 | 0.13 | 5.45E-23 | 5.71E-22 |
| ENSG00000136122 | BORA | -1.25 | 0.16 | 9.47E-16 | 7.24E-15 |
| ENSG00000244649 | LINC02086 | -1.25 | 0.46 | 2.05E-04 | 6.87E-04 |
| ENSG00000276633 | | -1.25 | 0.47 | 2.27E-04 | 7.57E-04 |
| ENSG00000144452 | ABCA12 | -1.25 | 0.62 | 9.70E-04 | 2.95E-03 |
| ENSG00000105968 | H2AZ2 | -1.24 | 0.05 | 3.34E-159 | 3.94E-157 |
| ENSG00000144231 | POLR2D | -1.24 | 0.06 | 7.73E-110 | 5.02E-108 |
| ENSG00000123080 | CDKN2C | -1.24 | 0.07 | 1.60E-72 | 5.75E-71 |
| ENSG00000188827 | SLX4 | -1.24 | 0.07 | 7.15E-72 | 2.54E-70 |
| ENSG00000073849 | ST6GAL1 | -1.24 | 0.08 | 5.04E-53 | 1.22E-51 |
| ENSG00000123106 | CCDC91 | -1.24 | 0.1 | 3.68E-36 | 5.99E-35 |
| ENSG00000037280 | FLT4 | -1.24 | 0.11 | 1.17E-31 | 1.65E-30 |
| ENSG00000102385 | DRP2 | -1.24 | 0.18 | 4.47E-13 | 2.99E-12 |
| ENSG00000237667 | LINC01115 | -1.24 | 0.18 | 8.41E-13 | 5.54E-12 |
| ENSG00000153246 | PLA2R1 | -1.24 | 0.31 | 2.46E-06 | 1.01E-05 |

| | | | | | |
|---|---|---|---|---|---|
| ENSG00000282393 | | -1.24 | 0.35 | 1.68E-05 | 6.36E-05 |
| ENSG00000072952 | IRAG1 | -1.24 | 0.47 | 2.50E-04 | 8.29E-04 |
| ENSG00000006210 | CX3CL1 | -1.24 | 0.54 | 5.76E-04 | 1.82E-03 |
| ENSG00000162745 | OLFML2B | -1.23 | 0.03 | 2.19E-281 | 9.25E-279 |
| ENSG00000076067 | RBMS2 | -1.23 | 0.06 | 1.68E-98 | 9.48E-97 |
| ENSG00000196154 | S100A4 | -1.23 | 0.06 | 1.25E-90 | 6.16E-89 |
| ENSG00000196177 | ACADSB | -1.23 | 0.07 | 5.32E-73 | 1.95E-71 |
| ENSG00000185745 | IFIT1 | -1.23 | 0.07 | 2.12E-65 | 6.61E-64 |
| ENSG00000168758 | SEMA4C | -1.23 | 0.07 | 1.17E-63 | 3.52E-62 |
| ENSG00000161692 | DBF4B | -1.23 | 0.08 | 6.26E-58 | 1.68E-56 |
| ENSG00000179833 | SERTAD2 | -1.23 | 0.08 | 6.90E-49 | 1.51E-47 |
| ENSG00000103550 | KNOP1 | -1.23 | 0.09 | 1.69E-46 | 3.51E-45 |
| ENSG00000205476 | CCDC85C | -1.23 | 0.09 | 4.27E-46 | 8.79E-45 |
| ENSG00000180818 | HOXC10 | -1.23 | 0.09 | 4.47E-46 | 9.19E-45 |
| ENSG00000176208 | ATAD5 | -1.23 | 0.09 | 3.04E-40 | 5.48E-39 |
| ENSG00000172348 | RCAN2 | -1.23 | 0.11 | 8.32E-33 | 1.22E-31 |
| ENSG00000146859 | TMEM140 | -1.23 | 0.11 | 9.89E-33 | 1.44E-31 |
| ENSG00000151136 | ABTB3 | -1.23 | 0.25 | 4.04E-08 | 1.93E-07 |
| ENSG00000284946 | | -1.23 | 0.25 | 5.12E-08 | 2.43E-07 |
| ENSG00000185652 | NTF3 | -1.23 | 0.29 | 6.26E-07 | 2.72E-06 |
| ENSG00000288903 | | -1.23 | 0.3 | 1.58E-06 | 6.57E-06 |
| ENSG00000270605 | | -1.23 | 0.36 | 2.33E-05 | 8.69E-05 |
| ENSG00000256540 | IQSEC3-AS1 | -1.23 | 0.94 | 5.72E-04 | 1.81E-03 |
| ENSG00000151208 | DLG5 | -1.22 | 0.05 | 4.38E-119 | 3.17E-117 |
| ENSG00000143924 | EML4 | -1.22 | 0.05 | 6.72E-117 | 4.75E-115 |
| ENSG00000068976 | PYGM | -1.22 | 0.06 | 1.66E-92 | 8.60E-91 |
| ENSG00000049759 | NEDD4L | -1.22 | 0.06 | 2.16E-90 | 1.06E-88 |
| ENSG00000149591 | TAGLN | -1.22 | 0.07 | 1.81E-75 | 6.92E-74 |
| ENSG00000204604 | ZNF468 | -1.22 | 0.07 | 1.38E-65 | 4.34E-64 |
| ENSG00000024422 | EHD2 | -1.22 | 0.08 | 6.69E-59 | 1.84E-57 |
| ENSG00000157657 | ZNF618 | -1.22 | 0.08 | 1.30E-58 | 3.53E-57 |
| ENSG00000081377 | CDC14B | -1.22 | 0.08 | 2.51E-51 | 5.83E-50 |
| ENSG00000101400 | SNTA1 | -1.22 | 0.16 | 7.84E-16 | 6.02E-15 |
| ENSG00000175874 | CREG2 | -1.22 | 0.23 | 3.26E-09 | 1.69E-08 |
| ENSG00000237489 | C10orf143 | -1.22 | 0.23 | 3.84E-09 | 1.98E-08 |
| ENSG00000166387 | PPFIBP2 | -1.22 | 0.24 | 1.31E-08 | 6.49E-08 |
| ENSG00000214872 | SMTNL1 | -1.22 | 0.32 | 4.38E-06 | 1.76E-05 |
| ENSG00000203706 | SERTAD4-AS1 | -1.22 | 0.46 | 2.68E-04 | 8.85E-04 |
| ENSG00000236740 | | -1.22 | 0.48 | 3.37E-04 | 1.09E-03 |
| ENSG00000137440 | FGFBP1 | -1.22 | 0.65 | 1.31E-03 | 3.92E-03 |
| ENSG00000113360 | DROSHA | -1.21 | 0.04 | 4.40E-180 | 6.69E-178 |
| ENSG00000116132 | PRRX1 | -1.21 | 0.05 | 7.36E-124 | 5.65E-122 |
| ENSG00000137831 | UACA | -1.21 | 0.06 | 1.99E-81 | 8.41E-80 |
| ENSG00000154642 | C21orf91 | -1.21 | 0.07 | 1.11E-68 | 3.69E-67 |
| ENSG00000007237 | GAS7 | -1.21 | 0.07 | 5.43E-62 | 1.57E-60 |
| ENSG00000185379 | RAD51D | -1.21 | 0.13 | 2.32E-22 | 2.37E-21 |
| ENSG00000081181 | ARG2 | -1.21 | 0.16 | 8.25E-15 | 6.07E-14 |
| ENSG00000138080 | EMILIN1 | -1.21 | 0.19 | 2.11E-11 | 1.28E-10 |
| ENSG00000128833 | MYO5C | -1.21 | 0.3 | 1.97E-06 | 8.14E-06 |
| ENSG00000253161 | LINC01605 | -1.21 | 0.33 | 7.60E-06 | 2.97E-05 |

| | | | | | |
|---|---|---|---|---|---|
| ENSG00000172794 | RAB37 | -1.21 | 0.33 | 8.32E-06 | 3.24E-05 |
| ENSG00000143257 | NR1I3 | -1.21 | 0.4 | 9.82E-05 | 3.42E-04 |
| ENSG00000225299 | | -1.21 | 0.65 | 1.46E-03 | 4.32E-03 |
| ENSG00000135821 | GLUL | -1.2 | 0.04 | 9.36E-238 | 2.75E-235 |
| ENSG00000116698 | SMG7 | -1.2 | 0.05 | 1.61E-140 | 1.54E-138 |
| ENSG00000111300 | NAA25 | -1.2 | 0.05 | 5.59E-112 | 3.73E-110 |
| ENSG00000119681 | LTBP2 | -1.2 | 0.06 | 5.60E-105 | 3.47E-103 |
| ENSG00000187772 | LIN28B | -1.2 | 0.06 | 6.85E-95 | 3.70E-93 |
| ENSG00000111696 | NT5DC3 | -1.2 | 0.06 | 5.53E-81 | 2.31E-79 |
| ENSG00000165490 | DDIAS | -1.2 | 0.07 | 1.72E-65 | 5.38E-64 |
| ENSG00000239474 | KLHL41 | -1.2 | 0.07 | 1.14E-63 | 3.43E-62 |
| ENSG00000198910 | L1CAM | -1.2 | 0.08 | 3.13E-56 | 8.11E-55 |
| ENSG00000176438 | SYNE3 | -1.2 | 0.08 | 4.74E-50 | 1.06E-48 |
| ENSG00000055147 | FAM114A2 | -1.2 | 0.09 | 1.31E-41 | 2.47E-40 |
| ENSG00000141441 | GAREM1 | -1.2 | 0.1 | 1.87E-33 | 2.80E-32 |
| ENSG00000198624 | CCDC69 | -1.2 | 0.12 | 5.72E-26 | 6.67E-25 |
| ENSG00000141622 | RNF165 | -1.2 | 0.13 | 3.08E-22 | 3.12E-21 |
| ENSG00000100311 | PDGFB | -1.2 | 0.17 | 1.47E-13 | 1.01E-12 |
| ENSG00000114166 | KAT2B | -1.2 | 0.18 | 6.09E-13 | 4.05E-12 |
| ENSG00000225613 | LINCMD1 | -1.2 | 0.23 | 7.99E-09 | 4.03E-08 |
| ENSG00000137558 | PI15 | -1.2 | 0.32 | 6.80E-06 | 2.67E-05 |
| ENSG00000229638 | RPL4P4 | -1.2 | 0.4 | 1.08E-04 | 3.73E-04 |
| ENSG00000260193 | LINC02846 | -1.2 | 0.42 | 1.52E-04 | 5.17E-04 |
| ENSG00000168453 | HR | -1.2 | 0.64 | 1.51E-03 | 4.46E-03 |
| ENSG00000107957 | SH3PXD2A | -1.19 | 0.04 | 1.87E-167 | 2.38E-165 |
| ENSG00000121966 | CXCR4 | -1.19 | 0.07 | 3.59E-70 | 1.24E-68 |
| ENSG00000165156 | ZHX1 | -1.19 | 0.07 | 1.85E-65 | 5.79E-64 |
| ENSG00000131435 | PDLIM4 | -1.19 | 0.07 | 4.88E-61 | 1.39E-59 |
| ENSG00000068001 | HYAL2 | -1.19 | 0.08 | 2.72E-57 | 7.20E-56 |
| ENSG00000155858 | LSM11 | -1.19 | 0.08 | 1.60E-54 | 4.03E-53 |
| ENSG00000135392 | DNAJC14 | -1.19 | 0.08 | 2.81E-52 | 6.68E-51 |
| ENSG00000058729 | RIOK2 | -1.19 | 0.08 | 3.80E-51 | 8.78E-50 |
| ENSG00000168283 | BMI1 | -1.19 | 0.09 | 1.55E-42 | 2.98E-41 |
| ENSG00000213799 | ZNF845 | -1.19 | 0.09 | 1.51E-41 | 2.83E-40 |
| ENSG00000261039 | LINC02544 | -1.19 | 0.09 | 2.83E-41 | 5.27E-40 |
| ENSG00000176909 | MAMSTR | -1.19 | 0.1 | 5.36E-36 | 8.69E-35 |
| ENSG00000163492 | CCDC141 | -1.19 | 0.11 | 7.67E-31 | 1.06E-29 |
| ENSG00000258952 | SALRNA1 | -1.19 | 0.12 | 2.28E-23 | 2.42E-22 |
| ENSG00000173641 | HSPB7 | -1.19 | 0.28 | 1.22E-06 | 5.13E-06 |
| ENSG00000183023 | SLC8A1 | -1.19 | 0.34 | 2.30E-05 | 8.59E-05 |
| ENSG00000231672 | DIRC3 | -1.19 | 0.43 | 1.91E-04 | 6.40E-04 |
| ENSG00000184900 | SUMO3 | -1.18 | 0.04 | 3.05E-175 | 4.34E-173 |
| ENSG00000155393 | HEATR3 | -1.18 | 0.06 | 7.31E-83 | 3.19E-81 |
| ENSG00000072422 | RHOBTB1 | -1.18 | 0.07 | 4.03E-66 | 1.28E-64 |
| ENSG00000111445 | RFC5 | -1.18 | 0.07 | 1.29E-59 | 3.61E-58 |
| ENSG00000138193 | PLCE1 | -1.18 | 0.08 | 4.23E-51 | 9.75E-50 |
| ENSG00000100292 | HMOX1 | -1.18 | 0.08 | 1.47E-48 | 3.19E-47 |
| ENSG00000139354 | GAS2L3 | -1.18 | 0.09 | 7.77E-40 | 1.38E-38 |
| ENSG00000138738 | PRDM5 | -1.18 | 0.12 | 3.06E-25 | 3.49E-24 |
| ENSG00000214756 | CSKMT | -1.18 | 0.12 | 1.04E-23 | 1.12E-22 |

| | | | | | |
|---|---|---|---|---|---|
| ENSG00000154920 | EME1 | -1.18 | 0.13 | 1.15E-21 | 1.14E-20 |
| ENSG00000280120 | | -1.18 | 0.15 | 9.35E-17 | 7.54E-16 |
| ENSG00000205755 | CRLF2 | -1.18 | 0.21 | 1.79E-09 | 9.48E-09 |
| ENSG00000010319 | SEMA3G | -1.18 | 0.24 | 3.10E-08 | 1.50E-07 |
| ENSG00000131386 | GALNT15 | -1.18 | 0.27 | 6.84E-07 | 2.96E-06 |
| ENSG00000144644 | GADL1 | -1.18 | 0.35 | 3.09E-05 | 1.14E-04 |
| ENSG00000176029 | C11orf16 | -1.18 | 0.48 | 5.04E-04 | 1.60E-03 |
| ENSG00000111602 | TIMELESS | -1.17 | 0.05 | 3.61E-141 | 3.51E-139 |
| ENSG00000196227 | FAM217B | -1.17 | 0.05 | 2.56E-106 | 1.62E-104 |
| ENSG00000136108 | CKAP2 | -1.17 | 0.06 | 3.57E-87 | 1.64E-85 |
| ENSG00000131351 | HAUS8 | -1.17 | 0.08 | 1.74E-54 | 4.37E-53 |
| ENSG00000126878 | AIF1L | -1.17 | 0.08 | 1.16E-46 | 2.43E-45 |
| ENSG00000127507 | ADGRE2 | -1.17 | 0.09 | 2.40E-43 | 4.69E-42 |
| ENSG00000102967 | DHODH | -1.17 | 0.09 | 5.91E-42 | 1.12E-40 |
| ENSG00000159445 | THEM4 | -1.17 | 0.09 | 1.44E-38 | 2.51E-37 |
| ENSG00000204054 | LINC00963 | -1.17 | 0.1 | 2.43E-33 | 3.61E-32 |
| ENSG00000119915 | ELOVL3 | -1.17 | 0.12 | 6.36E-23 | 6.65E-22 |
| ENSG00000181544 | FANCB | -1.17 | 0.13 | 1.95E-21 | 1.92E-20 |
| ENSG00000108387 | SEPTIN4 | -1.17 | 0.13 | 7.22E-20 | 6.68E-19 |
| ENSG00000139174 | PRICKLE1 | -1.17 | 0.15 | 1.47E-16 | 1.18E-15 |
| ENSG00000179921 | GPBAR1 | -1.17 | 0.22 | 6.07E-09 | 3.08E-08 |
| ENSG00000134716 | CYP2J2 | -1.17 | 0.38 | 8.76E-05 | 3.07E-04 |
| ENSG00000105810 | CDK6 | -1.16 | 0.05 | 1.26E-136 | 1.17E-134 |
| ENSG00000185482 | STAC3 | -1.16 | 0.05 | 1.05E-118 | 7.54E-117 |
| ENSG00000134369 | NAV1 | -1.16 | 0.05 | 2.02E-116 | 1.41E-114 |
| ENSG00000123473 | STIL | -1.16 | 0.06 | 1.99E-92 | 1.03E-90 |
| ENSG00000103005 | USB1 | -1.16 | 0.06 | 1.09E-90 | 5.38E-89 |
| ENSG00000181754 | AMIGO1 | -1.16 | 0.06 | 1.85E-76 | 7.22E-75 |
| ENSG00000166295 | ANAPC16 | -1.16 | 0.07 | 1.00E-68 | 3.33E-67 |
| ENSG00000213551 | DNAJC9 | -1.16 | 0.08 | 5.34E-54 | 1.33E-52 |
| ENSG00000214194 | SMIM30 | -1.16 | 0.08 | 1.80E-50 | 4.09E-49 |
| ENSG00000130304 | SLC27A1 | -1.16 | 0.1 | 3.17E-30 | 4.29E-29 |
| ENSG00000163170 | BOLA3 | -1.16 | 0.18 | 4.97E-12 | 3.12E-11 |
| ENSG00000196132 | MYT1 | -1.16 | 0.24 | 9.31E-08 | 4.33E-07 |
| ENSG00000248213 | CICP16 | -1.16 | 0.27 | 7.63E-07 | 3.28E-06 |
| ENSG00000164743 | C8orf48 | -1.16 | 0.29 | 2.59E-06 | 1.06E-05 |
| ENSG00000155792 | DEPTOR | -1.16 | 0.36 | 4.93E-05 | 1.77E-04 |
| ENSG00000279464 | | -1.16 | 0.52 | 8.24E-04 | 2.54E-03 |
| ENSG00000233581 | | -1.16 | 0.7 | 2.10E-03 | 6.05E-03 |
| ENSG00000152661 | GJA1 | -1.15 | 0.04 | 1.51E-154 | 1.67E-152 |
| ENSG00000250899 | | -1.15 | 0.05 | 1.94E-131 | 1.68E-129 |
| ENSG00000196850 | PPTC7 | -1.15 | 0.05 | 2.96E-127 | 2.40E-125 |
| ENSG00000147274 | RBMX | -1.15 | 0.06 | 4.82E-93 | 2.55E-91 |
| ENSG00000118971 | CCND2 | -1.15 | 0.06 | 5.90E-90 | 2.87E-88 |
| ENSG00000174606 | ANGEL2 | -1.15 | 0.06 | 3.80E-87 | 1.75E-85 |
| ENSG00000079616 | KIF22 | -1.15 | 0.06 | 2.57E-72 | 9.23E-71 |
| ENSG00000105357 | MYH14 | -1.15 | 0.07 | 4.32E-70 | 1.49E-68 |
| ENSG00000014824 | SLC30A9 | -1.15 | 0.07 | 1.42E-67 | 4.60E-66 |
| ENSG00000110442 | COMMD9 | -1.15 | 0.07 | 5.64E-64 | 1.71E-62 |
| ENSG00000144583 | MARCHF4 | -1.15 | 0.08 | 5.37E-50 | 1.20E-48 |

| | | | | | |
|---|---|---|---|---|---|
| ENSG00000185122 | HSF1 | -1.15 | 0.08 | 1.60E-48 | 3.45E-47 |
| ENSG00000152133 | GPATCH11 | -1.15 | 0.08 | 2.66E-45 | 5.40E-44 |
| ENSG00000125703 | ATG4C | -1.15 | 0.09 | 9.88E-40 | 1.76E-38 |
| ENSG00000178695 | KCTD12 | -1.15 | 0.1 | 2.90E-31 | 4.07E-30 |
| ENSG00000290292 | HOMEZ | -1.15 | 0.11 | 7.44E-25 | 8.37E-24 |
| ENSG00000276600 | RAB7B | -1.15 | 0.12 | 5.18E-23 | 5.43E-22 |
| ENSG00000127423 | AUNIP | -1.15 | 0.12 | 1.35E-22 | 1.39E-21 |
| ENSG00000187550 | SBK2 | -1.15 | 0.13 | 5.20E-20 | 4.84E-19 |
| ENSG00000178150 | ZNF114 | -1.15 | 0.2 | 3.51E-10 | 1.96E-09 |
| ENSG00000124212 | PTGIS | -1.15 | 0.25 | 2.93E-07 | 1.31E-06 |
| ENSG00000225706 | PTPRD-AS1 | -1.15 | 0.31 | 9.89E-06 | 3.83E-05 |
| ENSG00000058335 | RASGRF1 | -1.15 | 0.42 | 2.42E-04 | 8.01E-04 |
| ENSG00000118729 | CASQ2 | -1.15 | 0.9 | 1.01E-03 | 3.07E-03 |
| ENSG00000187862 | TTC24 | -1.15 | 0.54 | 1.21E-03 | 3.63E-03 |
| ENSG00000186854 | TRABD2A | -1.14 | 0.06 | 5.56E-88 | 2.63E-86 |
| ENSG00000006634 | DBF4 | -1.14 | 0.06 | 8.22E-88 | 3.84E-86 |
| ENSG00000132436 | FIGNL1 | -1.14 | 0.06 | 2.27E-84 | 1.01E-82 |
| ENSG00000131389 | SLC6A6 | -1.14 | 0.06 | 2.48E-83 | 1.10E-81 |
| ENSG00000065970 | FOXJ2 | -1.14 | 0.06 | 1.05E-72 | 3.80E-71 |
| ENSG00000167632 | TRAPPC9 | -1.14 | 0.08 | 4.72E-45 | 9.52E-44 |
| ENSG00000000460 | C1orf112 | -1.14 | 0.08 | 4.43E-44 | 8.78E-43 |
| ENSG00000182580 | EPHB3 | -1.14 | 0.09 | 5.71E-40 | 1.02E-38 |
| ENSG00000139618 | BRCA2 | -1.14 | 0.1 | 1.29E-32 | 1.87E-31 |
| ENSG00000149212 | SESN3 | -1.14 | 0.1 | 1.53E-29 | 2.02E-28 |
| ENSG00000123572 | NRK | -1.14 | 0.14 | 1.26E-17 | 1.06E-16 |
| ENSG00000272325 | NUDT3 | -1.14 | 0.15 | 1.14E-15 | 8.71E-15 |
| ENSG00000164342 | TLR3 | -1.14 | 0.18 | 1.31E-11 | 8.02E-11 |
| ENSG00000106538 | RARRES2 | -1.14 | 0.18 | 3.51E-11 | 2.09E-10 |
| ENSG00000134940 | ACRV1 | -1.14 | 0.38 | 1.08E-04 | 3.73E-04 |
| ENSG00000255853 | | -1.14 | 0.39 | 1.42E-04 | 4.85E-04 |
| ENSG00000174928 | C3orf33 | -1.14 | 0.41 | 2.20E-04 | 7.35E-04 |
| ENSG00000120262 | CCDC170 | -1.14 | 0.52 | 9.83E-04 | 2.99E-03 |
| ENSG00000153993 | SEMA3D | -1.13 | 0.06 | 1.57E-90 | 7.75E-89 |
| ENSG00000131381 | RBSN | -1.13 | 0.06 | 1.21E-77 | 4.80E-76 |
| ENSG00000198912 | C1orf174 | -1.13 | 0.06 | 2.56E-70 | 8.86E-69 |
| ENSG00000077522 | ACTN2 | -1.13 | 0.08 | 1.01E-52 | 2.44E-51 |
| ENSG00000271976 | | -1.13 | 0.11 | 2.08E-27 | 2.56E-26 |
| ENSG00000143858 | SYT2 | -1.13 | 0.12 | 1.32E-22 | 1.36E-21 |
| ENSG00000100433 | KCNK10 | -1.13 | 0.13 | 5.24E-19 | 4.65E-18 |
| ENSG00000183597 | TANGO2 | -1.13 | 0.14 | 1.48E-16 | 1.18E-15 |
| ENSG00000196872 | CRACDL | -1.13 | 0.3 | 9.42E-06 | 3.65E-05 |
| ENSG00000163017 | ACTG2 | -1.13 | 0.69 | 2.42E-03 | 6.93E-03 |
| ENSG00000053438 | NNAT | -1.12 | 0.04 | 2.07E-166 | 2.60E-164 |
| ENSG00000114738 | MAPKAPK3 | -1.12 | 0.05 | 1.10E-114 | 7.48E-113 |
| ENSG00000179242 | CDH4 | -1.12 | 0.05 | 9.79E-94 | 5.21E-92 |
| ENSG00000141084 | RANBP10 | -1.12 | 0.06 | 2.25E-84 | 1.01E-82 |
| ENSG00000196739 | COL27A1 | -1.12 | 0.06 | 9.08E-80 | 3.73E-78 |
| ENSG00000130962 | PRRG1 | -1.12 | 0.07 | 7.87E-56 | 2.03E-54 |
| ENSG00000091127 | PUS7 | -1.12 | 0.07 | 1.37E-52 | 3.29E-51 |
| ENSG00000157600 | TMEM164 | -1.12 | 0.08 | 1.55E-49 | 3.43E-48 |

| | | | | | |
|---|---|---|---|---|---|
| ENSG00000182054 | IDH2 | -1.12 | 0.08 | 1.06E-45 | 2.16E-44 |
| ENSG00000080839 | RBL1 | -1.12 | 0.09 | 3.47E-34 | 5.29E-33 |
| ENSG00000169891 | REPS2 | -1.12 | 0.1 | 1.14E-29 | 1.51E-28 |
| ENSG00000173894 | CBX2 | -1.12 | 0.11 | 2.38E-26 | 2.82E-25 |
| ENSG00000234160 | | -1.12 | 0.12 | 1.12E-23 | 1.21E-22 |
| ENSG00000283703 | VSIG10L2 | -1.12 | 0.18 | 1.57E-11 | 9.59E-11 |
| ENSG00000113657 | DPYSL3 | -1.12 | 0.2 | 6.78E-10 | 3.72E-09 |
| ENSG00000111305 | GSG1 | -1.12 | 0.2 | 1.66E-09 | 8.80E-09 |
| ENSG00000120217 | CD274 | -1.12 | 0.23 | 6.12E-08 | 2.88E-07 |
| ENSG00000183631 | PRR32 | -1.12 | 0.3 | 8.99E-06 | 3.49E-05 |
| ENSG00000228836 | CT45A5 | -1.12 | 0.63 | 2.40E-03 | 6.85E-03 |
| ENSG00000197043 | ANXA6 | -1.11 | 0.05 | 1.03E-121 | 7.78E-120 |
| ENSG00000145284 | SCD5 | -1.11 | 0.06 | 3.41E-76 | 1.33E-74 |
| ENSG00000132313 | MRPL35 | -1.11 | 0.07 | 1.77E-61 | 5.08E-60 |
| ENSG00000111666 | CHPT1 | -1.11 | 0.07 | 6.11E-54 | 1.52E-52 |
| ENSG00000124588 | NQO2 | -1.11 | 0.1 | 3.68E-30 | 4.96E-29 |
| ENSG00000141965 | FEM1A | -1.11 | 0.11 | 8.55E-24 | 9.24E-23 |
| ENSG00000289094 | | -1.11 | 0.21 | 1.07E-08 | 5.35E-08 |
| ENSG00000224239 | | -1.11 | 0.23 | 4.84E-08 | 2.30E-07 |
| ENSG00000285108 | | -1.11 | 0.26 | 9.90E-07 | 4.20E-06 |
| ENSG00000147394 | ZNF185 | -1.11 | 0.36 | 8.19E-05 | 2.88E-04 |
| ENSG00000134755 | DSC2 | -1.11 | 0.37 | 1.35E-04 | 4.63E-04 |
| ENSG00000070182 | SPTB | -1.11 | 0.45 | 5.59E-04 | 1.77E-03 |
| ENSG00000247877 | FAM174A-DT | -1.11 | 0.61 | 2.34E-03 | 6.71E-03 |
| ENSG00000274825 | | -1.11 | 0.63 | 2.35E-03 | 6.73E-03 |
| ENSG00000198570 | RD3 | -1.11 | 0.65 | 2.57E-03 | 7.30E-03 |
| ENSG00000259458 | MGC15885 | -1.11 | 0.7 | 2.74E-03 | 7.74E-03 |
| ENSG00000188042 | ARL4C | -1.1 | 0.04 | 7.45E-159 | 8.71E-157 |
| ENSG00000136478 | TEX2 | -1.1 | 0.05 | 3.57E-111 | 2.38E-109 |
| ENSG00000198743 | SLC5A3 | -1.1 | 0.07 | 1.34E-65 | 4.21E-64 |
| ENSG00000165675 | ENOX2 | -1.1 | 0.07 | 1.11E-58 | 3.04E-57 |
| ENSG00000184867 | ARMCX2 | -1.1 | 0.08 | 6.08E-50 | 1.36E-48 |
| ENSG00000136235 | GPNMB | -1.1 | 0.08 | 2.55E-48 | 5.49E-47 |
| ENSG00000175348 | TMEM9B | -1.1 | 0.08 | 6.51E-43 | 1.26E-41 |
| ENSG00000165209 | STRBP | -1.1 | 0.09 | 3.15E-35 | 4.96E-34 |
| ENSG00000215883 | CYB5RL | -1.1 | 0.09 | 8.54E-35 | 1.33E-33 |
| ENSG00000162591 | MEGF6 | -1.1 | 0.1 | 1.14E-29 | 1.52E-28 |
| ENSG00000198554 | WDHD1 | -1.1 | 0.11 | 1.83E-26 | 2.18E-25 |
| ENSG00000159184 | HOXB13 | -1.1 | 0.11 | 3.38E-24 | 3.71E-23 |
| ENSG00000079215 | SLC1A3 | -1.1 | 0.14 | 4.98E-17 | 4.06E-16 |
| ENSG00000198336 | MYL4 | -1.1 | 0.16 | 1.30E-12 | 8.45E-12 |
| ENSG00000259207 | ITGB3 | -1.1 | 0.21 | 7.19E-09 | 3.63E-08 |
| ENSG00000065717 | TLE2 | -1.1 | 0.25 | 6.32E-07 | 2.74E-06 |
| ENSG00000146005 | PSD2 | -1.1 | 0.47 | 9.12E-04 | 2.79E-03 |
| ENSG00000204542 | C6orf15 | -1.1 | 0.9 | 1.22E-03 | 3.65E-03 |
| ENSG00000248690 | HAS2-AS1 | -1.1 | 0.54 | 1.63E-03 | 4.78E-03 |
| ENSG00000064309 | CDON | -1.09 | 0.04 | 1.83E-200 | 3.60E-198 |
| ENSG00000127824 | TUBA4A | -1.09 | 0.04 | 6.61E-134 | 5.91E-132 |
| ENSG00000185686 | PRAME | -1.09 | 0.05 | 4.17E-128 | 3.45E-126 |
| ENSG00000182022 | CHST15 | -1.09 | 0.05 | 1.63E-110 | 1.08E-108 |

| | | | | | |
|---|---|---|---|---|---|
| ENSG00000085511 | MAP3K4 | -1.09 | 0.05 | 3.67E-110 | 2.41E-108 |
| ENSG00000001497 | LAS1L | -1.09 | 0.06 | 1.02E-82 | 4.42E-81 |
| ENSG00000149218 | ENDOD1 | -1.09 | 0.06 | 5.60E-75 | 2.12E-73 |
| ENSG00000135362 | PRR5L | -1.09 | 0.07 | 2.39E-56 | 6.20E-55 |
| ENSG00000184445 | KNTC1 | -1.09 | 0.07 | 1.65E-51 | 3.84E-50 |
| ENSG00000112029 | FBXO5 | -1.09 | 0.07 | 8.67E-51 | 1.98E-49 |
| ENSG00000075618 | FSCN1 | -1.09 | 0.07 | 8.77E-50 | 1.95E-48 |
| ENSG00000168916 | ZNF608 | -1.09 | 0.09 | 3.12E-32 | 4.51E-31 |
| ENSG00000012211 | PRICKLE3 | -1.09 | 0.12 | 3.83E-20 | 3.58E-19 |
| ENSG00000229563 | LINC01204 | -1.09 | 0.15 | 3.00E-15 | 2.25E-14 |
| ENSG00000185697 | MYBL1 | -1.09 | 0.16 | 8.71E-13 | 5.73E-12 |
| ENSG00000167074 | TEF | -1.09 | 0.16 | 2.00E-12 | 1.29E-11 |
| ENSG00000280119 | | -1.09 | 0.18 | 7.11E-11 | 4.16E-10 |
| ENSG00000134962 | KLB | -1.09 | 0.2 | 2.19E-09 | 1.16E-08 |
| ENSG00000242779 | ZNF702P | -1.09 | 0.21 | 9.91E-09 | 4.97E-08 |
| ENSG00000271851 | | -1.09 | 0.23 | 1.58E-07 | 7.21E-07 |
| ENSG00000159166 | LAD1 | -1.09 | 0.25 | 5.79E-07 | 2.52E-06 |
| ENSG00000290374 | FAM86EP | -1.09 | 0.27 | 2.31E-06 | 9.46E-06 |
| ENSG00000286638 | | -1.09 | 0.86 | 1.70E-03 | 4.97E-03 |
| ENSG00000267480 | LINC02979 | -1.09 | 0.56 | 2.07E-03 | 5.98E-03 |
| ENSG00000182158 | CREB3L2 | -1.08 | 0.04 | 2.39E-145 | 2.42E-143 |
| ENSG00000101294 | HM13 | -1.08 | 0.05 | 2.84E-102 | 1.69E-100 |
| ENSG00000174804 | FZD4 | -1.08 | 0.06 | 4.11E-68 | 1.35E-66 |
| ENSG00000112759 | SLC29A1 | -1.08 | 0.07 | 5.39E-62 | 1.56E-60 |
| ENSG00000079308 | TNS1 | -1.08 | 0.07 | 2.53E-57 | 6.71E-56 |
| ENSG00000129680 | MAP7D3 | -1.08 | 0.07 | 1.40E-53 | 3.45E-52 |
| ENSG00000137760 | ALKBH8 | -1.08 | 0.07 | 2.01E-50 | 4.56E-49 |
| ENSG00000105486 | LIG1 | -1.08 | 0.09 | 4.43E-38 | 7.60E-37 |
| ENSG00000183091 | NEB | -1.08 | 0.1 | 9.42E-30 | 1.26E-28 |
| ENSG00000129534 | MIS18BP1 | -1.08 | 0.1 | 4.02E-29 | 5.24E-28 |
| ENSG00000112246 | SIM1 | -1.08 | 0.1 | 6.93E-29 | 8.98E-28 |
| ENSG00000181938 | GINS3 | -1.08 | 0.1 | 3.56E-27 | 4.34E-26 |
| ENSG00000230487 | PSMG3-AS1 | -1.08 | 0.1 | 8.13E-26 | 9.44E-25 |
| ENSG00000157851 | DPYSL5 | -1.08 | 0.11 | 1.89E-24 | 2.09E-23 |
| ENSG00000280173 | | -1.08 | 0.11 | 1.34E-22 | 1.38E-21 |
| ENSG00000101412 | E2F1 | -1.08 | 0.13 | 2.39E-18 | 2.06E-17 |
| ENSG00000166896 | ATP23 | -1.08 | 0.14 | 7.83E-17 | 6.34E-16 |
| ENSG00000131650 | KREMEN2 | -1.08 | 0.17 | 1.43E-11 | 8.73E-11 |
| ENSG00000112137 | PHACTR1 | -1.08 | 0.17 | 3.19E-11 | 1.91E-10 |
| ENSG00000231274 | SBK3 | -1.08 | 0.18 | 2.09E-10 | 1.19E-09 |
| ENSG00000249577 | | -1.08 | 0.22 | 3.38E-08 | 1.63E-07 |
| ENSG00000101213 | PTK6 | -1.08 | 0.22 | 5.36E-08 | 2.54E-07 |
| ENSG00000285973 | | -1.08 | 0.24 | 3.47E-07 | 1.54E-06 |
| ENSG00000101605 | MYOM1 | -1.08 | 0.25 | 1.20E-06 | 5.07E-06 |
| ENSG00000133083 | DCLK1 | -1.08 | 0.28 | 6.61E-06 | 2.60E-05 |
| ENSG00000279838 | | -1.08 | 0.31 | 2.81E-05 | 1.04E-04 |
| ENSG00000289579 | | -1.08 | 0.33 | 6.24E-05 | 2.23E-04 |
| ENSG00000268055 | | -1.08 | 0.33 | 6.33E-05 | 2.26E-04 |
| ENSG00000179873 | NLRP11 | -1.08 | 0.36 | 1.55E-04 | 5.25E-04 |
| ENSG00000222035 | KIAA2012-AS1 | -1.08 | 0.38 | 2.31E-04 | 7.67E-04 |

| | | | | | |
|---|---|---|---|---|---|
| ENSG00000258311 | | -1.08 | 0.42 | 4.92E-04 | 1.57E-03 |
| ENSG00000260416 | | -1.08 | 0.88 | 1.45E-03 | 4.30E-03 |
| ENSG00000258777 | HIF1A-AS1 | -1.08 | 0.6 | 2.63E-03 | 7.47E-03 |
| ENSG00000169248 | CXCL11 | -1.08 | 0.66 | 3.16E-03 | 8.83E-03 |
| ENSG00000159217 | IGF2BP1 | -1.07 | 0.05 | 1.78E-120 | 1.32E-118 |
| ENSG00000137936 | BCAR3 | -1.07 | 0.05 | 1.28E-104 | 7.85E-103 |
| ENSG00000123908 | AGO2 | -1.07 | 0.05 | 1.75E-104 | 1.07E-102 |
| ENSG00000128050 | PAICS | -1.07 | 0.05 | 1.01E-100 | 5.88E-99 |
| ENSG00000069188 | SDK2 | -1.07 | 0.06 | 1.79E-84 | 8.04E-83 |
| ENSG00000105926 | PALS2 | -1.07 | 0.06 | 5.75E-67 | 1.85E-65 |
| ENSG00000143850 | PLEKHA6 | -1.07 | 0.06 | 1.28E-65 | 4.03E-64 |
| ENSG00000112658 | SRF | -1.07 | 0.06 | 3.16E-62 | 9.21E-61 |
| ENSG00000180694 | TMEM64 | -1.07 | 0.07 | 7.78E-61 | 2.22E-59 |
| ENSG00000204219 | TCEA3 | -1.07 | 0.08 | 7.74E-47 | 1.63E-45 |
| ENSG00000129152 | MYOD1 | -1.07 | 0.08 | 4.65E-44 | 9.22E-43 |
| ENSG00000112312 | GMNN | -1.07 | 0.08 | 6.43E-42 | 1.22E-40 |
| ENSG00000125378 | BMP4 | -1.07 | 0.1 | 6.35E-29 | 8.24E-28 |
| ENSG00000106351 | AGFG2 | -1.07 | 0.1 | 1.73E-27 | 2.14E-26 |
| ENSG00000165449 | SLC16A9 | -1.07 | 0.11 | 3.77E-25 | 4.27E-24 |
| ENSG00000125872 | LRRN4 | -1.07 | 0.19 | 7.73E-10 | 4.21E-09 |
| ENSG00000101306 | MYLK2 | -1.07 | 0.23 | 1.98E-07 | 9.00E-07 |
| ENSG00000290907 | | -1.07 | 0.27 | 5.43E-06 | 2.16E-05 |
| ENSG00000150637 | CD226 | -1.07 | 0.4 | 3.51E-04 | 1.14E-03 |
| ENSG00000130600 | H19 | -1.07 | 0.43 | 6.38E-04 | 2.00E-03 |
| ENSG00000181234 | TMEM132C | -1.07 | 0.88 | 1.49E-03 | 4.39E-03 |
| ENSG00000280399 | | -1.07 | 0.61 | 2.83E-03 | 7.97E-03 |
| ENSG00000237356 | | -1.07 | 0.68 | 3.39E-03 | 9.41E-03 |
| ENSG00000255363 | LINC02757 | -1.07 | 0.68 | 3.49E-03 | 9.67E-03 |
| ENSG00000065308 | TRAM2 | -1.06 | 0.04 | 2.38E-163 | 2.88E-161 |
| ENSG00000144867 | SRPRB | -1.06 | 0.06 | 1.14E-78 | 4.58E-77 |
| ENSG00000174243 | DDX23 | -1.06 | 0.06 | 1.02E-73 | 3.79E-72 |
| ENSG00000113504 | SLC12A7 | -1.06 | 0.06 | 2.56E-70 | 8.87E-69 |
| ENSG00000064115 | TM7SF3 | -1.06 | 0.06 | 5.41E-70 | 1.86E-68 |
| ENSG00000181991 | MRPS11 | -1.06 | 0.07 | 3.28E-60 | 9.22E-59 |
| ENSG00000171914 | TLN2 | -1.06 | 0.07 | 3.52E-58 | 9.47E-57 |
| ENSG00000101868 | POLA1 | -1.06 | 0.07 | 1.41E-46 | 2.95E-45 |
| ENSG00000170430 | MGMT | -1.06 | 0.07 | 2.42E-46 | 5.01E-45 |
| ENSG00000136237 | RAPGEF5 | -1.06 | 0.08 | 5.22E-42 | 9.94E-41 |
| ENSG00000099260 | PALMD | -1.06 | 0.08 | 1.78E-40 | 3.24E-39 |
| ENSG00000169499 | PLEKHA2 | -1.06 | 0.08 | 2.92E-38 | 5.04E-37 |
| ENSG00000173200 | PARP15 | -1.06 | 0.12 | 4.72E-19 | 4.20E-18 |
| ENSG00000116106 | EPHA4 | -1.06 | 0.13 | 1.07E-16 | 8.56E-16 |
| ENSG00000211584 | SLC48A1 | -1.06 | 0.17 | 2.31E-11 | 1.39E-10 |
| ENSG00000087253 | LPCAT2 | -1.06 | 0.17 | 8.52E-11 | 4.96E-10 |
| ENSG00000168062 | BATF2 | -1.06 | 0.21 | 1.79E-08 | 8.77E-08 |
| ENSG00000287356 | | -1.06 | 0.24 | 5.83E-07 | 2.54E-06 |
| ENSG00000188816 | HMX2 | -1.06 | 0.3 | 2.37E-05 | 8.81E-05 |
| ENSG00000272129 | | -1.06 | 0.35 | 1.35E-04 | 4.60E-04 |
| ENSG00000227620 | ALG1L8P | -1.06 | 0.78 | 3.10E-03 | 8.69E-03 |
| ENSG00000259250 | | -1.06 | 0.61 | 3.17E-03 | 8.86E-03 |

| | | | | | |
|---|---|---|---|---|---|
| ENSG00000289433 | | -1.06 | 0.72 | 3.67E-03 | 1.01E-02 |
| ENSG00000166949 | SMAD3 | -1.05 | 0.04 | 1.34E-164 | 1.65E-162 |
| ENSG00000100504 | PYGL | -1.05 | 0.04 | 7.17E-156 | 8.09E-154 |
| ENSG00000288701 | PRRC2B | -1.05 | 0.04 | 1.02E-137 | 9.49E-136 |
| ENSG00000110422 | HIPK3 | -1.05 | 0.04 | 5.59E-130 | 4.75E-128 |
| ENSG00000186468 | RPS23 | -1.05 | 0.05 | 6.76E-93 | 3.54E-91 |
| ENSG00000133424 | LARGE1 | -1.05 | 0.05 | 7.13E-88 | 3.35E-86 |
| ENSG00000135414 | GDF11 | -1.05 | 0.05 | 6.78E-83 | 2.97E-81 |
| ENSG00000148337 | CIZ1 | -1.05 | 0.06 | 9.53E-74 | 3.54E-72 |
| ENSG00000186472 | PCLO | -1.05 | 0.06 | 5.59E-70 | 1.92E-68 |
| ENSG00000198796 | ALPK2 | -1.05 | 0.06 | 3.10E-69 | 1.05E-67 |
| ENSG00000156103 | MMP16 | -1.05 | 0.07 | 2.93E-58 | 7.89E-57 |
| ENSG00000124784 | RIOK1 | -1.05 | 0.07 | 1.76E-56 | 4.59E-55 |
| ENSG00000138092 | CENPO | -1.05 | 0.07 | 3.35E-51 | 7.75E-50 |
| ENSG00000155629 | PIK3AP1 | -1.05 | 0.12 | 8.40E-20 | 7.75E-19 |
| ENSG00000183682 | BMP8A | -1.05 | 0.14 | 2.34E-14 | 1.68E-13 |
| ENSG00000067221 | STOML1 | -1.05 | 0.15 | 3.33E-13 | 2.25E-12 |
| ENSG00000267383 | | -1.05 | 0.18 | 1.61E-10 | 9.17E-10 |
| ENSG00000171517 | LPAR3 | -1.05 | 0.19 | 9.82E-10 | 5.30E-09 |
| ENSG00000249304 | | -1.05 | 0.23 | 2.23E-07 | 1.01E-06 |
| ENSG00000232439 | RPL18AP7 | -1.05 | 0.36 | 2.13E-04 | 7.11E-04 |
| ENSG00000197471 | SPN | -1.05 | 0.47 | 1.26E-03 | 3.76E-03 |
| ENSG00000176510 | OR10AC1 | -1.05 | 0.5 | 1.71E-03 | 4.99E-03 |
| ENSG00000166197 | NOLC1 | -1.04 | 0.04 | 4.01E-182 | 6.40E-180 |
| ENSG00000102900 | NUP93 | -1.04 | 0.04 | 3.08E-136 | 2.82E-134 |
| ENSG00000136383 | ALPK3 | -1.04 | 0.04 | 3.22E-124 | 2.49E-122 |
| ENSG00000134057 | CCNB1 | -1.04 | 0.04 | 1.73E-119 | 1.27E-117 |
| ENSG00000019549 | SNAI2 | -1.04 | 0.05 | 1.03E-114 | 7.06E-113 |
| ENSG00000171681 | ATF7IP | -1.04 | 0.05 | 8.16E-83 | 3.56E-81 |
| ENSG00000138835 | RGS3 | -1.04 | 0.06 | 3.60E-64 | 1.10E-62 |
| ENSG00000156802 | ATAD2 | -1.04 | 0.06 | 6.63E-62 | 1.92E-60 |
| ENSG00000154027 | AK5 | -1.04 | 0.07 | 3.00E-46 | 6.20E-45 |
| ENSG00000165548 | TMEM63C | -1.04 | 0.08 | 9.94E-39 | 1.74E-37 |
| ENSG00000092421 | SEMA6A | -1.04 | 0.08 | 2.36E-37 | 3.95E-36 |
| ENSG00000172340 | SUCLG2 | -1.04 | 0.09 | 2.01E-34 | 3.08E-33 |
| ENSG00000148516 | ZEB1 | -1.04 | 0.09 | 3.53E-30 | 4.78E-29 |
| ENSG00000139579 | NABP2 | -1.04 | 0.09 | 9.24E-30 | 1.23E-28 |
| ENSG00000157152 | SYN2 | -1.04 | 0.11 | 2.93E-22 | 2.98E-21 |
| ENSG00000069399 | BCL3 | -1.04 | 0.12 | 8.06E-20 | 7.44E-19 |
| ENSG00000056736 | IL17RB | -1.04 | 0.12 | 6.12E-18 | 5.18E-17 |
| ENSG00000119943 | PYROXD2 | -1.04 | 0.13 | 8.38E-18 | 7.06E-17 |
| ENSG00000261504 | LINC01686 | -1.04 | 0.2 | 1.91E-08 | 9.37E-08 |
| ENSG00000174807 | CD248 | -1.04 | 0.26 | 4.14E-06 | 1.66E-05 |
| ENSG00000254790 | | -1.04 | 0.37 | 3.06E-04 | 1.00E-03 |
| ENSG00000127325 | BEST3 | -1.04 | 0.43 | 8.35E-04 | 2.57E-03 |
| ENSG00000275591 | XKR5 | -1.04 | 0.44 | 1.00E-03 | 3.05E-03 |
| ENSG00000173809 | TDRD12 | -1.04 | 0.5 | 1.97E-03 | 5.73E-03 |
| ENSG00000285155 | | -1.04 | 0.58 | 3.22E-03 | 9.00E-03 |
| ENSG00000213892 | CEACAM16 | -1.04 | 0.66 | 3.92E-03 | 1.08E-02 |
| ENSG00000172315 | TP53RK | -1.03 | 0.05 | 2.28E-99 | 1.31E-97 |

| | | | | | |
|---|---|---|---|---|---|
| ENSG00000144730 | IL17RD | -1.03 | 0.05 | 1.01E-96 | 5.55E-95 |
| ENSG00000104964 | TLE5 | -1.03 | 0.06 | 1.43E-66 | 4.57E-65 |
| ENSG00000076356 | PLXNA2 | -1.03 | 0.06 | 4.01E-64 | 1.22E-62 |
| ENSG00000189184 | PCDH18 | -1.03 | 0.06 | 5.69E-64 | 1.72E-62 |
| ENSG00000164134 | NAA15 | -1.03 | 0.07 | 1.65E-53 | 4.05E-52 |
| ENSG00000173705 | SUSD5 | -1.03 | 0.07 | 3.77E-52 | 8.95E-51 |
| ENSG00000164051 | CCDC51 | -1.03 | 0.08 | 5.16E-40 | 9.23E-39 |
| ENSG00000089048 | ESF1 | -1.03 | 0.08 | 1.08E-37 | 1.83E-36 |
| ENSG00000128918 | ALDH1A2 | -1.03 | 0.08 | 1.96E-37 | 3.30E-36 |
| ENSG00000203760 | CENPW | -1.03 | 0.11 | 4.56E-21 | 4.43E-20 |
| ENSG00000167528 | ZNF641 | -1.03 | 0.12 | 2.75E-20 | 2.59E-19 |
| ENSG00000183486 | MX2 | -1.03 | 0.12 | 5.58E-19 | 4.94E-18 |
| ENSG00000016391 | CHDH | -1.03 | 0.13 | 4.15E-17 | 3.39E-16 |
| ENSG00000151470 | C4orf33 | -1.03 | 0.17 | 1.14E-10 | 6.57E-10 |
| ENSG00000169282 | KCNAB1 | -1.03 | 0.18 | 4.95E-10 | 2.73E-09 |
| ENSG00000288953 | | -1.03 | 0.19 | 2.26E-09 | 1.19E-08 |
| ENSG00000156966 | B3GNT7 | -1.03 | 0.2 | 1.23E-08 | 6.10E-08 |
| ENSG00000101162 | TUBB1 | -1.03 | 0.21 | 1.03E-07 | 4.78E-07 |
| ENSG00000170579 | DLGAP1 | -1.03 | 0.25 | 1.66E-06 | 6.92E-06 |
| ENSG00000225899 | FRG2B | -1.03 | 0.34 | 1.54E-04 | 5.23E-04 |
| ENSG00000233117 | LINC00702 | -1.03 | 0.34 | 1.57E-04 | 5.32E-04 |
| ENSG00000199080 | MIR133B | -1.03 | 0.43 | 9.20E-04 | 2.81E-03 |
| ENSG00000230479 | LNCTSI | -1.03 | 0.5 | 2.02E-03 | 5.85E-03 |
| ENSG00000120805 | ARL1 | -1.02 | 0.05 | 1.19E-81 | 5.05E-80 |
| ENSG00000103381 | CPPED1 | -1.02 | 0.06 | 2.59E-67 | 8.36E-66 |
| ENSG00000173848 | NET1 | -1.02 | 0.06 | 1.87E-65 | 5.84E-64 |
| ENSG00000164611 | PTTG1 | -1.02 | 0.07 | 1.38E-48 | 2.98E-47 |
| ENSG00000144406 | UNC80 | -1.02 | 0.07 | 1.49E-46 | 3.11E-45 |
| ENSG00000167130 | DOLPP1 | -1.02 | 0.08 | 3.03E-41 | 5.64E-40 |
| ENSG00000070159 | PTPN3 | -1.02 | 0.08 | 8.03E-37 | 1.33E-35 |
| ENSG00000072954 | TMEM38A | -1.02 | 0.08 | 3.20E-36 | 5.21E-35 |
| ENSG00000177614 | PGBD5 | -1.02 | 0.09 | 2.45E-33 | 3.65E-32 |
| ENSG00000182768 | NGRN | -1.02 | 0.09 | 1.25E-31 | 1.77E-30 |
| ENSG00000180233 | ZNRF2 | -1.02 | 0.09 | 5.55E-29 | 7.21E-28 |
| ENSG00000111269 | CREBL2 | -1.02 | 0.09 | 8.96E-29 | 1.15E-27 |
| ENSG00000168899 | VAMP5 | -1.02 | 0.11 | 5.37E-23 | 5.63E-22 |
| ENSG00000183230 | CTNNA3 | -1.02 | 0.12 | 1.26E-19 | 1.15E-18 |
| ENSG00000277972 | CISD3 | -1.02 | 0.12 | 3.61E-18 | 3.09E-17 |
| ENSG00000165617 | DACT1 | -1.02 | 0.14 | 1.70E-13 | 1.17E-12 |
| ENSG00000093217 | XYLB | -1.02 | 0.16 | 5.58E-12 | 3.49E-11 |
| ENSG00000133216 | EPHB2 | -1.02 | 0.17 | 3.80E-11 | 2.26E-10 |
| ENSG00000250519 | | -1.02 | 0.19 | 3.09E-09 | 1.61E-08 |
| ENSG00000179761 | PIPOX | -1.02 | 0.19 | 7.04E-09 | 3.56E-08 |
| ENSG00000237126 | | -1.02 | 0.24 | 1.23E-06 | 5.19E-06 |
| ENSG00000248671 | ALG1L9P | -1.02 | 0.46 | 1.43E-03 | 4.23E-03 |
| ENSG00000169129 | AFAP1L2 | -1.02 | 0.71 | 4.55E-03 | 1.23E-02 |
| ENSG00000147140 | NONO | -1.01 | 0.04 | 5.79E-156 | 6.56E-154 |
| ENSG00000162512 | SDC3 | -1.01 | 0.06 | 4.66E-73 | 1.71E-71 |
| ENSG00000047188 | YTHDC2 | -1.01 | 0.07 | 1.62E-52 | 3.87E-51 |
| ENSG00000171444 | MCC | -1.01 | 0.07 | 4.49E-50 | 1.01E-48 |

| | | | | | |
|---|---|---|---|---|---|
| ENSG00000172493 | AFF1 | -1.01 | 0.07 | 3.64E-44 | 7.24E-43 |
| ENSG00000157036 | EXOG | -1.01 | 0.08 | 1.43E-42 | 2.76E-41 |
| ENSG00000144857 | BOC | -1.01 | 0.08 | 2.10E-36 | 3.44E-35 |
| ENSG00000136295 | TTYH3 | -1.01 | 0.09 | 1.22E-33 | 1.84E-32 |
| ENSG00000164070 | HSPA4L | -1.01 | 0.09 | 5.35E-32 | 7.66E-31 |
| ENSG00000101945 | SUV39H1 | -1.01 | 0.09 | 3.38E-28 | 4.25E-27 |
| ENSG00000035499 | DEPDC1B | -1.01 | 0.1 | 6.52E-26 | 7.60E-25 |
| ENSG00000268798 | | -1.01 | 0.12 | 3.42E-17 | 2.81E-16 |
| ENSG00000233143 | DIRC3-AS1 | -1.01 | 0.17 | 9.29E-11 | 5.40E-10 |
| ENSG00000125898 | FAM110A | -1.01 | 0.19 | 4.66E-09 | 2.39E-08 |
| ENSG00000198406 | BZW1P2 | -1.01 | 0.21 | 6.39E-08 | 3.01E-07 |
| ENSG00000054690 | PLEKHH1 | -1.01 | 0.21 | 7.56E-08 | 3.54E-07 |
| ENSG00000160801 | PTH1R | -1.01 | 0.25 | 4.05E-06 | 1.62E-05 |
| ENSG00000251034 | | -1.01 | 0.26 | 8.47E-06 | 3.30E-05 |
| ENSG00000154914 | USP43 | -1.01 | 0.26 | 8.64E-06 | 3.36E-05 |
| ENSG00000100678 | SLC8A3 | -1.01 | 0.27 | 1.22E-05 | 4.69E-05 |
| ENSG00000277491 | | -1.01 | 0.3 | 4.85E-05 | 1.75E-04 |
| ENSG00000237523 | LINC00857 | -1.01 | 0.33 | 1.31E-04 | 4.47E-04 |
| ENSG00000114541 | FRMD4B | -1.01 | 0.35 | 2.33E-04 | 7.75E-04 |
| ENSG00000273004 | | -1.01 | 0.39 | 5.77E-04 | 1.82E-03 |
| ENSG00000025423 | HSD17B6 | -1.01 | 0.4 | 7.08E-04 | 2.21E-03 |
| ENSG00000279365 | | -1.01 | 0.9 | 1.41E-03 | 4.17E-03 |
| ENSG00000152102 | FAM168B | -1 | 0.04 | 7.03E-128 | 5.77E-126 |
| ENSG00000196141 | SPATS2L | -1 | 0.05 | 3.26E-108 | 2.11E-106 |
| ENSG00000170175 | CHRNB1 | -1 | 0.05 | 3.49E-101 | 2.06E-99 |
| ENSG00000169641 | LUZP1 | -1 | 0.05 | 4.79E-101 | 2.80E-99 |
| ENSG00000060749 | QSER1 | -1 | 0.06 | 7.94E-74 | 2.95E-72 |
| ENSG00000172869 | DMXL1 | -1 | 0.06 | 5.29E-60 | 1.48E-58 |
| ENSG00000136720 | HS6ST1 | -1 | 0.07 | 8.81E-49 | 1.92E-47 |
| ENSG00000156869 | FRRS1 | -1 | 0.08 | 2.08E-38 | 3.61E-37 |
| ENSG00000121671 | CRY2 | -1 | 0.08 | 6.22E-35 | 9.72E-34 |
| ENSG00000104490 | NCALD | -1 | 0.08 | 4.88E-33 | 7.19E-32 |
| ENSG00000088876 | ZNF343 | -1 | 0.09 | 1.12E-31 | 1.59E-30 |
| ENSG00000248746 | ACTN3 | -1 | 0.09 | 1.58E-30 | 2.17E-29 |
| ENSG00000138658 | ZGRF1 | -1 | 0.09 | 2.09E-28 | 2.64E-27 |
| ENSG00000136114 | THSD1 | -1 | 0.09 | 2.21E-27 | 2.72E-26 |
| ENSG00000143842 | SOX13 | -1 | 0.09 | 4.10E-27 | 5.00E-26 |
| ENSG00000179331 | RAB39A | -1 | 0.13 | 4.42E-16 | 3.45E-15 |
| ENSG00000172935 | MRGPRF | -1 | 0.15 | 4.67E-12 | 2.94E-11 |
| ENSG00000164296 | TIGD6 | -1 | 0.16 | 5.39E-11 | 3.18E-10 |
| ENSG00000135119 | RNFT2 | -1 | 0.17 | 2.65E-10 | 1.49E-09 |
| ENSG00000102362 | SYTL4 | -1 | 0.17 | 4.82E-10 | 2.66E-09 |
| ENSG00000269427 | | -1 | 0.18 | 1.73E-09 | 9.19E-09 |
| ENSG00000164841 | TMEM74 | -1 | 0.21 | 1.21E-07 | 5.58E-07 |
| ENSG00000188338 | SLC38A3 | -1 | 0.32 | 1.22E-04 | 4.18E-04 |
| ENSG00000161132 | PRODHLP | -1 | 0.32 | 1.29E-04 | 4.43E-04 |
| ENSG00000249889 | ALG1L11P | -1 | 0.45 | 1.65E-03 | 4.85E-03 |
| ENSG00000186007 | LEMD1 | -1 | 0.82 | 3.32E-03 | 9.24E-03 |
| ENSG00000162873 | KLHDC8A | -1 | 0.64 | 4.99E-03 | 1.34E-02 |
| ENSG00000216867 | RPL22P12 | -1 | 0.67 | 5.18E-03 | 1.39E-02 |

| | | | | | |
|---|---|---|---|---|---|
| ENSG00000168575 | SLC20A2 | -0.99 | 0.04 | 3.11E-172 | 4.24E-170 |
| ENSG00000163359 | COL6A3 | -0.99 | 0.04 | 3.52E-127 | 2.85E-125 |
| ENSG00000172936 | MYD88 | -0.99 | 0.05 | 3.30E-97 | 1.83E-95 |
| ENSG00000003056 | M6PR | -0.99 | 0.05 | 1.82E-89 | 8.83E-88 |
| ENSG00000116984 | MTR | -0.99 | 0.05 | 2.60E-89 | 1.26E-87 |
| ENSG00000110042 | DTX4 | -0.99 | 0.05 | 4.06E-77 | 1.61E-75 |
| ENSG00000076242 | MLH1 | -0.99 | 0.05 | 1.68E-73 | 6.20E-72 |
| ENSG00000120314 | WDR55 | -0.99 | 0.06 | 6.90E-73 | 2.51E-71 |
| ENSG00000179314 | WSCD1 | -0.99 | 0.06 | 1.05E-72 | 3.79E-71 |
| ENSG00000120686 | UFM1 | -0.99 | 0.06 | 7.00E-69 | 2.34E-67 |
| ENSG00000078328 | RBFOX1 | -0.99 | 0.06 | 2.58E-67 | 8.33E-66 |
| ENSG00000197905 | TEAD4 | -0.99 | 0.06 | 2.60E-57 | 6.89E-56 |
| ENSG00000163626 | COX18 | -0.99 | 0.06 | 9.07E-54 | 2.24E-52 |
| ENSG00000097046 | CDC7 | -0.99 | 0.07 | 5.41E-49 | 1.19E-47 |
| ENSG00000165138 | ANKS6 | -0.99 | 0.07 | 1.41E-43 | 2.78E-42 |
| ENSG00000063660 | GPC1 | -0.99 | 0.08 | 2.03E-37 | 3.41E-36 |
| ENSG00000114019 | AMOTL2 | -0.99 | 0.08 | 3.40E-37 | 5.68E-36 |
| ENSG00000204519 | ZNF551 | -0.99 | 0.08 | 9.19E-35 | 1.43E-33 |
| ENSG00000145198 | VWA5B2 | -0.99 | 0.13 | 1.05E-14 | 7.67E-14 |
| ENSG00000162999 | DUSP19 | -0.99 | 0.14 | 3.19E-13 | 2.16E-12 |
| ENSG00000176533 | GNG7 | -0.99 | 0.15 | 5.57E-12 | 3.49E-11 |
| ENSG00000168824 | NSG1 | -0.99 | 0.23 | 1.33E-06 | 5.60E-06 |
| ENSG00000143867 | OSR1 | -0.99 | 0.26 | 1.03E-05 | 3.99E-05 |
| ENSG00000164199 | ADGRV1 | -0.99 | 0.37 | 4.49E-04 | 1.44E-03 |
| ENSG00000182459 | TEX19 | -0.99 | 0.6 | 4.84E-03 | 1.31E-02 |
| ENSG00000067955 | CBFB | -0.98 | 0.04 | 1.55E-121 | 1.17E-119 |
| ENSG00000070814 | TCOF1 | -0.98 | 0.05 | 6.15E-96 | 3.35E-94 |
| ENSG00000164045 | CDC25A | -0.98 | 0.05 | 9.67E-92 | 4.92E-90 |
| ENSG00000073111 | MCM2 | -0.98 | 0.05 | 6.82E-81 | 2.84E-79 |
| ENSG00000076513 | ANKRD13A | -0.98 | 0.06 | 1.74E-70 | 6.04E-69 |
| ENSG00000133393 | CEP20 | -0.98 | 0.06 | 7.84E-57 | 2.06E-55 |
| ENSG00000172175 | MALT1 | -0.98 | 0.06 | 1.98E-54 | 4.98E-53 |
| ENSG00000133863 | TEX15 | -0.98 | 0.07 | 2.13E-52 | 5.08E-51 |
| ENSG00000137501 | SYTL2 | -0.98 | 0.07 | 4.95E-51 | 1.14E-49 |
| ENSG00000079277 | MKNK1 | -0.98 | 0.07 | 1.63E-46 | 3.39E-45 |
| ENSG00000156467 | UQCRB | -0.98 | 0.07 | 1.44E-43 | 2.83E-42 |
| ENSG00000136859 | ANGPTL2 | -0.98 | 0.08 | 4.97E-33 | 7.31E-32 |
| ENSG00000091482 | SMPX | -0.98 | 0.09 | 8.88E-30 | 1.19E-28 |
| ENSG00000198001 | IRAK4 | -0.98 | 0.1 | 1.12E-24 | 1.25E-23 |
| ENSG00000143375 | CGN | -0.98 | 0.1 | 1.25E-24 | 1.38E-23 |
| ENSG00000170801 | HTRA3 | -0.98 | 0.11 | 1.47E-19 | 1.33E-18 |
| ENSG00000158825 | CDA | -0.98 | 0.13 | 8.90E-16 | 6.82E-15 |
| ENSG00000162542 | TMCO4 | -0.98 | 0.14 | 4.82E-14 | 3.40E-13 |
| ENSG00000180739 | S1PR5 | -0.98 | 0.15 | 3.33E-12 | 2.12E-11 |
| ENSG00000291049 | FAM86B3P | -0.98 | 0.17 | 2.95E-10 | 1.65E-09 |
| ENSG00000224940 | PRRT4 | -0.98 | 0.17 | 3.48E-10 | 1.94E-09 |
| ENSG00000172995 | ARPP21 | -0.98 | 0.17 | 1.86E-09 | 9.84E-09 |
| ENSG00000186777 | ZNF732 | -0.98 | 0.2 | 5.77E-08 | 2.73E-07 |
| ENSG00000042832 | TG | -0.98 | 0.2 | 6.02E-08 | 2.84E-07 |
| ENSG00000232630 | PRPS1P2 | -0.98 | 0.2 | 7.79E-08 | 3.65E-07 |

| | | | | | |
|---|---|---|---|---|---|
| ENSG00000164651 | SP8 | -0.98 | 0.36 | 4.26E-04 | 1.37E-03 |
| ENSG00000148604 | RGR | -0.98 | 0.45 | 1.87E-03 | 5.45E-03 |
| ENSG00000010671 | BTK | -0.98 | 0.49 | 2.80E-03 | 7.90E-03 |
| ENSG00000167895 | TMC8 | -0.98 | 0.5 | 2.89E-03 | 8.14E-03 |
| ENSG00000251155 | SEPTIN14P4 | -0.98 | 0.53 | 3.45E-03 | 9.58E-03 |
| ENSG00000163632 | C3orf49 | -0.98 | 0.55 | 3.91E-03 | 1.07E-02 |
| ENSG00000123416 | TUBA1B | -0.97 | 0.05 | 1.71E-79 | 6.97E-78 |
| ENSG00000100077 | GRK3 | -0.97 | 0.06 | 1.11E-69 | 3.77E-68 |
| ENSG00000122591 | HYCC1 | -0.97 | 0.06 | 1.46E-64 | 4.50E-63 |
| ENSG00000014919 | COX15 | -0.97 | 0.06 | 1.04E-62 | 3.04E-61 |
| ENSG00000100505 | TRIM9 | -0.97 | 0.06 | 8.44E-55 | 2.14E-53 |
| ENSG00000134352 | IL6ST | -0.97 | 0.07 | 4.85E-49 | 1.07E-47 |
| ENSG00000078687 | TNRC6C | -0.97 | 0.07 | 1.89E-48 | 4.07E-47 |
| ENSG00000153253 | SCN3A | -0.97 | 0.08 | 2.07E-38 | 3.59E-37 |
| ENSG00000176971 | FIBIN | -0.97 | 0.08 | 1.13E-35 | 1.82E-34 |
| ENSG00000178966 | RMI1 | -0.97 | 0.09 | 1.55E-27 | 1.92E-26 |
| ENSG00000179954 | SSC5D | -0.97 | 0.1 | 6.12E-24 | 6.66E-23 |
| ENSG00000120896 | SORBS3 | -0.97 | 0.1 | 1.25E-23 | 1.34E-22 |
| ENSG00000120539 | MASTL | -0.97 | 0.1 | 6.28E-23 | 6.57E-22 |
| ENSG00000289731 | | -0.97 | 0.1 | 1.25E-22 | 1.29E-21 |
| ENSG00000172346 | CSDC2 | -0.97 | 0.11 | 2.09E-21 | 2.06E-20 |
| ENSG00000118898 | PPL | -0.97 | 0.11 | 8.71E-21 | 8.35E-20 |
| ENSG00000144061 | NPHP1 | -0.97 | 0.12 | 5.71E-18 | 4.85E-17 |
| ENSG00000021762 | OSBPL5 | -0.97 | 0.12 | 2.01E-17 | 1.67E-16 |
| ENSG00000151876 | FBXO4 | -0.97 | 0.13 | 3.77E-15 | 2.82E-14 |
| ENSG00000006283 | CACNA1G | -0.97 | 0.19 | 2.54E-08 | 1.23E-07 |
| ENSG00000266651 | | -0.97 | 0.25 | 8.76E-06 | 3.40E-05 |
| ENSG00000119514 | GALNT12 | -0.97 | 0.28 | 3.79E-05 | 1.38E-04 |
| ENSG00000256673 | | -0.97 | 0.36 | 5.03E-04 | 1.60E-03 |
| ENSG00000169271 | HSPB3 | -0.97 | 0.41 | 1.10E-03 | 3.32E-03 |
| ENSG00000238097 | LINC02037 | -0.97 | 0.89 | 1.61E-03 | 4.73E-03 |
| ENSG00000144596 | GRIP2 | -0.97 | 0.49 | 2.88E-03 | 8.10E-03 |
| ENSG00000143167 | GPA33 | -0.97 | 0.84 | 3.01E-03 | 8.47E-03 |
| ENSG00000271367 | | -0.97 | 0.68 | 5.96E-03 | 1.58E-02 |
| ENSG00000124541 | RRP36 | -0.96 | 0.05 | 1.44E-91 | 7.30E-90 |
| ENSG00000145604 | SKP2 | -0.96 | 0.05 | 4.52E-90 | 2.20E-88 |
| ENSG00000119321 | FKBP15 | -0.96 | 0.05 | 2.86E-82 | 1.23E-80 |
| ENSG00000068615 | REEP1 | -0.96 | 0.05 | 7.21E-76 | 2.79E-74 |
| ENSG00000101843 | PSMD10 | -0.96 | 0.05 | 4.90E-70 | 1.68E-68 |
| ENSG00000119616 | FCF1 | -0.96 | 0.06 | 8.65E-69 | 2.88E-67 |
| ENSG00000088387 | DOCK9 | -0.96 | 0.06 | 1.46E-65 | 4.58E-64 |
| ENSG00000146918 | NCAPG2 | -0.96 | 0.06 | 2.21E-62 | 6.47E-61 |
| ENSG00000170624 | SGCD | -0.96 | 0.06 | 7.84E-59 | 2.15E-57 |
| ENSG00000182263 | FIGN | -0.96 | 0.06 | 6.61E-52 | 1.56E-50 |
| ENSG00000265241 | RBM8A | -0.96 | 0.07 | 1.31E-45 | 2.68E-44 |
| ENSG00000102893 | PHKB | -0.96 | 0.07 | 1.07E-44 | 2.15E-43 |
| ENSG00000162458 | FBLIM1 | -0.96 | 0.08 | 5.50E-36 | 8.91E-35 |
| ENSG00000124785 | NRN1 | -0.96 | 0.08 | 3.65E-33 | 5.41E-32 |
| ENSG00000183943 | PRKX | -0.96 | 0.08 | 1.01E-31 | 1.43E-30 |
| ENSG00000111331 | OAS3 | -0.96 | 0.08 | 1.10E-31 | 1.56E-30 |

| | | | | | |
|---|---|---|---|---|---|
| ENSG00000140463 | BBS4 | -0.96 | 0.08 | 3.17E-31 | 4.45E-30 |
| ENSG00000116396 | KCNC4 | -0.96 | 0.09 | 2.53E-27 | 3.10E-26 |
| ENSG00000134326 | CMPK2 | -0.96 | 0.09 | 1.33E-25 | 1.54E-24 |
| ENSG00000130038 | CRACR2A | -0.96 | 0.1 | 2.77E-24 | 3.04E-23 |
| ENSG00000137700 | SLC37A4 | -0.96 | 0.12 | 5.39E-17 | 4.39E-16 |
| ENSG00000122592 | HOXA7 | -0.96 | 0.2 | 8.37E-08 | 3.90E-07 |
| ENSG00000175274 | TP53I11 | -0.96 | 0.21 | 5.05E-07 | 2.21E-06 |
| ENSG00000155926 | SLA | -0.96 | 0.26 | 1.85E-05 | 6.96E-05 |
| ENSG00000290728 | | -0.96 | 0.27 | 2.20E-05 | 8.21E-05 |
| ENSG00000273274 | ZBTB8B | -0.96 | 0.33 | 2.87E-04 | 9.43E-04 |
| ENSG00000268573 | | -0.96 | 0.34 | 3.83E-04 | 1.24E-03 |
| ENSG00000188282 | RUFY4 | -0.96 | 0.4 | 1.11E-03 | 3.34E-03 |
| ENSG00000188833 | ENTPD8 | -0.96 | 0.4 | 1.21E-03 | 3.63E-03 |
| ENSG00000171840 | NINJ2 | -0.96 | 0.44 | 1.86E-03 | 5.41E-03 |
| ENSG00000275088 | | -0.96 | 0.45 | 2.15E-03 | 6.20E-03 |
| ENSG00000117519 | CNN3 | -0.95 | 0.04 | 7.21E-141 | 6.93E-139 |
| ENSG00000171617 | ENC1 | -0.95 | 0.05 | 1.07E-92 | 5.57E-91 |
| ENSG00000174306 | ZHX3 | -0.95 | 0.05 | 1.37E-90 | 6.76E-89 |
| ENSG00000141556 | TBCD | -0.95 | 0.05 | 1.12E-88 | 5.35E-87 |
| ENSG00000147526 | TACC1 | -0.95 | 0.05 | 3.18E-82 | 1.36E-80 |
| ENSG00000103994 | ZNF106 | -0.95 | 0.05 | 8.14E-76 | 3.13E-74 |
| ENSG00000111615 | KRR1 | -0.95 | 0.05 | 9.19E-71 | 3.22E-69 |
| ENSG00000092607 | TBX15 | -0.95 | 0.05 | 1.98E-70 | 6.87E-69 |
| ENSG00000129347 | KRI1 | -0.95 | 0.06 | 3.82E-54 | 9.57E-53 |
| ENSG00000149636 | DSN1 | -0.95 | 0.07 | 8.25E-49 | 1.80E-47 |
| ENSG00000159363 | ATP13A2 | -0.95 | 0.09 | 1.47E-25 | 1.69E-24 |
| ENSG00000058404 | CAMK2B | -0.95 | 0.1 | 3.11E-21 | 3.05E-20 |
| ENSG00000184371 | CSF1 | -0.95 | 0.12 | 9.44E-17 | 7.61E-16 |
| ENSG00000244586 | WNT5A-AS1 | -0.95 | 0.13 | 3.49E-14 | 2.48E-13 |
| ENSG00000111424 | VDR | -0.95 | 0.14 | 2.60E-12 | 1.67E-11 |
| ENSG00000268940 | CT45A1 | -0.95 | 0.17 | 9.01E-10 | 4.87E-09 |
| ENSG00000168772 | CXXC4 | -0.95 | 0.17 | 1.27E-09 | 6.81E-09 |
| ENSG00000288940 | | -0.95 | 0.21 | 5.44E-07 | 2.37E-06 |
| ENSG00000162976 | SLC66A3 | -0.95 | 0.22 | 1.76E-06 | 7.30E-06 |
| ENSG00000224420 | ADM5 | -0.95 | 0.24 | 5.46E-06 | 2.17E-05 |
| ENSG00000226887 | ERVMER34-1 | -0.95 | 0.31 | 1.51E-04 | 5.15E-04 |
| ENSG00000260698 | | -0.95 | 0.32 | 2.19E-04 | 7.33E-04 |
| ENSG00000243709 | LEFTY1 | -0.95 | 0.4 | 1.21E-03 | 3.63E-03 |
| ENSG00000273812 | LINC02970 | -0.95 | 0.4 | 1.27E-03 | 3.78E-03 |
| ENSG00000142583 | SLC2A5 | -0.95 | 0.46 | 2.73E-03 | 7.73E-03 |
| ENSG00000276136 | | -0.95 | 0.48 | 3.27E-03 | 9.12E-03 |
| ENSG00000285677 | | -0.95 | 0.5 | 3.84E-03 | 1.06E-02 |
| ENSG00000236992 | RPL12P12 | -0.95 | 0.51 | 3.86E-03 | 1.06E-02 |
| ENSG00000049540 | ELN | -0.95 | 0.52 | 4.31E-03 | 1.17E-02 |
| ENSG00000286753 | | -0.95 | 0.54 | 4.89E-03 | 1.32E-02 |
| ENSG00000218725 | B3GALNT2P1 | -0.95 | 0.58 | 5.78E-03 | 1.54E-02 |
| ENSG00000006047 | YBX2 | -0.95 | 0.62 | 6.44E-03 | 1.69E-02 |
| ENSG00000154678 | PDE1C | -0.95 | 0.67 | 6.64E-03 | 1.74E-02 |
| ENSG00000260388 | LINC00562 | -0.95 | 0.63 | 6.66E-03 | 1.75E-02 |
| ENSG00000035862 | TIMP2 | -0.94 | 0.04 | 3.89E-135 | 3.51E-133 |

| | | | | | |
|---|---|---|---|---|---|
| ENSG00000106723 | SPIN1 | -0.94 | 0.04 | 8.54E-107 | 5.43E-105 |
| ENSG00000115561 | CHMP3 | -0.94 | 0.05 | 3.90E-76 | 1.51E-74 |
| ENSG00000107742 | SPOCK2 | -0.94 | 0.05 | 7.19E-73 | 2.60E-71 |
| ENSG00000120875 | DUSP4 | -0.94 | 0.05 | 7.02E-72 | 2.50E-70 |
| ENSG00000138073 | PREB | -0.94 | 0.05 | 1.21E-71 | 4.29E-70 |
| ENSG00000103356 | EARS2 | -0.94 | 0.06 | 3.75E-58 | 1.01E-56 |
| ENSG00000183808 | RBM12B | -0.94 | 0.06 | 5.20E-56 | 1.35E-54 |
| ENSG00000165406 | MARCHF8 | -0.94 | 0.06 | 3.86E-52 | 9.14E-51 |
| ENSG00000157450 | RNF111 | -0.94 | 0.06 | 9.83E-52 | 2.31E-50 |
| ENSG00000173456 | RNF26 | -0.94 | 0.08 | 1.25E-35 | 2.00E-34 |
| ENSG00000169188 | APEX2 | -0.94 | 0.08 | 6.00E-35 | 9.38E-34 |
| ENSG00000213186 | TRIM59 | -0.94 | 0.08 | 8.01E-34 | 1.21E-32 |
| ENSG00000186470 | BTN3A2 | -0.94 | 0.08 | 2.58E-33 | 3.83E-32 |
| ENSG00000273611 | ZNHIT3 | -0.94 | 0.08 | 8.70E-33 | 1.27E-31 |
| ENSG00000075131 | TIPIN | -0.94 | 0.08 | 5.21E-32 | 7.45E-31 |
| ENSG00000012171 | SEMA3B | -0.94 | 0.1 | 5.63E-23 | 5.89E-22 |
| ENSG00000130775 | THEMIS2 | -0.94 | 0.1 | 1.37E-22 | 1.41E-21 |
| ENSG00000171428 | NAT1 | -0.94 | 0.12 | 2.82E-16 | 2.22E-15 |
| ENSG00000188549 | CCDC9B | -0.94 | 0.17 | 3.18E-09 | 1.65E-08 |
| ENSG00000162009 | SSTR5 | -0.94 | 0.19 | 6.63E-08 | 3.12E-07 |
| ENSG00000281103 | TRG-AS1 | -0.94 | 0.2 | 1.82E-07 | 8.26E-07 |
| ENSG00000198768 | APCDD1L | -0.94 | 0.22 | 1.85E-06 | 7.67E-06 |
| ENSG00000236869 | ZKSCAN7-AS1 | -0.94 | 0.28 | 5.24E-05 | 1.88E-04 |
| ENSG00000004848 | ARX | -0.94 | 0.29 | 1.17E-04 | 4.05E-04 |
| ENSG00000188868 | ZNF563 | -0.94 | 0.32 | 2.62E-04 | 8.67E-04 |
| ENSG00000165475 | CRYL1 | -0.94 | 0.36 | 6.67E-04 | 2.08E-03 |
| ENSG00000162105 | SHANK2 | -0.94 | 0.39 | 1.20E-03 | 3.61E-03 |
| ENSG00000229647 | MYOSLID | -0.94 | 0.57 | 6.01E-03 | 1.59E-02 |
| ENSG00000272156 | | -0.94 | 0.68 | 6.87E-03 | 1.80E-02 |
| ENSG00000132646 | PCNA | -0.93 | 0.05 | 1.16E-74 | 4.35E-73 |
| ENSG00000076003 | MCM6 | -0.93 | 0.05 | 7.85E-70 | 2.68E-68 |
| ENSG00000182628 | SKA2 | -0.93 | 0.05 | 1.31E-68 | 4.35E-67 |
| ENSG00000152818 | UTRN | -0.93 | 0.05 | 2.46E-67 | 7.95E-66 |
| ENSG00000170322 | NFRKB | -0.93 | 0.06 | 1.98E-51 | 4.61E-50 |
| ENSG00000134321 | RSAD2 | -0.93 | 0.07 | 1.69E-41 | 3.17E-40 |
| ENSG00000166402 | TUB | -0.93 | 0.08 | 1.20E-34 | 1.86E-33 |
| ENSG00000171310 | CHST11 | -0.93 | 0.08 | 3.58E-30 | 4.84E-29 |
| ENSG00000134996 | OSTF1 | -0.93 | 0.09 | 2.47E-25 | 2.82E-24 |
| ENSG00000136546 | SCN7A | -0.93 | 0.09 | 3.28E-25 | 3.73E-24 |
| ENSG00000106333 | PCOLCE | -0.93 | 0.1 | 5.36E-22 | 5.40E-21 |
| ENSG00000125730 | C3 | -0.93 | 0.12 | 2.35E-16 | 1.86E-15 |
| ENSG00000122547 | EEPD1 | -0.93 | 0.2 | 2.63E-07 | 1.18E-06 |
| ENSG00000149527 | PLCH2 | -0.93 | 0.21 | 6.37E-07 | 2.76E-06 |
| ENSG00000287126 | | -0.93 | 0.21 | 1.34E-06 | 5.62E-06 |
| ENSG00000177570 | SAMD12 | -0.93 | 0.22 | 1.97E-06 | 8.13E-06 |
| ENSG00000133983 | COX16 | -0.93 | 0.23 | 6.24E-06 | 2.46E-05 |
| ENSG00000185942 | NKAIN3 | -0.93 | 0.26 | 2.35E-05 | 8.74E-05 |
| ENSG00000225756 | DBH-AS1 | -0.93 | 0.27 | 5.86E-05 | 2.09E-04 |
| ENSG00000158270 | COLEC12 | -0.93 | 0.36 | 7.07E-04 | 2.20E-03 |
| ENSG00000260743 | | -0.93 | 0.42 | 2.17E-03 | 6.25E-03 |

| | | | | | |
|---|---|---|---|---|---|
| ENSG00000269855 | RNF225 | -0.93 | 0.43 | 2.33E-03 | 6.69E-03 |
| ENSG00000287787 | | -0.93 | 0.44 | 2.65E-03 | 7.51E-03 |
| ENSG00000274752 | TRBV12-3 | -0.93 | 0.46 | 3.20E-03 | 8.94E-03 |
| ENSG00000187688 | TRPV2 | -0.93 | 0.49 | 4.08E-03 | 1.12E-02 |
| ENSG00000221539 | SNORD99 | -0.93 | 0.52 | 4.73E-03 | 1.28E-02 |
| ENSG00000158869 | FCER1G | -0.93 | 0.54 | 5.44E-03 | 1.45E-02 |
| ENSG00000272100 | | -0.93 | 0.57 | 6.25E-03 | 1.65E-02 |
| ENSG00000288580 | | -0.93 | 0.59 | 6.77E-03 | 1.78E-02 |
| ENSG00000151952 | TMEM132D | -0.93 | 0.61 | 6.84E-03 | 1.79E-02 |
| ENSG00000101138 | CSTF1 | -0.92 | 0.04 | 4.89E-106 | 3.07E-104 |
| ENSG00000071242 | RPS6KA2 | -0.92 | 0.05 | 6.41E-91 | 3.20E-89 |
| ENSG00000151576 | QTRT2 | -0.92 | 0.05 | 4.31E-75 | 1.63E-73 |
| ENSG00000173852 | DPY19L1 | -0.92 | 0.05 | 1.65E-68 | 5.45E-67 |
| ENSG00000102359 | SRPX2 | -0.92 | 0.06 | 4.11E-63 | 1.22E-61 |
| ENSG00000166822 | TMEM170A | -0.92 | 0.07 | 1.36E-45 | 2.77E-44 |
| ENSG00000106462 | EZH2 | -0.92 | 0.07 | 3.94E-44 | 7.81E-43 |
| ENSG00000275835 | TUBGCP5 | -0.92 | 0.07 | 5.65E-41 | 1.05E-39 |
| ENSG00000052850 | ALX4 | -0.92 | 0.08 | 7.59E-35 | 1.18E-33 |
| ENSG00000197444 | OGDHL | -0.92 | 0.09 | 2.14E-26 | 2.54E-25 |
| ENSG00000184208 | C22orf46P | -0.92 | 0.09 | 3.42E-26 | 4.02E-25 |
| ENSG00000147121 | KRBOX4 | -0.92 | 0.09 | 2.13E-24 | 2.35E-23 |
| ENSG00000164087 | POC1A | -0.92 | 0.09 | 1.46E-23 | 1.56E-22 |
| ENSG00000159259 | CHAF1B | -0.92 | 0.1 | 8.15E-23 | 8.46E-22 |
| ENSG00000168876 | ANKRD49 | -0.92 | 0.1 | 9.51E-22 | 9.48E-21 |
| ENSG00000099953 | MMP11 | -0.92 | 0.14 | 2.18E-11 | 1.32E-10 |
| ENSG00000196081 | ZNF724 | -0.92 | 0.18 | 1.85E-08 | 9.08E-08 |
| ENSG00000181029 | TRAPPC5 | -0.92 | 0.25 | 2.57E-05 | 9.50E-05 |
| ENSG00000260211 | | -0.92 | 0.31 | 2.82E-04 | 9.26E-04 |
| ENSG00000179057 | IGSF22 | -0.92 | 0.36 | 8.66E-04 | 2.66E-03 |
| ENSG00000169783 | LINGO1 | -0.92 | 0.42 | 2.03E-03 | 5.88E-03 |
| ENSG00000270457 | | -0.92 | 0.87 | 2.27E-03 | 6.52E-03 |
| ENSG00000267500 | ZNF887P | -0.92 | 0.44 | 2.96E-03 | 8.33E-03 |
| ENSG00000233714 | | -0.92 | 0.84 | 3.14E-03 | 8.80E-03 |
| ENSG00000234311 | LINC03068 | -0.92 | 0.46 | 3.48E-03 | 9.65E-03 |
| ENSG00000188162 | OTOG | -0.92 | 0.46 | 3.57E-03 | 9.88E-03 |
| ENSG00000287543 | | -0.92 | 0.53 | 5.67E-03 | 1.51E-02 |
| ENSG00000258902 | | -0.92 | 0.63 | 7.69E-03 | 1.99E-02 |
| ENSG00000206422 | LRRC30 | -0.92 | 0.67 | 7.75E-03 | 2.01E-02 |
| ENSG00000102317 | RBM3 | -0.91 | 0.04 | 1.77E-120 | 1.31E-118 |
| ENSG00000100320 | RBFOX2 | -0.91 | 0.04 | 2.16E-98 | 1.22E-96 |
| ENSG00000116199 | FAM20B | -0.91 | 0.04 | 7.64E-96 | 4.14E-94 |
| ENSG00000139209 | SLC38A4 | -0.91 | 0.05 | 8.17E-69 | 2.72E-67 |
| ENSG00000146674 | IGFBP3 | -0.91 | 0.05 | 7.74E-68 | 2.52E-66 |
| ENSG00000010803 | SCMH1 | -0.91 | 0.05 | 4.81E-62 | 1.40E-60 |
| ENSG00000133454 | MYO18B | -0.91 | 0.06 | 3.25E-53 | 7.91E-52 |
| ENSG00000155846 | PPARGC1B | -0.91 | 0.06 | 5.44E-49 | 1.19E-47 |
| ENSG00000129682 | FGF13 | -0.91 | 0.06 | 1.60E-48 | 3.45E-47 |
| ENSG00000196083 | IL1RAP | -0.91 | 0.07 | 7.32E-40 | 1.31E-38 |
| ENSG00000204267 | TAP2 | -0.91 | 0.07 | 1.77E-37 | 2.97E-36 |
| ENSG00000088756 | ARHGAP28 | -0.91 | 0.08 | 1.11E-28 | 1.43E-27 |

| | | | | | |
|---|---|---|---|---|---|
| ENSG00000102710 | SUPT20H | -0.91 | 0.09 | 6.76E-23 | 7.05E-22 |
| ENSG00000133678 | TMEM254 | -0.91 | 0.11 | 5.10E-19 | 4.53E-18 |
| ENSG00000150907 | FOXO1 | -0.91 | 0.12 | 3.58E-15 | 2.68E-14 |
| ENSG00000100162 | CENPM | -0.91 | 0.12 | 6.60E-15 | 4.88E-14 |
| ENSG00000010704 | HFE | -0.91 | 0.12 | 8.14E-15 | 5.99E-14 |
| ENSG00000081923 | ATP8B1 | -0.91 | 0.16 | 7.59E-10 | 4.13E-09 |
| ENSG00000106479 | ZNF862 | -0.91 | 0.16 | 1.01E-09 | 5.46E-09 |
| ENSG00000087116 | ADAMTS2 | -0.91 | 0.18 | 4.69E-08 | 2.23E-07 |
| ENSG00000187068 | C3orf70 | -0.91 | 0.2 | 7.14E-07 | 3.08E-06 |
| ENSG00000246859 | STARD4-AS1 | -0.91 | 0.21 | 2.23E-06 | 9.16E-06 |
| ENSG00000141404 | GNAL | -0.91 | 0.33 | 5.35E-04 | 1.70E-03 |
| ENSG00000251819 | RNU6-322P | -0.91 | 0.35 | 7.42E-04 | 2.30E-03 |
| ENSG00000248559 | | -0.91 | 0.48 | 4.32E-03 | 1.17E-02 |
| ENSG00000178217 | SH2D4B | -0.91 | 0.5 | 4.91E-03 | 1.32E-02 |
| ENSG00000198326 | TMEM239 | -0.91 | 0.74 | 7.07E-03 | 1.85E-02 |
| ENSG00000254303 | | -0.91 | 0.57 | 7.26E-03 | 1.89E-02 |
| ENSG00000100060 | MFNG | -0.91 | 0.72 | 7.64E-03 | 1.98E-02 |
| ENSG00000225458 | | -0.91 | 0.69 | 8.23E-03 | 2.12E-02 |
| ENSG00000145632 | PLK2 | -0.9 | 0.05 | 1.85E-87 | 8.54E-86 |
| ENSG00000138771 | SHROOM3 | -0.9 | 0.05 | 1.55E-84 | 6.94E-83 |
| ENSG00000113532 | ST8SIA4 | -0.9 | 0.05 | 1.10E-83 | 4.87E-82 |
| ENSG00000141385 | AFG3L2 | -0.9 | 0.05 | 4.73E-75 | 1.79E-73 |
| ENSG00000127124 | HIVEP3 | -0.9 | 0.05 | 3.21E-69 | 1.08E-67 |
| ENSG00000165025 | SYK | -0.9 | 0.06 | 1.48E-55 | 3.80E-54 |
| ENSG00000114993 | RTKN | -0.9 | 0.06 | 4.97E-53 | 1.21E-51 |
| ENSG00000244462 | RBM12 | -0.9 | 0.06 | 6.87E-53 | 1.66E-51 |
| ENSG00000100749 | VRK1 | -0.9 | 0.07 | 1.09E-41 | 2.06E-40 |
| ENSG00000112419 | PHACTR2 | -0.9 | 0.07 | 9.30E-41 | 1.71E-39 |
| ENSG00000131019 | ULBP3 | -0.9 | 0.08 | 8.84E-33 | 1.29E-31 |
| ENSG00000149474 | KAT14 | -0.9 | 0.08 | 7.58E-30 | 1.01E-28 |
| ENSG00000145439 | CBR4 | -0.9 | 0.08 | 5.49E-29 | 7.14E-28 |
| ENSG00000185252 | ZNF74 | -0.9 | 0.09 | 6.24E-27 | 7.55E-26 |
| ENSG00000048342 | CC2D2A | -0.9 | 0.1 | 1.43E-20 | 1.36E-19 |
| ENSG00000278619 | MRM1 | -0.9 | 0.13 | 2.35E-13 | 1.60E-12 |
| ENSG00000170074 | FAM153A | -0.9 | 0.14 | 1.47E-11 | 8.98E-11 |
| ENSG00000127720 | METTL25 | -0.9 | 0.15 | 4.80E-10 | 2.65E-09 |
| ENSG00000124191 | TOX2 | -0.9 | 0.16 | 2.11E-09 | 1.11E-08 |
| ENSG00000272077 | | -0.9 | 0.17 | 9.01E-09 | 4.52E-08 |
| ENSG00000188596 | CFAP54 | -0.9 | 0.18 | 7.75E-08 | 3.63E-07 |
| ENSG00000182185 | RAD51B | -0.9 | 0.19 | 1.88E-07 | 8.55E-07 |
| ENSG00000220343 | PPIAP74 | -0.9 | 0.2 | 3.45E-07 | 1.53E-06 |
| ENSG00000182389 | CACNB4 | -0.9 | 0.26 | 4.87E-05 | 1.75E-04 |
| ENSG00000198300 | PEG3 | -0.9 | 0.33 | 5.64E-04 | 1.78E-03 |
| ENSG00000238186 | | -0.9 | 0.33 | 6.11E-04 | 1.92E-03 |
| ENSG00000257953 | | -0.9 | 0.44 | 3.36E-03 | 9.34E-03 |
| ENSG00000115085 | ZAP70 | -0.9 | 0.5 | 5.21E-03 | 1.40E-02 |
| ENSG00000271776 | | -0.9 | 0.5 | 5.63E-03 | 1.50E-02 |
| ENSG00000182393 | IFNL1 | -0.9 | 0.52 | 6.26E-03 | 1.65E-02 |
| ENSG00000184544 | DHRS7C | -0.9 | 0.65 | 8.81E-03 | 2.26E-02 |
| ENSG00000231054 | | -0.9 | 0.66 | 8.86E-03 | 2.27E-02 |

| | | | | | |
|---|---|---|---|---|---|
| ENSG00000091409 | ITGA6 | -0.89 | 0.04 | 3.18E-120 | 2.34E-118 |
| ENSG00000012232 | EXTL3 | -0.89 | 0.04 | 5.41E-103 | 3.25E-101 |
| ENSG00000180801 | ARSJ | -0.89 | 0.04 | 8.40E-92 | 4.29E-90 |
| ENSG00000166272 | WBP1L | -0.89 | 0.05 | 5.43E-85 | 2.46E-83 |
| ENSG00000082781 | ITGB5 | -0.89 | 0.05 | 1.51E-62 | 4.42E-61 |
| ENSG00000165219 | GAPVD1 | -0.89 | 0.06 | 3.36E-54 | 8.45E-53 |
| ENSG00000106477 | CEP41 | -0.89 | 0.06 | 3.90E-45 | 7.88E-44 |
| ENSG00000101166 | PRELID3B | -0.89 | 0.06 | 8.89E-45 | 1.79E-43 |
| ENSG00000168502 | MTCL1 | -0.89 | 0.07 | 2.34E-40 | 4.23E-39 |
| ENSG00000132825 | PPP1R3D | -0.89 | 0.07 | 3.05E-37 | 5.09E-36 |
| ENSG00000163964 | PIGX | -0.89 | 0.07 | 5.96E-36 | 9.64E-35 |
| ENSG00000111335 | OAS2 | -0.89 | 0.08 | 7.77E-33 | 1.14E-31 |
| ENSG00000005469 | CROT | -0.89 | 0.08 | 2.60E-29 | 3.41E-28 |
| ENSG00000275004 | ZNF280B | -0.89 | 0.09 | 6.76E-25 | 7.61E-24 |
| ENSG00000056972 | TRAF3IP2 | -0.89 | 0.1 | 8.26E-21 | 7.93E-20 |
| ENSG00000100413 | POLR3H | -0.89 | 0.1 | 3.95E-20 | 3.69E-19 |
| ENSG00000174791 | RIN1 | -0.89 | 0.1 | 4.60E-20 | 4.29E-19 |
| ENSG00000152926 | ZNF117 | -0.89 | 0.1 | 2.96E-19 | 2.67E-18 |
| ENSG00000108439 | PNPO | -0.89 | 0.1 | 1.38E-18 | 1.20E-17 |
| ENSG00000160229 | ZNF66 | -0.89 | 0.13 | 6.22E-13 | 4.13E-12 |
| ENSG00000099377 | HSD3B7 | -0.89 | 0.13 | 7.13E-13 | 4.72E-12 |
| ENSG00000291186 | | -0.89 | 0.14 | 6.45E-12 | 4.02E-11 |
| ENSG00000115425 | PECR | -0.89 | 0.15 | 4.05E-10 | 2.25E-09 |
| ENSG00000280187 | | -0.89 | 0.16 | 1.16E-09 | 6.25E-09 |
| ENSG00000135678 | CPM | -0.89 | 0.2 | 5.92E-07 | 2.57E-06 |
| ENSG00000229380 | PRKAR1B-AS2 | -0.89 | 0.23 | 1.06E-05 | 4.09E-05 |
| ENSG00000265933 | LINC00668 | -0.89 | 0.23 | 1.31E-05 | 5.02E-05 |
| ENSG00000132274 | TRIM22 | -0.89 | 0.23 | 1.40E-05 | 5.33E-05 |
| ENSG00000134516 | DOCK2 | -0.89 | 0.25 | 3.19E-05 | 1.17E-04 |
| ENSG00000273343 | | -0.89 | 0.33 | 6.73E-04 | 2.10E-03 |
| ENSG00000134323 | MYCN | -0.89 | 0.42 | 2.83E-03 | 7.97E-03 |
| ENSG00000186628 | FSD2 | -0.89 | 0.45 | 4.23E-03 | 1.15E-02 |
| ENSG00000134215 | VAV3 | -0.89 | 0.46 | 4.32E-03 | 1.17E-02 |
| ENSG00000146955 | RAB19 | -0.89 | 0.49 | 5.46E-03 | 1.46E-02 |
| ENSG00000230637 | | -0.89 | 0.75 | 7.37E-03 | 1.92E-02 |
| ENSG00000175294 | CATSPER1 | -0.89 | 0.57 | 7.83E-03 | 2.03E-02 |
| ENSG00000272154 | | -0.89 | 0.59 | 8.77E-03 | 2.25E-02 |
| ENSG00000181355 | OFCC1 | -0.89 | 0.62 | 8.84E-03 | 2.26E-02 |
| ENSG00000179520 | SLC17A8 | -0.89 | 0.61 | 8.95E-03 | 2.29E-02 |
| ENSG00000141314 | RHBDL3 | -0.89 | 0.62 | 8.95E-03 | 2.29E-02 |
| ENSG00000110906 | KCTD10 | -0.88 | 0.04 | 2.14E-126 | 1.70E-124 |
| ENSG00000113719 | ERGIC1 | -0.88 | 0.04 | 7.74E-102 | 4.57E-100 |
| ENSG00000002549 | LAP3 | -0.88 | 0.05 | 1.07E-61 | 3.08E-60 |
| ENSG00000152242 | C18orf25 | -0.88 | 0.05 | 1.50E-58 | 4.06E-57 |
| ENSG00000131788 | PIAS3 | -0.88 | 0.06 | 1.90E-45 | 3.85E-44 |
| ENSG00000178921 | PFAS | -0.88 | 0.07 | 1.22E-40 | 2.23E-39 |
| ENSG00000185480 | PARPBP | -0.88 | 0.07 | 8.80E-37 | 1.46E-35 |
| ENSG00000105738 | SIPA1L3 | -0.88 | 0.07 | 2.93E-33 | 4.35E-32 |
| ENSG00000137502 | RAB30 | -0.88 | 0.08 | 2.35E-31 | 3.31E-30 |
| ENSG00000180071 | ANKRD18A | -0.88 | 0.1 | 3.57E-19 | 3.20E-18 |

| | | | | | |
|---|---|---|---|---|---|
| ENSG00000189362 | NEMP2 | -0.88 | 0.11 | 8.40E-16 | 6.44E-15 |
| ENSG00000165795 | NDRG2 | -0.88 | 0.12 | 1.50E-14 | 1.09E-13 |
| ENSG00000223508 | RPL23AP53 | -0.88 | 0.12 | 6.53E-14 | 4.58E-13 |
| ENSG00000197959 | DNM3 | -0.88 | 0.13 | 1.99E-12 | 1.28E-11 |
| ENSG00000197859 | ADAMTSL2 | -0.88 | 0.13 | 5.43E-12 | 3.40E-11 |
| ENSG00000061918 | GUCY1B1 | -0.88 | 0.14 | 1.16E-11 | 7.15E-11 |
| ENSG00000103241 | FOXF1 | -0.88 | 0.14 | 1.30E-10 | 7.48E-10 |
| ENSG00000138336 | TET1 | -0.88 | 0.16 | 4.47E-09 | 2.29E-08 |
| ENSG00000126561 | STAT5A | -0.88 | 0.17 | 2.38E-08 | 1.16E-07 |
| ENSG00000253802 | SIRLNT | -0.88 | 0.18 | 1.23E-07 | 5.67E-07 |
| ENSG00000103044 | HAS3 | -0.88 | 0.22 | 6.76E-06 | 2.66E-05 |
| ENSG00000179046 | TRIML2 | -0.88 | 0.24 | 3.27E-05 | 1.20E-04 |
| ENSG00000123095 | BHLHE41 | -0.88 | 0.28 | 1.82E-04 | 6.13E-04 |
| ENSG00000291212 | | -0.88 | 0.31 | 4.19E-04 | 1.35E-03 |
| ENSG00000261051 | | -0.88 | 0.4 | 2.61E-03 | 7.42E-03 |
| ENSG00000287064 | | -0.88 | 0.42 | 3.24E-03 | 9.04E-03 |
| ENSG00000215853 | RPTN | -0.88 | 0.81 | 5.12E-03 | 1.38E-02 |
| ENSG00000227203 | SUB1P1 | -0.88 | 0.79 | 6.16E-03 | 1.63E-02 |
| ENSG00000264964 | TWSG1-DT | -0.88 | 0.53 | 7.40E-03 | 1.93E-02 |
| ENSG00000162620 | LRRIQ3 | -0.88 | 0.75 | 7.93E-03 | 2.05E-02 |
| ENSG00000280916 | FOXCUT | -0.88 | 0.55 | 7.95E-03 | 2.06E-02 |
| ENSG00000279822 | | -0.88 | 0.74 | 8.39E-03 | 2.16E-02 |
| ENSG00000184828 | ZBTB7C | -0.88 | 0.68 | 9.59E-03 | 2.43E-02 |
| ENSG00000003393 | ALS2 | -0.87 | 0.04 | 7.05E-91 | 3.51E-89 |
| ENSG00000169057 | MECP2 | -0.87 | 0.04 | 4.92E-88 | 2.33E-86 |
| ENSG00000023902 | PLEKHO1 | -0.87 | 0.05 | 8.88E-71 | 3.11E-69 |
| ENSG00000120802 | TMPO | -0.87 | 0.05 | 2.23E-69 | 7.55E-68 |
| ENSG00000110400 | NECTIN1 | -0.87 | 0.05 | 9.02E-59 | 2.47E-57 |
| ENSG00000107263 | RAPGEF1 | -0.87 | 0.06 | 2.91E-55 | 7.45E-54 |
| ENSG00000197563 | PIGN | -0.87 | 0.06 | 6.99E-43 | 1.36E-41 |
| ENSG00000107104 | KANK1 | -0.87 | 0.08 | 3.32E-27 | 4.05E-26 |
| ENSG00000166825 | ANPEP | -0.87 | 0.09 | 1.18E-25 | 1.36E-24 |
| ENSG00000139304 | PTPRQ | -0.87 | 0.09 | 1.86E-25 | 2.13E-24 |
| ENSG00000198960 | ARMCX6 | -0.87 | 0.09 | 4.02E-25 | 4.55E-24 |
| ENSG00000103876 | FAH | -0.87 | 0.09 | 1.30E-24 | 1.45E-23 |
| ENSG00000139567 | ACVRL1 | -0.87 | 0.09 | 6.45E-23 | 6.74E-22 |
| ENSG00000176055 | MBLAC2 | -0.87 | 0.09 | 8.15E-23 | 8.46E-22 |
| ENSG00000165060 | FXN | -0.87 | 0.1 | 1.74E-18 | 1.51E-17 |
| ENSG00000186300 | ZNF555 | -0.87 | 0.11 | 2.56E-17 | 2.12E-16 |
| ENSG00000171608 | PIK3CD | -0.87 | 0.13 | 7.47E-13 | 4.94E-12 |
| ENSG00000123219 | CENPK | -0.87 | 0.13 | 7.84E-13 | 5.18E-12 |
| ENSG00000131094 | C1QL1 | -0.87 | 0.14 | 1.15E-11 | 7.06E-11 |
| ENSG00000133739 | LRRCC1 | -0.87 | 0.15 | 2.72E-10 | 1.53E-09 |
| ENSG00000131470 | PSMC3IP | -0.87 | 0.15 | 6.95E-10 | 3.80E-09 |
| ENSG00000125089 | SH3TC1 | -0.87 | 0.15 | 1.35E-09 | 7.22E-09 |
| ENSG00000058085 | LAMC2 | -0.87 | 0.17 | 7.47E-08 | 3.50E-07 |
| ENSG00000259994 | | -0.87 | 0.18 | 2.54E-07 | 1.14E-06 |
| ENSG00000226180 | | -0.87 | 0.2 | 1.51E-06 | 6.30E-06 |
| ENSG00000135299 | ANKRD6 | -0.87 | 0.21 | 2.53E-06 | 1.04E-05 |
| ENSG00000166821 | PEX11A | -0.87 | 0.21 | 2.63E-06 | 1.07E-05 |

| | | | | | |
|---|---|---|---|---|---|
| ENSG00000138615 | CILP | -0.87 | 0.21 | 2.77E-06 | 1.13E-05 |
| ENSG00000006016 | CRLF1 | -0.87 | 0.21 | 5.58E-06 | 2.21E-05 |
| ENSG00000006611 | USH1C | -0.87 | 0.23 | 1.38E-05 | 5.25E-05 |
| ENSG00000214960 | CRPPA | -0.87 | 0.31 | 5.73E-04 | 1.81E-03 |
| ENSG00000162643 | DNAI3 | -0.87 | 0.32 | 6.79E-04 | 2.12E-03 |
| ENSG00000259341 | LINC03080 | -0.87 | 0.49 | 6.29E-03 | 1.66E-02 |
| ENSG00000290888 | | -0.87 | 0.54 | 8.21E-03 | 2.11E-02 |
| ENSG00000287766 | | -0.87 | 0.59 | 9.90E-03 | 2.50E-02 |
| ENSG00000140545 | MFGE8 | -0.86 | 0.05 | 1.34E-64 | 4.14E-63 |
| ENSG00000138829 | FBN2 | -0.86 | 0.05 | 2.22E-63 | 6.61E-62 |
| ENSG00000153815 | CMIP | -0.86 | 0.06 | 7.89E-51 | 1.81E-49 |
| ENSG00000198265 | HELZ | -0.86 | 0.06 | 1.08E-49 | 2.41E-48 |
| ENSG00000072518 | MARK2 | -0.86 | 0.06 | 1.97E-47 | 4.19E-46 |
| ENSG00000221963 | APOL6 | -0.86 | 0.07 | 3.44E-37 | 5.74E-36 |
| ENSG00000174891 | RSRC1 | -0.86 | 0.07 | 2.83E-32 | 4.09E-31 |
| ENSG00000105559 | PLEKHA4 | -0.86 | 0.08 | 7.45E-29 | 9.62E-28 |
| ENSG00000164237 | CMBL | -0.86 | 0.08 | 1.65E-25 | 1.90E-24 |
| ENSG00000164494 | PDSS2 | -0.86 | 0.09 | 1.78E-24 | 1.97E-23 |
| ENSG00000146802 | TMEM168 | -0.86 | 0.09 | 2.47E-23 | 2.62E-22 |
| ENSG00000156535 | CD109 | -0.86 | 0.09 | 9.67E-23 | 1.00E-21 |
| ENSG00000099869 | IGF2-AS | -0.86 | 0.09 | 1.55E-21 | 1.54E-20 |
| ENSG00000140853 | NLRC5 | -0.86 | 0.09 | 8.83E-21 | 8.47E-20 |
| ENSG00000180914 | OXTR | -0.86 | 0.12 | 3.92E-14 | 2.79E-13 |
| ENSG00000196659 | TTC30B | -0.86 | 0.12 | 1.52E-13 | 1.05E-12 |
| ENSG00000198932 | GPRASP1 | -0.86 | 0.12 | 1.70E-13 | 1.17E-12 |
| ENSG00000132334 | PTPRE | -0.86 | 0.14 | 2.59E-10 | 1.46E-09 |
| ENSG00000145002 | FAM86B2 | -0.86 | 0.21 | 7.45E-06 | 2.92E-05 |
| ENSG00000144481 | TRPM8 | -0.86 | 0.22 | 1.25E-05 | 4.80E-05 |
| ENSG00000135363 | LMO2 | -0.86 | 0.23 | 2.46E-05 | 9.12E-05 |
| ENSG00000178038 | ALS2CL | -0.86 | 0.24 | 4.78E-05 | 1.72E-04 |
| ENSG00000187796 | CARD9 | -0.86 | 0.25 | 5.99E-05 | 2.14E-04 |
| ENSG00000289161 | | -0.86 | 0.27 | 1.24E-04 | 4.27E-04 |
| ENSG00000235092 | ID2-AS1 | -0.86 | 0.29 | 2.72E-04 | 8.98E-04 |
| ENSG00000260007 | | -0.86 | 0.3 | 3.73E-04 | 1.21E-03 |
| ENSG00000268686 | | -0.86 | 0.32 | 6.73E-04 | 2.10E-03 |
| ENSG00000227954 | TARID | -0.86 | 0.37 | 2.01E-03 | 5.81E-03 |
| ENSG00000250365 | HEATR5A-DT | -0.86 | 0.4 | 3.02E-03 | 8.48E-03 |
| ENSG00000184486 | POU3F2 | -0.86 | 0.5 | 6.92E-03 | 1.81E-02 |
| ENSG00000010282 | HHATL | -0.86 | 0.51 | 7.88E-03 | 2.04E-02 |
| ENSG00000259004 | LINC02285 | -0.86 | 0.73 | 9.18E-03 | 2.34E-02 |
| ENSG00000291133 | | -0.86 | 0.68 | 1.04E-02 | 2.63E-02 |
| ENSG00000284526 | | -0.86 | 0.61 | 1.05E-02 | 2.65E-02 |
| ENSG00000132386 | SERPINF1 | -0.85 | 0.04 | 2.31E-98 | 1.30E-96 |
| ENSG00000100625 | SIX4 | -0.85 | 0.06 | 4.06E-54 | 1.02E-52 |
| ENSG00000270647 | TAF15 | -0.85 | 0.06 | 2.24E-52 | 5.35E-51 |
| ENSG00000112578 | BYSL | -0.85 | 0.06 | 2.35E-43 | 4.59E-42 |
| ENSG00000163945 | UVSSA | -0.85 | 0.06 | 2.50E-42 | 4.78E-41 |
| ENSG00000070882 | OSBPL3 | -0.85 | 0.06 | 2.81E-42 | 5.37E-41 |
| ENSG00000140367 | UBE2Q2 | -0.85 | 0.07 | 4.15E-38 | 7.13E-37 |
| ENSG00000164649 | CDCA7L | -0.85 | 0.07 | 5.11E-38 | 8.75E-37 |

| | | | | | |
|---|---|---|---|---|---|
| ENSG00000135070 | ISCA1 | -0.85 | 0.07 | 2.35E-36 | 3.83E-35 |
| ENSG00000197183 | NOL4L | -0.85 | 0.07 | 2.64E-33 | 3.91E-32 |
| ENSG00000117020 | AKT3 | -0.85 | 0.07 | 4.36E-32 | 6.25E-31 |
| ENSG00000133805 | AMPD3 | -0.85 | 0.07 | 1.03E-30 | 1.42E-29 |
| ENSG00000163947 | ARHGEF3 | -0.85 | 0.08 | 5.48E-30 | 7.36E-29 |
| ENSG00000119969 | HELLS | -0.85 | 0.08 | 6.11E-29 | 7.93E-28 |
| ENSG00000085449 | WDFY1 | -0.85 | 0.08 | 2.74E-28 | 3.46E-27 |
| ENSG00000172878 | METAP1D | -0.85 | 0.08 | 1.29E-25 | 1.49E-24 |
| ENSG00000171766 | GATM | -0.85 | 0.08 | 1.56E-25 | 1.79E-24 |
| ENSG00000259781 | HMGB1P6 | -0.85 | 0.08 | 4.44E-25 | 5.02E-24 |
| ENSG00000100294 | MCAT | -0.85 | 0.09 | 9.26E-23 | 9.60E-22 |
| ENSG00000156398 | SFXN2 | -0.85 | 0.09 | 3.54E-22 | 3.58E-21 |
| ENSG00000103522 | IL21R | -0.85 | 0.09 | 3.83E-22 | 3.88E-21 |
| ENSG00000269343 | ZNF587B | -0.85 | 0.1 | 1.36E-18 | 1.19E-17 |
| ENSG00000242498 | ARPIN | -0.85 | 0.12 | 3.65E-13 | 2.46E-12 |
| ENSG00000107537 | PHYH | -0.85 | 0.14 | 1.97E-10 | 1.12E-09 |
| ENSG00000257176 | | -0.85 | 0.15 | 6.55E-10 | 3.59E-09 |
| ENSG00000027075 | PRKCH | -0.85 | 0.15 | 1.04E-09 | 5.59E-09 |
| ENSG00000116852 | KIF21B | -0.85 | 0.15 | 4.02E-09 | 2.07E-08 |
| ENSG00000123685 | BATF3 | -0.85 | 0.17 | 4.12E-08 | 1.97E-07 |
| ENSG00000164236 | ANKRD33B | -0.85 | 0.23 | 3.38E-05 | 1.24E-04 |
| ENSG00000267787 | | -0.85 | 0.37 | 2.03E-03 | 5.87E-03 |
| ENSG00000178562 | CD28 | -0.85 | 0.37 | 2.39E-03 | 6.84E-03 |
| ENSG00000184809 | B3GALT5-AS1 | -0.85 | 0.38 | 2.56E-03 | 7.29E-03 |
| ENSG00000185052 | SLC24A3 | -0.85 | 0.38 | 2.66E-03 | 7.54E-03 |
| ENSG00000162746 | FCRLB | -0.85 | 0.38 | 2.66E-03 | 7.54E-03 |
| ENSG00000119919 | NKX2-3 | -0.85 | 0.41 | 3.89E-03 | 1.07E-02 |
| ENSG00000004838 | ZMYND10 | -0.85 | 0.44 | 5.50E-03 | 1.47E-02 |
| ENSG00000258623 | | -0.85 | 0.47 | 6.38E-03 | 1.68E-02 |
| ENSG00000227811 | INKA2-AS1 | -0.85 | 0.48 | 6.81E-03 | 1.79E-02 |
| ENSG00000253891 | | -0.85 | 0.48 | 6.84E-03 | 1.79E-02 |
| ENSG00000100385 | IL2RB | -0.85 | 0.51 | 7.98E-03 | 2.06E-02 |
| ENSG00000188937 | NYX | -0.85 | 0.75 | 8.89E-03 | 2.27E-02 |
| ENSG00000290044 | | -0.85 | 0.74 | 9.38E-03 | 2.39E-02 |
| ENSG00000149646 | CNBD2 | -0.85 | 0.54 | 9.73E-03 | 2.47E-02 |
| ENSG00000286953 | | -0.85 | 0.55 | 9.95E-03 | 2.51E-02 |
| ENSG00000116748 | AMPD1 | -0.85 | 0.58 | 1.08E-02 | 2.70E-02 |
| ENSG00000240376 | | -0.85 | 0.69 | 1.09E-02 | 2.72E-02 |
| ENSG00000273786 | | -0.85 | 0.7 | 1.09E-02 | 2.73E-02 |
| ENSG00000186575 | NF2 | -0.84 | 0.05 | 6.83E-73 | 2.49E-71 |
| ENSG00000145592 | RPL37 | -0.84 | 0.05 | 8.26E-66 | 2.61E-64 |
| ENSG00000140526 | ABHD2 | -0.84 | 0.05 | 1.03E-56 | 2.69E-55 |
| ENSG00000165660 | ABRAXAS2 | -0.84 | 0.05 | 1.57E-53 | 3.86E-52 |
| ENSG00000139044 | B4GALNT3 | -0.84 | 0.06 | 2.00E-53 | 4.91E-52 |
| ENSG00000140416 | TPM1 | -0.84 | 0.06 | 9.15E-48 | 1.95E-46 |
| ENSG00000123737 | EXOSC9 | -0.84 | 0.06 | 4.22E-40 | 7.56E-39 |
| ENSG00000164941 | INTS8 | -0.84 | 0.07 | 1.83E-38 | 3.19E-37 |
| ENSG00000148400 | NOTCH1 | -0.84 | 0.07 | 1.71E-35 | 2.73E-34 |
| ENSG00000115525 | ST3GAL5 | -0.84 | 0.07 | 3.50E-34 | 5.33E-33 |
| ENSG00000118655 | DCLRE1B | -0.84 | 0.07 | 1.28E-33 | 1.92E-32 |

| | | | | | |
|---|---|---|---|---|---|
| ENSG00000101911 | PRPS2 | -0.84 | 0.07 | 1.04E-30 | 1.44E-29 |
| ENSG00000168765 | GSTM4 | -0.84 | 0.08 | 3.68E-27 | 4.49E-26 |
| ENSG00000205352 | PRR13 | -0.84 | 0.09 | 2.87E-23 | 3.03E-22 |
| ENSG00000153044 | CENPH | -0.84 | 0.09 | 1.13E-20 | 1.08E-19 |
| ENSG00000185477 | GPRIN3 | -0.84 | 0.09 | 1.30E-20 | 1.24E-19 |
| ENSG00000118894 | EEF2KMT | -0.84 | 0.1 | 1.94E-17 | 1.61E-16 |
| ENSG00000030304 | MUSK | -0.84 | 0.11 | 4.88E-15 | 3.63E-14 |
| ENSG00000175768 | TOMM5 | -0.84 | 0.13 | 5.95E-11 | 3.50E-10 |
| ENSG00000175643 | RMI2 | -0.84 | 0.15 | 5.15E-09 | 2.63E-08 |
| ENSG00000248713 | C4orf54 | -0.84 | 0.25 | 9.46E-05 | 3.30E-04 |
| ENSG00000273055 | | -0.84 | 0.29 | 4.62E-04 | 1.48E-03 |
| ENSG00000104833 | TUBB4A | -0.84 | 0.35 | 1.55E-03 | 4.57E-03 |
| ENSG00000077009 | NMRK2 | -0.84 | 0.42 | 4.37E-03 | 1.19E-02 |
| ENSG00000273007 | | -0.84 | 0.48 | 7.84E-03 | 2.03E-02 |
| ENSG00000266718 | | -0.84 | 0.77 | 8.05E-03 | 2.08E-02 |
| ENSG00000122585 | NPY | -0.84 | 0.52 | 9.45E-03 | 2.40E-02 |
| ENSG00000184361 | SPATA32 | -0.84 | 0.6 | 1.20E-02 | 2.97E-02 |
| ENSG00000173762 | CD7 | -0.84 | 0.63 | 1.24E-02 | 3.06E-02 |
| ENSG00000150760 | DOCK1 | -0.83 | 0.04 | 1.61E-91 | 8.13E-90 |
| ENSG00000148677 | ANKRD1 | -0.83 | 0.04 | 1.66E-87 | 7.67E-86 |
| ENSG00000162704 | ARPC5 | -0.83 | 0.04 | 2.52E-81 | 1.06E-79 |
| ENSG00000215301 | DDX3X | -0.83 | 0.04 | 1.16E-76 | 4.55E-75 |
| ENSG00000135341 | MAP3K7 | -0.83 | 0.05 | 1.66E-64 | 5.12E-63 |
| ENSG00000166716 | ZNF592 | -0.83 | 0.05 | 4.61E-64 | 1.40E-62 |
| ENSG00000108344 | PSMD3 | -0.83 | 0.05 | 4.17E-59 | 1.15E-57 |
| ENSG00000168785 | TSPAN5 | -0.83 | 0.06 | 7.56E-51 | 1.73E-49 |
| ENSG00000164062 | APEH | -0.83 | 0.06 | 1.83E-50 | 4.15E-49 |
| ENSG00000162989 | KCNJ3 | -0.83 | 0.06 | 6.32E-49 | 1.39E-47 |
| ENSG00000143190 | POU2F1 | -0.83 | 0.08 | 5.70E-28 | 7.11E-27 |
| ENSG00000166311 | SMPD1 | -0.83 | 0.09 | 1.27E-19 | 1.16E-18 |
| ENSG00000119630 | PGF | -0.83 | 0.1 | 2.10E-18 | 1.81E-17 |
| ENSG00000122483 | CCDC18 | -0.83 | 0.1 | 1.45E-17 | 1.21E-16 |
| ENSG00000174500 | GCSAM | -0.83 | 0.11 | 6.14E-16 | 4.76E-15 |
| ENSG00000144395 | CCDC150 | -0.83 | 0.13 | 3.97E-11 | 2.36E-10 |
| ENSG00000172456 | FGGY | -0.83 | 0.17 | 2.04E-07 | 9.23E-07 |
| ENSG00000164308 | ERAP2 | -0.83 | 0.18 | 2.63E-07 | 1.18E-06 |
| ENSG00000289507 | | -0.83 | 0.19 | 7.88E-07 | 3.38E-06 |
| ENSG00000157368 | IL34 | -0.83 | 0.23 | 2.96E-05 | 1.09E-04 |
| ENSG00000159387 | IRX6 | -0.83 | 0.23 | 4.58E-05 | 1.65E-04 |
| ENSG00000261594 | TPBGL | -0.83 | 0.24 | 6.99E-05 | 2.48E-04 |
| ENSG00000103544 | VPS35L | -0.83 | 0.26 | 1.57E-04 | 5.34E-04 |
| ENSG00000056558 | TRAF1 | -0.83 | 0.26 | 1.85E-04 | 6.23E-04 |
| ENSG00000137878 | GCOM1 | -0.83 | 0.34 | 1.59E-03 | 4.69E-03 |
| ENSG00000262358 | | -0.83 | 0.84 | 3.97E-03 | 1.09E-02 |
| ENSG00000119714 | GPR68 | -0.83 | 0.4 | 3.97E-03 | 1.09E-02 |
| ENSG00000158125 | XDH | -0.83 | 0.83 | 4.03E-03 | 1.10E-02 |
| ENSG00000166546 | BEAN1 | -0.83 | 0.43 | 5.24E-03 | 1.40E-02 |
| ENSG00000250995 | TMEM30BP1 | -0.83 | 0.46 | 7.15E-03 | 1.87E-02 |
| ENSG00000145087 | STXBP5L | -0.83 | 0.47 | 7.79E-03 | 2.02E-02 |
| ENSG00000216921 | FAM240C | -0.83 | 0.49 | 8.59E-03 | 2.20E-02 |

| | | | | | |
|---|---|---|---|---|---|
| ENSG00000278570 | NR2E3 | -0.83 | 0.54 | 1.07E-02 | 2.69E-02 |
| ENSG00000163814 | CDCP1 | -0.83 | 0.54 | 1.07E-02 | 2.70E-02 |
| ENSG00000105523 | FAM83E | -0.83 | 0.54 | 1.11E-02 | 2.77E-02 |
| ENSG00000272293 | | -0.83 | 0.59 | 1.24E-02 | 3.07E-02 |
| ENSG00000160714 | UBE2Q1 | -0.82 | 0.04 | 8.22E-77 | 3.25E-75 |
| ENSG00000115232 | ITGA4 | -0.82 | 0.05 | 8.25E-71 | 2.90E-69 |
| ENSG00000142207 | URB1 | -0.82 | 0.05 | 3.85E-59 | 1.07E-57 |
| ENSG00000103051 | COG4 | -0.82 | 0.05 | 1.51E-58 | 4.10E-57 |
| ENSG00000119403 | PHF19 | -0.82 | 0.05 | 1.62E-57 | 4.32E-56 |
| ENSG00000248905 | FMN1 | -0.82 | 0.05 | 1.25E-51 | 2.92E-50 |
| ENSG00000163507 | CIP2A | -0.82 | 0.06 | 1.60E-49 | 3.54E-48 |
| ENSG00000135953 | MFSD9 | -0.82 | 0.06 | 1.86E-44 | 3.71E-43 |
| ENSG00000145220 | LYAR | -0.82 | 0.06 | 9.66E-44 | 1.91E-42 |
| ENSG00000101811 | CSTF2 | -0.82 | 0.06 | 7.47E-41 | 1.37E-39 |
| ENSG00000136824 | SMC2 | -0.82 | 0.06 | 1.20E-40 | 2.19E-39 |
| ENSG00000115935 | WIPF1 | -0.82 | 0.06 | 2.03E-38 | 3.53E-37 |
| ENSG00000135316 | SYNCRIP | -0.82 | 0.06 | 3.60E-37 | 6.00E-36 |
| ENSG00000124615 | MOCS1 | -0.82 | 0.07 | 4.21E-34 | 6.40E-33 |
| ENSG00000144306 | SCRN3 | -0.82 | 0.07 | 4.70E-32 | 6.75E-31 |
| ENSG00000196214 | ZNF766 | -0.82 | 0.08 | 1.63E-26 | 1.94E-25 |
| ENSG00000167384 | ZNF180 | -0.82 | 0.08 | 2.68E-26 | 3.16E-25 |
| ENSG00000016082 | ISL1 | -0.82 | 0.09 | 2.22E-21 | 2.18E-20 |
| ENSG00000151611 | MMAA | -0.82 | 0.09 | 7.79E-21 | 7.49E-20 |
| ENSG00000111110 | PPM1H | -0.82 | 0.09 | 3.44E-20 | 3.23E-19 |
| ENSG00000103227 | LMF1 | -0.82 | 0.09 | 8.04E-20 | 7.42E-19 |
| ENSG00000196172 | ZNF681 | -0.82 | 0.1 | 1.23E-16 | 9.88E-16 |
| ENSG00000280323 | | -0.82 | 0.11 | 8.04E-14 | 5.62E-13 |
| ENSG00000165633 | VSTM4 | -0.82 | 0.11 | 1.07E-13 | 7.46E-13 |
| ENSG00000182923 | CEP63 | -0.82 | 0.13 | 1.34E-11 | 8.19E-11 |
| ENSG00000026297 | RNASET2 | -0.82 | 0.15 | 3.57E-09 | 1.85E-08 |
| ENSG00000125414 | MYH2 | -0.82 | 0.17 | 9.62E-08 | 4.47E-07 |
| ENSG00000163577 | EIF5A2 | -0.82 | 0.17 | 1.00E-07 | 4.65E-07 |
| ENSG00000198331 | HYLS1 | -0.82 | 0.19 | 1.39E-06 | 5.84E-06 |
| ENSG00000115648 | MLPH | -0.82 | 0.21 | 1.06E-05 | 4.08E-05 |
| ENSG00000138400 | MDH1B | -0.82 | 0.23 | 4.05E-05 | 1.47E-04 |
| ENSG00000128039 | SRD5A3 | -0.82 | 0.24 | 6.75E-05 | 2.40E-04 |
| ENSG00000169435 | RASSF6 | -0.82 | 0.29 | 5.21E-04 | 1.65E-03 |
| ENSG00000272263 | | -0.82 | 0.3 | 6.34E-04 | 1.99E-03 |
| ENSG00000168907 | PLA2G4F | -0.82 | 0.41 | 4.74E-03 | 1.28E-02 |
| ENSG00000274220 | | -0.82 | 0.42 | 5.62E-03 | 1.50E-02 |
| ENSG00000286520 | | -0.82 | 0.79 | 6.80E-03 | 1.78E-02 |
| ENSG00000259792 | | -0.82 | 0.55 | 1.22E-02 | 3.01E-02 |
| ENSG00000182253 | SYNM | -0.81 | 0.05 | 1.77E-59 | 4.94E-58 |
| ENSG00000087302 | RTRAF | -0.81 | 0.05 | 1.74E-58 | 4.73E-57 |
| ENSG00000146858 | ZC3HAV1L | -0.81 | 0.05 | 4.45E-57 | 1.17E-55 |
| ENSG00000106086 | PLEKHA8 | -0.81 | 0.05 | 5.18E-51 | 1.19E-49 |
| ENSG00000076248 | UNG | -0.81 | 0.06 | 1.73E-49 | 3.82E-48 |
| ENSG00000028137 | TNFRSF1B | -0.81 | 0.06 | 3.28E-49 | 7.22E-48 |
| ENSG00000110871 | COQ5 | -0.81 | 0.06 | 2.87E-46 | 5.93E-45 |
| ENSG00000165821 | SALL2 | -0.81 | 0.06 | 7.71E-38 | 1.32E-36 |

| | | | | | |
|---|---|---|---|---|---|
| ENSG00000163584 | RPL22L1 | -0.81 | 0.07 | 1.94E-33 | 2.91E-32 |
| ENSG00000196839 | ADA | -0.81 | 0.07 | 2.14E-33 | 3.19E-32 |
| ENSG00000116096 | SPR | -0.81 | 0.08 | 2.99E-27 | 3.66E-26 |
| ENSG00000107371 | EXOSC3 | -0.81 | 0.08 | 2.38E-23 | 2.52E-22 |
| ENSG00000230630 | DNM3OS | -0.81 | 0.08 | 1.99E-22 | 2.03E-21 |
| ENSG00000117305 | HMGCL | -0.81 | 0.09 | 6.94E-22 | 6.95E-21 |
| ENSG00000182901 | RGS7 | -0.81 | 0.09 | 1.69E-19 | 1.54E-18 |
| ENSG00000144749 | LRIG1 | -0.81 | 0.1 | 9.21E-16 | 7.05E-15 |
| ENSG00000162076 | FLYWCH2 | -0.81 | 0.11 | 9.55E-15 | 7.00E-14 |
| ENSG00000143891 | GALM | -0.81 | 0.11 | 1.77E-14 | 1.28E-13 |
| ENSG00000165891 | E2F7 | -0.81 | 0.12 | 5.28E-13 | 3.52E-12 |
| ENSG00000141750 | STAC2 | -0.81 | 0.13 | 2.09E-11 | 1.27E-10 |
| ENSG00000185340 | GAS2L1 | -0.81 | 0.14 | 5.61E-10 | 3.09E-09 |
| ENSG00000049769 | PPP1R3F | -0.81 | 0.14 | 1.91E-09 | 1.01E-08 |
| ENSG00000183850 | ZNF730 | -0.81 | 0.18 | 1.03E-06 | 4.36E-06 |
| ENSG00000279605 | | -0.81 | 0.2 | 7.45E-06 | 2.92E-05 |
| ENSG00000181408 | UTS2R | -0.81 | 0.21 | 1.00E-05 | 3.88E-05 |
| ENSG00000139160 | ETFBKMT | -0.81 | 0.22 | 3.39E-05 | 1.24E-04 |
| ENSG00000287666 | | -0.81 | 0.26 | 2.38E-04 | 7.90E-04 |
| ENSG00000196460 | RFX8 | -0.81 | 0.27 | 3.13E-04 | 1.02E-03 |
| ENSG00000232053 | | -0.81 | 0.41 | 5.38E-03 | 1.44E-02 |
| ENSG00000166664 | CHRFAM7A | -0.81 | 0.47 | 9.06E-03 | 2.31E-02 |
| ENSG00000135378 | PRRG4 | -0.81 | 0.5 | 1.07E-02 | 2.68E-02 |
| ENSG00000184908 | CLCNKB | -0.81 | 0.73 | 1.18E-02 | 2.94E-02 |
| ENSG00000280744 | LINC01173 | -0.81 | 0.54 | 1.22E-02 | 3.01E-02 |
| ENSG00000241644 | INMT | -0.81 | 0.7 | 1.27E-02 | 3.14E-02 |
| ENSG00000276292 | | -0.81 | 0.6 | 1.39E-02 | 3.40E-02 |
| ENSG00000279569 | | -0.81 | 0.62 | 1.41E-02 | 3.43E-02 |
| ENSG00000232528 | | -0.81 | 0.64 | 1.45E-02 | 3.52E-02 |
| ENSG00000239494 | RN7SL333P | -0.81 | 0.61 | 1.46E-02 | 3.55E-02 |
| ENSG00000169604 | ANTXR1 | -0.8 | 0.04 | 5.54E-113 | 3.73E-111 |
| ENSG00000128928 | IVD | -0.8 | 0.05 | 3.10E-69 | 1.05E-67 |
| ENSG00000009709 | PAX7 | -0.8 | 0.05 | 1.63E-64 | 5.04E-63 |
| ENSG00000134153 | EMC7 | -0.8 | 0.05 | 4.89E-64 | 1.48E-62 |
| ENSG00000168090 | COPS6 | -0.8 | 0.05 | 3.16E-63 | 9.41E-62 |
| ENSG00000109685 | NSD2 | -0.8 | 0.05 | 1.25E-58 | 3.41E-57 |
| ENSG00000186815 | TPCN1 | -0.8 | 0.05 | 1.20E-57 | 3.20E-56 |
| ENSG00000110987 | BCL7A | -0.8 | 0.05 | 2.93E-51 | 6.79E-50 |
| ENSG00000152601 | MBNL1 | -0.8 | 0.06 | 8.20E-49 | 1.79E-47 |
| ENSG00000197747 | S100A10 | -0.8 | 0.06 | 1.34E-46 | 2.80E-45 |
| ENSG00000205208 | C4orf46 | -0.8 | 0.06 | 2.72E-38 | 4.70E-37 |
| ENSG00000151612 | ZNF827 | -0.8 | 0.06 | 1.45E-37 | 2.44E-36 |
| ENSG00000168734 | PKIG | -0.8 | 0.07 | 3.02E-30 | 4.09E-29 |
| ENSG00000140365 | COMMD4 | -0.8 | 0.07 | 1.67E-29 | 2.20E-28 |
| ENSG00000137413 | TAF8 | -0.8 | 0.07 | 1.60E-28 | 2.04E-27 |
| ENSG00000178409 | BEND3 | -0.8 | 0.07 | 1.01E-27 | 1.26E-26 |
| ENSG00000095303 | PTGS1 | -0.8 | 0.08 | 6.85E-24 | 7.44E-23 |
| ENSG00000084072 | PPIE | -0.8 | 0.09 | 6.22E-21 | 6.02E-20 |
| ENSG00000291065 | | -0.8 | 0.09 | 4.33E-19 | 3.86E-18 |
| ENSG00000185869 | ZNF829 | -0.8 | 0.1 | 2.10E-16 | 1.66E-15 |

| | | | | | |
|---|---|---|---|---|---|
| ENSG00000065675 | PRKCQ | -0.8 | 0.11 | 1.10E-13 | 7.64E-13 |
| ENSG00000104154 | SLC30A4 | -0.8 | 0.15 | 6.41E-09 | 3.25E-08 |
| ENSG00000286177 | | -0.8 | 0.18 | 2.05E-06 | 8.47E-06 |
| ENSG00000163995 | ABLIM2 | -0.8 | 0.23 | 7.34E-05 | 2.59E-04 |
| ENSG00000261713 | SSTR5-AS1 | -0.8 | 0.3 | 1.03E-03 | 3.13E-03 |
| ENSG00000235033 | DAAM2-AS1 | -0.8 | 0.32 | 1.69E-03 | 4.94E-03 |
| ENSG00000152689 | RASGRP3 | -0.8 | 0.33 | 1.86E-03 | 5.41E-03 |
| ENSG00000237760 | | -0.8 | 0.83 | 4.66E-03 | 1.26E-02 |
| ENSG00000269416 | LINC01224 | -0.8 | 0.42 | 6.39E-03 | 1.68E-02 |
| ENSG00000286867 | | -0.8 | 0.8 | 6.62E-03 | 1.74E-02 |
| ENSG00000143195 | ILDR2 | -0.8 | 0.43 | 6.99E-03 | 1.83E-02 |
| ENSG00000090659 | CD209 | -0.8 | 0.45 | 8.79E-03 | 2.25E-02 |
| ENSG00000279289 | | -0.8 | 0.76 | 9.73E-03 | 2.47E-02 |
| ENSG00000280800 | | -0.8 | 0.51 | 1.17E-02 | 2.92E-02 |
| ENSG00000250107 | CACNA1G-AS1 | -0.8 | 0.54 | 1.37E-02 | 3.35E-02 |
| ENSG00000232453 | LINC02777 | -0.8 | 0.69 | 1.42E-02 | 3.46E-02 |
| ENSG00000273724 | | -0.8 | 0.58 | 1.44E-02 | 3.51E-02 |
| ENSG00000174171 | | -0.8 | 0.64 | 1.49E-02 | 3.62E-02 |
| ENSG00000078269 | SYNJ2 | -0.79 | 0.04 | 2.05E-103 | 1.24E-101 |
| ENSG00000145555 | MYO10 | -0.79 | 0.04 | 1.11E-87 | 5.14E-86 |
| ENSG00000143061 | IGSF3 | -0.79 | 0.04 | 1.34E-76 | 5.24E-75 |
| ENSG00000082516 | GEMIN5 | -0.79 | 0.04 | 9.62E-75 | 3.62E-73 |
| ENSG00000141367 | CLTC | -0.79 | 0.04 | 4.69E-73 | 1.72E-71 |
| ENSG00000136153 | LMO7 | -0.79 | 0.04 | 1.18E-72 | 4.23E-71 |
| ENSG00000113552 | GNPDA1 | -0.79 | 0.05 | 3.17E-64 | 9.72E-63 |
| ENSG00000073921 | PICALM | -0.79 | 0.05 | 2.34E-60 | 6.63E-59 |
| ENSG00000119638 | NEK9 | -0.79 | 0.05 | 7.20E-58 | 1.93E-56 |
| ENSG00000152556 | PFKM | -0.79 | 0.05 | 2.86E-54 | 7.20E-53 |
| ENSG00000172164 | SNTB1 | -0.79 | 0.05 | 1.74E-53 | 4.28E-52 |
| ENSG00000081087 | OSTM1 | -0.79 | 0.05 | 7.04E-53 | 1.70E-51 |
| ENSG00000160584 | SIK3 | -0.79 | 0.05 | 5.93E-51 | 1.36E-49 |
| ENSG00000141101 | NOB1 | -0.79 | 0.06 | 2.79E-47 | 5.89E-46 |
| ENSG00000029993 | HMGB3 | -0.79 | 0.06 | 4.90E-45 | 9.87E-44 |
| ENSG00000197381 | ADARB1 | -0.79 | 0.06 | 2.16E-42 | 4.13E-41 |
| ENSG00000137337 | MDC1 | -0.79 | 0.06 | 2.39E-38 | 4.13E-37 |
| ENSG00000139233 | LLPH | -0.79 | 0.07 | 1.01E-33 | 1.53E-32 |
| ENSG00000138796 | HADH | -0.79 | 0.07 | 6.77E-33 | 9.92E-32 |
| ENSG00000213390 | ARHGAP19 | -0.79 | 0.07 | 2.73E-27 | 3.35E-26 |
| ENSG00000105088 | OLFM2 | -0.79 | 0.08 | 1.75E-23 | 1.87E-22 |
| ENSG00000165084 | C8orf34 | -0.79 | 0.08 | 2.34E-23 | 2.49E-22 |
| ENSG00000167747 | C19orf48P | -0.79 | 0.08 | 1.58E-22 | 1.62E-21 |
| ENSG00000106772 | PRUNE2 | -0.79 | 0.09 | 5.04E-21 | 4.90E-20 |
| ENSG00000100065 | CARD10 | -0.79 | 0.09 | 1.71E-18 | 1.48E-17 |
| ENSG00000132570 | PCBD2 | -0.79 | 0.09 | 1.93E-18 | 1.67E-17 |
| ENSG00000155893 | PXYLP1 | -0.79 | 0.1 | 1.46E-16 | 1.16E-15 |
| ENSG00000197620 | EOLA1 | -0.79 | 0.1 | 4.00E-16 | 3.13E-15 |
| ENSG00000165501 | LRR1 | -0.79 | 0.1 | 2.32E-15 | 1.75E-14 |
| ENSG00000213793 | ZNF888 | -0.79 | 0.1 | 6.39E-15 | 4.73E-14 |
| ENSG00000123411 | IKZF4 | -0.79 | 0.11 | 7.57E-15 | 5.58E-14 |
| ENSG00000160957 | RECQL4 | -0.79 | 0.11 | 1.61E-14 | 1.16E-13 |

| | | | | | |
|---|---|---|---|---|---|
| ENSG00000187676 | B3GLCT | -0.79 | 0.14 | 1.56E-09 | 8.28E-09 |
| ENSG00000166265 | CYYR1 | -0.79 | 0.15 | 1.23E-08 | 6.10E-08 |
| ENSG00000129993 | CBFA2T3 | -0.79 | 0.16 | 4.76E-08 | 2.26E-07 |
| ENSG00000268087 | | -0.79 | 0.16 | 7.73E-08 | 3.62E-07 |
| ENSG00000137198 | GMPR | -0.79 | 0.16 | 9.96E-08 | 4.63E-07 |
| ENSG00000275074 | NUDT18 | -0.79 | 0.17 | 4.67E-07 | 2.05E-06 |
| ENSG00000165300 | SLITRK5 | -0.79 | 0.17 | 5.49E-07 | 2.40E-06 |
| ENSG00000175395 | ZNF25 | -0.79 | 0.18 | 1.18E-06 | 4.96E-06 |
| ENSG00000120324 | PCDHB10 | -0.79 | 0.19 | 3.50E-06 | 1.42E-05 |
| ENSG00000074276 | CDHR2 | -0.79 | 0.24 | 1.15E-04 | 3.97E-04 |
| ENSG00000287917 | | -0.79 | 0.24 | 1.54E-04 | 5.24E-04 |
| ENSG00000146038 | DCDC2 | -0.79 | 0.3 | 9.77E-04 | 2.98E-03 |
| ENSG00000267940 | | -0.79 | 0.35 | 2.71E-03 | 7.67E-03 |
| ENSG00000185483 | ROR1 | -0.79 | 0.35 | 3.24E-03 | 9.03E-03 |
| ENSG00000286535 | | -0.79 | 0.37 | 4.19E-03 | 1.14E-02 |
| ENSG00000224272 | | -0.79 | 0.38 | 4.62E-03 | 1.25E-02 |
| ENSG00000227158 | | -0.79 | 0.82 | 5.42E-03 | 1.45E-02 |
| ENSG00000166863 | TAC3 | -0.79 | 0.42 | 7.19E-03 | 1.88E-02 |
| ENSG00000184730 | APOBR | -0.79 | 0.78 | 8.50E-03 | 2.18E-02 |
| ENSG00000243746 | EEF1A1P10 | -0.79 | 0.49 | 1.19E-02 | 2.96E-02 |
| ENSG00000158748 | HTR6 | -0.79 | 0.58 | 1.51E-02 | 3.67E-02 |
| ENSG00000267737 | | -0.79 | 0.67 | 1.58E-02 | 3.81E-02 |
| ENSG00000091436 | MAP3K20 | -0.78 | 0.05 | 3.28E-63 | 9.75E-62 |
| ENSG00000164080 | RAD54L2 | -0.78 | 0.05 | 9.19E-60 | 2.57E-58 |
| ENSG00000185630 | PBX1 | -0.78 | 0.05 | 4.75E-55 | 1.21E-53 |
| ENSG00000152684 | PELO | -0.78 | 0.05 | 6.44E-54 | 1.60E-52 |
| ENSG00000132669 | RIN2 | -0.78 | 0.05 | 2.65E-47 | 5.60E-46 |
| ENSG00000080573 | COL5A3 | -0.78 | 0.06 | 3.96E-46 | 8.16E-45 |
| ENSG00000129675 | ARHGEF6 | -0.78 | 0.06 | 1.34E-45 | 2.73E-44 |
| ENSG00000107719 | PALD1 | -0.78 | 0.06 | 2.91E-43 | 5.68E-42 |
| ENSG00000156853 | ZNF689 | -0.78 | 0.06 | 7.76E-38 | 1.32E-36 |
| ENSG00000243156 | MICAL3 | -0.78 | 0.07 | 1.67E-32 | 2.42E-31 |
| ENSG00000198464 | ZNF480 | -0.78 | 0.07 | 7.97E-29 | 1.03E-27 |
| ENSG00000116198 | CEP104 | -0.78 | 0.07 | 3.41E-27 | 4.16E-26 |
| ENSG00000061337 | LZTS1 | -0.78 | 0.08 | 2.47E-25 | 2.82E-24 |
| ENSG00000108771 | DHX58 | -0.78 | 0.08 | 6.44E-24 | 7.00E-23 |
| ENSG00000272031 | ANKRD34A | -0.78 | 0.09 | 1.78E-18 | 1.54E-17 |
| ENSG00000163322 | ABRAXAS1 | -0.78 | 0.1 | 8.89E-17 | 7.18E-16 |
| ENSG00000146143 | PRIM2 | -0.78 | 0.1 | 9.70E-17 | 7.81E-16 |
| ENSG00000121691 | CAT | -0.78 | 0.1 | 4.08E-16 | 3.19E-15 |
| ENSG00000124279 | FASTKD3 | -0.78 | 0.1 | 9.00E-16 | 6.90E-15 |
| ENSG00000291102 | GUSBP1 | -0.78 | 0.1 | 1.27E-14 | 9.24E-14 |
| ENSG00000254093 | PINX1 | -0.78 | 0.11 | 4.09E-14 | 2.91E-13 |
| ENSG00000198959 | TGM2 | -0.78 | 0.12 | 3.52E-11 | 2.10E-10 |
| ENSG00000154511 | DIPK1A | -0.78 | 0.13 | 2.35E-10 | 1.33E-09 |
| ENSG00000111199 | TRPV4 | -0.78 | 0.13 | 3.66E-10 | 2.04E-09 |
| ENSG00000101265 | RASSF2 | -0.78 | 0.15 | 1.35E-08 | 6.67E-08 |
| ENSG00000100092 | SH3BP1 | -0.78 | 0.19 | 5.61E-06 | 2.22E-05 |
| ENSG00000285676 | | -0.78 | 0.19 | 6.61E-06 | 2.60E-05 |
| ENSG00000254290 | | -0.78 | 0.19 | 7.36E-06 | 2.88E-05 |

| | | | | | |
|---|---|---|---|---|---|
| ENSG00000267565 | | -0.78 | 0.25 | 2.38E-04 | 7.90E-04 |
| ENSG00000269397 | | -0.78 | 0.26 | 3.18E-04 | 1.04E-03 |
| ENSG00000287022 | | -0.78 | 0.82 | 5.48E-03 | 1.46E-02 |
| ENSG00000278626 | | -0.78 | 0.39 | 5.58E-03 | 1.49E-02 |
| ENSG00000278344 | | -0.78 | 0.39 | 5.80E-03 | 1.54E-02 |
| ENSG00000233538 | | -0.78 | 0.47 | 1.07E-02 | 2.70E-02 |
| ENSG00000232389 | | -0.78 | 0.74 | 1.20E-02 | 2.99E-02 |
| ENSG00000261273 | | -0.78 | 0.74 | 1.23E-02 | 3.03E-02 |
| ENSG00000250015 | | -0.78 | 0.51 | 1.31E-02 | 3.23E-02 |
| ENSG00000144152 | FBLN7 | -0.78 | 0.5 | 1.32E-02 | 3.25E-02 |
| ENSG00000078900 | TP73 | -0.78 | 0.53 | 1.44E-02 | 3.51E-02 |
| ENSG00000180644 | PRF1 | -0.78 | 0.56 | 1.63E-02 | 3.92E-02 |
| ENSG00000251689 | | -0.78 | 0.6 | 1.67E-02 | 4.00E-02 |
| ENSG00000160097 | FNDC5 | -0.77 | 0.04 | 2.03E-91 | 1.02E-89 |
| ENSG00000197852 | INKA2 | -0.77 | 0.05 | 2.56E-52 | 6.09E-51 |
| ENSG00000100418 | DESI1 | -0.77 | 0.05 | 2.11E-50 | 4.78E-49 |
| ENSG00000128510 | CPA4 | -0.77 | 0.05 | 2.09E-47 | 4.43E-46 |
| ENSG00000107819 | SFXN3 | -0.77 | 0.06 | 1.03E-43 | 2.03E-42 |
| ENSG00000109689 | STIM2 | -0.77 | 0.06 | 1.59E-40 | 2.90E-39 |
| ENSG00000117360 | PRPF3 | -0.77 | 0.06 | 6.75E-38 | 1.15E-36 |
| ENSG00000083123 | BCKDHB | -0.77 | 0.06 | 9.09E-38 | 1.55E-36 |
| ENSG00000132600 | PRMT7 | -0.77 | 0.06 | 1.60E-36 | 2.63E-35 |
| ENSG00000154743 | TSEN2 | -0.77 | 0.07 | 1.95E-29 | 2.58E-28 |
| ENSG00000166452 | AKIP1 | -0.77 | 0.07 | 1.36E-25 | 1.56E-24 |
| ENSG00000145996 | CDKAL1 | -0.77 | 0.09 | 5.57E-19 | 4.94E-18 |
| ENSG00000092208 | GEMIN2 | -0.77 | 0.09 | 1.28E-17 | 1.07E-16 |
| ENSG00000183018 | SPNS2 | -0.77 | 0.1 | 2.04E-15 | 1.54E-14 |
| ENSG00000166140 | ZFYVE19 | -0.77 | 0.11 | 1.30E-13 | 9.01E-13 |
| ENSG00000135315 | CEP162 | -0.77 | 0.11 | 1.73E-13 | 1.19E-12 |
| ENSG00000119698 | PPP4R4 | -0.77 | 0.13 | 4.87E-10 | 2.69E-09 |
| ENSG00000275395 | FCGBP | -0.77 | 0.17 | 9.44E-07 | 4.01E-06 |
| ENSG00000064787 | BCAS1 | -0.77 | 0.21 | 3.58E-05 | 1.30E-04 |
| ENSG00000280010 | | -0.77 | 0.31 | 1.96E-03 | 5.69E-03 |
| ENSG00000142185 | TRPM2 | -0.77 | 0.33 | 2.77E-03 | 7.84E-03 |
| ENSG00000260858 | | -0.77 | 0.79 | 7.54E-03 | 1.96E-02 |
| ENSG00000226851 | ZNF277-AS1 | -0.77 | 0.77 | 9.62E-03 | 2.44E-02 |
| ENSG00000126016 | AMOT | -0.77 | 0.45 | 1.13E-02 | 2.81E-02 |
| ENSG00000198732 | SMOC1 | -0.77 | 0.73 | 1.33E-02 | 3.26E-02 |
| ENSG00000288070 | | -0.77 | 0.51 | 1.40E-02 | 3.41E-02 |
| ENSG00000279347 | | -0.77 | 0.5 | 1.43E-02 | 3.48E-02 |
| ENSG00000005102 | MEOX1 | -0.77 | 0.56 | 1.73E-02 | 4.13E-02 |
| ENSG00000215210 | RBMXP2 | -0.77 | 0.59 | 1.79E-02 | 4.25E-02 |
| ENSG00000089327 | FXYD5 | -0.76 | 0.05 | 4.84E-64 | 1.47E-62 |
| ENSG00000078140 | UBE2K | -0.76 | 0.05 | 6.28E-47 | 1.32E-45 |
| ENSG00000205060 | SLC35B4 | -0.76 | 0.06 | 8.61E-43 | 1.66E-41 |
| ENSG00000120526 | NUDCD1 | -0.76 | 0.06 | 1.55E-41 | 2.90E-40 |
| ENSG00000137571 | SLCO5A1 | -0.76 | 0.06 | 1.13E-38 | 1.98E-37 |
| ENSG00000116138 | DNAJC16 | -0.76 | 0.06 | 3.04E-37 | 5.08E-36 |
| ENSG00000093000 | NUP50 | -0.76 | 0.06 | 1.02E-36 | 1.69E-35 |
| ENSG00000151131 | NOPCHAP1 | -0.76 | 0.06 | 3.43E-35 | 5.40E-34 |

| | | | | | |
|---|---|---|---|---|---|
| ENSG00000106034 | CPED1 | -0.76 | 0.06 | 5.10E-35 | 8.00E-34 |
| ENSG00000063601 | MTMR1 | -0.76 | 0.06 | 2.69E-34 | 4.11E-33 |
| ENSG00000117593 | DARS2 | -0.76 | 0.06 | 3.93E-34 | 5.98E-33 |
| ENSG00000108443 | RPS6KB1 | -0.76 | 0.06 | 6.69E-33 | 9.81E-32 |
| ENSG00000138379 | MSTN | -0.76 | 0.06 | 7.36E-33 | 1.08E-31 |
| ENSG00000196498 | NCOR2 | -0.76 | 0.07 | 2.35E-32 | 3.40E-31 |
| ENSG00000148296 | SURF6 | -0.76 | 0.07 | 1.97E-31 | 2.78E-30 |
| ENSG00000174903 | RAB1B | -0.76 | 0.07 | 3.43E-28 | 4.31E-27 |
| ENSG00000168813 | ZNF507 | -0.76 | 0.07 | 1.86E-27 | 2.30E-26 |
| ENSG00000171497 | PPID | -0.76 | 0.07 | 1.47E-26 | 1.76E-25 |
| ENSG00000133131 | MORC4 | -0.76 | 0.08 | 4.45E-24 | 4.86E-23 |
| ENSG00000275342 | PRAG1 | -0.76 | 0.08 | 4.34E-23 | 4.56E-22 |
| ENSG00000103047 | TANGO6 | -0.76 | 0.08 | 1.22E-22 | 1.26E-21 |
| ENSG00000027001 | MIPEP | -0.76 | 0.08 | 1.32E-22 | 1.36E-21 |
| ENSG00000054611 | TBC1D22A | -0.76 | 0.08 | 3.79E-22 | 3.84E-21 |
| ENSG00000164252 | AGGF1 | -0.76 | 0.08 | 4.76E-22 | 4.80E-21 |
| ENSG00000168016 | TRANK1 | -0.76 | 0.08 | 2.27E-20 | 2.15E-19 |
| ENSG00000125885 | MCM8 | -0.76 | 0.08 | 7.59E-20 | 7.01E-19 |
| ENSG00000184489 | PTP4A3 | -0.76 | 0.09 | 9.71E-20 | 8.93E-19 |
| ENSG00000213988 | ZNF90 | -0.76 | 0.1 | 3.11E-16 | 2.44E-15 |
| ENSG00000139734 | DIAPH3 | -0.76 | 0.1 | 3.96E-16 | 3.10E-15 |
| ENSG00000237765 | FAM200B | -0.76 | 0.1 | 6.54E-16 | 5.06E-15 |
| ENSG00000186174 | BCL9L | -0.76 | 0.1 | 7.37E-16 | 5.67E-15 |
| ENSG00000119946 | CNNM1 | -0.76 | 0.1 | 9.43E-16 | 7.22E-15 |
| ENSG00000134490 | TMEM241 | -0.76 | 0.1 | 7.78E-15 | 5.73E-14 |
| ENSG00000261379 | | -0.76 | 0.11 | 8.26E-13 | 5.45E-12 |
| ENSG00000198346 | ZNF813 | -0.76 | 0.11 | 1.95E-12 | 1.26E-11 |
| ENSG00000157502 | PWWP3B | -0.76 | 0.12 | 1.55E-10 | 8.89E-10 |
| ENSG00000269044 | | -0.76 | 0.13 | 9.94E-10 | 5.36E-09 |
| ENSG00000169372 | CRADD | -0.76 | 0.13 | 1.01E-09 | 5.44E-09 |
| ENSG00000106789 | CORO2A | -0.76 | 0.13 | 2.50E-09 | 1.31E-08 |
| ENSG00000167994 | RAB3IL1 | -0.76 | 0.14 | 1.23E-08 | 6.11E-08 |
| ENSG00000158186 | MRAS | -0.76 | 0.15 | 3.96E-08 | 1.89E-07 |
| ENSG00000180616 | SSTR2 | -0.76 | 0.16 | 1.49E-07 | 6.80E-07 |
| ENSG00000125384 | PTGER2 | -0.76 | 0.17 | 1.61E-06 | 6.72E-06 |
| ENSG00000151062 | CACNA2D4 | -0.76 | 0.19 | 7.30E-06 | 2.86E-05 |
| ENSG00000135605 | TEC | -0.76 | 0.22 | 8.51E-05 | 2.98E-04 |
| ENSG00000164406 | LEAP2 | -0.76 | 0.36 | 4.87E-03 | 1.31E-02 |
| ENSG00000286600 | | -0.76 | 0.8 | 6.86E-03 | 1.80E-02 |
| ENSG00000160505 | NLRP4 | -0.76 | 0.8 | 7.52E-03 | 1.96E-02 |
| ENSG00000277151 | | -0.76 | 0.77 | 1.05E-02 | 2.63E-02 |
| ENSG00000204022 | LIPJ | -0.76 | 0.75 | 1.16E-02 | 2.89E-02 |
| ENSG00000196420 | S100A5 | -0.76 | 0.45 | 1.21E-02 | 2.99E-02 |
| ENSG00000275476 | | -0.76 | 0.46 | 1.23E-02 | 3.05E-02 |
| ENSG00000187952 | HS6ST1P1 | -0.76 | 0.46 | 1.25E-02 | 3.09E-02 |
| ENSG00000263874 | LASP1NB | -0.76 | 0.49 | 1.46E-02 | 3.54E-02 |
| ENSG00000147481 | SNTG1 | -0.76 | 0.5 | 1.49E-02 | 3.62E-02 |
| ENSG00000286795 | | -0.76 | 0.53 | 1.73E-02 | 4.14E-02 |
| ENSG00000267391 | MIR122HG | -0.76 | 0.56 | 1.79E-02 | 4.26E-02 |
| ENSG00000250961 | RMEL3 | -0.76 | 0.64 | 1.91E-02 | 4.52E-02 |

| | | | | | |
|---|---|---|---|---|---|
| ENSG00000139926 | FRMD6 | -0.75 | 0.04 | 6.29E-72 | 2.25E-70 |
| ENSG00000166025 | AMOTL1 | -0.75 | 0.04 | 5.21E-69 | 1.75E-67 |
| ENSG00000088205 | DDX18 | -0.75 | 0.05 | 3.06E-51 | 7.08E-50 |
| ENSG00000146676 | PURB | -0.75 | 0.05 | 7.20E-49 | 1.57E-47 |
| ENSG00000171865 | RNASEH1 | -0.75 | 0.05 | 1.46E-46 | 3.04E-45 |
| ENSG00000165424 | ZCCHC24 | -0.75 | 0.05 | 8.35E-44 | 1.65E-42 |
| ENSG00000116128 | BCL9 | -0.75 | 0.05 | 1.14E-43 | 2.25E-42 |
| ENSG00000115053 | NCL | -0.75 | 0.05 | 1.24E-43 | 2.44E-42 |
| ENSG00000168685 | IL7R | -0.75 | 0.06 | 9.05E-41 | 1.66E-39 |
| ENSG00000176454 | LPCAT4 | -0.75 | 0.06 | 4.98E-39 | 8.75E-38 |
| ENSG00000076716 | GPC4 | -0.75 | 0.06 | 7.85E-39 | 1.38E-37 |
| ENSG00000153944 | MSI2 | -0.75 | 0.06 | 2.26E-33 | 3.37E-32 |
| ENSG00000164684 | ZNF704 | -0.75 | 0.06 | 1.49E-32 | 2.17E-31 |
| ENSG00000134882 | UBAC2 | -0.75 | 0.07 | 4.46E-29 | 5.80E-28 |
| ENSG00000170734 | POLH | -0.75 | 0.07 | 4.99E-26 | 5.84E-25 |
| ENSG00000115129 | TP53I3 | -0.75 | 0.07 | 5.49E-25 | 6.20E-24 |
| ENSG00000149782 | PLCB3 | -0.75 | 0.08 | 7.00E-24 | 7.59E-23 |
| ENSG00000167862 | MRPL58 | -0.75 | 0.08 | 7.91E-21 | 7.60E-20 |
| ENSG00000156787 | TBC1D31 | -0.75 | 0.08 | 2.19E-20 | 2.07E-19 |
| ENSG00000164626 | KCNK5 | -0.75 | 0.09 | 4.08E-19 | 3.65E-18 |
| ENSG00000089127 | OAS1 | -0.75 | 0.09 | 3.94E-18 | 3.36E-17 |
| ENSG00000177000 | MTHFR | -0.75 | 0.09 | 8.51E-18 | 7.16E-17 |
| ENSG00000167476 | JSRP1 | -0.75 | 0.1 | 3.19E-15 | 2.39E-14 |
| ENSG00000073050 | XRCC1 | -0.75 | 0.1 | 1.61E-14 | 1.16E-13 |
| ENSG00000104518 | GSDMD | -0.75 | 0.11 | 1.16E-11 | 7.13E-11 |
| ENSG00000164330 | EBF1 | -0.75 | 0.15 | 5.19E-08 | 2.46E-07 |
| ENSG00000172671 | ZFAND4 | -0.75 | 0.16 | 2.23E-07 | 1.01E-06 |
| ENSG00000230882 | | -0.75 | 0.17 | 8.78E-07 | 3.74E-06 |
| ENSG00000115194 | SLC30A3 | -0.75 | 0.18 | 6.83E-06 | 2.69E-05 |
| ENSG00000080007 | DDX43 | -0.75 | 0.18 | 7.66E-06 | 2.99E-05 |
| ENSG00000166578 | IQCD | -0.75 | 0.24 | 3.07E-04 | 1.00E-03 |
| ENSG00000259291 | ZNF710-AS1 | -0.75 | 0.31 | 2.00E-03 | 5.80E-03 |
| ENSG00000286186 | | -0.75 | 0.31 | 2.48E-03 | 7.07E-03 |
| ENSG00000237061 | | -0.75 | 0.81 | 6.20E-03 | 1.64E-02 |
| ENSG00000291096 | | -0.75 | 0.79 | 8.49E-03 | 2.18E-02 |
| ENSG00000180245 | RRH | -0.75 | 0.41 | 9.55E-03 | 2.42E-02 |
| ENSG00000290082 | | -0.75 | 0.42 | 9.83E-03 | 2.49E-02 |
| ENSG00000291071 | FAM86JP | -0.75 | 0.44 | 1.17E-02 | 2.92E-02 |
| ENSG00000169856 | ONECUT1 | -0.75 | 0.45 | 1.20E-02 | 2.98E-02 |
| ENSG00000280693 | SH3PXD2A-AS1 | -0.75 | 0.67 | 1.84E-02 | 4.38E-02 |
| ENSG00000085276 | MECOM | -0.75 | 0.67 | 1.86E-02 | 4.41E-02 |
| ENSG00000213013 | RPS15AP36 | -0.75 | 0.62 | 1.98E-02 | 4.68E-02 |
| ENSG00000212421 | | -0.75 | 0.64 | 1.99E-02 | 4.69E-02 |
| ENSG00000010292 | NCAPD2 | -0.74 | 0.04 | 7.56E-82 | 3.21E-80 |
| ENSG00000198700 | IPO9 | -0.74 | 0.05 | 1.30E-60 | 3.68E-59 |
| ENSG00000136261 | BZW2 | -0.74 | 0.05 | 1.29E-52 | 3.12E-51 |
| ENSG00000197321 | SVIL | -0.74 | 0.06 | 1.38E-37 | 2.34E-36 |
| ENSG00000071539 | TRIP13 | -0.74 | 0.06 | 1.27E-33 | 1.91E-32 |
| ENSG00000134871 | COL4A2 | -0.74 | 0.06 | 4.06E-32 | 5.82E-31 |
| ENSG00000119787 | ATL2 | -0.74 | 0.07 | 2.28E-30 | 3.10E-29 |

| | | | | | |
|---|---|---|---|---|---|
| ENSG00000171302 | CANT1 | -0.74 | 0.07 | 3.40E-29 | 4.45E-28 |
| ENSG00000060971 | ACAA1 | -0.74 | 0.07 | 4.19E-27 | 5.10E-26 |
| ENSG00000142634 | EFHD2 | -0.74 | 0.08 | 2.92E-23 | 3.08E-22 |
| ENSG00000104221 | BRF2 | -0.74 | 0.08 | 4.33E-23 | 4.55E-22 |
| ENSG00000114739 | ACVR2B | -0.74 | 0.08 | 1.35E-22 | 1.39E-21 |
| ENSG00000198521 | ZNF43 | -0.74 | 0.08 | 1.44E-22 | 1.48E-21 |
| ENSG00000081014 | AP4E1 | -0.74 | 0.08 | 2.94E-20 | 2.77E-19 |
| ENSG00000143633 | C1orf131 | -0.74 | 0.09 | 3.35E-18 | 2.87E-17 |
| ENSG00000167333 | TRIM68 | -0.74 | 0.09 | 3.39E-18 | 2.91E-17 |
| ENSG00000135838 | NPL | -0.74 | 0.09 | 1.62E-17 | 1.35E-16 |
| ENSG00000106077 | ABHD11 | -0.74 | 0.1 | 9.61E-16 | 7.35E-15 |
| ENSG00000092445 | TYRO3 | -0.74 | 0.09 | 1.15E-15 | 8.76E-15 |
| ENSG00000101311 | FERMT1 | -0.74 | 0.1 | 4.59E-15 | 3.41E-14 |
| ENSG00000186523 | FAM86B1 | -0.74 | 0.11 | 1.84E-12 | 1.19E-11 |
| ENSG00000112280 | COL9A1 | -0.74 | 0.12 | 2.05E-10 | 1.16E-09 |
| ENSG00000115041 | KCNIP3 | -0.74 | 0.17 | 1.56E-06 | 6.52E-06 |
| ENSG00000268119 | | -0.74 | 0.17 | 3.15E-06 | 1.28E-05 |
| ENSG00000228624 | HDAC2-AS2 | -0.74 | 0.18 | 5.23E-06 | 2.08E-05 |
| ENSG00000186051 | TAL2 | -0.74 | 0.2 | 2.30E-05 | 8.58E-05 |
| ENSG00000179071 | CCDC89 | -0.74 | 0.23 | 1.90E-04 | 6.39E-04 |
| ENSG00000128805 | ARHGAP22 | -0.74 | 0.23 | 2.17E-04 | 7.26E-04 |
| ENSG00000129167 | TPH1 | -0.74 | 0.27 | 1.00E-03 | 3.04E-03 |
| ENSG00000205683 | DPF3 | -0.74 | 0.29 | 1.49E-03 | 4.40E-03 |
| ENSG00000270248 | | -0.74 | 0.3 | 1.91E-03 | 5.56E-03 |
| ENSG00000129988 | LBP | -0.74 | 0.46 | 1.42E-02 | 3.46E-02 |
| ENSG00000266573 | | -0.74 | 0.51 | 1.80E-02 | 4.29E-02 |
| ENSG00000258378 | | -0.74 | 0.67 | 2.00E-02 | 4.70E-02 |
| ENSG00000196230 | TUBB | -0.73 | 0.03 | 7.73E-117 | 5.42E-115 |
| ENSG00000135372 | NAT10 | -0.73 | 0.04 | 2.19E-90 | 1.07E-88 |
| ENSG00000095319 | NUP188 | -0.73 | 0.04 | 1.33E-73 | 4.90E-72 |
| ENSG00000107566 | ERLIN1 | -0.73 | 0.04 | 2.45E-68 | 8.05E-67 |
| ENSG00000184117 | NIPSNAP1 | -0.73 | 0.04 | 2.36E-61 | 6.77E-60 |
| ENSG00000149532 | CPSF7 | -0.73 | 0.04 | 1.26E-60 | 3.58E-59 |
| ENSG00000064703 | DDX20 | -0.73 | 0.05 | 1.34E-52 | 3.21E-51 |
| ENSG00000155545 | MIER3 | -0.73 | 0.05 | 1.05E-47 | 2.23E-46 |
| ENSG00000146350 | TBC1D32 | -0.73 | 0.05 | 1.99E-45 | 4.05E-44 |
| ENSG00000173599 | PC | -0.73 | 0.05 | 5.73E-42 | 1.09E-40 |
| ENSG00000162664 | ZNF326 | -0.73 | 0.06 | 6.71E-41 | 1.24E-39 |
| ENSG00000159348 | CYB5R1 | -0.73 | 0.06 | 7.92E-41 | 1.46E-39 |
| ENSG00000082458 | DLG3 | -0.73 | 0.06 | 4.16E-35 | 6.55E-34 |
| ENSG00000163820 | FYCO1 | -0.73 | 0.06 | 1.72E-33 | 2.58E-32 |
| ENSG00000115355 | CCDC88A | -0.73 | 0.06 | 2.30E-31 | 3.24E-30 |
| ENSG00000042088 | TDP1 | -0.73 | 0.07 | 1.01E-29 | 1.34E-28 |
| ENSG00000142686 | C1orf216 | -0.73 | 0.07 | 8.32E-25 | 9.34E-24 |
| ENSG00000180901 | KCTD2 | -0.73 | 0.07 | 2.81E-24 | 3.09E-23 |
| ENSG00000104524 | PYCR3 | -0.73 | 0.08 | 6.31E-23 | 6.60E-22 |
| ENSG00000100285 | NEFH | -0.73 | 0.1 | 3.24E-14 | 2.31E-13 |
| ENSG00000054654 | SYNE2 | -0.73 | 0.1 | 6.71E-13 | 4.45E-12 |
| ENSG00000196793 | ZNF239 | -0.73 | 0.12 | 2.17E-10 | 1.23E-09 |
| ENSG00000011332 | DPF1 | -0.73 | 0.13 | 2.23E-09 | 1.17E-08 |

| | | | | | |
|---|---|---|---|---|---|
| ENSG00000248092 | NNT-AS1 | -0.73 | 0.13 | 3.88E-09 | 2.00E-08 |
| ENSG00000084636 | COL16A1 | -0.73 | 0.13 | 4.60E-09 | 2.35E-08 |
| ENSG00000114737 | CISH | -0.73 | 0.14 | 1.78E-08 | 8.75E-08 |
| ENSG00000106031 | HOXA13 | -0.73 | 0.14 | 4.61E-08 | 2.19E-07 |
| ENSG00000010626 | LRRC23 | -0.73 | 0.15 | 2.24E-07 | 1.01E-06 |
| ENSG00000169071 | ROR2 | -0.73 | 0.2 | 4.72E-05 | 1.70E-04 |
| ENSG00000111885 | MAN1A1 | -0.73 | 0.2 | 4.79E-05 | 1.72E-04 |
| ENSG00000230797 | YY2 | -0.73 | 0.2 | 5.79E-05 | 2.07E-04 |
| ENSG00000229321 | | -0.73 | 0.33 | 4.51E-03 | 1.22E-02 |
| ENSG00000198691 | ABCA4 | -0.73 | 0.35 | 5.10E-03 | 1.37E-02 |
| ENSG00000288079 | | -0.73 | 0.35 | 5.25E-03 | 1.41E-02 |
| ENSG00000176349 | | -0.73 | 0.36 | 6.43E-03 | 1.69E-02 |
| ENSG00000235169 | SMIM1 | -0.73 | 0.4 | 9.64E-03 | 2.44E-02 |
| ENSG00000200847 | Y_RNA | -0.73 | 0.68 | 1.99E-02 | 4.68E-02 |
| ENSG00000160710 | ADAR | -0.72 | 0.03 | 4.98E-99 | 2.85E-97 |
| ENSG00000140945 | CDH13 | -0.72 | 0.04 | 5.76E-64 | 1.73E-62 |
| ENSG00000129353 | SLC44A2 | -0.72 | 0.04 | 9.18E-63 | 2.70E-61 |
| ENSG00000180370 | PAK2 | -0.72 | 0.04 | 6.25E-61 | 1.78E-59 |
| ENSG00000154978 | VOPP1 | -0.72 | 0.04 | 1.37E-60 | 3.89E-59 |
| ENSG00000119977 | TCTN3 | -0.72 | 0.05 | 5.23E-53 | 1.27E-51 |
| ENSG00000153560 | UBP1 | -0.72 | 0.05 | 2.98E-52 | 7.08E-51 |
| ENSG00000171150 | SOCS5 | -0.72 | 0.06 | 5.08E-40 | 9.09E-39 |
| ENSG00000177238 | TRIM72 | -0.72 | 0.06 | 1.42E-37 | 2.40E-36 |
| ENSG00000123600 | METTL8 | -0.72 | 0.06 | 8.31E-31 | 1.15E-29 |
| ENSG00000102034 | ELF4 | -0.72 | 0.06 | 1.10E-30 | 1.52E-29 |
| ENSG00000140396 | NCOA2 | -0.72 | 0.07 | 1.15E-28 | 1.47E-27 |
| ENSG00000204131 | NHSL2 | -0.72 | 0.07 | 1.79E-28 | 2.28E-27 |
| ENSG00000184840 | TMED9 | -0.72 | 0.07 | 5.38E-28 | 6.72E-27 |
| ENSG00000166734 | GOLM2 | -0.72 | 0.07 | 8.40E-27 | 1.01E-25 |
| ENSG00000105447 | GRWD1 | -0.72 | 0.07 | 5.70E-26 | 6.66E-25 |
| ENSG00000125871 | MGME1 | -0.72 | 0.07 | 2.53E-25 | 2.89E-24 |
| ENSG00000178425 | NT5DC1 | -0.72 | 0.07 | 2.10E-24 | 2.32E-23 |
| ENSG00000124225 | PMEPA1 | -0.72 | 0.08 | 2.40E-22 | 2.45E-21 |
| ENSG00000084710 | EFR3B | -0.72 | 0.08 | 2.59E-22 | 2.64E-21 |
| ENSG00000171295 | ZNF440 | -0.72 | 0.08 | 4.61E-22 | 4.65E-21 |
| ENSG00000179041 | RRS1 | -0.72 | 0.08 | 2.22E-20 | 2.09E-19 |
| ENSG00000260807 | CEROX1 | -0.72 | 0.08 | 1.15E-19 | 1.05E-18 |
| ENSG00000242759 | LINC00882 | -0.72 | 0.08 | 1.22E-18 | 1.07E-17 |
| ENSG00000158122 | PRXL2C | -0.72 | 0.09 | 5.61E-18 | 4.76E-17 |
| ENSG00000163328 | GPR155 | -0.72 | 0.12 | 1.31E-09 | 7.01E-09 |
| ENSG00000173269 | MMRN2 | -0.72 | 0.13 | 1.19E-08 | 5.92E-08 |
| ENSG00000267904 | | -0.72 | 0.18 | 1.05E-05 | 4.04E-05 |
| ENSG00000246985 | SOCS2-AS1 | -0.72 | 0.22 | 1.96E-04 | 6.57E-04 |
| ENSG00000253741 | LNCOC1 | -0.72 | 0.24 | 4.87E-04 | 1.55E-03 |
| ENSG00000197275 | RAD54B | -0.72 | 0.25 | 5.95E-04 | 1.87E-03 |
| ENSG00000183621 | ZNF438 | -0.72 | 0.27 | 1.32E-03 | 3.92E-03 |
| ENSG00000082196 | C1QTNF3 | -0.72 | 0.29 | 2.01E-03 | 5.81E-03 |
| ENSG00000236308 | | -0.72 | 0.29 | 2.27E-03 | 6.52E-03 |
| ENSG00000165507 | DEPP1 | -0.72 | 0.31 | 3.39E-03 | 9.41E-03 |
| ENSG00000101443 | WFDC2 | -0.72 | 0.38 | 9.13E-03 | 2.33E-02 |

| | | | | | |
|---|---|---|---|---|---|
| ENSG00000185674 | LYG2 | -0.72 | 0.38 | 9.70E-03 | 2.46E-02 |
| ENSG00000204446 | LINC02872 | -0.72 | 0.39 | 9.86E-03 | 2.49E-02 |
| ENSG00000277806 | | -0.72 | 0.4 | 1.05E-02 | 2.63E-02 |
| ENSG00000233901 | LINC01503 | -0.72 | 0.4 | 1.12E-02 | 2.80E-02 |
| ENSG00000075429 | CACNG5 | -0.72 | 0.72 | 1.67E-02 | 3.99E-02 |
| ENSG00000143799 | PARP1 | -0.71 | 0.03 | 6.22E-105 | 3.84E-103 |
| ENSG00000114251 | WNT5A | -0.71 | 0.04 | 9.02E-58 | 2.42E-56 |
| ENSG00000198585 | NUDT16 | -0.71 | 0.05 | 9.02E-54 | 2.23E-52 |
| ENSG00000163938 | GNL3 | -0.71 | 0.05 | 1.03E-48 | 2.22E-47 |
| ENSG00000101224 | CDC25B | -0.71 | 0.05 | 1.89E-46 | 3.93E-45 |
| ENSG00000272622 | | -0.71 | 0.05 | 1.10E-44 | 2.20E-43 |
| ENSG00000161791 | FMNL3 | -0.71 | 0.05 | 1.58E-42 | 3.04E-41 |
| ENSG00000114125 | RNF7 | -0.71 | 0.05 | 2.01E-40 | 3.64E-39 |
| ENSG00000123268 | ATF1 | -0.71 | 0.06 | 1.73E-37 | 2.92E-36 |
| ENSG00000087502 | ERGIC2 | -0.71 | 0.06 | 6.33E-35 | 9.88E-34 |
| ENSG00000125827 | TMX4 | -0.71 | 0.06 | 1.62E-33 | 2.43E-32 |
| ENSG00000184640 | SEPTIN9 | -0.71 | 0.06 | 1.38E-28 | 1.76E-27 |
| ENSG00000164022 | AIMP1 | -0.71 | 0.07 | 4.18E-28 | 5.25E-27 |
| ENSG00000118600 | RXYLT1 | -0.71 | 0.07 | 2.38E-26 | 2.82E-25 |
| ENSG00000148943 | LIN7C | -0.71 | 0.07 | 2.56E-26 | 3.03E-25 |
| ENSG00000158636 | EMSY | -0.71 | 0.07 | 3.01E-25 | 3.43E-24 |
| ENSG00000133193 | FAM104A | -0.71 | 0.07 | 3.21E-25 | 3.65E-24 |
| ENSG00000126814 | TRMT5 | -0.71 | 0.08 | 5.27E-21 | 5.12E-20 |
| ENSG00000162849 | KIF26B | -0.71 | 0.08 | 2.17E-20 | 2.05E-19 |
| ENSG00000163545 | NUAK2 | -0.71 | 0.08 | 2.71E-19 | 2.45E-18 |
| ENSG00000186532 | SMYD4 | -0.71 | 0.08 | 1.42E-18 | 1.24E-17 |
| ENSG00000137962 | ARHGAP29 | -0.71 | 0.08 | 3.65E-18 | 3.12E-17 |
| ENSG00000152953 | STK32B | -0.71 | 0.09 | 5.49E-15 | 4.07E-14 |
| ENSG00000198794 | SCAMP5 | -0.71 | 0.1 | 1.32E-14 | 9.59E-14 |
| ENSG00000263528 | IKBKE | -0.71 | 0.1 | 3.78E-14 | 2.69E-13 |
| ENSG00000107611 | CUBN | -0.71 | 0.1 | 2.00E-13 | 1.37E-12 |
| ENSG00000112877 | CEP72 | -0.71 | 0.1 | 3.53E-13 | 2.38E-12 |
| ENSG00000237440 | ZNF737 | -0.71 | 0.12 | 2.40E-10 | 1.36E-09 |
| ENSG00000114107 | CEP70 | -0.71 | 0.12 | 3.18E-10 | 1.78E-09 |
| ENSG00000147813 | NAPRT | -0.71 | 0.12 | 5.18E-10 | 2.86E-09 |
| ENSG00000169598 | DFFB | -0.71 | 0.12 | 7.49E-10 | 4.08E-09 |
| ENSG00000111834 | RSPH4A | -0.71 | 0.14 | 2.84E-08 | 1.37E-07 |
| ENSG00000137877 | SPTBN5 | -0.71 | 0.14 | 4.45E-08 | 2.12E-07 |
| ENSG00000100319 | ZMAT5 | -0.71 | 0.19 | 2.81E-05 | 1.04E-04 |
| ENSG00000204923 | FBXO48 | -0.71 | 0.2 | 4.78E-05 | 1.72E-04 |
| ENSG00000198324 | PHETA1 | -0.71 | 0.25 | 6.54E-04 | 2.05E-03 |
| ENSG00000111261 | MANSC1 | -0.71 | 0.27 | 1.27E-03 | 3.79E-03 |
| ENSG00000261655 | | -0.71 | 0.28 | 1.72E-03 | 5.03E-03 |
| ENSG00000249550 | LINC01234 | -0.71 | 0.3 | 3.12E-03 | 8.73E-03 |
| ENSG00000242715 | CCDC169 | -0.71 | 0.31 | 3.59E-03 | 9.94E-03 |
| ENSG00000289026 | | -0.71 | 0.32 | 4.42E-03 | 1.20E-02 |
| ENSG00000271828 | | -0.71 | 0.32 | 4.71E-03 | 1.27E-02 |
| ENSG00000284693 | LINC02606 | -0.71 | 0.32 | 4.87E-03 | 1.31E-02 |
| ENSG00000212396 | RNA5SP323 | -0.71 | 0.8 | 7.46E-03 | 1.94E-02 |
| ENSG00000279805 | | -0.71 | 0.8 | 7.51E-03 | 1.95E-02 |

| | | | | | |
|---|---|---|---|---|---|
| ENSG00000143632 | ACTA1 | -0.71 | 0.35 | 7.53E-03 | 1.96E-02 |
| ENSG00000234503 | | -0.71 | 0.79 | 8.54E-03 | 2.19E-02 |
| ENSG00000264116 | | -0.71 | 0.37 | 8.90E-03 | 2.28E-02 |
| ENSG00000179023 | KLHDC7A | -0.71 | 0.79 | 9.12E-03 | 2.33E-02 |
| ENSG00000149571 | KIRREL3 | -0.71 | 0.4 | 1.16E-02 | 2.90E-02 |
| ENSG00000167553 | TUBA1C | -0.7 | 0.04 | 5.21E-68 | 1.70E-66 |
| ENSG00000113845 | TIMMDC1 | -0.7 | 0.04 | 2.69E-57 | 7.14E-56 |
| ENSG00000143153 | ATP1B1 | -0.7 | 0.04 | 5.60E-57 | 1.47E-55 |
| ENSG00000189403 | HMGB1 | -0.7 | 0.05 | 3.66E-50 | 8.25E-49 |
| ENSG00000170004 | CHD3 | -0.7 | 0.05 | 5.79E-46 | 1.19E-44 |
| ENSG00000111727 | HCFC2 | -0.7 | 0.05 | 7.07E-41 | 1.30E-39 |
| ENSG00000197312 | DDI2 | -0.7 | 0.05 | 2.80E-40 | 5.05E-39 |
| ENSG00000115652 | UXS1 | -0.7 | 0.05 | 4.29E-38 | 7.37E-37 |
| ENSG00000125741 | OPA3 | -0.7 | 0.06 | 1.12E-37 | 1.91E-36 |
| ENSG00000197894 | ADH5 | -0.7 | 0.06 | 2.22E-37 | 3.73E-36 |
| ENSG00000134982 | APC | -0.7 | 0.06 | 2.47E-37 | 4.14E-36 |
| ENSG00000147164 | SNX12 | -0.7 | 0.06 | 3.88E-37 | 6.46E-36 |
| ENSG00000234127 | TRIM26 | -0.7 | 0.06 | 2.10E-36 | 3.44E-35 |
| ENSG00000155760 | FZD7 | -0.7 | 0.06 | 1.47E-34 | 2.27E-33 |
| ENSG00000168883 | USP39 | -0.7 | 0.06 | 2.12E-34 | 3.25E-33 |
| ENSG00000132780 | NASP | -0.7 | 0.06 | 3.37E-34 | 5.14E-33 |
| ENSG00000163029 | SMC6 | -0.7 | 0.06 | 2.19E-32 | 3.18E-31 |
| ENSG00000164253 | WDR41 | -0.7 | 0.06 | 6.24E-32 | 8.91E-31 |
| ENSG00000068784 | SRBD1 | -0.7 | 0.06 | 1.33E-29 | 1.76E-28 |
| ENSG00000152495 | CAMK4 | -0.7 | 0.07 | 1.56E-27 | 1.93E-26 |
| ENSG00000173898 | SPTBN2 | -0.7 | 0.07 | 4.69E-25 | 5.29E-24 |
| ENSG00000143553 | SNAPIN | -0.7 | 0.07 | 1.60E-24 | 1.78E-23 |
| ENSG00000176542 | USF3 | -0.7 | 0.07 | 5.29E-24 | 5.75E-23 |
| ENSG00000104412 | EMC2 | -0.7 | 0.07 | 2.88E-23 | 3.04E-22 |
| ENSG00000128944 | KNSTRN | -0.7 | 0.07 | 2.61E-22 | 2.66E-21 |
| ENSG00000188486 | H2AX | -0.7 | 0.08 | 5.91E-21 | 5.72E-20 |
| ENSG00000106348 | IMPDH1 | -0.7 | 0.08 | 2.69E-20 | 2.53E-19 |
| ENSG00000085760 | MTIF2 | -0.7 | 0.08 | 3.04E-17 | 2.51E-16 |
| ENSG00000132394 | EEFSEC | -0.7 | 0.09 | 2.74E-16 | 2.16E-15 |
| ENSG00000167107 | ACSF2 | -0.7 | 0.11 | 2.82E-11 | 1.70E-10 |
| ENSG00000106003 | LFNG | -0.7 | 0.11 | 4.60E-11 | 2.72E-10 |
| ENSG00000232445 | EMSLR | -0.7 | 0.12 | 2.09E-10 | 1.19E-09 |
| ENSG00000155465 | SLC7A7 | -0.7 | 0.12 | 1.13E-09 | 6.06E-09 |
| ENSG00000124251 | TP53TG5 | -0.7 | 0.18 | 1.42E-05 | 5.42E-05 |
| ENSG00000146205 | ANO7 | -0.7 | 0.19 | 6.36E-05 | 2.26E-04 |
| ENSG00000241472 | PTPRG-AS1 | -0.7 | 0.24 | 7.12E-04 | 2.22E-03 |
| ENSG00000255121 | CENATAC-DT | -0.7 | 0.24 | 7.46E-04 | 2.31E-03 |
| ENSG00000131781 | FMO5 | -0.7 | 0.25 | 7.53E-04 | 2.33E-03 |
| ENSG00000187372 | PCDHB13 | -0.7 | 0.26 | 1.19E-03 | 3.58E-03 |
| ENSG00000286873 | | -0.7 | 0.26 | 1.49E-03 | 4.41E-03 |
| ENSG00000185668 | POU3F1 | -0.7 | 0.31 | 4.00E-03 | 1.10E-02 |
| ENSG00000289223 | | -0.7 | 0.33 | 5.79E-03 | 1.54E-02 |
| ENSG00000175591 | P2RY2 | -0.7 | 0.38 | 1.02E-02 | 2.57E-02 |
| ENSG00000287271 | | -0.7 | 0.38 | 1.05E-02 | 2.64E-02 |
| ENSG00000188039 | NWD1 | -0.7 | 0.78 | 1.06E-02 | 2.66E-02 |

| | | | | | |
|---|---|---|---|---|---|
| ENSG00000286603 | | -0.7 | 0.41 | 1.38E-02 | 3.38E-02 |
| ENSG00000112562 | SMOC2 | -0.7 | 0.4 | 1.38E-02 | 3.38E-02 |
| ENSG00000174640 | SLCO2A1 | -0.7 | 0.72 | 1.77E-02 | 4.22E-02 |
| ENSG00000198121 | LPAR1 | -0.7 | 0.71 | 2.02E-02 | 4.75E-02 |
| ENSG00000096384 | HSP90AB1 | -0.69 | 0.03 | 1.82E-90 | 8.96E-89 |
| ENSG00000099942 | CRKL | -0.69 | 0.04 | 2.16E-69 | 7.35E-68 |
| ENSG00000110713 | NUP98 | -0.69 | 0.04 | 2.23E-69 | 7.55E-68 |
| ENSG00000151617 | EDNRA | -0.69 | 0.04 | 6.80E-59 | 1.87E-57 |
| ENSG00000137054 | POLR1E | -0.69 | 0.04 | 3.08E-57 | 8.13E-56 |
| ENSG00000120800 | UTP20 | -0.69 | 0.04 | 1.56E-53 | 3.84E-52 |
| ENSG00000143320 | CRABP2 | -0.69 | 0.05 | 5.06E-52 | 1.20E-50 |
| ENSG00000132383 | RPA1 | -0.69 | 0.05 | 1.22E-51 | 2.86E-50 |
| ENSG00000109519 | GRPEL1 | -0.69 | 0.05 | 2.66E-51 | 6.18E-50 |
| ENSG00000064393 | HIPK2 | -0.69 | 0.05 | 7.55E-44 | 1.50E-42 |
| ENSG00000196730 | DAPK1 | -0.69 | 0.05 | 2.47E-43 | 4.82E-42 |
| ENSG00000085377 | PREP | -0.69 | 0.05 | 2.09E-42 | 4.01E-41 |
| ENSG00000119938 | PPP1R3C | -0.69 | 0.05 | 2.49E-39 | 4.40E-38 |
| ENSG00000148840 | PPRC1 | -0.69 | 0.06 | 3.26E-35 | 5.13E-34 |
| ENSG00000198648 | STK39 | -0.69 | 0.06 | 7.09E-35 | 1.11E-33 |
| ENSG00000183696 | UPP1 | -0.69 | 0.06 | 3.54E-34 | 5.39E-33 |
| ENSG00000091651 | ORC6 | -0.69 | 0.06 | 4.63E-31 | 6.47E-30 |
| ENSG00000112576 | CCND3 | -0.69 | 0.06 | 2.21E-29 | 2.90E-28 |
| ENSG00000075711 | DLG1 | -0.69 | 0.06 | 9.86E-29 | 1.27E-27 |
| ENSG00000130921 | MTRFR | -0.69 | 0.06 | 2.13E-27 | 2.62E-26 |
| ENSG00000247077 | PGAM5 | -0.69 | 0.07 | 9.72E-24 | 1.05E-22 |
| ENSG00000198879 | SFMBT2 | -0.69 | 0.07 | 6.49E-22 | 6.52E-21 |
| ENSG00000015133 | CCDC88C | -0.69 | 0.07 | 8.90E-22 | 8.89E-21 |
| ENSG00000177352 | CCDC71 | -0.69 | 0.08 | 7.23E-19 | 6.37E-18 |
| ENSG00000172469 | MANEA | -0.69 | 0.09 | 6.91E-17 | 5.61E-16 |
| ENSG00000156504 | PABIR2 | -0.69 | 0.09 | 8.33E-15 | 6.12E-14 |
| ENSG00000214013 | GANC | -0.69 | 0.1 | 7.67E-14 | 5.37E-13 |
| ENSG00000133302 | SLF1 | -0.69 | 0.1 | 5.38E-13 | 3.59E-12 |
| ENSG00000116990 | MYCL | -0.69 | 0.1 | 3.94E-12 | 2.50E-11 |
| ENSG00000159023 | EPB41 | -0.69 | 0.1 | 5.24E-12 | 3.28E-11 |
| ENSG00000266094 | RASSF5 | -0.69 | 0.11 | 3.18E-11 | 1.90E-10 |
| ENSG00000119862 | LGALSL | -0.69 | 0.11 | 4.63E-11 | 2.74E-10 |
| ENSG00000276644 | DACH1 | -0.69 | 0.13 | 1.44E-08 | 7.12E-08 |
| ENSG00000116574 | RHOU | -0.69 | 0.14 | 1.17E-07 | 5.40E-07 |
| ENSG00000104047 | DTWD1 | -0.69 | 0.14 | 1.90E-07 | 8.62E-07 |
| ENSG00000198429 | ZNF69 | -0.69 | 0.15 | 8.18E-07 | 3.50E-06 |
| ENSG00000273820 | USP27X | -0.69 | 0.15 | 1.58E-06 | 6.59E-06 |
| ENSG00000178573 | MAF | -0.69 | 0.16 | 1.71E-06 | 7.10E-06 |
| ENSG00000109832 | DDX25 | -0.69 | 0.16 | 2.92E-06 | 1.19E-05 |
| ENSG00000107614 | TRDMT1 | -0.69 | 0.18 | 1.55E-05 | 5.88E-05 |
| ENSG00000270127 | | -0.69 | 0.18 | 1.63E-05 | 6.18E-05 |
| ENSG00000152433 | ZNF547 | -0.69 | 0.18 | 3.46E-05 | 1.27E-04 |
| ENSG00000144712 | CAND2 | -0.69 | 0.23 | 5.79E-04 | 1.83E-03 |
| ENSG00000225889 | | -0.69 | 0.24 | 8.44E-04 | 2.60E-03 |
| ENSG00000160145 | KALRN | -0.69 | 0.25 | 9.06E-04 | 2.77E-03 |
| ENSG00000089101 | CFAP61 | -0.69 | 0.25 | 1.27E-03 | 3.79E-03 |

| | | | | | |
|---|---|---|---|---|---|
| ENSG00000226562 | CYP4F26P | -0.69 | 0.28 | 2.64E-03 | 7.49E-03 |
| ENSG00000008323 | PLEKHG6 | -0.69 | 0.29 | 3.60E-03 | 9.96E-03 |
| ENSG00000146006 | LRRTM2 | -0.69 | 0.35 | 9.10E-03 | 2.32E-02 |
| ENSG00000168367 | LINC00917 | -0.69 | 0.77 | 1.14E-02 | 2.83E-02 |
| ENSG00000263551 | | -0.69 | 0.76 | 1.29E-02 | 3.18E-02 |
| ENSG00000272010 | | -0.69 | 0.41 | 1.55E-02 | 3.74E-02 |
| ENSG00000198483 | ANKRD35 | -0.69 | 0.42 | 1.73E-02 | 4.12E-02 |
| ENSG00000111412 | SPRING1 | -0.68 | 0.03 | 5.52E-88 | 2.62E-86 |
| ENSG00000166226 | CCT2 | -0.68 | 0.04 | 2.04E-76 | 7.97E-75 |
| ENSG00000141076 | UTP4 | -0.68 | 0.04 | 2.29E-69 | 7.72E-68 |
| ENSG00000112118 | MCM3 | -0.68 | 0.04 | 1.09E-55 | 2.80E-54 |
| ENSG00000151503 | NCAPD3 | -0.68 | 0.04 | 5.11E-53 | 1.24E-51 |
| ENSG00000121210 | TMEM131L | -0.68 | 0.05 | 1.08E-42 | 2.07E-41 |
| ENSG00000115993 | TRAK2 | -0.68 | 0.05 | 7.07E-40 | 1.26E-38 |
| ENSG00000084676 | NCOA1 | -0.68 | 0.05 | 8.99E-39 | 1.57E-37 |
| ENSG00000136451 | VEZF1 | -0.68 | 0.06 | 1.01E-35 | 1.62E-34 |
| ENSG00000174197 | MGA | -0.68 | 0.06 | 7.63E-35 | 1.19E-33 |
| ENSG00000271601 | LIX1L | -0.68 | 0.06 | 6.63E-31 | 9.21E-30 |
| ENSG00000089818 | NECAP1 | -0.68 | 0.06 | 7.03E-31 | 9.75E-30 |
| ENSG00000230989 | HSBP1 | -0.68 | 0.06 | 1.12E-30 | 1.54E-29 |
| ENSG00000105355 | PLIN3 | -0.68 | 0.06 | 1.14E-30 | 1.57E-29 |
| ENSG00000117632 | STMN1 | -0.68 | 0.06 | 8.75E-29 | 1.13E-27 |
| ENSG00000186184 | POLR1D | -0.68 | 0.07 | 1.05E-24 | 1.17E-23 |
| ENSG00000176393 | RNPEP | -0.68 | 0.07 | 3.34E-21 | 3.27E-20 |
| ENSG00000172167 | MTBP | -0.68 | 0.08 | 1.17E-18 | 1.02E-17 |
| ENSG00000196923 | PDLIM7 | -0.68 | 0.08 | 1.59E-17 | 1.32E-16 |
| ENSG00000116863 | ADPRS | -0.68 | 0.09 | 1.13E-15 | 8.63E-15 |
| ENSG00000102554 | KLF5 | -0.68 | 0.09 | 1.17E-15 | 8.94E-15 |
| ENSG00000133943 | DGLUCY | -0.68 | 0.09 | 7.25E-14 | 5.08E-13 |
| ENSG00000162337 | LRP5 | -0.68 | 0.09 | 1.17E-13 | 8.11E-13 |
| ENSG00000284837 | | -0.68 | 0.1 | 2.72E-13 | 1.84E-12 |
| ENSG00000197971 | MBP | -0.68 | 0.1 | 8.36E-13 | 5.51E-12 |
| ENSG00000178685 | PARP10 | -0.68 | 0.1 | 8.68E-13 | 5.71E-12 |
| ENSG00000196678 | ERI2 | -0.68 | 0.11 | 4.02E-10 | 2.23E-09 |
| ENSG00000204305 | AGER | -0.68 | 0.11 | 4.03E-10 | 2.24E-09 |
| ENSG00000188428 | BLOC1S5 | -0.68 | 0.12 | 9.72E-10 | 5.25E-09 |
| ENSG00000136104 | RNASEH2B | -0.68 | 0.12 | 4.49E-09 | 2.30E-08 |
| ENSG00000197769 | MAP1LC3C | -0.68 | 0.13 | 1.53E-08 | 7.57E-08 |
| ENSG00000162878 | PKDCC | -0.68 | 0.14 | 2.57E-07 | 1.15E-06 |
| ENSG00000267107 | PCAT19 | -0.68 | 0.15 | 6.72E-07 | 2.91E-06 |
| ENSG00000241316 | SUCLG2-DT | -0.68 | 0.15 | 7.31E-07 | 3.15E-06 |
| ENSG00000147231 | RADX | -0.68 | 0.15 | 1.62E-06 | 6.73E-06 |
| ENSG00000161835 | TAMALIN | -0.68 | 0.18 | 4.40E-05 | 1.59E-04 |
| ENSG00000271361 | HTATSF1P2 | -0.68 | 0.2 | 1.34E-04 | 4.58E-04 |
| ENSG00000176842 | IRX5 | -0.68 | 0.21 | 1.71E-04 | 5.76E-04 |
| ENSG00000275620 | FLJ16779 | -0.68 | 0.21 | 2.53E-04 | 8.38E-04 |
| ENSG00000129194 | SOX15 | -0.68 | 0.21 | 2.75E-04 | 9.07E-04 |
| ENSG00000258077 | | -0.68 | 0.22 | 3.48E-04 | 1.13E-03 |
| ENSG00000198807 | PAX9 | -0.68 | 0.27 | 2.19E-03 | 6.29E-03 |
| ENSG00000169418 | NPR1 | -0.68 | 0.27 | 2.30E-03 | 6.61E-03 |

| | | | | | |
|---|---|---|---|---|---|
| ENSG00000136881 | BAAT | -0.68 | 0.31 | 5.36E-03 | 1.43E-02 |
| ENSG00000184492 | FOXD4L1 | -0.68 | 0.32 | 6.09E-03 | 1.61E-02 |
| ENSG00000232906 | | -0.68 | 0.79 | 7.88E-03 | 2.04E-02 |
| ENSG00000183324 | REC114 | -0.68 | 0.78 | 9.33E-03 | 2.37E-02 |
| ENSG00000272153 | | -0.68 | 0.35 | 9.64E-03 | 2.44E-02 |
| ENSG00000200246 | RNA5SP263 | -0.68 | 0.36 | 1.11E-02 | 2.78E-02 |
| ENSG00000171115 | GIMAP8 | -0.68 | 0.76 | 1.31E-02 | 3.23E-02 |
| ENSG00000234373 | SNX18P7 | -0.68 | 0.39 | 1.39E-02 | 3.38E-02 |
| ENSG00000131187 | F12 | -0.68 | 0.43 | 1.83E-02 | 4.34E-02 |
| ENSG00000129009 | ISLR | -0.68 | 0.44 | 2.10E-02 | 4.92E-02 |
| ENSG00000166833 | NAV2 | -0.67 | 0.03 | 9.05E-85 | 4.09E-83 |
| ENSG00000131873 | CHSY1 | -0.67 | 0.04 | 7.99E-76 | 3.08E-74 |
| ENSG00000126778 | SIX1 | -0.67 | 0.04 | 9.31E-67 | 2.98E-65 |
| ENSG00000185722 | ANKFY1 | -0.67 | 0.04 | 3.75E-64 | 1.14E-62 |
| ENSG00000070614 | NDST1 | -0.67 | 0.05 | 1.81E-48 | 3.89E-47 |
| ENSG00000152990 | ADGRA3 | -0.67 | 0.05 | 8.29E-46 | 1.70E-44 |
| ENSG00000241258 | CRCP | -0.67 | 0.05 | 1.74E-44 | 3.48E-43 |
| ENSG00000197448 | GSTK1 | -0.67 | 0.05 | 6.87E-42 | 1.30E-40 |
| ENSG00000196218 | RYR1 | -0.67 | 0.05 | 4.73E-40 | 8.47E-39 |
| ENSG00000095261 | PSMD5 | -0.67 | 0.05 | 5.07E-37 | 8.42E-36 |
| ENSG00000163166 | IWS1 | -0.67 | 0.05 | 7.30E-36 | 1.18E-34 |
| ENSG00000204304 | PBX2 | -0.67 | 0.06 | 2.32E-33 | 3.46E-32 |
| ENSG00000120029 | ARMH3 | -0.67 | 0.06 | 3.50E-32 | 5.04E-31 |
| ENSG00000106012 | IQCE | -0.67 | 0.06 | 3.77E-32 | 5.42E-31 |
| ENSG00000111271 | ACAD10 | -0.67 | 0.06 | 5.42E-32 | 7.75E-31 |
| ENSG00000175110 | MRPS22 | -0.67 | 0.06 | 1.18E-29 | 1.56E-28 |
| ENSG00000128829 | EIF2AK4 | -0.67 | 0.06 | 1.03E-28 | 1.33E-27 |
| ENSG00000168924 | LETM1 | -0.67 | 0.06 | 3.02E-28 | 3.81E-27 |
| ENSG00000071243 | ING3 | -0.67 | 0.06 | 1.93E-27 | 2.38E-26 |
| ENSG00000103540 | CCP110 | -0.67 | 0.06 | 1.32E-26 | 1.59E-25 |
| ENSG00000112655 | PTK7 | -0.67 | 0.06 | 2.73E-26 | 3.23E-25 |
| ENSG00000138347 | MYPN | -0.67 | 0.07 | 6.95E-24 | 7.54E-23 |
| ENSG00000165895 | ARHGAP42 | -0.67 | 0.07 | 1.02E-23 | 1.09E-22 |
| ENSG00000005700 | IBTK | -0.67 | 0.07 | 1.92E-23 | 2.05E-22 |
| ENSG00000122376 | SHLD2 | -0.67 | 0.07 | 2.00E-23 | 2.13E-22 |
| ENSG00000197977 | ELOVL2 | -0.67 | 0.07 | 4.86E-22 | 4.90E-21 |
| ENSG00000244509 | APOBEC3C | -0.67 | 0.07 | 6.52E-20 | 6.04E-19 |
| ENSG00000177542 | SLC25A22 | -0.67 | 0.07 | 1.17E-19 | 1.07E-18 |
| ENSG00000280385 | | -0.67 | 0.08 | 5.15E-19 | 4.57E-18 |
| ENSG00000144959 | NCEH1 | -0.67 | 0.08 | 6.12E-18 | 5.18E-17 |
| ENSG00000184557 | SOCS3 | -0.67 | 0.09 | 1.33E-14 | 9.69E-14 |
| ENSG00000124813 | RUNX2 | -0.67 | 0.09 | 1.45E-14 | 1.05E-13 |
| ENSG00000104915 | STX10 | -0.67 | 0.09 | 1.55E-13 | 1.07E-12 |
| ENSG00000188811 | NHLRC3 | -0.67 | 0.1 | 4.69E-13 | 3.14E-12 |
| ENSG00000101986 | ABCD1 | -0.67 | 0.1 | 7.82E-13 | 5.17E-12 |
| ENSG00000177409 | SAMD9L | -0.67 | 0.1 | 3.56E-12 | 2.26E-11 |
| ENSG00000198298 | ZNF485 | -0.67 | 0.1 | 1.24E-11 | 7.63E-11 |
| ENSG00000171202 | TMEM126A | -0.67 | 0.11 | 5.57E-10 | 3.07E-09 |
| ENSG00000214367 | HAUS3 | -0.67 | 0.12 | 1.81E-09 | 9.57E-09 |
| ENSG00000183628 | DGCR6 | -0.67 | 0.13 | 3.06E-08 | 1.48E-07 |

| | | | | | |
|---|---|---|---|---|---|
| ENSG00000080503 | SMARCA2 | -0.67 | 0.13 | 5.98E-08 | 2.82E-07 |
| ENSG00000014138 | POLA2 | -0.67 | 0.14 | 8.11E-07 | 3.47E-06 |
| ENSG00000289810 | | -0.67 | 0.15 | 2.79E-06 | 1.14E-05 |
| ENSG00000115457 | IGFBP2 | -0.67 | 0.17 | 1.08E-05 | 4.16E-05 |
| ENSG00000162881 | OXER1 | -0.67 | 0.18 | 5.39E-05 | 1.93E-04 |
| ENSG00000166448 | TMEM130 | -0.67 | 0.2 | 1.65E-04 | 5.59E-04 |
| ENSG00000145703 | IQGAP2 | -0.67 | 0.21 | 3.02E-04 | 9.87E-04 |
| ENSG00000233967 | | -0.67 | 0.23 | 5.84E-04 | 1.84E-03 |
| ENSG00000133106 | EPSTI1 | -0.67 | 0.22 | 5.85E-04 | 1.85E-03 |
| ENSG00000127152 | BCL11B | -0.67 | 0.24 | 8.67E-04 | 2.66E-03 |
| ENSG00000135144 | DTX1 | -0.67 | 0.24 | 8.84E-04 | 2.71E-03 |
| ENSG00000260293 | | -0.67 | 0.24 | 1.01E-03 | 3.07E-03 |
| ENSG00000143631 | FLG | -0.67 | 0.24 | 1.08E-03 | 3.25E-03 |
| ENSG00000227487 | NCAM1-AS1 | -0.67 | 0.27 | 2.83E-03 | 7.99E-03 |
| ENSG00000283031 | | -0.67 | 0.32 | 6.99E-03 | 1.83E-02 |
| ENSG00000272072 | | -0.67 | 0.34 | 9.02E-03 | 2.30E-02 |
| ENSG00000271420 | | -0.67 | 0.78 | 9.44E-03 | 2.40E-02 |
| ENSG00000240796 | MTND2P14 | -0.67 | 0.78 | 9.95E-03 | 2.51E-02 |
| ENSG00000225330 | | -0.67 | 0.78 | 1.00E-02 | 2.54E-02 |
| ENSG00000182050 | MGAT4C | -0.67 | 0.78 | 1.03E-02 | 2.59E-02 |
| ENSG00000164099 | PRSS12 | -0.67 | 0.4 | 1.62E-02 | 3.89E-02 |
| ENSG00000196660 | SLC30A10 | -0.67 | 0.42 | 1.96E-02 | 4.64E-02 |
| ENSG00000168906 | MAT2A | -0.66 | 0.04 | 1.22E-61 | 3.52E-60 |
| ENSG00000115875 | SRSF7 | -0.66 | 0.05 | 3.39E-46 | 6.98E-45 |
| ENSG00000101407 | TTI1 | -0.66 | 0.05 | 1.82E-45 | 3.71E-44 |
| ENSG00000265354 | TIMM23 | -0.66 | 0.05 | 4.14E-38 | 7.12E-37 |
| ENSG00000178802 | MPI | -0.66 | 0.05 | 1.10E-36 | 1.82E-35 |
| ENSG00000125648 | SLC25A23 | -0.66 | 0.05 | 1.56E-35 | 2.50E-34 |
| ENSG00000196628 | TCF4 | -0.66 | 0.05 | 2.35E-35 | 3.74E-34 |
| ENSG00000128512 | DOCK4 | -0.66 | 0.05 | 2.41E-35 | 3.82E-34 |
| ENSG00000112146 | FBXO9 | -0.66 | 0.06 | 7.20E-31 | 9.98E-30 |
| ENSG00000094631 | HDAC6 | -0.66 | 0.06 | 5.54E-30 | 7.44E-29 |
| ENSG00000172380 | GNG12 | -0.66 | 0.06 | 2.46E-27 | 3.02E-26 |
| ENSG00000170234 | PWWP2A | -0.66 | 0.06 | 4.81E-26 | 5.63E-25 |
| ENSG00000128335 | APOL2 | -0.66 | 0.07 | 1.11E-24 | 1.24E-23 |
| ENSG00000256229 | ZNF486 | -0.66 | 0.06 | 1.20E-24 | 1.33E-23 |
| ENSG00000050405 | LIMA1 | -0.66 | 0.07 | 2.43E-22 | 2.48E-21 |
| ENSG00000142627 | EPHA2 | -0.66 | 0.07 | 2.90E-22 | 2.95E-21 |
| ENSG00000182544 | MFSD5 | -0.66 | 0.07 | 2.03E-21 | 2.00E-20 |
| ENSG00000143314 | MRPL24 | -0.66 | 0.07 | 1.38E-19 | 1.26E-18 |
| ENSG00000115977 | AAK1 | -0.66 | 0.07 | 1.67E-19 | 1.52E-18 |
| ENSG00000185298 | CCDC137 | -0.66 | 0.08 | 2.72E-19 | 2.45E-18 |
| ENSG00000135093 | USP30 | -0.66 | 0.08 | 1.03E-18 | 9.03E-18 |
| ENSG00000182179 | UBA7 | -0.66 | 0.08 | 1.71E-18 | 1.48E-17 |
| ENSG00000141560 | FN3KRP | -0.66 | 0.08 | 3.33E-18 | 2.85E-17 |
| ENSG00000130702 | LAMA5 | -0.66 | 0.08 | 7.01E-16 | 5.41E-15 |
| ENSG00000281332 | LINC00997 | -0.66 | 0.09 | 3.89E-15 | 2.90E-14 |
| ENSG00000130299 | GTPBP3 | -0.66 | 0.09 | 1.50E-13 | 1.03E-12 |
| ENSG00000170873 | MTSS1 | -0.66 | 0.09 | 5.49E-13 | 3.66E-12 |
| ENSG00000100263 | RHBDD3 | -0.66 | 0.09 | 5.78E-13 | 3.85E-12 |

| | | | | | |
|---|---|---|---|---|---|
| ENSG00000149346 | SLX4IP | -0.66 | 0.12 | 3.42E-09 | 1.77E-08 |
| ENSG00000118420 | UBE3D | -0.66 | 0.12 | 1.51E-08 | 7.44E-08 |
| ENSG00000265688 | MILIP | -0.66 | 0.13 | 2.94E-08 | 1.42E-07 |
| ENSG00000086717 | PPEF1 | -0.66 | 0.13 | 3.44E-08 | 1.65E-07 |
| ENSG00000136999 | CCN3 | -0.66 | 0.14 | 6.20E-07 | 2.69E-06 |
| ENSG00000225791 | TRAM2-AS1 | -0.66 | 0.15 | 1.90E-06 | 7.87E-06 |
| ENSG00000271936 | | -0.66 | 0.16 | 6.34E-06 | 2.50E-05 |
| ENSG00000168301 | KCTD6 | -0.66 | 0.19 | 8.95E-05 | 3.13E-04 |
| ENSG00000255468 | B4GAT1-DT | -0.66 | 0.21 | 2.79E-04 | 9.17E-04 |
| ENSG00000254536 | | -0.66 | 0.21 | 3.45E-04 | 1.12E-03 |
| ENSG00000099812 | MISP | -0.66 | 0.22 | 4.82E-04 | 1.54E-03 |
| ENSG00000247746 | USP51 | -0.66 | 0.23 | 7.43E-04 | 2.31E-03 |
| ENSG00000174327 | SLC16A13 | -0.66 | 0.23 | 8.50E-04 | 2.61E-03 |
| ENSG00000279456 | | -0.66 | 0.27 | 2.63E-03 | 7.46E-03 |
| ENSG00000188763 | FZD9 | -0.66 | 0.29 | 4.13E-03 | 1.13E-02 |
| ENSG00000169918 | OTUD7A | -0.66 | 0.3 | 4.88E-03 | 1.32E-02 |
| ENSG00000205309 | NT5M | -0.66 | 0.3 | 5.29E-03 | 1.42E-02 |
| ENSG00000130182 | ZSCAN10 | -0.66 | 0.31 | 6.32E-03 | 1.67E-02 |
| ENSG00000260862 | | -0.66 | 0.33 | 8.37E-03 | 2.15E-02 |
| ENSG00000110799 | VWF | -0.66 | 0.34 | 9.92E-03 | 2.51E-02 |
| ENSG00000224452 | RSL24D1P6 | -0.66 | 0.34 | 1.01E-02 | 2.54E-02 |
| ENSG00000255308 | CSRP3-AS1 | -0.66 | 0.76 | 1.42E-02 | 3.46E-02 |
| ENSG00000248626 | GAPDHP40 | -0.66 | 0.75 | 1.48E-02 | 3.60E-02 |
| ENSG00000233198 | RNF224 | -0.66 | 0.4 | 1.76E-02 | 4.19E-02 |
| ENSG00000280149 | | -0.66 | 0.73 | 1.92E-02 | 4.55E-02 |
| ENSG00000149294 | NCAM1 | -0.65 | 0.04 | 5.58E-70 | 1.91E-68 |
| ENSG00000159216 | RUNX1 | -0.65 | 0.04 | 9.79E-49 | 2.13E-47 |
| ENSG00000070601 | FRMPD1 | -0.65 | 0.05 | 1.51E-40 | 2.76E-39 |
| ENSG00000171824 | EXOSC10 | -0.65 | 0.05 | 3.64E-40 | 6.54E-39 |
| ENSG00000119771 | KLHL29 | -0.65 | 0.06 | 1.70E-31 | 2.40E-30 |
| ENSG00000170802 | FOXN2 | -0.65 | 0.06 | 3.14E-31 | 4.41E-30 |
| ENSG00000175183 | CSRP2 | -0.65 | 0.06 | 8.73E-31 | 1.21E-29 |
| ENSG00000204611 | ZNF616 | -0.65 | 0.06 | 4.57E-26 | 5.36E-25 |
| ENSG00000168612 | ZSWIM1 | -0.65 | 0.06 | 3.45E-24 | 3.78E-23 |
| ENSG00000196526 | AFAP1 | -0.65 | 0.07 | 3.64E-23 | 3.83E-22 |
| ENSG00000153814 | JAZF1 | -0.65 | 0.07 | 2.48E-22 | 2.54E-21 |
| ENSG00000152061 | RABGAP1L | -0.65 | 0.07 | 1.06E-19 | 9.71E-19 |
| ENSG00000180098 | TRNAU1AP | -0.65 | 0.07 | 4.71E-19 | 4.19E-18 |
| ENSG00000135709 | KIAA0513 | -0.65 | 0.07 | 6.07E-19 | 5.37E-18 |
| ENSG00000103707 | MTFMT | -0.65 | 0.08 | 1.27E-16 | 1.02E-15 |
| ENSG00000070047 | PHRF1 | -0.65 | 0.08 | 6.55E-16 | 5.07E-15 |
| ENSG00000198924 | DCLRE1A | -0.65 | 0.09 | 1.11E-14 | 8.09E-14 |
| ENSG00000288905 | LSP1P5 | -0.65 | 0.09 | 1.56E-13 | 1.08E-12 |
| ENSG00000121957 | GPSM2 | -0.65 | 0.09 | 8.42E-13 | 5.55E-12 |
| ENSG00000198885 | ITPRIPL1 | -0.65 | 0.1 | 1.98E-12 | 1.28E-11 |
| ENSG00000082126 | MPP4 | -0.65 | 0.1 | 5.21E-12 | 3.27E-11 |
| ENSG00000162645 | GBP2 | -0.65 | 0.11 | 3.14E-10 | 1.75E-09 |
| ENSG00000132967 | HMGB1P5 | -0.65 | 0.11 | 3.70E-09 | 1.91E-08 |
| ENSG00000291130 | | -0.65 | 0.12 | 5.04E-09 | 2.57E-08 |
| ENSG00000083544 | TDRD3 | -0.65 | 0.15 | 3.66E-06 | 1.48E-05 |

| | | | | | |
|---|---|---|---|---|---|
| ENSG00000124802 | EEF1E1 | -0.65 | 0.15 | 3.86E-06 | 1.55E-05 |
| ENSG00000160298 | C21orf58 | -0.65 | 0.15 | 3.91E-06 | 1.57E-05 |
| ENSG00000099957 | P2RX6 | -0.65 | 0.16 | 8.75E-06 | 3.40E-05 |
| ENSG00000272142 | LYRM4-AS1 | -0.65 | 0.18 | 7.02E-05 | 2.49E-04 |
| ENSG00000259205 | PRKXP1 | -0.65 | 0.21 | 4.68E-04 | 1.50E-03 |
| ENSG00000077092 | RARB | -0.65 | 0.23 | 1.17E-03 | 3.51E-03 |
| ENSG00000140807 | NKD1 | -0.65 | 0.25 | 2.11E-03 | 6.09E-03 |
| ENSG00000129170 | CSRP3 | -0.65 | 0.25 | 2.16E-03 | 6.23E-03 |
| ENSG00000099998 | GGT5 | -0.65 | 0.28 | 4.17E-03 | 1.14E-02 |
| ENSG00000182165 | TP53TG1 | -0.65 | 0.29 | 5.10E-03 | 1.37E-02 |
| ENSG00000283439 | SPEM3 | -0.65 | 0.77 | 1.16E-02 | 2.89E-02 |
| ENSG00000197291 | RAMP2-AS1 | -0.65 | 0.35 | 1.22E-02 | 3.02E-02 |
| ENSG00000260269 | | -0.65 | 0.38 | 1.75E-02 | 4.18E-02 |
| ENSG00000279722 | | -0.65 | 0.39 | 1.81E-02 | 4.31E-02 |
| ENSG00000167193 | CRK | -0.64 | 0.04 | 3.56E-63 | 1.06E-61 |
| ENSG00000082898 | XPO1 | -0.64 | 0.04 | 8.93E-62 | 2.58E-60 |
| ENSG00000109971 | HSPA8 | -0.64 | 0.04 | 2.10E-55 | 5.37E-54 |
| ENSG00000108039 | XPNPEP1 | -0.64 | 0.05 | 1.82E-43 | 3.56E-42 |
| ENSG00000140374 | ETFA | -0.64 | 0.05 | 3.96E-38 | 6.82E-37 |
| ENSG00000140262 | TCF12 | -0.64 | 0.05 | 1.70E-36 | 2.78E-35 |
| ENSG00000107443 | CCNJ | -0.64 | 0.06 | 5.24E-30 | 7.04E-29 |
| ENSG00000157538 | VPS26C | -0.64 | 0.06 | 5.77E-28 | 7.21E-27 |
| ENSG00000124496 | TRERF1 | -0.64 | 0.06 | 1.32E-25 | 1.53E-24 |
| ENSG00000170425 | ADORA2B | -0.64 | 0.06 | 4.55E-25 | 5.14E-24 |
| ENSG00000149930 | TAOK2 | -0.64 | 0.06 | 1.80E-24 | 1.99E-23 |
| ENSG00000051825 | MPHOSPH9 | -0.64 | 0.07 | 2.62E-23 | 2.78E-22 |
| ENSG00000126775 | ATG14 | -0.64 | 0.07 | 8.39E-23 | 8.71E-22 |
| ENSG00000122707 | RECK | -0.64 | 0.07 | 9.75E-23 | 1.01E-21 |
| ENSG00000130023 | ERMARD | -0.64 | 0.08 | 3.50E-18 | 2.99E-17 |
| ENSG00000075891 | PAX2 | -0.64 | 0.08 | 1.03E-16 | 8.28E-16 |
| ENSG00000101638 | ST8SIA5 | -0.64 | 0.08 | 2.85E-16 | 2.25E-15 |
| ENSG00000081138 | CDH7 | -0.64 | 0.08 | 2.27E-15 | 1.72E-14 |
| ENSG00000176834 | VSIG10 | -0.64 | 0.08 | 2.66E-15 | 2.00E-14 |
| ENSG00000125863 | MKKS | -0.64 | 0.08 | 3.11E-15 | 2.33E-14 |
| ENSG00000143786 | CNIH3 | -0.64 | 0.09 | 6.58E-14 | 4.62E-13 |
| ENSG00000155657 | TTN | -0.64 | 0.09 | 5.90E-13 | 3.92E-12 |
| ENSG00000119699 | TGFB3 | -0.64 | 0.1 | 3.33E-11 | 1.99E-10 |
| ENSG00000213020 | ZNF611 | -0.64 | 0.1 | 7.52E-11 | 4.39E-10 |
| ENSG00000106689 | LHX2 | -0.64 | 0.11 | 4.47E-10 | 2.47E-09 |
| ENSG00000099203 | TMED1 | -0.64 | 0.12 | 1.27E-08 | 6.32E-08 |
| ENSG00000131944 | FAAP24 | -0.64 | 0.12 | 1.82E-08 | 8.94E-08 |
| ENSG00000110801 | PSMD9 | -0.64 | 0.12 | 5.25E-08 | 2.49E-07 |
| ENSG00000164188 | RANBP3L | -0.64 | 0.13 | 1.58E-07 | 7.20E-07 |
| ENSG00000106211 | HSPB1 | -0.64 | 0.14 | 8.01E-07 | 3.44E-06 |
| ENSG00000291029 | FKBP9P1 | -0.64 | 0.15 | 8.15E-06 | 3.18E-05 |
| ENSG00000163689 | CFAP20DC | -0.64 | 0.24 | 1.56E-03 | 4.59E-03 |
| ENSG00000123297 | TSFM | -0.64 | 0.29 | 6.01E-03 | 1.59E-02 |
| ENSG00000185432 | METTL7A | -0.64 | 0.31 | 9.25E-03 | 2.36E-02 |
| ENSG00000109956 | B3GAT1 | -0.64 | 0.75 | 1.61E-02 | 3.88E-02 |
| ENSG00000138161 | CUZD1 | -0.64 | 0.36 | 1.63E-02 | 3.91E-02 |

| | | | | | |
|---|---|---|---|---|---|
| ENSG00000165181 | SHOC1 | -0.64 | 0.38 | 1.94E-02 | 4.58E-02 |
| ENSG00000272417 | | -0.64 | 0.73 | 2.06E-02 | 4.84E-02 |
| ENSG00000135905 | DOCK10 | -0.63 | 0.04 | 3.31E-50 | 7.47E-49 |
| ENSG00000124193 | SRSF6 | -0.63 | 0.04 | 1.37E-49 | 3.03E-48 |
| ENSG00000162377 | COA7 | -0.63 | 0.04 | 8.22E-48 | 1.76E-46 |
| ENSG00000145191 | EIF2B5 | -0.63 | 0.05 | 1.03E-42 | 1.98E-41 |
| ENSG00000147010 | SH3KBP1 | -0.63 | 0.05 | 5.08E-39 | 8.91E-38 |
| ENSG00000151176 | PLBD2 | -0.63 | 0.05 | 1.15E-34 | 1.78E-33 |
| ENSG00000225778 | PROSER2-AS1 | -0.63 | 0.05 | 6.64E-34 | 1.00E-32 |
| ENSG00000009844 | VTA1 | -0.63 | 0.05 | 1.47E-32 | 2.13E-31 |
| ENSG00000114850 | SSR3 | -0.63 | 0.05 | 4.15E-31 | 5.81E-30 |
| ENSG00000143653 | SCCPDH | -0.63 | 0.05 | 5.05E-31 | 7.04E-30 |
| ENSG00000136044 | APPL2 | -0.63 | 0.06 | 2.50E-30 | 3.40E-29 |
| ENSG00000168393 | DTYMK | -0.63 | 0.06 | 7.54E-30 | 1.01E-28 |
| ENSG00000176887 | SOX11 | -0.63 | 0.06 | 3.52E-28 | 4.42E-27 |
| ENSG00000108175 | ZMIZ1 | -0.63 | 0.06 | 1.08E-27 | 1.35E-26 |
| ENSG00000178295 | GEN1 | -0.63 | 0.06 | 2.76E-25 | 3.15E-24 |
| ENSG00000112592 | TBP | -0.63 | 0.06 | 8.17E-24 | 8.83E-23 |
| ENSG00000142453 | CARM1 | -0.63 | 0.07 | 1.71E-22 | 1.75E-21 |
| ENSG00000137628 | DDX60 | -0.63 | 0.07 | 1.90E-22 | 1.95E-21 |
| ENSG00000212719 | LINC02693 | -0.63 | 0.07 | 2.10E-21 | 2.06E-20 |
| ENSG00000019144 | PHLDB1 | -0.63 | 0.07 | 3.36E-21 | 3.29E-20 |
| ENSG00000151806 | GUF1 | -0.63 | 0.07 | 4.58E-21 | 4.46E-20 |
| ENSG00000172985 | SH3RF3 | -0.63 | 0.07 | 1.10E-18 | 9.65E-18 |
| ENSG00000015532 | XYLT2 | -0.63 | 0.08 | 1.83E-16 | 1.46E-15 |
| ENSG00000215305 | VPS16 | -0.63 | 0.08 | 1.91E-16 | 1.51E-15 |
| ENSG00000136830 | NIBAN2 | -0.63 | 0.08 | 3.82E-16 | 2.99E-15 |
| ENSG00000189046 | ALKBH2 | -0.63 | 0.08 | 4.53E-16 | 3.53E-15 |
| ENSG00000198538 | ZNF28 | -0.63 | 0.08 | 5.59E-16 | 4.34E-15 |
| ENSG00000174720 | LARP7 | -0.63 | 0.09 | 1.03E-13 | 7.14E-13 |
| ENSG00000103852 | TTC23 | -0.63 | 0.09 | 2.70E-12 | 1.73E-11 |
| ENSG00000188312 | CENPP | -0.63 | 0.11 | 1.29E-09 | 6.91E-09 |
| ENSG00000133321 | PLAAT4 | -0.63 | 0.11 | 2.67E-09 | 1.40E-08 |
| ENSG00000177873 | ZNF619 | -0.63 | 0.13 | 1.08E-07 | 5.00E-07 |
| ENSG00000105894 | PTN | -0.63 | 0.13 | 1.98E-07 | 8.99E-07 |
| ENSG00000049246 | PER3 | -0.63 | 0.15 | 7.68E-06 | 3.00E-05 |
| ENSG00000257859 | CASC18 | -0.63 | 0.24 | 1.93E-03 | 5.60E-03 |
| ENSG00000261616 | TTC23-AS1 | -0.63 | 0.29 | 7.32E-03 | 1.91E-02 |
| ENSG00000229474 | PATL2 | -0.63 | 0.35 | 1.50E-02 | 3.63E-02 |
| ENSG00000166278 | C2 | -0.63 | 0.37 | 1.88E-02 | 4.46E-02 |
| ENSG00000138435 | CHRNA1 | -0.62 | 0.04 | 6.55E-67 | 2.10E-65 |
| ENSG00000135074 | ADAM19 | -0.62 | 0.04 | 7.45E-49 | 1.63E-47 |
| ENSG00000153214 | TMEM87B | -0.62 | 0.04 | 4.26E-45 | 8.59E-44 |
| ENSG00000164904 | ALDH7A1 | -0.62 | 0.05 | 3.41E-41 | 6.34E-40 |
| ENSG00000153310 | CYRIB | -0.62 | 0.05 | 4.89E-39 | 8.60E-38 |
| ENSG00000198176 | TFDP1 | -0.62 | 0.05 | 1.66E-36 | 2.72E-35 |
| ENSG00000113569 | NUP155 | -0.62 | 0.05 | 2.13E-36 | 3.48E-35 |
| ENSG00000117899 | MESD | -0.62 | 0.05 | 3.00E-36 | 4.88E-35 |
| ENSG00000213965 | NUDT19 | -0.62 | 0.05 | 4.49E-36 | 7.28E-35 |
| ENSG00000125445 | MRPS7 | -0.62 | 0.05 | 1.10E-35 | 1.77E-34 |

| | | | | | |
|---|---|---|---|---|---|
| ENSG00000109171 | SLAIN2 | -0.62 | 0.05 | 2.57E-35 | 4.08E-34 |
| ENSG00000140263 | SORD | -0.62 | 0.05 | 2.97E-35 | 4.70E-34 |
| ENSG00000074603 | DPP8 | -0.62 | 0.06 | 2.54E-30 | 3.45E-29 |
| ENSG00000124532 | MRS2 | -0.62 | 0.06 | 2.96E-28 | 3.73E-27 |
| ENSG00000033867 | SLC4A7 | -0.62 | 0.06 | 3.45E-25 | 3.91E-24 |
| ENSG00000114686 | MRPL3 | -0.62 | 0.06 | 1.53E-24 | 1.69E-23 |
| ENSG00000120949 | TNFRSF8 | -0.62 | 0.06 | 3.41E-24 | 3.73E-23 |
| ENSG00000171723 | GPHN | -0.62 | 0.06 | 1.62E-22 | 1.66E-21 |
| ENSG00000070214 | SLC44A1 | -0.62 | 0.07 | 3.56E-19 | 3.19E-18 |
| ENSG00000075188 | NUP37 | -0.62 | 0.07 | 5.70E-19 | 5.04E-18 |
| ENSG00000180185 | FAHD1 | -0.62 | 0.08 | 6.03E-15 | 4.47E-14 |
| ENSG00000088035 | ALG6 | -0.62 | 0.08 | 1.13E-14 | 8.24E-14 |
| ENSG00000235173 | HGH1 | -0.62 | 0.08 | 3.27E-14 | 2.34E-13 |
| ENSG00000116205 | TCEANC2 | -0.62 | 0.09 | 3.61E-13 | 2.43E-12 |
| ENSG00000101417 | PXMP4 | -0.62 | 0.09 | 4.17E-12 | 2.64E-11 |
| ENSG00000163344 | PMVK | -0.62 | 0.1 | 1.05E-10 | 6.10E-10 |
| ENSG00000185875 | THNSL1 | -0.62 | 0.1 | 2.45E-10 | 1.38E-09 |
| ENSG00000204536 | CCHCR1 | -0.62 | 0.11 | 1.20E-09 | 6.45E-09 |
| ENSG00000261150 | EPPK1 | -0.62 | 0.12 | 8.32E-08 | 3.88E-07 |
| ENSG00000156050 | FAM161B | -0.62 | 0.12 | 1.14E-07 | 5.26E-07 |
| ENSG00000165275 | TRMT10B | -0.62 | 0.12 | 1.20E-07 | 5.53E-07 |
| ENSG00000120662 | MTRF1 | -0.62 | 0.14 | 3.70E-06 | 1.49E-05 |
| ENSG00000064199 | SPA17 | -0.62 | 0.16 | 1.39E-05 | 5.30E-05 |
| ENSG00000081189 | MEF2C | -0.62 | 0.18 | 1.74E-04 | 5.87E-04 |
| ENSG00000254064 | PIWIL2-DT | -0.62 | 0.19 | 2.35E-04 | 7.81E-04 |
| ENSG00000253106 | | -0.62 | 0.22 | 1.14E-03 | 3.44E-03 |
| ENSG00000248866 | USP46-DT | -0.62 | 0.24 | 2.50E-03 | 7.13E-03 |
| ENSG00000237181 | PRKAR1B-AS1 | -0.62 | 0.26 | 4.27E-03 | 1.16E-02 |
| ENSG00000160408 | ST6GALNAC6 | -0.62 | 0.26 | 4.32E-03 | 1.17E-02 |
| ENSG00000157613 | CREB3L1 | -0.62 | 0.28 | 6.41E-03 | 1.69E-02 |
| ENSG00000181433 | SAGE1 | -0.62 | 0.29 | 7.16E-03 | 1.87E-02 |
| ENSG00000246308 | | -0.62 | 0.29 | 7.85E-03 | 2.03E-02 |
| ENSG00000113231 | PDE8B | -0.62 | 0.32 | 1.13E-02 | 2.82E-02 |
| ENSG00000236819 | LINC01563 | -0.62 | 0.77 | 1.27E-02 | 3.12E-02 |
| ENSG00000207805 | MIR483 | -0.62 | 0.34 | 1.50E-02 | 3.63E-02 |
| ENSG00000108861 | DUSP3 | -0.61 | 0.04 | 1.01E-43 | 1.99E-42 |
| ENSG00000130816 | DNMT1 | -0.61 | 0.05 | 5.03E-42 | 9.58E-41 |
| ENSG00000110841 | PPFIBP1 | -0.61 | 0.05 | 2.89E-38 | 4.99E-37 |
| ENSG00000197077 | KIAA1671 | -0.61 | 0.05 | 1.77E-35 | 2.83E-34 |
| ENSG00000107554 | DNMBP | -0.61 | 0.05 | 5.25E-33 | 7.72E-32 |
| ENSG00000112319 | EYA4 | -0.61 | 0.05 | 4.20E-31 | 5.88E-30 |
| ENSG00000135486 | HNRNPA1 | -0.61 | 0.05 | 1.96E-30 | 2.68E-29 |
| ENSG00000091656 | ZFHX4 | -0.61 | 0.05 | 3.36E-30 | 4.55E-29 |
| ENSG00000170571 | EMB | -0.61 | 0.05 | 1.16E-29 | 1.54E-28 |
| ENSG00000100714 | MTHFD1 | -0.61 | 0.05 | 2.54E-29 | 3.33E-28 |
| ENSG00000120868 | APAF1 | -0.61 | 0.06 | 4.40E-29 | 5.74E-28 |
| ENSG00000104356 | POP1 | -0.61 | 0.06 | 7.32E-29 | 9.47E-28 |
| ENSG00000114745 | GORASP1 | -0.61 | 0.06 | 1.53E-28 | 1.94E-27 |
| ENSG00000065882 | TBC1D1 | -0.61 | 0.06 | 7.42E-27 | 8.95E-26 |
| ENSG00000174705 | SH3PXD2B | -0.61 | 0.06 | 1.50E-26 | 1.80E-25 |

| | | | | | |
|---|---|---|---|---|---|
| ENSG00000100034 | PPM1F | -0.61 | 0.06 | 1.86E-26 | 2.21E-25 |
| ENSG00000159593 | NAE1 | -0.61 | 0.06 | 1.74E-25 | 1.99E-24 |
| ENSG00000182492 | BGN | -0.61 | 0.06 | 9.75E-25 | 1.09E-23 |
| ENSG00000087266 | SH3BP2 | -0.61 | 0.06 | 1.55E-24 | 1.72E-23 |
| ENSG00000135956 | TMEM127 | -0.61 | 0.06 | 1.81E-24 | 2.00E-23 |
| ENSG00000196576 | PLXNB2 | -0.61 | 0.06 | 4.81E-24 | 5.24E-23 |
| ENSG00000158321 | AUTS2 | -0.61 | 0.06 | 1.73E-23 | 1.84E-22 |
| ENSG00000165801 | ARHGEF40 | -0.61 | 0.07 | 5.94E-21 | 5.75E-20 |
| ENSG00000118246 | FASTKD2 | -0.61 | 0.07 | 1.82E-19 | 1.65E-18 |
| ENSG00000255529 | POLR2M | -0.61 | 0.07 | 3.24E-19 | 2.91E-18 |
| ENSG00000122484 | RPAP2 | -0.61 | 0.07 | 2.34E-18 | 2.02E-17 |
| ENSG00000166165 | CKB | -0.61 | 0.08 | 3.16E-16 | 2.48E-15 |
| ENSG00000158092 | NCK1 | -0.61 | 0.08 | 7.20E-16 | 5.56E-15 |
| ENSG00000155287 | SLC25A28 | -0.61 | 0.08 | 4.29E-15 | 3.20E-14 |
| ENSG00000136367 | ZFHX2 | -0.61 | 0.08 | 4.34E-14 | 3.08E-13 |
| ENSG00000129474 | AJUBA | -0.61 | 0.08 | 1.36E-13 | 9.38E-13 |
| ENSG00000101247 | NDUFAF5 | -0.61 | 0.09 | 9.98E-13 | 6.55E-12 |
| ENSG00000183647 | ZNF530 | -0.61 | 0.1 | 1.37E-10 | 7.88E-10 |
| ENSG00000182175 | RGMA | -0.61 | 0.1 | 1.69E-10 | 9.66E-10 |
| ENSG00000196459 | TRAPPC2 | -0.61 | 0.1 | 4.51E-10 | 2.50E-09 |
| ENSG00000182742 | HOXB4 | -0.61 | 0.1 | 8.99E-10 | 4.87E-09 |
| ENSG00000254585 | MAGEL2 | -0.61 | 0.11 | 1.29E-08 | 6.42E-08 |
| ENSG00000170017 | ALCAM | -0.61 | 0.12 | 3.62E-08 | 1.73E-07 |
| ENSG00000240891 | PLCXD2 | -0.61 | 0.12 | 2.46E-07 | 1.11E-06 |
| ENSG00000197472 | ZNF695 | -0.61 | 0.15 | 1.35E-05 | 5.15E-05 |
| ENSG00000156218 | ADAMTSL3 | -0.61 | 0.17 | 8.09E-05 | 2.85E-04 |
| ENSG00000137474 | MYO7A | -0.61 | 0.2 | 4.66E-04 | 1.49E-03 |
| ENSG00000184785 | SMIM10 | -0.61 | 0.25 | 3.91E-03 | 1.07E-02 |
| ENSG00000184574 | LPAR5 | -0.61 | 0.26 | 4.64E-03 | 1.26E-02 |
| ENSG00000225930 | LINC02249 | -0.61 | 0.31 | 1.15E-02 | 2.87E-02 |
| ENSG00000187123 | LYPD6 | -0.61 | 0.33 | 1.59E-02 | 3.83E-02 |
| ENSG00000181790 | ADGRB1 | -0.61 | 0.35 | 1.95E-02 | 4.61E-02 |
| ENSG00000152127 | MGAT5 | -0.6 | 0.03 | 5.98E-75 | 2.26E-73 |
| ENSG00000114491 | UMPS | -0.6 | 0.05 | 1.47E-37 | 2.48E-36 |
| ENSG00000129351 | ILF3 | -0.6 | 0.05 | 3.47E-32 | 5.01E-31 |
| ENSG00000196547 | MAN2A2 | -0.6 | 0.05 | 3.40E-29 | 4.45E-28 |
| ENSG00000169764 | UGP2 | -0.6 | 0.06 | 2.10E-26 | 2.50E-25 |
| ENSG00000183723 | CMTM4 | -0.6 | 0.06 | 5.64E-22 | 5.68E-21 |
| ENSG00000204394 | VARS1 | -0.6 | 0.07 | 6.77E-21 | 6.52E-20 |
| ENSG00000087008 | ACOX3 | -0.6 | 0.07 | 1.82E-20 | 1.73E-19 |
| ENSG00000184752 | NDUFA12 | -0.6 | 0.07 | 1.19E-19 | 1.09E-18 |
| ENSG00000169105 | CHST14 | -0.6 | 0.07 | 1.38E-17 | 1.15E-16 |
| ENSG00000078403 | MLLT10 | -0.6 | 0.07 | 7.94E-17 | 6.43E-16 |
| ENSG00000120093 | HOXB3 | -0.6 | 0.07 | 1.91E-16 | 1.51E-15 |
| ENSG00000176623 | RMDN1 | -0.6 | 0.08 | 7.60E-16 | 5.84E-15 |
| ENSG00000179604 | CDC42EP4 | -0.6 | 0.08 | 1.89E-15 | 1.43E-14 |
| ENSG00000152240 | HAUS1 | -0.6 | 0.08 | 2.43E-15 | 1.83E-14 |
| ENSG00000221829 | FANCG | -0.6 | 0.08 | 6.89E-14 | 4.83E-13 |
| ENSG00000125846 | ZNF133 | -0.6 | 0.09 | 3.54E-12 | 2.25E-11 |
| ENSG00000165917 | RAPSN | -0.6 | 0.09 | 3.88E-12 | 2.46E-11 |

| | | | | | |
|---|---|---|---|---|---|
| ENSG00000102935 | ZNF423 | -0.6 | 0.09 | 8.93E-12 | 5.53E-11 |
| ENSG00000106268 | NUDT1 | -0.6 | 0.1 | 8.72E-11 | 5.08E-10 |
| ENSG00000076555 | ACACB | -0.6 | 0.1 | 8.79E-11 | 5.11E-10 |
| ENSG00000117228 | GBP1 | -0.6 | 0.1 | 1.48E-10 | 8.49E-10 |
| ENSG00000151725 | CENPU | -0.6 | 0.1 | 2.24E-10 | 1.27E-09 |
| ENSG00000154721 | JAM2 | -0.6 | 0.1 | 3.89E-10 | 2.16E-09 |
| ENSG00000151849 | CENPJ | -0.6 | 0.1 | 1.98E-09 | 1.05E-08 |
| ENSG00000070404 | FSTL3 | -0.6 | 0.11 | 6.40E-09 | 3.25E-08 |
| ENSG00000047648 | ARHGAP6 | -0.6 | 0.11 | 1.70E-08 | 8.38E-08 |
| ENSG00000212123 | PRR22 | -0.6 | 0.11 | 3.40E-08 | 1.63E-07 |
| ENSG00000186615 | KTN1-AS1 | -0.6 | 0.12 | 9.38E-08 | 4.36E-07 |
| ENSG00000176244 | ACBD7 | -0.6 | 0.14 | 6.47E-06 | 2.55E-05 |
| ENSG00000187486 | KCNJ11 | -0.6 | 0.14 | 6.92E-06 | 2.72E-05 |
| ENSG00000132639 | SNAP25 | -0.6 | 0.15 | 1.45E-05 | 5.53E-05 |
| ENSG00000227252 | COPS8-DT | -0.6 | 0.16 | 3.48E-05 | 1.27E-04 |
| ENSG00000102879 | CORO1A | -0.6 | 0.17 | 9.19E-05 | 3.21E-04 |
| ENSG00000231680 | LINC02723 | -0.6 | 0.17 | 1.32E-04 | 4.52E-04 |
| ENSG00000079337 | RAPGEF3 | -0.6 | 0.2 | 5.62E-04 | 1.78E-03 |
| ENSG00000177335 | LINC02904 | -0.6 | 0.2 | 6.69E-04 | 2.09E-03 |
| ENSG00000229140 | CCDC26 | -0.6 | 0.22 | 1.98E-03 | 5.73E-03 |
| ENSG00000267325 | LINC01415 | -0.6 | 0.23 | 2.47E-03 | 7.05E-03 |
| ENSG00000185010 | F8 | -0.6 | 0.25 | 4.44E-03 | 1.21E-02 |
| ENSG00000113205 | PCDHB3 | -0.6 | 0.25 | 4.83E-03 | 1.30E-02 |
| ENSG00000187595 | ZNF385C | -0.6 | 0.28 | 8.01E-03 | 2.07E-02 |
| ENSG00000146054 | TRIM7 | -0.6 | 0.29 | 8.93E-03 | 2.28E-02 |
| ENSG00000007952 | NOX1 | -0.6 | 0.3 | 1.17E-02 | 2.90E-02 |
| ENSG00000091536 | MYO15A | -0.6 | 0.3 | 1.19E-02 | 2.95E-02 |
| ENSG00000132623 | ANKEF1 | -0.6 | 0.3 | 1.23E-02 | 3.04E-02 |
| ENSG00000271414 | | -0.6 | 0.76 | 1.51E-02 | 3.67E-02 |
| ENSG00000252992 | | -0.6 | 0.75 | 1.58E-02 | 3.80E-02 |
| ENSG00000153936 | HS2ST1 | -0.59 | 0.04 | 2.15E-47 | 4.56E-46 |
| ENSG00000086015 | MAST2 | -0.59 | 0.04 | 5.77E-45 | 1.16E-43 |
| ENSG00000088833 | NSFL1C | -0.59 | 0.04 | 8.44E-41 | 1.55E-39 |
| ENSG00000198663 | C6orf89 | -0.59 | 0.04 | 1.14E-40 | 2.08E-39 |
| ENSG00000183283 | DAZAP2 | -0.59 | 0.04 | 1.99E-40 | 3.61E-39 |
| ENSG00000071575 | TRIB2 | -0.59 | 0.05 | 3.80E-37 | 6.33E-36 |
| ENSG00000145782 | ATG12 | -0.59 | 0.05 | 6.22E-31 | 8.65E-30 |
| ENSG00000140382 | HMG20A | -0.59 | 0.05 | 3.81E-30 | 5.14E-29 |
| ENSG00000131844 | MCCC2 | -0.59 | 0.05 | 1.10E-29 | 1.47E-28 |
| ENSG00000004864 | SLC25A13 | -0.59 | 0.05 | 2.23E-29 | 2.94E-28 |
| ENSG00000101452 | DHX35 | -0.59 | 0.06 | 9.61E-27 | 1.16E-25 |
| ENSG00000163320 | CGGBP1 | -0.59 | 0.06 | 2.84E-26 | 3.35E-25 |
| ENSG00000132155 | RAF1 | -0.59 | 0.06 | 2.07E-25 | 2.37E-24 |
| ENSG00000110047 | EHD1 | -0.59 | 0.06 | 1.18E-20 | 1.12E-19 |
| ENSG00000156990 | RPUSD3 | -0.59 | 0.06 | 4.26E-20 | 3.97E-19 |
| ENSG00000119927 | GPAM | -0.59 | 0.07 | 4.66E-19 | 4.15E-18 |
| ENSG00000118418 | HMGN3 | -0.59 | 0.07 | 8.19E-18 | 6.91E-17 |
| ENSG00000125962 | ARMCX5 | -0.59 | 0.07 | 1.00E-16 | 8.08E-16 |
| ENSG00000133119 | RFC3 | -0.59 | 0.08 | 3.26E-15 | 2.45E-14 |
| ENSG00000171033 | PKIA | -0.59 | 0.08 | 1.08E-14 | 7.87E-14 |

| | | | | | |
|---|---|---|---|---|---|
| ENSG00000115548 | KDM3A | -0.59 | 0.08 | 1.69E-14 | 1.23E-13 |
| ENSG00000056277 | ZNF280C | -0.59 | 0.08 | 5.94E-14 | 4.18E-13 |
| ENSG00000105750 | ZNF85 | -0.59 | 0.08 | 4.63E-13 | 3.10E-12 |
| ENSG00000258634 | | -0.59 | 0.09 | 1.57E-12 | 1.02E-11 |
| ENSG00000123407 | HOXC12 | -0.59 | 0.09 | 3.95E-12 | 2.50E-11 |
| ENSG00000205413 | SAMD9 | -0.59 | 0.09 | 2.13E-11 | 1.29E-10 |
| ENSG00000028116 | VRK2 | -0.59 | 0.09 | 2.52E-11 | 1.52E-10 |
| ENSG00000137404 | NRM | -0.59 | 0.09 | 3.10E-11 | 1.86E-10 |
| ENSG00000203875 | SNHG5 | -0.59 | 0.1 | 1.45E-09 | 7.73E-09 |
| ENSG00000163755 | HPS3 | -0.59 | 0.1 | 1.66E-09 | 8.83E-09 |
| ENSG00000090447 | TFAP4 | -0.59 | 0.1 | 2.70E-09 | 1.41E-08 |
| ENSG00000137872 | SEMA6D | -0.59 | 0.11 | 4.17E-09 | 2.14E-08 |
| ENSG00000128394 | APOBEC3F | -0.59 | 0.11 | 7.67E-09 | 3.87E-08 |
| ENSG00000189114 | BLOC1S3 | -0.59 | 0.11 | 3.10E-08 | 1.49E-07 |
| ENSG00000144821 | MYH15 | -0.59 | 0.11 | 3.84E-08 | 1.84E-07 |
| ENSG00000125247 | TMTC4 | -0.59 | 0.12 | 1.11E-07 | 5.14E-07 |
| ENSG00000164089 | ETNPPL | -0.59 | 0.12 | 1.31E-07 | 6.02E-07 |
| ENSG00000126749 | EMG1 | -0.59 | 0.12 | 3.09E-07 | 1.38E-06 |
| ENSG00000033170 | FUT8 | -0.59 | 0.16 | 4.42E-05 | 1.60E-04 |
| ENSG00000184185 | KCNJ12 | -0.59 | 0.22 | 1.72E-03 | 5.04E-03 |
| ENSG00000185261 | KIAA0825 | -0.59 | 0.22 | 2.43E-03 | 6.95E-03 |
| ENSG00000081059 | TCF7 | -0.59 | 0.24 | 3.71E-03 | 1.02E-02 |
| ENSG00000250508 | LINC02701 | -0.59 | 0.24 | 3.87E-03 | 1.06E-02 |
| ENSG00000275401 | | -0.59 | 0.31 | 1.34E-02 | 3.29E-02 |
| ENSG00000227479 | | -0.59 | 0.74 | 2.10E-02 | 4.92E-02 |
| ENSG00000099901 | RANBP1 | -0.58 | 0.04 | 4.77E-49 | 1.05E-47 |
| ENSG00000166747 | AP1G1 | -0.58 | 0.04 | 3.76E-40 | 6.76E-39 |
| ENSG00000135040 | NAA35 | -0.58 | 0.05 | 8.28E-30 | 1.11E-28 |
| ENSG00000116459 | ATP5PB | -0.58 | 0.05 | 3.39E-29 | 4.43E-28 |
| ENSG00000109917 | ZPR1 | -0.58 | 0.05 | 6.66E-29 | 8.64E-28 |
| ENSG00000137168 | PPIL1 | -0.58 | 0.05 | 1.11E-28 | 1.42E-27 |
| ENSG00000113407 | TARS1 | -0.58 | 0.05 | 2.00E-28 | 2.53E-27 |
| ENSG00000137563 | GGH | -0.58 | 0.05 | 2.50E-28 | 3.16E-27 |
| ENSG00000162607 | USP1 | -0.58 | 0.06 | 1.48E-25 | 1.70E-24 |
| ENSG00000179456 | ZBTB18 | -0.58 | 0.06 | 1.22E-24 | 1.36E-23 |
| ENSG00000167394 | ZNF668 | -0.58 | 0.06 | 1.72E-23 | 1.83E-22 |
| ENSG00000005059 | MCUB | -0.58 | 0.06 | 6.20E-21 | 5.99E-20 |
| ENSG00000120156 | TEK | -0.58 | 0.06 | 9.48E-20 | 8.72E-19 |
| ENSG00000213024 | NUP62 | -0.58 | 0.07 | 1.52E-18 | 1.32E-17 |
| ENSG00000116353 | MECR | -0.58 | 0.07 | 1.87E-18 | 1.62E-17 |
| ENSG00000188643 | S100A16 | -0.58 | 0.07 | 2.57E-18 | 2.21E-17 |
| ENSG00000103485 | QPRT | -0.58 | 0.07 | 3.86E-18 | 3.30E-17 |
| ENSG00000124688 | MAD2L1BP | -0.58 | 0.07 | 1.07E-17 | 8.98E-17 |
| ENSG00000135913 | USP37 | -0.58 | 0.07 | 4.94E-17 | 4.04E-16 |
| ENSG00000173442 | EHBP1L1 | -0.58 | 0.07 | 6.95E-17 | 5.64E-16 |
| ENSG00000065320 | NTN1 | -0.58 | 0.07 | 9.05E-16 | 6.93E-15 |
| ENSG00000128973 | CLN6 | -0.58 | 0.08 | 9.28E-15 | 6.81E-14 |
| ENSG00000184675 | AMER1 | -0.58 | 0.08 | 3.98E-14 | 2.82E-13 |
| ENSG00000168427 | KLHL30 | -0.58 | 0.08 | 2.67E-13 | 1.81E-12 |
| ENSG00000142961 | MOB3C | -0.58 | 0.08 | 1.43E-12 | 9.27E-12 |

| | | | | | |
|---|---|---|---|---|---|
| ENSG00000167549 | CORO6 | -0.58 | 0.09 | 2.57E-12 | 1.64E-11 |
| ENSG00000110080 | ST3GAL4 | -0.58 | 0.09 | 4.61E-12 | 2.91E-11 |
| ENSG00000142794 | NBPF3 | -0.58 | 0.09 | 4.47E-11 | 2.65E-10 |
| ENSG00000161714 | PLCD3 | -0.58 | 0.09 | 5.27E-11 | 3.11E-10 |
| ENSG00000023608 | SNAPC1 | -0.58 | 0.09 | 1.13E-10 | 6.54E-10 |
| ENSG00000246763 | RGMB-AS1 | -0.58 | 0.09 | 2.24E-10 | 1.27E-09 |
| ENSG00000178726 | THBD | -0.58 | 0.1 | 4.38E-09 | 2.25E-08 |
| ENSG00000067840 | PDZD4 | -0.58 | 0.11 | 1.10E-08 | 5.52E-08 |
| ENSG00000171016 | PYGO1 | -0.58 | 0.11 | 2.02E-08 | 9.90E-08 |
| ENSG00000164048 | ZNF589 | -0.58 | 0.12 | 3.53E-07 | 1.57E-06 |
| ENSG00000184898 | RBM43 | -0.58 | 0.12 | 3.55E-07 | 1.58E-06 |
| ENSG00000066056 | TIE1 | -0.58 | 0.12 | 3.67E-07 | 1.63E-06 |
| ENSG00000166352 | IFTAP | -0.58 | 0.13 | 1.67E-06 | 6.95E-06 |
| ENSG00000164855 | TMEM184A | -0.58 | 0.13 | 3.40E-06 | 1.37E-05 |
| ENSG00000260588 | | -0.58 | 0.15 | 2.37E-05 | 8.81E-05 |
| ENSG00000106992 | AK1 | -0.58 | 0.15 | 3.97E-05 | 1.44E-04 |
| ENSG00000100078 | PLA2G3 | -0.58 | 0.16 | 4.94E-05 | 1.78E-04 |
| ENSG00000041353 | RAB27B | -0.58 | 0.16 | 8.25E-05 | 2.90E-04 |
| ENSG00000285816 | | -0.58 | 0.18 | 3.31E-04 | 1.08E-03 |
| ENSG00000159208 | CIART | -0.58 | 0.19 | 6.65E-04 | 2.08E-03 |
| ENSG00000241547 | ACTG1P20 | -0.58 | 0.21 | 1.56E-03 | 4.60E-03 |
| ENSG00000079691 | CARMIL1 | -0.58 | 0.23 | 3.21E-03 | 8.97E-03 |
| ENSG00000259673 | IQCH-AS1 | -0.58 | 0.24 | 4.00E-03 | 1.10E-02 |
| ENSG00000248538 | PPP1R3B-DT | -0.58 | 0.27 | 8.84E-03 | 2.26E-02 |
| ENSG00000180884 | ZNF792 | -0.58 | 0.27 | 9.80E-03 | 2.48E-02 |
| ENSG00000290571 | GOLGA2P5 | -0.58 | 0.29 | 1.11E-02 | 2.77E-02 |
| ENSG00000163235 | TGFA | -0.58 | 0.29 | 1.22E-02 | 3.01E-02 |
| ENSG00000248969 | | -0.58 | 0.76 | 1.47E-02 | 3.57E-02 |
| ENSG00000259294 | | -0.58 | 0.76 | 1.52E-02 | 3.67E-02 |
| ENSG00000154099 | DNAAF1 | -0.58 | 0.31 | 1.64E-02 | 3.92E-02 |
| ENSG00000145794 | MEGF10 | -0.57 | 0.03 | 1.57E-82 | 6.81E-81 |
| ENSG00000104738 | MCM4 | -0.57 | 0.04 | 2.73E-49 | 6.01E-48 |
| ENSG00000122705 | CLTA | -0.57 | 0.04 | 3.40E-42 | 6.50E-41 |
| ENSG00000182944 | EWSR1 | -0.57 | 0.04 | 1.05E-41 | 1.99E-40 |
| ENSG00000091483 | FH | -0.57 | 0.04 | 1.89E-37 | 3.18E-36 |
| ENSG00000113282 | CLINT1 | -0.57 | 0.05 | 2.75E-35 | 4.35E-34 |
| ENSG00000143515 | ATP8B2 | -0.57 | 0.05 | 1.29E-34 | 2.00E-33 |
| ENSG00000151693 | ASAP2 | -0.57 | 0.05 | 1.26E-32 | 1.84E-31 |
| ENSG00000122741 | DCAF10 | -0.57 | 0.05 | 2.58E-28 | 3.26E-27 |
| ENSG00000141568 | FOXK2 | -0.57 | 0.05 | 5.80E-28 | 7.24E-27 |
| ENSG00000146109 | ABT1 | -0.57 | 0.05 | 1.14E-27 | 1.41E-26 |
| ENSG00000168411 | RFWD3 | -0.57 | 0.06 | 9.80E-25 | 1.10E-23 |
| ENSG00000110583 | NAA40 | -0.57 | 0.06 | 1.13E-24 | 1.26E-23 |
| ENSG00000153823 | PID1 | -0.57 | 0.06 | 1.22E-23 | 1.31E-22 |
| ENSG00000162722 | TRIM58 | -0.57 | 0.06 | 3.80E-21 | 3.72E-20 |
| ENSG00000196968 | FUT11 | -0.57 | 0.06 | 6.49E-21 | 6.27E-20 |
| ENSG00000135636 | DYSF | -0.57 | 0.06 | 1.14E-20 | 1.09E-19 |
| ENSG00000175414 | ARL10 | -0.57 | 0.06 | 3.14E-19 | 2.82E-18 |
| ENSG00000135974 | C2orf49 | -0.57 | 0.06 | 3.91E-19 | 3.49E-18 |
| ENSG00000138185 | ENTPD1 | -0.57 | 0.07 | 3.12E-17 | 2.57E-16 |

| | | | | | |
|---|---|---|---|---|---|
| ENSG00000213339 | QTRT1 | -0.57 | 0.07 | 4.46E-17 | 3.65E-16 |
| ENSG00000171604 | CXXC5 | -0.57 | 0.07 | 3.69E-16 | 2.89E-15 |
| ENSG00000198160 | MIER1 | -0.57 | 0.07 | 4.97E-16 | 3.87E-15 |
| ENSG00000128185 | DGCR6L | -0.57 | 0.08 | 7.70E-15 | 5.67E-14 |
| ENSG00000276234 | TADA2A | -0.57 | 0.08 | 3.38E-14 | 2.41E-13 |
| ENSG00000104427 | ZC2HC1A | -0.57 | 0.08 | 2.19E-13 | 1.50E-12 |
| ENSG00000197989 | SNHG12 | -0.57 | 0.08 | 2.27E-13 | 1.55E-12 |
| ENSG00000120068 | HOXB8 | -0.57 | 0.08 | 2.76E-13 | 1.87E-12 |
| ENSG00000170364 | SETMAR | -0.57 | 0.08 | 7.15E-13 | 4.74E-12 |
| ENSG00000173193 | PARP14 | -0.57 | 0.08 | 4.14E-12 | 2.62E-11 |
| ENSG00000131378 | RFTN1 | -0.57 | 0.09 | 1.37E-11 | 8.38E-11 |
| ENSG00000180329 | CCDC43 | -0.57 | 0.09 | 2.16E-11 | 1.31E-10 |
| ENSG00000145375 | SPATA5 | -0.57 | 0.09 | 7.16E-11 | 4.18E-10 |
| ENSG00000147251 | DOCK11 | -0.57 | 0.09 | 1.14E-10 | 6.60E-10 |
| ENSG00000111012 | CYP27B1 | -0.57 | 0.09 | 2.27E-10 | 1.29E-09 |
| ENSG00000168556 | ING2 | -0.57 | 0.1 | 4.24E-09 | 2.18E-08 |
| ENSG00000164647 | STEAP1 | -0.57 | 0.1 | 5.04E-09 | 2.58E-08 |
| ENSG00000099139 | PCSK5 | -0.57 | 0.12 | 2.44E-07 | 1.10E-06 |
| ENSG00000180257 | ZNF816 | -0.57 | 0.12 | 7.53E-07 | 3.24E-06 |
| ENSG00000259768 | ZFHX3-AS1 | -0.57 | 0.12 | 8.12E-07 | 3.48E-06 |
| ENSG00000110057 | UNC93B1 | -0.57 | 0.12 | 1.45E-06 | 6.08E-06 |
| ENSG00000145982 | FARS2 | -0.57 | 0.13 | 1.60E-06 | 6.69E-06 |
| ENSG00000140280 | LYSMD2 | -0.57 | 0.15 | 2.28E-05 | 8.52E-05 |
| ENSG00000228649 | SNHG26 | -0.57 | 0.15 | 2.81E-05 | 1.04E-04 |
| ENSG00000180758 | GPR157 | -0.57 | 0.15 | 2.86E-05 | 1.06E-04 |
| ENSG00000196470 | SIAH1 | -0.57 | 0.15 | 3.41E-05 | 1.25E-04 |
| ENSG00000130830 | MPP1 | -0.57 | 0.15 | 4.38E-05 | 1.58E-04 |
| ENSG00000230359 | TPI1P2 | -0.57 | 0.16 | 7.53E-05 | 2.66E-04 |
| ENSG00000107954 | NEURL1 | -0.57 | 0.16 | 8.37E-05 | 2.94E-04 |
| ENSG00000186666 | BCDIN3D | -0.57 | 0.19 | 8.06E-04 | 2.49E-03 |
| ENSG00000165757 | JCAD | -0.57 | 0.2 | 1.13E-03 | 3.41E-03 |
| ENSG00000272800 | | -0.57 | 0.21 | 1.90E-03 | 5.53E-03 |
| ENSG00000151575 | TEX9 | -0.57 | 0.23 | 3.80E-03 | 1.05E-02 |
| ENSG00000170381 | SEMA3E | -0.57 | 0.26 | 7.21E-03 | 1.88E-02 |
| ENSG00000176761 | ZNF285BP | -0.57 | 0.27 | 1.01E-02 | 2.56E-02 |
| ENSG00000169621 | APLF | -0.57 | 0.31 | 1.79E-02 | 4.25E-02 |
| ENSG00000275874 | PICSAR | -0.57 | 0.74 | 1.93E-02 | 4.57E-02 |
| ENSG00000071054 | MAP4K4 | -0.56 | 0.04 | 1.08E-52 | 2.61E-51 |
| ENSG00000103742 | IGDCC4 | -0.56 | 0.05 | 4.74E-31 | 6.61E-30 |
| ENSG00000121486 | TRMT1L | -0.56 | 0.05 | 1.60E-29 | 2.11E-28 |
| ENSG00000171307 | ZDHHC16 | -0.56 | 0.05 | 2.08E-29 | 2.73E-28 |
| ENSG00000152795 | HNRNPDL | -0.56 | 0.05 | 1.29E-28 | 1.65E-27 |
| ENSG00000134313 | KIDINS220 | -0.56 | 0.05 | 4.36E-27 | 5.31E-26 |
| ENSG00000132773 | TOE1 | -0.56 | 0.05 | 2.03E-26 | 2.42E-25 |
| ENSG00000117133 | RPF1 | -0.56 | 0.05 | 9.08E-26 | 1.05E-24 |
| ENSG00000164211 | STARD4 | -0.56 | 0.05 | 4.85E-25 | 5.47E-24 |
| ENSG00000130935 | NOL11 | -0.56 | 0.06 | 2.02E-24 | 2.23E-23 |
| ENSG00000164902 | PHAX | -0.56 | 0.06 | 6.21E-24 | 6.75E-23 |
| ENSG00000179981 | TSHZ1 | -0.56 | 0.06 | 1.83E-20 | 1.73E-19 |
| ENSG00000100234 | TIMP3 | -0.56 | 0.06 | 5.54E-20 | 5.14E-19 |

| | | | | | |
|---|---|---|---|---|---|
| ENSG00000163961 | RNF168 | -0.56 | 0.07 | 3.65E-17 | 2.99E-16 |
| ENSG00000113810 | SMC4 | -0.56 | 0.07 | 9.40E-17 | 7.57E-16 |
| ENSG00000128951 | DUT | -0.56 | 0.07 | 9.91E-17 | 7.98E-16 |
| ENSG00000112297 | CRYBG1 | -0.56 | 0.07 | 2.54E-16 | 2.00E-15 |
| ENSG00000277161 | PIGW | -0.56 | 0.07 | 3.36E-15 | 2.52E-14 |
| ENSG00000121621 | KIF18A | -0.56 | 0.07 | 3.42E-15 | 2.56E-14 |
| ENSG00000168268 | NT5DC2 | -0.56 | 0.07 | 1.12E-14 | 8.17E-14 |
| ENSG00000174032 | SLC25A30 | -0.56 | 0.07 | 1.41E-14 | 1.02E-13 |
| ENSG00000152147 | GEMIN6 | -0.56 | 0.08 | 2.79E-13 | 1.89E-12 |
| ENSG00000082014 | SMARCD3 | -0.56 | 0.08 | 4.45E-12 | 2.80E-11 |
| ENSG00000062822 | POLD1 | -0.56 | 0.08 | 6.15E-12 | 3.84E-11 |
| ENSG00000130787 | HIP1R | -0.56 | 0.08 | 7.81E-12 | 4.85E-11 |
| ENSG00000077514 | POLD3 | -0.56 | 0.09 | 1.59E-11 | 9.71E-11 |
| ENSG00000133315 | MACROD1 | -0.56 | 0.09 | 1.85E-11 | 1.13E-10 |
| ENSG00000225614 | ZNF469 | -0.56 | 0.09 | 2.50E-10 | 1.41E-09 |
| ENSG00000233491 | CDHR17P | -0.56 | 0.09 | 4.88E-10 | 2.69E-09 |
| ENSG00000226137 | BAIAP2-DT | -0.56 | 0.09 | 5.00E-10 | 2.76E-09 |
| ENSG00000127191 | TRAF2 | -0.56 | 0.09 | 6.56E-10 | 3.59E-09 |
| ENSG00000156876 | SASS6 | -0.56 | 0.09 | 8.79E-10 | 4.76E-09 |
| ENSG00000111554 | MDM1 | -0.56 | 0.09 | 1.04E-09 | 5.61E-09 |
| ENSG00000136378 | ADAMTS7 | -0.56 | 0.1 | 1.79E-09 | 9.50E-09 |
| ENSG00000130592 | LSP1 | -0.56 | 0.1 | 7.04E-09 | 3.56E-08 |
| ENSG00000073711 | PPP2R3A | -0.56 | 0.11 | 3.59E-08 | 1.72E-07 |
| ENSG00000265817 | FSBP | -0.56 | 0.12 | 7.32E-07 | 3.15E-06 |
| ENSG00000172465 | TCEAL1 | -0.56 | 0.12 | 8.88E-07 | 3.79E-06 |
| ENSG00000181004 | BBS12 | -0.56 | 0.14 | 1.63E-05 | 6.17E-05 |
| ENSG00000148288 | GBGT1 | -0.56 | 0.14 | 2.56E-05 | 9.46E-05 |
| ENSG00000185924 | RTN4RL1 | -0.56 | 0.16 | 9.23E-05 | 3.22E-04 |
| ENSG00000103150 | MLYCD | -0.56 | 0.18 | 4.10E-04 | 1.32E-03 |
| ENSG00000132821 | VSTM2L | -0.56 | 0.18 | 5.57E-04 | 1.76E-03 |
| ENSG00000169026 | SLC49A3 | -0.56 | 0.19 | 7.04E-04 | 2.19E-03 |
| ENSG00000153012 | LGI2 | -0.56 | 0.2 | 1.54E-03 | 4.55E-03 |
| ENSG00000267493 | CIRBP-AS1 | -0.56 | 0.21 | 2.58E-03 | 7.33E-03 |
| ENSG00000255050 | | -0.56 | 0.22 | 2.89E-03 | 8.15E-03 |
| ENSG00000197417 | SHPK | -0.56 | 0.23 | 4.20E-03 | 1.15E-02 |
| ENSG00000158683 | PKD1L1 | -0.56 | 0.23 | 4.46E-03 | 1.21E-02 |
| ENSG00000197385 | ZNF860 | -0.56 | 0.23 | 4.50E-03 | 1.22E-02 |
| ENSG00000286606 | | -0.56 | 0.24 | 5.53E-03 | 1.48E-02 |
| ENSG00000172197 | MBOAT1 | -0.56 | 0.26 | 9.41E-03 | 2.39E-02 |
| ENSG00000221994 | ZNF630 | -0.56 | 0.29 | 1.61E-02 | 3.87E-02 |
| ENSG00000262454 | MIR193BHG | -0.56 | 0.3 | 1.69E-02 | 4.05E-02 |
| ENSG00000172005 | MAL | -0.56 | 0.75 | 1.70E-02 | 4.06E-02 |
| ENSG00000142623 | PADI1 | -0.56 | 0.74 | 1.90E-02 | 4.50E-02 |
| ENSG00000172379 | ARNT2 | -0.56 | 0.32 | 2.08E-02 | 4.88E-02 |
| ENSG00000064042 | LIMCH1 | -0.55 | 0.04 | 4.64E-39 | 8.16E-38 |
| ENSG00000151892 | GFRA1 | -0.55 | 0.05 | 4.09E-33 | 6.03E-32 |
| ENSG00000155090 | KLF10 | -0.55 | 0.05 | 3.55E-32 | 5.10E-31 |
| ENSG00000145623 | OSMR | -0.55 | 0.05 | 7.22E-32 | 1.03E-30 |
| ENSG00000176994 | SMCR8 | -0.55 | 0.05 | 1.73E-28 | 2.20E-27 |
| ENSG00000142669 | SH3BGRL3 | -0.55 | 0.05 | 5.79E-27 | 7.01E-26 |

| | | | | | |
|---|---|---|---|---|---|
| ENSG00000213699 | SLC35F6 | -0.55 | 0.06 | 1.34E-23 | 1.43E-22 |
| ENSG00000204209 | DAXX | -0.55 | 0.06 | 3.24E-23 | 3.41E-22 |
| ENSG00000102908 | NFAT5 | -0.55 | 0.06 | 7.13E-23 | 7.42E-22 |
| ENSG00000119042 | SATB2 | -0.55 | 0.06 | 2.57E-22 | 2.62E-21 |
| ENSG00000180228 | PRKRA | -0.55 | 0.06 | 6.14E-21 | 5.94E-20 |
| ENSG00000166923 | GREM1 | -0.55 | 0.06 | 7.68E-18 | 6.49E-17 |
| ENSG00000112852 | PCDHB2 | -0.55 | 0.07 | 2.98E-17 | 2.46E-16 |
| ENSG00000128714 | HOXD13 | -0.55 | 0.07 | 3.02E-16 | 2.38E-15 |
| ENSG00000196743 | GM2A | -0.55 | 0.07 | 2.40E-15 | 1.81E-14 |
| ENSG00000112130 | RNF8 | -0.55 | 0.07 | 5.34E-15 | 3.96E-14 |
| ENSG00000168818 | STX18 | -0.55 | 0.07 | 2.92E-14 | 2.10E-13 |
| ENSG00000095539 | SEMA4G | -0.55 | 0.08 | 7.08E-14 | 4.96E-13 |
| ENSG00000166004 | CEP295 | -0.55 | 0.08 | 1.78E-13 | 1.22E-12 |
| ENSG00000256771 | ZNF253 | -0.55 | 0.08 | 3.23E-12 | 2.06E-11 |
| ENSG00000128578 | STRIP2 | -0.55 | 0.08 | 6.45E-12 | 4.02E-11 |
| ENSG00000152422 | XRCC4 | -0.55 | 0.08 | 9.01E-12 | 5.58E-11 |
| ENSG00000164167 | LSM6 | -0.55 | 0.09 | 5.72E-11 | 3.36E-10 |
| ENSG00000173163 | COMMD1 | -0.55 | 0.09 | 6.01E-11 | 3.53E-10 |
| ENSG00000147180 | ZNF711 | -0.55 | 0.09 | 9.60E-11 | 5.58E-10 |
| ENSG00000138468 | SENP7 | -0.55 | 0.09 | 3.81E-10 | 2.12E-09 |
| ENSG00000146409 | SLC18B1 | -0.55 | 0.09 | 8.01E-10 | 4.35E-09 |
| ENSG00000272405 | | -0.55 | 0.09 | 8.47E-10 | 4.59E-09 |
| ENSG00000117877 | POLR1G | -0.55 | 0.1 | 1.88E-09 | 9.93E-09 |
| ENSG00000240583 | AQP1 | -0.55 | 0.1 | 4.01E-09 | 2.07E-08 |
| ENSG00000213626 | LBH | -0.55 | 0.1 | 1.78E-08 | 8.75E-08 |
| ENSG00000189190 | ZNF600 | -0.55 | 0.1 | 4.36E-08 | 2.08E-07 |
| ENSG00000031081 | ARHGAP31 | -0.55 | 0.11 | 9.03E-08 | 4.21E-07 |
| ENSG00000186654 | PRR5 | -0.55 | 0.11 | 1.60E-07 | 7.28E-07 |
| ENSG00000129968 | ABHD17A | -0.55 | 0.11 | 1.68E-07 | 7.66E-07 |
| ENSG00000135744 | AGT | -0.55 | 0.11 | 2.81E-07 | 1.26E-06 |
| ENSG00000183876 | ARSI | -0.55 | 0.12 | 5.97E-07 | 2.60E-06 |
| ENSG00000283154 | IQCJ-SCHIP1 | -0.55 | 0.12 | 6.59E-07 | 2.85E-06 |
| ENSG00000111850 | SMIM8 | -0.55 | 0.12 | 6.87E-07 | 2.97E-06 |
| ENSG00000245522 | LINC02709 | -0.55 | 0.12 | 9.07E-07 | 3.87E-06 |
| ENSG00000173214 | MFSD4B | -0.55 | 0.12 | 1.91E-06 | 7.88E-06 |
| ENSG00000146197 | SCUBE3 | -0.55 | 0.13 | 6.38E-06 | 2.51E-05 |
| ENSG00000178401 | DNAJC22 | -0.55 | 0.13 | 6.68E-06 | 2.63E-05 |
| ENSG00000160999 | SH2B2 | -0.55 | 0.14 | 2.02E-05 | 7.59E-05 |
| ENSG00000113296 | THBS4 | -0.55 | 0.17 | 3.58E-04 | 1.16E-03 |
| ENSG00000158525 | CPA5 | -0.55 | 0.19 | 1.46E-03 | 4.33E-03 |
| ENSG00000154133 | ROBO4 | -0.55 | 0.2 | 2.22E-03 | 6.38E-03 |
| ENSG00000157601 | MX1 | -0.55 | 0.21 | 2.77E-03 | 7.84E-03 |
| ENSG00000128011 | LRFN1 | -0.55 | 0.22 | 3.71E-03 | 1.02E-02 |
| ENSG00000257446 | ZNF878 | -0.55 | 0.23 | 5.10E-03 | 1.37E-02 |
| ENSG00000174599 | TRAM1L1 | -0.55 | 0.24 | 6.11E-03 | 1.62E-02 |
| ENSG00000225096 | | -0.55 | 0.27 | 1.14E-02 | 2.84E-02 |
| ENSG00000101361 | NOP56 | -0.54 | 0.04 | 8.17E-47 | 1.71E-45 |
| ENSG00000073792 | IGF2BP2 | -0.54 | 0.04 | 1.25E-40 | 2.28E-39 |
| ENSG00000114648 | KLHL18 | -0.54 | 0.04 | 2.12E-38 | 3.68E-37 |
| ENSG00000102910 | LONP2 | -0.54 | 0.04 | 3.60E-34 | 5.48E-33 |

| | | | | | |
|---|---|---|---|---|---|
| ENSG00000087303 | NID2 | -0.54 | 0.05 | 9.16E-33 | 1.34E-31 |
| ENSG00000104299 | INTS9 | -0.54 | 0.05 | 3.89E-28 | 4.87E-27 |
| ENSG00000146281 | PM20D2 | -0.54 | 0.05 | 7.74E-25 | 8.70E-24 |
| ENSG00000011376 | LARS2 | -0.54 | 0.05 | 2.24E-24 | 2.46E-23 |
| ENSG00000165494 | PCF11 | -0.54 | 0.05 | 6.77E-24 | 7.35E-23 |
| ENSG00000162129 | CLPB | -0.54 | 0.05 | 8.28E-24 | 8.95E-23 |
| ENSG00000173950 | XXYLT1 | -0.54 | 0.05 | 1.75E-23 | 1.87E-22 |
| ENSG00000163539 | CLASP2 | -0.54 | 0.06 | 6.67E-22 | 6.69E-21 |
| ENSG00000155096 | AZIN1 | -0.54 | 0.06 | 4.94E-19 | 4.39E-18 |
| ENSG00000198837 | DENND4B | -0.54 | 0.06 | 8.34E-18 | 7.03E-17 |
| ENSG00000137414 | FAM8A1 | -0.54 | 0.07 | 1.01E-16 | 8.11E-16 |
| ENSG00000135424 | ITGA7 | -0.54 | 0.07 | 3.92E-15 | 2.93E-14 |
| ENSG00000120318 | ARAP3 | -0.54 | 0.08 | 6.73E-13 | 4.47E-12 |
| ENSG00000196187 | TMEM63A | -0.54 | 0.08 | 7.51E-13 | 4.96E-12 |
| ENSG00000198039 | ZNF273 | -0.54 | 0.08 | 1.10E-12 | 7.18E-12 |
| ENSG00000173218 | VANGL1 | -0.54 | 0.08 | 1.13E-12 | 7.36E-12 |
| ENSG00000160877 | NACC1 | -0.54 | 0.08 | 2.70E-12 | 1.73E-11 |
| ENSG00000160193 | WDR4 | -0.54 | 0.08 | 4.41E-12 | 2.78E-11 |
| ENSG00000136877 | FPGS | -0.54 | 0.08 | 7.63E-12 | 4.74E-11 |
| ENSG00000107816 | LZTS2 | -0.54 | 0.08 | 7.95E-12 | 4.93E-11 |
| ENSG00000173638 | SLC19A1 | -0.54 | 0.08 | 1.26E-11 | 7.73E-11 |
| ENSG00000115687 | PASK | -0.54 | 0.08 | 3.18E-11 | 1.90E-10 |
| ENSG00000182287 | AP1S2 | -0.54 | 0.08 | 6.24E-11 | 3.66E-10 |
| ENSG00000196639 | HRH1 | -0.54 | 0.09 | 1.27E-09 | 6.81E-09 |
| ENSG00000267041 | ZNF850 | -0.54 | 0.09 | 2.22E-09 | 1.16E-08 |
| ENSG00000100726 | TELO2 | -0.54 | 0.09 | 4.53E-09 | 2.32E-08 |
| ENSG00000160796 | NBEAL2 | -0.54 | 0.1 | 7.34E-09 | 3.71E-08 |
| ENSG00000165118 | C9orf64 | -0.54 | 0.1 | 2.55E-08 | 1.24E-07 |
| ENSG00000133069 | TMCC2 | -0.54 | 0.1 | 3.81E-08 | 1.83E-07 |
| ENSG00000241852 | C8orf58 | -0.54 | 0.1 | 5.60E-08 | 2.65E-07 |
| ENSG00000073670 | ADAM11 | -0.54 | 0.11 | 1.17E-07 | 5.40E-07 |
| ENSG00000109576 | AADAT | -0.54 | 0.11 | 2.34E-07 | 1.06E-06 |
| ENSG00000114473 | IQCG | -0.54 | 0.12 | 1.45E-06 | 6.08E-06 |
| ENSG00000183963 | SMTN | -0.54 | 0.13 | 4.84E-06 | 1.93E-05 |
| ENSG00000157911 | PEX10 | -0.54 | 0.12 | 5.66E-06 | 2.24E-05 |
| ENSG00000136944 | LMX1B | -0.54 | 0.13 | 9.26E-06 | 3.59E-05 |
| ENSG00000112139 | MDGA1 | -0.54 | 0.13 | 1.93E-05 | 7.26E-05 |
| ENSG00000254087 | LYN | -0.54 | 0.14 | 2.42E-05 | 9.01E-05 |
| ENSG00000275764 | | -0.54 | 0.14 | 3.87E-05 | 1.41E-04 |
| ENSG00000153531 | ADPRHL1 | -0.54 | 0.15 | 8.93E-05 | 3.12E-04 |
| ENSG00000215492 | HNRNPA1P7 | -0.54 | 0.15 | 1.21E-04 | 4.15E-04 |
| ENSG00000154146 | NRGN | -0.54 | 0.15 | 1.31E-04 | 4.48E-04 |
| ENSG00000251615 | | -0.54 | 0.15 | 1.32E-04 | 4.53E-04 |
| ENSG00000286019 | NOTCH2NLB | -0.54 | 0.17 | 4.39E-04 | 1.41E-03 |
| ENSG00000261168 | | -0.54 | 0.21 | 3.25E-03 | 9.07E-03 |
| ENSG00000139971 | ARMH4 | -0.54 | 0.21 | 4.00E-03 | 1.10E-02 |
| ENSG00000113361 | CDH6 | -0.54 | 0.22 | 4.01E-03 | 1.10E-02 |
| ENSG00000101230 | ISM1 | -0.54 | 0.27 | 1.23E-02 | 3.05E-02 |
| ENSG00000130595 | TNNT3 | -0.54 | 0.26 | 1.28E-02 | 3.15E-02 |
| ENSG00000248015 | | -0.54 | 0.27 | 1.43E-02 | 3.49E-02 |

| | | | | | |
|---|---|---|---|---|---|
| ENSG00000100206 | DMC1 | -0.54 | 0.29 | 1.93E-02 | 4.57E-02 |
| ENSG00000161249 | DMKN | -0.54 | 0.29 | 1.99E-02 | 4.69E-02 |
| ENSG00000166250 | CLMP | -0.53 | 0.04 | 6.70E-38 | 1.15E-36 |
| ENSG00000187079 | TEAD1 | -0.53 | 0.04 | 1.56E-36 | 2.55E-35 |
| ENSG00000143376 | SNX27 | -0.53 | 0.05 | 2.35E-26 | 2.78E-25 |
| ENSG00000116679 | IVNS1ABP | -0.53 | 0.05 | 9.25E-25 | 1.04E-23 |
| ENSG00000198730 | CTR9 | -0.53 | 0.05 | 1.13E-24 | 1.27E-23 |
| ENSG00000271503 | CCL5 | -0.53 | 0.06 | 5.82E-21 | 5.64E-20 |
| ENSG00000086475 | SEPHS1 | -0.53 | 0.06 | 1.90E-18 | 1.65E-17 |
| ENSG00000197256 | KANK2 | -0.53 | 0.06 | 2.73E-18 | 2.35E-17 |
| ENSG00000144535 | DIS3L2 | -0.53 | 0.06 | 1.94E-17 | 1.62E-16 |
| ENSG00000198146 | ZNF770 | -0.53 | 0.06 | 1.21E-16 | 9.71E-16 |
| ENSG00000119917 | IFIT3 | -0.53 | 0.07 | 4.07E-16 | 3.18E-15 |
| ENSG00000125772 | GPCPD1 | -0.53 | 0.07 | 4.49E-16 | 3.50E-15 |
| ENSG00000132763 | MMACHC | -0.53 | 0.07 | 6.25E-16 | 4.84E-15 |
| ENSG00000204469 | PRRC2A | -0.53 | 0.07 | 8.06E-16 | 6.19E-15 |
| ENSG00000175283 | DOLK | -0.53 | 0.07 | 5.26E-15 | 3.90E-14 |
| ENSG00000077942 | FBLN1 | -0.53 | 0.07 | 8.53E-15 | 6.27E-14 |
| ENSG00000106624 | AEBP1 | -0.53 | 0.07 | 1.14E-14 | 8.33E-14 |
| ENSG00000005884 | ITGA3 | -0.53 | 0.07 | 2.11E-14 | 1.52E-13 |
| ENSG00000113318 | MSH3 | -0.53 | 0.07 | 1.56E-13 | 1.07E-12 |
| ENSG00000162065 | TBC1D24 | -0.53 | 0.07 | 1.62E-13 | 1.11E-12 |
| ENSG00000166845 | C18orf54 | -0.53 | 0.07 | 3.52E-13 | 2.37E-12 |
| ENSG00000119685 | TTLL5 | -0.53 | 0.07 | 6.82E-13 | 4.52E-12 |
| ENSG00000133195 | SLC39A11 | -0.53 | 0.08 | 2.13E-12 | 1.37E-11 |
| ENSG00000065923 | SLC9A7 | -0.53 | 0.08 | 6.73E-11 | 3.94E-10 |
| ENSG00000100104 | SRRD | -0.53 | 0.09 | 2.02E-10 | 1.15E-09 |
| ENSG00000205268 | PDE7A | -0.53 | 0.09 | 3.44E-10 | 1.92E-09 |
| ENSG00000169213 | RAB3B | -0.53 | 0.09 | 2.19E-09 | 1.15E-08 |
| ENSG00000183172 | SMDT1 | -0.53 | 0.09 | 3.38E-09 | 1.75E-08 |
| ENSG00000243927 | MRPS6 | -0.53 | 0.09 | 3.90E-09 | 2.01E-08 |
| ENSG00000134780 | DAGLA | -0.53 | 0.1 | 1.31E-08 | 6.48E-08 |
| ENSG00000124596 | OARD1 | -0.53 | 0.11 | 1.98E-07 | 8.97E-07 |
| ENSG00000175093 | SPSB4 | -0.53 | 0.12 | 1.63E-06 | 6.77E-06 |
| ENSG00000139832 | RAB20 | -0.53 | 0.12 | 2.81E-06 | 1.14E-05 |
| ENSG00000170153 | RNF150 | -0.53 | 0.13 | 1.81E-05 | 6.83E-05 |
| ENSG00000152582 | SPEF2 | -0.53 | 0.14 | 3.52E-05 | 1.29E-04 |
| ENSG00000215241 | LINC02449 | -0.53 | 0.15 | 1.76E-04 | 5.94E-04 |
| ENSG00000169884 | WNT10B | -0.53 | 0.16 | 2.36E-04 | 7.85E-04 |
| ENSG00000269973 | | -0.53 | 0.16 | 2.79E-04 | 9.18E-04 |
| ENSG00000204315 | FKBPL | -0.53 | 0.16 | 3.69E-04 | 1.19E-03 |
| ENSG00000077616 | NAALAD2 | -0.53 | 0.2 | 2.61E-03 | 7.41E-03 |
| ENSG00000261338 | | -0.53 | 0.21 | 4.29E-03 | 1.17E-02 |
| ENSG00000196159 | FAT4 | -0.53 | 0.22 | 5.04E-03 | 1.36E-02 |
| ENSG00000227001 | NBPF2P | -0.53 | 0.22 | 5.61E-03 | 1.49E-02 |
| ENSG00000162241 | SLC25A45 | -0.53 | 0.22 | 5.79E-03 | 1.54E-02 |
| ENSG00000169989 | TIGD4 | -0.53 | 0.23 | 7.74E-03 | 2.01E-02 |
| ENSG00000270959 | LPP-AS2 | -0.53 | 0.24 | 9.06E-03 | 2.31E-02 |
| ENSG00000105880 | DLX5 | -0.53 | 0.25 | 1.20E-02 | 2.98E-02 |
| ENSG00000204882 | GPR20 | -0.53 | 0.25 | 1.24E-02 | 3.06E-02 |

| | | | | | |
|---|---|---|---|---|---|
| ENSG00000248469 | | -0.53 | 0.28 | 1.72E-02 | 4.10E-02 |
| ENSG00000110427 | KIAA1549L | -0.53 | 0.29 | 2.02E-02 | 4.75E-02 |
| ENSG00000159176 | CSRP1 | -0.52 | 0.03 | 4.06E-53 | 9.87E-52 |
| ENSG00000171793 | CTPS1 | -0.52 | 0.04 | 1.56E-46 | 3.25E-45 |
| ENSG00000079819 | EPB41L2 | -0.52 | 0.04 | 1.05E-42 | 2.02E-41 |
| ENSG00000182197 | EXT1 | -0.52 | 0.04 | 6.35E-40 | 1.13E-38 |
| ENSG00000164951 | PDP1 | -0.52 | 0.04 | 2.79E-39 | 4.92E-38 |
| ENSG00000134531 | EMP1 | -0.52 | 0.04 | 4.04E-37 | 6.71E-36 |
| ENSG00000049245 | VAMP3 | -0.52 | 0.05 | 2.25E-30 | 3.06E-29 |
| ENSG00000169490 | TM2D2 | -0.52 | 0.05 | 7.48E-30 | 1.00E-28 |
| ENSG00000130294 | KIF1A | -0.52 | 0.05 | 1.17E-29 | 1.56E-28 |
| ENSG00000131469 | RPL27 | -0.52 | 0.05 | 1.44E-29 | 1.91E-28 |
| ENSG00000144021 | CIAO1 | -0.52 | 0.05 | 3.43E-29 | 4.48E-28 |
| ENSG00000003989 | SLC7A2 | -0.52 | 0.05 | 2.18E-28 | 2.76E-27 |
| ENSG00000183864 | TOB2 | -0.52 | 0.05 | 1.34E-25 | 1.54E-24 |
| ENSG00000126947 | ARMCX1 | -0.52 | 0.05 | 9.17E-24 | 9.89E-23 |
| ENSG00000110367 | DDX6 | -0.52 | 0.05 | 1.46E-23 | 1.56E-22 |
| ENSG00000118564 | FBXL5 | -0.52 | 0.05 | 3.17E-23 | 3.34E-22 |
| ENSG00000143155 | TIPRL | -0.52 | 0.05 | 1.99E-22 | 2.04E-21 |
| ENSG00000166508 | MCM7 | -0.52 | 0.06 | 1.97E-21 | 1.94E-20 |
| ENSG00000167447 | SMG8 | -0.52 | 0.05 | 2.24E-21 | 2.20E-20 |
| ENSG00000186522 | SEPTIN10 | -0.52 | 0.06 | 3.25E-21 | 3.19E-20 |
| ENSG00000161638 | ITGA5 | -0.52 | 0.06 | 1.43E-20 | 1.36E-19 |
| ENSG00000101442 | ACTR5 | -0.52 | 0.06 | 1.45E-20 | 1.38E-19 |
| ENSG00000165338 | HECTD2 | -0.52 | 0.06 | 1.66E-20 | 1.58E-19 |
| ENSG00000160803 | UBQLN4 | -0.52 | 0.06 | 3.84E-20 | 3.59E-19 |
| ENSG00000117697 | NSL1 | -0.52 | 0.06 | 3.76E-19 | 3.37E-18 |
| ENSG00000172775 | PSME3IP1 | -0.52 | 0.06 | 8.80E-19 | 7.73E-18 |
| ENSG00000183826 | BTBD9 | -0.52 | 0.06 | 1.06E-18 | 9.32E-18 |
| ENSG00000152642 | GPD1L | -0.52 | 0.06 | 7.71E-18 | 6.51E-17 |
| ENSG00000011198 | ABHD5 | -0.52 | 0.06 | 1.32E-17 | 1.10E-16 |
| ENSG00000124422 | USP22 | -0.52 | 0.06 | 1.61E-17 | 1.34E-16 |
| ENSG00000125733 | TRIP10 | -0.52 | 0.06 | 3.11E-17 | 2.56E-16 |
| ENSG00000090924 | PLEKHG2 | -0.52 | 0.06 | 5.77E-17 | 4.70E-16 |
| ENSG00000157778 | PSMG3 | -0.52 | 0.06 | 2.54E-16 | 2.00E-15 |
| ENSG00000132356 | PRKAA1 | -0.52 | 0.06 | 3.30E-16 | 2.59E-15 |
| ENSG00000137106 | GRHPR | -0.52 | 0.07 | 2.33E-15 | 1.76E-14 |
| ENSG00000165322 | ARHGAP12 | -0.52 | 0.07 | 3.32E-14 | 2.37E-13 |
| ENSG00000173166 | RAPH1 | -0.52 | 0.07 | 1.70E-13 | 1.17E-12 |
| ENSG00000175595 | ERCC4 | -0.52 | 0.07 | 5.07E-13 | 3.39E-12 |
| ENSG00000047621 | C12orf4 | -0.52 | 0.07 | 1.03E-12 | 6.72E-12 |
| ENSG00000163918 | RFC4 | -0.52 | 0.08 | 1.29E-12 | 8.42E-12 |
| ENSG00000171621 | SPSB1 | -0.52 | 0.08 | 7.22E-11 | 4.22E-10 |
| ENSG00000163362 | INAVA | -0.52 | 0.09 | 1.87E-09 | 9.89E-09 |
| ENSG00000119397 | CNTRL | -0.52 | 0.09 | 2.20E-09 | 1.16E-08 |
| ENSG00000160963 | COL26A1 | -0.52 | 0.09 | 3.22E-09 | 1.67E-08 |
| ENSG00000099910 | KLHL22 | -0.52 | 0.09 | 3.35E-09 | 1.74E-08 |
| ENSG00000031691 | CENPQ | -0.52 | 0.09 | 4.49E-09 | 2.30E-08 |
| ENSG00000156136 | DCK | -0.52 | 0.1 | 2.04E-08 | 1.00E-07 |
| ENSG00000057704 | TMCC3 | -0.52 | 0.1 | 2.80E-08 | 1.36E-07 |

| | | | | | |
|---|---|---|---|---|---|
| ENSG00000171466 | ZNF562 | -0.52 | 0.1 | 4.11E-08 | 1.96E-07 |
| ENSG00000127586 | CHTF18 | -0.52 | 0.1 | 5.64E-08 | 2.67E-07 |
| ENSG00000143971 | ETAA1 | -0.52 | 0.1 | 1.29E-07 | 5.94E-07 |
| ENSG00000122971 | ACADS | -0.52 | 0.11 | 3.03E-07 | 1.35E-06 |
| ENSG00000214826 | DDX12P | -0.52 | 0.11 | 1.57E-06 | 6.55E-06 |
| ENSG00000131849 | ZNF132 | -0.52 | 0.12 | 2.24E-06 | 9.19E-06 |
| ENSG00000177839 | PCDHB9 | -0.52 | 0.14 | 7.94E-05 | 2.80E-04 |
| ENSG00000282164 | PEG13 | -0.52 | 0.15 | 1.15E-04 | 3.96E-04 |
| ENSG00000291202 | | -0.52 | 0.18 | 1.10E-03 | 3.32E-03 |
| ENSG00000135929 | CYP27A1 | -0.52 | 0.18 | 1.30E-03 | 3.88E-03 |
| ENSG00000180867 | PDIA3P1 | -0.52 | 0.18 | 1.34E-03 | 4.00E-03 |
| ENSG00000267508 | ZNF285 | -0.52 | 0.2 | 2.95E-03 | 8.30E-03 |
| ENSG00000164627 | KIF6 | -0.52 | 0.23 | 7.62E-03 | 1.98E-02 |
| ENSG00000155265 | GOLGA7B | -0.52 | 0.27 | 1.90E-02 | 4.50E-02 |
| ENSG00000142687 | KIAA0319L | -0.51 | 0.04 | 3.57E-39 | 6.30E-38 |
| ENSG00000114302 | PRKAR2A | -0.51 | 0.04 | 1.27E-35 | 2.04E-34 |
| ENSG00000164574 | GALNT10 | -0.51 | 0.04 | 1.04E-31 | 1.48E-30 |
| ENSG00000064999 | ANKS1A | -0.51 | 0.04 | 2.66E-31 | 3.74E-30 |
| ENSG00000018510 | AGPS | -0.51 | 0.05 | 1.75E-30 | 2.39E-29 |
| ENSG00000139990 | DCAF5 | -0.51 | 0.04 | 2.06E-30 | 2.81E-29 |
| ENSG00000145041 | DCAF1 | -0.51 | 0.05 | 1.53E-29 | 2.02E-28 |
| ENSG00000171456 | ASXL1 | -0.51 | 0.05 | 2.79E-29 | 3.67E-28 |
| ENSG00000198890 | PRMT6 | -0.51 | 0.05 | 7.95E-28 | 9.89E-27 |
| ENSG00000148120 | AOPEP | -0.51 | 0.05 | 6.63E-27 | 8.02E-26 |
| ENSG00000130147 | SH3BP4 | -0.51 | 0.05 | 9.86E-27 | 1.19E-25 |
| ENSG00000100281 | HMGXB4 | -0.51 | 0.05 | 2.26E-26 | 2.68E-25 |
| ENSG00000141298 | SSH2 | -0.51 | 0.05 | 2.67E-25 | 3.05E-24 |
| ENSG00000135823 | STX6 | -0.51 | 0.05 | 1.95E-23 | 2.08E-22 |
| ENSG00000165650 | PDZD8 | -0.51 | 0.05 | 2.18E-23 | 2.31E-22 |
| ENSG00000154124 | OTULIN | -0.51 | 0.05 | 7.36E-22 | 7.37E-21 |
| ENSG00000180776 | ZDHHC20 | -0.51 | 0.06 | 1.21E-20 | 1.15E-19 |
| ENSG00000062725 | APPBP2 | -0.51 | 0.06 | 3.50E-18 | 2.99E-17 |
| ENSG00000107130 | NCS1 | -0.51 | 0.06 | 1.56E-17 | 1.30E-16 |
| ENSG00000166411 | IDH3A | -0.51 | 0.06 | 9.31E-17 | 7.51E-16 |
| ENSG00000122335 | SERAC1 | -0.51 | 0.06 | 1.72E-16 | 1.37E-15 |
| ENSG00000184083 | FAM120C | -0.51 | 0.07 | 3.35E-15 | 2.51E-14 |
| ENSG00000132612 | VPS4A | -0.51 | 0.07 | 4.32E-15 | 3.22E-14 |
| ENSG00000198825 | INPP5F | -0.51 | 0.07 | 6.94E-15 | 5.12E-14 |
| ENSG00000139291 | TMEM19 | -0.51 | 0.07 | 5.67E-14 | 4.00E-13 |
| ENSG00000109881 | CCDC34 | -0.51 | 0.08 | 3.34E-12 | 2.12E-11 |
| ENSG00000065615 | CYB5R4 | -0.51 | 0.08 | 4.08E-12 | 2.58E-11 |
| ENSG00000214941 | ZSWIM7 | -0.51 | 0.08 | 1.10E-11 | 6.78E-11 |
| ENSG00000156521 | TYSND1 | -0.51 | 0.08 | 1.80E-11 | 1.10E-10 |
| ENSG00000257267 | ZNF271P | -0.51 | 0.08 | 2.47E-11 | 1.49E-10 |
| ENSG00000290429 | | -0.51 | 0.08 | 4.18E-11 | 2.48E-10 |
| ENSG00000120334 | CENPL | -0.51 | 0.08 | 4.40E-11 | 2.61E-10 |
| ENSG00000137145 | DENND4C | -0.51 | 0.08 | 7.06E-11 | 4.13E-10 |
| ENSG00000011143 | MKS1 | -0.51 | 0.08 | 2.45E-10 | 1.39E-09 |
| ENSG00000108666 | C17orf75 | -0.51 | 0.08 | 3.22E-10 | 1.80E-09 |
| ENSG00000198353 | HOXC4 | -0.51 | 0.09 | 1.48E-09 | 7.88E-09 |

| | | | | | |
|---|---|---|---|---|---|
| ENSG00000137166 | FOXP4 | -0.51 | 0.09 | 1.13E-08 | 5.62E-08 |
| ENSG00000153029 | MR1 | -0.51 | 0.09 | 1.13E-08 | 5.62E-08 |
| ENSG00000139133 | ALG10 | -0.51 | 0.1 | 3.39E-08 | 1.63E-07 |
| ENSG00000185761 | ADAMTSL5 | -0.51 | 0.1 | 2.29E-07 | 1.03E-06 |
| ENSG00000178694 | NSUN3 | -0.51 | 0.1 | 2.69E-07 | 1.21E-06 |
| ENSG00000112874 | NUDT12 | -0.51 | 0.1 | 3.90E-07 | 1.73E-06 |
| ENSG00000112234 | FBXL4 | -0.51 | 0.11 | 6.54E-07 | 2.83E-06 |
| ENSG00000117013 | KCNQ4 | -0.51 | 0.12 | 5.60E-06 | 2.22E-05 |
| ENSG00000156804 | FBXO32 | -0.51 | 0.12 | 9.81E-06 | 3.80E-05 |
| ENSG00000185347 | TEDC1 | -0.51 | 0.14 | 9.54E-05 | 3.32E-04 |
| ENSG00000173083 | HPSE | -0.51 | 0.15 | 3.05E-04 | 9.98E-04 |
| ENSG00000291064 | RPL23AP7 | -0.51 | 0.17 | 8.23E-04 | 2.54E-03 |
| ENSG00000187231 | SESTD1 | -0.51 | 0.19 | 2.48E-03 | 7.09E-03 |
| ENSG00000072163 | LIMS2 | -0.51 | 0.19 | 2.81E-03 | 7.94E-03 |
| ENSG00000279696 | | -0.51 | 0.2 | 3.01E-03 | 8.46E-03 |
| ENSG00000278784 | | -0.51 | 0.21 | 5.89E-03 | 1.56E-02 |
| ENSG00000270039 | | -0.51 | 0.26 | 1.71E-02 | 4.09E-02 |
| ENSG00000213225 | NOC2LP1 | -0.51 | 0.27 | 2.13E-02 | 4.99E-02 |
| ENSG00000120733 | KDM3B | -0.5 | 0.04 | 5.59E-34 | 8.47E-33 |
| ENSG00000172292 | CERS6 | -0.5 | 0.04 | 1.40E-33 | 2.10E-32 |
| ENSG00000174840 | PDE12 | -0.5 | 0.05 | 1.95E-28 | 2.47E-27 |
| ENSG00000172965 | MIR4435-2HG | -0.5 | 0.05 | 2.61E-26 | 3.08E-25 |
| ENSG00000196914 | ARHGEF12 | -0.5 | 0.05 | 3.60E-26 | 4.23E-25 |
| ENSG00000205726 | ITSN1 | -0.5 | 0.05 | 2.27E-24 | 2.50E-23 |
| ENSG00000143379 | SETDB1 | -0.5 | 0.05 | 1.55E-23 | 1.65E-22 |
| ENSG00000113621 | TXNDC15 | -0.5 | 0.05 | 4.07E-22 | 4.11E-21 |
| ENSG00000060688 | SNRNP40 | -0.5 | 0.05 | 6.08E-22 | 6.12E-21 |
| ENSG00000197461 | PDGFA | -0.5 | 0.05 | 8.77E-22 | 8.76E-21 |
| ENSG00000156299 | TIAM1 | -0.5 | 0.05 | 7.01E-21 | 6.75E-20 |
| ENSG00000164402 | SEPTIN8 | -0.5 | 0.06 | 5.13E-20 | 4.77E-19 |
| ENSG00000135018 | UBQLN1 | -0.5 | 0.06 | 1.34E-19 | 1.22E-18 |
| ENSG00000174151 | CYB561D1 | -0.5 | 0.06 | 1.52E-19 | 1.38E-18 |
| ENSG00000263753 | LINC00667 | -0.5 | 0.06 | 6.74E-19 | 5.95E-18 |
| ENSG00000012963 | UBR7 | -0.5 | 0.06 | 7.69E-19 | 6.76E-18 |
| ENSG00000101955 | SRPX | -0.5 | 0.06 | 1.11E-18 | 9.73E-18 |
| ENSG00000094916 | CBX5 | -0.5 | 0.06 | 2.46E-18 | 2.12E-17 |
| ENSG00000116157 | GPX7 | -0.5 | 0.06 | 6.01E-18 | 5.10E-17 |
| ENSG00000168002 | POLR2G | -0.5 | 0.06 | 7.98E-18 | 6.73E-17 |
| ENSG00000121316 | PLBD1 | -0.5 | 0.06 | 1.96E-17 | 1.63E-16 |
| ENSG00000013288 | MAN2B2 | -0.5 | 0.06 | 3.67E-17 | 3.01E-16 |
| ENSG00000117543 | DPH5 | -0.5 | 0.06 | 4.81E-16 | 3.75E-15 |
| ENSG00000135439 | AGAP2 | -0.5 | 0.06 | 3.43E-15 | 2.57E-14 |
| ENSG00000100568 | VTI1B | -0.5 | 0.07 | 1.92E-14 | 1.39E-13 |
| ENSG00000158290 | CUL4B | -0.5 | 0.07 | 2.89E-14 | 2.07E-13 |
| ENSG00000185920 | PTCH1 | -0.5 | 0.07 | 1.39E-13 | 9.56E-13 |
| ENSG00000198093 | ZNF649 | -0.5 | 0.07 | 5.89E-13 | 3.92E-12 |
| ENSG00000181026 | AEN | -0.5 | 0.07 | 1.77E-12 | 1.14E-11 |
| ENSG00000095015 | MAP3K1 | -0.5 | 0.07 | 2.07E-12 | 1.33E-11 |
| ENSG00000113389 | NPR3 | -0.5 | 0.07 | 2.41E-12 | 1.54E-11 |
| ENSG00000130818 | ZNF426 | -0.5 | 0.07 | 4.29E-12 | 2.71E-11 |

| | | | | | |
|---|---|---|---|---|---|
| ENSG00000115325 | DOK1 | -0.5 | 0.07 | 1.02E-11 | 6.28E-11 |
| ENSG00000099284 | MACROH2A2 | -0.5 | 0.08 | 2.24E-11 | 1.36E-10 |
| ENSG00000149582 | TMEM25 | -0.5 | 0.08 | 2.22E-10 | 1.26E-09 |
| ENSG00000233757 | ZNF892 | -0.5 | 0.08 | 2.68E-10 | 1.51E-09 |
| ENSG00000163659 | TIPARP | -0.5 | 0.08 | 2.70E-10 | 1.52E-09 |
| ENSG00000141959 | PFKL | -0.5 | 0.08 | 3.60E-10 | 2.01E-09 |
| ENSG00000130021 | PUDP | -0.5 | 0.08 | 4.34E-10 | 2.41E-09 |
| ENSG00000129317 | PUS7L | -0.5 | 0.08 | 1.04E-09 | 5.60E-09 |
| ENSG00000177683 | THAP5 | -0.5 | 0.09 | 2.82E-09 | 1.47E-08 |
| ENSG00000288658 | | -0.5 | 0.09 | 6.31E-09 | 3.20E-08 |
| ENSG00000256268 | LINC02454 | -0.5 | 0.09 | 1.61E-08 | 7.95E-08 |
| ENSG00000213445 | SIPA1 | -0.5 | 0.09 | 1.96E-08 | 9.61E-08 |
| ENSG00000182919 | C11orf54 | -0.5 | 0.1 | 5.54E-08 | 2.62E-07 |
| ENSG00000103145 | HCFC1R1 | -0.5 | 0.1 | 7.83E-08 | 3.66E-07 |
| ENSG00000006062 | MAP3K14 | -0.5 | 0.1 | 1.58E-07 | 7.22E-07 |
| ENSG00000237945 | LINC00649 | -0.5 | 0.1 | 2.20E-07 | 9.96E-07 |
| ENSG00000152455 | SUV39H2 | -0.5 | 0.1 | 4.85E-07 | 2.13E-06 |
| ENSG00000162390 | ACOT11 | -0.5 | 0.11 | 6.90E-07 | 2.98E-06 |
| ENSG00000143942 | CHAC2 | -0.5 | 0.11 | 8.31E-07 | 3.56E-06 |
| ENSG00000153790 | C7orf31 | -0.5 | 0.12 | 1.62E-05 | 6.12E-05 |
| ENSG00000117010 | ZNF684 | -0.5 | 0.13 | 3.20E-05 | 1.17E-04 |
| ENSG00000136098 | NEK3 | -0.5 | 0.13 | 3.68E-05 | 1.34E-04 |
| ENSG00000168779 | SHOX2 | -0.5 | 0.13 | 4.71E-05 | 1.70E-04 |
| ENSG00000230453 | ANKRD18B | -0.5 | 0.14 | 6.95E-05 | 2.46E-04 |
| ENSG00000271147 | ARMCX5-GPRASI | -0.5 | 0.14 | 1.09E-04 | 3.78E-04 |
| ENSG00000185305 | ARL15 | -0.5 | 0.14 | 1.39E-04 | 4.75E-04 |
| ENSG00000259959 | | -0.5 | 0.14 | 1.42E-04 | 4.85E-04 |
| ENSG00000262246 | CORO7 | -0.5 | 0.15 | 2.08E-04 | 6.96E-04 |
| ENSG00000164484 | TMEM200A | -0.5 | 0.15 | 2.81E-04 | 9.24E-04 |
| ENSG00000149403 | GRIK4 | -0.5 | 0.15 | 4.03E-04 | 1.30E-03 |
| ENSG00000181418 | DDN | -0.5 | 0.16 | 5.65E-04 | 1.79E-03 |
| ENSG00000105516 | DBP | -0.5 | 0.18 | 2.18E-03 | 6.27E-03 |
| ENSG00000204314 | PRRT1 | -0.5 | 0.22 | 8.59E-03 | 2.20E-02 |
| ENSG00000171631 | P2RY6 | -0.5 | 0.26 | 1.89E-02 | 4.48E-02 |
| ENSG00000167984 | NLRC3 | -0.5 | 0.26 | 2.03E-02 | 4.79E-02 |
| ENSG00000143569 | UBAP2L | -0.49 | 0.03 | 2.92E-55 | 7.48E-54 |
| ENSG00000169032 | MAP2K1 | -0.49 | 0.04 | 1.58E-32 | 2.29E-31 |
| ENSG00000124160 | NCOA5 | -0.49 | 0.05 | 3.28E-27 | 4.01E-26 |
| ENSG00000107779 | BMPR1A | -0.49 | 0.05 | 1.61E-26 | 1.92E-25 |
| ENSG00000198830 | HMGN2 | -0.49 | 0.05 | 2.81E-26 | 3.32E-25 |
| ENSG00000119392 | GLE1 | -0.49 | 0.05 | 6.89E-23 | 7.17E-22 |
| ENSG00000136158 | SPRY2 | -0.49 | 0.05 | 1.43E-22 | 1.47E-21 |
| ENSG00000152492 | CCDC50 | -0.49 | 0.05 | 1.09E-21 | 1.08E-20 |
| ENSG00000071859 | FAM50A | -0.49 | 0.05 | 1.83E-21 | 1.81E-20 |
| ENSG00000139370 | SLC15A4 | -0.49 | 0.05 | 7.24E-21 | 6.96E-20 |
| ENSG00000114999 | TTL | -0.49 | 0.05 | 7.12E-20 | 6.59E-19 |
| ENSG00000178163 | ZNF518B | -0.49 | 0.06 | 3.92E-19 | 3.50E-18 |
| ENSG00000155827 | RNF20 | -0.49 | 0.06 | 2.28E-18 | 1.97E-17 |
| ENSG00000189306 | RRP7A | -0.49 | 0.06 | 1.03E-17 | 8.67E-17 |
| ENSG00000204356 | NELFE | -0.49 | 0.06 | 2.02E-17 | 1.67E-16 |

| | | | | | |
|---|---|---|---|---|---|
| ENSG00000163541 | SUCLG1 | -0.49 | 0.06 | 2.97E-17 | 2.45E-16 |
| ENSG00000065457 | ADAT1 | -0.49 | 0.06 | 3.66E-16 | 2.87E-15 |
| ENSG00000160953 | PWWP3A | -0.49 | 0.06 | 5.05E-15 | 3.75E-14 |
| ENSG00000198951 | NAGA | -0.49 | 0.06 | 2.12E-14 | 1.53E-13 |
| ENSG00000120837 | NFYB | -0.49 | 0.07 | 3.41E-14 | 2.43E-13 |
| ENSG00000138399 | FASTKD1 | -0.49 | 0.07 | 5.94E-14 | 4.18E-13 |
| ENSG00000197056 | ZMYM1 | -0.49 | 0.07 | 8.87E-14 | 6.19E-13 |
| ENSG00000130717 | UCK1 | -0.49 | 0.07 | 2.06E-13 | 1.41E-12 |
| ENSG00000118518 | RNF146 | -0.49 | 0.07 | 8.24E-13 | 5.43E-12 |
| ENSG00000116809 | ZBTB17 | -0.49 | 0.08 | 3.88E-10 | 2.16E-09 |
| ENSG00000005073 | HOXA11 | -0.49 | 0.08 | 1.04E-09 | 5.60E-09 |
| ENSG00000163661 | PTX3 | -0.49 | 0.08 | 1.16E-09 | 6.26E-09 |
| ENSG00000212978 | C2orf74-DT | -0.49 | 0.08 | 1.52E-09 | 8.10E-09 |
| ENSG00000196935 | SRGAP1 | -0.49 | 0.09 | 2.86E-09 | 1.49E-08 |
| ENSG00000158301 | GPRASP2 | -0.49 | 0.09 | 8.12E-09 | 4.09E-08 |
| ENSG00000169635 | HIC2 | -0.49 | 0.09 | 2.81E-08 | 1.36E-07 |
| ENSG00000250317 | SMIM20 | -0.49 | 0.09 | 3.33E-08 | 1.60E-07 |
| ENSG00000152518 | ZFP36L2 | -0.49 | 0.09 | 6.70E-08 | 3.15E-07 |
| ENSG00000165632 | TAF3 | -0.49 | 0.09 | 8.10E-08 | 3.79E-07 |
| ENSG00000198113 | TOR4A | -0.49 | 0.1 | 3.48E-07 | 1.55E-06 |
| ENSG00000156795 | NTAQ1 | -0.49 | 0.1 | 6.75E-07 | 2.92E-06 |
| ENSG00000228065 | LINC01515 | -0.49 | 0.1 | 1.02E-06 | 4.30E-06 |
| ENSG00000108960 | MMD | -0.49 | 0.11 | 1.79E-06 | 7.42E-06 |
| ENSG00000087077 | TRIP6 | -0.49 | 0.11 | 2.43E-06 | 9.96E-06 |
| ENSG00000173085 | COQ2 | -0.49 | 0.12 | 2.38E-05 | 8.84E-05 |
| ENSG00000153406 | NMRAL1 | -0.49 | 0.15 | 3.12E-04 | 1.02E-03 |
| ENSG00000226919 | | -0.49 | 0.16 | 1.14E-03 | 3.43E-03 |
| ENSG00000184307 | ZDHHC23 | -0.49 | 0.18 | 2.06E-03 | 5.95E-03 |
| ENSG00000286532 | PARTICL | -0.49 | 0.18 | 2.09E-03 | 6.02E-03 |
| ENSG00000198948 | MFAP3L | -0.49 | 0.19 | 4.26E-03 | 1.16E-02 |
| ENSG00000143578 | CREB3L4 | -0.49 | 0.2 | 5.15E-03 | 1.38E-02 |
| ENSG00000150477 | KIAA1328 | -0.49 | 0.2 | 6.01E-03 | 1.59E-02 |
| ENSG00000291220 | | -0.49 | 0.2 | 6.09E-03 | 1.61E-02 |
| ENSG00000240356 | RPL23AP7 | -0.49 | 0.21 | 6.10E-03 | 1.61E-02 |
| ENSG00000125864 | BFSP1 | -0.49 | 0.22 | 8.64E-03 | 2.22E-02 |
| ENSG00000157429 | ZNF19 | -0.49 | 0.22 | 9.38E-03 | 2.39E-02 |
| ENSG00000271781 | | -0.49 | 0.24 | 1.62E-02 | 3.89E-02 |
| ENSG00000095587 | TLL2 | -0.48 | 0.03 | 1.13E-44 | 2.25E-43 |
| ENSG00000106829 | TLE4 | -0.48 | 0.04 | 2.71E-39 | 4.79E-38 |
| ENSG00000132688 | NES | -0.48 | 0.04 | 8.04E-30 | 1.08E-28 |
| ENSG00000173598 | NUDT4 | -0.48 | 0.05 | 1.55E-26 | 1.86E-25 |
| ENSG00000275700 | AATF | -0.48 | 0.05 | 8.71E-26 | 1.01E-24 |
| ENSG00000105974 | CAV1 | -0.48 | 0.05 | 1.25E-23 | 1.34E-22 |
| ENSG00000198467 | TPM2 | -0.48 | 0.05 | 2.57E-23 | 2.72E-22 |
| ENSG00000066136 | NFYC | -0.48 | 0.05 | 3.03E-22 | 3.08E-21 |
| ENSG00000105171 | POP4 | -0.48 | 0.05 | 5.54E-22 | 5.58E-21 |
| ENSG00000128567 | PODXL | -0.48 | 0.05 | 1.49E-21 | 1.47E-20 |
| ENSG00000114529 | C3orf52 | -0.48 | 0.05 | 5.11E-19 | 4.54E-18 |
| ENSG00000138375 | SMARCAL1 | -0.48 | 0.06 | 1.43E-17 | 1.20E-16 |
| ENSG00000111052 | LIN7A | -0.48 | 0.06 | 1.85E-17 | 1.54E-16 |

| | | | | | |
|---|---|---|---|---|---|
| ENSG00000101384 | JAG1 | -0.48 | 0.06 | 8.06E-17 | 6.53E-16 |
| ENSG00000136450 | SRSF1 | -0.48 | 0.06 | 1.14E-16 | 9.13E-16 |
| ENSG00000101639 | CEP192 | -0.48 | 0.06 | 3.76E-16 | 2.94E-15 |
| ENSG00000119421 | NDUFA8 | -0.48 | 0.06 | 5.45E-16 | 4.24E-15 |
| ENSG00000167378 | IRGQ | -0.48 | 0.06 | 7.26E-16 | 5.60E-15 |
| ENSG00000079931 | MOXD1 | -0.48 | 0.06 | 2.59E-15 | 1.95E-14 |
| ENSG00000119471 | HSDL2 | -0.48 | 0.06 | 7.59E-15 | 5.59E-14 |
| ENSG00000060656 | PTPRU | -0.48 | 0.06 | 2.59E-14 | 1.86E-13 |
| ENSG00000106399 | RPA3 | -0.48 | 0.06 | 3.14E-14 | 2.24E-13 |
| ENSG00000164961 | WASHC5 | -0.48 | 0.06 | 3.41E-14 | 2.43E-13 |
| ENSG00000164219 | PGGT1B | -0.48 | 0.06 | 7.85E-14 | 5.49E-13 |
| ENSG00000188283 | ZNF383 | -0.48 | 0.07 | 3.16E-12 | 2.02E-11 |
| ENSG00000105928 | GSDME | -0.48 | 0.07 | 6.40E-12 | 3.99E-11 |
| ENSG00000100601 | ALKBH1 | -0.48 | 0.07 | 1.56E-11 | 9.48E-11 |
| ENSG00000077684 | JADE1 | -0.48 | 0.07 | 5.70E-11 | 3.36E-10 |
| ENSG00000175416 | CLTB | -0.48 | 0.08 | 8.23E-10 | 4.47E-09 |
| ENSG00000272674 | PCDHB16 | -0.48 | 0.08 | 1.26E-09 | 6.77E-09 |
| ENSG00000135776 | ABCB10 | -0.48 | 0.08 | 1.96E-09 | 1.04E-08 |
| ENSG00000020426 | MNAT1 | -0.48 | 0.08 | 2.90E-09 | 1.51E-08 |
| ENSG00000152767 | FARP1 | -0.48 | 0.09 | 1.30E-08 | 6.47E-08 |
| ENSG00000203705 | TATDN3 | -0.48 | 0.09 | 2.95E-08 | 1.43E-07 |
| ENSG00000127328 | RAB3IP | -0.48 | 0.09 | 3.15E-08 | 1.52E-07 |
| ENSG00000185100 | ADSS1 | -0.48 | 0.09 | 3.48E-08 | 1.67E-07 |
| ENSG00000151466 | SCLT1 | -0.48 | 0.1 | 3.96E-07 | 1.75E-06 |
| ENSG00000169093 | ASMTL | -0.48 | 0.1 | 5.61E-07 | 2.44E-06 |
| ENSG00000197858 | GPAA1 | -0.48 | 0.1 | 7.88E-07 | 3.38E-06 |
| ENSG00000196636 | SDHAF3 | -0.48 | 0.1 | 9.59E-07 | 4.08E-06 |
| ENSG00000291201 | | -0.48 | 0.11 | 3.05E-06 | 1.24E-05 |
| ENSG00000124313 | IQSEC2 | -0.48 | 0.11 | 5.63E-06 | 2.23E-05 |
| ENSG00000023892 | DEF6 | -0.48 | 0.11 | 6.96E-06 | 2.73E-05 |
| ENSG00000125319 | HROB | -0.48 | 0.11 | 8.47E-06 | 3.30E-05 |
| ENSG00000250479 | CHCHD10 | -0.48 | 0.12 | 2.90E-05 | 1.07E-04 |
| ENSG00000159958 | TNFRSF13C | -0.48 | 0.12 | 3.95E-05 | 1.44E-04 |
| ENSG00000167992 | VWCE | -0.48 | 0.13 | 1.34E-04 | 4.58E-04 |
| ENSG00000100314 | CABP7 | -0.48 | 0.15 | 4.09E-04 | 1.32E-03 |
| ENSG00000163040 | CCDC74A | -0.48 | 0.15 | 5.57E-04 | 1.76E-03 |
| ENSG00000257167 | TMPO-AS1 | -0.48 | 0.15 | 6.18E-04 | 1.94E-03 |
| ENSG00000115616 | SLC9A2 | -0.48 | 0.15 | 8.32E-04 | 2.56E-03 |
| ENSG00000139914 | FITM1 | -0.48 | 0.17 | 1.88E-03 | 5.48E-03 |
| ENSG00000115271 | GCA | -0.48 | 0.2 | 6.71E-03 | 1.76E-02 |
| ENSG00000200534 | SNORA33 | -0.48 | 0.24 | 1.75E-02 | 4.17E-02 |
| ENSG00000122378 | PRXL2A | -0.47 | 0.04 | 9.72E-31 | 1.34E-29 |
| ENSG00000108312 | UBTF | -0.47 | 0.04 | 1.56E-27 | 1.93E-26 |
| ENSG00000134333 | LDHA | -0.47 | 0.05 | 3.25E-25 | 3.70E-24 |
| ENSG00000089280 | FUS | -0.47 | 0.05 | 2.11E-23 | 2.24E-22 |
| ENSG00000144645 | OSBPL10 | -0.47 | 0.05 | 2.69E-23 | 2.84E-22 |
| ENSG00000272886 | DCP1A | -0.47 | 0.05 | 2.82E-23 | 2.98E-22 |
| ENSG00000172943 | PHF8 | -0.47 | 0.05 | 5.14E-23 | 5.39E-22 |
| ENSG00000182963 | GJC1 | -0.47 | 0.05 | 6.79E-23 | 7.07E-22 |
| ENSG00000159267 | HLCS | -0.47 | 0.05 | 1.10E-22 | 1.13E-21 |

| | | | | | |
|---|---|---|---|---|---|
| ENSG00000101079 | NDRG3 | -0.47 | 0.05 | 1.06E-21 | 1.06E-20 |
| ENSG00000112739 | PRPF4B | -0.47 | 0.05 | 2.82E-21 | 2.77E-20 |
| ENSG00000174780 | SRP72 | -0.47 | 0.05 | 1.92E-20 | 1.82E-19 |
| ENSG00000065613 | SLK | -0.47 | 0.05 | 7.99E-20 | 7.38E-19 |
| ENSG00000124762 | CDKN1A | -0.47 | 0.05 | 1.09E-18 | 9.58E-18 |
| ENSG00000186469 | GNG2 | -0.47 | 0.05 | 2.03E-18 | 1.75E-17 |
| ENSG00000167491 | GATAD2A | -0.47 | 0.07 | 2.04E-13 | 1.39E-12 |
| ENSG00000182552 | RWDD4 | -0.47 | 0.06 | 2.50E-13 | 1.70E-12 |
| ENSG00000120690 | ELF1 | -0.47 | 0.07 | 4.73E-13 | 3.17E-12 |
| ENSG00000168769 | TET2 | -0.47 | 0.07 | 5.48E-13 | 3.65E-12 |
| ENSG00000142798 | HSPG2 | -0.47 | 0.07 | 1.12E-12 | 7.29E-12 |
| ENSG00000149269 | PAK1 | -0.47 | 0.07 | 1.30E-12 | 8.49E-12 |
| ENSG00000188647 | PTAR1 | -0.47 | 0.07 | 5.43E-12 | 3.40E-11 |
| ENSG00000146426 | TIAM2 | -0.47 | 0.07 | 9.50E-12 | 5.87E-11 |
| ENSG00000108784 | NAGLU | -0.47 | 0.07 | 1.53E-11 | 9.36E-11 |
| ENSG00000074266 | EED | -0.47 | 0.07 | 1.54E-11 | 9.37E-11 |
| ENSG00000001084 | GCLC | -0.47 | 0.07 | 3.22E-11 | 1.92E-10 |
| ENSG00000183978 | COA3 | -0.47 | 0.07 | 4.49E-11 | 2.66E-10 |
| ENSG00000196123 | MATCAP1 | -0.47 | 0.07 | 1.03E-10 | 5.96E-10 |
| ENSG00000121417 | ZNF211 | -0.47 | 0.07 | 1.24E-10 | 7.16E-10 |
| ENSG00000141569 | TRIM65 | -0.47 | 0.08 | 2.93E-10 | 1.64E-09 |
| ENSG00000221923 | ZNF880 | -0.47 | 0.08 | 3.20E-10 | 1.79E-09 |
| ENSG00000065989 | PDE4A | -0.47 | 0.08 | 5.84E-10 | 3.21E-09 |
| ENSG00000120549 | KIAA1217 | -0.47 | 0.08 | 2.25E-09 | 1.18E-08 |
| ENSG00000260917 | | -0.47 | 0.09 | 2.21E-08 | 1.08E-07 |
| ENSG00000131773 | KHDRBS3 | -0.47 | 0.09 | 6.55E-08 | 3.08E-07 |
| ENSG00000146828 | SLC12A9 | -0.47 | 0.09 | 7.24E-08 | 3.40E-07 |
| ENSG00000203326 | ZNF525 | -0.47 | 0.09 | 8.65E-08 | 4.03E-07 |
| ENSG00000185361 | TNFAIP8L1 | -0.47 | 0.09 | 1.07E-07 | 4.97E-07 |
| ENSG00000069509 | FUNDC1 | -0.47 | 0.09 | 1.85E-07 | 8.39E-07 |
| ENSG00000106714 | CNTNAP3 | -0.47 | 0.09 | 2.26E-07 | 1.02E-06 |
| ENSG00000213465 | ARL2 | -0.47 | 0.09 | 2.75E-07 | 1.23E-06 |
| ENSG00000172830 | SSH3 | -0.47 | 0.11 | 5.83E-06 | 2.31E-05 |
| ENSG00000010361 | FUZ | -0.47 | 0.12 | 2.61E-05 | 9.67E-05 |
| ENSG00000175322 | ZNF519 | -0.47 | 0.12 | 2.77E-05 | 1.02E-04 |
| ENSG00000135407 | AVIL | -0.47 | 0.12 | 4.71E-05 | 1.70E-04 |
| ENSG00000160062 | ZBTB8A | -0.47 | 0.13 | 8.66E-05 | 3.03E-04 |
| ENSG00000138617 | PARP16 | -0.47 | 0.13 | 1.09E-04 | 3.77E-04 |
| ENSG00000091592 | NLRP1 | -0.47 | 0.15 | 6.10E-04 | 1.92E-03 |
| ENSG00000165029 | ABCA1 | -0.47 | 0.15 | 6.33E-04 | 1.99E-03 |
| ENSG00000247796 | MOCS2-DT | -0.47 | 0.16 | 1.93E-03 | 5.61E-03 |
| ENSG00000228175 | GEMIN8P4 | -0.47 | 0.17 | 2.30E-03 | 6.60E-03 |
| ENSG00000138735 | PDE5A | -0.47 | 0.18 | 3.31E-03 | 9.22E-03 |
| ENSG00000130433 | CACNG6 | -0.47 | 0.21 | 1.18E-02 | 2.93E-02 |
| ENSG00000272970 | | -0.47 | 0.23 | 1.45E-02 | 3.53E-02 |
| ENSG00000060237 | WNK1 | -0.46 | 0.04 | 3.06E-35 | 4.83E-34 |
| ENSG00000164091 | WDR82 | -0.46 | 0.04 | 6.16E-34 | 9.32E-33 |
| ENSG00000188690 | UROS | -0.46 | 0.04 | 1.46E-29 | 1.93E-28 |
| ENSG00000113712 | CSNK1A1 | -0.46 | 0.04 | 1.19E-28 | 1.53E-27 |
| ENSG00000205730 | ITPRIPL2 | -0.46 | 0.04 | 1.84E-28 | 2.34E-27 |

| | | | | | |
|---|---|---|---|---|---|
| ENSG00000126883 | NUP214 | -0.46 | 0.04 | 1.38E-26 | 1.66E-25 |
| ENSG00000171208 | NETO2 | -0.46 | 0.04 | 3.07E-26 | 3.61E-25 |
| ENSG00000057252 | SOAT1 | -0.46 | 0.04 | 1.43E-25 | 1.65E-24 |
| ENSG00000166326 | TRIM44 | -0.46 | 0.04 | 2.16E-25 | 2.47E-24 |
| ENSG00000167881 | SRP68 | -0.46 | 0.05 | 2.74E-23 | 2.90E-22 |
| ENSG00000077238 | IL4R | -0.46 | 0.05 | 8.00E-23 | 8.31E-22 |
| ENSG00000136861 | CDK5RAP2 | -0.46 | 0.05 | 1.22E-22 | 1.26E-21 |
| ENSG00000049541 | RFC2 | -0.46 | 0.05 | 4.74E-21 | 4.61E-20 |
| ENSG00000139645 | ANKRD52 | -0.46 | 0.05 | 1.36E-19 | 1.24E-18 |
| ENSG00000169251 | NMD3 | -0.46 | 0.05 | 1.42E-17 | 1.19E-16 |
| ENSG00000164172 | MOCS2 | -0.46 | 0.05 | 2.80E-17 | 2.31E-16 |
| ENSG00000156017 | CARNMT1 | -0.46 | 0.06 | 3.45E-17 | 2.83E-16 |
| ENSG00000117500 | TMED5 | -0.46 | 0.06 | 2.83E-16 | 2.23E-15 |
| ENSG00000164347 | GFM2 | -0.46 | 0.06 | 5.71E-16 | 4.43E-15 |
| ENSG00000169299 | PGM2 | -0.46 | 0.06 | 2.32E-15 | 1.75E-14 |
| ENSG00000100522 | GNPNAT1 | -0.46 | 0.06 | 4.58E-15 | 3.41E-14 |
| ENSG00000119912 | IDE | -0.46 | 0.06 | 3.34E-14 | 2.38E-13 |
| ENSG00000172534 | HCFC1 | -0.46 | 0.06 | 3.62E-14 | 2.57E-13 |
| ENSG00000103034 | NDRG4 | -0.46 | 0.06 | 5.66E-14 | 3.99E-13 |
| ENSG00000160688 | FLAD1 | -0.46 | 0.06 | 6.77E-13 | 4.49E-12 |
| ENSG00000075239 | ACAT1 | -0.46 | 0.07 | 1.40E-12 | 9.07E-12 |
| ENSG00000107821 | KAZALD1 | -0.46 | 0.07 | 3.24E-11 | 1.94E-10 |
| ENSG00000174989 | FBXW8 | -0.46 | 0.07 | 7.21E-11 | 4.21E-10 |
| ENSG00000123064 | DDX54 | -0.46 | 0.07 | 3.44E-10 | 1.92E-09 |
| ENSG00000173436 | MICOS10 | -0.46 | 0.08 | 2.06E-09 | 1.09E-08 |
| ENSG00000166133 | RPUSD2 | -0.46 | 0.08 | 2.62E-09 | 1.37E-08 |
| ENSG00000163528 | CHCHD4 | -0.46 | 0.08 | 4.11E-09 | 2.12E-08 |
| ENSG00000176974 | SHMT1 | -0.46 | 0.08 | 1.20E-08 | 5.97E-08 |
| ENSG00000181381 | DDX60L | -0.46 | 0.08 | 1.81E-08 | 8.87E-08 |
| ENSG00000197594 | ENPP1 | -0.46 | 0.08 | 2.12E-08 | 1.03E-07 |
| ENSG00000129596 | CDO1 | -0.46 | 0.09 | 8.27E-08 | 3.86E-07 |
| ENSG00000104332 | SFRP1 | -0.46 | 0.09 | 2.15E-07 | 9.73E-07 |
| ENSG00000166206 | GABRB3 | -0.46 | 0.09 | 2.46E-07 | 1.11E-06 |
| ENSG00000083635 | NUFIP1 | -0.46 | 0.09 | 2.99E-07 | 1.34E-06 |
| ENSG00000119509 | INVS | -0.46 | 0.09 | 4.25E-07 | 1.87E-06 |
| ENSG00000198736 | MSRB1 | -0.46 | 0.09 | 4.34E-07 | 1.91E-06 |
| ENSG00000158882 | TOMM40L | -0.46 | 0.1 | 4.01E-06 | 1.61E-05 |
| ENSG00000185664 | PMEL | -0.46 | 0.11 | 1.76E-05 | 6.64E-05 |
| ENSG00000085871 | MGST2 | -0.46 | 0.14 | 2.46E-04 | 8.15E-04 |
| ENSG00000121851 | POLR3GL | -0.46 | 0.14 | 2.97E-04 | 9.72E-04 |
| ENSG00000170745 | KCNS3 | -0.46 | 0.14 | 6.15E-04 | 1.93E-03 |
| ENSG00000088882 | CPXM1 | -0.46 | 0.14 | 6.23E-04 | 1.96E-03 |
| ENSG00000118004 | COLEC11 | -0.46 | 0.16 | 1.49E-03 | 4.40E-03 |
| ENSG00000163082 | SGPP2 | -0.46 | 0.16 | 1.67E-03 | 4.88E-03 |
| ENSG00000168243 | GNG4 | -0.46 | 0.16 | 1.87E-03 | 5.46E-03 |
| ENSG00000204860 | FAM201A | -0.46 | 0.17 | 2.56E-03 | 7.28E-03 |
| ENSG00000275183 | LENG9 | -0.46 | 0.17 | 2.84E-03 | 8.01E-03 |
| ENSG00000171962 | DRC3 | -0.46 | 0.2 | 8.15E-03 | 2.10E-02 |
| ENSG00000180155 | LYNX1 | -0.46 | 0.21 | 1.21E-02 | 2.99E-02 |
| ENSG00000044115 | CTNNA1 | -0.45 | 0.04 | 1.45E-30 | 1.99E-29 |

| | | | | | |
|---|---|---|---|---|---|
| ENSG00000169021 | UQCRFS1 | -0.45 | 0.04 | 1.73E-29 | 2.28E-28 |
| ENSG00000173402 | DAG1 | -0.45 | 0.04 | 2.43E-29 | 3.20E-28 |
| ENSG00000135506 | OS9 | -0.45 | 0.04 | 1.04E-25 | 1.20E-24 |
| ENSG00000167721 | TSR1 | -0.45 | 0.04 | 1.51E-23 | 1.61E-22 |
| ENSG00000188229 | TUBB4B | -0.45 | 0.05 | 1.62E-23 | 1.73E-22 |
| ENSG00000118689 | FOXO3 | -0.45 | 0.05 | 6.43E-22 | 6.47E-21 |
| ENSG00000087274 | ADD1 | -0.45 | 0.05 | 1.48E-21 | 1.47E-20 |
| ENSG00000130414 | NDUFA10 | -0.45 | 0.05 | 1.61E-21 | 1.59E-20 |
| ENSG00000138801 | PAPSS1 | -0.45 | 0.05 | 4.22E-21 | 4.11E-20 |
| ENSG00000181163 | NPM1 | -0.45 | 0.05 | 3.36E-17 | 2.76E-16 |
| ENSG00000162928 | PEX13 | -0.45 | 0.05 | 6.66E-17 | 5.42E-16 |
| ENSG00000005156 | LIG3 | -0.45 | 0.06 | 1.39E-16 | 1.11E-15 |
| ENSG00000198954 | KIFBP | -0.45 | 0.06 | 4.22E-16 | 3.29E-15 |
| ENSG00000109099 | PMP22 | -0.45 | 0.06 | 5.61E-16 | 4.36E-15 |
| ENSG00000102870 | ZNF629 | -0.45 | 0.06 | 2.48E-15 | 1.87E-14 |
| ENSG00000164654 | MIOS | -0.45 | 0.06 | 4.81E-15 | 3.58E-14 |
| ENSG00000147604 | RPL7 | -0.45 | 0.06 | 5.96E-15 | 4.41E-14 |
| ENSG00000138642 | HERC6 | -0.45 | 0.06 | 9.47E-15 | 6.95E-14 |
| ENSG00000125970 | RALY | -0.45 | 0.06 | 3.33E-14 | 2.38E-13 |
| ENSG00000147419 | CCDC25 | -0.45 | 0.06 | 4.30E-14 | 3.04E-13 |
| ENSG00000184428 | TOP1MT | -0.45 | 0.06 | 1.77E-13 | 1.21E-12 |
| ENSG00000170485 | NPAS2 | -0.45 | 0.06 | 1.89E-13 | 1.29E-12 |
| ENSG00000143819 | EPHX1 | -0.45 | 0.06 | 2.94E-13 | 1.99E-12 |
| ENSG00000166451 | CENPN | -0.45 | 0.06 | 4.10E-13 | 2.75E-12 |
| ENSG00000115808 | STRN | -0.45 | 0.06 | 2.69E-12 | 1.72E-11 |
| ENSG00000144791 | LIMD1 | -0.45 | 0.07 | 1.15E-11 | 7.07E-11 |
| ENSG00000122870 | BICC1 | -0.45 | 0.07 | 2.86E-11 | 1.72E-10 |
| ENSG00000132481 | TRIM47 | -0.45 | 0.07 | 3.24E-11 | 1.94E-10 |
| ENSG00000121864 | ZNF639 | -0.45 | 0.07 | 3.52E-11 | 2.10E-10 |
| ENSG00000166938 | DIS3L | -0.45 | 0.07 | 5.68E-11 | 3.34E-10 |
| ENSG00000181830 | SLC35C1 | -0.45 | 0.07 | 1.32E-10 | 7.59E-10 |
| ENSG00000135090 | TAOK3 | -0.45 | 0.07 | 2.45E-10 | 1.38E-09 |
| ENSG00000165458 | INPPL1 | -0.45 | 0.07 | 5.88E-10 | 3.23E-09 |
| ENSG00000275832 | ARHGAP23 | -0.45 | 0.08 | 2.69E-09 | 1.41E-08 |
| ENSG00000164086 | DUSP7 | -0.45 | 0.08 | 1.12E-08 | 5.57E-08 |
| ENSG00000187741 | FANCA | -0.45 | 0.08 | 3.27E-08 | 1.57E-07 |
| ENSG00000197355 | UAP1L1 | -0.45 | 0.09 | 6.57E-08 | 3.09E-07 |
| ENSG00000129159 | KCNC1 | -0.45 | 0.09 | 7.94E-08 | 3.71E-07 |
| ENSG00000172638 | EFEMP2 | -0.45 | 0.09 | 2.48E-07 | 1.11E-06 |
| ENSG00000254535 | PABPC4L | -0.45 | 0.1 | 4.45E-06 | 1.78E-05 |
| ENSG00000258289 | CHURC1 | -0.45 | 0.1 | 6.41E-06 | 2.53E-05 |
| ENSG00000142459 | EVI5L | -0.45 | 0.1 | 6.66E-06 | 2.62E-05 |
| ENSG00000148200 | NR6A1 | -0.45 | 0.1 | 7.05E-06 | 2.77E-05 |
| ENSG00000021355 | SERPINB1 | -0.45 | 0.11 | 9.70E-06 | 3.76E-05 |
| ENSG00000185475 | TMEM179B | -0.45 | 0.11 | 3.28E-05 | 1.20E-04 |
| ENSG00000161513 | FDXR | -0.45 | 0.12 | 8.32E-05 | 2.92E-04 |
| ENSG00000118507 | AKAP7 | -0.45 | 0.13 | 2.18E-04 | 7.26E-04 |
| ENSG00000167889 | MGAT5B | -0.45 | 0.13 | 2.47E-04 | 8.19E-04 |
| ENSG00000177640 | CASC2 | -0.45 | 0.15 | 9.36E-04 | 2.86E-03 |
| ENSG00000270112 | ST8SIA5-DT | -0.45 | 0.16 | 1.59E-03 | 4.68E-03 |

| | | | | | |
|---|---|---|---|---|---|
| ENSG00000183323 | CCDC125 | -0.45 | 0.16 | 1.86E-03 | 5.41E-03 |
| ENSG00000280303 | ERICD | -0.45 | 0.16 | 1.95E-03 | 5.66E-03 |
| ENSG00000231107 | LINC01508 | -0.45 | 0.16 | 2.05E-03 | 5.91E-03 |
| ENSG00000288559 | | -0.45 | 0.16 | 2.60E-03 | 7.38E-03 |
| ENSG00000198203 | SULT1C2 | -0.45 | 0.17 | 3.31E-03 | 9.22E-03 |
| ENSG00000120899 | PTK2B | -0.45 | 0.18 | 5.48E-03 | 1.46E-02 |
| ENSG00000168026 | TTC21A | -0.45 | 0.18 | 5.59E-03 | 1.49E-02 |
| ENSG00000183208 | GDPGP1 | -0.45 | 0.2 | 1.07E-02 | 2.69E-02 |
| ENSG00000287299 | | -0.45 | 0.21 | 1.32E-02 | 3.24E-02 |
| ENSG00000167601 | AXL | -0.44 | 0.04 | 9.69E-30 | 1.29E-28 |
| ENSG00000082293 | COL19A1 | -0.44 | 0.04 | 5.22E-28 | 6.52E-27 |
| ENSG00000137573 | SULF1 | -0.44 | 0.04 | 2.46E-27 | 3.02E-26 |
| ENSG00000127616 | SMARCA4 | -0.44 | 0.05 | 8.98E-22 | 8.96E-21 |
| ENSG00000134744 | TUT4 | -0.44 | 0.05 | 2.06E-21 | 2.03E-20 |
| ENSG00000181090 | EHMT1 | -0.44 | 0.05 | 4.18E-20 | 3.90E-19 |
| ENSG00000108021 | TASOR2 | -0.44 | 0.05 | 1.71E-19 | 1.55E-18 |
| ENSG00000182827 | ACBD3 | -0.44 | 0.05 | 3.29E-18 | 2.82E-17 |
| ENSG00000113594 | LIFR | -0.44 | 0.05 | 3.65E-17 | 2.99E-16 |
| ENSG00000163811 | WDR43 | -0.44 | 0.05 | 1.25E-16 | 9.96E-16 |
| ENSG00000158985 | CDC42SE2 | -0.44 | 0.05 | 1.42E-16 | 1.13E-15 |
| ENSG00000143486 | EIF2D | -0.44 | 0.06 | 8.34E-15 | 6.13E-14 |
| ENSG00000100644 | HIF1A | -0.44 | 0.06 | 4.13E-14 | 2.93E-13 |
| ENSG00000132768 | DPH2 | -0.44 | 0.06 | 1.30E-12 | 8.46E-12 |
| ENSG00000164916 | FOXK1 | -0.44 | 0.06 | 3.10E-12 | 1.98E-11 |
| ENSG00000278053 | DDX52 | -0.44 | 0.06 | 3.26E-12 | 2.07E-11 |
| ENSG00000130772 | MED18 | -0.44 | 0.06 | 5.35E-12 | 3.35E-11 |
| ENSG00000151883 | PARP8 | -0.44 | 0.07 | 1.63E-11 | 9.93E-11 |
| ENSG00000196465 | MYL6B | -0.44 | 0.07 | 2.10E-11 | 1.27E-10 |
| ENSG00000132382 | MYBBP1A | -0.44 | 0.07 | 3.95E-11 | 2.35E-10 |
| ENSG00000116525 | TRIM62 | -0.44 | 0.07 | 1.21E-10 | 6.98E-10 |
| ENSG00000128534 | LSM8 | -0.44 | 0.07 | 6.84E-10 | 3.74E-09 |
| ENSG00000119929 | CUTC | -0.44 | 0.07 | 1.23E-09 | 6.57E-09 |
| ENSG00000187790 | FANCM | -0.44 | 0.08 | 1.93E-09 | 1.02E-08 |
| ENSG00000104884 | ERCC2 | -0.44 | 0.08 | 2.19E-09 | 1.15E-08 |
| ENSG00000249992 | TMEM158 | -0.44 | 0.08 | 2.35E-09 | 1.23E-08 |
| ENSG00000081721 | DUSP12 | -0.44 | 0.08 | 5.21E-09 | 2.66E-08 |
| ENSG00000049239 | H6PD | -0.44 | 0.08 | 6.44E-09 | 3.27E-08 |
| ENSG00000104325 | DECR1 | -0.44 | 0.08 | 2.61E-08 | 1.27E-07 |
| ENSG00000049089 | COL9A2 | -0.44 | 0.08 | 3.43E-08 | 1.65E-07 |
| ENSG00000170417 | TMEM182 | -0.44 | 0.08 | 5.02E-08 | 2.39E-07 |
| ENSG00000171827 | ZNF570 | -0.44 | 0.08 | 7.19E-08 | 3.38E-07 |
| ENSG00000101190 | TCFL5 | -0.44 | 0.09 | 2.05E-07 | 9.28E-07 |
| ENSG00000148481 | MINDY3 | -0.44 | 0.09 | 3.06E-07 | 1.36E-06 |
| ENSG00000166436 | TRIM66 | -0.44 | 0.09 | 4.61E-07 | 2.02E-06 |
| ENSG00000158445 | KCNB1 | -0.44 | 0.09 | 5.33E-07 | 2.33E-06 |
| ENSG00000181804 | SLC9A9 | -0.44 | 0.1 | 1.23E-05 | 4.73E-05 |
| ENSG00000182405 | PGBD4 | -0.44 | 0.11 | 1.80E-05 | 6.78E-05 |
| ENSG00000197766 | CFD | -0.44 | 0.12 | 1.49E-04 | 5.08E-04 |
| ENSG00000214223 | HNRNPA1P10 | -0.44 | 0.12 | 1.97E-04 | 6.60E-04 |
| ENSG00000162062 | TEDC2 | -0.44 | 0.13 | 2.50E-04 | 8.27E-04 |

| | | | | | |
|---|---|---|---|---|---|
| ENSG00000286596 | | -0.44 | 0.14 | 4.73E-04 | 1.51E-03 |
| ENSG00000118922 | KLF12 | -0.44 | 0.13 | 4.77E-04 | 1.53E-03 |
| ENSG00000189144 | ZNF573 | -0.44 | 0.14 | 7.44E-04 | 2.31E-03 |
| ENSG00000166192 | SENP8 | -0.44 | 0.15 | 1.07E-03 | 3.23E-03 |
| ENSG00000181404 | WASHC1 | -0.44 | 0.15 | 1.26E-03 | 3.77E-03 |
| ENSG00000236901 | MIR600HG | -0.44 | 0.17 | 4.75E-03 | 1.28E-02 |
| ENSG00000102805 | CLN5 | -0.44 | 0.17 | 4.91E-03 | 1.32E-02 |
| ENSG00000264343 | NOTCH2NLA | -0.44 | 0.18 | 6.01E-03 | 1.59E-02 |
| ENSG00000105963 | ADAP1 | -0.44 | 0.18 | 7.67E-03 | 1.99E-02 |
| ENSG00000111581 | NUP107 | -0.43 | 0.04 | 1.92E-25 | 2.20E-24 |
| ENSG00000141027 | NCOR1 | -0.43 | 0.04 | 3.79E-24 | 4.14E-23 |
| ENSG00000138363 | ATIC | -0.43 | 0.04 | 1.25E-23 | 1.34E-22 |
| ENSG00000085998 | POMGNT1 | -0.43 | 0.05 | 1.98E-20 | 1.88E-19 |
| ENSG00000132792 | CTNNBL1 | -0.43 | 0.05 | 3.88E-20 | 3.63E-19 |
| ENSG00000137073 | UBAP2 | -0.43 | 0.05 | 5.39E-20 | 5.01E-19 |
| ENSG00000124222 | STX16 | -0.43 | 0.05 | 7.62E-19 | 6.70E-18 |
| ENSG00000144827 | ABHD10 | -0.43 | 0.05 | 3.51E-18 | 3.00E-17 |
| ENSG00000164715 | LMTK2 | -0.43 | 0.05 | 7.81E-18 | 6.59E-17 |
| ENSG00000178980 | SELENOW | -0.43 | 0.05 | 3.06E-17 | 2.52E-16 |
| ENSG00000068366 | ACSL4 | -0.43 | 0.05 | 3.32E-17 | 2.73E-16 |
| ENSG00000115946 | PNO1 | -0.43 | 0.05 | 3.98E-17 | 3.26E-16 |
| ENSG00000095002 | MSH2 | -0.43 | 0.05 | 1.60E-16 | 1.28E-15 |
| ENSG00000010165 | METTL13 | -0.43 | 0.05 | 5.54E-16 | 4.30E-15 |
| ENSG00000154727 | GABPA | -0.43 | 0.05 | 7.32E-16 | 5.64E-15 |
| ENSG00000089195 | TRMT6 | -0.43 | 0.05 | 3.32E-15 | 2.49E-14 |
| ENSG00000154310 | TNIK | -0.43 | 0.06 | 5.80E-15 | 4.30E-14 |
| ENSG00000120992 | LYPLA1 | -0.43 | 0.06 | 2.64E-14 | 1.90E-13 |
| ENSG00000115942 | ORC2 | -0.43 | 0.06 | 3.99E-14 | 2.83E-13 |
| ENSG00000141232 | TOB1 | -0.43 | 0.06 | 6.28E-14 | 4.41E-13 |
| ENSG00000117408 | IPO13 | -0.43 | 0.06 | 6.47E-14 | 4.54E-13 |
| ENSG00000166348 | USP54 | -0.43 | 0.06 | 7.02E-14 | 4.92E-13 |
| ENSG00000139263 | LRIG3 | -0.43 | 0.06 | 1.10E-13 | 7.63E-13 |
| ENSG00000139436 | GIT2 | -0.43 | 0.06 | 1.21E-13 | 8.38E-13 |
| ENSG00000102178 | UBL4A | -0.43 | 0.06 | 8.12E-13 | 5.36E-12 |
| ENSG00000104133 | SPG11 | -0.43 | 0.06 | 4.39E-12 | 2.77E-11 |
| ENSG00000092094 | OSGEP | -0.43 | 0.06 | 5.91E-12 | 3.69E-11 |
| ENSG00000169083 | AR | -0.43 | 0.06 | 1.36E-11 | 8.32E-11 |
| ENSG00000103319 | EEF2K | -0.43 | 0.06 | 1.96E-11 | 1.19E-10 |
| ENSG00000103197 | TSC2 | -0.43 | 0.07 | 4.59E-11 | 2.72E-10 |
| ENSG00000154917 | RAB6B | -0.43 | 0.07 | 1.38E-10 | 7.94E-10 |
| ENSG00000178033 | CALHM5 | -0.43 | 0.07 | 2.05E-10 | 1.17E-09 |
| ENSG00000164751 | PEX2 | -0.43 | 0.07 | 2.49E-10 | 1.40E-09 |
| ENSG00000147874 | HAUS6 | -0.43 | 0.07 | 2.91E-10 | 1.63E-09 |
| ENSG00000038210 | PI4K2B | -0.43 | 0.07 | 6.61E-10 | 3.62E-09 |
| ENSG00000171132 | PRKCE | -0.43 | 0.07 | 6.82E-10 | 3.73E-09 |
| ENSG00000105948 | TTC26 | -0.43 | 0.07 | 7.52E-10 | 4.10E-09 |
| ENSG00000105793 | GTPBP10 | -0.43 | 0.07 | 1.02E-09 | 5.48E-09 |
| ENSG00000113758 | DBN1 | -0.43 | 0.07 | 2.31E-09 | 1.21E-08 |
| ENSG00000033011 | ALG1 | -0.43 | 0.07 | 2.67E-09 | 1.39E-08 |
| ENSG00000175455 | CCDC14 | -0.43 | 0.08 | 1.05E-08 | 5.28E-08 |

| | | | | | |
|---|---|---|---|---|---|
| ENSG00000172183 | ISG20 | -0.43 | 0.08 | 1.34E-08 | 6.64E-08 |
| ENSG00000130348 | QRSL1 | -0.43 | 0.08 | 2.09E-08 | 1.02E-07 |
| ENSG00000163013 | FBXO41 | -0.43 | 0.08 | 2.32E-08 | 1.13E-07 |
| ENSG00000108821 | COL1A1 | -0.43 | 0.08 | 2.49E-08 | 1.21E-07 |
| ENSG00000291171 | | -0.43 | 0.08 | 3.63E-08 | 1.74E-07 |
| ENSG00000253293 | HOXA10 | -0.43 | 0.08 | 5.98E-08 | 2.82E-07 |
| ENSG00000089682 | RBM41 | -0.43 | 0.08 | 6.14E-08 | 2.89E-07 |
| ENSG00000175575 | PAAF1 | -0.43 | 0.08 | 6.31E-08 | 2.97E-07 |
| ENSG00000167244 | IGF2 | -0.43 | 0.08 | 1.27E-07 | 5.85E-07 |
| ENSG00000122783 | CYREN | -0.43 | 0.09 | 2.76E-07 | 1.23E-06 |
| ENSG00000170315 | UBB | -0.43 | 0.09 | 3.00E-07 | 1.34E-06 |
| ENSG00000156239 | N6AMT1 | -0.43 | 0.09 | 8.93E-07 | 3.80E-06 |
| ENSG00000102078 | SLC25A14 | -0.43 | 0.1 | 3.83E-06 | 1.54E-05 |
| ENSG00000153140 | CETN3 | -0.43 | 0.1 | 6.40E-06 | 2.52E-05 |
| ENSG00000103245 | CIAO3 | -0.43 | 0.1 | 7.90E-06 | 3.08E-05 |
| ENSG00000025434 | NR1H3 | -0.43 | 0.11 | 3.41E-05 | 1.25E-04 |
| ENSG00000168938 | PPIC | -0.43 | 0.12 | 1.02E-04 | 3.54E-04 |
| ENSG00000022976 | ZNF839 | -0.43 | 0.12 | 1.02E-04 | 3.54E-04 |
| ENSG00000285517 | | -0.43 | 0.12 | 1.68E-04 | 5.69E-04 |
| ENSG00000099785 | MARCHF2 | -0.43 | 0.12 | 1.77E-04 | 5.97E-04 |
| ENSG00000046647 | GEMIN8 | -0.43 | 0.13 | 4.36E-04 | 1.40E-03 |
| ENSG00000221886 | FAM200C | -0.43 | 0.14 | 9.36E-04 | 2.86E-03 |
| ENSG00000117242 | PINK1-AS | -0.43 | 0.16 | 2.76E-03 | 7.80E-03 |
| ENSG00000120756 | PLS1 | -0.43 | 0.16 | 4.14E-03 | 1.13E-02 |
| ENSG00000263731 | | -0.43 | 0.17 | 5.21E-03 | 1.40E-02 |
| ENSG00000258183 | LINC02392 | -0.43 | 0.18 | 7.20E-03 | 1.88E-02 |
| ENSG00000184029 | DSCR4 | -0.43 | 0.19 | 1.05E-02 | 2.64E-02 |
| ENSG00000147642 | SYBU | -0.43 | 0.21 | 1.74E-02 | 4.15E-02 |
| ENSG00000144028 | SNRNP200 | -0.42 | 0.03 | 1.76E-39 | 3.13E-38 |
| ENSG00000188517 | COL25A1 | -0.42 | 0.04 | 6.38E-32 | 9.10E-31 |
| ENSG00000116962 | NID1 | -0.42 | 0.04 | 5.28E-26 | 6.17E-25 |
| ENSG00000148848 | ADAM12 | -0.42 | 0.04 | 5.80E-23 | 6.07E-22 |
| ENSG00000165283 | STOML2 | -0.42 | 0.05 | 5.58E-21 | 5.41E-20 |
| ENSG00000134014 | ELP3 | -0.42 | 0.05 | 1.01E-19 | 9.31E-19 |
| ENSG00000167202 | TBC1D2B | -0.42 | 0.05 | 1.43E-19 | 1.30E-18 |
| ENSG00000133313 | CNDP2 | -0.42 | 0.05 | 1.50E-17 | 1.25E-16 |
| ENSG00000152700 | SAR1B | -0.42 | 0.05 | 4.93E-17 | 4.03E-16 |
| ENSG00000100345 | MYH9 | -0.42 | 0.05 | 1.50E-16 | 1.20E-15 |
| ENSG00000122779 | TRIM24 | -0.42 | 0.05 | 2.40E-16 | 1.90E-15 |
| ENSG00000148019 | CEP78 | -0.42 | 0.05 | 2.62E-16 | 2.07E-15 |
| ENSG00000169762 | TAPT1 | -0.42 | 0.05 | 2.25E-15 | 1.70E-14 |
| ENSG00000107949 | BCCIP | -0.42 | 0.05 | 1.21E-14 | 8.86E-14 |
| ENSG00000168827 | GFM1 | -0.42 | 0.06 | 3.21E-14 | 2.29E-13 |
| ENSG00000198795 | ZNF521 | -0.42 | 0.06 | 3.32E-14 | 2.37E-13 |
| ENSG00000188706 | ZDHHC9 | -0.42 | 0.06 | 4.40E-14 | 3.11E-13 |
| ENSG00000117385 | P3H1 | -0.42 | 0.06 | 4.45E-14 | 3.15E-13 |
| ENSG00000105048 | TNNT1 | -0.42 | 0.06 | 5.03E-14 | 3.55E-13 |
| ENSG00000179409 | GEMIN4 | -0.42 | 0.06 | 6.25E-14 | 4.39E-13 |
| ENSG00000122034 | GTF3A | -0.42 | 0.06 | 2.16E-13 | 1.47E-12 |
| ENSG00000116001 | TIA1 | -0.42 | 0.06 | 1.76E-12 | 1.14E-11 |

| | | | | | |
|---|---|---|---|---|---|
| ENSG00000131732 | ZCCHC9 | -0.42 | 0.06 | 4.80E-12 | 3.02E-11 |
| ENSG00000049883 | PTCD2 | -0.42 | 0.06 | 1.23E-11 | 7.57E-11 |
| ENSG00000099622 | CIRBP | -0.42 | 0.06 | 1.38E-11 | 8.45E-11 |
| ENSG00000181704 | YIPF6 | -0.42 | 0.06 | 5.64E-11 | 3.32E-10 |
| ENSG00000174206 | KICS2 | -0.42 | 0.06 | 7.11E-11 | 4.16E-10 |
| ENSG00000066629 | EML1 | -0.42 | 0.06 | 8.00E-11 | 4.67E-10 |
| ENSG00000138442 | WDR12 | -0.42 | 0.07 | 9.43E-11 | 5.48E-10 |
| ENSG00000137074 | APTX | -0.42 | 0.07 | 1.08E-10 | 6.25E-10 |
| ENSG00000070950 | RAD18 | -0.42 | 0.07 | 1.36E-10 | 7.84E-10 |
| ENSG00000180447 | GAS1 | -0.42 | 0.07 | 1.72E-10 | 9.84E-10 |
| ENSG00000090470 | PDCD7 | -0.42 | 0.07 | 4.46E-10 | 2.47E-09 |
| ENSG00000138035 | PNPT1 | -0.42 | 0.07 | 8.28E-10 | 4.49E-09 |
| ENSG00000131462 | TUBG1 | -0.42 | 0.07 | 1.47E-09 | 7.86E-09 |
| ENSG00000116194 | ANGPTL1 | -0.42 | 0.07 | 3.18E-09 | 1.65E-08 |
| ENSG00000150867 | PIP4K2A | -0.42 | 0.07 | 3.37E-09 | 1.75E-08 |
| ENSG00000046653 | GPM6B | -0.42 | 0.07 | 5.32E-09 | 2.71E-08 |
| ENSG00000154240 | CEP112 | -0.42 | 0.07 | 5.76E-09 | 2.93E-08 |
| ENSG00000105483 | CARD8 | -0.42 | 0.07 | 6.00E-09 | 3.06E-08 |
| ENSG00000185238 | PRMT3 | -0.42 | 0.07 | 7.49E-09 | 3.78E-08 |
| ENSG00000154814 | OXNAD1 | -0.42 | 0.08 | 1.33E-08 | 6.60E-08 |
| ENSG00000079387 | SENP1 | -0.42 | 0.08 | 5.85E-08 | 2.76E-07 |
| ENSG00000128463 | EMC4 | -0.42 | 0.08 | 1.66E-07 | 7.54E-07 |
| ENSG00000271122 | HERPUD2-AS1 | -0.42 | 0.08 | 2.55E-07 | 1.14E-06 |
| ENSG00000203995 | ZYG11A | -0.42 | 0.09 | 4.88E-07 | 2.14E-06 |
| ENSG00000102781 | KATNAL1 | -0.42 | 0.09 | 6.02E-07 | 2.62E-06 |
| ENSG00000161048 | NAPEPLD | -0.42 | 0.09 | 7.15E-07 | 3.08E-06 |
| ENSG00000228672 | PROB1 | -0.42 | 0.09 | 8.62E-07 | 3.68E-06 |
| ENSG00000100577 | GSTZ1 | -0.42 | 0.09 | 1.25E-06 | 5.27E-06 |
| ENSG00000121310 | ECHDC2 | -0.42 | 0.09 | 1.51E-06 | 6.30E-06 |
| ENSG00000280071 | | -0.42 | 0.09 | 2.46E-06 | 1.01E-05 |
| ENSG00000112365 | ZBTB24 | -0.42 | 0.09 | 3.39E-06 | 1.37E-05 |
| ENSG00000149679 | CABLES2 | -0.42 | 0.09 | 4.08E-06 | 1.64E-05 |
| ENSG00000180891 | CUEDC1 | -0.42 | 0.1 | 1.12E-05 | 4.29E-05 |
| ENSG00000198719 | DLL1 | -0.42 | 0.1 | 1.70E-05 | 6.42E-05 |
| ENSG00000227036 | LINC00511 | -0.42 | 0.11 | 2.98E-05 | 1.10E-04 |
| ENSG00000117676 | RPS6KA1 | -0.42 | 0.11 | 7.65E-05 | 2.70E-04 |
| ENSG00000135835 | KIAA1614 | -0.42 | 0.11 | 9.47E-05 | 3.30E-04 |
| ENSG00000144792 | ZNF660 | -0.42 | 0.12 | 2.32E-04 | 7.72E-04 |
| ENSG00000104081 | BMF | -0.42 | 0.13 | 8.99E-04 | 2.75E-03 |
| ENSG00000233101 | HOXB-AS3 | -0.42 | 0.15 | 2.07E-03 | 5.98E-03 |
| ENSG00000117791 | MTARC2 | -0.42 | 0.15 | 2.81E-03 | 7.94E-03 |
| ENSG00000176490 | DIRAS1 | -0.42 | 0.16 | 2.93E-03 | 8.25E-03 |
| ENSG00000100842 | EFS | -0.42 | 0.16 | 3.06E-03 | 8.58E-03 |
| ENSG00000196427 | NBPF4 | -0.42 | 0.16 | 3.33E-03 | 9.27E-03 |
| ENSG00000133401 | PDZD2 | -0.42 | 0.16 | 3.38E-03 | 9.41E-03 |
| ENSG00000140678 | ITGAX | -0.42 | 0.16 | 3.88E-03 | 1.07E-02 |
| ENSG00000230896 | | -0.42 | 0.17 | 6.97E-03 | 1.82E-02 |
| ENSG00000112293 | GPLD1 | -0.42 | 0.19 | 1.34E-02 | 3.28E-02 |
| ENSG00000011590 | ZBTB32 | -0.42 | 0.19 | 1.38E-02 | 3.38E-02 |
| ENSG00000290927 | | -0.42 | 0.2 | 1.47E-02 | 3.57E-02 |

| | | | | | |
|---|---|---|---|---|---|
| ENSG00000168495 | POLR3D | -0.41 | 0.04 | 1.37E-26 | 1.64E-25 |
| ENSG00000146223 | RPL7L1 | -0.41 | 0.04 | 1.29E-25 | 1.49E-24 |
| ENSG00000189159 | JPT1 | -0.41 | 0.04 | 2.53E-25 | 2.89E-24 |
| ENSG00000135829 | DHX9 | -0.41 | 0.05 | 2.48E-19 | 2.24E-18 |
| ENSG00000132434 | LANCL2 | -0.41 | 0.05 | 2.90E-17 | 2.40E-16 |
| ENSG00000007062 | PROM1 | -0.41 | 0.05 | 2.92E-17 | 2.41E-16 |
| ENSG00000122729 | ACO1 | -0.41 | 0.05 | 6.66E-17 | 5.42E-16 |
| ENSG00000153187 | HNRNPU | -0.41 | 0.05 | 7.71E-17 | 6.25E-16 |
| ENSG00000219481 | NBPF1 | -0.41 | 0.05 | 1.50E-16 | 1.20E-15 |
| ENSG00000075884 | ARHGAP15 | -0.41 | 0.05 | 5.44E-15 | 4.03E-14 |
| ENSG00000142208 | AKT1 | -0.41 | 0.06 | 2.89E-13 | 1.96E-12 |
| ENSG00000163348 | PYGO2 | -0.41 | 0.06 | 1.06E-12 | 6.97E-12 |
| ENSG00000146842 | TMEM209 | -0.41 | 0.06 | 1.93E-12 | 1.24E-11 |
| ENSG00000173065 | FAM222B | -0.41 | 0.06 | 4.77E-12 | 3.00E-11 |
| ENSG00000198142 | SOWAHC | -0.41 | 0.06 | 2.97E-11 | 1.78E-10 |
| ENSG00000134900 | TPP2 | -0.41 | 0.06 | 4.17E-11 | 2.47E-10 |
| ENSG00000155329 | ZCCHC10 | -0.41 | 0.06 | 4.86E-11 | 2.87E-10 |
| ENSG00000143363 | PRUNE1 | -0.41 | 0.06 | 1.01E-10 | 5.86E-10 |
| ENSG00000115107 | STEAP3 | -0.41 | 0.07 | 3.05E-10 | 1.71E-09 |
| ENSG00000005175 | RPAP3 | -0.41 | 0.07 | 4.18E-10 | 2.32E-09 |
| ENSG00000105971 | CAV2 | -0.41 | 0.07 | 4.28E-10 | 2.38E-09 |
| ENSG00000088247 | KHSRP | -0.41 | 0.07 | 4.73E-10 | 2.61E-09 |
| ENSG00000180340 | FZD2 | -0.41 | 0.07 | 7.17E-10 | 3.91E-09 |
| ENSG00000135778 | NTPCR | -0.41 | 0.07 | 2.38E-09 | 1.25E-08 |
| ENSG00000078070 | MCCC1 | -0.41 | 0.07 | 2.88E-09 | 1.50E-08 |
| ENSG00000100324 | TAB1 | -0.41 | 0.07 | 3.87E-09 | 2.00E-08 |
| ENSG00000104936 | DMPK | -0.41 | 0.07 | 4.32E-09 | 2.22E-08 |
| ENSG00000161267 | BDH1 | -0.41 | 0.07 | 7.37E-09 | 3.72E-08 |
| ENSG00000134597 | RBMX2 | -0.41 | 0.07 | 1.08E-08 | 5.40E-08 |
| ENSG00000149499 | EML3 | -0.41 | 0.07 | 1.66E-08 | 8.16E-08 |
| ENSG00000166024 | R3HCC1L | -0.41 | 0.08 | 3.18E-08 | 1.53E-07 |
| ENSG00000002016 | RAD52 | -0.41 | 0.08 | 3.78E-08 | 1.81E-07 |
| ENSG00000146707 | POMZP3 | -0.41 | 0.08 | 3.18E-07 | 1.42E-06 |
| ENSG00000140950 | MEAK7 | -0.41 | 0.08 | 5.13E-07 | 2.24E-06 |
| ENSG00000148429 | USP6NL | -0.41 | 0.09 | 9.70E-07 | 4.12E-06 |
| ENSG00000143537 | ADAM15 | -0.41 | 0.09 | 1.08E-06 | 4.58E-06 |
| ENSG00000121988 | ZRANB3 | -0.41 | 0.09 | 1.10E-06 | 4.66E-06 |
| ENSG00000126705 | AHDC1 | -0.41 | 0.09 | 1.89E-06 | 7.81E-06 |
| ENSG00000152457 | DCLRE1C | -0.41 | 0.09 | 2.58E-06 | 1.05E-05 |
| ENSG00000185880 | TRIM69 | -0.41 | 0.09 | 2.91E-06 | 1.18E-05 |
| ENSG00000181007 | ZFP82 | -0.41 | 0.09 | 3.89E-06 | 1.57E-05 |
| ENSG00000130881 | LRP3 | -0.41 | 0.09 | 6.18E-06 | 2.44E-05 |
| ENSG00000196417 | ZNF765 | -0.41 | 0.09 | 6.23E-06 | 2.46E-05 |
| ENSG00000167264 | DUS2 | -0.41 | 0.1 | 1.62E-05 | 6.13E-05 |
| ENSG00000115459 | ELMOD3 | -0.41 | 0.1 | 1.71E-05 | 6.44E-05 |
| ENSG00000100209 | HSCB | -0.41 | 0.1 | 2.94E-05 | 1.08E-04 |
| ENSG00000157551 | KCNJ15 | -0.41 | 0.1 | 2.98E-05 | 1.10E-04 |
| ENSG00000095370 | SH2D3C | -0.41 | 0.1 | 3.08E-05 | 1.13E-04 |
| ENSG00000112039 | FANCE | -0.41 | 0.1 | 3.24E-05 | 1.19E-04 |
| ENSG00000105997 | HOXA3 | -0.41 | 0.14 | 1.47E-03 | 4.35E-03 |

| | | | | | |
|---|---|---|---|---|---|
| ENSG00000178035 | IMPDH2 | -0.41 | 0.14 | 1.71E-03 | 5.02E-03 |
| ENSG00000164362 | TERT | -0.41 | 0.16 | 5.28E-03 | 1.42E-02 |
| ENSG00000289294 | | -0.41 | 0.18 | 9.84E-03 | 2.49E-02 |
| ENSG00000188060 | RAB42 | -0.41 | 0.18 | 1.03E-02 | 2.59E-02 |
| ENSG00000159307 | SCUBE1 | -0.41 | 0.18 | 1.04E-02 | 2.61E-02 |
| ENSG00000160219 | GAB3 | -0.41 | 0.18 | 1.14E-02 | 2.84E-02 |
| ENSG00000214944 | ARHGEF28 | -0.41 | 0.18 | 1.22E-02 | 3.01E-02 |
| ENSG00000289436 | | -0.41 | 0.19 | 1.52E-02 | 3.67E-02 |
| ENSG00000103342 | GSPT1 | -0.4 | 0.04 | 3.40E-28 | 4.28E-27 |
| ENSG00000092820 | EZR | -0.4 | 0.04 | 1.26E-26 | 1.51E-25 |
| ENSG00000072501 | SMC1A | -0.4 | 0.04 | 1.99E-24 | 2.20E-23 |
| ENSG00000133687 | TMTC1 | -0.4 | 0.04 | 2.62E-24 | 2.87E-23 |
| ENSG00000107372 | ZFAND5 | -0.4 | 0.04 | 2.01E-23 | 2.13E-22 |
| ENSG00000182481 | KPNA2 | -0.4 | 0.04 | 2.08E-23 | 2.21E-22 |
| ENSG00000185650 | ZFP36L1 | -0.4 | 0.04 | 2.10E-20 | 1.99E-19 |
| ENSG00000112062 | MAPK14 | -0.4 | 0.04 | 1.03E-19 | 9.48E-19 |
| ENSG00000135900 | MRPL44 | -0.4 | 0.05 | 1.35E-18 | 1.18E-17 |
| ENSG00000166557 | TMED3 | -0.4 | 0.05 | 2.41E-18 | 2.07E-17 |
| ENSG00000169432 | SCN9A | -0.4 | 0.05 | 2.83E-17 | 2.34E-16 |
| ENSG00000160194 | NDUFV3 | -0.4 | 0.05 | 2.94E-17 | 2.42E-16 |
| ENSG00000138279 | ANXA7 | -0.4 | 0.05 | 3.29E-17 | 2.70E-16 |
| ENSG00000124207 | CSE1L | -0.4 | 0.05 | 5.41E-17 | 4.41E-16 |
| ENSG00000156052 | GNAQ | -0.4 | 0.05 | 7.50E-17 | 6.08E-16 |
| ENSG00000156304 | SCAF4 | -0.4 | 0.05 | 8.26E-17 | 6.68E-16 |
| ENSG00000136643 | RPS6KC1 | -0.4 | 0.05 | 1.31E-16 | 1.05E-15 |
| ENSG00000132604 | TERF2 | -0.4 | 0.05 | 6.61E-16 | 5.12E-15 |
| ENSG00000064313 | TAF2 | -0.4 | 0.05 | 9.69E-16 | 7.41E-15 |
| ENSG00000165072 | MAMDC2 | -0.4 | 0.05 | 6.01E-15 | 4.45E-14 |
| ENSG00000082146 | STRADB | -0.4 | 0.05 | 2.69E-14 | 1.93E-13 |
| ENSG00000137802 | MAPKBP1 | -0.4 | 0.05 | 6.13E-14 | 4.31E-13 |
| ENSG00000105953 | OGDH | -0.4 | 0.05 | 9.39E-14 | 6.55E-13 |
| ENSG00000177646 | ACAD9 | -0.4 | 0.06 | 4.03E-13 | 2.71E-12 |
| ENSG00000173273 | TNKS | -0.4 | 0.06 | 4.90E-13 | 3.27E-12 |
| ENSG00000183579 | ZNRF3 | -0.4 | 0.06 | 7.49E-13 | 4.95E-12 |
| ENSG00000168172 | HOOK3 | -0.4 | 0.06 | 1.78E-12 | 1.15E-11 |
| ENSG00000070495 | JMJD6 | -0.4 | 0.06 | 2.39E-12 | 1.53E-11 |
| ENSG00000214022 | REPIN1 | -0.4 | 0.06 | 4.06E-12 | 2.57E-11 |
| ENSG00000114054 | PCCB | -0.4 | 0.06 | 1.41E-11 | 8.61E-11 |
| ENSG00000091527 | CDV3 | -0.4 | 0.06 | 3.46E-11 | 2.06E-10 |
| ENSG00000173545 | ZNF622 | -0.4 | 0.06 | 3.76E-11 | 2.24E-10 |
| ENSG00000149182 | ARFGAP2 | -0.4 | 0.06 | 1.37E-10 | 7.90E-10 |
| ENSG00000106266 | SNX8 | -0.4 | 0.06 | 1.98E-10 | 1.13E-09 |
| ENSG00000180104 | EXOC3 | -0.4 | 0.07 | 7.55E-10 | 4.11E-09 |
| ENSG00000127995 | CASD1 | -0.4 | 0.07 | 1.18E-09 | 6.33E-09 |
| ENSG00000138780 | GSTCD | -0.4 | 0.07 | 1.36E-09 | 7.27E-09 |
| ENSG00000156983 | BRPF1 | -0.4 | 0.07 | 2.37E-09 | 1.24E-08 |
| ENSG00000059588 | TARBP1 | -0.4 | 0.07 | 2.64E-09 | 1.38E-08 |
| ENSG00000101846 | STS | -0.4 | 0.07 | 3.13E-09 | 1.63E-08 |
| ENSG00000160410 | SHKBP1 | -0.4 | 0.07 | 3.44E-09 | 1.78E-08 |
| ENSG00000112343 | TRIM38 | -0.4 | 0.07 | 5.19E-09 | 2.65E-08 |

| | | | | | |
|---|---|---|---|---|---|
| ENSG00000103932 | RPAP1 | -0.4 | 0.07 | 5.47E-09 | 2.79E-08 |
| ENSG00000150347 | ARID5B | -0.4 | 0.07 | 5.59E-09 | 2.85E-08 |
| ENSG00000007944 | MYLIP | -0.4 | 0.07 | 1.77E-08 | 8.72E-08 |
| ENSG00000086504 | MRPL28 | -0.4 | 0.07 | 2.15E-08 | 1.05E-07 |
| ENSG00000178585 | CTNNBIP1 | -0.4 | 0.07 | 2.39E-08 | 1.16E-07 |
| ENSG00000063241 | ISOC2 | -0.4 | 0.07 | 3.33E-08 | 1.60E-07 |
| ENSG00000164430 | CGAS | -0.4 | 0.08 | 7.40E-08 | 3.47E-07 |
| ENSG00000083720 | OXCT1 | -0.4 | 0.08 | 1.01E-07 | 4.69E-07 |
| ENSG00000168803 | ADAL | -0.4 | 0.08 | 1.45E-07 | 6.63E-07 |
| ENSG00000138756 | BMP2K | -0.4 | 0.08 | 1.52E-07 | 6.93E-07 |
| ENSG00000188313 | PLSCR1 | -0.4 | 0.08 | 1.57E-07 | 7.18E-07 |
| ENSG00000171530 | TBCA | -0.4 | 0.08 | 2.68E-07 | 1.20E-06 |
| ENSG00000158691 | ZSCAN12 | -0.4 | 0.08 | 4.48E-07 | 1.97E-06 |
| ENSG00000160767 | ENTREP3 | -0.4 | 0.08 | 4.54E-07 | 1.99E-06 |
| ENSG00000119547 | ONECUT2 | -0.4 | 0.08 | 5.22E-07 | 2.28E-06 |
| ENSG00000171943 | SRGAP2C | -0.4 | 0.08 | 5.29E-07 | 2.31E-06 |
| ENSG00000072195 | SPEG | -0.4 | 0.08 | 5.58E-07 | 2.43E-06 |
| ENSG00000173171 | MTX1 | -0.4 | 0.08 | 9.12E-07 | 3.89E-06 |
| ENSG00000079150 | FKBP7 | -0.4 | 0.09 | 1.09E-06 | 4.62E-06 |
| ENSG00000184979 | USP18 | -0.4 | 0.09 | 1.24E-06 | 5.20E-06 |
| ENSG00000166199 | ALKBH3 | -0.4 | 0.09 | 3.21E-06 | 1.30E-05 |
| ENSG00000056050 | HPF1 | -0.4 | 0.09 | 3.39E-06 | 1.37E-05 |
| ENSG00000148459 | PDSS1 | -0.4 | 0.09 | 4.84E-06 | 1.93E-05 |
| ENSG00000188343 | CIBAR1 | -0.4 | 0.09 | 6.48E-06 | 2.55E-05 |
| ENSG00000198081 | ZBTB14 | -0.4 | 0.1 | 1.54E-05 | 5.85E-05 |
| ENSG00000064652 | SNX24 | -0.4 | 0.1 | 1.88E-05 | 7.07E-05 |
| ENSG00000236088 | COX10-DT | -0.4 | 0.1 | 2.85E-05 | 1.05E-04 |
| ENSG00000100564 | PIGH | -0.4 | 0.1 | 6.17E-05 | 2.20E-04 |
| ENSG00000111860 | CEP85L | -0.4 | 0.1 | 6.34E-05 | 2.26E-04 |
| ENSG00000162882 | HAAO | -0.4 | 0.11 | 1.60E-04 | 5.42E-04 |
| ENSG00000065057 | NTHL1 | -0.4 | 0.11 | 2.26E-04 | 7.53E-04 |
| ENSG00000213963 | | -0.4 | 0.12 | 2.64E-04 | 8.72E-04 |
| ENSG00000090776 | EFNB1 | -0.4 | 0.12 | 3.12E-04 | 1.02E-03 |
| ENSG00000155666 | KDM8 | -0.4 | 0.12 | 4.49E-04 | 1.44E-03 |
| ENSG00000177051 | FBXO46 | -0.4 | 0.12 | 6.10E-04 | 1.92E-03 |
| ENSG00000170412 | GPRC5C | -0.4 | 0.13 | 8.09E-04 | 2.50E-03 |
| ENSG00000158089 | GALNT14 | -0.4 | 0.13 | 9.12E-04 | 2.79E-03 |
| ENSG00000104369 | JPH1 | -0.4 | 0.13 | 9.86E-04 | 3.00E-03 |
| ENSG00000167216 | KATNAL2 | -0.4 | 0.13 | 1.02E-03 | 3.08E-03 |
| ENSG00000183337 | BCOR | -0.4 | 0.13 | 1.18E-03 | 3.55E-03 |
| ENSG00000186684 | CYP27C1 | -0.4 | 0.14 | 2.71E-03 | 7.67E-03 |
| ENSG00000227496 | | -0.4 | 0.16 | 5.60E-03 | 1.49E-02 |
| ENSG00000111254 | AKAP3 | -0.4 | 0.16 | 7.04E-03 | 1.84E-02 |
| ENSG00000288924 | | -0.4 | 0.17 | 1.09E-02 | 2.72E-02 |
| ENSG00000158615 | PPP1R15B | -0.39 | 0.04 | 4.70E-26 | 5.51E-25 |
| ENSG00000117713 | ARID1A | -0.39 | 0.04 | 5.53E-25 | 6.24E-24 |
| ENSG00000154518 | ATP5MC3 | -0.39 | 0.04 | 6.83E-23 | 7.11E-22 |
| ENSG00000135912 | TTLL4 | -0.39 | 0.04 | 1.02E-21 | 1.02E-20 |
| ENSG00000130826 | DKC1 | -0.39 | 0.04 | 7.43E-19 | 6.54E-18 |
| ENSG00000080371 | RAB21 | -0.39 | 0.04 | 7.61E-19 | 6.69E-18 |

| | | | | | |
|---|---|---|---|---|---|
| ENSG00000168256 | NKIRAS2 | -0.39 | 0.05 | 3.25E-18 | 2.79E-17 |
| ENSG00000181104 | F2R | -0.39 | 0.04 | 3.29E-18 | 2.82E-17 |
| ENSG00000198815 | FOXJ3 | -0.39 | 0.05 | 5.92E-18 | 5.02E-17 |
| ENSG00000104388 | RAB2A | -0.39 | 0.05 | 6.84E-18 | 5.78E-17 |
| ENSG00000132561 | MATN2 | -0.39 | 0.05 | 2.15E-15 | 1.63E-14 |
| ENSG00000114978 | MOB1A | -0.39 | 0.05 | 6.32E-15 | 4.68E-14 |
| ENSG00000119326 | CTNNAL1 | -0.39 | 0.05 | 1.41E-14 | 1.02E-13 |
| ENSG00000180530 | NRIP1 | -0.39 | 0.05 | 9.91E-14 | 6.90E-13 |
| ENSG00000117597 | UTP25 | -0.39 | 0.05 | 1.93E-13 | 1.32E-12 |
| ENSG00000127527 | EPS15L1 | -0.39 | 0.05 | 5.77E-13 | 3.85E-12 |
| ENSG00000075292 | ZNF638 | -0.39 | 0.06 | 1.77E-12 | 1.15E-11 |
| ENSG00000196535 | MYO18A | -0.39 | 0.06 | 3.74E-12 | 2.37E-11 |
| ENSG00000173821 | RNF213 | -0.39 | 0.06 | 6.84E-12 | 4.25E-11 |
| ENSG00000115159 | GPD2 | -0.39 | 0.06 | 6.98E-12 | 4.34E-11 |
| ENSG00000164597 | COG5 | -0.39 | 0.06 | 7.64E-12 | 4.74E-11 |
| ENSG00000196705 | ZNF431 | -0.39 | 0.06 | 1.53E-11 | 9.35E-11 |
| ENSG00000072682 | P4HA2 | -0.39 | 0.06 | 2.38E-11 | 1.44E-10 |
| ENSG00000125676 | THOC2 | -0.39 | 0.06 | 3.71E-11 | 2.21E-10 |
| ENSG00000172500 | FIBP | -0.39 | 0.06 | 3.96E-11 | 2.35E-10 |
| ENSG00000163950 | SLBP | -0.39 | 0.06 | 4.05E-11 | 2.40E-10 |
| ENSG00000198690 | FAN1 | -0.39 | 0.06 | 5.47E-11 | 3.22E-10 |
| ENSG00000162623 | TYW3 | -0.39 | 0.07 | 8.85E-10 | 4.79E-09 |
| ENSG00000171467 | ZNF318 | -0.39 | 0.07 | 1.66E-09 | 8.83E-09 |
| ENSG00000184584 | STING1 | -0.39 | 0.07 | 1.96E-09 | 1.04E-08 |
| ENSG00000152749 | GPR180 | -0.39 | 0.07 | 2.28E-09 | 1.20E-08 |
| ENSG00000107951 | MTPAP | -0.39 | 0.07 | 6.07E-08 | 2.86E-07 |
| ENSG00000186918 | ZNF395 | -0.39 | 0.08 | 2.05E-07 | 9.28E-07 |
| ENSG00000144161 | ZC3H8 | -0.39 | 0.08 | 3.20E-07 | 1.42E-06 |
| ENSG00000095321 | CRAT | -0.39 | 0.08 | 3.21E-07 | 1.43E-06 |
| ENSG00000151229 | SLC2A13 | -0.39 | 0.08 | 4.33E-07 | 1.91E-06 |
| ENSG00000187325 | TAF9B | -0.39 | 0.08 | 7.00E-07 | 3.02E-06 |
| ENSG00000155330 | C16orf87 | -0.39 | 0.08 | 7.40E-07 | 3.18E-06 |
| ENSG00000162522 | KIAA1522 | -0.39 | 0.08 | 7.80E-07 | 3.35E-06 |
| ENSG00000170577 | SIX2 | -0.39 | 0.08 | 8.47E-07 | 3.62E-06 |
| ENSG00000171791 | BCL2 | -0.39 | 0.08 | 9.11E-07 | 3.88E-06 |
| ENSG00000013364 | MVP | -0.39 | 0.08 | 1.21E-06 | 5.10E-06 |
| ENSG00000290445 | CCNYL2 | -0.39 | 0.09 | 2.57E-06 | 1.05E-05 |
| ENSG00000162650 | ATXN7L2 | -0.39 | 0.09 | 2.87E-06 | 1.17E-05 |
| ENSG00000158292 | GPR153 | -0.39 | 0.09 | 2.95E-06 | 1.20E-05 |
| ENSG00000121481 | RNF2 | -0.39 | 0.09 | 3.06E-06 | 1.24E-05 |
| ENSG00000132199 | ENOSF1 | -0.39 | 0.09 | 3.19E-06 | 1.29E-05 |
| ENSG00000130529 | TRPM4 | -0.39 | 0.09 | 6.59E-06 | 2.60E-05 |
| ENSG00000196689 | TRPV1 | -0.39 | 0.09 | 9.25E-06 | 3.59E-05 |
| ENSG00000168476 | REEP4 | -0.39 | 0.09 | 1.03E-05 | 3.96E-05 |
| ENSG00000147118 | ZNF182 | -0.39 | 0.09 | 1.37E-05 | 5.23E-05 |
| ENSG00000134504 | KCTD1 | -0.39 | 0.09 | 1.56E-05 | 5.93E-05 |
| ENSG00000179094 | PER1 | -0.39 | 0.1 | 3.94E-05 | 1.43E-04 |
| ENSG00000164638 | SLC29A4 | -0.39 | 0.1 | 7.61E-05 | 2.68E-04 |
| ENSG00000120860 | WASHC3 | -0.39 | 0.1 | 1.25E-04 | 4.30E-04 |
| ENSG00000168152 | THAP9 | -0.39 | 0.11 | 1.28E-04 | 4.40E-04 |

| | | | | | |
|---|---|---|---|---|---|
| ENSG00000166394 | CYB5R2 | -0.39 | 0.13 | 1.36E-03 | 4.03E-03 |
| ENSG00000198721 | ECI2 | -0.39 | 0.14 | 2.32E-03 | 6.66E-03 |
| ENSG00000135338 | LCA5 | -0.39 | 0.15 | 3.63E-03 | 1.00E-02 |
| ENSG00000005249 | PRKAR2B | -0.39 | 0.16 | 7.77E-03 | 2.01E-02 |
| ENSG00000196456 | ZNF775 | -0.39 | 0.17 | 8.98E-03 | 2.30E-02 |
| ENSG00000249853 | HS3ST5 | -0.39 | 0.17 | 9.15E-03 | 2.33E-02 |
| ENSG00000257135 | ODC1-DT | -0.39 | 0.18 | 1.70E-02 | 4.07E-02 |
| ENSG00000205937 | RNPS1 | -0.38 | 0.04 | 2.37E-25 | 2.71E-24 |
| ENSG00000168710 | AHCYL1 | -0.38 | 0.04 | 2.63E-19 | 2.37E-18 |
| ENSG00000172845 | SP3 | -0.38 | 0.04 | 1.62E-17 | 1.35E-16 |
| ENSG00000004478 | FKBP4 | -0.38 | 0.05 | 2.22E-13 | 1.51E-12 |
| ENSG00000084774 | CAD | -0.38 | 0.05 | 7.41E-13 | 4.90E-12 |
| ENSG00000035681 | NSMAF | -0.38 | 0.05 | 1.47E-12 | 9.55E-12 |
| ENSG00000186660 | ZFP91 | -0.38 | 0.05 | 1.49E-12 | 9.66E-12 |
| ENSG00000159842 | ABR | -0.38 | 0.06 | 3.52E-12 | 2.24E-11 |
| ENSG00000106066 | CPVL | -0.38 | 0.06 | 3.81E-12 | 2.41E-11 |
| ENSG00000115539 | PDCL3 | -0.38 | 0.06 | 6.20E-12 | 3.87E-11 |
| ENSG00000008513 | ST3GAL1 | -0.38 | 0.06 | 1.30E-11 | 7.95E-11 |
| ENSG00000105576 | TNPO2 | -0.38 | 0.06 | 2.35E-11 | 1.42E-10 |
| ENSG00000122512 | PMS2 | -0.38 | 0.06 | 6.03E-11 | 3.55E-10 |
| ENSG00000090376 | IRAK3 | -0.38 | 0.06 | 6.58E-11 | 3.85E-10 |
| ENSG00000100325 | ASCC2 | -0.38 | 0.06 | 1.25E-10 | 7.21E-10 |
| ENSG00000256269 | HMBS | -0.38 | 0.06 | 1.50E-10 | 8.58E-10 |
| ENSG00000103275 | UBE2I | -0.38 | 0.06 | 1.95E-10 | 1.11E-09 |
| ENSG00000213625 | LEPROT | -0.38 | 0.06 | 3.59E-10 | 2.00E-09 |
| ENSG00000102755 | FLT1 | -0.38 | 0.06 | 6.24E-10 | 3.42E-09 |
| ENSG00000157181 | ODR4 | -0.38 | 0.06 | 2.17E-09 | 1.14E-08 |
| ENSG00000129055 | ANAPC13 | -0.38 | 0.06 | 2.27E-09 | 1.19E-08 |
| ENSG00000110074 | FOXRED1 | -0.38 | 0.07 | 3.67E-09 | 1.90E-08 |
| ENSG00000146414 | SHPRH | -0.38 | 0.07 | 5.57E-09 | 2.84E-08 |
| ENSG00000141540 | TTYH2 | -0.38 | 0.07 | 5.68E-09 | 2.89E-08 |
| ENSG00000167461 | RAB8A | -0.38 | 0.07 | 1.51E-08 | 7.47E-08 |
| ENSG00000156162 | DPY19L4 | -0.38 | 0.07 | 3.12E-08 | 1.50E-07 |
| ENSG00000171729 | TMEM51 | -0.38 | 0.07 | 3.14E-08 | 1.51E-07 |
| ENSG00000157617 | C2CD2 | -0.38 | 0.07 | 3.53E-08 | 1.69E-07 |
| ENSG00000177733 | HNRNPA0 | -0.38 | 0.07 | 1.82E-07 | 8.29E-07 |
| ENSG00000183386 | FHL3 | -0.38 | 0.08 | 3.17E-07 | 1.41E-06 |
| ENSG00000136874 | STX17 | -0.38 | 0.08 | 3.32E-07 | 1.48E-06 |
| ENSG00000176225 | RTTN | -0.38 | 0.08 | 6.83E-07 | 2.95E-06 |
| ENSG00000124641 | MED20 | -0.38 | 0.08 | 1.11E-06 | 4.67E-06 |
| ENSG00000176390 | CRLF3 | -0.38 | 0.08 | 2.17E-06 | 8.90E-06 |
| ENSG00000213672 | NCKIPSD | -0.38 | 0.08 | 2.78E-06 | 1.13E-05 |
| ENSG00000197763 | TXNRD3 | -0.38 | 0.09 | 4.59E-06 | 1.83E-05 |
| ENSG00000155008 | APOOL | -0.38 | 0.09 | 4.78E-06 | 1.91E-05 |
| ENSG00000197779 | ZNF81 | -0.38 | 0.09 | 8.22E-06 | 3.21E-05 |
| ENSG00000248008 | NRAV | -0.38 | 0.09 | 1.07E-05 | 4.12E-05 |
| ENSG00000162396 | PARS2 | -0.38 | 0.09 | 1.11E-05 | 4.26E-05 |
| ENSG00000149476 | TKFC | -0.38 | 0.09 | 1.34E-05 | 5.12E-05 |
| ENSG00000103642 | LACTB | -0.38 | 0.09 | 1.68E-05 | 6.36E-05 |
| ENSG00000089820 | ARHGAP4 | -0.38 | 0.09 | 2.13E-05 | 7.96E-05 |

| | | | | | |
|---|---|---|---|---|---|
| ENSG00000111799 | COL12A1 | -0.38 | 0.09 | 2.55E-05 | 9.45E-05 |
| ENSG00000137338 | PGBD1 | -0.38 | 0.09 | 3.24E-05 | 1.19E-04 |
| ENSG00000090013 | BLVRB | -0.38 | 0.1 | 4.32E-05 | 1.56E-04 |
| ENSG00000244165 | P2RY11 | -0.38 | 0.1 | 5.29E-05 | 1.90E-04 |
| ENSG00000180855 | ZNF443 | -0.38 | 0.11 | 2.33E-04 | 7.74E-04 |
| ENSG00000063761 | ADCK1 | -0.38 | 0.11 | 2.66E-04 | 8.77E-04 |
| ENSG00000173614 | NMNAT1 | -0.38 | 0.11 | 2.88E-04 | 9.45E-04 |
| ENSG00000126215 | XRCC3 | -0.38 | 0.11 | 5.30E-04 | 1.68E-03 |
| ENSG00000090539 | CHRD | -0.38 | 0.13 | 1.48E-03 | 4.37E-03 |
| ENSG00000156384 | SFR1 | -0.38 | 0.14 | 3.40E-03 | 9.45E-03 |
| ENSG00000089775 | ZBTB25 | -0.38 | 0.15 | 5.60E-03 | 1.49E-02 |
| ENSG00000129295 | DNAAF11 | -0.38 | 0.15 | 6.61E-03 | 1.74E-02 |
| ENSG00000248475 | | -0.38 | 0.16 | 9.28E-03 | 2.36E-02 |
| ENSG00000175536 | LIPT2 | -0.38 | 0.16 | 9.36E-03 | 2.38E-02 |
| ENSG00000138759 | FRAS1 | -0.38 | 0.18 | 1.61E-02 | 3.86E-02 |
| ENSG00000108671 | PSMD11 | -0.37 | 0.03 | 6.16E-26 | 7.18E-25 |
| ENSG00000153250 | RBMS1 | -0.37 | 0.04 | 2.60E-20 | 2.45E-19 |
| ENSG00000134686 | PHC2 | -0.37 | 0.04 | 9.43E-20 | 8.68E-19 |
| ENSG00000139318 | DUSP6 | -0.37 | 0.04 | 7.12E-17 | 5.77E-16 |
| ENSG00000007923 | DNAJC11 | -0.37 | 0.05 | 7.31E-16 | 5.63E-15 |
| ENSG00000110851 | PRDM4 | -0.37 | 0.05 | 2.08E-15 | 1.57E-14 |
| ENSG00000138074 | SLC5A6 | -0.37 | 0.05 | 9.15E-15 | 6.71E-14 |
| ENSG00000175084 | DES | -0.37 | 0.05 | 1.09E-14 | 7.95E-14 |
| ENSG00000054793 | ATP9A | -0.37 | 0.05 | 1.71E-14 | 1.24E-13 |
| ENSG00000206527 | HACD2 | -0.37 | 0.05 | 1.18E-13 | 8.17E-13 |
| ENSG00000095209 | TMEM38B | -0.37 | 0.05 | 4.18E-13 | 2.80E-12 |
| ENSG00000169567 | HINT1 | -0.37 | 0.05 | 6.81E-13 | 4.52E-12 |
| ENSG00000160014 | CALM3 | -0.37 | 0.05 | 6.96E-13 | 4.62E-12 |
| ENSG00000067248 | DHX29 | -0.37 | 0.05 | 7.38E-13 | 4.89E-12 |
| ENSG00000115970 | THADA | -0.37 | 0.05 | 1.53E-12 | 9.90E-12 |
| ENSG00000152558 | TMEM123 | -0.37 | 0.05 | 2.22E-12 | 1.43E-11 |
| ENSG00000163900 | TMEM41A | -0.37 | 0.05 | 2.66E-12 | 1.70E-11 |
| ENSG00000196591 | HDAC2 | -0.37 | 0.06 | 2.28E-11 | 1.38E-10 |
| ENSG00000160199 | PKNOX1 | -0.37 | 0.06 | 1.39E-10 | 7.99E-10 |
| ENSG00000074657 | ZNF532 | -0.37 | 0.06 | 3.97E-10 | 2.21E-09 |
| ENSG00000153767 | GTF2E1 | -0.37 | 0.06 | 5.34E-10 | 2.94E-09 |
| ENSG00000141564 | RPTOR | -0.37 | 0.06 | 7.25E-10 | 3.96E-09 |
| ENSG00000132429 | POPDC3 | -0.37 | 0.06 | 7.28E-10 | 3.97E-09 |
| ENSG00000172037 | LAMB2 | -0.37 | 0.06 | 7.70E-10 | 4.19E-09 |
| ENSG00000183684 | ALYREF | -0.37 | 0.06 | 7.89E-10 | 4.29E-09 |
| ENSG00000101493 | ZNF516 | -0.37 | 0.06 | 1.21E-09 | 6.51E-09 |
| ENSG00000132153 | DHX30 | -0.37 | 0.06 | 2.25E-09 | 1.18E-08 |
| ENSG00000172765 | TMCC1 | -0.37 | 0.06 | 6.25E-09 | 3.17E-08 |
| ENSG00000126953 | TIMM8A | -0.37 | 0.06 | 8.10E-09 | 4.08E-08 |
| ENSG00000134905 | CARS2 | -0.37 | 0.07 | 8.34E-09 | 4.20E-08 |
| ENSG00000112773 | TENT5A | -0.37 | 0.07 | 4.63E-08 | 2.20E-07 |
| ENSG00000135454 | B4GALNT1 | -0.37 | 0.07 | 5.96E-08 | 2.81E-07 |
| ENSG00000104825 | NFKBIB | -0.37 | 0.07 | 1.14E-07 | 5.26E-07 |
| ENSG00000105618 | PRPF31 | -0.37 | 0.07 | 3.22E-07 | 1.43E-06 |
| ENSG00000271643 | PDCD6IP-DT | -0.37 | 0.08 | 3.60E-07 | 1.59E-06 |

| | | | | | |
|---|---|---|---|---|---|
| ENSG00000139350 | NEDD1 | -0.37 | 0.07 | 4.96E-07 | 2.17E-06 |
| ENSG00000101098 | RIMS4 | -0.37 | 0.08 | 8.25E-07 | 3.53E-06 |
| ENSG00000204392 | LSM2 | -0.37 | 0.08 | 1.00E-06 | 4.25E-06 |
| ENSG00000197479 | PCDHB11 | -0.37 | 0.08 | 1.24E-06 | 5.21E-06 |
| ENSG00000115392 | FANCL | -0.37 | 0.08 | 1.49E-06 | 6.23E-06 |
| ENSG00000167716 | WDR81 | -0.37 | 0.08 | 2.09E-06 | 8.59E-06 |
| ENSG00000173917 | HOXB2 | -0.37 | 0.08 | 2.86E-06 | 1.17E-05 |
| ENSG00000151065 | DCP1B | -0.37 | 0.08 | 3.07E-06 | 1.25E-05 |
| ENSG00000132906 | CASP9 | -0.37 | 0.08 | 6.67E-06 | 2.63E-05 |
| ENSG00000270629 | NBPF14 | -0.37 | 0.09 | 9.02E-06 | 3.50E-05 |
| ENSG00000160949 | TONSL | -0.37 | 0.09 | 1.29E-05 | 4.92E-05 |
| ENSG00000206190 | ATP10A | -0.37 | 0.09 | 1.59E-05 | 6.04E-05 |
| ENSG00000060491 | OGFR | -0.37 | 0.09 | 1.69E-05 | 6.37E-05 |
| ENSG00000185798 | WDR53 | -0.37 | 0.09 | 2.59E-05 | 9.59E-05 |
| ENSG00000140682 | TGFB1I1 | -0.37 | 0.09 | 2.83E-05 | 1.04E-04 |
| ENSG00000163273 | NPPC | -0.37 | 0.11 | 4.25E-04 | 1.37E-03 |
| ENSG00000161298 | ZNF382 | -0.37 | 0.11 | 4.68E-04 | 1.50E-03 |
| ENSG00000186862 | PDZD7 | -0.37 | 0.11 | 5.04E-04 | 1.60E-03 |
| ENSG00000188566 | NDOR1 | -0.37 | 0.11 | 7.10E-04 | 2.21E-03 |
| ENSG00000145723 | GIN1 | -0.37 | 0.13 | 1.96E-03 | 5.70E-03 |
| ENSG00000185215 | TNFAIP2 | -0.37 | 0.13 | 2.00E-03 | 5.79E-03 |
| ENSG00000164162 | ANAPC10 | -0.37 | 0.13 | 2.29E-03 | 6.56E-03 |
| ENSG00000141642 | ELAC1 | -0.37 | 0.13 | 2.72E-03 | 7.70E-03 |
| ENSG00000120696 | KBTBD7 | -0.37 | 0.13 | 3.03E-03 | 8.50E-03 |
| ENSG00000160703 | NLRX1 | -0.37 | 0.13 | 3.42E-03 | 9.49E-03 |
| ENSG00000097096 | SYDE2 | -0.37 | 0.13 | 3.55E-03 | 9.82E-03 |
| ENSG00000288066 | | -0.37 | 0.14 | 3.68E-03 | 1.02E-02 |
| ENSG00000269934 | | -0.37 | 0.14 | 5.33E-03 | 1.43E-02 |
| ENSG00000171126 | KCNG3 | -0.37 | 0.15 | 7.98E-03 | 2.06E-02 |
| ENSG00000178075 | GRAMD1C | -0.37 | 0.16 | 9.90E-03 | 2.50E-02 |
| ENSG00000226200 | SGMS1-AS1 | -0.37 | 0.16 | 9.97E-03 | 2.52E-02 |
| ENSG00000138380 | CARF | -0.37 | 0.16 | 1.09E-02 | 2.73E-02 |
| ENSG00000289858 | | -0.37 | 0.16 | 1.32E-02 | 3.24E-02 |
| ENSG00000267260 | | -0.37 | 0.17 | 1.46E-02 | 3.54E-02 |
| ENSG00000258913 | LINC02691 | -0.37 | 0.17 | 1.47E-02 | 3.58E-02 |
| ENSG00000136463 | TACO1 | -0.37 | 0.17 | 1.49E-02 | 3.61E-02 |
| ENSG00000115414 | FN1 | -0.36 | 0.04 | 2.57E-23 | 2.72E-22 |
| ENSG00000120451 | SNX19 | -0.36 | 0.04 | 1.47E-21 | 1.45E-20 |
| ENSG00000125944 | HNRNPR | -0.36 | 0.04 | 9.25E-17 | 7.46E-16 |
| ENSG00000197555 | SIPA1L1 | -0.36 | 0.04 | 3.30E-16 | 2.59E-15 |
| ENSG00000132466 | ANKRD17 | -0.36 | 0.05 | 3.64E-15 | 2.72E-14 |
| ENSG00000204843 | DCTN1 | -0.36 | 0.05 | 3.25E-14 | 2.33E-13 |
| ENSG00000156471 | PTDSS1 | -0.36 | 0.05 | 3.45E-13 | 2.33E-12 |
| ENSG00000111817 | DSE | -0.36 | 0.05 | 4.14E-13 | 2.77E-12 |
| ENSG00000096060 | FKBP5 | -0.36 | 0.05 | 3.03E-12 | 1.93E-11 |
| ENSG00000187257 | RSBN1L | -0.36 | 0.05 | 9.63E-12 | 5.95E-11 |
| ENSG00000180011 | PTGR3 | -0.36 | 0.05 | 2.63E-11 | 1.58E-10 |
| ENSG00000198522 | GPN1 | -0.36 | 0.05 | 3.54E-11 | 2.11E-10 |
| ENSG00000137965 | IFI44 | -0.36 | 0.06 | 9.42E-11 | 5.47E-10 |
| ENSG00000086061 | DNAJA1 | -0.36 | 0.06 | 9.43E-11 | 5.48E-10 |

| | | | | | |
|---|---|---|---|---|---|
| ENSG00000197329 | PELI1 | -0.36 | 0.06 | 1.15E-10 | 6.66E-10 |
| ENSG00000116991 | SIPA1L2 | -0.36 | 0.06 | 7.95E-10 | 4.32E-09 |
| ENSG00000036257 | CUL3 | -0.36 | 0.06 | 8.08E-10 | 4.39E-09 |
| ENSG00000132591 | ERAL1 | -0.36 | 0.06 | 9.25E-10 | 5.00E-09 |
| ENSG00000196652 | ZKSCAN5 | -0.36 | 0.06 | 2.28E-09 | 1.19E-08 |
| ENSG00000077782 | FGFR1 | -0.36 | 0.06 | 2.63E-09 | 1.38E-08 |
| ENSG00000111196 | MAGOHB | -0.36 | 0.06 | 3.94E-09 | 2.03E-08 |
| ENSG00000281398 | SNHG4 | -0.36 | 0.06 | 5.36E-09 | 2.74E-08 |
| ENSG00000124782 | RREB1 | -0.36 | 0.07 | 6.97E-08 | 3.27E-07 |
| ENSG00000196876 | SCN8A | -0.36 | 0.07 | 1.42E-07 | 6.50E-07 |
| ENSG00000172269 | DPAGT1 | -0.36 | 0.07 | 4.82E-07 | 2.11E-06 |
| ENSG00000143797 | MBOAT2 | -0.36 | 0.07 | 6.12E-07 | 2.66E-06 |
| ENSG00000083097 | DOP1A | -0.36 | 0.08 | 1.15E-06 | 4.86E-06 |
| ENSG00000162688 | AGL | -0.36 | 0.08 | 1.49E-06 | 6.25E-06 |
| ENSG00000113597 | TRAPPC13 | -0.36 | 0.08 | 4.11E-06 | 1.65E-05 |
| ENSG00000105875 | WDR91 | -0.36 | 0.08 | 5.44E-06 | 2.16E-05 |
| ENSG00000180448 | ARHGAP45 | -0.36 | 0.08 | 5.85E-06 | 2.31E-05 |
| ENSG00000164053 | ATRIP | -0.36 | 0.08 | 7.48E-06 | 2.93E-05 |
| ENSG00000153774 | CFDP1 | -0.36 | 0.08 | 1.09E-05 | 4.20E-05 |
| ENSG00000061273 | HDAC7 | -0.36 | 0.08 | 1.34E-05 | 5.11E-05 |
| ENSG00000108823 | SGCA | -0.36 | 0.09 | 1.97E-05 | 7.40E-05 |
| ENSG00000115267 | IFIH1 | -0.36 | 0.09 | 2.72E-05 | 1.01E-04 |
| ENSG00000128294 | TPST2 | -0.36 | 0.09 | 2.81E-05 | 1.04E-04 |
| ENSG00000178537 | SLC25A20 | -0.36 | 0.09 | 4.99E-05 | 1.79E-04 |
| ENSG00000074416 | MGLL | -0.36 | 0.1 | 8.35E-05 | 2.93E-04 |
| ENSG00000162302 | RPS6KA4 | -0.36 | 0.1 | 2.91E-04 | 9.55E-04 |
| ENSG00000144026 | ZNF514 | -0.36 | 0.11 | 5.06E-04 | 1.61E-03 |
| ENSG00000135272 | MDFIC | -0.36 | 0.12 | 1.70E-03 | 4.99E-03 |
| ENSG00000274349 | ZNF658 | -0.36 | 0.13 | 3.82E-03 | 1.05E-02 |
| ENSG00000243742 | RPLP0P2 | -0.36 | 0.14 | 4.72E-03 | 1.28E-02 |
| ENSG00000175344 | CHRNA7 | -0.36 | 0.16 | 1.68E-02 | 4.03E-02 |
| ENSG00000151006 | PRSS53 | -0.36 | 0.17 | 1.90E-02 | 4.49E-02 |
| ENSG00000151287 | TEX30 | -0.36 | 0.17 | 1.97E-02 | 4.66E-02 |
| ENSG00000117751 | PPP1R8 | -0.35 | 0.04 | 5.44E-20 | 5.05E-19 |
| ENSG00000101040 | ZMYND8 | -0.35 | 0.04 | 1.11E-18 | 9.71E-18 |
| ENSG00000113649 | TCERG1 | -0.35 | 0.04 | 3.06E-17 | 2.52E-16 |
| ENSG00000146731 | CCT6A | -0.35 | 0.04 | 6.89E-17 | 5.60E-16 |
| ENSG00000154473 | BUB3 | -0.35 | 0.04 | 2.29E-16 | 1.81E-15 |
| ENSG00000065150 | IPO5 | -0.35 | 0.04 | 4.81E-16 | 3.75E-15 |
| ENSG00000177084 | POLE | -0.35 | 0.04 | 4.98E-16 | 3.88E-15 |
| ENSG00000184634 | MED12 | -0.35 | 0.04 | 6.30E-16 | 4.88E-15 |
| ENSG00000183853 | KIRREL1 | -0.35 | 0.04 | 1.31E-15 | 9.97E-15 |
| ENSG00000157895 | C12orf43 | -0.35 | 0.05 | 9.24E-15 | 6.78E-14 |
| ENSG00000198873 | GRK5 | -0.35 | 0.05 | 1.27E-14 | 9.28E-14 |
| ENSG00000109189 | USP46 | -0.35 | 0.05 | 3.20E-14 | 2.29E-13 |
| ENSG00000176485 | PLAAT3 | -0.35 | 0.05 | 1.12E-13 | 7.76E-13 |
| ENSG00000106025 | TSPAN12 | -0.35 | 0.05 | 6.48E-13 | 4.30E-12 |
| ENSG00000120942 | UBIAD1 | -0.35 | 0.05 | 2.29E-12 | 1.47E-11 |
| ENSG00000108788 | MLX | -0.35 | 0.05 | 3.00E-12 | 1.92E-11 |
| ENSG00000058668 | ATP2B4 | -0.35 | 0.05 | 8.52E-12 | 5.28E-11 |

| | | | | | |
|---|---|---|---|---|---|
| ENSG00000100426 | ZBED4 | -0.35 | 0.05 | 2.93E-11 | 1.76E-10 |
| ENSG00000164209 | SLC25A46 | -0.35 | 0.05 | 1.42E-10 | 8.17E-10 |
| ENSG00000102225 | CDK16 | -0.35 | 0.06 | 7.80E-10 | 4.24E-09 |
| ENSG00000068097 | HEATR6 | -0.35 | 0.06 | 2.66E-09 | 1.39E-08 |
| ENSG00000083444 | PLOD1 | -0.35 | 0.06 | 3.05E-09 | 1.59E-08 |
| ENSG00000093144 | ECHDC1 | -0.35 | 0.06 | 1.24E-08 | 6.16E-08 |
| ENSG00000138386 | NAB1 | -0.35 | 0.06 | 1.37E-08 | 6.79E-08 |
| ENSG00000145725 | PPIP5K2 | -0.35 | 0.06 | 4.36E-08 | 2.08E-07 |
| ENSG00000152778 | IFIT5 | -0.35 | 0.07 | 4.45E-08 | 2.12E-07 |
| ENSG00000183166 | CALN1 | -0.35 | 0.07 | 5.82E-08 | 2.75E-07 |
| ENSG00000129219 | PLD2 | -0.35 | 0.07 | 1.23E-07 | 5.67E-07 |
| ENSG00000116667 | C1orf21 | -0.35 | 0.07 | 4.14E-07 | 1.83E-06 |
| ENSG00000170035 | UBE2E3 | -0.35 | 0.07 | 4.49E-07 | 1.97E-06 |
| ENSG00000174744 | BRMS1 | -0.35 | 0.07 | 7.31E-07 | 3.15E-06 |
| ENSG00000152284 | TCF7L1 | -0.35 | 0.07 | 9.70E-07 | 4.12E-06 |
| ENSG00000136444 | RSAD1 | -0.35 | 0.07 | 1.39E-06 | 5.82E-06 |
| ENSG00000149091 | DGKZ | -0.35 | 0.08 | 1.42E-06 | 5.97E-06 |
| ENSG00000144736 | SHQ1 | -0.35 | 0.07 | 1.51E-06 | 6.30E-06 |
| ENSG00000113593 | PPWD1 | -0.35 | 0.07 | 1.56E-06 | 6.53E-06 |
| ENSG00000150967 | ABCB9 | -0.35 | 0.08 | 3.17E-06 | 1.28E-05 |
| ENSG00000288558 | DUS4L-BCAP29 | -0.35 | 0.08 | 3.75E-06 | 1.51E-05 |
| ENSG00000174173 | TRMT10C | -0.35 | 0.08 | 4.37E-06 | 1.75E-05 |
| ENSG00000188807 | TMEM201 | -0.35 | 0.08 | 4.68E-06 | 1.87E-05 |
| ENSG00000171448 | ZBTB26 | -0.35 | 0.08 | 7.63E-06 | 2.99E-05 |
| ENSG00000111859 | NEDD9 | -0.35 | 0.09 | 3.18E-05 | 1.17E-04 |
| ENSG00000070371 | CLTCL1 | -0.35 | 0.09 | 3.29E-05 | 1.21E-04 |
| ENSG00000160336 | ZNF761 | -0.35 | 0.09 | 3.77E-05 | 1.37E-04 |
| ENSG00000204977 | TRIM13 | -0.35 | 0.09 | 6.31E-05 | 2.25E-04 |
| ENSG00000188917 | TRMT2B | -0.35 | 0.09 | 6.98E-05 | 2.48E-04 |
| ENSG00000131652 | THOC6 | -0.35 | 0.09 | 1.01E-04 | 3.51E-04 |
| ENSG00000235954 | TTC28-AS1 | -0.35 | 0.1 | 1.88E-04 | 6.33E-04 |
| ENSG00000144893 | MED12L | -0.35 | 0.1 | 2.30E-04 | 7.64E-04 |
| ENSG00000106009 | BRAT1 | -0.35 | 0.1 | 2.72E-04 | 8.98E-04 |
| ENSG00000241015 | TPM3P9 | -0.35 | 0.1 | 3.32E-04 | 1.08E-03 |
| ENSG00000198785 | GRIN3A | -0.35 | 0.1 | 3.65E-04 | 1.18E-03 |
| ENSG00000186806 | VSIG10L | -0.35 | 0.1 | 4.52E-04 | 1.45E-03 |
| ENSG00000175066 | GK5 | -0.35 | 0.1 | 4.72E-04 | 1.51E-03 |
| ENSG00000180787 | ZFP3 | -0.35 | 0.11 | 5.20E-04 | 1.65E-03 |
| ENSG00000006740 | ARHGAP44 | -0.35 | 0.11 | 1.01E-03 | 3.07E-03 |
| ENSG00000003137 | CYP26B1 | -0.35 | 0.11 | 1.12E-03 | 3.37E-03 |
| ENSG00000169951 | ZNF764 | -0.35 | 0.11 | 1.22E-03 | 3.66E-03 |
| ENSG00000143458 | GABPB2 | -0.35 | 0.12 | 2.02E-03 | 5.85E-03 |
| ENSG00000289811 | FAM95A | -0.35 | 0.13 | 3.19E-03 | 8.92E-03 |
| ENSG00000214485 | RPL7P1 | -0.35 | 0.13 | 5.57E-03 | 1.49E-02 |
| ENSG00000288886 | | -0.35 | 0.14 | 8.16E-03 | 2.10E-02 |
| ENSG00000129911 | KLF16 | -0.35 | 0.14 | 9.21E-03 | 2.35E-02 |
| ENSG00000108465 | CDK5RAP3 | -0.35 | 0.15 | 9.82E-03 | 2.49E-02 |
| ENSG00000143554 | SLC27A3 | -0.35 | 0.16 | 2.11E-02 | 4.95E-02 |
| ENSG00000200087 | SNORA73B | -0.35 | 0.17 | 2.12E-02 | 4.98E-02 |
| ENSG00000078668 | VDAC3 | -0.34 | 0.04 | 4.20E-18 | 3.58E-17 |

| | | | | | |
|---|---|---|---|---|---|
| ENSG00000119772 | DNMT3A | -0.34 | 0.04 | 1.95E-15 | 1.47E-14 |
| ENSG00000182149 | IST1 | -0.34 | 0.04 | 2.44E-15 | 1.84E-14 |
| ENSG00000153107 | ANAPC1 | -0.34 | 0.04 | 1.06E-14 | 7.78E-14 |
| ENSG00000135763 | URB2 | -0.34 | 0.04 | 1.45E-14 | 1.05E-13 |
| ENSG00000103365 | GGA2 | -0.34 | 0.04 | 2.21E-14 | 1.59E-13 |
| ENSG00000139323 | POC1B | -0.34 | 0.05 | 3.82E-13 | 2.57E-12 |
| ENSG00000101391 | CDK5RAP1 | -0.34 | 0.05 | 7.70E-13 | 5.09E-12 |
| ENSG00000158161 | EYA3 | -0.34 | 0.05 | 1.13E-12 | 7.37E-12 |
| ENSG00000153827 | TRIP12 | -0.34 | 0.05 | 2.75E-12 | 1.76E-11 |
| ENSG00000232956 | SNHG15 | -0.34 | 0.05 | 2.92E-12 | 1.86E-11 |
| ENSG00000117174 | ZNHIT6 | -0.34 | 0.05 | 1.37E-11 | 8.36E-11 |
| ENSG00000108578 | BLMH | -0.34 | 0.05 | 6.25E-11 | 3.67E-10 |
| ENSG00000050130 | JKAMP | -0.34 | 0.06 | 4.41E-10 | 2.44E-09 |
| ENSG00000187778 | MCRS1 | -0.34 | 0.05 | 6.90E-10 | 3.78E-09 |
| ENSG00000125450 | NUP85 | -0.34 | 0.06 | 8.81E-10 | 4.77E-09 |
| ENSG00000166261 | ZNF202 | -0.34 | 0.06 | 9.33E-10 | 5.05E-09 |
| ENSG00000235162 | C12orf75 | -0.34 | 0.06 | 1.67E-09 | 8.86E-09 |
| ENSG00000186141 | POLR3C | -0.34 | 0.06 | 2.25E-09 | 1.18E-08 |
| ENSG00000170088 | TMEM192 | -0.34 | 0.06 | 2.80E-09 | 1.46E-08 |
| ENSG00000165288 | BRWD3 | -0.34 | 0.06 | 4.17E-09 | 2.14E-08 |
| ENSG00000105849 | POLR1F | -0.34 | 0.06 | 4.80E-09 | 2.45E-08 |
| ENSG00000115368 | WDR75 | -0.34 | 0.06 | 1.68E-08 | 8.27E-08 |
| ENSG00000188191 | PRKAR1B | -0.34 | 0.06 | 1.96E-08 | 9.61E-08 |
| ENSG00000064666 | CNN2 | -0.34 | 0.06 | 2.27E-08 | 1.11E-07 |
| ENSG00000143847 | PPFIA4 | -0.34 | 0.06 | 6.11E-08 | 2.88E-07 |
| ENSG00000058056 | USP13 | -0.34 | 0.07 | 1.77E-07 | 8.07E-07 |
| ENSG00000177034 | MTX3 | -0.34 | 0.07 | 2.12E-07 | 9.61E-07 |
| ENSG00000151468 | CCDC3 | -0.34 | 0.07 | 3.26E-07 | 1.45E-06 |
| ENSG00000206562 | METTL6 | -0.34 | 0.07 | 3.39E-07 | 1.51E-06 |
| ENSG00000139219 | COL2A1 | -0.34 | 0.07 | 4.67E-07 | 2.05E-06 |
| ENSG00000151304 | SRFBP1 | -0.34 | 0.07 | 9.13E-07 | 3.89E-06 |
| ENSG00000136143 | SUCLA2 | -0.34 | 0.07 | 1.11E-06 | 4.68E-06 |
| ENSG00000061938 | TNK2 | -0.34 | 0.07 | 1.83E-06 | 7.56E-06 |
| ENSG00000100473 | COCH | -0.34 | 0.07 | 2.34E-06 | 9.59E-06 |
| ENSG00000103351 | CLUAP1 | -0.34 | 0.08 | 2.01E-05 | 7.56E-05 |
| ENSG00000180318 | ALX1 | -0.34 | 0.08 | 3.41E-05 | 1.25E-04 |
| ENSG00000125351 | UPF3B | -0.34 | 0.09 | 3.85E-05 | 1.40E-04 |
| ENSG00000043039 | BARX2 | -0.34 | 0.08 | 4.11E-05 | 1.49E-04 |
| ENSG00000238227 | TMEM250 | -0.34 | 0.09 | 5.56E-05 | 1.99E-04 |
| ENSG00000091073 | DTX2 | -0.34 | 0.09 | 7.08E-05 | 2.51E-04 |
| ENSG00000148357 | HMCN2 | -0.34 | 0.09 | 8.26E-05 | 2.90E-04 |
| ENSG00000174917 | MICOS13 | -0.34 | 0.09 | 8.30E-05 | 2.91E-04 |
| ENSG00000108852 | MPP2 | -0.34 | 0.09 | 9.93E-05 | 3.46E-04 |
| ENSG00000187097 | ENTPD5 | -0.34 | 0.1 | 2.05E-04 | 6.87E-04 |
| ENSG00000104722 | NEFM | -0.34 | 0.1 | 3.70E-04 | 1.20E-03 |
| ENSG00000212916 | MAP10 | -0.34 | 0.1 | 5.51E-04 | 1.75E-03 |
| ENSG00000131061 | ZNF341 | -0.34 | 0.11 | 8.11E-04 | 2.50E-03 |
| ENSG00000050555 | LAMC3 | -0.34 | 0.12 | 2.81E-03 | 7.93E-03 |
| ENSG00000166484 | MAPK7 | -0.34 | 0.12 | 2.93E-03 | 8.25E-03 |
| ENSG00000104881 | PPP1R13L | -0.34 | 0.13 | 5.45E-03 | 1.46E-02 |

| | | | | | |
|---|---|---|---|---|---|
| ENSG00000127399 | LRRC61 | -0.34 | 0.14 | 7.44E-03 | 1.94E-02 |
| ENSG00000250992 | | -0.34 | 0.14 | 8.49E-03 | 2.18E-02 |
| ENSG00000088881 | EBF4 | -0.34 | 0.14 | 9.09E-03 | 2.32E-02 |
| ENSG00000155158 | TTC39B | -0.34 | 0.15 | 1.43E-02 | 3.49E-02 |
| ENSG00000137203 | TFAP2A | -0.34 | 0.15 | 1.59E-02 | 3.83E-02 |
| ENSG00000091136 | LAMB1 | -0.33 | 0.03 | 1.99E-24 | 2.20E-23 |
| ENSG00000030066 | NUP160 | -0.33 | 0.04 | 1.95E-14 | 1.41E-13 |
| ENSG00000100422 | CERK | -0.33 | 0.05 | 1.02E-12 | 6.66E-12 |
| ENSG00000139719 | VPS33A | -0.33 | 0.05 | 1.30E-12 | 8.44E-12 |
| ENSG00000147573 | TRIM55 | -0.33 | 0.05 | 6.98E-12 | 4.34E-11 |
| ENSG00000114503 | NCBP2 | -0.33 | 0.05 | 7.22E-12 | 4.49E-11 |
| ENSG00000167977 | KCTD5 | -0.33 | 0.05 | 3.09E-11 | 1.85E-10 |
| ENSG00000113812 | ACTR8 | -0.33 | 0.05 | 4.64E-11 | 2.75E-10 |
| ENSG00000138085 | ATRAID | -0.33 | 0.05 | 8.09E-11 | 4.72E-10 |
| ENSG00000152977 | ZIC1 | -0.33 | 0.05 | 1.03E-10 | 6.00E-10 |
| ENSG00000213593 | TMX2 | -0.33 | 0.05 | 1.72E-10 | 9.81E-10 |
| ENSG00000197037 | ZSCAN25 | -0.33 | 0.05 | 1.79E-10 | 1.02E-09 |
| ENSG00000048162 | NOP16 | -0.33 | 0.05 | 3.65E-10 | 2.03E-09 |
| ENSG00000198301 | SDAD1 | -0.33 | 0.05 | 3.90E-10 | 2.17E-09 |
| ENSG00000215193 | PEX26 | -0.33 | 0.05 | 4.53E-10 | 2.51E-09 |
| ENSG00000254726 | MEX3A | -0.33 | 0.05 | 5.45E-10 | 3.00E-09 |
| ENSG00000121058 | COIL | -0.33 | 0.06 | 1.19E-09 | 6.39E-09 |
| ENSG00000058091 | CDK14 | -0.33 | 0.06 | 1.67E-09 | 8.88E-09 |
| ENSG00000140332 | TLE3 | -0.33 | 0.06 | 2.44E-09 | 1.27E-08 |
| ENSG00000086200 | IPO11 | -0.33 | 0.06 | 3.85E-09 | 1.98E-08 |
| ENSG00000108506 | INTS2 | -0.33 | 0.06 | 4.83E-09 | 2.47E-08 |
| ENSG00000180008 | SOCS4 | -0.33 | 0.06 | 6.17E-09 | 3.14E-08 |
| ENSG00000129292 | PHF20L1 | -0.33 | 0.06 | 6.51E-09 | 3.30E-08 |
| ENSG00000051620 | HEBP2 | -0.33 | 0.06 | 1.53E-08 | 7.56E-08 |
| ENSG00000119203 | CPSF3 | -0.33 | 0.06 | 1.81E-08 | 8.90E-08 |
| ENSG00000215251 | FASTKD5 | -0.33 | 0.06 | 3.04E-08 | 1.47E-07 |
| ENSG00000183765 | CHEK2 | -0.33 | 0.06 | 4.86E-08 | 2.31E-07 |
| ENSG00000109445 | ZNF330 | -0.33 | 0.06 | 8.92E-08 | 4.16E-07 |
| ENSG00000147439 | BIN3 | -0.33 | 0.06 | 2.10E-07 | 9.52E-07 |
| ENSG00000197170 | PSMD12 | -0.33 | 0.06 | 2.85E-07 | 1.28E-06 |
| ENSG00000171163 | ZNF692 | -0.33 | 0.07 | 4.80E-07 | 2.11E-06 |
| ENSG00000136159 | NUDT15 | -0.33 | 0.07 | 4.90E-07 | 2.15E-06 |
| ENSG00000178974 | FBXO34 | -0.33 | 0.07 | 6.11E-07 | 2.65E-06 |
| ENSG00000162885 | B3GALNT2 | -0.33 | 0.07 | 9.97E-07 | 4.23E-06 |
| ENSG00000144744 | UBA3 | -0.33 | 0.07 | 1.00E-06 | 4.24E-06 |
| ENSG00000111450 | STX2 | -0.33 | 0.07 | 1.36E-06 | 5.73E-06 |
| ENSG00000115750 | TAF1B | -0.33 | 0.07 | 1.48E-06 | 6.20E-06 |
| ENSG00000221968 | FADS3 | -0.33 | 0.07 | 1.90E-06 | 7.84E-06 |
| ENSG00000247556 | OIP5-AS1 | -0.33 | 0.07 | 5.76E-06 | 2.28E-05 |
| ENSG00000010072 | SPRTN | -0.33 | 0.07 | 5.82E-06 | 2.30E-05 |
| ENSG00000143469 | SYT14 | -0.33 | 0.07 | 8.62E-06 | 3.35E-05 |
| ENSG00000106991 | ENG | -0.33 | 0.08 | 1.00E-05 | 3.87E-05 |
| ENSG00000197885 | NKIRAS1 | -0.33 | 0.08 | 1.09E-05 | 4.19E-05 |
| ENSG00000169189 | NSMCE1 | -0.33 | 0.08 | 1.39E-05 | 5.29E-05 |
| ENSG00000147654 | EBAG9 | -0.33 | 0.08 | 1.41E-05 | 5.38E-05 |

| | | | | | |
|---|---|---|---|---|---|
| ENSG00000173480 | ZNF417 | -0.33 | 0.08 | 2.19E-05 | 8.20E-05 |
| ENSG00000124104 | SNX21 | -0.33 | 0.08 | 3.84E-05 | 1.40E-04 |
| ENSG00000014164 | ZC3H3 | -0.33 | 0.09 | 8.01E-05 | 2.82E-04 |
| ENSG00000099256 | PRTFDC1 | -0.33 | 0.09 | 9.49E-05 | 3.31E-04 |
| ENSG00000099864 | PALM | -0.33 | 0.09 | 1.37E-04 | 4.69E-04 |
| ENSG00000169019 | COMMD8 | -0.33 | 0.09 | 1.72E-04 | 5.80E-04 |
| ENSG00000188931 | CFAP126 | -0.33 | 0.09 | 2.01E-04 | 6.73E-04 |
| ENSG00000088305 | DNMT3B | -0.33 | 0.09 | 2.03E-04 | 6.79E-04 |
| ENSG00000131370 | SH3BP5 | -0.33 | 0.09 | 2.75E-04 | 9.05E-04 |
| ENSG00000288670 | | -0.33 | 0.09 | 3.11E-04 | 1.02E-03 |
| ENSG00000169193 | CCDC126 | -0.33 | 0.1 | 7.23E-04 | 2.25E-03 |
| ENSG00000255624 | C10orf88B | -0.33 | 0.1 | 7.94E-04 | 2.46E-03 |
| ENSG00000277462 | ZNF670 | -0.33 | 0.11 | 1.16E-03 | 3.49E-03 |
| ENSG00000150455 | TIRAP | -0.33 | 0.1 | 1.16E-03 | 3.50E-03 |
| ENSG00000140511 | HAPLN3 | -0.33 | 0.11 | 1.39E-03 | 4.12E-03 |
| ENSG00000111801 | BTN3A3 | -0.33 | 0.11 | 1.89E-03 | 5.50E-03 |
| ENSG00000204439 | C6orf47 | -0.33 | 0.11 | 2.04E-03 | 5.89E-03 |
| ENSG00000140025 | EFCAB11 | -0.33 | 0.12 | 3.00E-03 | 8.43E-03 |
| ENSG00000186603 | HPDL | -0.33 | 0.12 | 3.50E-03 | 9.69E-03 |
| ENSG00000165898 | ISCA2 | -0.33 | 0.12 | 3.75E-03 | 1.03E-02 |
| ENSG00000239713 | APOBEC3G | -0.33 | 0.13 | 8.40E-03 | 2.16E-02 |
| ENSG00000261236 | BOP1 | -0.33 | 0.13 | 8.52E-03 | 2.19E-02 |
| ENSG00000204934 | ATP6V0E2-AS1 | -0.33 | 0.14 | 1.16E-02 | 2.89E-02 |
| ENSG00000145365 | TIFA | -0.33 | 0.15 | 1.79E-02 | 4.25E-02 |
| ENSG00000166086 | JAM3 | -0.32 | 0.04 | 1.21E-18 | 1.06E-17 |
| ENSG00000120694 | HSPH1 | -0.32 | 0.04 | 4.00E-18 | 3.41E-17 |
| ENSG00000140632 | GLYR1 | -0.32 | 0.04 | 1.15E-16 | 9.23E-16 |
| ENSG00000169851 | PCDH7 | -0.32 | 0.04 | 1.21E-15 | 9.19E-15 |
| ENSG00000121579 | NAA50 | -0.32 | 0.04 | 2.16E-14 | 1.55E-13 |
| ENSG00000145741 | BTF3 | -0.32 | 0.04 | 2.67E-14 | 1.92E-13 |
| ENSG00000087111 | PIGS | -0.32 | 0.04 | 7.62E-14 | 5.34E-13 |
| ENSG00000215021 | PHB2 | -0.32 | 0.04 | 7.74E-14 | 5.41E-13 |
| ENSG00000123505 | AMD1 | -0.32 | 0.04 | 1.40E-13 | 9.66E-13 |
| ENSG00000184677 | ZBTB40 | -0.32 | 0.04 | 4.76E-13 | 3.18E-12 |
| ENSG00000065183 | WDR3 | -0.32 | 0.04 | 1.38E-12 | 8.96E-12 |
| ENSG00000166130 | IKBIP | -0.32 | 0.05 | 4.29E-12 | 2.71E-11 |
| ENSG00000161526 | SAP30BP | -0.32 | 0.05 | 4.32E-12 | 2.72E-11 |
| ENSG00000163697 | APBB2 | -0.32 | 0.05 | 1.27E-11 | 7.80E-11 |
| ENSG00000143624 | INTS3 | -0.32 | 0.05 | 2.32E-11 | 1.40E-10 |
| ENSG00000197724 | PHF2 | -0.32 | 0.05 | 2.64E-11 | 1.59E-10 |
| ENSG00000125686 | MED1 | -0.32 | 0.05 | 6.87E-11 | 4.02E-10 |
| ENSG00000185129 | PURA | -0.32 | 0.05 | 2.10E-10 | 1.19E-09 |
| ENSG00000205581 | HMGN1 | -0.32 | 0.05 | 2.11E-10 | 1.20E-09 |
| ENSG00000183530 | PRR14L | -0.32 | 0.05 | 1.01E-09 | 5.45E-09 |
| ENSG00000161203 | AP2M1 | -0.32 | 0.05 | 1.09E-09 | 5.87E-09 |
| ENSG00000180628 | PCGF5 | -0.32 | 0.05 | 1.18E-09 | 6.33E-09 |
| ENSG00000143368 | SF3B4 | -0.32 | 0.05 | 2.70E-09 | 1.41E-08 |
| ENSG00000075213 | SEMA3A | -0.32 | 0.05 | 2.79E-09 | 1.45E-08 |
| ENSG00000129128 | SPCS3 | -0.32 | 0.06 | 6.43E-09 | 3.26E-08 |
| ENSG00000013503 | POLR3B | -0.32 | 0.06 | 1.07E-08 | 5.37E-08 |

| | | | | | |
|---|---|---|---|---|---|
| ENSG00000172586 | CHCHD1 | -0.32 | 0.06 | 1.13E-08 | 5.62E-08 |
| ENSG00000134453 | RBM17 | -0.32 | 0.06 | 2.73E-08 | 1.32E-07 |
| ENSG00000125482 | TTF1 | -0.32 | 0.06 | 4.34E-08 | 2.07E-07 |
| ENSG00000116922 | C1orf109 | -0.32 | 0.06 | 7.65E-08 | 3.58E-07 |
| ENSG00000167081 | PBX3 | -0.32 | 0.06 | 8.31E-08 | 3.88E-07 |
| ENSG00000124608 | AARS2 | -0.32 | 0.06 | 1.13E-07 | 5.22E-07 |
| ENSG00000115866 | DARS1 | -0.32 | 0.06 | 5.86E-07 | 2.55E-06 |
| ENSG00000120798 | NR2C1 | -0.32 | 0.07 | 6.81E-07 | 2.94E-06 |
| ENSG00000005020 | SKAP2 | -0.32 | 0.07 | 8.02E-07 | 3.44E-06 |
| ENSG00000136518 | ACTL6A | -0.32 | 0.07 | 8.70E-07 | 3.71E-06 |
| ENSG00000139722 | VPS37B | -0.32 | 0.07 | 9.20E-07 | 3.92E-06 |
| ENSG00000174442 | ZWILCH | -0.32 | 0.07 | 1.01E-06 | 4.29E-06 |
| ENSG00000145388 | METTL14 | -0.32 | 0.07 | 2.26E-06 | 9.27E-06 |
| ENSG00000118965 | WDR35 | -0.32 | 0.07 | 2.36E-06 | 9.68E-06 |
| ENSG00000112685 | EXOC2 | -0.32 | 0.07 | 3.03E-06 | 1.23E-05 |
| ENSG00000100605 | ITPK1 | -0.32 | 0.07 | 4.97E-06 | 1.98E-05 |
| ENSG00000259956 | RBM15B | -0.32 | 0.07 | 6.16E-06 | 2.43E-05 |
| ENSG00000177054 | ZDHHC13 | -0.32 | 0.07 | 1.30E-05 | 4.98E-05 |
| ENSG00000198399 | ITSN2 | -0.32 | 0.08 | 1.40E-05 | 5.33E-05 |
| ENSG00000183814 | LIN9 | -0.32 | 0.08 | 2.48E-05 | 9.20E-05 |
| ENSG00000186318 | BACE1 | -0.32 | 0.08 | 3.26E-05 | 1.19E-04 |
| ENSG00000240694 | PNMA2 | -0.32 | 0.08 | 3.92E-05 | 1.42E-04 |
| ENSG00000178773 | CPNE7 | -0.32 | 0.09 | 1.28E-04 | 4.38E-04 |
| ENSG00000175806 | MSRA | -0.32 | 0.09 | 1.47E-04 | 4.99E-04 |
| ENSG00000182518 | FAM104B | -0.32 | 0.09 | 4.19E-04 | 1.35E-03 |
| ENSG00000185220 | PGBD2 | -0.32 | 0.1 | 6.58E-04 | 2.06E-03 |
| ENSG00000266074 | BAHCC1 | -0.32 | 0.1 | 7.04E-04 | 2.19E-03 |
| ENSG00000198157 | HMGN5 | -0.32 | 0.1 | 7.25E-04 | 2.26E-03 |
| ENSG00000180902 | D2HGDH | -0.32 | 0.1 | 7.29E-04 | 2.26E-03 |
| ENSG00000050344 | NFE2L3 | -0.32 | 0.11 | 3.32E-03 | 9.24E-03 |
| ENSG00000113971 | NPHP3 | -0.32 | 0.13 | 8.85E-03 | 2.27E-02 |
| ENSG00000249353 | NPM1P27 | -0.32 | 0.14 | 1.26E-02 | 3.11E-02 |
| ENSG00000134463 | ECHDC3 | -0.32 | 0.15 | 2.04E-02 | 4.81E-02 |
| ENSG00000198492 | YTHDF2 | -0.31 | 0.04 | 2.14E-16 | 1.70E-15 |
| ENSG00000067798 | NAV3 | -0.31 | 0.04 | 3.65E-15 | 2.72E-14 |
| ENSG00000143179 | UCK2 | -0.31 | 0.04 | 7.48E-14 | 5.24E-13 |
| ENSG00000182534 | MXRA7 | -0.31 | 0.04 | 2.06E-13 | 1.41E-12 |
| ENSG00000143621 | ILF2 | -0.31 | 0.04 | 6.74E-13 | 4.47E-12 |
| ENSG00000183495 | EP400 | -0.31 | 0.04 | 1.08E-12 | 7.06E-12 |
| ENSG00000065427 | KARS1 | -0.31 | 0.05 | 2.05E-11 | 1.24E-10 |
| ENSG00000171109 | MFN1 | -0.31 | 0.05 | 9.96E-11 | 5.78E-10 |
| ENSG00000166479 | TMX3 | -0.31 | 0.05 | 3.24E-10 | 1.81E-09 |
| ENSG00000100461 | RBM23 | -0.31 | 0.05 | 6.13E-10 | 3.36E-09 |
| ENSG00000049618 | ARID1B | -0.31 | 0.05 | 2.10E-09 | 1.11E-08 |
| ENSG00000074800 | ENO1 | -0.31 | 0.05 | 4.03E-09 | 2.08E-08 |
| ENSG00000147257 | GPC3 | -0.31 | 0.05 | 4.20E-09 | 2.16E-08 |
| ENSG00000070785 | EIF2B3 | -0.31 | 0.05 | 6.11E-09 | 3.11E-08 |
| ENSG00000124831 | LRRFIP1 | -0.31 | 0.06 | 1.33E-08 | 6.60E-08 |
| ENSG00000234420 | ZNF37BP | -0.31 | 0.06 | 4.61E-08 | 2.20E-07 |
| ENSG00000111731 | C2CD5 | -0.31 | 0.06 | 4.81E-08 | 2.29E-07 |

| | | | | | |
|---|---|---|---|---|---|
| ENSG00000122565 | CBX3 | -0.31 | 0.06 | 5.06E-08 | 2.40E-07 |
| ENSG00000137992 | DBT | -0.31 | 0.06 | 1.02E-07 | 4.72E-07 |
| ENSG00000148187 | MRRF | -0.31 | 0.06 | 1.09E-07 | 5.04E-07 |
| ENSG00000168487 | BMP1 | -0.31 | 0.06 | 1.16E-07 | 5.36E-07 |
| ENSG00000134291 | TMEM106C | -0.31 | 0.06 | 1.51E-07 | 6.91E-07 |
| ENSG00000186063 | AIDA | -0.31 | 0.06 | 1.56E-07 | 7.11E-07 |
| ENSG00000153721 | CNKSR3 | -0.31 | 0.06 | 1.60E-07 | 7.30E-07 |
| ENSG00000095485 | CWF19L1 | -0.31 | 0.06 | 1.66E-07 | 7.54E-07 |
| ENSG00000115211 | EIF2B4 | -0.31 | 0.06 | 3.70E-07 | 1.64E-06 |
| ENSG00000129691 | ASH2L | -0.31 | 0.06 | 4.02E-07 | 1.78E-06 |
| ENSG00000227057 | WDR46 | -0.31 | 0.06 | 4.94E-07 | 2.16E-06 |
| ENSG00000136997 | MYC | -0.31 | 0.06 | 5.85E-07 | 2.55E-06 |
| ENSG00000115641 | FHL2 | -0.31 | 0.06 | 7.27E-07 | 3.13E-06 |
| ENSG00000138640 | FAM13A | -0.31 | 0.06 | 7.47E-07 | 3.21E-06 |
| ENSG00000060339 | CCAR1 | -0.31 | 0.06 | 9.91E-07 | 4.21E-06 |
| ENSG00000127980 | PEX1 | -0.31 | 0.06 | 1.05E-06 | 4.45E-06 |
| ENSG00000241878 | PISD | -0.31 | 0.07 | 2.85E-06 | 1.16E-05 |
| ENSG00000106355 | LSM5 | -0.31 | 0.07 | 4.18E-06 | 1.68E-05 |
| ENSG00000004399 | PLXND1 | -0.31 | 0.07 | 6.86E-06 | 2.70E-05 |
| ENSG00000039523 | RIPOR1 | -0.31 | 0.07 | 8.49E-06 | 3.30E-05 |
| ENSG00000131269 | ABCB7 | -0.31 | 0.07 | 8.73E-06 | 3.39E-05 |
| ENSG00000133107 | TRPC4 | -0.31 | 0.07 | 9.70E-06 | 3.76E-05 |
| ENSG00000014216 | CAPN1 | -0.31 | 0.07 | 9.73E-06 | 3.77E-05 |
| ENSG00000153006 | SREK1IP1 | -0.31 | 0.07 | 1.62E-05 | 6.12E-05 |
| ENSG00000198189 | HSD17B11 | -0.31 | 0.07 | 1.62E-05 | 6.14E-05 |
| ENSG00000167191 | GPRC5B | -0.31 | 0.07 | 2.32E-05 | 8.63E-05 |
| ENSG00000148153 | INIP | -0.31 | 0.07 | 2.53E-05 | 9.39E-05 |
| ENSG00000148572 | NRBF2 | -0.31 | 0.08 | 3.28E-05 | 1.20E-04 |
| ENSG00000130303 | BST2 | -0.31 | 0.08 | 4.21E-05 | 1.53E-04 |
| ENSG00000137601 | NEK1 | -0.31 | 0.08 | 4.42E-05 | 1.60E-04 |
| ENSG00000078967 | UBE2D4 | -0.31 | 0.08 | 4.47E-05 | 1.61E-04 |
| ENSG00000189077 | TMEM120A | -0.31 | 0.08 | 4.55E-05 | 1.64E-04 |
| ENSG00000130810 | PPAN | -0.31 | 0.08 | 1.31E-04 | 4.47E-04 |
| ENSG00000159788 | RGS12 | -0.31 | 0.08 | 1.57E-04 | 5.33E-04 |
| ENSG00000187699 | C2orf88 | -0.31 | 0.09 | 6.11E-04 | 1.92E-03 |
| ENSG00000074047 | GLI2 | -0.31 | 0.09 | 6.56E-04 | 2.05E-03 |
| ENSG00000198945 | L3MBTL3 | -0.31 | 0.1 | 1.83E-03 | 5.35E-03 |
| ENSG00000167645 | YIF1B | -0.31 | 0.1 | 1.94E-03 | 5.64E-03 |
| ENSG00000081026 | MAGI3 | -0.31 | 0.11 | 1.97E-03 | 5.71E-03 |
| ENSG00000167771 | RCOR2 | -0.31 | 0.11 | 2.61E-03 | 7.43E-03 |
| ENSG00000164403 | SHROOM1 | -0.31 | 0.11 | 4.96E-03 | 1.34E-02 |
| ENSG00000203791 | EEF1AKMT2 | -0.31 | 0.12 | 5.10E-03 | 1.37E-02 |
| ENSG00000169084 | DHRSX | -0.31 | 0.12 | 6.14E-03 | 1.62E-02 |
| ENSG00000187626 | ZKSCAN4 | -0.31 | 0.12 | 7.29E-03 | 1.90E-02 |
| ENSG00000188033 | ZNF490 | -0.31 | 0.13 | 1.14E-02 | 2.83E-02 |
| ENSG00000122386 | ZNF205 | -0.31 | 0.14 | 1.56E-02 | 3.77E-02 |
| ENSG00000123728 | RAP2C | -0.3 | 0.04 | 5.09E-12 | 3.20E-11 |
| ENSG00000092010 | PSME1 | -0.3 | 0.05 | 2.08E-11 | 1.26E-10 |
| ENSG00000182307 | C8orf33 | -0.3 | 0.05 | 2.54E-11 | 1.53E-10 |
| ENSG00000112110 | MRPL18 | -0.3 | 0.05 | 3.13E-11 | 1.88E-10 |

| | | | | | |
|---|---|---|---|---|---|
| ENSG00000136504 | KAT7 | -0.3 | 0.05 | 5.23E-11 | 3.09E-10 |
| ENSG00000008838 | MED24 | -0.3 | 0.05 | 1.07E-10 | 6.17E-10 |
| ENSG00000106603 | COA1 | -0.3 | 0.05 | 2.78E-10 | 1.56E-09 |
| ENSG00000161654 | LSM12 | -0.3 | 0.05 | 3.10E-10 | 1.73E-09 |
| ENSG00000119402 | FBXW2 | -0.3 | 0.05 | 3.66E-10 | 2.04E-09 |
| ENSG00000169946 | ZFPM2 | -0.3 | 0.05 | 7.04E-10 | 3.85E-09 |
| ENSG00000119318 | RAD23B | -0.3 | 0.05 | 8.92E-10 | 4.83E-09 |
| ENSG00000147224 | PRPS1 | -0.3 | 0.05 | 1.12E-09 | 6.05E-09 |
| ENSG00000058804 | NDC1 | -0.3 | 0.05 | 1.80E-09 | 9.51E-09 |
| ENSG00000185565 | LSAMP | -0.3 | 0.05 | 1.86E-09 | 9.85E-09 |
| ENSG00000223501 | VPS52 | -0.3 | 0.05 | 5.19E-09 | 2.65E-08 |
| ENSG00000103415 | HMOX2 | -0.3 | 0.05 | 6.77E-09 | 3.43E-08 |
| ENSG00000091039 | OSBPL8 | -0.3 | 0.05 | 6.88E-09 | 3.48E-08 |
| ENSG00000165410 | CFL2 | -0.3 | 0.05 | 1.15E-08 | 5.72E-08 |
| ENSG00000156587 | UBE2L6 | -0.3 | 0.05 | 1.35E-08 | 6.70E-08 |
| ENSG00000105732 | ZNF574 | -0.3 | 0.06 | 5.59E-08 | 2.64E-07 |
| ENSG00000143033 | MTF2 | -0.3 | 0.06 | 9.13E-08 | 4.25E-07 |
| ENSG00000144848 | ATG3 | -0.3 | 0.06 | 1.05E-07 | 4.86E-07 |
| ENSG00000171552 | BCL2L1 | -0.3 | 0.06 | 1.32E-07 | 6.05E-07 |
| ENSG00000101444 | AHCY | -0.3 | 0.06 | 2.29E-07 | 1.03E-06 |
| ENSG00000067533 | RRP15 | -0.3 | 0.06 | 2.45E-07 | 1.10E-06 |
| ENSG00000110108 | TMEM109 | -0.3 | 0.06 | 3.37E-07 | 1.50E-06 |
| ENSG00000175054 | ATR | -0.3 | 0.06 | 3.46E-07 | 1.54E-06 |
| ENSG00000125257 | ABCC4 | -0.3 | 0.06 | 4.63E-07 | 2.03E-06 |
| ENSG00000111880 | RNGTT | -0.3 | 0.06 | 6.62E-07 | 2.86E-06 |
| ENSG00000149554 | CHEK1 | -0.3 | 0.06 | 7.38E-07 | 3.18E-06 |
| ENSG00000239306 | RBM14 | -0.3 | 0.06 | 1.75E-06 | 7.26E-06 |
| ENSG00000163104 | SMARCAD1 | -0.3 | 0.06 | 1.80E-06 | 7.45E-06 |
| ENSG00000112659 | CUL9 | -0.3 | 0.06 | 2.13E-06 | 8.76E-06 |
| ENSG00000076650 | GPATCH1 | -0.3 | 0.07 | 5.65E-06 | 2.24E-05 |
| ENSG00000100304 | TTLL12 | -0.3 | 0.07 | 6.52E-06 | 2.57E-05 |
| ENSG00000087245 | MMP2 | -0.3 | 0.07 | 7.82E-06 | 3.05E-05 |
| ENSG00000161980 | POLR3K | -0.3 | 0.07 | 8.07E-06 | 3.15E-05 |
| ENSG00000119669 | IRF2BPL | -0.3 | 0.07 | 8.46E-06 | 3.30E-05 |
| ENSG00000198805 | PNP | -0.3 | 0.07 | 9.58E-06 | 3.71E-05 |
| ENSG00000184178 | SCFD2 | -0.3 | 0.07 | 1.01E-05 | 3.91E-05 |
| ENSG00000163754 | GYG1 | -0.3 | 0.07 | 1.10E-05 | 4.24E-05 |
| ENSG00000175104 | TRAF6 | -0.3 | 0.07 | 1.69E-05 | 6.39E-05 |
| ENSG00000182173 | TSEN54 | -0.3 | 0.07 | 1.69E-05 | 6.39E-05 |
| ENSG00000102572 | STK24 | -0.3 | 0.07 | 1.76E-05 | 6.62E-05 |
| ENSG00000163364 | LINC01116 | -0.3 | 0.07 | 2.41E-05 | 8.95E-05 |
| ENSG00000083093 | PALB2 | -0.3 | 0.07 | 2.97E-05 | 1.09E-04 |
| ENSG00000214706 | IFRD2 | -0.3 | 0.07 | 3.00E-05 | 1.11E-04 |
| ENSG00000166257 | SCN3B | -0.3 | 0.07 | 3.31E-05 | 1.21E-04 |
| ENSG00000011677 | GABRA3 | -0.3 | 0.07 | 3.99E-05 | 1.45E-04 |
| ENSG00000196405 | EVL | -0.3 | 0.08 | 4.82E-05 | 1.74E-04 |
| ENSG00000020922 | MRE11 | -0.3 | 0.08 | 7.41E-05 | 2.62E-04 |
| ENSG00000143995 | MEIS1 | -0.3 | 0.08 | 7.54E-05 | 2.66E-04 |
| ENSG00000205356 | TECPR1 | -0.3 | 0.08 | 1.04E-04 | 3.62E-04 |
| ENSG00000204371 | EHMT2 | -0.3 | 0.08 | 1.10E-04 | 3.81E-04 |

| | | | | | |
|---|---|---|---|---|---|
| ENSG00000106100 | NOD1 | -0.3 | 0.08 | 1.16E-04 | 3.99E-04 |
| ENSG00000186073 | CDIN1 | -0.3 | 0.08 | 1.60E-04 | 5.44E-04 |
| ENSG00000141013 | GAS8 | -0.3 | 0.08 | 1.81E-04 | 6.09E-04 |
| ENSG00000173457 | PPP1R14B | -0.3 | 0.08 | 1.93E-04 | 6.47E-04 |
| ENSG00000132109 | TRIM21 | -0.3 | 0.08 | 2.47E-04 | 8.18E-04 |
| ENSG00000171161 | ZNF672 | -0.3 | 0.08 | 2.69E-04 | 8.89E-04 |
| ENSG00000123178 | SPRYD7 | -0.3 | 0.09 | 3.07E-04 | 1.00E-03 |
| ENSG00000155729 | KCTD18 | -0.3 | 0.09 | 3.89E-04 | 1.26E-03 |
| ENSG00000213463 | SYNJ2BP | -0.3 | 0.09 | 4.81E-04 | 1.53E-03 |
| ENSG00000121690 | DEPDC7 | -0.3 | 0.09 | 6.80E-04 | 2.12E-03 |
| ENSG00000108423 | TUBD1 | -0.3 | 0.09 | 8.91E-04 | 2.73E-03 |
| ENSG00000180035 | ZNF48 | -0.3 | 0.09 | 9.30E-04 | 2.84E-03 |
| ENSG00000104522 | GFUS | -0.3 | 0.09 | 1.01E-03 | 3.05E-03 |
| ENSG00000273604 | EPOP | -0.3 | 0.1 | 1.57E-03 | 4.63E-03 |
| ENSG00000175895 | PLEKHF2 | -0.3 | 0.1 | 2.20E-03 | 6.31E-03 |
| ENSG00000187630 | DHRS4L2 | -0.3 | 0.1 | 2.46E-03 | 7.01E-03 |
| ENSG00000140470 | ADAMTS17 | -0.3 | 0.11 | 5.01E-03 | 1.35E-02 |
| ENSG00000155957 | TMBIM4 | -0.3 | 0.11 | 5.51E-03 | 1.47E-02 |
| ENSG00000177842 | ZNF620 | -0.3 | 0.11 | 6.18E-03 | 1.63E-02 |
| ENSG00000129559 | NEDD8 | -0.3 | 0.13 | 1.15E-02 | 2.86E-02 |
| ENSG00000089163 | SIRT4 | -0.3 | 0.13 | 1.31E-02 | 3.22E-02 |
| ENSG00000128710 | HOXD10 | -0.3 | 0.13 | 1.68E-02 | 4.02E-02 |
| ENSG00000187091 | PLCD1 | -0.3 | 0.14 | 2.08E-02 | 4.88E-02 |
| ENSG00000158710 | TAGLN2 | -0.29 | 0.03 | 2.28E-20 | 2.16E-19 |
| ENSG00000133030 | MPRIP | -0.29 | 0.03 | 5.59E-17 | 4.55E-16 |
| ENSG00000136193 | SCRN1 | -0.29 | 0.04 | 2.50E-15 | 1.88E-14 |
| ENSG00000085662 | AKR1B1 | -0.29 | 0.04 | 2.68E-15 | 2.02E-14 |
| ENSG00000035403 | VCL | -0.29 | 0.04 | 8.24E-14 | 5.75E-13 |
| ENSG00000121060 | TRIM25 | -0.29 | 0.04 | 9.99E-13 | 6.55E-12 |
| ENSG00000171365 | CLCN5 | -0.29 | 0.04 | 2.78E-11 | 1.67E-10 |
| ENSG00000198231 | DDX42 | -0.29 | 0.05 | 1.07E-10 | 6.18E-10 |
| ENSG00000152104 | PTPN14 | -0.29 | 0.05 | 2.70E-10 | 1.52E-09 |
| ENSG00000178904 | DPY19L3 | -0.29 | 0.05 | 8.09E-10 | 4.39E-09 |
| ENSG00000100823 | APEX1 | -0.29 | 0.05 | 1.27E-09 | 6.81E-09 |
| ENSG00000109586 | GALNT7 | -0.29 | 0.05 | 1.70E-09 | 9.04E-09 |
| ENSG00000135632 | SMYD5 | -0.29 | 0.05 | 5.73E-09 | 2.92E-08 |
| ENSG00000121297 | TSHZ3 | -0.29 | 0.05 | 9.16E-09 | 4.60E-08 |
| ENSG00000005194 | CIAPIN1 | -0.29 | 0.05 | 1.17E-08 | 5.81E-08 |
| ENSG00000016864 | GLT8D1 | -0.29 | 0.05 | 3.88E-08 | 1.86E-07 |
| ENSG00000139514 | SLC7A1 | -0.29 | 0.05 | 7.76E-08 | 3.63E-07 |
| ENSG00000138166 | DUSP5 | -0.29 | 0.06 | 3.82E-07 | 1.69E-06 |
| ENSG00000168014 | C2CD3 | -0.29 | 0.06 | 6.17E-07 | 2.68E-06 |
| ENSG00000105176 | URI1 | -0.29 | 0.06 | 1.23E-06 | 5.16E-06 |
| ENSG00000148180 | GSN | -0.29 | 0.06 | 1.40E-06 | 5.87E-06 |
| ENSG00000275464 | | -0.29 | 0.06 | 1.64E-06 | 6.81E-06 |
| ENSG00000119965 | C10orf88 | -0.29 | 0.06 | 2.10E-06 | 8.64E-06 |
| ENSG00000144909 | OSBPL11 | -0.29 | 0.06 | 2.38E-06 | 9.76E-06 |
| ENSG00000164983 | TMEM65 | -0.29 | 0.06 | 2.59E-06 | 1.06E-05 |
| ENSG00000135945 | REV1 | -0.29 | 0.06 | 2.93E-06 | 1.19E-05 |
| ENSG00000183762 | KREMEN1 | -0.29 | 0.06 | 4.78E-06 | 1.91E-05 |

| | | | | | |
|---|---|---|---|---|---|
| ENSG00000133641 | C12orf29 | -0.29 | 0.06 | 5.51E-06 | 2.19E-05 |
| ENSG00000123545 | NDUFAF4 | -0.29 | 0.07 | 1.17E-05 | 4.48E-05 |
| ENSG00000106459 | NRF1 | -0.29 | 0.07 | 1.65E-05 | 6.25E-05 |
| ENSG00000162804 | SNED1 | -0.29 | 0.07 | 2.69E-05 | 9.93E-05 |
| ENSG00000090989 | EXOC1 | -0.29 | 0.07 | 2.94E-05 | 1.08E-04 |
| ENSG00000101464 | PIGU | -0.29 | 0.07 | 3.27E-05 | 1.20E-04 |
| ENSG00000100056 | ESS2 | -0.29 | 0.08 | 1.08E-04 | 3.73E-04 |
| ENSG00000183513 | COA5 | -0.29 | 0.08 | 1.15E-04 | 3.97E-04 |
| ENSG00000148704 | VAX1 | -0.29 | 0.08 | 1.35E-04 | 4.61E-04 |
| ENSG00000197619 | ZNF615 | -0.29 | 0.08 | 2.58E-04 | 8.53E-04 |
| ENSG00000164136 | IL15 | -0.29 | 0.08 | 3.68E-04 | 1.19E-03 |
| ENSG00000176720 | BOK | -0.29 | 0.09 | 6.32E-04 | 1.98E-03 |
| ENSG00000167280 | ENGASE | -0.29 | 0.1 | 2.05E-03 | 5.93E-03 |
| ENSG00000174165 | ZDHHC24 | -0.29 | 0.1 | 3.46E-03 | 9.60E-03 |
| ENSG00000164011 | ZNF691 | -0.29 | 0.11 | 6.73E-03 | 1.77E-02 |
| ENSG00000172889 | EGFL7 | -0.29 | 0.11 | 7.38E-03 | 1.92E-02 |
| ENSG00000151665 | PIGF | -0.29 | 0.12 | 9.36E-03 | 2.38E-02 |
| ENSG00000278730 | | -0.29 | 0.13 | 1.81E-02 | 4.30E-02 |
| ENSG00000237943 | PRKCQ-AS1 | -0.29 | 0.13 | 1.85E-02 | 4.38E-02 |
| ENSG00000131467 | PSME3 | -0.28 | 0.03 | 3.47E-17 | 2.85E-16 |
| ENSG00000137955 | RABGGTB | -0.28 | 0.04 | 2.30E-12 | 1.48E-11 |
| ENSG00000088888 | MAVS | -0.28 | 0.04 | 9.15E-12 | 5.66E-11 |
| ENSG00000134697 | GNL2 | -0.28 | 0.04 | 2.02E-10 | 1.15E-09 |
| ENSG00000266028 | SRGAP2 | -0.28 | 0.05 | 1.35E-09 | 7.22E-09 |
| ENSG00000102024 | PLS3 | -0.28 | 0.05 | 1.67E-09 | 8.88E-09 |
| ENSG00000120885 | CLU | -0.28 | 0.05 | 1.97E-09 | 1.04E-08 |
| ENSG00000172840 | PDP2 | -0.28 | 0.05 | 4.71E-09 | 2.41E-08 |
| ENSG00000144381 | HSPD1 | -0.28 | 0.05 | 2.14E-08 | 1.04E-07 |
| ENSG00000153707 | PTPRD | -0.28 | 0.05 | 3.18E-08 | 1.53E-07 |
| ENSG00000112081 | SRSF3 | -0.28 | 0.05 | 7.26E-08 | 3.40E-07 |
| ENSG00000147471 | PLPBP | -0.28 | 0.05 | 7.44E-08 | 3.49E-07 |
| ENSG00000119720 | NRDE2 | -0.28 | 0.06 | 3.76E-07 | 1.66E-06 |
| ENSG00000117859 | OSBPL9 | -0.28 | 0.06 | 3.83E-07 | 1.69E-06 |
| ENSG00000135801 | TAF5L | -0.28 | 0.06 | 3.98E-07 | 1.76E-06 |
| ENSG00000117528 | ABCD3 | -0.28 | 0.06 | 4.17E-07 | 1.84E-06 |
| ENSG00000150275 | PCDH15 | -0.28 | 0.06 | 4.72E-07 | 2.07E-06 |
| ENSG00000196943 | NOP9 | -0.28 | 0.06 | 1.12E-06 | 4.74E-06 |
| ENSG00000147316 | MCPH1 | -0.28 | 0.06 | 2.39E-06 | 9.79E-06 |
| ENSG00000067177 | PHKA1 | -0.28 | 0.06 | 3.19E-06 | 1.29E-05 |
| ENSG00000070778 | PTPN21 | -0.28 | 0.06 | 3.79E-06 | 1.53E-05 |
| ENSG00000002834 | LASP1 | -0.28 | 0.06 | 3.88E-06 | 1.56E-05 |
| ENSG00000100207 | TCF20 | -0.28 | 0.06 | 4.22E-06 | 1.69E-05 |
| ENSG00000121879 | PIK3CA | -0.28 | 0.06 | 4.41E-06 | 1.76E-05 |
| ENSG00000121903 | ZSCAN20 | -0.28 | 0.06 | 4.66E-06 | 1.86E-05 |
| ENSG00000068383 | INPP5A | -0.28 | 0.06 | 4.83E-06 | 1.93E-05 |
| ENSG00000167515 | TRAPPC2L | -0.28 | 0.06 | 8.59E-06 | 3.34E-05 |
| ENSG00000181827 | RFX7 | -0.28 | 0.06 | 1.01E-05 | 3.91E-05 |
| ENSG00000073536 | NLE1 | -0.28 | 0.07 | 1.14E-05 | 4.40E-05 |
| ENSG00000145293 | ENOPH1 | -0.28 | 0.06 | 1.46E-05 | 5.55E-05 |
| ENSG00000143772 | ITPKB | -0.28 | 0.07 | 1.60E-05 | 6.07E-05 |

| | | | | | |
|---|---|---|---|---|---|
| ENSG00000196865 | NHLRC2 | -0.28 | 0.07 | 2.57E-05 | 9.50E-05 |
| ENSG00000184220 | CMSS1 | -0.28 | 0.07 | 5.33E-05 | 1.91E-04 |
| ENSG00000167182 | SP2 | -0.28 | 0.07 | 8.04E-05 | 2.83E-04 |
| ENSG00000103591 | AAGAB | -0.28 | 0.07 | 9.47E-05 | 3.30E-04 |
| ENSG00000154767 | XPC | -0.28 | 0.07 | 9.77E-05 | 3.40E-04 |
| ENSG00000171148 | TADA3 | -0.28 | 0.07 | 1.19E-04 | 4.11E-04 |
| ENSG00000173540 | GMPPB | -0.28 | 0.08 | 1.27E-04 | 4.36E-04 |
| ENSG00000083642 | PDS5B | -0.28 | 0.07 | 1.42E-04 | 4.85E-04 |
| ENSG00000134184 | GSTM1 | -0.28 | 0.07 | 1.46E-04 | 4.99E-04 |
| ENSG00000127989 | MTERF1 | -0.28 | 0.08 | 1.56E-04 | 5.28E-04 |
| ENSG00000148291 | SURF2 | -0.28 | 0.08 | 2.26E-04 | 7.54E-04 |
| ENSG00000197429 | IPP | -0.28 | 0.08 | 2.27E-04 | 7.57E-04 |
| ENSG00000107959 | PITRM1 | -0.28 | 0.08 | 4.00E-04 | 1.29E-03 |
| ENSG00000141040 | ZNF287 | -0.28 | 0.09 | 6.28E-04 | 1.97E-03 |
| ENSG00000134809 | TIMM10 | -0.28 | 0.08 | 7.04E-04 | 2.20E-03 |
| ENSG00000185236 | RAB11B | -0.28 | 0.08 | 7.83E-04 | 2.42E-03 |
| ENSG00000182378 | PLCXD1 | -0.28 | 0.09 | 1.05E-03 | 3.17E-03 |
| ENSG00000112167 | SAYSD1 | -0.28 | 0.09 | 1.11E-03 | 3.35E-03 |
| ENSG00000186166 | CENATAC | -0.28 | 0.09 | 1.48E-03 | 4.37E-03 |
| ENSG00000102390 | PBDC1 | -0.28 | 0.09 | 1.85E-03 | 5.38E-03 |
| ENSG00000034533 | ASTE1 | -0.28 | 0.1 | 2.49E-03 | 7.11E-03 |
| ENSG00000150756 | ATPSCKMT | -0.28 | 0.1 | 2.77E-03 | 7.82E-03 |
| ENSG00000188171 | ZNF626 | -0.28 | 0.1 | 2.79E-03 | 7.89E-03 |
| ENSG00000138101 | DTNB | -0.28 | 0.1 | 3.10E-03 | 8.69E-03 |
| ENSG00000150433 | TMEM218 | -0.28 | 0.1 | 3.17E-03 | 8.87E-03 |
| ENSG00000104894 | CD37 | -0.28 | 0.1 | 5.66E-03 | 1.51E-02 |
| ENSG00000164093 | PITX2 | -0.28 | 0.11 | 5.95E-03 | 1.58E-02 |
| ENSG00000168792 | ABHD15 | -0.28 | 0.11 | 6.68E-03 | 1.75E-02 |
| ENSG00000108984 | MAP2K6 | -0.28 | 0.11 | 8.02E-03 | 2.07E-02 |
| ENSG00000150456 | EEF1AKMT1 | -0.28 | 0.12 | 1.07E-02 | 2.68E-02 |
| ENSG00000179886 | TIGD5 | -0.28 | 0.12 | 1.49E-02 | 3.61E-02 |
| ENSG00000204366 | ZBTB12 | -0.28 | 0.12 | 1.53E-02 | 3.70E-02 |
| ENSG00000171004 | HS6ST2 | -0.27 | 0.04 | 2.82E-14 | 2.02E-13 |
| ENSG00000164683 | HEY1 | -0.27 | 0.04 | 3.19E-14 | 2.28E-13 |
| ENSG00000155561 | NUP205 | -0.27 | 0.04 | 1.06E-12 | 6.93E-12 |
| ENSG00000168439 | STIP1 | -0.27 | 0.04 | 4.06E-12 | 2.57E-11 |
| ENSG00000161547 | SRSF2 | -0.27 | 0.04 | 2.12E-10 | 1.21E-09 |
| ENSG00000133606 | MKRN1 | -0.27 | 0.04 | 4.85E-10 | 2.68E-09 |
| ENSG00000071967 | CYBRD1 | -0.27 | 0.04 | 8.34E-10 | 4.52E-09 |
| ENSG00000085978 | ATG16L1 | -0.27 | 0.05 | 3.17E-09 | 1.65E-08 |
| ENSG00000010270 | STARD3NL | -0.27 | 0.05 | 4.55E-09 | 2.33E-08 |
| ENSG00000070770 | CSNK2A2 | -0.27 | 0.05 | 8.55E-09 | 4.30E-08 |
| ENSG00000077097 | TOP2B | -0.27 | 0.05 | 1.11E-08 | 5.57E-08 |
| ENSG00000116161 | CACYBP | -0.27 | 0.05 | 1.71E-08 | 8.41E-08 |
| ENSG00000226479 | TMEM185B | -0.27 | 0.05 | 2.10E-08 | 1.03E-07 |
| ENSG00000087191 | PSMC5 | -0.27 | 0.05 | 3.19E-08 | 1.54E-07 |
| ENSG00000204592 | HLA-E | -0.27 | 0.05 | 3.21E-08 | 1.54E-07 |
| ENSG00000091986 | CCDC80 | -0.27 | 0.05 | 6.53E-08 | 3.07E-07 |
| ENSG00000131149 | GSE1 | -0.27 | 0.05 | 9.78E-08 | 4.54E-07 |
| ENSG00000075568 | TMEM131 | -0.27 | 0.05 | 1.02E-07 | 4.74E-07 |

| | | | | | |
|---|---|---|---|---|---|
| ENSG00000133104 | SPART | -0.27 | 0.05 | 4.55E-07 | 2.00E-06 |
| ENSG00000181610 | MRPS23 | -0.27 | 0.06 | 9.84E-07 | 4.18E-06 |
| ENSG00000066135 | KDM4A | -0.27 | 0.06 | 1.24E-06 | 5.20E-06 |
| ENSG00000247626 | MARS2 | -0.27 | 0.06 | 1.35E-06 | 5.66E-06 |
| ENSG00000124795 | DEK | -0.27 | 0.06 | 6.40E-06 | 2.52E-05 |
| ENSG00000142892 | PIGK | -0.27 | 0.06 | 7.42E-06 | 2.90E-05 |
| ENSG00000184708 | EIF4ENIF1 | -0.27 | 0.06 | 7.45E-06 | 2.92E-05 |
| ENSG00000167315 | ACAA2 | -0.27 | 0.06 | 8.29E-06 | 3.23E-05 |
| ENSG00000122359 | ANXA11 | -0.27 | 0.06 | 8.50E-06 | 3.31E-05 |
| ENSG00000107863 | ARHGAP21 | -0.27 | 0.06 | 1.20E-05 | 4.59E-05 |
| ENSG00000164190 | NIPBL | -0.27 | 0.06 | 1.20E-05 | 4.62E-05 |
| ENSG00000134318 | ROCK2 | -0.27 | 0.06 | 1.39E-05 | 5.30E-05 |
| ENSG00000105676 | ARMC6 | -0.27 | 0.06 | 1.44E-05 | 5.46E-05 |
| ENSG00000066654 | THUMPD1 | -0.27 | 0.06 | 1.83E-05 | 6.90E-05 |
| ENSG00000102699 | PARP4 | -0.27 | 0.06 | 2.17E-05 | 8.11E-05 |
| ENSG00000010539 | ZNF200 | -0.27 | 0.07 | 2.42E-05 | 9.00E-05 |
| ENSG00000100523 | DDHD1 | -0.27 | 0.07 | 3.29E-05 | 1.21E-04 |
| ENSG00000123975 | CKS2 | -0.27 | 0.07 | 3.37E-05 | 1.23E-04 |
| ENSG00000138593 | SECISBP2L | -0.27 | 0.07 | 3.67E-05 | 1.34E-04 |
| ENSG00000170522 | ELOVL6 | -0.27 | 0.07 | 3.80E-05 | 1.38E-04 |
| ENSG00000236345 | SCAT8 | -0.27 | 0.07 | 3.95E-05 | 1.44E-04 |
| ENSG00000159055 | MIS18A | -0.27 | 0.07 | 4.31E-05 | 1.56E-04 |
| ENSG00000185222 | TCEAL9 | -0.27 | 0.07 | 4.80E-05 | 1.73E-04 |
| ENSG00000254986 | DPP3 | -0.27 | 0.07 | 4.88E-05 | 1.76E-04 |
| ENSG00000040275 | SPDL1 | -0.27 | 0.07 | 1.03E-04 | 3.58E-04 |
| ENSG00000114767 | RRP9 | -0.27 | 0.07 | 1.03E-04 | 3.58E-04 |
| ENSG00000170191 | NANP | -0.27 | 0.07 | 1.19E-04 | 4.10E-04 |
| ENSG00000197818 | SLC9A8 | -0.27 | 0.07 | 1.27E-04 | 4.37E-04 |
| ENSG00000075407 | ZNF37A | -0.27 | 0.07 | 1.46E-04 | 4.96E-04 |
| ENSG00000037042 | TUBG2 | -0.27 | 0.07 | 1.67E-04 | 5.66E-04 |
| ENSG00000143622 | RIT1 | -0.27 | 0.08 | 2.26E-04 | 7.53E-04 |
| ENSG00000173409 | ARV1 | -0.27 | 0.08 | 2.35E-04 | 7.82E-04 |
| ENSG00000187866 | PABIR1 | -0.27 | 0.08 | 2.79E-04 | 9.17E-04 |
| ENSG00000131845 | ZNF304 | -0.27 | 0.08 | 3.20E-04 | 1.05E-03 |
| ENSG00000139405 | RITA1 | -0.27 | 0.08 | 3.50E-04 | 1.14E-03 |
| ENSG00000113838 | TBCCD1 | -0.27 | 0.08 | 4.40E-04 | 1.41E-03 |
| ENSG00000213366 | GSTM2 | -0.27 | 0.08 | 5.32E-04 | 1.69E-03 |
| ENSG00000181751 | MACIR | -0.27 | 0.08 | 5.87E-04 | 1.85E-03 |
| ENSG00000184939 | ZFP90 | -0.27 | 0.08 | 5.91E-04 | 1.86E-03 |
| ENSG00000160360 | GPSM1 | -0.27 | 0.08 | 5.98E-04 | 1.88E-03 |
| ENSG00000169599 | NFU1 | -0.27 | 0.08 | 7.21E-04 | 2.24E-03 |
| ENSG00000134115 | CNTN6 | -0.27 | 0.08 | 7.31E-04 | 2.27E-03 |
| ENSG00000187244 | BCAM | -0.27 | 0.08 | 8.60E-04 | 2.64E-03 |
| ENSG00000176396 | EID2 | -0.27 | 0.08 | 9.03E-04 | 2.76E-03 |
| ENSG00000107902 | LHPP | -0.27 | 0.08 | 9.77E-04 | 2.98E-03 |
| ENSG00000162869 | PPP1R21 | -0.27 | 0.09 | 1.27E-03 | 3.79E-03 |
| ENSG00000142082 | SIRT3 | -0.27 | 0.09 | 1.52E-03 | 4.49E-03 |
| ENSG00000148335 | NTMT1 | -0.27 | 0.09 | 2.17E-03 | 6.25E-03 |
| ENSG00000177875 | CCDC184 | -0.27 | 0.09 | 2.53E-03 | 7.21E-03 |
| ENSG00000165066 | NKX6-3 | -0.27 | 0.1 | 3.99E-03 | 1.09E-02 |

| | | | | | |
|---|---|---|---|---|---|
| ENSG00000139211 | AMIGO2 | -0.27 | 0.1 | 4.01E-03 | 1.10E-02 |
| ENSG00000130347 | RTN4IP1 | -0.27 | 0.1 | 6.39E-03 | 1.68E-02 |
| ENSG00000173153 | ESRRA | -0.27 | 0.11 | 7.62E-03 | 1.98E-02 |
| ENSG00000066735 | KIF26A | -0.27 | 0.11 | 8.13E-03 | 2.10E-02 |
| ENSG00000172086 | KRCC1 | -0.27 | 0.11 | 9.21E-03 | 2.35E-02 |
| ENSG00000103152 | MPG | -0.27 | 0.11 | 9.30E-03 | 2.37E-02 |
| ENSG00000122490 | SLC66A2 | -0.27 | 0.11 | 9.53E-03 | 2.42E-02 |
| ENSG00000167460 | TPM4 | -0.26 | 0.03 | 4.73E-16 | 3.69E-15 |
| ENSG00000111145 | ELK3 | -0.26 | 0.04 | 1.66E-12 | 1.07E-11 |
| ENSG00000124571 | XPO5 | -0.26 | 0.04 | 1.14E-11 | 7.00E-11 |
| ENSG00000116729 | WLS | -0.26 | 0.04 | 1.69E-10 | 9.63E-10 |
| ENSG00000134910 | STT3A | -0.26 | 0.04 | 2.99E-10 | 1.68E-09 |
| ENSG00000135932 | CAB39 | -0.26 | 0.04 | 3.08E-10 | 1.73E-09 |
| ENSG00000135387 | CAPRIN1 | -0.26 | 0.04 | 3.27E-10 | 1.83E-09 |
| ENSG00000114867 | EIF4G1 | -0.26 | 0.04 | 2.21E-09 | 1.16E-08 |
| ENSG00000173207 | CKS1B | -0.26 | 0.05 | 1.53E-08 | 7.56E-08 |
| ENSG00000153786 | ZDHHC7 | -0.26 | 0.05 | 2.43E-08 | 1.18E-07 |
| ENSG00000124205 | EDN3 | -0.26 | 0.05 | 9.63E-08 | 4.48E-07 |
| ENSG00000017797 | RALBP1 | -0.26 | 0.05 | 1.60E-07 | 7.30E-07 |
| ENSG00000130640 | TUBGCP2 | -0.26 | 0.05 | 2.28E-07 | 1.03E-06 |
| ENSG00000215012 | RTL10 | -0.26 | 0.05 | 2.50E-07 | 1.12E-06 |
| ENSG00000144711 | IQSEC1 | -0.26 | 0.05 | 8.47E-07 | 3.62E-06 |
| ENSG00000011275 | RNF216 | -0.26 | 0.06 | 1.97E-06 | 8.14E-06 |
| ENSG00000134278 | SPIRE1 | -0.26 | 0.06 | 2.35E-06 | 9.64E-06 |
| ENSG00000107643 | MAPK8 | -0.26 | 0.06 | 3.21E-06 | 1.30E-05 |
| ENSG00000186283 | TOR3A | -0.26 | 0.06 | 4.00E-06 | 1.61E-05 |
| ENSG00000174953 | DHX36 | -0.26 | 0.06 | 4.50E-06 | 1.80E-05 |
| ENSG00000069275 | NUCKS1 | -0.26 | 0.06 | 1.50E-05 | 5.70E-05 |
| ENSG00000103111 | MON1B | -0.26 | 0.06 | 1.55E-05 | 5.87E-05 |
| ENSG00000123415 | SMUG1 | -0.26 | 0.06 | 1.65E-05 | 6.25E-05 |
| ENSG00000112379 | ARFGEF3 | -0.26 | 0.06 | 1.86E-05 | 7.01E-05 |
| ENSG00000085274 | MYNN | -0.26 | 0.06 | 2.96E-05 | 1.09E-04 |
| ENSG00000136840 | ST6GALNAC4 | -0.26 | 0.07 | 4.40E-05 | 1.59E-04 |
| ENSG00000112237 | CCNC | -0.26 | 0.06 | 5.37E-05 | 1.93E-04 |
| ENSG00000178951 | ZBTB7A | -0.26 | 0.07 | 6.28E-05 | 2.24E-04 |
| ENSG00000156469 | MTERF3 | -0.26 | 0.07 | 7.09E-05 | 2.51E-04 |
| ENSG00000101333 | PLCB4 | -0.26 | 0.07 | 7.54E-05 | 2.66E-04 |
| ENSG00000138663 | COPS4 | -0.26 | 0.07 | 9.07E-05 | 3.17E-04 |
| ENSG00000213995 | NAXD | -0.26 | 0.07 | 1.00E-04 | 3.49E-04 |
| ENSG00000187720 | THSD4 | -0.26 | 0.07 | 1.05E-04 | 3.65E-04 |
| ENSG00000053900 | ANAPC4 | -0.26 | 0.07 | 1.13E-04 | 3.91E-04 |
| ENSG00000107338 | SHB | -0.26 | 0.07 | 1.22E-04 | 4.19E-04 |
| ENSG00000176273 | SLC35G1 | -0.26 | 0.07 | 1.39E-04 | 4.73E-04 |
| ENSG00000173145 | NOC3L | -0.26 | 0.07 | 1.55E-04 | 5.25E-04 |
| ENSG00000185278 | ZBTB37 | -0.26 | 0.07 | 1.64E-04 | 5.56E-04 |
| ENSG00000198042 | MAK16 | -0.26 | 0.07 | 1.65E-04 | 5.60E-04 |
| ENSG00000148334 | PTGES2 | -0.26 | 0.07 | 2.01E-04 | 6.75E-04 |
| ENSG00000145012 | LPP | -0.26 | 0.07 | 2.69E-04 | 8.90E-04 |
| ENSG00000189007 | ADAT2 | -0.26 | 0.08 | 3.82E-04 | 1.24E-03 |
| ENSG00000130726 | TRIM28 | -0.26 | 0.08 | 4.89E-04 | 1.56E-03 |

| | | | | | |
|---|---|---|---|---|---|
| ENSG00000183161 | FANCF | -0.26 | 0.08 | 6.32E-04 | 1.98E-03 |
| ENSG00000183751 | TBL3 | -0.26 | 0.08 | 7.10E-04 | 2.21E-03 |
| ENSG00000123427 | EEF1AKMT3 | -0.26 | 0.08 | 7.48E-04 | 2.32E-03 |
| ENSG00000175467 | SART1 | -0.26 | 0.08 | 8.54E-04 | 2.63E-03 |
| ENSG00000183828 | NUDT14 | -0.26 | 0.08 | 1.36E-03 | 4.05E-03 |
| ENSG00000174365 | SNHG11 | -0.26 | 0.09 | 3.02E-03 | 8.47E-03 |
| ENSG00000006757 | PNPLA4 | -0.26 | 0.09 | 3.31E-03 | 9.22E-03 |
| ENSG00000135045 | C9orf40 | -0.26 | 0.09 | 4.42E-03 | 1.20E-02 |
| ENSG00000196814 | MVB12B | -0.26 | 0.11 | 1.45E-02 | 3.52E-02 |
| ENSG00000165046 | LETM2 | -0.26 | 0.12 | 1.80E-02 | 4.29E-02 |
| ENSG00000214562 | NUTM2D | -0.26 | 0.12 | 1.81E-02 | 4.31E-02 |
| ENSG00000112769 | LAMA4 | -0.25 | 0.04 | 3.65E-12 | 2.31E-11 |
| ENSG00000164733 | CTSB | -0.25 | 0.04 | 7.60E-10 | 4.14E-09 |
| ENSG00000135624 | CCT7 | -0.25 | 0.04 | 8.40E-10 | 4.55E-09 |
| ENSG00000130713 | EXOSC2 | -0.25 | 0.04 | 2.14E-09 | 1.13E-08 |
| ENSG00000127483 | HP1BP3 | -0.25 | 0.04 | 3.31E-09 | 1.72E-08 |
| ENSG00000178202 | POGLUT3 | -0.25 | 0.04 | 4.98E-09 | 2.54E-08 |
| ENSG00000117868 | ESYT2 | -0.25 | 0.04 | 5.65E-09 | 2.88E-08 |
| ENSG00000197771 | MCMBP | -0.25 | 0.05 | 2.60E-08 | 1.26E-07 |
| ENSG00000165732 | DDX21 | -0.25 | 0.05 | 1.07E-07 | 4.94E-07 |
| ENSG00000136709 | WDR33 | -0.25 | 0.05 | 3.35E-07 | 1.49E-06 |
| ENSG00000188612 | SUMO2 | -0.25 | 0.05 | 4.71E-07 | 2.07E-06 |
| ENSG00000143093 | STRIP1 | -0.25 | 0.05 | 9.69E-07 | 4.12E-06 |
| ENSG00000132300 | PTCD3 | -0.25 | 0.05 | 9.73E-07 | 4.13E-06 |
| ENSG00000100350 | FOXRED2 | -0.25 | 0.05 | 1.15E-06 | 4.86E-06 |
| ENSG00000100266 | PACSIN2 | -0.25 | 0.05 | 1.40E-06 | 5.87E-06 |
| ENSG00000142875 | PRKACB | -0.25 | 0.05 | 1.52E-06 | 6.35E-06 |
| ENSG00000147130 | ZMYM3 | -0.25 | 0.05 | 1.73E-06 | 7.18E-06 |
| ENSG00000198874 | TYW1 | -0.25 | 0.05 | 2.18E-06 | 8.98E-06 |
| ENSG00000180263 | FGD6 | -0.25 | 0.05 | 2.73E-06 | 1.12E-05 |
| ENSG00000204852 | TCTN1 | -0.25 | 0.05 | 2.83E-06 | 1.15E-05 |
| ENSG00000065559 | MAP2K4 | -0.25 | 0.06 | 5.67E-06 | 2.24E-05 |
| ENSG00000068305 | MEF2A | -0.25 | 0.06 | 7.17E-06 | 2.81E-05 |
| ENSG00000140718 | FTO | -0.25 | 0.06 | 7.96E-06 | 3.11E-05 |
| ENSG00000125651 | GTF2F1 | -0.25 | 0.06 | 1.15E-05 | 4.44E-05 |
| ENSG00000155100 | OTUD6B | -0.25 | 0.06 | 1.21E-05 | 4.65E-05 |
| ENSG00000100911 | PSME2 | -0.25 | 0.06 | 1.64E-05 | 6.21E-05 |
| ENSG00000137513 | NARS2 | -0.25 | 0.06 | 1.93E-05 | 7.26E-05 |
| ENSG00000165282 | PIGO | -0.25 | 0.06 | 2.19E-05 | 8.19E-05 |
| ENSG00000155959 | VBP1 | -0.25 | 0.06 | 3.53E-05 | 1.29E-04 |
| ENSG00000143401 | ANP32E | -0.25 | 0.06 | 4.36E-05 | 1.58E-04 |
| ENSG00000010017 | RANBP9 | -0.25 | 0.06 | 4.94E-05 | 1.78E-04 |
| ENSG00000169231 | THBS3 | -0.25 | 0.06 | 5.70E-05 | 2.04E-04 |
| ENSG00000153933 | DGKE | -0.25 | 0.06 | 6.28E-05 | 2.24E-04 |
| ENSG00000136925 | TSTD2 | -0.25 | 0.06 | 6.95E-05 | 2.46E-04 |
| ENSG00000165359 | INTS6L | -0.25 | 0.06 | 8.46E-05 | 2.97E-04 |
| ENSG00000178057 | NDUFAF3 | -0.25 | 0.06 | 9.41E-05 | 3.28E-04 |
| ENSG00000111321 | LTBR | -0.25 | 0.06 | 9.52E-05 | 3.32E-04 |
| ENSG00000138685 | FGF2 | -0.25 | 0.07 | 1.32E-04 | 4.52E-04 |
| ENSG00000076604 | TRAF4 | -0.25 | 0.07 | 1.55E-04 | 5.25E-04 |

| | | | | | |
|---|---|---|---|---|---|
| ENSG00000198440 | ZNF583 | -0.25 | 0.07 | 2.01E-04 | 6.73E-04 |
| ENSG00000197024 | ZNF398 | -0.25 | 0.07 | 2.35E-04 | 7.82E-04 |
| ENSG00000126653 | NSRP1 | -0.25 | 0.07 | 3.92E-04 | 1.26E-03 |
| ENSG00000042493 | CAPG | -0.25 | 0.07 | 5.64E-04 | 1.78E-03 |
| ENSG00000135899 | SP110 | -0.25 | 0.08 | 6.64E-04 | 2.08E-03 |
| ENSG00000135698 | MPHOSPH6 | -0.25 | 0.07 | 7.70E-04 | 2.38E-03 |
| ENSG00000063244 | U2AF2 | -0.25 | 0.08 | 8.83E-04 | 2.71E-03 |
| ENSG00000167323 | STIM1 | -0.25 | 0.08 | 9.90E-04 | 3.01E-03 |
| ENSG00000110060 | PUS3 | -0.25 | 0.08 | 1.38E-03 | 4.10E-03 |
| ENSG00000152404 | CWF19L2 | -0.25 | 0.08 | 1.98E-03 | 5.73E-03 |
| ENSG00000196236 | XPNPEP3 | -0.25 | 0.08 | 2.00E-03 | 5.80E-03 |
| ENSG00000155621 | C9orf85 | -0.25 | 0.08 | 2.42E-03 | 6.92E-03 |
| ENSG00000167775 | CD320 | -0.25 | 0.08 | 2.49E-03 | 7.11E-03 |
| ENSG00000185024 | BRF1 | -0.25 | 0.09 | 2.71E-03 | 7.67E-03 |
| ENSG00000104866 | PPP1R37 | -0.25 | 0.09 | 3.29E-03 | 9.17E-03 |
| ENSG00000005436 | GCFC2 | -0.25 | 0.09 | 3.53E-03 | 9.78E-03 |
| ENSG00000141179 | PCTP | -0.25 | 0.09 | 4.97E-03 | 1.34E-02 |
| ENSG00000160447 | PKN3 | -0.25 | 0.1 | 7.83E-03 | 2.03E-02 |
| ENSG00000186868 | MAPT | -0.25 | 0.1 | 8.11E-03 | 2.09E-02 |
| ENSG00000102547 | CAB39L | -0.25 | 0.1 | 8.22E-03 | 2.12E-02 |
| ENSG00000130005 | GAMT | -0.25 | 0.1 | 8.43E-03 | 2.17E-02 |
| ENSG00000047230 | CTPS2 | -0.25 | 0.1 | 9.64E-03 | 2.44E-02 |
| ENSG00000196418 | ZNF124 | -0.25 | 0.1 | 1.26E-02 | 3.11E-02 |
| ENSG00000185947 | ZNF267 | -0.25 | 0.11 | 1.47E-02 | 3.57E-02 |
| ENSG00000197566 | ZNF624 | -0.25 | 0.11 | 1.56E-02 | 3.76E-02 |
| ENSG00000169740 | ZNF32 | -0.25 | 0.11 | 1.69E-02 | 4.04E-02 |
| ENSG00000102302 | FGD1 | -0.25 | 0.11 | 1.74E-02 | 4.16E-02 |
| ENSG00000221866 | PLXNA4 | -0.25 | 0.11 | 1.85E-02 | 4.38E-02 |
| ENSG00000181619 | GPR135 | -0.25 | 0.11 | 1.87E-02 | 4.44E-02 |
| ENSG00000120438 | TCP1 | -0.24 | 0.04 | 2.28E-10 | 1.29E-09 |
| ENSG00000125166 | GOT2 | -0.24 | 0.04 | 2.76E-10 | 1.55E-09 |
| ENSG00000157654 | PALM2AKAP2 | -0.24 | 0.04 | 3.07E-09 | 1.60E-08 |
| ENSG00000153207 | AHCTF1 | -0.24 | 0.04 | 4.29E-09 | 2.20E-08 |
| ENSG00000167258 | CDK12 | -0.24 | 0.04 | 5.76E-09 | 2.93E-08 |
| ENSG00000006744 | ELAC2 | -0.24 | 0.04 | 1.22E-08 | 6.07E-08 |
| ENSG00000134954 | ETS1 | -0.24 | 0.04 | 4.90E-08 | 2.33E-07 |
| ENSG00000011114 | BTBD7 | -0.24 | 0.05 | 8.95E-08 | 4.17E-07 |
| ENSG00000162402 | USP24 | -0.24 | 0.04 | 1.02E-07 | 4.74E-07 |
| ENSG00000125841 | NRSN2 | -0.24 | 0.05 | 2.40E-07 | 1.08E-06 |
| ENSG00000112996 | MRPS30 | -0.24 | 0.05 | 3.95E-07 | 1.75E-06 |
| ENSG00000186298 | PPP1CC | -0.24 | 0.05 | 7.06E-07 | 3.05E-06 |
| ENSG00000099783 | HNRNPM | -0.24 | 0.05 | 7.98E-07 | 3.42E-06 |
| ENSG00000049860 | HEXB | -0.24 | 0.05 | 1.16E-06 | 4.88E-06 |
| ENSG00000168461 | RAB31 | -0.24 | 0.05 | 1.56E-06 | 6.52E-06 |
| ENSG00000101577 | LPIN2 | -0.24 | 0.05 | 1.70E-06 | 7.07E-06 |
| ENSG00000119777 | TMEM214 | -0.24 | 0.05 | 2.42E-06 | 9.90E-06 |
| ENSG00000111142 | METAP2 | -0.24 | 0.05 | 3.55E-06 | 1.43E-05 |
| ENSG00000075336 | TIMM21 | -0.24 | 0.05 | 6.06E-06 | 2.40E-05 |
| ENSG00000012983 | MAP4K5 | -0.24 | 0.05 | 6.13E-06 | 2.42E-05 |
| ENSG00000177565 | TBL1XR1 | -0.24 | 0.05 | 8.03E-06 | 3.14E-05 |

| | | | | | |
|---|---|---|---|---|---|
| ENSG00000149196 | HIKESHI | -0.24 | 0.06 | 1.26E-05 | 4.84E-05 |
| ENSG00000015475 | BID | -0.24 | 0.06 | 1.29E-05 | 4.92E-05 |
| ENSG00000150093 | ITGB1 | -0.24 | 0.06 | 1.32E-05 | 5.03E-05 |
| ENSG00000108395 | TRIM37 | -0.24 | 0.06 | 1.92E-05 | 7.24E-05 |
| ENSG00000116752 | BCAS2 | -0.24 | 0.06 | 2.23E-05 | 8.32E-05 |
| ENSG00000116191 | RALGPS2 | -0.24 | 0.06 | 2.73E-05 | 1.01E-04 |
| ENSG00000140395 | SKIC8 | -0.24 | 0.06 | 3.52E-05 | 1.29E-04 |
| ENSG00000182372 | CLN8 | -0.24 | 0.06 | 3.61E-05 | 1.32E-04 |
| ENSG00000174718 | RESF1 | -0.24 | 0.06 | 4.45E-05 | 1.61E-04 |
| ENSG00000163026 | WDCP | -0.24 | 0.06 | 4.52E-05 | 1.63E-04 |
| ENSG00000112640 | PPP2R5D | -0.24 | 0.06 | 4.96E-05 | 1.78E-04 |
| ENSG00000126777 | KTN1 | -0.24 | 0.06 | 8.03E-05 | 2.82E-04 |
| ENSG00000099817 | POLR2E | -0.24 | 0.06 | 9.09E-05 | 3.18E-04 |
| ENSG00000101680 | LAMA1 | -0.24 | 0.06 | 9.71E-05 | 3.38E-04 |
| ENSG00000183117 | CSMD1 | -0.24 | 0.07 | 2.10E-04 | 7.01E-04 |
| ENSG00000095380 | NANS | -0.24 | 0.07 | 2.29E-04 | 7.63E-04 |
| ENSG00000170037 | CNTROB | -0.24 | 0.07 | 2.79E-04 | 9.17E-04 |
| ENSG00000099991 | CABIN1 | -0.24 | 0.07 | 2.84E-04 | 9.34E-04 |
| ENSG00000104164 | BLOC1S6 | -0.24 | 0.07 | 2.92E-04 | 9.59E-04 |
| ENSG00000122863 | CHST3 | -0.24 | 0.07 | 3.09E-04 | 1.01E-03 |
| ENSG00000100014 | SPECC1L | -0.24 | 0.07 | 4.17E-04 | 1.34E-03 |
| ENSG00000011105 | TSPAN9 | -0.24 | 0.07 | 4.36E-04 | 1.40E-03 |
| ENSG00000129250 | KIF1C | -0.24 | 0.07 | 4.45E-04 | 1.43E-03 |
| ENSG00000172590 | MRPL52 | -0.24 | 0.07 | 4.90E-04 | 1.56E-03 |
| ENSG00000179152 | TCAIM | -0.24 | 0.07 | 4.97E-04 | 1.59E-03 |
| ENSG00000103148 | NPRL3 | -0.24 | 0.07 | 5.18E-04 | 1.65E-03 |
| ENSG00000166685 | COG1 | -0.24 | 0.07 | 5.44E-04 | 1.73E-03 |
| ENSG00000066557 | LRRC40 | -0.24 | 0.07 | 5.52E-04 | 1.75E-03 |
| ENSG00000168005 | SPINDOC | -0.24 | 0.07 | 6.25E-04 | 1.96E-03 |
| ENSG00000181744 | DIPK2A | -0.24 | 0.07 | 6.77E-04 | 2.11E-03 |
| ENSG00000152620 | NADK2 | -0.24 | 0.07 | 8.29E-04 | 2.55E-03 |
| ENSG00000213066 | CEP43 | -0.24 | 0.07 | 8.30E-04 | 2.56E-03 |
| ENSG00000135686 | KLHL36 | -0.24 | 0.07 | 8.47E-04 | 2.60E-03 |
| ENSG00000176619 | LMNB2 | -0.24 | 0.07 | 1.00E-03 | 3.04E-03 |
| ENSG00000100442 | FKBP3 | -0.24 | 0.07 | 1.02E-03 | 3.08E-03 |
| ENSG00000113761 | ZNF346 | -0.24 | 0.08 | 1.07E-03 | 3.24E-03 |
| ENSG00000138190 | EXOC6 | -0.24 | 0.08 | 1.25E-03 | 3.75E-03 |
| ENSG00000136144 | RCBTB1 | -0.24 | 0.08 | 1.30E-03 | 3.87E-03 |
| ENSG00000079393 | DUSP13 | -0.24 | 0.08 | 2.18E-03 | 6.28E-03 |
| ENSG00000178229 | ZNF543 | -0.24 | 0.08 | 3.05E-03 | 8.55E-03 |
| ENSG00000146376 | ARHGAP18 | -0.24 | 0.08 | 3.24E-03 | 9.04E-03 |
| ENSG00000139974 | SLC38A6 | -0.24 | 0.08 | 3.50E-03 | 9.70E-03 |
| ENSG00000138286 | FAM149B1 | -0.24 | 0.08 | 3.67E-03 | 1.01E-02 |
| ENSG00000164169 | PRMT9 | -0.24 | 0.09 | 4.45E-03 | 1.21E-02 |
| ENSG00000074855 | ANO8 | -0.24 | 0.09 | 4.92E-03 | 1.33E-02 |
| ENSG00000165240 | ATP7A | -0.24 | 0.09 | 5.02E-03 | 1.35E-02 |
| ENSG00000136770 | DNAJC1 | -0.24 | 0.09 | 5.69E-03 | 1.51E-02 |
| ENSG00000167646 | DNAAF3 | -0.24 | 0.1 | 8.05E-03 | 2.08E-02 |
| ENSG00000120910 | PPP3CC | -0.24 | 0.09 | 9.47E-03 | 2.40E-02 |
| ENSG00000100379 | KCTD17 | -0.24 | 0.1 | 1.00E-02 | 2.53E-02 |

| | | | | | |
|---|---|---|---|---|---|
| ENSG00000126254 | RBM42 | -0.24 | 0.1 | 1.12E-02 | 2.79E-02 |
| ENSG00000184343 | SRPK3 | -0.24 | 0.1 | 1.12E-02 | 2.81E-02 |
| ENSG00000171700 | RGS19 | -0.24 | 0.1 | 1.29E-02 | 3.17E-02 |
| ENSG00000120688 | WBP4 | -0.24 | 0.1 | 1.35E-02 | 3.31E-02 |
| ENSG00000140022 | STON2 | -0.24 | 0.1 | 1.73E-02 | 4.13E-02 |
| ENSG00000160191 | PDE9A | -0.24 | 0.11 | 1.96E-02 | 4.63E-02 |
| ENSG00000261488 | TBILA | -0.24 | 0.11 | 1.98E-02 | 4.67E-02 |
| ENSG00000186462 | NAP1L2 | -0.24 | 0.11 | 2.11E-02 | 4.95E-02 |
| ENSG00000130396 | AFDN | -0.23 | 0.03 | 5.01E-12 | 3.15E-11 |
| ENSG00000068654 | POLR1A | -0.23 | 0.04 | 2.82E-11 | 1.69E-10 |
| ENSG00000155380 | SLC16A1 | -0.23 | 0.04 | 6.42E-11 | 3.77E-10 |
| ENSG00000170445 | HARS1 | -0.23 | 0.04 | 2.96E-09 | 1.54E-08 |
| ENSG00000084112 | SSH1 | -0.23 | 0.04 | 3.81E-09 | 1.96E-08 |
| ENSG00000143442 | POGZ | -0.23 | 0.04 | 1.37E-08 | 6.80E-08 |
| ENSG00000127334 | DYRK2 | -0.23 | 0.04 | 2.77E-08 | 1.34E-07 |
| ENSG00000133028 | SCO1 | -0.23 | 0.04 | 4.38E-08 | 2.08E-07 |
| ENSG00000115484 | CCT4 | -0.23 | 0.04 | 6.88E-08 | 3.23E-07 |
| ENSG00000132359 | RAP1GAP2 | -0.23 | 0.04 | 2.05E-07 | 9.27E-07 |
| ENSG00000006607 | FARP2 | -0.23 | 0.05 | 8.51E-07 | 3.64E-06 |
| ENSG00000156697 | UTP14A | -0.23 | 0.05 | 9.38E-07 | 3.99E-06 |
| ENSG00000132275 | RRP8 | -0.23 | 0.05 | 1.55E-06 | 6.48E-06 |
| ENSG00000169554 | ZEB2 | -0.23 | 0.05 | 1.94E-06 | 8.02E-06 |
| ENSG00000090273 | NUDC | -0.23 | 0.05 | 3.64E-06 | 1.47E-05 |
| ENSG00000138018 | SELENOI | -0.23 | 0.05 | 5.52E-06 | 2.19E-05 |
| ENSG00000176087 | SLC35A4 | -0.23 | 0.05 | 5.79E-06 | 2.29E-05 |
| ENSG00000160563 | MED27 | -0.23 | 0.05 | 8.83E-06 | 3.43E-05 |
| ENSG00000168066 | SF1 | -0.23 | 0.05 | 1.13E-05 | 4.33E-05 |
| ENSG00000136731 | UGGT1 | -0.23 | 0.05 | 1.25E-05 | 4.80E-05 |
| ENSG00000163785 | RYK | -0.23 | 0.05 | 1.52E-05 | 5.77E-05 |
| ENSG00000182986 | ZNF320 | -0.23 | 0.05 | 1.58E-05 | 5.98E-05 |
| ENSG00000177426 | TGIF1 | -0.23 | 0.05 | 1.59E-05 | 6.01E-05 |
| ENSG00000138768 | USO1 | -0.23 | 0.05 | 1.96E-05 | 7.37E-05 |
| ENSG00000121057 | AKAP1 | -0.23 | 0.05 | 2.06E-05 | 7.73E-05 |
| ENSG00000118007 | STAG1 | -0.23 | 0.06 | 2.49E-05 | 9.22E-05 |
| ENSG00000116906 | GNPAT | -0.23 | 0.06 | 4.10E-05 | 1.49E-04 |
| ENSG00000064102 | INTS13 | -0.23 | 0.06 | 7.46E-05 | 2.64E-04 |
| ENSG00000143369 | ECM1 | -0.23 | 0.06 | 7.79E-05 | 2.74E-04 |
| ENSG00000157483 | MYO1E | -0.23 | 0.06 | 8.04E-05 | 2.83E-04 |
| ENSG00000039123 | MTREX | -0.23 | 0.06 | 8.37E-05 | 2.94E-04 |
| ENSG00000138646 | HERC5 | -0.23 | 0.06 | 8.91E-05 | 3.12E-04 |
| ENSG00000104880 | ARHGEF18 | -0.23 | 0.06 | 1.87E-04 | 6.30E-04 |
| ENSG00000172687 | ZNF738 | -0.23 | 0.06 | 2.53E-04 | 8.36E-04 |
| ENSG00000104756 | KCTD9 | -0.23 | 0.06 | 2.76E-04 | 9.08E-04 |
| ENSG00000123124 | WWP1 | -0.23 | 0.06 | 3.73E-04 | 1.21E-03 |
| ENSG00000100028 | SNRPD3 | -0.23 | 0.07 | 3.86E-04 | 1.25E-03 |
| ENSG00000185247 | MAGEA11 | -0.23 | 0.07 | 4.60E-04 | 1.47E-03 |
| ENSG00000133627 | ACTR3B | -0.23 | 0.07 | 4.98E-04 | 1.59E-03 |
| ENSG00000237190 | CDKN2AIPNL | -0.23 | 0.07 | 5.96E-04 | 1.88E-03 |
| ENSG00000157978 | LDLRAP1 | -0.23 | 0.07 | 8.09E-04 | 2.50E-03 |
| ENSG00000071189 | SNX13 | -0.23 | 0.07 | 9.70E-04 | 2.96E-03 |

| | | | | | |
|---|---|---|---|---|---|
| ENSG00000181896 | ZNF101 | -0.23 | 0.07 | 1.01E-03 | 3.05E-03 |
| ENSG00000197785 | ATAD3A | -0.23 | 0.07 | 1.06E-03 | 3.21E-03 |
| ENSG00000123607 | TTC21B | -0.23 | 0.07 | 1.13E-03 | 3.41E-03 |
| ENSG00000121931 | LRIF1 | -0.23 | 0.07 | 1.15E-03 | 3.47E-03 |
| ENSG00000197496 | SLC2A10 | -0.23 | 0.07 | 1.15E-03 | 3.47E-03 |
| ENSG00000167965 | MLST8 | -0.23 | 0.07 | 1.36E-03 | 4.04E-03 |
| ENSG00000157869 | RAB28 | -0.23 | 0.07 | 1.43E-03 | 4.25E-03 |
| ENSG00000118762 | PKD2 | -0.23 | 0.07 | 1.61E-03 | 4.72E-03 |
| ENSG00000135903 | PAX3 | -0.23 | 0.07 | 1.77E-03 | 5.17E-03 |
| ENSG00000125877 | ITPA | -0.23 | 0.08 | 1.98E-03 | 5.75E-03 |
| ENSG00000160117 | ANKLE1 | -0.23 | 0.08 | 1.99E-03 | 5.77E-03 |
| ENSG00000131653 | TRAF7 | -0.23 | 0.08 | 2.07E-03 | 5.98E-03 |
| ENSG00000010278 | CD9 | -0.23 | 0.08 | 2.10E-03 | 6.06E-03 |
| ENSG00000273136 | NBPF26 | -0.23 | 0.08 | 3.05E-03 | 8.56E-03 |
| ENSG00000168538 | TRAPPC11 | -0.23 | 0.08 | 3.14E-03 | 8.79E-03 |
| ENSG00000129667 | RHBDF2 | -0.23 | 0.08 | 3.47E-03 | 9.63E-03 |
| ENSG00000066827 | ZFAT | -0.23 | 0.08 | 3.65E-03 | 1.01E-02 |
| ENSG00000110693 | SOX6 | -0.23 | 0.08 | 3.74E-03 | 1.03E-02 |
| ENSG00000165959 | CLMN | -0.23 | 0.08 | 3.91E-03 | 1.08E-02 |
| ENSG00000100097 | LGALS1 | -0.23 | 0.08 | 3.93E-03 | 1.08E-02 |
| ENSG00000051382 | PIK3CB | -0.23 | 0.08 | 4.79E-03 | 1.29E-02 |
| ENSG00000166016 | ABTB2 | -0.23 | 0.09 | 5.42E-03 | 1.45E-02 |
| ENSG00000198919 | DZIP3 | -0.23 | 0.09 | 6.17E-03 | 1.63E-02 |
| ENSG00000289353 | | -0.23 | 0.09 | 6.47E-03 | 1.70E-02 |
| ENSG00000181450 | ZNF678 | -0.23 | 0.09 | 8.05E-03 | 2.08E-02 |
| ENSG00000159685 | CHCHD6 | -0.23 | 0.09 | 8.93E-03 | 2.28E-02 |
| ENSG00000108106 | UBE2S | -0.23 | 0.09 | 9.92E-03 | 2.51E-02 |
| ENSG00000149016 | TUT1 | -0.23 | 0.09 | 1.11E-02 | 2.79E-02 |
| ENSG00000187118 | CMC1 | -0.23 | 0.09 | 1.21E-02 | 3.01E-02 |
| ENSG00000171169 | NAIF1 | -0.23 | 0.1 | 1.23E-02 | 3.05E-02 |
| ENSG00000110881 | ASIC1 | -0.23 | 0.1 | 1.56E-02 | 3.76E-02 |
| ENSG00000197021 | EOLA2 | -0.23 | 0.1 | 1.59E-02 | 3.84E-02 |
| ENSG00000060642 | PIGV | -0.23 | 0.1 | 1.62E-02 | 3.89E-02 |
| ENSG00000096654 | ZNF184 | -0.23 | 0.1 | 1.70E-02 | 4.08E-02 |
| ENSG00000189164 | ZNF527 | -0.23 | 0.1 | 1.79E-02 | 4.26E-02 |
| ENSG00000100890 | PRORP | -0.23 | 0.1 | 1.82E-02 | 4.33E-02 |
| ENSG00000155868 | MED7 | -0.23 | 0.1 | 1.93E-02 | 4.57E-02 |
| ENSG00000067704 | IARS2 | -0.22 | 0.04 | 1.49E-08 | 7.39E-08 |
| ENSG00000197702 | PARVA | -0.22 | 0.04 | 2.89E-08 | 1.40E-07 |
| ENSG00000066044 | ELAVL1 | -0.22 | 0.04 | 5.90E-08 | 2.79E-07 |
| ENSG00000112242 | E2F3 | -0.22 | 0.04 | 8.63E-08 | 4.03E-07 |
| ENSG00000108592 | FTSJ3 | -0.22 | 0.04 | 4.29E-07 | 1.89E-06 |
| ENSG00000138162 | TACC2 | -0.22 | 0.04 | 7.29E-07 | 3.14E-06 |
| ENSG00000055044 | NOP58 | -0.22 | 0.05 | 7.94E-07 | 3.41E-06 |
| ENSG00000134684 | YARS1 | -0.22 | 0.05 | 1.23E-06 | 5.16E-06 |
| ENSG00000113460 | BRIX1 | -0.22 | 0.05 | 2.31E-06 | 9.48E-06 |
| ENSG00000049323 | LTBP1 | -0.22 | 0.05 | 4.97E-06 | 1.98E-05 |
| ENSG00000144724 | PTPRG | -0.22 | 0.05 | 5.22E-06 | 2.08E-05 |
| ENSG00000124766 | SOX4 | -0.22 | 0.05 | 8.15E-06 | 3.18E-05 |
| ENSG00000136718 | IMP4 | -0.22 | 0.05 | 1.03E-05 | 3.98E-05 |

| | | | | | |
|---|---|---|---|---|---|
| ENSG00000085415 | SEH1L | -0.22 | 0.05 | 1.07E-05 | 4.13E-05 |
| ENSG00000140740 | UQCRC2 | -0.22 | 0.05 | 1.08E-05 | 4.17E-05 |
| ENSG00000139437 | TCHP | -0.22 | 0.05 | 1.60E-05 | 6.05E-05 |
| ENSG00000118900 | UBN1 | -0.22 | 0.05 | 2.44E-05 | 9.06E-05 |
| ENSG00000156976 | EIF4A2 | -0.22 | 0.05 | 2.46E-05 | 9.14E-05 |
| ENSG00000180182 | MED14 | -0.22 | 0.05 | 3.10E-05 | 1.14E-04 |
| ENSG00000091732 | ZC3HC1 | -0.22 | 0.05 | 3.77E-05 | 1.37E-04 |
| ENSG00000119041 | GTF3C3 | -0.22 | 0.05 | 4.65E-05 | 1.68E-04 |
| ENSG00000133703 | KRAS | -0.22 | 0.06 | 6.97E-05 | 2.47E-04 |
| ENSG00000197728 | RPS26 | -0.22 | 0.06 | 8.94E-05 | 3.13E-04 |
| ENSG00000164338 | UTP15 | -0.22 | 0.06 | 1.05E-04 | 3.64E-04 |
| ENSG00000139218 | SCAF11 | -0.22 | 0.06 | 1.27E-04 | 4.35E-04 |
| ENSG00000135473 | PAN2 | -0.22 | 0.06 | 1.47E-04 | 5.02E-04 |
| ENSG00000107874 | CUEDC2 | -0.22 | 0.06 | 1.51E-04 | 5.14E-04 |
| ENSG00000139197 | PEX5 | -0.22 | 0.06 | 1.55E-04 | 5.26E-04 |
| ENSG00000148143 | ZNF462 | -0.22 | 0.06 | 1.61E-04 | 5.46E-04 |
| ENSG00000100280 | AP1B1 | -0.22 | 0.06 | 1.75E-04 | 5.91E-04 |
| ENSG00000168264 | IRF2BP2 | -0.22 | 0.06 | 1.98E-04 | 6.63E-04 |
| ENSG00000204070 | SYS1 | -0.22 | 0.06 | 2.52E-04 | 8.34E-04 |
| ENSG00000223482 | NUTM2A-AS1 | -0.22 | 0.06 | 2.74E-04 | 9.03E-04 |
| ENSG00000159399 | HK2 | -0.22 | 0.06 | 2.78E-04 | 9.17E-04 |
| ENSG00000185753 | CXorf38 | -0.22 | 0.06 | 2.82E-04 | 9.26E-04 |
| ENSG00000137776 | SLTM | -0.22 | 0.06 | 3.40E-04 | 1.10E-03 |
| ENSG00000215252 | GOLGA8B | -0.22 | 0.06 | 3.51E-04 | 1.14E-03 |
| ENSG00000163636 | PSMD6 | -0.22 | 0.06 | 3.88E-04 | 1.25E-03 |
| ENSG00000107897 | ACBD5 | -0.22 | 0.06 | 4.26E-04 | 1.37E-03 |
| ENSG00000178996 | SNX18 | -0.22 | 0.06 | 4.37E-04 | 1.40E-03 |
| ENSG00000092931 | MFSD11 | -0.22 | 0.06 | 5.61E-04 | 1.78E-03 |
| ENSG00000163510 | CWC22 | -0.22 | 0.07 | 6.39E-04 | 2.00E-03 |
| ENSG00000106692 | FKTN | -0.22 | 0.07 | 7.16E-04 | 2.23E-03 |
| ENSG00000186951 | PPARA | -0.22 | 0.07 | 8.06E-04 | 2.49E-03 |
| ENSG00000169155 | ZBTB43 | -0.22 | 0.07 | 8.41E-04 | 2.59E-03 |
| ENSG00000071205 | ARHGAP10 | -0.22 | 0.07 | 8.52E-04 | 2.62E-03 |
| ENSG00000170085 | SIMC1 | -0.22 | 0.07 | 8.62E-04 | 2.65E-03 |
| ENSG00000163214 | DHX57 | -0.22 | 0.07 | 8.93E-04 | 2.74E-03 |
| ENSG00000119185 | ITGB1BP1 | -0.22 | 0.07 | 9.98E-04 | 3.03E-03 |
| ENSG00000138942 | RNF185 | -0.22 | 0.07 | 1.04E-03 | 3.16E-03 |
| ENSG00000100626 | GALNT16 | -0.22 | 0.07 | 1.15E-03 | 3.46E-03 |
| ENSG00000115504 | EHBP1 | -0.22 | 0.07 | 1.15E-03 | 3.47E-03 |
| ENSG00000081320 | STK17B | -0.22 | 0.07 | 1.21E-03 | 3.64E-03 |
| ENSG00000166477 | LEO1 | -0.22 | 0.07 | 1.24E-03 | 3.71E-03 |
| ENSG00000048028 | USP28 | -0.22 | 0.07 | 1.25E-03 | 3.74E-03 |
| ENSG00000018610 | STEEP1 | -0.22 | 0.07 | 1.60E-03 | 4.71E-03 |
| ENSG00000164885 | CDK5 | -0.22 | 0.07 | 1.63E-03 | 4.78E-03 |
| ENSG00000215041 | NEURL4 | -0.22 | 0.07 | 1.89E-03 | 5.50E-03 |
| ENSG00000147862 | NFIB | -0.22 | 0.07 | 2.09E-03 | 6.02E-03 |
| ENSG00000221909 | FAM200A | -0.22 | 0.07 | 2.34E-03 | 6.72E-03 |
| ENSG00000164284 | GRPEL2 | -0.22 | 0.08 | 2.91E-03 | 8.20E-03 |
| ENSG00000243232 | PCDHAC2 | -0.22 | 0.08 | 3.54E-03 | 9.80E-03 |
| ENSG00000110811 | P3H3 | -0.22 | 0.08 | 4.62E-03 | 1.25E-02 |

| | | | | | |
|---|---|---|---|---|---|
| ENSG00000015153 | YAF2 | -0.22 | 0.08 | 4.64E-03 | 1.26E-02 |
| ENSG00000108963 | DPH1 | -0.22 | 0.08 | 4.72E-03 | 1.28E-02 |
| ENSG00000117475 | BLZF1 | -0.22 | 0.08 | 5.49E-03 | 1.47E-02 |
| ENSG00000139178 | C1RL | -0.22 | 0.08 | 5.59E-03 | 1.49E-02 |
| ENSG00000163975 | MELTF | -0.22 | 0.08 | 5.65E-03 | 1.50E-02 |
| ENSG00000111845 | PAK1IP1 | -0.22 | 0.08 | 6.50E-03 | 1.71E-02 |
| ENSG00000160392 | C19orf47 | -0.22 | 0.09 | 1.15E-02 | 2.86E-02 |
| ENSG00000167779 | IGFBP6 | -0.22 | 0.09 | 1.31E-02 | 3.23E-02 |
| ENSG00000241127 | YAE1 | -0.22 | 0.09 | 1.59E-02 | 3.83E-02 |
| ENSG00000159403 | C1R | -0.22 | 0.09 | 1.88E-02 | 4.46E-02 |
| ENSG00000146834 | MEPCE | -0.22 | 0.1 | 2.11E-02 | 4.95E-02 |
| ENSG00000129187 | DCTD | -0.21 | 0.04 | 2.49E-08 | 1.21E-07 |
| ENSG00000104331 | BPNT2 | -0.21 | 0.04 | 2.19E-07 | 9.91E-07 |
| ENSG00000132002 | DNAJB1 | -0.21 | 0.04 | 8.52E-07 | 3.64E-06 |
| ENSG00000197892 | KIF13B | -0.21 | 0.04 | 1.69E-06 | 7.02E-06 |
| ENSG00000131043 | AAR2 | -0.21 | 0.04 | 1.70E-06 | 7.07E-06 |
| ENSG00000122786 | CALD1 | -0.21 | 0.04 | 1.98E-06 | 8.16E-06 |
| ENSG00000085721 | RRN3 | -0.21 | 0.05 | 2.77E-06 | 1.13E-05 |
| ENSG00000085982 | USP40 | -0.21 | 0.05 | 3.84E-06 | 1.55E-05 |
| ENSG00000101544 | ADNP2 | -0.21 | 0.05 | 4.92E-06 | 1.96E-05 |
| ENSG00000146457 | WTAP | -0.21 | 0.05 | 1.22E-05 | 4.69E-05 |
| ENSG00000163597 | SNHG16 | -0.21 | 0.05 | 1.84E-05 | 6.93E-05 |
| ENSG00000151929 | BAG3 | -0.21 | 0.05 | 3.25E-05 | 1.19E-04 |
| ENSG00000109452 | INPP4B | -0.21 | 0.05 | 3.52E-05 | 1.29E-04 |
| ENSG00000109113 | RAB34 | -0.21 | 0.05 | 3.58E-05 | 1.31E-04 |
| ENSG00000104626 | ERI1 | -0.21 | 0.05 | 6.94E-05 | 2.46E-04 |
| ENSG00000147679 | UTP23 | -0.21 | 0.05 | 8.82E-05 | 3.09E-04 |
| ENSG00000079156 | OSBPL6 | -0.21 | 0.05 | 1.12E-04 | 3.87E-04 |
| ENSG00000100647 | SUSD6 | -0.21 | 0.06 | 1.13E-04 | 3.91E-04 |
| ENSG00000129255 | MPDU1 | -0.21 | 0.06 | 1.18E-04 | 4.08E-04 |
| ENSG00000156253 | RWDD2B | -0.21 | 0.06 | 1.52E-04 | 5.17E-04 |
| ENSG00000068024 | HDAC4 | -0.21 | 0.06 | 1.54E-04 | 5.22E-04 |
| ENSG00000119446 | RBM18 | -0.21 | 0.06 | 1.74E-04 | 5.86E-04 |
| ENSG00000019485 | PRDM11 | -0.21 | 0.06 | 1.77E-04 | 5.97E-04 |
| ENSG00000134874 | DZIP1 | -0.21 | 0.06 | 4.10E-04 | 1.32E-03 |
| ENSG00000211445 | GPX3 | -0.21 | 0.06 | 4.86E-04 | 1.55E-03 |
| ENSG00000204560 | DHX16 | -0.21 | 0.06 | 5.18E-04 | 1.65E-03 |
| ENSG00000171574 | ZNF584 | -0.21 | 0.06 | 5.78E-04 | 1.83E-03 |
| ENSG00000100994 | PYGB | -0.21 | 0.06 | 6.24E-04 | 1.96E-03 |
| ENSG00000112339 | HBS1L | -0.21 | 0.06 | 7.09E-04 | 2.21E-03 |
| ENSG00000074582 | BCS1L | -0.21 | 0.06 | 7.43E-04 | 2.30E-03 |
| ENSG00000035141 | FAM136A | -0.21 | 0.06 | 7.94E-04 | 2.46E-03 |
| ENSG00000122299 | ZC3H7A | -0.21 | 0.06 | 8.24E-04 | 2.54E-03 |
| ENSG00000144647 | POMGNT2 | -0.21 | 0.06 | 8.46E-04 | 2.60E-03 |
| ENSG00000113456 | RAD1 | -0.21 | 0.06 | 8.55E-04 | 2.63E-03 |
| ENSG00000188986 | NELFB | -0.21 | 0.06 | 9.18E-04 | 2.81E-03 |
| ENSG00000151474 | FRMD4A | -0.21 | 0.07 | 1.08E-03 | 3.27E-03 |
| ENSG00000147044 | CASK | -0.21 | 0.07 | 1.37E-03 | 4.08E-03 |
| ENSG00000178927 | CYBC1 | -0.21 | 0.07 | 1.64E-03 | 4.82E-03 |
| ENSG00000159210 | SNF8 | -0.21 | 0.07 | 1.95E-03 | 5.67E-03 |

| | | | | | |
|---|---|---|---|---|---|
| ENSG00000160439 | RDH13 | -0.21 | 0.07 | 1.97E-03 | 5.72E-03 |
| ENSG00000184916 | JAG2 | -0.21 | 0.07 | 3.11E-03 | 8.71E-03 |
| ENSG00000141068 | KSR1 | -0.21 | 0.07 | 3.26E-03 | 9.08E-03 |
| ENSG00000171503 | ETFDH | -0.21 | 0.07 | 3.27E-03 | 9.12E-03 |
| ENSG00000162627 | SNX7 | -0.21 | 0.07 | 3.81E-03 | 1.05E-02 |
| ENSG00000122008 | POLK | -0.21 | 0.08 | 5.75E-03 | 1.53E-02 |
| ENSG00000110429 | FBXO3 | -0.21 | 0.08 | 6.03E-03 | 1.60E-02 |
| ENSG00000138172 | CALHM2 | -0.21 | 0.08 | 6.32E-03 | 1.67E-02 |
| ENSG00000213096 | ZNF254 | -0.21 | 0.08 | 7.69E-03 | 2.00E-02 |
| ENSG00000175548 | ALG10B | -0.21 | 0.08 | 8.08E-03 | 2.08E-02 |
| ENSG00000130204 | TOMM40 | -0.21 | 0.08 | 8.22E-03 | 2.12E-02 |
| ENSG00000178209 | PLEC | -0.21 | 0.08 | 8.93E-03 | 2.28E-02 |
| ENSG00000007080 | CCDC124 | -0.21 | 0.08 | 1.00E-02 | 2.53E-02 |
| ENSG00000260027 | HOXB7 | -0.21 | 0.09 | 1.67E-02 | 4.01E-02 |
| ENSG00000133794 | BMAL1 | -0.21 | 0.09 | 1.93E-02 | 4.57E-02 |
| ENSG00000115677 | HDLBP | -0.2 | 0.04 | 6.20E-08 | 2.92E-07 |
| ENSG00000175792 | RUVBL1 | -0.2 | 0.05 | 1.26E-05 | 4.81E-05 |
| ENSG00000055130 | CUL1 | -0.2 | 0.05 | 7.10E-05 | 2.51E-04 |
| ENSG00000018408 | WWTR1 | -0.2 | 0.05 | 7.40E-05 | 2.61E-04 |
| ENSG00000169814 | BTD | -0.2 | 0.05 | 1.24E-04 | 4.25E-04 |
| ENSG00000158545 | ZC3H18 | -0.2 | 0.05 | 1.24E-04 | 4.26E-04 |
| ENSG00000158435 | CNOT11 | -0.2 | 0.05 | 1.29E-04 | 4.43E-04 |
| ENSG00000103449 | SALL1 | -0.2 | 0.06 | 2.72E-04 | 8.98E-04 |
| ENSG00000115234 | SNX17 | -0.2 | 0.05 | 2.73E-04 | 8.99E-04 |
| ENSG00000105879 | CBLL1 | -0.2 | 0.06 | 2.80E-04 | 9.22E-04 |
| ENSG00000165271 | NOL6 | -0.2 | 0.06 | 3.06E-04 | 1.00E-03 |
| ENSG00000010322 | NISCH | -0.2 | 0.06 | 4.59E-04 | 1.47E-03 |
| ENSG00000185551 | NR2F2 | -0.2 | 0.06 | 4.86E-04 | 1.55E-03 |
| ENSG00000131981 | LGALS3 | -0.2 | 0.06 | 5.51E-04 | 1.75E-03 |
| ENSG00000143499 | SMYD2 | -0.2 | 0.06 | 8.53E-04 | 2.62E-03 |
| ENSG00000106976 | DNM1 | -0.2 | 0.06 | 9.32E-04 | 2.84E-03 |
| ENSG00000145675 | PIK3R1 | -0.2 | 0.06 | 9.42E-04 | 2.88E-03 |
| ENSG00000177485 | ZBTB33 | -0.2 | 0.06 | 9.97E-04 | 3.03E-03 |
| ENSG00000035928 | RFC1 | -0.2 | 0.06 | 1.04E-03 | 3.14E-03 |
| ENSG00000130363 | RSPH3 | -0.2 | 0.06 | 1.19E-03 | 3.56E-03 |
| ENSG00000067334 | DNTTIP2 | -0.2 | 0.06 | 1.19E-03 | 3.57E-03 |
| ENSG00000144366 | GULP1 | -0.2 | 0.06 | 1.25E-03 | 3.74E-03 |
| ENSG00000120833 | SOCS2 | -0.2 | 0.06 | 1.27E-03 | 3.80E-03 |
| ENSG00000148498 | PARD3 | -0.2 | 0.06 | 1.48E-03 | 4.39E-03 |
| ENSG00000139531 | SUOX | -0.2 | 0.06 | 1.59E-03 | 4.68E-03 |
| ENSG00000186812 | ZNF397 | -0.2 | 0.07 | 1.77E-03 | 5.17E-03 |
| ENSG00000170421 | KRT8 | -0.2 | 0.06 | 1.77E-03 | 5.18E-03 |
| ENSG00000166788 | SAAL1 | -0.2 | 0.07 | 2.07E-03 | 5.98E-03 |
| ENSG00000101187 | SLCO4A1 | -0.2 | 0.07 | 2.35E-03 | 6.73E-03 |
| ENSG00000177981 | ASB8 | -0.2 | 0.07 | 2.41E-03 | 6.88E-03 |
| ENSG00000262879 | | -0.2 | 0.07 | 2.56E-03 | 7.29E-03 |
| ENSG00000198836 | OPA1 | -0.2 | 0.07 | 2.69E-03 | 7.62E-03 |
| ENSG00000160789 | LMNA | -0.2 | 0.07 | 3.07E-03 | 8.61E-03 |
| ENSG00000101181 | MTG2 | -0.2 | 0.07 | 3.71E-03 | 1.02E-02 |
| ENSG00000225630 | MTND2P28 | -0.2 | 0.07 | 3.78E-03 | 1.04E-02 |

| | | | | | |
|---|---|---|---|---|---|
| ENSG00000134077 | THUMPD3 | -0.2 | 0.07 | 4.20E-03 | 1.15E-02 |
| ENSG00000052795 | FNIP2 | -0.2 | 0.07 | 4.96E-03 | 1.33E-02 |
| ENSG00000168806 | LCMT2 | -0.2 | 0.08 | 6.14E-03 | 1.62E-02 |
| ENSG00000150990 | DHX37 | -0.2 | 0.08 | 6.20E-03 | 1.64E-02 |
| ENSG00000196150 | ZNF250 | -0.2 | 0.07 | 6.41E-03 | 1.69E-02 |
| ENSG00000157426 | AASDH | -0.2 | 0.07 | 6.43E-03 | 1.69E-02 |
| ENSG00000166439 | RNF169 | -0.2 | 0.07 | 7.31E-03 | 1.90E-02 |
| ENSG00000110013 | SIAE | -0.2 | 0.08 | 8.72E-03 | 2.23E-02 |
| ENSG00000101773 | RBBP8 | -0.2 | 0.08 | 8.72E-03 | 2.24E-02 |
| ENSG00000120158 | RCL1 | -0.2 | 0.08 | 1.00E-02 | 2.53E-02 |
| ENSG00000163006 | CCDC138 | -0.2 | 0.08 | 1.01E-02 | 2.55E-02 |
| ENSG00000105135 | ILVBL | -0.2 | 0.08 | 1.16E-02 | 2.89E-02 |
| ENSG00000103043 | VAC14 | -0.2 | 0.08 | 1.36E-02 | 3.33E-02 |
| ENSG00000167625 | ZNF526 | -0.2 | 0.08 | 1.40E-02 | 3.42E-02 |
| ENSG00000219545 | UMAD1 | -0.2 | 0.09 | 1.76E-02 | 4.19E-02 |
| ENSG00000184635 | ZNF93 | -0.2 | 0.09 | 1.92E-02 | 4.55E-02 |
| ENSG00000198105 | ZNF248 | -0.2 | 0.09 | 1.94E-02 | 4.59E-02 |
| ENSG00000127463 | EMC1 | -0.19 | 0.03 | 1.91E-08 | 9.36E-08 |
| ENSG00000132341 | RAN | -0.19 | 0.04 | 1.03E-07 | 4.78E-07 |
| ENSG00000100353 | EIF3D | -0.19 | 0.04 | 2.30E-07 | 1.04E-06 |
| ENSG00000165689 | ENTR1 | -0.19 | 0.04 | 3.33E-06 | 1.35E-05 |
| ENSG00000196262 | PPIA | -0.19 | 0.04 | 4.01E-06 | 1.61E-05 |
| ENSG00000114982 | KANSL3 | -0.19 | 0.04 | 5.45E-06 | 2.17E-05 |
| ENSG00000111358 | GTF2H3 | -0.19 | 0.04 | 8.19E-06 | 3.19E-05 |
| ENSG00000102054 | RBBP7 | -0.19 | 0.04 | 8.23E-06 | 3.21E-05 |
| ENSG00000117505 | DR1 | -0.19 | 0.04 | 8.42E-06 | 3.28E-05 |
| ENSG00000115183 | TANC1 | -0.19 | 0.04 | 8.42E-06 | 3.28E-05 |
| ENSG00000104824 | HNRNPL | -0.19 | 0.05 | 1.66E-05 | 6.28E-05 |
| ENSG00000136045 | PWP1 | -0.19 | 0.05 | 2.95E-05 | 1.09E-04 |
| ENSG00000117748 | RPA2 | -0.19 | 0.05 | 3.90E-05 | 1.42E-04 |
| ENSG00000189060 | H1-0 | -0.19 | 0.05 | 4.74E-05 | 1.71E-04 |
| ENSG00000072062 | PRKACA | -0.19 | 0.05 | 4.91E-05 | 1.76E-04 |
| ENSG00000265491 | RNF115 | -0.19 | 0.05 | 5.41E-05 | 1.94E-04 |
| ENSG00000182606 | TRAK1 | -0.19 | 0.05 | 7.67E-05 | 2.71E-04 |
| ENSG00000146909 | NOM1 | -0.19 | 0.05 | 8.63E-05 | 3.02E-04 |
| ENSG00000139921 | TMX1 | -0.19 | 0.05 | 1.00E-04 | 3.48E-04 |
| ENSG00000133961 | NUMB | -0.19 | 0.05 | 1.47E-04 | 5.02E-04 |
| ENSG00000110274 | CEP164 | -0.19 | 0.05 | 1.58E-04 | 5.36E-04 |
| ENSG00000103855 | CD276 | -0.19 | 0.05 | 1.64E-04 | 5.56E-04 |
| ENSG00000127955 | GNAI1 | -0.19 | 0.05 | 2.26E-04 | 7.53E-04 |
| ENSG00000133706 | LARS1 | -0.19 | 0.05 | 2.79E-04 | 9.17E-04 |
| ENSG00000151247 | EIF4E | -0.19 | 0.05 | 2.81E-04 | 9.24E-04 |
| ENSG00000165526 | RPUSD4 | -0.19 | 0.05 | 4.13E-04 | 1.33E-03 |
| ENSG00000044524 | EPHA3 | -0.19 | 0.05 | 4.27E-04 | 1.37E-03 |
| ENSG00000111802 | TDP2 | -0.19 | 0.05 | 4.41E-04 | 1.41E-03 |
| ENSG00000164924 | YWHAZ | -0.19 | 0.05 | 4.45E-04 | 1.43E-03 |
| ENSG00000108094 | CUL2 | -0.19 | 0.05 | 4.72E-04 | 1.51E-03 |
| ENSG00000099810 | MTAP | -0.19 | 0.05 | 4.76E-04 | 1.52E-03 |
| ENSG00000089057 | SLC23A2 | -0.19 | 0.06 | 8.72E-04 | 2.67E-03 |
| ENSG00000175334 | BANF1 | -0.19 | 0.06 | 8.95E-04 | 2.74E-03 |

| | | | | | |
|---|---|---|---|---|---|
| ENSG00000117298 | ECE1 | -0.19 | 0.06 | 9.39E-04 | 2.87E-03 |
| ENSG00000122958 | VPS26A | -0.19 | 0.06 | 9.69E-04 | 2.95E-03 |
| ENSG00000136891 | TEX10 | -0.19 | 0.06 | 1.04E-03 | 3.14E-03 |
| ENSG00000060762 | MPC1 | -0.19 | 0.06 | 1.38E-03 | 4.10E-03 |
| ENSG00000197586 | ENTPD6 | -0.19 | 0.06 | 1.47E-03 | 4.36E-03 |
| ENSG00000144591 | GMPPA | -0.19 | 0.06 | 1.64E-03 | 4.81E-03 |
| ENSG00000157240 | FZD1 | -0.19 | 0.06 | 1.66E-03 | 4.87E-03 |
| ENSG00000162378 | ZYG11B | -0.19 | 0.06 | 1.89E-03 | 5.50E-03 |
| ENSG00000128626 | MRPS12 | -0.19 | 0.06 | 2.18E-03 | 6.26E-03 |
| ENSG00000158793 | NIT1 | -0.19 | 0.07 | 2.72E-03 | 7.69E-03 |
| ENSG00000156675 | RAB11FIP1 | -0.19 | 0.06 | 2.87E-03 | 8.08E-03 |
| ENSG00000186340 | THBS2 | -0.19 | 0.06 | 2.94E-03 | 8.28E-03 |
| ENSG00000136848 | DAB2IP | -0.19 | 0.07 | 3.35E-03 | 9.33E-03 |
| ENSG00000153975 | ZUP1 | -0.19 | 0.07 | 3.65E-03 | 1.01E-02 |
| ENSG00000079974 | RABL2B | -0.19 | 0.07 | 4.09E-03 | 1.12E-02 |
| ENSG00000177150 | FAM210A | -0.19 | 0.07 | 4.18E-03 | 1.14E-02 |
| ENSG00000161981 | SNRNP25 | -0.19 | 0.07 | 4.25E-03 | 1.16E-02 |
| ENSG00000171204 | TMEM126B | -0.19 | 0.07 | 4.31E-03 | 1.17E-02 |
| ENSG00000160214 | RRP1 | -0.19 | 0.07 | 4.92E-03 | 1.32E-02 |
| ENSG00000196781 | TLE1 | -0.19 | 0.07 | 6.52E-03 | 1.71E-02 |
| ENSG00000162086 | ZNF75A | -0.19 | 0.07 | 6.77E-03 | 1.78E-02 |
| ENSG00000188352 | FOCAD | -0.19 | 0.07 | 6.86E-03 | 1.80E-02 |
| ENSG00000100575 | TIMM9 | -0.19 | 0.07 | 7.44E-03 | 1.94E-02 |
| ENSG00000166341 | DCHS1 | -0.19 | 0.08 | 8.43E-03 | 2.17E-02 |
| ENSG00000155636 | RBM45 | -0.19 | 0.08 | 9.97E-03 | 2.52E-02 |
| ENSG00000158169 | FANCC | -0.19 | 0.08 | 1.04E-02 | 2.62E-02 |
| ENSG00000173889 | PHC3 | -0.19 | 0.07 | 1.16E-02 | 2.90E-02 |
| ENSG00000126391 | FRMD8 | -0.19 | 0.08 | 1.21E-02 | 3.00E-02 |
| ENSG00000090612 | ZNF268 | -0.19 | 0.08 | 1.26E-02 | 3.10E-02 |
| ENSG00000286388 | | -0.19 | 0.08 | 1.28E-02 | 3.16E-02 |
| ENSG00000246273 | SBF2-AS1 | -0.19 | 0.08 | 1.49E-02 | 3.61E-02 |
| ENSG00000274070 | CASTOR2 | -0.19 | 0.08 | 1.68E-02 | 4.03E-02 |
| ENSG00000150764 | DIXDC1 | -0.19 | 0.08 | 1.75E-02 | 4.17E-02 |
| ENSG00000124380 | SNRNP27 | -0.19 | 0.08 | 1.85E-02 | 4.39E-02 |
| ENSG00000131759 | RARA | -0.19 | 0.08 | 1.96E-02 | 4.62E-02 |
| ENSG00000049167 | ERCC8 | -0.19 | 0.08 | 1.98E-02 | 4.67E-02 |
| ENSG00000270816 | LINC00221 | -0.19 | 0.08 | 2.03E-02 | 4.78E-02 |
| ENSG00000134308 | YWHAQ | -0.18 | 0.04 | 4.91E-07 | 2.15E-06 |
| ENSG00000173706 | HEG1 | -0.18 | 0.04 | 1.85E-06 | 7.64E-06 |
| ENSG00000138814 | PPP3CA | -0.18 | 0.04 | 7.49E-06 | 2.93E-05 |
| ENSG00000266412 | NCOA4 | -0.18 | 0.05 | 6.08E-05 | 2.17E-04 |
| ENSG00000118495 | PLAGL1 | -0.18 | 0.05 | 7.74E-05 | 2.73E-04 |
| ENSG00000156642 | NPTN | -0.18 | 0.05 | 8.11E-05 | 2.85E-04 |
| ENSG00000136381 | IREB2 | -0.18 | 0.05 | 1.08E-04 | 3.75E-04 |
| ENSG00000061987 | MON2 | -0.18 | 0.05 | 1.29E-04 | 4.43E-04 |
| ENSG00000028310 | BRD9 | -0.18 | 0.05 | 1.84E-04 | 6.20E-04 |
| ENSG00000124181 | PLCG1 | -0.18 | 0.05 | 1.88E-04 | 6.33E-04 |
| ENSG00000240184 | PCDHGC3 | -0.18 | 0.05 | 2.29E-04 | 7.62E-04 |
| ENSG00000100109 | TFIP11 | -0.18 | 0.05 | 2.70E-04 | 8.92E-04 |
| ENSG00000170881 | RNF139 | -0.18 | 0.05 | 3.62E-04 | 1.17E-03 |

| | | | | | |
|---|---|---|---|---|---|
| ENSG00000163399 | ATP1A1 | -0.18 | 0.05 | 4.15E-04 | 1.33E-03 |
| ENSG00000108946 | PRKAR1A | -0.18 | 0.05 | 4.85E-04 | 1.55E-03 |
| ENSG00000086827 | ZW10 | -0.18 | 0.05 | 4.87E-04 | 1.55E-03 |
| ENSG00000131374 | TBC1D5 | -0.18 | 0.05 | 5.05E-04 | 1.61E-03 |
| ENSG00000130695 | CEP85 | -0.18 | 0.05 | 5.33E-04 | 1.69E-03 |
| ENSG00000255690 | TRIL | -0.18 | 0.05 | 5.44E-04 | 1.73E-03 |
| ENSG00000072110 | ACTN1 | -0.18 | 0.05 | 6.52E-04 | 2.04E-03 |
| ENSG00000089876 | DHX32 | -0.18 | 0.05 | 6.63E-04 | 2.07E-03 |
| ENSG00000143374 | TARS2 | -0.18 | 0.06 | 8.07E-04 | 2.49E-03 |
| ENSG00000186416 | NKRF | -0.18 | 0.06 | 8.74E-04 | 2.68E-03 |
| ENSG00000135966 | TGFBRAP1 | -0.18 | 0.06 | 9.20E-04 | 2.81E-03 |
| ENSG00000148688 | RPP30 | -0.18 | 0.05 | 1.02E-03 | 3.10E-03 |
| ENSG00000183605 | SFXN4 | -0.18 | 0.06 | 1.03E-03 | 3.11E-03 |
| ENSG00000110880 | CORO1C | -0.18 | 0.05 | 1.21E-03 | 3.62E-03 |
| ENSG00000139620 | KANSL2 | -0.18 | 0.06 | 1.23E-03 | 3.69E-03 |
| ENSG00000101752 | MIB1 | -0.18 | 0.06 | 1.28E-03 | 3.81E-03 |
| ENSG00000198040 | ZNF84 | -0.18 | 0.06 | 1.31E-03 | 3.90E-03 |
| ENSG00000142227 | EMP3 | -0.18 | 0.06 | 1.31E-03 | 3.92E-03 |
| ENSG00000073417 | PDE8A | -0.18 | 0.06 | 1.51E-03 | 4.47E-03 |
| ENSG00000274523 | RCC1L | -0.18 | 0.06 | 1.51E-03 | 4.47E-03 |
| ENSG00000100395 | L3MBTL2 | -0.18 | 0.06 | 1.55E-03 | 4.55E-03 |
| ENSG00000120963 | ZNF706 | -0.18 | 0.06 | 1.88E-03 | 5.47E-03 |
| ENSG00000178028 | DMAP1 | -0.18 | 0.06 | 2.38E-03 | 6.82E-03 |
| ENSG00000175832 | ETV4 | -0.18 | 0.06 | 2.72E-03 | 7.70E-03 |
| ENSG00000114098 | ARMC8 | -0.18 | 0.06 | 2.74E-03 | 7.76E-03 |
| ENSG00000138231 | DBR1 | -0.18 | 0.06 | 3.00E-03 | 8.44E-03 |
| ENSG00000143183 | TMCO1 | -0.18 | 0.06 | 3.23E-03 | 9.01E-03 |
| ENSG00000188976 | NOC2L | -0.18 | 0.06 | 3.26E-03 | 9.08E-03 |
| ENSG00000048471 | SNX29 | -0.18 | 0.06 | 3.48E-03 | 9.64E-03 |
| ENSG00000115840 | SLC25A12 | -0.18 | 0.06 | 3.56E-03 | 9.86E-03 |
| ENSG00000104320 | NBN | -0.18 | 0.06 | 3.76E-03 | 1.04E-02 |
| ENSG00000108515 | ENO3 | -0.18 | 0.06 | 3.93E-03 | 1.08E-02 |
| ENSG00000136710 | CCDC115 | -0.18 | 0.06 | 4.03E-03 | 1.10E-02 |
| ENSG00000127603 | MACF1 | -0.18 | 0.06 | 4.15E-03 | 1.13E-02 |
| ENSG00000163933 | RFT1 | -0.18 | 0.07 | 4.26E-03 | 1.16E-02 |
| ENSG00000156931 | VPS8 | -0.18 | 0.06 | 4.70E-03 | 1.27E-02 |
| ENSG00000126581 | BECN1 | -0.18 | 0.07 | 4.82E-03 | 1.30E-02 |
| ENSG00000143811 | PYCR2 | -0.18 | 0.06 | 5.31E-03 | 1.42E-02 |
| ENSG00000082068 | WDR70 | -0.18 | 0.07 | 5.35E-03 | 1.43E-02 |
| ENSG00000119778 | ATAD2B | -0.18 | 0.07 | 6.03E-03 | 1.60E-02 |
| ENSG00000100372 | SLC25A17 | -0.18 | 0.07 | 6.35E-03 | 1.67E-02 |
| ENSG00000118620 | ZNF430 | -0.18 | 0.07 | 7.68E-03 | 1.99E-02 |
| ENSG00000180964 | TCEAL8 | -0.18 | 0.07 | 9.35E-03 | 2.38E-02 |
| ENSG00000136682 | ZNG1B | -0.18 | 0.07 | 1.06E-02 | 2.67E-02 |
| ENSG00000168286 | THAP11 | -0.18 | 0.07 | 1.24E-02 | 3.08E-02 |
| ENSG00000112210 | RAB23 | -0.18 | 0.07 | 1.32E-02 | 3.24E-02 |
| ENSG00000103653 | CSK | -0.18 | 0.07 | 1.35E-02 | 3.31E-02 |
| ENSG00000132661 | NXT1 | -0.18 | 0.07 | 1.40E-02 | 3.41E-02 |
| ENSG00000091947 | TMEM101 | -0.18 | 0.08 | 1.41E-02 | 3.43E-02 |
| ENSG00000156603 | MED19 | -0.18 | 0.07 | 1.41E-02 | 3.44E-02 |

| | | | | | |
|---|---|---|---|---|---|
| ENSG00000005801 | ZNF195 | -0.18 | 0.07 | 1.49E-02 | 3.61E-02 |
| ENSG00000132952 | USPL1 | -0.18 | 0.08 | 1.59E-02 | 3.82E-02 |
| ENSG00000135828 | RNASEL | -0.18 | 0.08 | 1.67E-02 | 4.01E-02 |
| ENSG00000145780 | FEM1C | -0.18 | 0.08 | 1.77E-02 | 4.21E-02 |
| ENSG00000137656 | BUD13 | -0.18 | 0.08 | 1.77E-02 | 4.21E-02 |
| ENSG00000108509 | CAMTA2 | -0.18 | 0.08 | 1.82E-02 | 4.33E-02 |
| ENSG00000188677 | PARVB | -0.18 | 0.08 | 1.98E-02 | 4.68E-02 |
| ENSG00000067992 | PDK3 | -0.18 | 0.08 | 2.07E-02 | 4.86E-02 |
| ENSG00000162783 | IER5 | -0.18 | 0.08 | 2.10E-02 | 4.92E-02 |
| ENSG00000116560 | SFPQ | -0.17 | 0.03 | 4.96E-07 | 2.17E-06 |
| ENSG00000133026 | MYH10 | -0.17 | 0.04 | 7.51E-07 | 3.23E-06 |
| ENSG00000170275 | CRTAP | -0.17 | 0.04 | 4.45E-06 | 1.78E-05 |
| ENSG00000177889 | UBE2N | -0.17 | 0.04 | 5.51E-06 | 2.19E-05 |
| ENSG00000110344 | UBE4A | -0.17 | 0.04 | 8.54E-06 | 3.32E-05 |
| ENSG00000163527 | STT3B | -0.17 | 0.04 | 1.49E-05 | 5.65E-05 |
| ENSG00000089737 | DDX24 | -0.17 | 0.04 | 2.14E-05 | 8.02E-05 |
| ENSG00000224078 | SNHG14 | -0.17 | 0.04 | 2.55E-05 | 9.45E-05 |
| ENSG00000198947 | DMD | -0.17 | 0.04 | 2.89E-05 | 1.06E-04 |
| ENSG00000100393 | EP300 | -0.17 | 0.04 | 4.14E-05 | 1.50E-04 |
| ENSG00000163812 | ZDHHC3 | -0.17 | 0.04 | 5.64E-05 | 2.02E-04 |
| ENSG00000176624 | MEX3C | -0.17 | 0.04 | 1.17E-04 | 4.03E-04 |
| ENSG00000073008 | PVR | -0.17 | 0.04 | 1.22E-04 | 4.20E-04 |
| ENSG00000263001 | GTF2I | -0.17 | 0.05 | 2.64E-04 | 8.72E-04 |
| ENSG00000172888 | ZNF621 | -0.17 | 0.05 | 2.89E-04 | 9.49E-04 |
| ENSG00000157259 | GATAD1 | -0.17 | 0.05 | 2.93E-04 | 9.60E-04 |
| ENSG00000185414 | MRPL30 | -0.17 | 0.05 | 5.86E-04 | 1.85E-03 |
| ENSG00000100347 | SAMM50 | -0.17 | 0.05 | 6.24E-04 | 1.96E-03 |
| ENSG00000143367 | TUFT1 | -0.17 | 0.05 | 6.95E-04 | 2.17E-03 |
| ENSG00000104408 | EIF3E | -0.17 | 0.05 | 9.43E-04 | 2.88E-03 |
| ENSG00000139131 | YARS2 | -0.17 | 0.05 | 1.02E-03 | 3.08E-03 |
| ENSG00000128641 | MYO1B | -0.17 | 0.05 | 1.03E-03 | 3.11E-03 |
| ENSG00000162408 | NOL9 | -0.17 | 0.05 | 1.24E-03 | 3.70E-03 |
| ENSG00000196313 | POM121 | -0.17 | 0.05 | 1.28E-03 | 3.81E-03 |
| ENSG00000198162 | MAN1A2 | -0.17 | 0.05 | 1.37E-03 | 4.08E-03 |
| ENSG00000171720 | HDAC3 | -0.17 | 0.05 | 1.57E-03 | 4.63E-03 |
| ENSG00000124574 | ABCC10 | -0.17 | 0.05 | 1.62E-03 | 4.76E-03 |
| ENSG00000115685 | PPP1R7 | -0.17 | 0.05 | 1.66E-03 | 4.87E-03 |
| ENSG00000047932 | GOPC | -0.17 | 0.05 | 1.90E-03 | 5.54E-03 |
| ENSG00000154222 | CC2D1B | -0.17 | 0.06 | 2.01E-03 | 5.81E-03 |
| ENSG00000177200 | CHD9 | -0.17 | 0.06 | 2.10E-03 | 6.04E-03 |
| ENSG00000112208 | BAG2 | -0.17 | 0.06 | 2.11E-03 | 6.08E-03 |
| ENSG00000204580 | DDR1 | -0.17 | 0.06 | 2.51E-03 | 7.16E-03 |
| ENSG00000135404 | CD63 | -0.17 | 0.06 | 2.63E-03 | 7.46E-03 |
| ENSG00000117461 | PIK3R3 | -0.17 | 0.06 | 2.78E-03 | 7.84E-03 |
| ENSG00000124570 | SERPINB6 | -0.17 | 0.06 | 2.90E-03 | 8.16E-03 |
| ENSG00000126457 | PRMT1 | -0.17 | 0.06 | 3.08E-03 | 8.63E-03 |
| ENSG00000173786 | CNP | -0.17 | 0.06 | 3.16E-03 | 8.83E-03 |
| ENSG00000138303 | ASCC1 | -0.17 | 0.06 | 3.21E-03 | 8.97E-03 |
| ENSG00000141698 | NT5C3B | -0.17 | 0.06 | 3.22E-03 | 9.00E-03 |
| ENSG00000157110 | RBPMS | -0.17 | 0.06 | 3.33E-03 | 9.28E-03 |

| | | | | | |
|---|---|---|---|---|---|
| ENSG00000080854 | IGSF9B | -0.17 | 0.06 | 3.34E-03 | 9.30E-03 |
| ENSG00000130402 | ACTN4 | -0.17 | 0.06 | 4.07E-03 | 1.11E-02 |
| ENSG00000154719 | MRPL39 | -0.17 | 0.06 | 4.11E-03 | 1.12E-02 |
| ENSG00000126698 | DNAJC8 | -0.17 | 0.06 | 4.20E-03 | 1.15E-02 |
| ENSG00000137713 | PPP2R1B | -0.17 | 0.06 | 4.53E-03 | 1.23E-02 |
| ENSG00000135775 | COG2 | -0.17 | 0.06 | 5.08E-03 | 1.37E-02 |
| ENSG00000131848 | ZSCAN5A | -0.17 | 0.06 | 5.20E-03 | 1.39E-02 |
| ENSG00000134255 | CEPT1 | -0.17 | 0.06 | 5.24E-03 | 1.40E-02 |
| ENSG00000106628 | POLD2 | -0.17 | 0.06 | 5.64E-03 | 1.50E-02 |
| ENSG00000157625 | TAB3 | -0.17 | 0.06 | 5.66E-03 | 1.51E-02 |
| ENSG00000023697 | DERA | -0.17 | 0.06 | 6.03E-03 | 1.60E-02 |
| ENSG00000157212 | PAXIP1 | -0.17 | 0.06 | 6.10E-03 | 1.61E-02 |
| ENSG00000143256 | PFDN2 | -0.17 | 0.06 | 7.06E-03 | 1.85E-02 |
| ENSG00000018699 | TTC27 | -0.17 | 0.07 | 8.51E-03 | 2.19E-02 |
| ENSG00000177728 | TMEM94 | -0.17 | 0.07 | 9.28E-03 | 2.36E-02 |
| ENSG00000214753 | HNRNPUL2 | -0.17 | 0.07 | 9.31E-03 | 2.37E-02 |
| ENSG00000272602 | ZNF595 | -0.17 | 0.07 | 9.55E-03 | 2.42E-02 |
| ENSG00000100342 | APOL1 | -0.17 | 0.07 | 9.57E-03 | 2.43E-02 |
| ENSG00000141258 | SGSM2 | -0.17 | 0.07 | 1.26E-02 | 3.11E-02 |
| ENSG00000107186 | MPDZ | -0.17 | 0.07 | 1.52E-02 | 3.67E-02 |
| ENSG00000104343 | UBE2W | -0.17 | 0.07 | 1.89E-02 | 4.49E-02 |
| ENSG00000079999 | KEAP1 | -0.17 | 0.07 | 1.91E-02 | 4.53E-02 |
| ENSG00000048828 | FAM120A | -0.16 | 0.03 | 2.56E-06 | 1.05E-05 |
| ENSG00000078018 | MAP2 | -0.16 | 0.04 | 2.65E-05 | 9.80E-05 |
| ENSG00000253729 | PRKDC | -0.16 | 0.04 | 1.19E-04 | 4.11E-04 |
| ENSG00000116288 | PARK7 | -0.16 | 0.04 | 1.40E-04 | 4.78E-04 |
| ENSG00000100220 | RTCB | -0.16 | 0.05 | 5.03E-04 | 1.60E-03 |
| ENSG00000075624 | ACTB | -0.16 | 0.05 | 5.89E-04 | 1.86E-03 |
| ENSG00000163798 | SLC4A1AP | -0.16 | 0.05 | 7.56E-04 | 2.34E-03 |
| ENSG00000091164 | TXNL1 | -0.16 | 0.05 | 1.17E-03 | 3.52E-03 |
| ENSG00000177469 | CAVIN1 | -0.16 | 0.05 | 1.35E-03 | 4.03E-03 |
| ENSG00000116120 | FARSB | -0.16 | 0.05 | 1.35E-03 | 4.03E-03 |
| ENSG00000170921 | TANC2 | -0.16 | 0.05 | 1.38E-03 | 4.10E-03 |
| ENSG00000134283 | PPHLN1 | -0.16 | 0.05 | 1.45E-03 | 4.28E-03 |
| ENSG00000013374 | NUB1 | -0.16 | 0.05 | 1.88E-03 | 5.48E-03 |
| ENSG00000164024 | METAP1 | -0.16 | 0.05 | 1.93E-03 | 5.62E-03 |
| ENSG00000120008 | WDR11 | -0.16 | 0.05 | 1.97E-03 | 5.71E-03 |
| ENSG00000119718 | EIF2B2 | -0.16 | 0.05 | 1.97E-03 | 5.72E-03 |
| ENSG00000141219 | C17orf80 | -0.16 | 0.05 | 2.20E-03 | 6.33E-03 |
| ENSG00000101343 | CRNKL1 | -0.16 | 0.05 | 2.32E-03 | 6.64E-03 |
| ENSG00000177370 | TIMM22 | -0.16 | 0.05 | 2.58E-03 | 7.35E-03 |
| ENSG00000170113 | NIPA1 | -0.16 | 0.06 | 3.66E-03 | 1.01E-02 |
| ENSG00000178691 | SUZ12 | -0.16 | 0.06 | 4.04E-03 | 1.11E-02 |
| ENSG00000123575 | FAM199X | -0.16 | 0.06 | 4.35E-03 | 1.18E-02 |
| ENSG00000137312 | FLOT1 | -0.16 | 0.06 | 4.79E-03 | 1.29E-02 |
| ENSG00000128609 | NDUFA5 | -0.16 | 0.06 | 5.99E-03 | 1.59E-02 |
| ENSG00000169826 | CSGALNACT2 | -0.16 | 0.06 | 6.21E-03 | 1.64E-02 |
| ENSG00000080845 | DLGAP4 | -0.16 | 0.06 | 6.69E-03 | 1.76E-02 |
| ENSG00000101096 | NFATC2 | -0.16 | 0.06 | 8.56E-03 | 2.20E-02 |
| ENSG00000077157 | PPP1R12B | -0.16 | 0.06 | 8.60E-03 | 2.21E-02 |

| | | | | | |
|---|---|---|---|---|---|
| ENSG00000204264 | PSMB8 | -0.16 | 0.06 | 8.74E-03 | 2.24E-02 |
| ENSG00000101347 | SAMHD1 | -0.16 | 0.06 | 9.00E-03 | 2.30E-02 |
| ENSG00000004059 | ARF5 | -0.16 | 0.06 | 9.32E-03 | 2.37E-02 |
| ENSG00000151353 | TMEM18 | -0.16 | 0.06 | 9.46E-03 | 2.40E-02 |
| ENSG00000103942 | HOMER2 | -0.16 | 0.06 | 9.76E-03 | 2.47E-02 |
| ENSG00000138668 | HNRNPD | -0.16 | 0.07 | 1.04E-02 | 2.63E-02 |
| ENSG00000145414 | NAF1 | -0.16 | 0.06 | 1.08E-02 | 2.71E-02 |
| ENSG00000242485 | MRPL20 | -0.16 | 0.06 | 1.08E-02 | 2.71E-02 |
| ENSG00000116212 | LRRC42 | -0.16 | 0.07 | 1.38E-02 | 3.37E-02 |
| ENSG00000105829 | BET1 | -0.16 | 0.06 | 1.39E-02 | 3.38E-02 |
| ENSG00000011451 | WIZ | -0.16 | 0.07 | 1.46E-02 | 3.56E-02 |
| ENSG00000164610 | RP9 | -0.16 | 0.06 | 1.50E-02 | 3.64E-02 |
| ENSG00000213160 | KLHL23 | -0.16 | 0.07 | 1.67E-02 | 4.00E-02 |
| ENSG00000166794 | PPIB | -0.16 | 0.07 | 2.07E-02 | 4.87E-02 |
| ENSG00000151148 | UBE3B | -0.15 | 0.03 | 3.25E-06 | 1.32E-05 |
| ENSG00000134138 | MEIS2 | -0.15 | 0.04 | 5.73E-05 | 2.05E-04 |
| ENSG00000166963 | MAP1A | -0.15 | 0.04 | 1.03E-04 | 3.58E-04 |
| ENSG00000138095 | LRPPRC | -0.15 | 0.04 | 1.91E-04 | 6.43E-04 |
| ENSG00000158864 | NDUFS2 | -0.15 | 0.04 | 2.02E-04 | 6.77E-04 |
| ENSG00000130508 | PXDN | -0.15 | 0.04 | 2.55E-04 | 8.42E-04 |
| ENSG00000218336 | TENM3 | -0.15 | 0.04 | 2.88E-04 | 9.46E-04 |
| ENSG00000156026 | MCU | -0.15 | 0.04 | 6.09E-04 | 1.92E-03 |
| ENSG00000204138 | PHACTR4 | -0.15 | 0.04 | 7.54E-04 | 2.34E-03 |
| ENSG00000111540 | RAB5B | -0.15 | 0.05 | 1.06E-03 | 3.21E-03 |
| ENSG00000125354 | SEPTIN6 | -0.15 | 0.05 | 1.33E-03 | 3.95E-03 |
| ENSG00000173418 | NAA20 | -0.15 | 0.05 | 1.42E-03 | 4.21E-03 |
| ENSG00000109919 | MTCH2 | -0.15 | 0.05 | 1.49E-03 | 4.40E-03 |
| ENSG00000079332 | SAR1A | -0.15 | 0.05 | 1.75E-03 | 5.11E-03 |
| ENSG00000177425 | PAWR | -0.15 | 0.05 | 1.77E-03 | 5.16E-03 |
| ENSG00000119487 | MAPKAP1 | -0.15 | 0.05 | 1.82E-03 | 5.32E-03 |
| ENSG00000111652 | COPS7A | -0.15 | 0.05 | 2.25E-03 | 6.46E-03 |
| ENSG00000138107 | ACTR1A | -0.15 | 0.05 | 2.28E-03 | 6.54E-03 |
| ENSG00000166167 | BTRC | -0.15 | 0.05 | 2.29E-03 | 6.57E-03 |
| ENSG00000117523 | PRRC2C | -0.15 | 0.05 | 2.44E-03 | 6.96E-03 |
| ENSG00000018189 | RUFY3 | -0.15 | 0.05 | 2.66E-03 | 7.55E-03 |
| ENSG00000166181 | API5 | -0.15 | 0.05 | 2.87E-03 | 8.08E-03 |
| ENSG00000062650 | WAPL | -0.15 | 0.05 | 2.97E-03 | 8.35E-03 |
| ENSG00000135597 | REPS1 | -0.15 | 0.05 | 3.17E-03 | 8.86E-03 |
| ENSG00000163482 | STK36 | -0.15 | 0.05 | 3.62E-03 | 1.00E-02 |
| ENSG00000177302 | TOP3A | -0.15 | 0.05 | 4.26E-03 | 1.16E-02 |
| ENSG00000160310 | PRMT2 | -0.15 | 0.05 | 4.36E-03 | 1.18E-02 |
| ENSG00000163930 | BAP1 | -0.15 | 0.05 | 4.58E-03 | 1.24E-02 |
| ENSG00000132953 | XPO4 | -0.15 | 0.05 | 4.70E-03 | 1.27E-02 |
| ENSG00000132716 | DCAF8 | -0.15 | 0.05 | 5.40E-03 | 1.44E-02 |
| ENSG00000172466 | ZNF24 | -0.15 | 0.06 | 6.21E-03 | 1.64E-02 |
| ENSG00000004700 | RECQL | -0.15 | 0.06 | 8.19E-03 | 2.11E-02 |
| ENSG00000076321 | KLHL20 | -0.15 | 0.06 | 8.22E-03 | 2.12E-02 |
| ENSG00000155115 | GTF3C6 | -0.15 | 0.06 | 9.35E-03 | 2.38E-02 |
| ENSG00000136940 | PDCL | -0.15 | 0.06 | 9.85E-03 | 2.49E-02 |
| ENSG00000161813 | LARP4 | -0.15 | 0.06 | 9.91E-03 | 2.51E-02 |

| | | | | | |
|---|---|---|---|---|---|
| ENSG00000169919 | GUSB | -0.15 | 0.06 | 1.01E-02 | 2.56E-02 |
| ENSG00000112837 | TBX18 | -0.15 | 0.06 | 1.06E-02 | 2.66E-02 |
| ENSG00000169047 | IRS1 | -0.15 | 0.06 | 1.07E-02 | 2.70E-02 |
| ENSG00000115419 | GLS | -0.15 | 0.06 | 1.11E-02 | 2.78E-02 |
| ENSG00000099940 | SNAP29 | -0.15 | 0.06 | 1.14E-02 | 2.84E-02 |
| ENSG00000123094 | RASSF8 | -0.15 | 0.06 | 1.22E-02 | 3.03E-02 |
| ENSG00000151458 | ANKRD50 | -0.15 | 0.06 | 1.29E-02 | 3.17E-02 |
| ENSG00000163946 | TASOR | -0.15 | 0.06 | 1.29E-02 | 3.18E-02 |
| ENSG00000163536 | SERPINI1 | -0.15 | 0.06 | 1.53E-02 | 3.70E-02 |
| ENSG00000122778 | KIAA1549 | -0.15 | 0.06 | 1.56E-02 | 3.76E-02 |
| ENSG00000173926 | MARCHF3 | -0.15 | 0.06 | 1.67E-02 | 4.00E-02 |
| ENSG00000119599 | DCAF4 | -0.15 | 0.06 | 1.77E-02 | 4.21E-02 |
| ENSG00000164815 | ORC5 | -0.15 | 0.06 | 1.80E-02 | 4.28E-02 |
| ENSG00000150054 | MPP7 | -0.15 | 0.06 | 1.81E-02 | 4.29E-02 |
| ENSG00000129933 | MAU2 | -0.15 | 0.06 | 1.86E-02 | 4.42E-02 |
| ENSG00000102081 | FMR1 | -0.15 | 0.06 | 1.99E-02 | 4.70E-02 |
| ENSG00000151689 | INPP1 | -0.15 | 0.07 | 2.04E-02 | 4.80E-02 |
| ENSG00000183258 | DDX41 | -0.15 | 0.07 | 2.05E-02 | 4.83E-02 |
| ENSG00000196569 | LAMA2 | -0.14 | 0.03 | 3.57E-05 | 1.30E-04 |
| ENSG00000155463 | OXA1L | -0.14 | 0.04 | 3.05E-04 | 9.98E-04 |
| ENSG00000159131 | GART | -0.14 | 0.04 | 6.07E-04 | 1.91E-03 |
| ENSG00000145349 | CAMK2D | -0.14 | 0.04 | 1.47E-03 | 4.35E-03 |
| ENSG00000116062 | MSH6 | -0.14 | 0.05 | 1.93E-03 | 5.60E-03 |
| ENSG00000127804 | METTL16 | -0.14 | 0.05 | 2.53E-03 | 7.21E-03 |
| ENSG00000150768 | DLAT | -0.14 | 0.05 | 3.23E-03 | 9.03E-03 |
| ENSG00000136811 | ODF2 | -0.14 | 0.05 | 3.36E-03 | 9.34E-03 |
| ENSG00000204574 | ABCF1 | -0.14 | 0.05 | 3.81E-03 | 1.05E-02 |
| ENSG00000204568 | MRPS18B | -0.14 | 0.05 | 3.86E-03 | 1.06E-02 |
| ENSG00000137200 | CMTR1 | -0.14 | 0.05 | 4.15E-03 | 1.13E-02 |
| ENSG00000257218 | GATC | -0.14 | 0.05 | 4.31E-03 | 1.17E-02 |
| ENSG00000173692 | PSMD1 | -0.14 | 0.05 | 4.36E-03 | 1.18E-02 |
| ENSG00000141543 | EIF4A3 | -0.14 | 0.05 | 4.85E-03 | 1.31E-02 |
| ENSG00000184743 | ATL3 | -0.14 | 0.05 | 4.97E-03 | 1.34E-02 |
| ENSG00000001167 | NFYA | -0.14 | 0.05 | 5.55E-03 | 1.48E-02 |
| ENSG00000110172 | CHORDC1 | -0.14 | 0.05 | 5.99E-03 | 1.59E-02 |
| ENSG00000073969 | NSF | -0.14 | 0.05 | 6.54E-03 | 1.72E-02 |
| ENSG00000088682 | COQ9 | -0.14 | 0.05 | 6.82E-03 | 1.79E-02 |
| ENSG00000004534 | RBM6 | -0.14 | 0.05 | 7.58E-03 | 1.97E-02 |
| ENSG00000109654 | TRIM2 | -0.14 | 0.05 | 8.66E-03 | 2.22E-02 |
| ENSG00000160293 | VAV2 | -0.14 | 0.06 | 1.00E-02 | 2.53E-02 |
| ENSG00000115904 | SOS1 | -0.14 | 0.06 | 1.05E-02 | 2.63E-02 |
| ENSG00000165055 | METTL2B | -0.14 | 0.06 | 1.09E-02 | 2.73E-02 |
| ENSG00000072121 | ZFYVE26 | -0.14 | 0.06 | 1.20E-02 | 2.98E-02 |
| ENSG00000107937 | GTPBP4 | -0.14 | 0.05 | 1.22E-02 | 3.03E-02 |
| ENSG00000007392 | LUC7L | -0.14 | 0.06 | 1.33E-02 | 3.26E-02 |
| ENSG00000138750 | NUP54 | -0.14 | 0.06 | 1.37E-02 | 3.35E-02 |
| ENSG00000167699 | GLOD4 | -0.14 | 0.06 | 1.38E-02 | 3.38E-02 |
| ENSG00000055208 | TAB2 | -0.14 | 0.06 | 1.49E-02 | 3.61E-02 |
| ENSG00000163939 | PBRM1 | -0.14 | 0.06 | 1.56E-02 | 3.77E-02 |
| ENSG00000126804 | ZBTB1 | -0.14 | 0.06 | 1.68E-02 | 4.02E-02 |

| | | | | | |
|---|---|---|---|---|---|
| ENSG00000124217 | MOCS3 | -0.14 | 0.06 | 1.83E-02 | 4.35E-02 |
| ENSG00000037897 | METTL1 | -0.14 | 0.06 | 1.97E-02 | 4.64E-02 |
| ENSG00000108055 | SMC3 | -0.14 | 0.06 | 2.06E-02 | 4.85E-02 |
| ENSG00000163191 | S100A11 | -0.13 | 0.04 | 2.55E-04 | 8.44E-04 |
| ENSG00000196419 | XRCC6 | -0.13 | 0.03 | 3.03E-04 | 9.93E-04 |
| ENSG00000130741 | EIF2S3 | -0.13 | 0.04 | 6.33E-04 | 1.98E-03 |
| ENSG00000165119 | HNRNPK | -0.13 | 0.04 | 7.01E-04 | 2.19E-03 |
| ENSG00000162909 | CAPN2 | -0.13 | 0.04 | 7.41E-04 | 2.30E-03 |
| ENSG00000090621 | PABPC4 | -0.13 | 0.04 | 1.22E-03 | 3.67E-03 |
| ENSG00000106609 | TMEM248 | -0.13 | 0.04 | 1.33E-03 | 3.96E-03 |
| ENSG00000234741 | GAS5 | -0.13 | 0.04 | 1.84E-03 | 5.38E-03 |
| ENSG00000124789 | NUP153 | -0.13 | 0.04 | 1.96E-03 | 5.69E-03 |
| ENSG00000114942 | EEF1B2 | -0.13 | 0.04 | 2.08E-03 | 5.99E-03 |
| ENSG00000115568 | ZNF142 | -0.13 | 0.04 | 2.15E-03 | 6.21E-03 |
| ENSG00000079785 | DDX1 | -0.13 | 0.04 | 2.20E-03 | 6.32E-03 |
| ENSG00000137547 | MRPL15 | -0.13 | 0.04 | 2.58E-03 | 7.34E-03 |
| ENSG00000135930 | EIF4E2 | -0.13 | 0.04 | 2.76E-03 | 7.80E-03 |
| ENSG00000104447 | TRPS1 | -0.13 | 0.04 | 3.50E-03 | 9.71E-03 |
| ENSG00000147650 | LRP12 | -0.13 | 0.05 | 3.75E-03 | 1.03E-02 |
| ENSG00000040933 | INPP4A | -0.13 | 0.05 | 4.72E-03 | 1.28E-02 |
| ENSG00000110090 | CPT1A | -0.13 | 0.05 | 4.89E-03 | 1.32E-02 |
| ENSG00000136026 | CKAP4 | -0.13 | 0.05 | 5.02E-03 | 1.35E-02 |
| ENSG00000138758 | SEPTIN11 | -0.13 | 0.05 | 5.69E-03 | 1.51E-02 |
| ENSG00000111670 | GNPTAB | -0.13 | 0.05 | 6.22E-03 | 1.64E-02 |
| ENSG00000111057 | KRT18 | -0.13 | 0.05 | 6.37E-03 | 1.68E-02 |
| ENSG00000089006 | SNX5 | -0.13 | 0.05 | 7.42E-03 | 1.93E-02 |
| ENSG00000096070 | BRPF3 | -0.13 | 0.05 | 7.96E-03 | 2.06E-02 |
| ENSG00000169410 | PTPN9 | -0.13 | 0.05 | 8.00E-03 | 2.07E-02 |
| ENSG00000107262 | BAG1 | -0.13 | 0.05 | 8.02E-03 | 2.07E-02 |
| ENSG00000095574 | IKZF5 | -0.13 | 0.05 | 8.10E-03 | 2.09E-02 |
| ENSG00000108518 | PFN1 | -0.13 | 0.05 | 1.05E-02 | 2.64E-02 |
| ENSG00000101000 | PROCR | -0.13 | 0.05 | 1.08E-02 | 2.71E-02 |
| ENSG00000185591 | SP1 | -0.13 | 0.05 | 1.12E-02 | 2.80E-02 |
| ENSG00000144369 | FAM171B | -0.13 | 0.05 | 1.15E-02 | 2.86E-02 |
| ENSG00000166575 | TMEM135 | -0.13 | 0.05 | 1.21E-02 | 3.00E-02 |
| ENSG00000151150 | ANK3 | -0.13 | 0.05 | 1.34E-02 | 3.28E-02 |
| ENSG00000196586 | MYO6 | -0.13 | 0.05 | 1.36E-02 | 3.32E-02 |
| ENSG00000159202 | UBE2Z | -0.13 | 0.05 | 1.51E-02 | 3.66E-02 |
| ENSG00000110768 | GTF2H1 | -0.13 | 0.05 | 1.57E-02 | 3.78E-02 |
| ENSG00000003756 | RBM5 | -0.13 | 0.05 | 1.69E-02 | 4.04E-02 |
| ENSG00000117153 | KLHL12 | -0.13 | 0.06 | 2.07E-02 | 4.86E-02 |
| ENSG00000156011 | PSD3 | -0.13 | 0.06 | 2.09E-02 | 4.91E-02 |
| ENSG00000132182 | NUP210 | -0.13 | 0.05 | 2.10E-02 | 4.93E-02 |
| ENSG00000118705 | RPN2 | -0.12 | 0.03 | 3.09E-04 | 1.01E-03 |
| ENSG00000242265 | PEG10 | -0.12 | 0.04 | 9.90E-04 | 3.01E-03 |
| ENSG00000148843 | PDCD11 | -0.12 | 0.04 | 1.14E-03 | 3.43E-03 |
| ENSG00000153956 | CACNA2D1 | -0.12 | 0.04 | 1.47E-03 | 4.35E-03 |
| ENSG00000163349 | HIPK1 | -0.12 | 0.04 | 1.82E-03 | 5.32E-03 |
| ENSG00000171940 | ZNF217 | -0.12 | 0.04 | 2.85E-03 | 8.05E-03 |
| ENSG00000155438 | NIFK | -0.12 | 0.04 | 3.97E-03 | 1.09E-02 |

| | | | | | |
|---|---|---|---|---|---|
| ENSG00000164741 | DLC1 | -0.12 | 0.04 | 5.82E-03 | 1.55E-02 |
| ENSG00000198668 | CALM1 | -0.12 | 0.05 | 6.46E-03 | 1.70E-02 |
| ENSG00000165699 | TSC1 | -0.12 | 0.04 | 7.16E-03 | 1.87E-02 |
| ENSG00000105939 | ZC3HAV1 | -0.12 | 0.05 | 7.59E-03 | 1.97E-02 |
| ENSG00000257923 | CUX1 | -0.12 | 0.04 | 7.85E-03 | 2.03E-02 |
| ENSG00000116237 | ICMT | -0.12 | 0.04 | 8.47E-03 | 2.18E-02 |
| ENSG00000128059 | PPAT | -0.12 | 0.05 | 9.19E-03 | 2.34E-02 |
| ENSG00000070061 | ELP1 | -0.12 | 0.05 | 9.81E-03 | 2.49E-02 |
| ENSG00000101558 | VAPA | -0.12 | 0.05 | 1.05E-02 | 2.64E-02 |
| ENSG00000061936 | SFSWAP | -0.12 | 0.05 | 1.12E-02 | 2.79E-02 |
| ENSG00000116221 | MRPL37 | -0.12 | 0.05 | 1.28E-02 | 3.15E-02 |
| ENSG00000143933 | CALM2 | -0.12 | 0.05 | 1.30E-02 | 3.20E-02 |
| ENSG00000100029 | PES1 | -0.12 | 0.05 | 1.52E-02 | 3.69E-02 |
| ENSG00000147133 | TAF1 | -0.12 | 0.05 | 1.55E-02 | 3.74E-02 |
| ENSG00000022277 | RTF2 | -0.12 | 0.05 | 1.75E-02 | 4.18E-02 |
| ENSG00000154553 | PDLIM3 | -0.11 | 0.04 | 2.24E-03 | 6.44E-03 |
| ENSG00000082641 | NFE2L1 | -0.11 | 0.04 | 2.24E-03 | 6.45E-03 |
| ENSG00000186591 | UBE2H | -0.11 | 0.04 | 2.28E-03 | 6.55E-03 |
| ENSG00000107862 | GBF1 | -0.11 | 0.04 | 2.46E-03 | 7.01E-03 |
| ENSG00000115758 | ODC1 | -0.11 | 0.04 | 3.53E-03 | 9.77E-03 |
| ENSG00000128607 | KLHDC10 | -0.11 | 0.04 | 4.37E-03 | 1.19E-02 |
| ENSG00000165678 | GHITM | -0.11 | 0.04 | 4.47E-03 | 1.21E-02 |
| ENSG00000171316 | CHD7 | -0.11 | 0.04 | 4.86E-03 | 1.31E-02 |
| ENSG00000103512 | NOMO1 | -0.11 | 0.04 | 5.21E-03 | 1.40E-02 |
| ENSG00000163872 | YEATS2 | -0.11 | 0.04 | 5.53E-03 | 1.48E-02 |
| ENSG00000067225 | PKM | -0.11 | 0.04 | 6.09E-03 | 1.61E-02 |
| ENSG00000119682 | AREL1 | -0.11 | 0.04 | 6.18E-03 | 1.63E-02 |
| ENSG00000105202 | FBL | -0.11 | 0.04 | 6.44E-03 | 1.70E-02 |
| ENSG00000144283 | PKP4 | -0.11 | 0.04 | 8.80E-03 | 2.25E-02 |
| ENSG00000181915 | ADO | -0.11 | 0.04 | 9.44E-03 | 2.40E-02 |
| ENSG00000138434 | ITPRID2 | -0.11 | 0.04 | 9.96E-03 | 2.52E-02 |
| ENSG00000159352 | PSMD4 | -0.11 | 0.04 | 1.03E-02 | 2.60E-02 |
| ENSG00000138246 | DNAJC13 | -0.11 | 0.04 | 1.22E-02 | 3.02E-02 |
| ENSG00000130699 | TAF4 | -0.11 | 0.05 | 1.35E-02 | 3.31E-02 |
| ENSG00000101019 | UQCC1 | -0.11 | 0.05 | 1.37E-02 | 3.36E-02 |
| ENSG00000115241 | PPM1G | -0.11 | 0.05 | 1.39E-02 | 3.39E-02 |
| ENSG00000189337 | KAZN | -0.11 | 0.05 | 1.86E-02 | 4.42E-02 |
| ENSG00000133019 | CHRM3 | -0.11 | 0.05 | 1.96E-02 | 4.64E-02 |
| ENSG00000152894 | PTPRK | -0.11 | 0.05 | 2.07E-02 | 4.85E-02 |
| ENSG00000197343 | ZNF655 | -0.11 | 0.05 | 2.09E-02 | 4.90E-02 |
| ENSG00000101346 | POFUT1 | -0.1 | 0.04 | 9.51E-03 | 2.41E-02 |
| ENSG00000145730 | PAM | -0.1 | 0.04 | 9.78E-03 | 2.48E-02 |
| ENSG00000106484 | MEST | -0.1 | 0.04 | 1.18E-02 | 2.94E-02 |
| ENSG00000118257 | NRP2 | -0.1 | 0.04 | 1.41E-02 | 3.44E-02 |
| ENSG00000169045 | HNRNPH1 | -0.1 | 0.04 | 1.50E-02 | 3.64E-02 |
| ENSG00000278845 | MRPL45 | -0.1 | 0.04 | 1.51E-02 | 3.65E-02 |
| ENSG00000108349 | CASC3 | -0.1 | 0.04 | 1.64E-02 | 3.95E-02 |
| ENSG00000088986 | DYNLL1 | -0.09 | 0.04 | 1.02E-02 | 2.58E-02 |
| ENSG00000105058 | FAM32A | -0.09 | 0.04 | 1.63E-02 | 3.92E-02 |
| ENSG00000151491 | EPS8 | -0.09 | 0.04 | 1.90E-02 | 4.50E-02 |

| | | | | | |
|---|---|---|---|---|---|
| ENSG00000110107 | PRPF19 | -0.09 | 0.04 | 2.05E-02 | 4.82E-02 |
| ENSG00000076108 | BAZ2A | -0.08 | 0.03 | 1.68E-02 | 4.02E-02 |

**Table S3. Human primers used for RT-qPCR analysis.**

| | PRIMER (5’-3’) |
|---|---|
| DHCR7 forward | TCCACAGCCATGTGACCAATGC |
| DHCR7 reverse | CGAAGTGGTCATGGCAGATGTC |
| HMGCR forward | ACTTCGTGTTCATGACTTTC |
| HMGCR reverse | GACATAATCATCTTGACCCTC |
| PERK forward | GTCCCAAGGCTTTGGAATCTGTC |
| PERK reverse | CCTACCAAGACAGGAGTTCTGG |
| ATF4 forward | TTCTCCAGCGACAAGGCTAAGG |
| ATF4 reverse | CTCCAACATCCAATCTGTCCCG |
| CHOP forward | GGTATGAGGACCTGCAAGAGGT |
| CHOP reverse | CTTGTGACCTCTGCTGGTTCTG |
| GADD34 forward | TCCGACTGCAAAGGCGGCTCA |
| GADD34 reverse | CAGCCAGGAAATGGACAGTGAC |
| PROX1 forward | CTGAAGACCTACTTCTCCGACG |
| PROX1 reverse | GATGGCTTGACGTGCGTACTTC |
| HPRT forward | CTCATGGACTGATTATGGACAGGAC |
| HPRT reverse | GCAGGTCAGCAAAGAACTTATAGCC |

**Table S2. Upregulated genes in DHCR7 silenced RD cells**

| | symbol | log2FoldChange | lfcSE | pvalue | padj |
|---|---|---|---|---|---|
| ENSG00000290862 | NMRAL2P | 5.64 | 0.68 | 1.21E-22 | 1.25E-21 |
| ENSG00000187479 | C11orf96 | 4.03 | 0.14 | 7.41E-187 | 1.24E-184 |
| ENSG00000173110 | HSPA6 | 3.8 | 0.14 | 1.19E-174 | 1.68E-172 |
| ENSG00000151012 | SLC7A11 | 3.77 | 0.06 | 0.00E+00 | 0.00E+00 |
| ENSG00000228955 | RPL7AP82 | 3.75 | 0.28 | 1.28E-46 | 2.67E-45 |
| ENSG00000128965 | CHAC1 | 3.69 | 0.12 | 1.13E-212 | 2.60E-210 |
| ENSG00000152463 | OLAH | 3.29 | 0.39 | 9.41E-21 | 9.01E-20 |
| ENSG00000130766 | SESN2 | 3.25 | 0.09 | 4.55E-300 | 2.25E-297 |
| ENSG00000196517 | SLC6A9 | 3.18 | 0.1 | 4.09E-235 | 1.17E-232 |
| ENSG00000105327 | BBC3 | 3.17 | 0.29 | 1.81E-31 | 2.55E-30 |
| ENSG00000128342 | LIF | 3.16 | 0.06 | 0.00E+00 | 0.00E+00 |
| ENSG00000128165 | ADM2 | 3.13 | 0.12 | 2.14E-155 | 2.40E-153 |
| ENSG00000070669 | ASNS | 2.92 | 0.2 | 7.44E-50 | 1.66E-48 |
| ENSG00000101255 | TRIB3 | 2.88 | 0.08 | 3.72E-273 | 1.45E-270 |
| ENSG00000015592 | STMN4 | 2.87 | 0.35 | 2.06E-18 | 1.78E-17 |
| ENSG00000269688 | | 2.8 | 0.18 | 6.98E-59 | 1.92E-57 |
| ENSG00000222389 | RNU2-28P | 2.79 | 1.2 | 4.12E-07 | 1.82E-06 |
| ENSG00000121552 | CSTA | 2.76 | 0.17 | 5.70E-64 | 1.72E-62 |
| ENSG00000124194 | GDAP1L1 | 2.76 | 0.43 | 3.41E-13 | 2.30E-12 |
| ENSG00000136010 | ALDH1L2 | 2.73 | 0.06 | 0.00E+00 | 0.00E+00 |
| ENSG00000124731 | TREM1 | 2.73 | 0.48 | 3.44E-11 | 2.05E-10 |
| ENSG00000081041 | CXCL2 | 2.66 | 0.1 | 3.51E-159 | 4.12E-157 |
| ENSG00000248927 | | 2.65 | 0.58 | 8.20E-09 | 4.12E-08 |
| ENSG00000268812 | LIF-AS2 | 2.56 | 0.27 | 3.12E-23 | 3.29E-22 |
| ENSG00000073150 | PANX2 | 2.52 | 0.1 | 5.93E-132 | 5.15E-130 |
| ENSG00000105550 | FGF21 | 2.52 | 1.09 | 1.36E-06 | 5.73E-06 |
| ENSG00000115009 | CCL20 | 2.48 | 0.08 | 2.79E-205 | 5.94E-203 |
| ENSG00000186480 | INSIG1 | 2.44 | 0.04 | 0.00E+00 | 0.00E+00 |
| ENSG00000188396 | DYNLT4 | 2.41 | 0.17 | 5.20E-50 | 1.17E-48 |
| ENSG00000176907 | TCIM | 2.4 | 0.13 | 4.09E-81 | 1.71E-79 |
| ENSG00000130513 | GDF15 | 2.39 | 0.08 | 1.67E-178 | 2.48E-176 |
| ENSG00000285873 | | 2.39 | 0.43 | 1.19E-10 | 6.89E-10 |
| ENSG00000184678 | H2BC21 | 2.37 | 0.08 | 5.70E-203 | 1.17E-200 |
| ENSG00000237719 | | 2.37 | 0.35 | 7.47E-14 | 5.24E-13 |
| ENSG00000112972 | HMGCS1 | 2.36 | 0.05 | 0.00E+00 | 0.00E+00 |
| ENSG00000139269 | INHBE | 2.36 | 0.09 | 3.59E-166 | 4.48E-164 |
| ENSG00000168209 | DDIT4 | 2.34 | 0.09 | 6.12E-150 | 6.41E-148 |
| ENSG00000289530 | | 2.33 | 0.15 | 7.74E-54 | 1.92E-52 |
| ENSG00000223802 | CERS1 | 2.32 | 0.19 | 2.88E-35 | 4.56E-34 |
| ENSG00000290041 | | 2.31 | 0.18 | 2.06E-40 | 3.73E-39 |
| ENSG00000235513 | L3MBTL2-AS1 | 2.31 | 0.2 | 2.53E-33 | 3.77E-32 |
| ENSG00000251127 | | 2.3 | 0.37 | 3.66E-12 | 2.32E-11 |
| ENSG00000206417 | H1-10-AS1 | 2.29 | 0.31 | 7.52E-16 | 5.79E-15 |
| ENSG00000290398 | ASNSP1 | 2.29 | 0.55 | 1.16E-07 | 5.33E-07 |
| ENSG00000290923 | PGM5P2 | 2.28 | 0.12 | 1.67E-82 | 7.21E-81 |
| ENSG00000175197 | DDIT3 | 2.26 | 0.09 | 2.36E-153 | 2.57E-151 |
| ENSG00000245532 | NEAT1 | 2.22 | 0.08 | 3.65E-163 | 4.39E-161 |
| ENSG00000158079 | PTPDC1 | 2.21 | 0.06 | 0.00E+00 | 0.00E+00 |

| | | | | | |
|---|---|---|---|---|---|
| ENSG00000174456 | C12orf76 | 2.21 | 0.07 | 1.78E-230 | 4.85E-228 |
| ENSG00000115902 | SLC1A4 | 2.21 | 0.07 | 2.24E-201 | 4.51E-199 |
| ENSG00000100276 | RASL10A | 2.21 | 0.25 | 3.87E-21 | 3.78E-20 |
| ENSG00000265972 | TXNIP | 2.2 | 0.04 | 0.00E+00 | 0.00E+00 |
| ENSG00000250033 | SLC7A11-AS1 | 2.19 | 0.68 | 3.00E-06 | 1.22E-05 |
| ENSG00000111981 | ULBP1 | 2.18 | 0.11 | 7.35E-88 | 3.44E-86 |
| ENSG00000266401 | | 2.18 | 0.65 | 2.12E-06 | 8.74E-06 |
| ENSG00000161013 | MGAT4B | 2.17 | 0.07 | 9.08E-216 | 2.14E-213 |
| ENSG00000259518 | LINC01583 | 2.17 | 0.45 | 1.01E-08 | 5.05E-08 |
| ENSG00000251350 | LINC02475 | 2.17 | 0.66 | 2.67E-06 | 1.09E-05 |
| ENSG00000169429 | CXCL8 | 2.16 | 0.06 | 7.44E-291 | 3.48E-288 |
| ENSG00000172216 | CEBPB | 2.16 | 0.08 | 4.37E-173 | 6.05E-171 |
| ENSG00000144485 | HES6 | 2.15 | 0.12 | 4.72E-73 | 1.73E-71 |
| ENSG00000223770 | CACNA2D1-AS1 | 2.15 | 0.15 | 7.77E-46 | 1.59E-44 |
| ENSG00000228626 | | 2.15 | 1.08 | 7.39E-06 | 2.90E-05 |
| ENSG00000139547 | RDH16 | 2.14 | 0.27 | 2.27E-17 | 1.88E-16 |
| ENSG00000121769 | FABP3 | 2.13 | 0.05 | 0.00E+00 | 0.00E+00 |
| ENSG00000236255 | | 2.13 | 0.12 | 1.63E-71 | 5.74E-70 |
| ENSG00000164438 | TLX3 | 2.13 | 0.48 | 5.50E-08 | 2.60E-07 |
| ENSG00000166123 | GPT2 | 2.12 | 0.07 | 2.05E-180 | 3.14E-178 |
| ENSG00000127561 | SYNGR3 | 2.12 | 0.14 | 2.47E-57 | 6.57E-56 |
| ENSG00000163347 | CLDN1 | 2.12 | 0.22 | 8.31E-24 | 8.97E-23 |
| ENSG00000197580 | BCO2 | 2.12 | 0.34 | 4.38E-12 | 2.76E-11 |
| ENSG00000172318 | B3GALT1 | 2.11 | 0.13 | 4.52E-58 | 1.22E-56 |
| ENSG00000182379 | NXPH4 | 2.11 | 0.14 | 2.02E-56 | 5.24E-55 |
| ENSG00000273802 | H2BC8 | 2.11 | 0.22 | 5.26E-23 | 5.51E-22 |
| ENSG00000168386 | FILIP1L | 2.1 | 0.06 | 0.00E+00 | 0.00E+00 |
| ENSG00000288880 | | 2.09 | 0.3 | 4.12E-14 | 2.92E-13 |
| ENSG00000225828 | FAM229A | 2.08 | 0.15 | 3.72E-46 | 7.66E-45 |
| ENSG00000287796 | | 2.07 | 0.88 | 1.52E-05 | 5.79E-05 |
| ENSG00000172201 | ID4 | 2.06 | 0.12 | 4.88E-63 | 1.44E-61 |
| ENSG00000185022 | MAFF | 2.05 | 0.08 | 6.33E-150 | 6.61E-148 |
| ENSG00000285588 | | 2.05 | 0.32 | 2.22E-12 | 1.43E-11 |
| ENSG00000276075 | | 2.05 | 0.6 | 2.90E-06 | 1.18E-05 |
| ENSG00000230131 | LINC02641 | 2.05 | 0.71 | 9.72E-06 | 3.76E-05 |
| ENSG00000226744 | | 2.04 | 0.33 | 7.42E-12 | 4.61E-11 |
| ENSG00000105825 | TFPI2 | 2.03 | 0.07 | 4.62E-198 | 8.83E-196 |
| ENSG00000241218 | CSP2 | 2.03 | 0.57 | 1.79E-06 | 7.41E-06 |
| ENSG00000171533 | MAP6 | 2.02 | 0.26 | 2.72E-16 | 2.15E-15 |
| ENSG00000279672 | | 2 | 0.2 | 9.23E-25 | 1.04E-23 |
| ENSG00000170006 | TMEM154 | 2 | 0.22 | 5.36E-21 | 5.20E-20 |
| ENSG00000273017 | | 2 | 0.37 | 5.85E-10 | 3.21E-09 |
| ENSG00000287839 | | 2 | 0.4 | 8.02E-09 | 4.04E-08 |
| ENSG00000130522 | JUND | 1.99 | 0.09 | 2.61E-120 | 1.93E-118 |
| ENSG00000166682 | TMPRSS5 | 1.99 | 0.32 | 4.28E-12 | 2.71E-11 |
| ENSG00000163092 | XIRP2 | 1.98 | 0.1 | 1.80E-95 | 9.74E-94 |
| ENSG00000180044 | C3orf80 | 1.98 | 0.24 | 1.81E-18 | 1.57E-17 |
| ENSG00000178531 | CTXN1 | 1.96 | 0.19 | 7.04E-26 | 8.19E-25 |
| ENSG00000185633 | NDUFA4L2 | 1.96 | 0.21 | 9.44E-23 | 9.79E-22 |
| ENSG00000229431 | MED8-AS1 | 1.96 | 0.24 | 7.40E-18 | 6.26E-17 |

| | | | | | |
|---|---|---|---|---|---|
| ENSG00000170909 | OSCAR | 1.95 | 0.2 | 1.91E-24 | 2.12E-23 |
| ENSG00000262119 | | 1.95 | 0.65 | 1.12E-05 | 4.30E-05 |
| ENSG00000290877 | ROCK1P1 | 1.94 | 0.32 | 2.94E-11 | 1.77E-10 |
| ENSG00000255920 | CCND2-AS1 | 1.94 | 0.34 | 2.57E-10 | 1.45E-09 |
| ENSG00000272574 | | 1.94 | 0.39 | 6.50E-09 | 3.29E-08 |
| ENSG00000147852 | VLDLR | 1.93 | 0.1 | 1.66E-92 | 8.60E-91 |
| ENSG00000198496 | NBR2 | 1.92 | 0.15 | 5.35E-38 | 9.16E-37 |
| ENSG00000125538 | IL1B | 1.91 | 0.29 | 4.80E-13 | 3.21E-12 |
| ENSG00000129990 | SYT5 | 1.9 | 0.18 | 8.16E-27 | 9.84E-26 |
| ENSG00000198535 | C2CD4A | 1.9 | 0.28 | 1.04E-13 | 7.23E-13 |
| ENSG00000165071 | TMEM71 | 1.9 | 0.43 | 1.29E-07 | 5.91E-07 |
| ENSG00000135069 | PSAT1 | 1.89 | 0.04 | 0.00E+00 | 0.00E+00 |
| ENSG00000215105 | TTC3P1 | 1.89 | 0.12 | 3.65E-54 | 9.15E-53 |
| ENSG00000172071 | EIF2AK3 | 1.87 | 0.06 | 7.67E-232 | 2.11E-229 |
| ENSG00000234233 | KCNH1-IT1 | 1.87 | 0.16 | 6.03E-35 | 9.42E-34 |
| ENSG00000130487 | KLHDC7B | 1.87 | 0.29 | 2.90E-12 | 1.85E-11 |
| ENSG00000272267 | | 1.87 | 0.31 | 3.50E-11 | 2.09E-10 |
| ENSG00000272079 | | 1.87 | 0.38 | 9.97E-09 | 4.99E-08 |
| ENSG00000200152 | RNU6-216P | 1.87 | 0.44 | 2.58E-07 | 1.16E-06 |
| ENSG00000213015 | ZNF580 | 1.86 | 0.11 | 1.31E-60 | 3.71E-59 |
| ENSG00000249626 | PPP4R2P4 | 1.86 | 0.48 | 1.09E-06 | 4.61E-06 |
| ENSG00000160013 | PTGIR | 1.85 | 0.11 | 1.17E-70 | 4.09E-69 |
| ENSG00000163393 | SLC22A15 | 1.85 | 0.27 | 9.85E-14 | 6.86E-13 |
| ENSG00000289183 | | 1.83 | 0.67 | 3.02E-05 | 1.11E-04 |
| ENSG00000167508 | MVD | 1.82 | 0.09 | 7.17E-89 | 3.43E-87 |
| ENSG00000161011 | SQSTM1 | 1.81 | 0.05 | 0.00E+00 | 0.00E+00 |
| ENSG00000165807 | PPP1R36 | 1.81 | 0.36 | 5.22E-09 | 2.66E-08 |
| ENSG00000111653 | ING4 | 1.79 | 0.06 | 3.05E-170 | 4.05E-168 |
| ENSG00000104856 | RELB | 1.79 | 0.09 | 1.19E-93 | 6.29E-92 |
| ENSG00000181218 | H2AC25 | 1.79 | 0.13 | 1.08E-41 | 2.04E-40 |
| ENSG00000290241 | | 1.79 | 0.54 | 7.43E-06 | 2.91E-05 |
| ENSG00000155980 | KIF5A | 1.78 | 0.07 | 2.07E-165 | 2.57E-163 |
| ENSG00000158406 | H4C8 | 1.78 | 0.12 | 1.43E-51 | 3.33E-50 |
| ENSG00000128564 | VGF | 1.78 | 0.18 | 1.54E-25 | 1.77E-24 |
| ENSG00000092621 | PHGDH | 1.77 | 0.08 | 4.90E-117 | 3.47E-115 |
| ENSG00000116183 | PAPPA2 | 1.77 | 0.29 | 1.45E-11 | 8.86E-11 |
| ENSG00000258096 | SLC38A2-AS1 | 1.76 | 0.41 | 2.06E-07 | 9.32E-07 |
| ENSG00000180573 | H2AC6 | 1.75 | 0.09 | 2.66E-94 | 1.42E-92 |
| ENSG00000178809 | TRIM73 | 1.75 | 0.4 | 1.66E-07 | 7.55E-07 |
| ENSG00000219507 | FTH1P8 | 1.75 | 0.43 | 7.07E-07 | 3.05E-06 |
| ENSG00000279699 | | 1.75 | 0.45 | 1.18E-06 | 4.97E-06 |
| ENSG00000251867 | | 1.75 | 0.47 | 2.02E-06 | 8.33E-06 |
| ENSG00000049130 | KITLG | 1.74 | 0.09 | 9.00E-77 | 3.55E-75 |
| ENSG00000172738 | TMEM217 | 1.74 | 0.16 | 1.35E-29 | 1.79E-28 |
| ENSG00000145390 | USP53 | 1.73 | 0.06 | 2.08E-188 | 3.56E-186 |
| ENSG00000274565 | | 1.73 | 0.33 | 1.92E-09 | 1.02E-08 |
| ENSG00000076826 | CAMSAP3 | 1.73 | 0.33 | 2.26E-09 | 1.19E-08 |
| ENSG00000262001 | DLGAP1-AS2 | 1.72 | 0.1 | 5.57E-74 | 2.08E-72 |
| ENSG00000184545 | DUSP8 | 1.72 | 0.13 | 1.87E-42 | 3.58E-41 |
| ENSG00000290989 | | 1.72 | 0.14 | 3.69E-35 | 5.82E-34 |

| | | | | | |
|---|---|---|---|---|---|
| ENSG00000290833 | | 1.72 | 0.2 | 9.21E-19 | 8.09E-18 |
| ENSG00000189229 | | 1.72 | 0.21 | 4.94E-18 | 4.20E-17 |
| ENSG00000162687 | KCNT2 | 1.72 | 0.51 | 8.12E-06 | 3.17E-05 |
| ENSG00000269694 | | 1.72 | 0.53 | 1.19E-05 | 4.56E-05 |
| ENSG00000104413 | ESRP1 | 1.72 | 0.54 | 1.33E-05 | 5.09E-05 |
| ENSG00000138675 | FGF5 | 1.71 | 0.09 | 3.98E-83 | 1.75E-81 |
| ENSG00000267002 | | 1.71 | 0.1 | 6.61E-69 | 2.21E-67 |
| ENSG00000254682 | | 1.7 | 0.16 | 2.95E-29 | 3.86E-28 |
| ENSG00000290999 | | 1.7 | 0.36 | 2.92E-08 | 1.41E-07 |
| ENSG00000289506 | | 1.7 | 0.57 | 2.46E-05 | 9.11E-05 |
| ENSG00000269821 | KCNQ1OT1 | 1.69 | 0.12 | 3.81E-44 | 7.57E-43 |
| ENSG00000277449 | CEBPB-AS1 | 1.69 | 0.25 | 1.74E-13 | 1.19E-12 |
| ENSG00000225177 | NHSL1-AS1 | 1.69 | 0.32 | 1.47E-09 | 7.86E-09 |
| ENSG00000187840 | EIF4EBP1 | 1.68 | 0.09 | 4.94E-82 | 2.10E-80 |
| ENSG00000120437 | ACAT2 | 1.67 | 0.06 | 3.88E-190 | 6.77E-188 |
| ENSG00000174951 | FUT1 | 1.67 | 0.2 | 1.79E-18 | 1.55E-17 |
| ENSG00000052802 | MSMO1 | 1.66 | 0.04 | 0.00E+00 | 0.00E+00 |
| ENSG00000113161 | HMGCR | 1.66 | 0.04 | 0.00E+00 | 0.00E+00 |
| ENSG00000175894 | TSPEAR | 1.66 | 0.13 | 1.59E-41 | 2.98E-40 |
| ENSG00000271551 | | 1.66 | 0.45 | 3.06E-06 | 1.24E-05 |
| ENSG00000260941 | LINC00622 | 1.66 | 0.52 | 1.51E-05 | 5.73E-05 |
| ENSG00000100889 | PCK2 | 1.65 | 0.11 | 1.19E-56 | 3.10E-55 |
| ENSG00000176593 | | 1.65 | 0.11 | 6.57E-52 | 1.55E-50 |
| ENSG00000272841 | MAP3K4-AS1 | 1.65 | 0.12 | 3.37E-48 | 7.24E-47 |
| ENSG00000087494 | PTHLH | 1.65 | 0.23 | 1.22E-14 | 8.89E-14 |
| ENSG00000269952 | | 1.65 | 0.32 | 4.30E-09 | 2.21E-08 |
| ENSG00000112715 | VEGFA | 1.64 | 0.06 | 1.08E-149 | 1.12E-147 |
| ENSG00000127585 | FBXL16 | 1.64 | 0.13 | 3.10E-39 | 5.47E-38 |
| ENSG00000173930 | SLCO4C1 | 1.64 | 0.19 | 1.85E-19 | 1.68E-18 |
| ENSG00000166183 | ASPG | 1.64 | 0.28 | 9.95E-11 | 5.77E-10 |
| ENSG00000275221 | H2AC15 | 1.64 | 0.42 | 1.61E-06 | 6.70E-06 |
| ENSG00000111266 | DUSP16 | 1.63 | 0.05 | 8.75E-196 | 1.64E-193 |
| ENSG00000167615 | LENG8 | 1.63 | 0.08 | 8.07E-104 | 4.91E-102 |
| ENSG00000159200 | RCAN1 | 1.62 | 0.04 | 0.00E+00 | 0.00E+00 |
| ENSG00000047346 | ATOSA | 1.62 | 0.09 | 3.99E-72 | 1.43E-70 |
| ENSG00000289153 | | 1.62 | 0.22 | 4.31E-15 | 3.21E-14 |
| ENSG00000250284 | | 1.62 | 0.23 | 4.41E-14 | 3.12E-13 |
| ENSG00000272931 | LRRC8D-DT | 1.62 | 0.33 | 2.07E-08 | 1.01E-07 |
| ENSG00000283108 | | 1.62 | 0.33 | 2.36E-08 | 1.15E-07 |
| ENSG00000067064 | IDI1 | 1.61 | 0.05 | 5.66E-257 | 1.95E-254 |
| ENSG00000113070 | HBEGF | 1.61 | 0.05 | 1.01E-197 | 1.91E-195 |
| ENSG00000130164 | LDLR | 1.61 | 0.05 | 3.64E-194 | 6.62E-192 |
| ENSG00000184205 | TSPYL2 | 1.61 | 0.06 | 5.44E-138 | 5.12E-136 |
| ENSG00000247400 | DNAJC3-DT | 1.61 | 0.21 | 3.01E-16 | 2.37E-15 |
| ENSG00000146733 | PSPH | 1.6 | 0.06 | 7.19E-168 | 9.23E-166 |
| ENSG00000132196 | HSD17B7 | 1.6 | 0.06 | 6.13E-148 | 6.32E-146 |
| ENSG00000184988 | TMEM106A | 1.6 | 0.15 | 7.56E-29 | 9.76E-28 |
| ENSG00000099840 | IZUMO4 | 1.6 | 0.25 | 3.75E-12 | 2.38E-11 |
| ENSG00000261020 | | 1.6 | 0.26 | 1.42E-11 | 8.65E-11 |
| ENSG00000276107 | THBS1-IT1 | 1.6 | 0.33 | 2.72E-08 | 1.32E-07 |

| | | | | | |
|---|---|---|---|---|---|
| ENSG00000225217 | HSPA7 | 1.6 | 0.69 | 1.48E-04 | 5.05E-04 |
| ENSG00000073146 | MOV10L1 | 1.6 | 0.82 | 2.01E-04 | 6.74E-04 |
| ENSG00000140941 | MAP1LC3B | 1.59 | 0.05 | 1.21E-191 | 2.15E-189 |
| ENSG00000215440 | NPEPL1 | 1.59 | 0.11 | 1.70E-45 | 3.45E-44 |
| ENSG00000133134 | BEX2 | 1.59 | 0.15 | 1.11E-26 | 1.34E-25 |
| ENSG00000158887 | MPZ | 1.59 | 0.22 | 3.22E-14 | 2.30E-13 |
| ENSG00000227815 | | 1.59 | 1 | 1.02E-04 | 3.54E-04 |
| ENSG00000087086 | FTL | 1.58 | 0.06 | 1.51E-176 | 2.20E-174 |
| ENSG00000105856 | HBP1 | 1.58 | 0.06 | 2.02E-164 | 2.47E-162 |
| ENSG00000108176 | DNAJC12 | 1.58 | 0.11 | 3.84E-45 | 7.77E-44 |
| ENSG00000106113 | CRHR2 | 1.58 | 0.39 | 8.26E-07 | 3.53E-06 |
| ENSG00000051108 | HERPUD1 | 1.57 | 0.04 | 0.00E+00 | 0.00E+00 |
| ENSG00000154319 | FAM167A | 1.57 | 0.2 | 4.04E-17 | 3.31E-16 |
| ENSG00000165349 | SLC7A3 | 1.57 | 0.22 | 1.35E-14 | 9.79E-14 |
| ENSG00000271966 | ARFGEF1-DT | 1.57 | 0.38 | 6.05E-07 | 2.63E-06 |
| ENSG00000179454 | KLHL28 | 1.56 | 0.06 | 6.57E-157 | 7.52E-155 |
| ENSG00000287593 | | 1.56 | 0.25 | 1.61E-11 | 9.80E-11 |
| ENSG00000234694 | CDC20-DT | 1.56 | 0.46 | 1.05E-05 | 4.04E-05 |
| ENSG00000106105 | GARS1 | 1.55 | 0.03 | 0.00E+00 | 0.00E+00 |
| ENSG00000148408 | CACNA1B | 1.55 | 0.07 | 6.83E-113 | 4.59E-111 |
| ENSG00000184838 | PRR16 | 1.55 | 0.09 | 2.13E-75 | 8.14E-74 |
| ENSG00000276791 | | 1.55 | 0.18 | 1.14E-18 | 9.97E-18 |
| ENSG00000197405 | C5AR1 | 1.55 | 0.3 | 6.52E-09 | 3.31E-08 |
| ENSG00000273355 | | 1.55 | 0.37 | 6.49E-07 | 2.81E-06 |
| ENSG00000261652 | PIERCE2 | 1.55 | 0.64 | 1.54E-04 | 5.24E-04 |
| ENSG00000259943 | ZMPSTE24-DT | 1.54 | 0.1 | 1.59E-52 | 3.81E-51 |
| ENSG00000105146 | AURKC | 1.54 | 0.21 | 1.61E-15 | 1.22E-14 |
| ENSG00000276334 | | 1.54 | 0.32 | 3.13E-08 | 1.51E-07 |
| ENSG00000251639 | | 1.54 | 0.35 | 2.67E-07 | 1.20E-06 |
| ENSG00000117983 | MUC5B | 1.54 | 0.52 | 4.06E-05 | 1.48E-04 |
| ENSG00000128590 | DNAJB9 | 1.53 | 0.06 | 2.97E-165 | 3.67E-163 |
| ENSG00000120129 | DUSP1 | 1.53 | 0.08 | 4.02E-82 | 1.71E-80 |
| ENSG00000167996 | FTH1 | 1.53 | 0.1 | 6.65E-59 | 1.83E-57 |
| ENSG00000273344 | PAXIP1-DT | 1.53 | 0.12 | 1.33E-36 | 2.19E-35 |
| ENSG00000180535 | BHLHA15 | 1.53 | 0.23 | 4.79E-13 | 3.20E-12 |
| ENSG00000287200 | | 1.53 | 0.39 | 1.42E-06 | 5.95E-06 |
| ENSG00000286230 | | 1.53 | 0.43 | 6.14E-06 | 2.43E-05 |
| ENSG00000044574 | HSPA5 | 1.52 | 0.03 | 0.00E+00 | 0.00E+00 |
| ENSG00000273238 | TMEM271 | 1.52 | 0.13 | 1.17E-33 | 1.76E-32 |
| ENSG00000277128 | | 1.52 | 0.2 | 7.94E-16 | 6.10E-15 |
| ENSG00000248476 | BACH1-IT1 | 1.52 | 0.22 | 6.58E-14 | 4.62E-13 |
| ENSG00000223638 | RFPL4A | 1.52 | 0.24 | 6.48E-12 | 4.04E-11 |
| ENSG00000167995 | BEST1 | 1.52 | 0.41 | 3.80E-06 | 1.53E-05 |
| ENSG00000290707 | MED15P9 | 1.52 | 0.43 | 6.96E-06 | 2.73E-05 |
| ENSG00000258581 | | 1.52 | 0.81 | 3.10E-04 | 1.01E-03 |
| ENSG00000197822 | OCLN | 1.51 | 0.1 | 2.24E-49 | 4.93E-48 |
| ENSG00000183307 | TMEM121B | 1.51 | 0.1 | 9.51E-49 | 2.07E-47 |
| ENSG00000290832 | | 1.51 | 0.14 | 5.62E-27 | 6.82E-26 |
| ENSG00000230183 | CNOT6LP1 | 1.51 | 1 | 1.38E-04 | 4.71E-04 |
| ENSG00000166173 | LARP6 | 1.5 | 0.08 | 1.64E-87 | 7.60E-86 |

| | | | | | |
|---|---|---|---|---|---|
| ENSG00000237310 | LINC03011 | 1.5 | 0.28 | 1.26E-09 | 6.78E-09 |
| ENSG00000108309 | RUNDC3A | 1.5 | 0.28 | 1.64E-09 | 8.71E-09 |
| ENSG00000279283 | | 1.5 | 0.49 | 3.78E-05 | 1.38E-04 |
| ENSG00000213316 | LTC4S | 1.5 | 0.61 | 1.75E-04 | 5.91E-04 |
| ENSG00000166770 | ZNF667-AS1 | 1.49 | 0.06 | 7.33E-128 | 6.00E-126 |
| ENSG00000249592 | PCGF3-AS1 | 1.49 | 0.1 | 7.50E-53 | 1.81E-51 |
| ENSG00000163734 | CXCL3 | 1.49 | 0.12 | 2.22E-38 | 3.84E-37 |
| ENSG00000233937 | CTC-338M12.4 | 1.49 | 0.3 | 2.22E-08 | 1.08E-07 |
| ENSG00000287211 | | 1.49 | 0.35 | 4.93E-07 | 2.16E-06 |
| ENSG00000236824 | BCYRN1 | 1.49 | 0.58 | 1.41E-04 | 4.82E-04 |
| ENSG00000153234 | NR4A2 | 1.48 | 0.09 | 5.10E-62 | 1.48E-60 |
| ENSG00000134899 | ERCC5 | 1.48 | 0.23 | 5.01E-12 | 3.15E-11 |
| ENSG00000204228 | HSD17B8 | 1.48 | 0.3 | 2.97E-08 | 1.44E-07 |
| ENSG00000256258 | | 1.48 | 0.37 | 1.25E-06 | 5.27E-06 |
| ENSG00000288753 | | 1.48 | 0.41 | 5.40E-06 | 2.15E-05 |
| ENSG00000125787 | GNRH2 | 1.48 | 0.53 | 8.37E-05 | 2.94E-04 |
| ENSG00000172432 | GTPBP2 | 1.47 | 0.05 | 1.06E-175 | 1.54E-173 |
| ENSG00000006459 | KDM7A | 1.47 | 0.09 | 4.39E-62 | 1.28E-60 |
| ENSG00000143630 | HCN3 | 1.47 | 0.15 | 1.13E-23 | 1.21E-22 |
| ENSG00000130590 | SAMD10 | 1.47 | 0.15 | 1.23E-23 | 1.32E-22 |
| ENSG00000196118 | CFAP119 | 1.47 | 0.31 | 3.89E-08 | 1.86E-07 |
| ENSG00000273542 | H4C12 | 1.47 | 0.4 | 4.85E-06 | 1.93E-05 |
| ENSG00000085514 | PILRA | 1.47 | 0.43 | 1.14E-05 | 4.37E-05 |
| ENSG00000122861 | PLAU | 1.46 | 0.04 | 0.00E+00 | 0.00E+00 |
| ENSG00000248810 | LINC02432 | 1.46 | 0.2 | 1.04E-14 | 7.63E-14 |
| ENSG00000198915 | RASGEF1A | 1.46 | 0.37 | 1.55E-06 | 6.46E-06 |
| ENSG00000141854 | MISP3 | 1.46 | 0.39 | 4.71E-06 | 1.88E-05 |
| ENSG00000287059 | | 1.46 | 0.83 | 4.21E-04 | 1.35E-03 |
| ENSG00000137801 | THBS1 | 1.45 | 0.05 | 7.23E-205 | 1.53E-202 |
| ENSG00000149809 | TM7SF2 | 1.45 | 0.1 | 1.06E-50 | 2.43E-49 |
| ENSG00000248275 | TRIM52-AS1 | 1.45 | 0.12 | 2.40E-37 | 4.02E-36 |
| ENSG00000152467 | ZSCAN1 | 1.45 | 0.2 | 1.10E-14 | 8.01E-14 |
| ENSG00000079459 | FDFT1 | 1.44 | 0.04 | 0.00E+00 | 0.00E+00 |
| ENSG00000033030 | ZCCHC8 | 1.44 | 0.05 | 1.13E-163 | 1.38E-161 |
| ENSG00000134070 | IRAK2 | 1.44 | 0.1 | 1.89E-46 | 3.92E-45 |
| ENSG00000140044 | JDP2 | 1.44 | 0.11 | 2.01E-41 | 3.76E-40 |
| ENSG00000182326 | C1S | 1.44 | 0.13 | 1.34E-29 | 1.78E-28 |
| ENSG00000236423 | LINC01134 | 1.44 | 0.21 | 2.86E-13 | 1.94E-12 |
| ENSG00000227799 | CDPF1P1 | 1.44 | 0.28 | 8.75E-09 | 4.40E-08 |
| ENSG00000247950 | SEC24B-AS1 | 1.44 | 0.33 | 2.88E-07 | 1.29E-06 |
| ENSG00000286863 | | 1.44 | 0.36 | 1.57E-06 | 6.56E-06 |
| ENSG00000228218 | ATF4P3 | 1.44 | 0.4 | 5.62E-06 | 2.23E-05 |
| ENSG00000266010 | GATA6-AS1 | 1.44 | 0.85 | 4.38E-04 | 1.41E-03 |
| ENSG00000110921 | MVK | 1.43 | 0.08 | 2.97E-72 | 1.07E-70 |
| ENSG00000095752 | IL11 | 1.43 | 0.1 | 1.67E-52 | 4.00E-51 |
| ENSG00000106948 | AKNA | 1.43 | 0.1 | 3.73E-47 | 7.87E-46 |
| ENSG00000225855 | RUSC1-AS1 | 1.43 | 0.13 | 5.32E-29 | 6.93E-28 |
| ENSG00000291174 | | 1.43 | 0.17 | 4.07E-18 | 3.47E-17 |
| ENSG00000272086 | GOLPH3-DT | 1.43 | 0.28 | 8.78E-09 | 4.41E-08 |
| ENSG00000279692 | | 1.43 | 0.4 | 8.84E-06 | 3.43E-05 |

| | | | | | |
|---|---|---|---|---|---|
| ENSG00000260949 | | 1.43 | 0.44 | 2.27E-05 | 8.46E-05 |
| ENSG00000256955 | | 1.43 | 0.61 | 2.83E-04 | 9.31E-04 |
| ENSG00000232442 | MHENCR | 1.42 | 0.14 | 9.34E-25 | 1.05E-23 |
| ENSG00000254995 | STX16-NPEPL1 | 1.42 | 0.16 | 7.48E-20 | 6.91E-19 |
| ENSG00000255284 | GATD1-DT | 1.42 | 0.21 | 1.38E-13 | 9.51E-13 |
| ENSG00000250240 | | 1.42 | 0.3 | 7.88E-08 | 3.69E-07 |
| ENSG00000198363 | ASPH | 1.41 | 0.04 | 9.78E-250 | 3.08E-247 |
| ENSG00000108639 | SYNGR2 | 1.41 | 0.07 | 4.30E-102 | 2.55E-100 |
| ENSG00000170345 | FOS | 1.41 | 0.09 | 2.27E-57 | 6.03E-56 |
| ENSG00000123610 | TNFAIP6 | 1.41 | 0.23 | 1.80E-11 | 1.09E-10 |
| ENSG00000207425 | Y_RNA | 1.41 | 0.26 | 2.21E-09 | 1.16E-08 |
| ENSG00000107736 | CDH23 | 1.41 | 0.3 | 8.10E-08 | 3.78E-07 |
| ENSG00000273218 | | 1.41 | 0.35 | 1.50E-06 | 6.29E-06 |
| ENSG00000177989 | ODF3B | 1.41 | 0.36 | 2.26E-06 | 9.27E-06 |
| ENSG00000067715 | SYT1 | 1.41 | 0.46 | 4.42E-05 | 1.60E-04 |
| ENSG00000230606 | APPAT | 1.41 | 0.47 | 5.88E-05 | 2.10E-04 |
| ENSG00000225138 | SLC9A3-AS1 | 1.4 | 0.14 | 2.49E-24 | 2.74E-23 |
| ENSG00000260236 | PTPN23-DT | 1.4 | 0.19 | 5.09E-15 | 3.78E-14 |
| ENSG00000267321 | SNHG30 | 1.4 | 0.2 | 2.66E-14 | 1.91E-13 |
| ENSG00000257126 | FOXG1-AS1 | 1.4 | 0.26 | 2.86E-09 | 1.49E-08 |
| ENSG00000277531 | PNMA8C | 1.4 | 0.31 | 1.45E-07 | 6.65E-07 |
| ENSG00000236404 | VLDLR-AS1 | 1.4 | 0.38 | 6.13E-06 | 2.42E-05 |
| ENSG00000269386 | RAB11B-AS1 | 1.4 | 0.4 | 1.07E-05 | 4.12E-05 |
| ENSG00000164898 | FMC1 | 1.4 | 0.4 | 1.08E-05 | 4.15E-05 |
| ENSG00000135842 | NIBAN1 | 1.39 | 0.07 | 2.56E-96 | 1.40E-94 |
| ENSG00000109083 | IFT20 | 1.39 | 0.1 | 4.44E-48 | 9.53E-47 |
| ENSG00000168300 | PCMTD1 | 1.39 | 0.11 | 2.68E-41 | 5.01E-40 |
| ENSG00000117862 | TXNDC12 | 1.39 | 0.13 | 1.43E-28 | 1.82E-27 |
| ENSG00000290654 | | 1.39 | 0.16 | 1.85E-19 | 1.68E-18 |
| ENSG00000290951 | | 1.39 | 0.17 | 4.54E-17 | 3.72E-16 |
| ENSG00000255566 | | 1.39 | 0.55 | 2.01E-04 | 6.74E-04 |
| ENSG00000267595 | BRCA1P1 | 1.39 | 0.59 | 2.93E-04 | 9.60E-04 |
| ENSG00000104808 | DHDH | 1.39 | 0.62 | 3.83E-04 | 1.24E-03 |
| ENSG00000134548 | SPX | 1.39 | 0.69 | 5.62E-04 | 1.78E-03 |
| ENSG00000128272 | ATF4 | 1.38 | 0.04 | 8.19E-302 | 4.12E-299 |
| ENSG00000224116 | INHBA-AS1 | 1.38 | 0.21 | 2.42E-12 | 1.55E-11 |
| ENSG00000256092 | SBNO1-AS1 | 1.38 | 0.23 | 1.23E-10 | 7.10E-10 |
| ENSG00000261485 | PAN3-AS1 | 1.38 | 0.4 | 1.27E-05 | 4.84E-05 |
| ENSG00000164106 | SCRG1 | 1.38 | 0.61 | 3.93E-04 | 1.27E-03 |
| ENSG00000106366 | SERPINE1 | 1.37 | 0.03 | 0.00E+00 | 0.00E+00 |
| ENSG00000261824 | LINC00662 | 1.37 | 0.05 | 3.90E-203 | 8.04E-201 |
| ENSG00000178752 | ERFE | 1.37 | 0.06 | 1.72E-136 | 1.59E-134 |
| ENSG00000289586 | | 1.37 | 0.07 | 3.35E-91 | 1.68E-89 |
| ENSG00000107281 | NPDC1 | 1.37 | 0.13 | 7.18E-26 | 8.35E-25 |
| ENSG00000280213 | UCKL1-AS1 | 1.37 | 0.14 | 3.38E-25 | 3.84E-24 |
| ENSG00000124713 | GNMT | 1.37 | 0.37 | 5.58E-06 | 2.22E-05 |
| ENSG00000107593 | PKD2L1 | 1.37 | 0.86 | 5.92E-04 | 1.87E-03 |
| ENSG00000244953 | | 1.37 | 0.77 | 6.93E-04 | 2.16E-03 |
| ENSG00000153879 | CEBPG | 1.36 | 0.06 | 2.53E-133 | 2.24E-131 |
| ENSG00000147883 | CDKN2B | 1.36 | 0.06 | 4.45E-122 | 3.39E-120 |

| | | | | | |
|---|---|---|---|---|---|
| ENSG00000156510 | HKDC1 | 1.36 | 0.17 | 3.90E-17 | 3.19E-16 |
| ENSG00000278600 | | 1.36 | 0.2 | 2.85E-13 | 1.93E-12 |
| ENSG00000288596 | C8orf44 | 1.36 | 0.21 | 1.38E-12 | 8.96E-12 |
| ENSG00000289869 | | 1.36 | 0.49 | 1.14E-04 | 3.94E-04 |
| ENSG00000270760 | | 1.36 | 0.58 | 3.67E-04 | 1.19E-03 |
| ENSG00000115266 | APC2 | 1.35 | 0.11 | 2.05E-35 | 3.26E-34 |
| ENSG00000232931 | LINC00342 | 1.35 | 0.13 | 6.52E-28 | 8.13E-27 |
| ENSG00000273382 | TMEM167B-DT | 1.35 | 0.13 | 4.53E-26 | 5.32E-25 |
| ENSG00000258102 | MAP1LC3B2 | 1.35 | 0.19 | 3.24E-14 | 2.32E-13 |
| ENSG00000271344 | | 1.35 | 0.55 | 2.78E-04 | 9.15E-04 |
| ENSG00000259771 | | 1.35 | 0.65 | 5.84E-04 | 1.84E-03 |
| ENSG00000247572 | CKMT2-AS1 | 1.34 | 0.08 | 4.87E-63 | 1.44E-61 |
| ENSG00000152527 | PLEKHH2 | 1.34 | 0.08 | 4.61E-61 | 1.32E-59 |
| ENSG00000203950 | RTL8A | 1.34 | 0.09 | 4.14E-54 | 1.03E-52 |
| ENSG00000278535 | DHRS11 | 1.34 | 0.12 | 1.77E-32 | 2.56E-31 |
| ENSG00000243279 | PRAF2 | 1.34 | 0.17 | 2.78E-17 | 2.30E-16 |
| ENSG00000233822 | H2BC15 | 1.34 | 0.2 | 3.21E-13 | 2.17E-12 |
| ENSG00000279529 | | 1.34 | 0.26 | 6.25E-09 | 3.17E-08 |
| ENSG00000261526 | | 1.34 | 0.32 | 8.33E-07 | 3.56E-06 |
| ENSG00000235927 | NEXN-AS1 | 1.34 | 0.35 | 3.43E-06 | 1.39E-05 |
| ENSG00000110619 | CARS1 | 1.33 | 0.04 | 3.42E-194 | 6.31E-192 |
| ENSG00000013588 | GPRC5A | 1.33 | 0.04 | 3.46E-194 | 6.33E-192 |
| ENSG00000162772 | ATF3 | 1.33 | 0.06 | 1.11E-127 | 9.08E-126 |
| ENSG00000179085 | DPM3 | 1.33 | 0.11 | 6.65E-34 | 1.01E-32 |
| ENSG00000132003 | ZSWIM4 | 1.33 | 0.12 | 1.16E-29 | 1.54E-28 |
| ENSG00000133110 | POSTN | 1.33 | 0.13 | 9.15E-25 | 1.03E-23 |
| ENSG00000203814 | H2BC18 | 1.33 | 0.32 | 9.87E-07 | 4.19E-06 |
| ENSG00000258982 | | 1.33 | 0.4 | 2.38E-05 | 8.85E-05 |
| ENSG00000114796 | KLHL24 | 1.32 | 0.06 | 3.59E-116 | 2.49E-114 |
| ENSG00000111674 | ENO2 | 1.32 | 0.07 | 7.39E-76 | 2.86E-74 |
| ENSG00000173846 | PLK3 | 1.32 | 0.08 | 1.71E-70 | 5.97E-69 |
| ENSG00000101670 | LIPG | 1.32 | 0.09 | 9.69E-56 | 2.49E-54 |
| ENSG00000163683 | SMIM14 | 1.32 | 0.09 | 3.34E-54 | 8.38E-53 |
| ENSG00000166401 | SERPINB8 | 1.32 | 0.12 | 2.99E-31 | 4.19E-30 |
| ENSG00000123329 | ARHGAP9 | 1.32 | 0.12 | 5.45E-29 | 7.09E-28 |
| ENSG00000174871 | CNIH2 | 1.32 | 0.13 | 4.59E-27 | 5.57E-26 |
| ENSG00000247137 | ANKRD42-DT | 1.32 | 0.17 | 6.81E-16 | 5.26E-15 |
| ENSG00000279455 | | 1.32 | 0.27 | 3.80E-08 | 1.82E-07 |
| ENSG00000172426 | RSPH9 | 1.32 | 0.29 | 2.03E-07 | 9.21E-07 |
| ENSG00000272669 | | 1.32 | 0.35 | 4.59E-06 | 1.83E-05 |
| ENSG00000205791 | LOH12CR2 | 1.32 | 0.37 | 1.13E-05 | 4.35E-05 |
| ENSG00000108306 | FBXL20 | 1.31 | 0.07 | 5.27E-71 | 1.85E-69 |
| ENSG00000268205 | | 1.31 | 0.08 | 1.35E-57 | 3.61E-56 |
| ENSG00000171160 | MORN4 | 1.31 | 0.09 | 8.91E-55 | 2.26E-53 |
| ENSG00000278828 | H3C10 | 1.31 | 0.23 | 4.56E-10 | 2.52E-09 |
| ENSG00000228412 | LNC-LBCS | 1.31 | 0.27 | 2.91E-08 | 1.41E-07 |
| ENSG00000221883 | ARIH2OS | 1.31 | 0.29 | 2.22E-07 | 1.00E-06 |
| ENSG00000140839 | CLEC18B | 1.31 | 0.49 | 1.91E-04 | 6.43E-04 |
| ENSG00000251391 | | 1.31 | 0.97 | 3.22E-04 | 1.05E-03 |
| ENSG00000288930 | | 1.31 | 0.62 | 6.47E-04 | 2.03E-03 |

| | | | | | |
|---|---|---|---|---|---|
| ENSG00000100219 | XBP1 | 1.3 | 0.05 | 2.04E-177 | 3.01E-175 |
| ENSG00000115295 | CLIP4 | 1.3 | 0.07 | 2.53E-80 | 1.04E-78 |
| ENSG00000267100 | ILF3-DT | 1.3 | 0.08 | 4.81E-64 | 1.46E-62 |
| ENSG00000257704 | INAFM1 | 1.3 | 0.13 | 1.96E-23 | 2.09E-22 |
| ENSG00000128346 | C22orf23 | 1.3 | 0.15 | 9.80E-20 | 9.02E-19 |
| ENSG00000137331 | IER3 | 1.3 | 0.16 | 1.54E-17 | 1.29E-16 |
| ENSG00000137449 | CPEB2 | 1.3 | 0.21 | 5.30E-11 | 3.13E-10 |
| ENSG00000288899 | | 1.3 | 0.23 | 4.32E-10 | 2.39E-09 |
| ENSG00000226981 | ABHD17AP6 | 1.3 | 0.28 | 1.13E-07 | 5.22E-07 |
| ENSG00000148600 | CDHR1 | 1.3 | 0.3 | 6.28E-07 | 2.72E-06 |
| ENSG00000246263 | UBR5-DT | 1.3 | 0.39 | 2.16E-05 | 8.09E-05 |
| ENSG00000237781 | ADAMTSL4-AS2 | 1.3 | 0.45 | 1.14E-04 | 3.95E-04 |
| ENSG00000274492 | | 1.3 | 0.96 | 3.66E-04 | 1.19E-03 |
| ENSG00000104549 | SQLE | 1.29 | 0.03 | 0.00E+00 | 0.00E+00 |
| ENSG00000153443 | UBALD1 | 1.29 | 0.11 | 1.93E-35 | 3.07E-34 |
| ENSG00000274750 | H3C6 | 1.29 | 0.13 | 1.10E-23 | 1.18E-22 |
| ENSG00000230699 | | 1.29 | 0.15 | 2.43E-18 | 2.09E-17 |
| ENSG00000288983 | | 1.29 | 0.19 | 8.95E-13 | 5.88E-12 |
| ENSG00000279541 | | 1.29 | 0.33 | 3.59E-06 | 1.45E-05 |
| ENSG00000283930 | PLD5P1 | 1.29 | 0.34 | 4.97E-06 | 1.98E-05 |
| ENSG00000187242 | KRT12 | 1.29 | 0.84 | 9.26E-04 | 2.83E-03 |
| ENSG00000132718 | SYT11 | 1.28 | 0.04 | 2.03E-275 | 8.16E-273 |
| ENSG00000065060 | BLTP3A | 1.28 | 0.05 | 2.17E-133 | 1.93E-131 |
| ENSG00000170689 | HOXB9 | 1.28 | 0.07 | 1.69E-78 | 6.82E-77 |
| ENSG00000184602 | SNN | 1.28 | 0.08 | 8.46E-59 | 2.32E-57 |
| ENSG00000245849 | RAD51-AS1 | 1.28 | 0.14 | 8.26E-21 | 7.93E-20 |
| ENSG00000253304 | TMEM200B | 1.28 | 0.15 | 3.12E-19 | 2.80E-18 |
| ENSG00000261094 | | 1.28 | 0.17 | 6.35E-15 | 4.70E-14 |
| ENSG00000253716 | MINCR | 1.28 | 0.18 | 1.83E-14 | 1.32E-13 |
| ENSG00000273702 | | 1.28 | 0.31 | 1.22E-06 | 5.16E-06 |
| ENSG00000271913 | TAGAP-AS1 | 1.28 | 0.34 | 4.57E-06 | 1.83E-05 |
| ENSG00000288106 | | 1.28 | 0.42 | 6.06E-05 | 2.16E-04 |
| ENSG00000130711 | PRDM12 | 1.28 | 0.5 | 2.87E-04 | 9.41E-04 |
| ENSG00000273253 | TRABD-AS1 | 1.28 | 0.52 | 3.33E-04 | 1.08E-03 |
| ENSG00000259605 | | 1.28 | 0.77 | 1.09E-03 | 3.30E-03 |
| ENSG00000179774 | ATOH7 | 1.28 | 0.7 | 1.10E-03 | 3.33E-03 |
| ENSG00000142949 | PTPRF | 1.27 | 0.06 | 2.40E-102 | 1.43E-100 |
| ENSG00000125148 | MT2A | 1.27 | 0.07 | 1.21E-75 | 4.63E-74 |
| ENSG00000267519 | MIR23AHG | 1.27 | 0.11 | 3.23E-31 | 4.52E-30 |
| ENSG00000266714 | MYO15B | 1.27 | 0.21 | 6.73E-11 | 3.94E-10 |
| ENSG00000260285 | TRMT61A-DT | 1.27 | 0.23 | 8.58E-10 | 4.65E-09 |
| ENSG00000226416 | MRPL23-AS1 | 1.27 | 0.31 | 1.68E-06 | 6.98E-06 |
| ENSG00000268945 | | 1.27 | 0.32 | 2.06E-06 | 8.48E-06 |
| ENSG00000289929 | | 1.27 | 0.42 | 7.79E-05 | 2.74E-04 |
| ENSG00000278041 | | 1.27 | 0.47 | 2.08E-04 | 6.96E-04 |
| ENSG00000270021 | | 1.27 | 0.48 | 2.29E-04 | 7.61E-04 |
| ENSG00000261114 | | 1.27 | 0.5 | 2.95E-04 | 9.65E-04 |
| ENSG00000131080 | EDA2R | 1.27 | 0.55 | 4.89E-04 | 1.56E-03 |
| ENSG00000249755 | | 1.27 | 0.61 | 8.25E-04 | 2.54E-03 |
| ENSG00000105376 | ICAM5 | 1.26 | 0.1 | 3.98E-36 | 6.47E-35 |

| | | | | | |
|---|---|---|---|---|---|
| ENSG00000273117 | INSIG1-DT | 1.26 | 0.18 | 1.32E-13 | 9.13E-13 |
| ENSG00000102409 | BEX4 | 1.26 | 0.21 | 4.38E-11 | 2.60E-10 |
| ENSG00000167103 | PIP5KL1 | 1.26 | 0.23 | 2.63E-09 | 1.37E-08 |
| ENSG00000275437 | | 1.26 | 0.26 | 5.22E-08 | 2.47E-07 |
| ENSG00000271151 | | 1.26 | 0.46 | 1.93E-04 | 6.50E-04 |
| ENSG00000146216 | TTBK1 | 1.26 | 0.6 | 7.96E-04 | 2.46E-03 |
| ENSG00000261592 | | 1.26 | 0.84 | 9.94E-04 | 3.02E-03 |
| ENSG00000213028 | | 1.26 | 0.69 | 1.18E-03 | 3.54E-03 |
| ENSG00000177427 | MIEF2 | 1.25 | 0.14 | 1.52E-19 | 1.38E-18 |
| ENSG00000281501 | SEPSECS-AS1 | 1.25 | 0.35 | 1.11E-05 | 4.27E-05 |
| ENSG00000276916 | | 1.25 | 0.54 | 5.21E-04 | 1.66E-03 |
| ENSG00000105443 | CYTH2 | 1.24 | 0.07 | 1.40E-71 | 4.94E-70 |
| ENSG00000179523 | EIF3J-DT | 1.24 | 0.11 | 1.82E-28 | 2.31E-27 |
| ENSG00000272686 | WASL-DT | 1.24 | 0.12 | 1.62E-25 | 1.86E-24 |
| ENSG00000163624 | CDS1 | 1.24 | 0.28 | 3.51E-07 | 1.56E-06 |
| ENSG00000163293 | NIPAL1 | 1.24 | 0.3 | 1.11E-06 | 4.70E-06 |
| ENSG00000117143 | UAP1 | 1.23 | 0.05 | 3.53E-128 | 2.93E-126 |
| ENSG00000118523 | CCN2 | 1.23 | 0.05 | 4.93E-118 | 3.53E-116 |
| ENSG00000196182 | STK40 | 1.23 | 0.06 | 5.77E-93 | 3.04E-91 |
| ENSG00000102003 | SYP | 1.23 | 0.12 | 1.33E-24 | 1.47E-23 |
| ENSG00000177337 | DLGAP1-AS1 | 1.23 | 0.13 | 3.70E-21 | 3.62E-20 |
| ENSG00000180596 | H2BC4 | 1.23 | 0.16 | 2.93E-16 | 2.31E-15 |
| ENSG00000188710 | QRFP | 1.23 | 0.17 | 1.25E-14 | 9.11E-14 |
| ENSG00000256073 | URB1-AS1 | 1.23 | 0.19 | 1.17E-12 | 7.61E-12 |
| ENSG00000125551 | PLGLB2 | 1.23 | 0.29 | 8.64E-07 | 3.69E-06 |
| ENSG00000235888 | LINC02940 | 1.23 | 0.38 | 4.04E-05 | 1.47E-04 |
| ENSG00000197599 | CCDC154 | 1.23 | 0.4 | 7.52E-05 | 2.65E-04 |
| ENSG00000179141 | MTUS2-AS1 | 1.23 | 0.72 | 1.50E-03 | 4.42E-03 |
| ENSG00000123240 | OPTN | 1.22 | 0.06 | 1.13E-91 | 5.71E-90 |
| ENSG00000077150 | NFKB2 | 1.22 | 0.07 | 7.91E-74 | 2.95E-72 |
| ENSG00000188760 | TMEM198 | 1.22 | 0.1 | 3.82E-33 | 5.65E-32 |
| ENSG00000228393 | LINC01004 | 1.22 | 0.18 | 3.25E-13 | 2.19E-12 |
| ENSG00000214198 | TTC41P | 1.22 | 0.21 | 1.16E-10 | 6.67E-10 |
| ENSG00000272501 | | 1.22 | 0.21 | 2.68E-10 | 1.51E-09 |
| ENSG00000248161 | | 1.22 | 0.41 | 1.12E-04 | 3.86E-04 |
| ENSG00000165215 | CLDN3 | 1.22 | 0.48 | 3.24E-04 | 1.06E-03 |
| ENSG00000234500 | | 1.22 | 0.48 | 3.50E-04 | 1.14E-03 |
| ENSG00000260101 | AFTPH-DT | 1.22 | 0.51 | 5.01E-04 | 1.60E-03 |
| ENSG00000177462 | OR2T8 | 1.22 | 0.94 | 5.75E-04 | 1.81E-03 |
| ENSG00000238260 | | 1.22 | 0.72 | 1.55E-03 | 4.57E-03 |
| ENSG00000272582 | | 1.22 | 0.72 | 1.58E-03 | 4.65E-03 |
| ENSG00000031698 | SARS1 | 1.21 | 0.04 | 1.06E-199 | 2.06E-197 |
| ENSG00000164620 | RELL2 | 1.21 | 0.11 | 1.48E-30 | 2.04E-29 |
| ENSG00000204186 | ZDBF2 | 1.21 | 0.12 | 5.04E-26 | 5.89E-25 |
| ENSG00000174125 | TLR1 | 1.21 | 0.33 | 7.88E-06 | 3.08E-05 |
| ENSG00000136404 | TM6SF1 | 1.21 | 0.35 | 2.23E-05 | 8.34E-05 |
| ENSG00000228436 | RRAGC-DT | 1.21 | 0.35 | 2.30E-05 | 8.58E-05 |
| ENSG00000156042 | CFAP70 | 1.21 | 0.36 | 2.68E-05 | 9.92E-05 |
| ENSG00000259514 | | 1.21 | 0.39 | 7.59E-05 | 2.68E-04 |
| ENSG00000228817 | BACH1-IT2 | 1.21 | 0.45 | 2.40E-04 | 7.96E-04 |

| | | | | | |
|---|---|---|---|---|---|
| ENSG00000235545 | DOCK7-DT | 1.21 | 0.46 | 2.87E-04 | 9.42E-04 |
| ENSG00000185792 | NLRP9 | 1.21 | 0.55 | 7.35E-04 | 2.28E-03 |
| ENSG00000280402 | | 1.21 | 0.55 | 8.06E-04 | 2.49E-03 |
| ENSG00000011021 | CLCN6 | 1.2 | 0.07 | 2.63E-75 | 9.99E-74 |
| ENSG00000137496 | IL18BP | 1.2 | 0.08 | 1.13E-54 | 2.85E-53 |
| ENSG00000176658 | MYO1D | 1.2 | 0.1 | 8.66E-35 | 1.35E-33 |
| ENSG00000130477 | UNC13A | 1.2 | 0.11 | 2.80E-31 | 3.93E-30 |
| ENSG00000174721 | FGFBP3 | 1.2 | 0.14 | 2.18E-19 | 1.97E-18 |
| ENSG00000232388 | SMIM26 | 1.2 | 0.17 | 1.34E-13 | 9.24E-13 |
| ENSG00000256806 | C17orf100 | 1.2 | 0.24 | 3.90E-08 | 1.86E-07 |
| ENSG00000232450 | | 1.2 | 0.25 | 1.15E-07 | 5.33E-07 |
| ENSG00000243738 | RN7SL181P | 1.2 | 0.27 | 2.85E-07 | 1.27E-06 |
| ENSG00000289515 | | 1.2 | 0.31 | 5.27E-06 | 2.09E-05 |
| ENSG00000250486 | FAM218A | 1.2 | 0.33 | 1.08E-05 | 4.17E-05 |
| ENSG00000286162 | | 1.2 | 0.4 | 1.00E-04 | 3.49E-04 |
| ENSG00000257433 | RPAP3-DT | 1.2 | 0.4 | 1.06E-04 | 3.67E-04 |
| ENSG00000241769 | EOLA1-DT | 1.2 | 0.45 | 2.52E-04 | 8.35E-04 |
| ENSG00000198156 | NPIPB6 | 1.2 | 0.47 | 3.59E-04 | 1.16E-03 |
| ENSG00000206625 | RNU6-1 | 1.2 | 0.55 | 8.56E-04 | 2.63E-03 |
| ENSG00000139182 | CLSTN3 | 1.19 | 0.05 | 8.71E-117 | 6.10E-115 |
| ENSG00000127184 | COX7C | 1.19 | 0.06 | 2.53E-96 | 1.38E-94 |
| ENSG00000215808 | LINC01139 | 1.19 | 0.06 | 4.84E-93 | 2.56E-91 |
| ENSG00000234608 | MAPKAPK5-AS1 | 1.19 | 0.07 | 3.37E-65 | 1.05E-63 |
| ENSG00000206573 | THUMPD3-AS1 | 1.19 | 0.07 | 2.49E-64 | 7.64E-63 |
| ENSG00000214193 | SH3D21 | 1.19 | 0.07 | 9.01E-59 | 2.47E-57 |
| ENSG00000099875 | MKNK2 | 1.19 | 0.09 | 7.56E-42 | 1.43E-40 |
| ENSG00000088899 | LZTS3 | 1.19 | 0.12 | 4.22E-26 | 4.96E-25 |
| ENSG00000245556 | SCAMP1-AS1 | 1.19 | 0.13 | 1.57E-21 | 1.56E-20 |
| ENSG00000187860 | CCDC157 | 1.19 | 0.16 | 1.55E-15 | 1.18E-14 |
| ENSG00000221947 | XKR9 | 1.19 | 0.3 | 2.88E-06 | 1.17E-05 |
| ENSG00000271303 | SRXN1 | 1.19 | 0.35 | 2.52E-05 | 9.34E-05 |
| ENSG00000188738 | FSIP2 | 1.19 | 0.42 | 1.52E-04 | 5.18E-04 |
| ENSG00000256588 | | 1.19 | 0.91 | 8.31E-04 | 2.56E-03 |
| ENSG00000269481 | | 1.19 | 0.54 | 8.81E-04 | 2.70E-03 |
| ENSG00000135127 | BICDL1 | 1.18 | 0.07 | 4.14E-68 | 1.36E-66 |
| ENSG00000163738 | MTHFD2L | 1.18 | 0.07 | 2.51E-60 | 7.08E-59 |
| ENSG00000232533 | FAM131B-AS2 | 1.18 | 0.09 | 1.43E-38 | 2.49E-37 |
| ENSG00000205155 | PSENEN | 1.18 | 0.18 | 7.22E-12 | 4.49E-11 |
| ENSG00000235703 | EOLA2-DT | 1.18 | 0.25 | 9.34E-08 | 4.35E-07 |
| ENSG00000206337 | HCP5 | 1.18 | 0.28 | 1.10E-06 | 4.64E-06 |
| ENSG00000286138 | | 1.18 | 0.34 | 2.27E-05 | 8.48E-05 |
| ENSG00000129654 | FOXJ1 | 1.18 | 0.48 | 4.79E-04 | 1.53E-03 |
| ENSG00000279075 | | 1.18 | 0.57 | 1.13E-03 | 3.41E-03 |
| ENSG00000136628 | EPRS1 | 1.17 | 0.05 | 6.09E-128 | 5.02E-126 |
| ENSG00000174749 | FAM241A | 1.17 | 0.07 | 3.07E-59 | 8.51E-58 |
| ENSG00000182796 | TMEM198B | 1.17 | 0.11 | 3.65E-26 | 4.29E-25 |
| ENSG00000284747 | | 1.17 | 0.14 | 6.91E-19 | 6.09E-18 |
| ENSG00000176809 | LRRC37A3 | 1.17 | 0.17 | 2.01E-13 | 1.38E-12 |
| ENSG00000267080 | ASB16-AS1 | 1.17 | 0.17 | 3.61E-13 | 2.43E-12 |
| ENSG00000125965 | GDF5 | 1.17 | 0.28 | 1.41E-06 | 5.90E-06 |

| | | | | | |
|---|---|---|---|---|---|
| ENSG00000251562 | MALAT1 | 1.17 | 0.36 | 4.42E-05 | 1.60E-04 |
| ENSG00000197140 | ADAM32 | 1.17 | 0.37 | 5.40E-05 | 1.93E-04 |
| ENSG00000289428 | | 1.17 | 0.59 | 1.38E-03 | 4.09E-03 |
| ENSG00000132837 | DMGDH | 1.17 | 0.64 | 1.72E-03 | 5.03E-03 |
| ENSG00000111291 | GPRC5D | 1.17 | 0.71 | 2.04E-03 | 5.90E-03 |
| ENSG00000087074 | PPP1R15A | 1.16 | 0.05 | 4.20E-103 | 2.54E-101 |
| ENSG00000187954 | ZFTRAF1 | 1.16 | 0.11 | 1.40E-26 | 1.68E-25 |
| ENSG00000287260 | | 1.16 | 0.18 | 5.30E-12 | 3.33E-11 |
| ENSG00000130208 | APOC1 | 1.16 | 0.2 | 3.21E-10 | 1.80E-09 |
| ENSG00000255135 | EMSY-DT | 1.16 | 0.23 | 1.72E-08 | 8.46E-08 |
| ENSG00000257543 | | 1.16 | 0.29 | 2.97E-06 | 1.21E-05 |
| ENSG00000283464 | | 1.16 | 0.3 | 6.67E-06 | 2.62E-05 |
| ENSG00000258789 | | 1.16 | 0.32 | 1.59E-05 | 6.03E-05 |
| ENSG00000264769 | | 1.16 | 0.34 | 3.16E-05 | 1.16E-04 |
| ENSG00000287088 | | 1.16 | 0.39 | 1.12E-04 | 3.87E-04 |
| ENSG00000257335 | MGAM | 1.16 | 0.44 | 3.00E-04 | 9.83E-04 |
| ENSG00000181634 | TNFSF15 | 1.16 | 0.49 | 6.37E-04 | 2.00E-03 |
| ENSG00000181016 | LSMEM1 | 1.16 | 0.51 | 8.35E-04 | 2.57E-03 |
| ENSG00000258884 | LINC02321 | 1.16 | 0.77 | 1.98E-03 | 5.73E-03 |
| ENSG00000237916 | RPL37P12 | 1.16 | 0.73 | 2.11E-03 | 6.08E-03 |
| ENSG00000260144 | | 1.16 | 0.73 | 2.17E-03 | 6.23E-03 |
| ENSG00000198380 | GFPT1 | 1.15 | 0.04 | 3.08E-191 | 5.44E-189 |
| ENSG00000162733 | DDR2 | 1.15 | 0.04 | 6.43E-176 | 9.35E-174 |
| ENSG00000137216 | TMEM63B | 1.15 | 0.06 | 5.56E-97 | 3.07E-95 |
| ENSG00000117616 | RSRP1 | 1.15 | 0.07 | 8.64E-66 | 2.73E-64 |
| ENSG00000163739 | CXCL1 | 1.15 | 0.07 | 3.46E-60 | 9.74E-59 |
| ENSG00000178761 | FAM219B | 1.15 | 0.09 | 1.27E-36 | 2.09E-35 |
| ENSG00000196998 | WDR45 | 1.15 | 0.09 | 1.49E-35 | 2.39E-34 |
| ENSG00000235609 | | 1.15 | 0.11 | 3.13E-27 | 3.83E-26 |
| ENSG00000197714 | ZNF460 | 1.15 | 0.21 | 3.74E-09 | 1.93E-08 |
| ENSG00000207721 | MIR186 | 1.15 | 0.23 | 2.43E-08 | 1.19E-07 |
| ENSG00000273002 | ARHGEF2-AS2 | 1.15 | 0.27 | 1.03E-06 | 4.35E-06 |
| ENSG00000121410 | A1BG | 1.15 | 0.3 | 4.68E-06 | 1.87E-05 |
| ENSG00000278611 | ZNF426-DT | 1.15 | 0.35 | 4.55E-05 | 1.64E-04 |
| ENSG00000278467 | | 1.15 | 0.36 | 5.84E-05 | 2.09E-04 |
| ENSG00000249249 | TICAM2-AS1 | 1.15 | 0.36 | 6.37E-05 | 2.27E-04 |
| ENSG00000261270 | | 1.15 | 0.44 | 3.56E-04 | 1.16E-03 |
| ENSG00000232104 | RFX3-DT | 1.15 | 0.48 | 6.74E-04 | 2.11E-03 |
| ENSG00000279057 | | 1.15 | 0.59 | 1.61E-03 | 4.73E-03 |
| ENSG00000233155 | HMGA1P8 | 1.15 | 0.71 | 2.23E-03 | 6.40E-03 |
| ENSG00000196449 | YRDC | 1.14 | 0.04 | 1.16E-154 | 1.29E-152 |
| ENSG00000105855 | ITGB8 | 1.14 | 0.06 | 2.49E-71 | 8.77E-70 |
| ENSG00000177225 | GATD1 | 1.14 | 0.07 | 2.02E-59 | 5.63E-58 |
| ENSG00000177169 | ULK1 | 1.14 | 0.08 | 7.60E-43 | 1.47E-41 |
| ENSG00000171443 | ZNF524 | 1.14 | 0.17 | 3.59E-12 | 2.28E-11 |
| ENSG00000268182 | SMIM17 | 1.14 | 0.19 | 6.66E-11 | 3.90E-10 |
| ENSG00000269696 | ZNF470-DT | 1.14 | 0.21 | 3.16E-09 | 1.64E-08 |
| ENSG00000164849 | GPR146 | 1.14 | 0.24 | 1.13E-07 | 5.21E-07 |
| ENSG00000198468 | FLVCR1-DT | 1.14 | 0.26 | 8.11E-07 | 3.47E-06 |
| ENSG00000251279 | SMIM15-AS1 | 1.14 | 0.28 | 2.35E-06 | 9.61E-06 |

| | | | | | |
|---|---|---|---|---|---|
| ENSG00000286834 | | 1.14 | 0.3 | 6.01E-06 | 2.38E-05 |
| ENSG00000284308 | C2orf81 | 1.14 | 0.32 | 1.64E-05 | 6.21E-05 |
| ENSG00000179862 | CITED4 | 1.14 | 0.33 | 2.54E-05 | 9.40E-05 |
| ENSG00000133636 | NTS | 1.14 | 0.47 | 5.97E-04 | 1.88E-03 |
| ENSG00000289206 | | 1.14 | 0.53 | 1.18E-03 | 3.54E-03 |
| ENSG00000290808 | | 1.14 | 0.57 | 1.53E-03 | 4.50E-03 |
| ENSG00000272523 | LINC01023 | 1.14 | 0.62 | 1.98E-03 | 5.74E-03 |
| ENSG00000133935 | ERG28 | 1.13 | 0.05 | 1.04E-134 | 9.31E-133 |
| ENSG00000160216 | AGPAT3 | 1.13 | 0.06 | 1.11E-92 | 5.80E-91 |
| ENSG00000198598 | MMP17 | 1.13 | 0.11 | 1.83E-27 | 2.25E-26 |
| ENSG00000223745 | CCDC18-AS1 | 1.13 | 0.15 | 1.84E-15 | 1.39E-14 |
| ENSG00000102981 | PARD6A | 1.13 | 0.21 | 2.28E-09 | 1.19E-08 |
| ENSG00000135951 | TSGA10 | 1.13 | 0.23 | 5.32E-08 | 2.52E-07 |
| ENSG00000273472 | | 1.13 | 0.23 | 7.99E-08 | 3.74E-07 |
| ENSG00000178201 | VN1R1 | 1.13 | 0.28 | 2.88E-06 | 1.17E-05 |
| ENSG00000259212 | | 1.13 | 0.4 | 2.28E-04 | 7.60E-04 |
| ENSG00000251131 | | 1.13 | 0.44 | 4.20E-04 | 1.35E-03 |
| ENSG00000269533 | | 1.13 | 0.53 | 1.16E-03 | 3.49E-03 |
| ENSG00000288056 | | 1.13 | 0.53 | 1.25E-03 | 3.74E-03 |
| ENSG00000239462 | | 1.13 | 0.54 | 1.30E-03 | 3.88E-03 |
| ENSG00000164305 | CASP3 | 1.12 | 0.06 | 2.93E-88 | 1.39E-86 |
| ENSG00000147400 | CETN2 | 1.12 | 0.06 | 1.34E-76 | 5.25E-75 |
| ENSG00000182648 | RNF32-DT | 1.12 | 0.09 | 4.15E-40 | 7.43E-39 |
| ENSG00000278133 | | 1.12 | 0.11 | 1.17E-27 | 1.45E-26 |
| ENSG00000162066 | AMDHD2 | 1.12 | 0.13 | 2.88E-20 | 2.71E-19 |
| ENSG00000001630 | CYP51A1 | 1.12 | 0.13 | 2.95E-19 | 2.66E-18 |
| ENSG00000165124 | SVEP1 | 1.12 | 0.14 | 1.01E-16 | 8.14E-16 |
| ENSG00000183397 | TEKTIP1 | 1.12 | 0.14 | 1.54E-16 | 1.23E-15 |
| ENSG00000116675 | DNAJC6 | 1.12 | 0.18 | 5.13E-11 | 3.03E-10 |
| ENSG00000175155 | YPEL2 | 1.12 | 0.18 | 7.53E-11 | 4.40E-10 |
| ENSG00000267053 | | 1.12 | 0.27 | 2.13E-06 | 8.76E-06 |
| ENSG00000260978 | | 1.12 | 0.38 | 1.32E-04 | 4.53E-04 |
| ENSG00000234975 | FTH1P2 | 1.12 | 0.39 | 1.69E-04 | 5.71E-04 |
| ENSG00000249125 | | 1.12 | 0.43 | 3.86E-04 | 1.25E-03 |
| ENSG00000279259 | | 1.12 | 0.43 | 3.93E-04 | 1.27E-03 |
| ENSG00000201616 | RNU1-91P | 1.12 | 0.92 | 9.18E-04 | 2.81E-03 |
| ENSG00000285770 | | 1.12 | 0.5 | 9.84E-04 | 2.99E-03 |
| ENSG00000259172 | SNRPA1-DT | 1.12 | 0.64 | 2.33E-03 | 6.69E-03 |
| ENSG00000267892 | | 1.12 | 0.69 | 2.59E-03 | 7.36E-03 |
| ENSG00000181019 | NQO1 | 1.11 | 0.05 | 5.03E-129 | 4.24E-127 |
| ENSG00000197903 | H2BC12 | 1.11 | 0.07 | 7.11E-64 | 2.14E-62 |
| ENSG00000171451 | DSEL | 1.11 | 0.09 | 1.90E-38 | 3.30E-37 |
| ENSG00000284753 | EEF1AKMT4 | 1.11 | 0.19 | 7.68E-10 | 4.18E-09 |
| ENSG00000260563 | | 1.11 | 0.21 | 3.76E-09 | 1.94E-08 |
| ENSG00000139597 | N4BP2L1 | 1.11 | 0.26 | 9.26E-07 | 3.94E-06 |
| ENSG00000184389 | A3GALT2 | 1.11 | 0.4 | 2.65E-04 | 8.75E-04 |
| ENSG00000167077 | MEI1 | 1.11 | 0.42 | 3.41E-04 | 1.11E-03 |
| ENSG00000237927 | | 1.11 | 0.53 | 1.37E-03 | 4.08E-03 |
| ENSG00000262140 | | 1.11 | 0.85 | 1.70E-03 | 4.98E-03 |
| ENSG00000251050 | RBX1P2 | 1.11 | 0.56 | 1.75E-03 | 5.12E-03 |

| | | | | | |
|---|---|---|---|---|---|
| ENSG00000163083 | INHBB | 1.11 | 0.79 | 2.47E-03 | 7.05E-03 |
| ENSG00000263177 | MTND1P8 | 1.11 | 0.77 | 2.60E-03 | 7.39E-03 |
| ENSG00000207359 | RNU6-925P | 1.11 | 0.66 | 2.62E-03 | 7.45E-03 |
| ENSG00000261396 | | 1.11 | 0.72 | 2.79E-03 | 7.89E-03 |
| ENSG00000144802 | NFKBIZ | 1.1 | 0.05 | 1.65E-101 | 9.75E-100 |
| ENSG00000198855 | FICD | 1.1 | 0.07 | 6.14E-50 | 1.37E-48 |
| ENSG00000139636 | LMBR1L | 1.1 | 0.1 | 3.47E-31 | 4.85E-30 |
| ENSG00000229320 | KRT8P12 | 1.1 | 0.11 | 4.42E-25 | 5.01E-24 |
| ENSG00000246067 | RAB30-DT | 1.1 | 0.12 | 5.48E-22 | 5.52E-21 |
| ENSG00000111678 | C12orf57 | 1.1 | 0.12 | 1.30E-21 | 1.29E-20 |
| ENSG00000039139 | DNAH5 | 1.1 | 0.12 | 3.67E-20 | 3.44E-19 |
| ENSG00000116701 | NCF2 | 1.1 | 0.13 | 9.40E-19 | 8.25E-18 |
| ENSG00000279495 | | 1.1 | 0.14 | 5.90E-16 | 4.58E-15 |
| ENSG00000107872 | FBXL15 | 1.1 | 0.14 | 7.14E-16 | 5.51E-15 |
| ENSG00000273619 | | 1.1 | 0.15 | 5.13E-14 | 3.62E-13 |
| ENSG00000174501 | ANKRD36C | 1.1 | 0.16 | 7.54E-13 | 4.99E-12 |
| ENSG00000188620 | HMX3 | 1.1 | 0.19 | 3.70E-10 | 2.06E-09 |
| ENSG00000174564 | IL20RB | 1.1 | 0.25 | 6.37E-07 | 2.76E-06 |
| ENSG00000267871 | ZNF460-AS1 | 1.1 | 0.31 | 2.21E-05 | 8.27E-05 |
| ENSG00000162032 | SPSB3 | 1.1 | 0.36 | 9.35E-05 | 3.26E-04 |
| ENSG00000100429 | HDAC10 | 1.1 | 0.37 | 1.56E-04 | 5.28E-04 |
| ENSG00000251417 | | 1.1 | 0.38 | 1.95E-04 | 6.53E-04 |
| ENSG00000158815 | FGF17 | 1.1 | 0.44 | 5.25E-04 | 1.67E-03 |
| ENSG00000254872 | LINC02688 | 1.1 | 0.63 | 2.63E-03 | 7.47E-03 |
| ENSG00000280069 | | 1.1 | 0.66 | 2.88E-03 | 8.12E-03 |
| ENSG00000158470 | B4GALT5 | 1.09 | 0.03 | 3.36E-249 | 1.05E-246 |
| ENSG00000113083 | LOX | 1.09 | 0.04 | 4.29E-187 | 7.24E-185 |
| ENSG00000104635 | SLC39A14 | 1.09 | 0.04 | 2.54E-181 | 3.98E-179 |
| ENSG00000131711 | MAP1B | 1.09 | 0.04 | 2.72E-136 | 2.50E-134 |
| ENSG00000131018 | SYNE1 | 1.09 | 0.05 | 1.19E-99 | 6.90E-98 |
| ENSG00000197063 | MAFG | 1.09 | 0.07 | 3.45E-64 | 1.06E-62 |
| ENSG00000238129 | | 1.09 | 0.07 | 1.15E-55 | 2.95E-54 |
| ENSG00000019991 | HGF | 1.09 | 0.09 | 5.29E-32 | 7.57E-31 |
| ENSG00000160972 | PPP1R16A | 1.09 | 0.1 | 4.30E-30 | 5.79E-29 |
| ENSG00000080298 | RFX3 | 1.09 | 0.11 | 3.43E-25 | 3.89E-24 |
| ENSG00000109927 | TECTA | 1.09 | 0.13 | 8.64E-19 | 7.60E-18 |
| ENSG00000280077 | | 1.09 | 0.14 | 7.95E-17 | 6.43E-16 |
| ENSG00000166046 | TCP11L2 | 1.09 | 0.19 | 1.48E-09 | 7.87E-09 |
| ENSG00000291015 | | 1.09 | 0.26 | 1.78E-06 | 7.38E-06 |
| ENSG00000279520 | | 1.09 | 0.29 | 8.76E-06 | 3.40E-05 |
| ENSG00000260352 | SMG1-DT | 1.09 | 0.48 | 1.00E-03 | 3.05E-03 |
| ENSG00000253180 | | 1.09 | 0.89 | 1.39E-03 | 4.12E-03 |
| ENSG00000189348 | FAM90A27P | 1.09 | 0.85 | 1.80E-03 | 5.25E-03 |
| ENSG00000208028 | MIR616 | 1.09 | 0.56 | 2.05E-03 | 5.92E-03 |
| ENSG00000285382 | | 1.09 | 0.77 | 2.83E-03 | 7.99E-03 |
| ENSG00000280212 | | 1.09 | 0.76 | 2.94E-03 | 8.26E-03 |
| ENSG00000120053 | GOT1 | 1.08 | 0.05 | 1.80E-126 | 1.44E-124 |
| ENSG00000260032 | NORAD | 1.08 | 0.05 | 2.39E-124 | 1.85E-122 |
| ENSG00000023287 | RB1CC1 | 1.08 | 0.05 | 1.67E-105 | 1.04E-103 |
| ENSG00000131069 | ACSS2 | 1.08 | 0.06 | 6.83E-75 | 2.57E-73 |

| | | | | | |
|---|---|---|---|---|---|
| ENSG00000227115 | LINC01630 | 1.08 | 0.06 | 2.09E-64 | 6.43E-63 |
| ENSG00000124702 | KLHDC3 | 1.08 | 0.08 | 2.57E-47 | 5.44E-46 |
| ENSG00000130733 | YIPF2 | 1.08 | 0.08 | 6.76E-47 | 1.42E-45 |
| ENSG00000114544 | SLC41A3 | 1.08 | 0.08 | 5.78E-39 | 1.01E-37 |
| ENSG00000204348 | DXO | 1.08 | 0.09 | 5.25E-35 | 8.22E-34 |
| ENSG00000168936 | TMEM129 | 1.08 | 0.09 | 3.51E-32 | 5.05E-31 |
| ENSG00000115649 | CNPPD1 | 1.08 | 0.1 | 1.11E-28 | 1.42E-27 |
| ENSG00000155367 | PPM1J | 1.08 | 0.16 | 1.49E-12 | 9.65E-12 |
| ENSG00000256436 | TAS2R31 | 1.08 | 0.27 | 2.70E-06 | 1.10E-05 |
| ENSG00000284879 | NCAL1 | 1.08 | 0.29 | 1.17E-05 | 4.49E-05 |
| ENSG00000251580 | LINC02482 | 1.08 | 0.32 | 4.36E-05 | 1.58E-04 |
| ENSG00000279048 | | 1.08 | 0.36 | 1.58E-04 | 5.34E-04 |
| ENSG00000260618 | | 1.08 | 0.88 | 1.54E-03 | 4.53E-03 |
| ENSG00000274444 | | 1.08 | 0.51 | 1.58E-03 | 4.66E-03 |
| ENSG00000205869 | KRTAP5-1 | 1.08 | 0.86 | 1.75E-03 | 5.13E-03 |
| ENSG00000288990 | | 1.08 | 0.85 | 1.98E-03 | 5.73E-03 |
| ENSG00000232065 | LINC01063 | 1.08 | 0.56 | 2.22E-03 | 6.39E-03 |
| ENSG00000235586 | | 1.08 | 0.57 | 2.33E-03 | 6.67E-03 |
| ENSG00000120137 | PANK3 | 1.07 | 0.05 | 1.32E-105 | 8.23E-104 |
| ENSG00000205269 | TMEM170B | 1.07 | 0.06 | 4.43E-75 | 1.68E-73 |
| ENSG00000107798 | LIPA | 1.07 | 0.06 | 3.49E-66 | 1.11E-64 |
| ENSG00000157693 | TMEM268 | 1.07 | 0.07 | 2.77E-58 | 7.48E-57 |
| ENSG00000152256 | PDK1 | 1.07 | 0.07 | 2.66E-56 | 6.90E-55 |
| ENSG00000165030 | NFIL3 | 1.07 | 0.07 | 1.00E-47 | 2.13E-46 |
| ENSG00000102096 | PIM2 | 1.07 | 0.09 | 3.89E-33 | 5.74E-32 |
| ENSG00000099326 | MZF1 | 1.07 | 0.09 | 1.24E-30 | 1.70E-29 |
| ENSG00000069966 | GNB5 | 1.07 | 0.1 | 4.43E-30 | 5.96E-29 |
| ENSG00000225663 | MCRIP1 | 1.07 | 0.12 | 1.13E-20 | 1.08E-19 |
| ENSG00000158792 | SPATA2L | 1.07 | 0.13 | 2.43E-18 | 2.09E-17 |
| ENSG00000054356 | PTPRN | 1.07 | 0.13 | 4.44E-18 | 3.78E-17 |
| ENSG00000240498 | CDKN2B-AS1 | 1.07 | 0.16 | 5.33E-13 | 3.56E-12 |
| ENSG00000227775 | | 1.07 | 0.16 | 2.22E-12 | 1.43E-11 |
| ENSG00000053524 | MCF2L2 | 1.07 | 0.25 | 1.55E-06 | 6.48E-06 |
| ENSG00000224790 | HLCS-AS1 | 1.07 | 0.26 | 2.24E-06 | 9.20E-06 |
| ENSG00000250746 | | 1.07 | 0.56 | 2.37E-03 | 6.79E-03 |
| ENSG00000238140 | | 1.07 | 0.75 | 3.26E-03 | 9.08E-03 |
| ENSG00000164463 | CREBRF | 1.06 | 0.08 | 3.49E-37 | 5.81E-36 |
| ENSG00000288851 | | 1.06 | 0.11 | 1.48E-22 | 1.52E-21 |
| ENSG00000168273 | UQCC5 | 1.06 | 0.12 | 1.99E-21 | 1.96E-20 |
| ENSG00000151917 | BEND6 | 1.06 | 0.12 | 6.85E-21 | 6.60E-20 |
| ENSG00000213420 | GPC2 | 1.06 | 0.13 | 2.75E-18 | 2.36E-17 |
| ENSG00000040608 | RTN4R | 1.06 | 0.16 | 1.09E-12 | 7.16E-12 |
| ENSG00000247934 | | 1.06 | 0.17 | 1.69E-11 | 1.03E-10 |
| ENSG00000268575 | | 1.06 | 0.2 | 9.89E-09 | 4.96E-08 |
| ENSG00000268006 | PTOV1-AS1 | 1.06 | 0.22 | 6.27E-08 | 2.95E-07 |
| ENSG00000290126 | | 1.06 | 0.23 | 1.49E-07 | 6.83E-07 |
| ENSG00000135116 | HRK | 1.06 | 0.23 | 4.22E-07 | 1.86E-06 |
| ENSG00000290791 | | 1.06 | 0.24 | 6.14E-07 | 2.66E-06 |
| ENSG00000290086 | | 1.06 | 0.39 | 3.32E-04 | 1.08E-03 |
| ENSG00000279140 | | 1.06 | 0.43 | 7.17E-04 | 2.23E-03 |

| | | | | | |
|---|---|---|---|---|---|
| ENSG00000232859 | LYRM9 | 1.06 | 0.46 | 1.02E-03 | 3.09E-03 |
| ENSG00000271856 | LINC01215 | 1.06 | 0.51 | 1.65E-03 | 4.83E-03 |
| ENSG00000274578 | | 1.06 | 0.52 | 1.86E-03 | 5.42E-03 |
| ENSG00000272597 | | 1.06 | 0.85 | 2.07E-03 | 5.98E-03 |
| ENSG00000162755 | KLHDC9 | 1.06 | 0.61 | 3.21E-03 | 8.97E-03 |
| ENSG00000033627 | ATP6V0A1 | 1.05 | 0.04 | 2.78E-156 | 3.17E-154 |
| ENSG00000164830 | OXR1 | 1.05 | 0.05 | 1.06E-87 | 4.94E-86 |
| ENSG00000141425 | RPRD1A | 1.05 | 0.05 | 4.77E-87 | 2.18E-85 |
| ENSG00000185658 | BRWD1 | 1.05 | 0.05 | 3.30E-83 | 1.46E-81 |
| ENSG00000120738 | EGR1 | 1.05 | 0.06 | 5.64E-82 | 2.40E-80 |
| ENSG00000135241 | PNPLA8 | 1.05 | 0.07 | 3.37E-55 | 8.60E-54 |
| ENSG00000145016 | RUBCN | 1.05 | 0.07 | 2.84E-52 | 6.74E-51 |
| ENSG00000138036 | DYNC2LI1 | 1.05 | 0.07 | 2.35E-46 | 4.87E-45 |
| ENSG00000186222 | BLOC1S4 | 1.05 | 0.08 | 4.04E-43 | 7.85E-42 |
| ENSG00000167797 | CDK2AP2 | 1.05 | 0.09 | 2.44E-32 | 3.52E-31 |
| ENSG00000246705 | H2AJ | 1.05 | 0.09 | 1.71E-30 | 2.35E-29 |
| ENSG00000139200 | PIANP | 1.05 | 0.11 | 2.44E-23 | 2.58E-22 |
| ENSG00000158373 | H2BC5 | 1.05 | 0.11 | 2.30E-22 | 2.35E-21 |
| ENSG00000290032 | | 1.05 | 0.12 | 6.87E-21 | 6.62E-20 |
| ENSG00000291117 | | 1.05 | 0.14 | 2.49E-14 | 1.79E-13 |
| ENSG00000290762 | | 1.05 | 0.18 | 5.13E-10 | 2.83E-09 |
| ENSG00000172508 | CARNS1 | 1.05 | 0.29 | 2.25E-05 | 8.38E-05 |
| ENSG00000160111 | CPAMD8 | 1.05 | 0.36 | 2.01E-04 | 6.73E-04 |
| ENSG00000102931 | ARL2BP | 1.05 | 0.4 | 5.01E-04 | 1.60E-03 |
| ENSG00000275719 | | 1.05 | 0.7 | 3.85E-03 | 1.06E-02 |
| ENSG00000142188 | TMEM50B | 1.04 | 0.06 | 1.00E-75 | 3.85E-74 |
| ENSG00000152520 | PAN3 | 1.04 | 0.06 | 2.38E-59 | 6.62E-58 |
| ENSG00000150593 | PDCD4 | 1.04 | 0.06 | 3.03E-59 | 8.43E-58 |
| ENSG00000182195 | LDOC1 | 1.04 | 0.08 | 7.27E-43 | 1.41E-41 |
| ENSG00000090971 | NAT14 | 1.04 | 0.09 | 4.08E-33 | 6.02E-32 |
| ENSG00000164542 | MATCAP2 | 1.04 | 0.09 | 9.59E-33 | 1.40E-31 |
| ENSG00000184949 | FAM227A | 1.04 | 0.1 | 3.96E-26 | 4.64E-25 |
| ENSG00000171159 | BBLN | 1.04 | 0.11 | 2.38E-23 | 2.52E-22 |
| ENSG00000112406 | HECA | 1.04 | 0.11 | 6.50E-21 | 6.27E-20 |
| ENSG00000204618 | RNF39 | 1.04 | 0.13 | 7.44E-16 | 5.73E-15 |
| ENSG00000289004 | | 1.04 | 0.29 | 2.17E-05 | 8.10E-05 |
| ENSG00000101222 | SPEF1 | 1.04 | 0.3 | 2.85E-05 | 1.05E-04 |
| ENSG00000124440 | HIF3A | 1.04 | 0.31 | 4.02E-05 | 1.46E-04 |
| ENSG00000215769 | ARHGAP27P1-BPTFP1 | 1.04 | 0.36 | 2.26E-04 | 7.54E-04 |
| ENSG00000289855 | | 1.04 | 0.84 | 2.49E-03 | 7.10E-03 |
| ENSG00000234036 | TXNP6 | 1.04 | 0.61 | 3.51E-03 | 9.71E-03 |
| ENSG00000151893 | CACUL1 | 1.03 | 0.04 | 9.48E-136 | 8.62E-134 |
| ENSG00000067082 | KLF6 | 1.03 | 0.04 | 7.59E-122 | 5.76E-120 |
| ENSG00000174010 | KLHL15 | 1.03 | 0.06 | 1.71E-68 | 5.66E-67 |
| ENSG00000178607 | ERN1 | 1.03 | 0.07 | 5.92E-55 | 1.51E-53 |
| ENSG00000168887 | C2orf68 | 1.03 | 0.07 | 3.49E-51 | 8.08E-50 |
| ENSG00000166971 | AKTIP | 1.03 | 0.09 | 2.22E-33 | 3.31E-32 |
| ENSG00000260852 | FBXL19-AS1 | 1.03 | 0.09 | 3.34E-33 | 4.95E-32 |
| ENSG00000143344 | RGL1 | 1.03 | 0.09 | 1.31E-29 | 1.73E-28 |
| ENSG00000261188 | TFIP11-DT | 1.03 | 0.15 | 1.37E-12 | 8.93E-12 |

| | | | | | |
|---|---|---|---|---|---|
| ENSG00000141933 | TPGS1 | 1.03 | 0.16 | 1.37E-11 | 8.36E-11 |
| ENSG00000155849 | ELMO1 | 1.03 | 0.18 | 1.17E-09 | 6.31E-09 |
| ENSG00000090238 | YPEL3 | 1.03 | 0.19 | 2.38E-09 | 1.25E-08 |
| ENSG00000274020 | LINC01138 | 1.03 | 0.25 | 2.56E-06 | 1.05E-05 |
| ENSG00000229124 | VIM-AS1 | 1.03 | 0.27 | 1.03E-05 | 3.99E-05 |
| ENSG00000099769 | IGFALS | 1.03 | 0.32 | 7.37E-05 | 2.60E-04 |
| ENSG00000196890 | H2BC26 | 1.03 | 0.35 | 1.70E-04 | 5.74E-04 |
| ENSG00000135525 | MAP7 | 1.03 | 0.38 | 4.36E-04 | 1.40E-03 |
| ENSG00000176046 | NUPR1 | 1.03 | 0.41 | 6.18E-04 | 1.94E-03 |
| ENSG00000120337 | TNFSF18 | 1.03 | 0.62 | 3.95E-03 | 1.08E-02 |
| ENSG00000259024 | TVP23C-CDRT4 | 1.03 | 0.72 | 4.21E-03 | 1.15E-02 |
| ENSG00000120889 | TNFRSF10B | 1.02 | 0.03 | 3.42E-226 | 8.78E-224 |
| ENSG00000141458 | NPC1 | 1.02 | 0.04 | 1.58E-173 | 2.21E-171 |
| ENSG00000173221 | GLRX | 1.02 | 0.05 | 3.74E-99 | 2.15E-97 |
| ENSG00000032219 | ARID4A | 1.02 | 0.08 | 1.25E-39 | 2.23E-38 |
| ENSG00000162236 | STX5 | 1.02 | 0.09 | 2.36E-33 | 3.51E-32 |
| ENSG00000171222 | SCAND1 | 1.02 | 0.1 | 5.45E-27 | 6.61E-26 |
| ENSG00000273015 | LINC00938 | 1.02 | 0.1 | 1.09E-24 | 1.21E-23 |
| ENSG00000181274 | FRAT2 | 1.02 | 0.14 | 1.88E-14 | 1.36E-13 |
| ENSG00000147174 | GCNA | 1.02 | 0.16 | 6.47E-12 | 4.03E-11 |
| ENSG00000159182 | PRAC1 | 1.02 | 0.19 | 6.08E-09 | 3.09E-08 |
| ENSG00000124635 | H2BC11 | 1.02 | 0.2 | 1.19E-08 | 5.93E-08 |
| ENSG00000254554 | ADM-DT | 1.02 | 0.23 | 4.36E-07 | 1.92E-06 |
| ENSG00000289440 | | 1.02 | 0.24 | 1.68E-06 | 6.99E-06 |
| ENSG00000161664 | ASB16 | 1.02 | 0.33 | 1.31E-04 | 4.50E-04 |
| ENSG00000289370 | | 1.02 | 0.42 | 9.07E-04 | 2.78E-03 |
| ENSG00000247498 | GPRC5D-AS1 | 1.02 | 0.43 | 1.03E-03 | 3.11E-03 |
| ENSG00000231663 | COA6-AS1 | 1.02 | 0.48 | 1.80E-03 | 5.24E-03 |
| ENSG00000225434 | LINC01504 | 1.02 | 0.61 | 3.94E-03 | 1.08E-02 |
| ENSG00000132680 | KHDC4 | 1.01 | 0.04 | 4.47E-125 | 3.49E-123 |
| ENSG00000168917 | SLC35G2 | 1.01 | 0.05 | 2.50E-81 | 1.06E-79 |
| ENSG00000119729 | RHOQ | 1.01 | 0.06 | 1.68E-66 | 5.35E-65 |
| ENSG00000113739 | STC2 | 1.01 | 0.06 | 2.19E-64 | 6.74E-63 |
| ENSG00000227195 | MIR663AHG | 1.01 | 0.09 | 1.32E-29 | 1.75E-28 |
| ENSG00000154639 | CXADR | 1.01 | 0.1 | 6.78E-25 | 7.63E-24 |
| ENSG00000099385 | BCL7C | 1.01 | 0.11 | 2.05E-21 | 2.02E-20 |
| ENSG00000249898 | MCPH1-AS1 | 1.01 | 0.12 | 2.93E-19 | 2.64E-18 |
| ENSG00000059915 | PSD | 1.01 | 0.12 | 4.78E-18 | 4.07E-17 |
| ENSG00000247092 | SNHG10 | 1.01 | 0.14 | 6.68E-14 | 4.68E-13 |
| ENSG00000271780 | | 1.01 | 0.27 | 1.41E-05 | 5.38E-05 |
| ENSG00000237491 | LINC01409 | 1.01 | 0.29 | 4.16E-05 | 1.51E-04 |
| ENSG00000076344 | RGS11 | 1.01 | 0.34 | 2.05E-04 | 6.85E-04 |
| ENSG00000120664 | SPART-AS1 | 1.01 | 0.36 | 2.88E-04 | 9.45E-04 |
| ENSG00000196787 | H2AC11 | 1.01 | 0.39 | 5.51E-04 | 1.75E-03 |
| ENSG00000275630 | | 1.01 | 0.39 | 5.61E-04 | 1.78E-03 |
| ENSG00000010379 | SLC6A13 | 1.01 | 0.42 | 9.80E-04 | 2.98E-03 |
| ENSG00000184860 | SDR42E1 | 1.01 | 0.42 | 1.02E-03 | 3.09E-03 |
| ENSG00000230658 | KLHL7-DT | 1.01 | 0.52 | 2.68E-03 | 7.60E-03 |
| ENSG00000267649 | | 1.01 | 0.54 | 3.01E-03 | 8.46E-03 |
| ENSG00000264235 | MYL12-AS1 | 1.01 | 0.79 | 3.64E-03 | 1.01E-02 |

| | | | | | |
|---|---|---|---|---|---|
| ENSG00000155307 | SAMSN1 | 1.01 | 0.78 | 3.93E-03 | 1.08E-02 |
| ENSG00000136868 | SLC31A1 | 1 | 0.04 | 3.63E-129 | 3.07E-127 |
| ENSG00000109084 | TMEM97 | 1 | 0.04 | 1.49E-113 | 1.01E-111 |
| ENSG00000003436 | TFPI | 1 | 0.05 | 2.15E-102 | 1.28E-100 |
| ENSG00000111229 | ARPC3 | 1 | 0.05 | 4.41E-90 | 2.15E-88 |
| ENSG00000068912 | ERLEC1 | 1 | 0.05 | 2.77E-79 | 1.13E-77 |
| ENSG00000112679 | DUSP22 | 1 | 0.09 | 2.64E-28 | 3.33E-27 |
| ENSG00000145949 | MYLK4 | 1 | 0.11 | 1.33E-20 | 1.26E-19 |
| ENSG00000167702 | KIFC2 | 1 | 0.12 | 1.88E-18 | 1.63E-17 |
| ENSG00000164181 | ELOVL7 | 1 | 0.17 | 4.64E-10 | 2.57E-09 |
| ENSG00000274423 | SEC22B2P | 1 | 0.23 | 8.48E-07 | 3.63E-06 |
| ENSG00000171786 | NHLH1 | 1 | 0.23 | 1.02E-06 | 4.32E-06 |
| ENSG00000108839 | ALOX12 | 1 | 0.24 | 1.62E-06 | 6.75E-06 |
| ENSG00000136244 | IL6 | 1 | 0.25 | 3.14E-06 | 1.27E-05 |
| ENSG00000205041 | | 1 | 0.29 | 3.84E-05 | 1.40E-04 |
| ENSG00000279086 | | 1 | 0.3 | 5.62E-05 | 2.01E-04 |
| ENSG00000101203 | COL20A1 | 1 | 0.3 | 6.12E-05 | 2.18E-04 |
| ENSG00000125804 | FAM182A | 1 | 0.4 | 7.44E-04 | 2.31E-03 |
| ENSG00000253667 | | 1 | 0.45 | 1.66E-03 | 4.86E-03 |
| ENSG00000288720 | | 1 | 0.48 | 2.05E-03 | 5.91E-03 |
| ENSG00000288970 | | 1 | 0.85 | 2.40E-03 | 6.85E-03 |
| ENSG00000121594 | CD80 | 1 | 0.59 | 4.11E-03 | 1.12E-02 |
| ENSG00000235831 | BHLHE40-AS1 | 1 | 0.66 | 4.99E-03 | 1.34E-02 |
| ENSG00000112308 | C6orf62 | 0.99 | 0.05 | 7.15E-92 | 3.66E-90 |
| ENSG00000141446 | ESCO1 | 0.99 | 0.06 | 4.74E-56 | 1.23E-54 |
| ENSG00000204130 | RUFY2 | 0.99 | 0.07 | 1.19E-48 | 2.57E-47 |
| ENSG00000244879 | GABPB1-AS1 | 0.99 | 0.07 | 7.89E-43 | 1.53E-41 |
| ENSG00000125826 | RBCK1 | 0.99 | 0.07 | 4.05E-42 | 7.71E-41 |
| ENSG00000237441 | RGL2 | 0.99 | 0.08 | 1.15E-37 | 1.95E-36 |
| ENSG00000162545 | CAMK2N1 | 0.99 | 0.1 | 6.38E-24 | 6.94E-23 |
| ENSG00000135211 | TMEM60 | 0.99 | 0.1 | 8.04E-23 | 8.36E-22 |
| ENSG00000277283 | RAB35-AS1 | 0.99 | 0.12 | 1.71E-16 | 1.36E-15 |
| ENSG00000291043 | GLUD1P3 | 0.99 | 0.16 | 5.27E-11 | 3.11E-10 |
| ENSG00000212664 | RPL17P39 | 0.99 | 0.26 | 7.46E-06 | 2.92E-05 |
| ENSG00000246339 | EXTL3-AS1 | 0.99 | 0.26 | 9.67E-06 | 3.75E-05 |
| ENSG00000181798 | LINC00471 | 0.99 | 0.26 | 1.19E-05 | 4.56E-05 |
| ENSG00000203392 | | 0.99 | 0.33 | 1.83E-04 | 6.15E-04 |
| ENSG00000267938 | EIF1P6 | 0.99 | 0.34 | 2.23E-04 | 7.43E-04 |
| ENSG00000273240 | | 0.99 | 0.41 | 9.92E-04 | 3.02E-03 |
| ENSG00000226067 | LINC00623 | 0.99 | 0.42 | 1.22E-03 | 3.65E-03 |
| ENSG00000261386 | | 0.99 | 0.48 | 2.40E-03 | 6.85E-03 |
| ENSG00000249476 | | 0.99 | 0.57 | 4.01E-03 | 1.10E-02 |
| ENSG00000163606 | CD200R1 | 0.99 | 0.71 | 5.39E-03 | 1.44E-02 |
| ENSG00000132824 | SERINC3 | 0.98 | 0.04 | 8.18E-121 | 6.10E-119 |
| ENSG00000151414 | NEK7 | 0.98 | 0.05 | 8.74E-88 | 4.08E-86 |
| ENSG00000196139 | AKR1C3 | 0.98 | 0.07 | 9.48E-49 | 2.06E-47 |
| ENSG00000120727 | PAIP2 | 0.98 | 0.08 | 5.87E-38 | 1.01E-36 |
| ENSG00000242802 | AP5Z1 | 0.98 | 0.08 | 2.74E-35 | 4.34E-34 |
| ENSG00000276399 | FLJ36000 | 0.98 | 0.09 | 3.51E-31 | 4.91E-30 |
| ENSG00000128016 | ZFP36 | 0.98 | 0.11 | 1.34E-19 | 1.23E-18 |

| | | | | | |
|---|---|---|---|---|---|
| ENSG00000119138 | KLF9 | 0.98 | 0.12 | 1.59E-17 | 1.33E-16 |
| ENSG00000112182 | BACH2 | 0.98 | 0.12 | 6.90E-17 | 5.61E-16 |
| ENSG00000250790 | | 0.98 | 0.2 | 8.05E-08 | 3.76E-07 |
| ENSG00000223573 | TINCR | 0.98 | 0.24 | 3.21E-06 | 1.30E-05 |
| ENSG00000265982 | | 0.98 | 0.25 | 8.20E-06 | 3.20E-05 |
| ENSG00000257599 | OVCH1-AS1 | 0.98 | 0.26 | 1.31E-05 | 5.01E-05 |
| ENSG00000141497 | ZMYND15 | 0.98 | 0.33 | 2.16E-04 | 7.20E-04 |
| ENSG00000288756 | | 0.98 | 0.44 | 1.55E-03 | 4.56E-03 |
| ENSG00000170296 | GABARAP | 0.98 | 0.52 | 3.52E-03 | 9.74E-03 |
| ENSG00000240922 | LSAMP-AS1 | 0.98 | 0.68 | 5.68E-03 | 1.51E-02 |
| ENSG00000196305 | IARS1 | 0.97 | 0.04 | 8.73E-133 | 7.67E-131 |
| ENSG00000149428 | HYOU1 | 0.97 | 0.05 | 4.26E-101 | 2.50E-99 |
| ENSG00000120306 | CYSTM1 | 0.97 | 0.06 | 2.30E-68 | 7.58E-67 |
| ENSG00000196693 | ZNF33B | 0.97 | 0.06 | 6.66E-67 | 2.14E-65 |
| ENSG00000112033 | PPARD | 0.97 | 0.06 | 6.66E-54 | 1.65E-52 |
| ENSG00000141503 | MINK1 | 0.97 | 0.06 | 1.17E-51 | 2.75E-50 |
| ENSG00000171928 | TVP23B | 0.97 | 0.08 | 1.55E-38 | 2.70E-37 |
| ENSG00000220205 | VAMP2 | 0.97 | 0.09 | 2.13E-30 | 2.91E-29 |
| ENSG00000170634 | ACYP2 | 0.97 | 0.11 | 1.16E-18 | 1.01E-17 |
| ENSG00000090339 | ICAM1 | 0.97 | 0.13 | 8.76E-16 | 6.72E-15 |
| ENSG00000226824 | | 0.97 | 0.17 | 1.51E-09 | 8.06E-09 |
| ENSG00000092068 | SLC7A8 | 0.97 | 0.19 | 3.43E-08 | 1.65E-07 |
| ENSG00000225206 | MIR137HG | 0.97 | 0.21 | 4.36E-07 | 1.92E-06 |
| ENSG00000109107 | ALDOC | 0.97 | 0.23 | 1.71E-06 | 7.11E-06 |
| ENSG00000205090 | TMEM240 | 0.97 | 0.24 | 4.47E-06 | 1.79E-05 |
| ENSG00000269737 | | 0.97 | 0.24 | 5.67E-06 | 2.25E-05 |
| ENSG00000272269 | NUP153-AS1 | 0.97 | 0.29 | 5.40E-05 | 1.93E-04 |
| ENSG00000258081 | | 0.97 | 0.31 | 1.17E-04 | 4.03E-04 |
| ENSG00000286735 | | 0.97 | 0.41 | 1.19E-03 | 3.58E-03 |
| ENSG00000284830 | | 0.97 | 0.41 | 1.25E-03 | 3.74E-03 |
| ENSG00000289997 | | 0.97 | 0.48 | 2.85E-03 | 8.04E-03 |
| ENSG00000182557 | SPNS3 | 0.97 | 0.8 | 4.11E-03 | 1.12E-02 |
| ENSG00000256030 | CBX3P4 | 0.97 | 0.56 | 4.65E-03 | 1.26E-02 |
| ENSG00000205323 | SARNP | 0.97 | 0.59 | 5.05E-03 | 1.36E-02 |
| ENSG00000156006 | NAT2 | 0.97 | 0.64 | 5.96E-03 | 1.58E-02 |
| ENSG00000130779 | CLIP1 | 0.96 | 0.05 | 6.57E-89 | 3.15E-87 |
| ENSG00000109929 | SC5D | 0.96 | 0.05 | 7.27E-81 | 3.02E-79 |
| ENSG00000112514 | CUTA | 0.96 | 0.06 | 1.11E-64 | 3.44E-63 |
| ENSG00000114784 | EIF1B | 0.96 | 0.08 | 3.00E-33 | 4.45E-32 |
| ENSG00000187601 | MAGEH1 | 0.96 | 0.08 | 1.48E-32 | 2.16E-31 |
| ENSG00000160753 | RUSC1 | 0.96 | 0.08 | 7.11E-31 | 9.86E-30 |
| ENSG00000092969 | TGFB2 | 0.96 | 0.09 | 9.75E-28 | 1.21E-26 |
| ENSG00000132004 | FBXW9 | 0.96 | 0.11 | 1.29E-18 | 1.12E-17 |
| ENSG00000159556 | ISL2 | 0.96 | 0.12 | 1.54E-15 | 1.17E-14 |
| ENSG00000149927 | DOC2A | 0.96 | 0.23 | 2.01E-06 | 8.30E-06 |
| ENSG00000232656 | IDI2-AS1 | 0.96 | 0.23 | 2.23E-06 | 9.16E-06 |
| ENSG00000170092 | SPDYE5 | 0.96 | 0.25 | 8.56E-06 | 3.33E-05 |
| ENSG00000211454 | AKR7L | 0.96 | 0.27 | 3.86E-05 | 1.40E-04 |
| ENSG00000203709 | MIR29B2CHG | 0.96 | 0.34 | 2.99E-04 | 9.79E-04 |
| ENSG00000248734 | | 0.96 | 0.37 | 7.11E-04 | 2.21E-03 |

| | | | | | |
|---|---|---|---|---|---|
| ENSG00000264107 | MIR4733HG | 0.96 | 0.44 | 1.97E-03 | 5.72E-03 |
| ENSG00000254162 | | 0.96 | 0.47 | 2.66E-03 | 7.54E-03 |
| ENSG00000254208 | | 0.96 | 0.47 | 2.72E-03 | 7.70E-03 |
| ENSG00000197915 | HRNR | 0.96 | 0.47 | 2.80E-03 | 7.92E-03 |
| ENSG00000284430 | | 0.96 | 0.49 | 3.18E-03 | 8.89E-03 |
| ENSG00000163050 | COQ8A | 0.96 | 0.53 | 4.16E-03 | 1.13E-02 |
| ENSG00000236924 | | 0.96 | 0.59 | 5.54E-03 | 1.48E-02 |
| ENSG00000043591 | ADRB1 | 0.96 | 0.67 | 6.45E-03 | 1.70E-02 |
| ENSG00000150961 | SEC24D | 0.95 | 0.04 | 2.63E-136 | 2.42E-134 |
| ENSG00000214548 | MEG3 | 0.95 | 0.05 | 1.67E-94 | 8.91E-93 |
| ENSG00000187735 | TCEA1 | 0.95 | 0.05 | 1.01E-92 | 5.30E-91 |
| ENSG00000243701 | DUBR | 0.95 | 0.05 | 3.57E-91 | 1.79E-89 |
| ENSG00000179918 | SEPHS2 | 0.95 | 0.05 | 1.15E-70 | 4.00E-69 |
| ENSG00000118058 | KMT2A | 0.95 | 0.06 | 1.15E-64 | 3.56E-63 |
| ENSG00000168280 | KIF5C | 0.95 | 0.06 | 3.98E-54 | 9.96E-53 |
| ENSG00000146278 | PNRC1 | 0.95 | 0.07 | 2.41E-47 | 5.10E-46 |
| ENSG00000139718 | SETD1B | 0.95 | 0.07 | 5.10E-47 | 1.08E-45 |
| ENSG00000204231 | RXRB | 0.95 | 0.07 | 9.92E-45 | 1.99E-43 |
| ENSG00000162298 | SYVN1 | 0.95 | 0.08 | 1.37E-37 | 2.33E-36 |
| ENSG00000173530 | TNFRSF10D | 0.95 | 0.09 | 6.25E-28 | 7.79E-27 |
| ENSG00000251136 | RIPK2-DT | 0.95 | 0.1 | 7.69E-24 | 8.32E-23 |
| ENSG00000223891 | OSER1-DT | 0.95 | 0.14 | 9.74E-13 | 6.39E-12 |
| ENSG00000123358 | NR4A1 | 0.95 | 0.15 | 3.26E-11 | 1.95E-10 |
| ENSG00000260260 | SNHG19 | 0.95 | 0.17 | 7.17E-10 | 3.91E-09 |
| ENSG00000140398 | NEIL1 | 0.95 | 0.17 | 3.48E-09 | 1.80E-08 |
| ENSG00000259972 | | 0.95 | 0.19 | 4.41E-08 | 2.10E-07 |
| ENSG00000261770 | | 0.95 | 0.2 | 3.07E-07 | 1.37E-06 |
| ENSG00000177045 | SIX5 | 0.95 | 0.23 | 2.08E-06 | 8.58E-06 |
| ENSG00000288253 | | 0.95 | 0.3 | 1.00E-04 | 3.49E-04 |
| ENSG00000231742 | LINC01273 | 0.95 | 0.3 | 1.13E-04 | 3.91E-04 |
| ENSG00000099977 | DDT | 0.95 | 0.31 | 1.82E-04 | 6.14E-04 |
| ENSG00000288868 | | 0.95 | 0.34 | 4.38E-04 | 1.41E-03 |
| ENSG00000255198 | SNHG9 | 0.95 | 0.46 | 2.54E-03 | 7.24E-03 |
| ENSG00000280767 | | 0.95 | 0.85 | 2.76E-03 | 7.81E-03 |
| ENSG00000232884 | | 0.95 | 0.5 | 3.53E-03 | 9.77E-03 |
| ENSG00000221916 | C19orf73 | 0.95 | 0.51 | 4.23E-03 | 1.15E-02 |
| ENSG00000258940 | MIA2-AS1 | 0.95 | 0.75 | 5.80E-03 | 1.54E-02 |
| ENSG00000134109 | EDEM1 | 0.94 | 0.05 | 2.22E-92 | 1.15E-90 |
| ENSG00000106299 | WASL | 0.94 | 0.05 | 3.44E-83 | 1.52E-81 |
| ENSG00000095794 | CREM | 0.94 | 0.06 | 1.63E-65 | 5.09E-64 |
| ENSG00000130340 | SNX9 | 0.94 | 0.06 | 1.98E-63 | 5.92E-62 |
| ENSG00000164823 | OSGIN2 | 0.94 | 0.07 | 1.37E-38 | 2.39E-37 |
| ENSG00000087903 | RFX2 | 0.94 | 0.09 | 2.04E-28 | 2.58E-27 |
| ENSG00000109680 | TBC1D19 | 0.94 | 0.1 | 6.98E-22 | 7.00E-21 |
| ENSG00000177042 | TMEM80 | 0.94 | 0.11 | 2.11E-17 | 1.75E-16 |
| ENSG00000250742 | LINC02381 | 0.94 | 0.12 | 1.31E-16 | 1.05E-15 |
| ENSG00000101084 | RAB5IF | 0.94 | 0.13 | 1.83E-14 | 1.32E-13 |
| ENSG00000290792 | | 0.94 | 0.13 | 8.66E-14 | 6.04E-13 |
| ENSG00000160271 | RALGDS | 0.94 | 0.15 | 8.04E-11 | 4.69E-10 |
| ENSG00000205746 | PKD1P4 | 0.94 | 0.17 | 3.59E-09 | 1.86E-08 |

| | | | | | |
|---|---|---|---|---|---|
| ENSG00000250220 | ANKRD17-DT | 0.94 | 0.19 | 7.62E-08 | 3.57E-07 |
| ENSG00000023839 | ABCC2 | 0.94 | 0.2 | 2.01E-07 | 9.13E-07 |
| ENSG00000214309 | MBLAC1 | 0.94 | 0.21 | 5.97E-07 | 2.60E-06 |
| ENSG00000290793 | | 0.94 | 0.22 | 2.45E-06 | 1.00E-05 |
| ENSG00000069696 | DRD4 | 0.94 | 0.23 | 4.30E-06 | 1.72E-05 |
| ENSG00000142700 | DMRTA2 | 0.94 | 0.23 | 5.22E-06 | 2.08E-05 |
| ENSG00000226696 | LENG8-AS1 | 0.94 | 0.24 | 9.44E-06 | 3.66E-05 |
| ENSG00000271743 | | 0.94 | 0.27 | 3.67E-05 | 1.34E-04 |
| ENSG00000167614 | TTYH1 | 0.94 | 0.29 | 8.61E-05 | 3.02E-04 |
| ENSG00000186919 | ZACN | 0.94 | 0.33 | 3.05E-04 | 9.96E-04 |
| ENSG00000267796 | LIN37 | 0.94 | 0.39 | 1.22E-03 | 3.67E-03 |
| ENSG00000249684 | | 0.94 | 0.43 | 2.25E-03 | 6.46E-03 |
| ENSG00000272323 | TTC23L-AS1 | 0.94 | 0.45 | 2.61E-03 | 7.40E-03 |
| ENSG00000279852 | | 0.94 | 0.45 | 2.66E-03 | 7.55E-03 |
| ENSG00000285427 | SOD2-OT1 | 0.94 | 0.45 | 2.70E-03 | 7.64E-03 |
| ENSG00000248498 | ASNSP1 | 0.94 | 0.84 | 3.25E-03 | 9.07E-03 |
| ENSG00000158315 | RHBDL2 | 0.94 | 0.49 | 3.97E-03 | 1.09E-02 |
| ENSG00000179571 | NBPF17P | 0.94 | 0.78 | 5.26E-03 | 1.41E-02 |
| ENSG00000256574 | OR13A1 | 0.94 | 0.77 | 5.58E-03 | 1.49E-02 |
| ENSG00000269836 | | 0.94 | 0.58 | 5.91E-03 | 1.57E-02 |
| ENSG00000229950 | TFAP2A-AS1 | 0.94 | 0.6 | 6.47E-03 | 1.70E-02 |
| ENSG00000258548 | LINC00645 | 0.94 | 0.6 | 6.63E-03 | 1.74E-02 |
| ENSG00000184347 | SLIT3 | 0.94 | 0.67 | 7.19E-03 | 1.88E-02 |
| ENSG00000006576 | PHTF2 | 0.93 | 0.05 | 5.47E-83 | 2.40E-81 |
| ENSG00000265808 | SEC22B | 0.93 | 0.06 | 2.28E-61 | 6.53E-60 |
| ENSG00000139428 | MMAB | 0.93 | 0.06 | 7.04E-56 | 1.82E-54 |
| ENSG00000160781 | PAQR6 | 0.93 | 0.1 | 6.82E-23 | 7.11E-22 |
| ENSG00000250571 | GLI4 | 0.93 | 0.1 | 5.88E-20 | 5.45E-19 |
| ENSG00000183718 | TRIM52 | 0.93 | 0.11 | 3.99E-18 | 3.40E-17 |
| ENSG00000268471 | MIR4453HG | 0.93 | 0.14 | 1.48E-12 | 9.60E-12 |
| ENSG00000188219 | POTEE | 0.93 | 0.15 | 2.80E-11 | 1.68E-10 |
| ENSG00000260464 | | 0.93 | 0.18 | 9.53E-09 | 4.78E-08 |
| ENSG00000160813 | PPP1R35 | 0.93 | 0.18 | 3.38E-08 | 1.62E-07 |
| ENSG00000203780 | FANK1 | 0.93 | 0.2 | 2.43E-07 | 1.09E-06 |
| ENSG00000154589 | LY96 | 0.93 | 0.21 | 6.57E-07 | 2.84E-06 |
| ENSG00000152763 | DNAI4 | 0.93 | 0.21 | 9.38E-07 | 3.99E-06 |
| ENSG00000280254 | | 0.93 | 0.23 | 5.80E-06 | 2.30E-05 |
| ENSG00000239332 | LINC01119 | 0.93 | 0.26 | 2.63E-05 | 9.73E-05 |
| ENSG00000223485 | LINC01615 | 0.93 | 0.27 | 4.22E-05 | 1.53E-04 |
| ENSG00000273576 | | 0.93 | 0.28 | 8.06E-05 | 2.84E-04 |
| ENSG00000183929 | DUSP5P1 | 0.93 | 0.3 | 1.52E-04 | 5.16E-04 |
| ENSG00000229419 | RALGAPA1P1 | 0.93 | 0.31 | 2.06E-04 | 6.89E-04 |
| ENSG00000185304 | RGPD2 | 0.93 | 0.33 | 4.42E-04 | 1.42E-03 |
| ENSG00000270808 | | 0.93 | 0.34 | 4.58E-04 | 1.47E-03 |
| ENSG00000258521 | | 0.93 | 0.34 | 5.87E-04 | 1.85E-03 |
| ENSG00000236138 | DUX4L26 | 0.93 | 0.39 | 1.32E-03 | 3.95E-03 |
| ENSG00000183281 | PLGLB1 | 0.93 | 0.41 | 1.60E-03 | 4.71E-03 |
| ENSG00000162931 | TRIM17 | 0.93 | 0.42 | 2.06E-03 | 5.95E-03 |
| ENSG00000228288 | PCAT6 | 0.93 | 0.45 | 2.92E-03 | 8.23E-03 |
| ENSG00000260778 | MIR3677HG | 0.93 | 0.48 | 3.78E-03 | 1.04E-02 |

| | | | | | |
|---|---|---|---|---|---|
| ENSG00000236796 | | 0.93 | 0.77 | 5.77E-03 | 1.54E-02 |
| ENSG00000286369 | | 0.93 | 0.68 | 7.27E-03 | 1.90E-02 |
| ENSG00000090861 | AARS1 | 0.92 | 0.05 | 7.96E-89 | 3.80E-87 |
| ENSG00000134294 | SLC38A2 | 0.92 | 0.05 | 4.18E-87 | 1.92E-85 |
| ENSG00000107290 | SETX | 0.92 | 0.05 | 8.08E-72 | 2.87E-70 |
| ENSG00000138078 | PREPL | 0.92 | 0.05 | 3.84E-70 | 1.32E-68 |
| ENSG00000103257 | SLC7A5 | 0.92 | 0.06 | 3.82E-47 | 8.06E-46 |
| ENSG00000178188 | SH2B1 | 0.92 | 0.08 | 3.12E-35 | 4.93E-34 |
| ENSG00000168003 | SLC3A2 | 0.92 | 0.08 | 3.06E-28 | 3.85E-27 |
| ENSG00000269834 | ZNF528-AS1 | 0.92 | 0.09 | 4.74E-26 | 5.55E-25 |
| ENSG00000100968 | NFATC4 | 0.92 | 0.09 | 2.37E-23 | 2.51E-22 |
| ENSG00000236753 | MKLN1-AS | 0.92 | 0.16 | 5.68E-10 | 3.12E-09 |
| ENSG00000163803 | PLB1 | 0.92 | 0.19 | 6.13E-08 | 2.89E-07 |
| ENSG00000260097 | SPDYE6 | 0.92 | 0.25 | 2.46E-05 | 9.14E-05 |
| ENSG00000286509 | | 0.92 | 0.27 | 7.01E-05 | 2.49E-04 |
| ENSG00000262089 | | 0.92 | 0.29 | 1.39E-04 | 4.75E-04 |
| ENSG00000249803 | | 0.92 | 0.33 | 5.00E-04 | 1.59E-03 |
| ENSG00000225655 | | 0.92 | 0.33 | 5.23E-04 | 1.66E-03 |
| ENSG00000289594 | | 0.92 | 0.36 | 9.22E-04 | 2.82E-03 |
| ENSG00000279108 | | 0.92 | 0.37 | 1.15E-03 | 3.47E-03 |
| ENSG00000289909 | | 0.92 | 0.4 | 1.62E-03 | 4.75E-03 |
| ENSG00000267577 | DNAAF3-AS1 | 0.92 | 0.41 | 1.91E-03 | 5.56E-03 |
| ENSG00000237605 | DYRK3-AS1 | 0.92 | 0.41 | 2.10E-03 | 6.06E-03 |
| ENSG00000272172 | | 0.92 | 0.52 | 5.00E-03 | 1.34E-02 |
| ENSG00000253317 | | 0.92 | 0.7 | 7.38E-03 | 1.92E-02 |
| ENSG00000267091 | CTBP2P7 | 0.92 | 0.7 | 7.66E-03 | 1.99E-02 |
| ENSG00000149187 | CELF1 | 0.91 | 0.03 | 3.78E-170 | 5.00E-168 |
| ENSG00000156639 | ZFAND3 | 0.91 | 0.04 | 3.34E-127 | 2.71E-125 |
| ENSG00000115944 | COX7A2L | 0.91 | 0.05 | 3.56E-84 | 1.58E-82 |
| ENSG00000160058 | BSDC1 | 0.91 | 0.05 | 1.94E-68 | 6.41E-67 |
| ENSG00000223768 | LINC00205 | 0.91 | 0.05 | 4.82E-64 | 1.46E-62 |
| ENSG00000119801 | YPEL5 | 0.91 | 0.06 | 5.41E-52 | 1.28E-50 |
| ENSG00000165861 | ZFYVE1 | 0.91 | 0.06 | 1.17E-49 | 2.61E-48 |
| ENSG00000095951 | HIVEP1 | 0.91 | 0.06 | 9.97E-49 | 2.16E-47 |
| ENSG00000128596 | CCDC136 | 0.91 | 0.07 | 1.54E-36 | 2.53E-35 |
| ENSG00000109089 | CDR2L | 0.91 | 0.08 | 2.29E-28 | 2.89E-27 |
| ENSG00000172922 | RNASEH2C | 0.91 | 0.09 | 7.27E-27 | 8.78E-26 |
| ENSG00000128000 | ZNF780B | 0.91 | 0.09 | 1.94E-23 | 2.06E-22 |
| ENSG00000158106 | RHPN1 | 0.91 | 0.1 | 5.22E-21 | 5.07E-20 |
| ENSG00000176597 | B3GNT5 | 0.91 | 0.1 | 2.18E-19 | 1.97E-18 |
| ENSG00000185158 | LRRC37B | 0.91 | 0.12 | 3.44E-15 | 2.58E-14 |
| ENSG00000185386 | MAPK11 | 0.91 | 0.12 | 4.36E-15 | 3.25E-14 |
| ENSG00000175106 | TVP23C | 0.91 | 0.16 | 7.24E-10 | 3.95E-09 |
| ENSG00000274180 | NATD1 | 0.91 | 0.17 | 1.94E-08 | 9.52E-08 |
| ENSG00000110944 | IL23A | 0.91 | 0.19 | 1.31E-07 | 6.02E-07 |
| ENSG00000258932 | MIR3171HG | 0.91 | 0.23 | 6.66E-06 | 2.62E-05 |
| ENSG00000260774 | | 0.91 | 0.26 | 4.74E-05 | 1.71E-04 |
| ENSG00000273080 | | 0.91 | 0.27 | 8.26E-05 | 2.90E-04 |
| ENSG00000281404 | LINC01176 | 0.91 | 0.3 | 2.48E-04 | 8.21E-04 |
| ENSG00000227959 | EPHA2-AS1 | 0.91 | 0.4 | 1.69E-03 | 4.94E-03 |

| | | | | | |
|---|---|---|---|---|---|
| ENSG00000198221 | AFDN-DT | 0.91 | 0.41 | 2.09E-03 | 6.03E-03 |
| ENSG00000289401 | | 0.91 | 0.44 | 3.12E-03 | 8.74E-03 |
| ENSG00000203288 | TDRKH-AS1 | 0.91 | 0.48 | 4.31E-03 | 1.17E-02 |
| ENSG00000267142 | | 0.91 | 0.49 | 4.74E-03 | 1.28E-02 |
| ENSG00000178130 | FAM27E5 | 0.91 | 0.5 | 5.10E-03 | 1.37E-02 |
| ENSG00000255867 | DENND5B-AS1 | 0.91 | 0.51 | 5.44E-03 | 1.45E-02 |
| ENSG00000279161 | | 0.91 | 0.52 | 5.67E-03 | 1.51E-02 |
| ENSG00000099974 | DDTL | 0.91 | 0.55 | 6.75E-03 | 1.77E-02 |
| ENSG00000253214 | | 0.91 | 0.55 | 6.76E-03 | 1.77E-02 |
| ENSG00000256508 | MRGPRF-AS1 | 0.91 | 0.63 | 8.37E-03 | 2.15E-02 |
| ENSG00000173812 | EIF1 | 0.9 | 0.04 | 9.20E-127 | 7.39E-125 |
| ENSG00000232677 | LINC00665 | 0.9 | 0.04 | 3.57E-97 | 1.97E-95 |
| ENSG00000012822 | CALCOCO1 | 0.9 | 0.05 | 2.05E-82 | 8.85E-81 |
| ENSG00000184575 | XPOT | 0.9 | 0.05 | 2.81E-70 | 9.71E-69 |
| ENSG00000176903 | PNMA1 | 0.9 | 0.06 | 1.19E-58 | 3.25E-57 |
| ENSG00000146592 | CREB5 | 0.9 | 0.06 | 1.00E-54 | 2.53E-53 |
| ENSG00000164327 | RICTOR | 0.9 | 0.06 | 5.16E-48 | 1.11E-46 |
| ENSG00000166946 | CCNDBP1 | 0.9 | 0.06 | 7.83E-46 | 1.60E-44 |
| ENSG00000163728 | TTC14 | 0.9 | 0.07 | 8.47E-37 | 1.40E-35 |
| ENSG00000100485 | SOS2 | 0.9 | 0.07 | 1.84E-35 | 2.94E-34 |
| ENSG00000008517 | IL32 | 0.9 | 0.09 | 2.70E-23 | 2.86E-22 |
| ENSG00000205707 | ETFRF1 | 0.9 | 0.1 | 8.32E-21 | 7.98E-20 |
| ENSG00000185818 | NAT8L | 0.9 | 0.11 | 1.56E-17 | 1.30E-16 |
| ENSG00000251247 | ZNF345 | 0.9 | 0.13 | 1.92E-12 | 1.24E-11 |
| ENSG00000229644 | NAMPTP1 | 0.9 | 0.15 | 7.15E-11 | 4.18E-10 |
| ENSG00000272667 | | 0.9 | 0.15 | 1.05E-10 | 6.09E-10 |
| ENSG00000166924 | NYAP1 | 0.9 | 0.15 | 3.05E-10 | 1.71E-09 |
| ENSG00000233527 | ZNF529-AS1 | 0.9 | 0.15 | 7.21E-10 | 3.94E-09 |
| ENSG00000258768 | | 0.9 | 0.19 | 3.60E-07 | 1.60E-06 |
| ENSG00000165879 | FRAT1 | 0.9 | 0.2 | 4.38E-07 | 1.93E-06 |
| ENSG00000176945 | MUC20 | 0.9 | 0.2 | 8.03E-07 | 3.44E-06 |
| ENSG00000261879 | ZNF594-DT | 0.9 | 0.23 | 7.00E-06 | 2.74E-05 |
| ENSG00000289975 | | 0.9 | 0.25 | 3.08E-05 | 1.13E-04 |
| ENSG00000023445 | BIRC3 | 0.9 | 0.27 | 7.24E-05 | 2.56E-04 |
| ENSG00000165899 | OTOGL | 0.9 | 0.35 | 8.73E-04 | 2.68E-03 |
| ENSG00000173080 | RXFP4 | 0.9 | 0.38 | 1.59E-03 | 4.68E-03 |
| ENSG00000228382 | ITPKB-IT1 | 0.9 | 0.85 | 2.93E-03 | 8.25E-03 |
| ENSG00000272696 | | 0.9 | 0.43 | 2.97E-03 | 8.36E-03 |
| ENSG00000280214 | | 0.9 | 0.43 | 3.05E-03 | 8.56E-03 |
| ENSG00000269038 | SF1-DT | 0.9 | 0.46 | 4.02E-03 | 1.10E-02 |
| ENSG00000231154 | MORF4L2-AS1 | 0.9 | 0.46 | 4.15E-03 | 1.13E-02 |
| ENSG00000273264 | | 0.9 | 0.48 | 4.79E-03 | 1.29E-02 |
| ENSG00000197993 | KEL | 0.9 | 0.48 | 4.80E-03 | 1.30E-02 |
| ENSG00000284700 | | 0.9 | 0.8 | 5.09E-03 | 1.37E-02 |
| ENSG00000174482 | LINGO2 | 0.9 | 0.51 | 5.96E-03 | 1.58E-02 |
| ENSG00000270130 | | 0.9 | 0.55 | 6.92E-03 | 1.81E-02 |
| ENSG00000172650 | AGAP5 | 0.9 | 0.55 | 7.28E-03 | 1.90E-02 |
| ENSG00000222043 | | 0.9 | 0.56 | 7.46E-03 | 1.94E-02 |
| ENSG00000269961 | ERBIN-DT | 0.9 | 0.57 | 7.53E-03 | 1.96E-02 |
| ENSG00000275092 | | 0.9 | 0.71 | 8.26E-03 | 2.13E-02 |

| | | | | | |
|---|---|---|---|---|---|
| ENSG00000148584 | A1CF | 0.9 | 0.61 | 8.35E-03 | 2.15E-02 |
| ENSG00000108854 | SMURF2 | 0.89 | 0.04 | 1.88E-126 | 1.50E-124 |
| ENSG00000073614 | KDM5A | 0.89 | 0.05 | 2.50E-80 | 1.03E-78 |
| ENSG00000026652 | AGPAT4 | 0.89 | 0.05 | 6.89E-63 | 2.03E-61 |
| ENSG00000111912 | NCOA7 | 0.89 | 0.05 | 1.04E-61 | 3.00E-60 |
| ENSG00000114554 | PLXNA1 | 0.89 | 0.06 | 1.32E-45 | 2.70E-44 |
| ENSG00000144218 | AFF3 | 0.89 | 0.07 | 1.54E-39 | 2.73E-38 |
| ENSG00000143862 | ARL8A | 0.89 | 0.07 | 1.41E-35 | 2.26E-34 |
| ENSG00000236279 | CLEC2L | 0.89 | 0.07 | 5.82E-35 | 9.10E-34 |
| ENSG00000170846 | | 0.89 | 0.08 | 2.31E-31 | 3.26E-30 |
| ENSG00000026559 | KCNG1 | 0.89 | 0.09 | 3.33E-25 | 3.78E-24 |
| ENSG00000080031 | PTPRH | 0.89 | 0.1 | 1.17E-19 | 1.07E-18 |
| ENSG00000088340 | FER1L4 | 0.89 | 0.1 | 5.97E-19 | 5.28E-18 |
| ENSG00000203993 | ARRDC1-AS1 | 0.89 | 0.11 | 2.91E-18 | 2.50E-17 |
| ENSG00000248323 | LUCAT1 | 0.89 | 0.13 | 2.79E-13 | 1.89E-12 |
| ENSG00000225361 | PPP1R26-AS1 | 0.89 | 0.15 | 1.09E-10 | 6.28E-10 |
| ENSG00000234327 | ZNF232-AS1 | 0.89 | 0.26 | 6.69E-05 | 2.38E-04 |
| ENSG00000265666 | RARA-AS1 | 0.89 | 0.27 | 1.02E-04 | 3.55E-04 |
| ENSG00000166126 | AMN | 0.89 | 0.29 | 1.85E-04 | 6.23E-04 |
| ENSG00000278993 | | 0.89 | 0.3 | 3.34E-04 | 1.09E-03 |
| ENSG00000266017 | MIR4477B | 0.89 | 0.33 | 7.29E-04 | 2.26E-03 |
| ENSG00000222046 | DCDC2B | 0.89 | 0.36 | 1.12E-03 | 3.38E-03 |
| ENSG00000286818 | | 0.89 | 0.4 | 2.17E-03 | 6.25E-03 |
| ENSG00000275576 | | 0.89 | 0.85 | 3.16E-03 | 8.84E-03 |
| ENSG00000234636 | MED14OS | 0.89 | 0.44 | 3.48E-03 | 9.63E-03 |
| ENSG00000290146 | | 0.89 | 0.45 | 4.00E-03 | 1.10E-02 |
| ENSG00000290760 | UBE2Q2P13 | 0.89 | 0.48 | 4.87E-03 | 1.31E-02 |
| ENSG00000254756 | | 0.89 | 0.49 | 5.21E-03 | 1.40E-02 |
| ENSG00000253695 | PINX1-DT | 0.89 | 0.76 | 7.05E-03 | 1.84E-02 |
| ENSG00000252172 | RNU6-720P | 0.89 | 0.54 | 7.30E-03 | 1.90E-02 |
| ENSG00000271265 | | 0.89 | 0.57 | 8.04E-03 | 2.08E-02 |
| ENSG00000163554 | SPTA1 | 0.89 | 0.69 | 8.91E-03 | 2.28E-02 |
| ENSG00000274698 | | 0.89 | 0.62 | 9.15E-03 | 2.33E-02 |
| ENSG00000286337 | | 0.89 | 0.65 | 9.43E-03 | 2.40E-02 |
| ENSG00000133858 | ZFC3H1 | 0.88 | 0.05 | 3.52E-76 | 1.37E-74 |
| ENSG00000160211 | G6PD | 0.88 | 0.06 | 7.32E-54 | 1.81E-52 |
| ENSG00000158158 | CNNM4 | 0.88 | 0.07 | 3.73E-36 | 6.06E-35 |
| ENSG00000103249 | CLCN7 | 0.88 | 0.07 | 7.39E-35 | 1.15E-33 |
| ENSG00000071246 | VASH1 | 0.88 | 0.07 | 5.98E-34 | 9.07E-33 |
| ENSG00000149932 | TMEM219 | 0.88 | 0.07 | 2.58E-33 | 3.84E-32 |
| ENSG00000155961 | RAB39B | 0.88 | 0.09 | 1.22E-22 | 1.26E-21 |
| ENSG00000167106 | EEIG1 | 0.88 | 0.09 | 1.85E-22 | 1.90E-21 |
| ENSG00000114268 | PFKFB4 | 0.88 | 0.11 | 1.57E-15 | 1.19E-14 |
| ENSG00000213222 | TOMM40P4 | 0.88 | 0.12 | 4.54E-15 | 3.38E-14 |
| ENSG00000135766 | EGLN1 | 0.88 | 0.13 | 2.96E-13 | 2.00E-12 |
| ENSG00000160055 | TMEM234 | 0.88 | 0.13 | 1.20E-12 | 7.85E-12 |
| ENSG00000254681 | PKD1P5 | 0.88 | 0.14 | 4.34E-11 | 2.57E-10 |
| ENSG00000154654 | NCAM2 | 0.88 | 0.15 | 1.80E-10 | 1.02E-09 |
| ENSG00000287407 | | 0.88 | 0.15 | 6.94E-10 | 3.80E-09 |
| ENSG00000142765 | SYTL1 | 0.88 | 0.17 | 1.78E-08 | 8.74E-08 |

| | | | | | |
|---|---|---|---|---|---|
| ENSG00000286656 | | 0.88 | 0.2 | 9.35E-07 | 3.98E-06 |
| ENSG00000186235 | LINC02610 | 0.88 | 0.24 | 2.62E-05 | 9.71E-05 |
| ENSG00000231527 | FAM27C | 0.88 | 0.25 | 5.11E-05 | 1.84E-04 |
| ENSG00000178342 | KCNG2 | 0.88 | 0.26 | 7.60E-05 | 2.68E-04 |
| ENSG00000272468 | | 0.88 | 0.35 | 1.19E-03 | 3.57E-03 |
| ENSG00000196747 | H2AC13 | 0.88 | 0.43 | 3.38E-03 | 9.38E-03 |
| ENSG00000155428 | TRIM74 | 0.88 | 0.47 | 5.32E-03 | 1.42E-02 |
| ENSG00000186466 | AQP7P1 | 0.88 | 0.51 | 6.70E-03 | 1.76E-02 |
| ENSG00000250582 | SMAD1-AS2 | 0.88 | 0.74 | 8.31E-03 | 2.14E-02 |
| ENSG00000241280 | | 0.88 | 0.56 | 8.35E-03 | 2.15E-02 |
| ENSG00000239282 | CASTOR1 | 0.88 | 0.64 | 9.46E-03 | 2.40E-02 |
| ENSG00000116133 | DHCR24 | 0.87 | 0.04 | 4.22E-108 | 2.72E-106 |
| ENSG00000059804 | SLC2A3 | 0.87 | 0.04 | 2.26E-84 | 1.01E-82 |
| ENSG00000133818 | RRAS2 | 0.87 | 0.05 | 6.34E-72 | 2.26E-70 |
| ENSG00000167522 | ANKRD11 | 0.87 | 0.05 | 1.72E-66 | 5.47E-65 |
| ENSG00000105835 | NAMPT | 0.87 | 0.06 | 2.43E-53 | 5.95E-52 |
| ENSG00000188554 | NBR1 | 0.87 | 0.06 | 7.07E-53 | 1.71E-51 |
| ENSG00000109572 | CLCN3 | 0.87 | 0.06 | 4.33E-50 | 9.71E-49 |
| ENSG00000138623 | SEMA7A | 0.87 | 0.06 | 1.43E-49 | 3.17E-48 |
| ENSG00000274211 | SOCS7 | 0.87 | 0.06 | 2.20E-49 | 4.85E-48 |
| ENSG00000160741 | CRTC2 | 0.87 | 0.07 | 3.14E-35 | 4.95E-34 |
| ENSG00000129946 | SHC2 | 0.87 | 0.08 | 1.35E-29 | 1.79E-28 |
| ENSG00000141582 | CBX4 | 0.87 | 0.09 | 9.10E-24 | 9.82E-23 |
| ENSG00000131480 | AOC2 | 0.87 | 0.09 | 1.96E-21 | 1.93E-20 |
| ENSG00000291118 | ZNF767P | 0.87 | 0.12 | 5.23E-15 | 3.88E-14 |
| ENSG00000133247 | KMT5C | 0.87 | 0.14 | 2.82E-11 | 1.69E-10 |
| ENSG00000275131 | PDE4DIPP2 | 0.87 | 0.17 | 3.23E-08 | 1.56E-07 |
| ENSG00000272812 | | 0.87 | 0.24 | 2.69E-05 | 9.95E-05 |
| ENSG00000124215 | CDH26 | 0.87 | 0.25 | 5.01E-05 | 1.80E-04 |
| ENSG00000247595 | MISFA | 0.87 | 0.35 | 1.29E-03 | 3.85E-03 |
| ENSG00000267421 | ZFP28-DT | 0.87 | 0.37 | 1.74E-03 | 5.08E-03 |
| ENSG00000286974 | | 0.87 | 0.39 | 2.35E-03 | 6.73E-03 |
| ENSG00000180458 | | 0.87 | 0.84 | 3.43E-03 | 9.53E-03 |
| ENSG00000127774 | EMC6 | 0.87 | 0.43 | 4.11E-03 | 1.12E-02 |
| ENSG00000226312 | CFLAR-AS1 | 0.87 | 0.43 | 4.28E-03 | 1.16E-02 |
| ENSG00000270996 | | 0.87 | 0.8 | 5.69E-03 | 1.52E-02 |
| ENSG00000227908 | IL6ST-DT | 0.87 | 0.52 | 7.63E-03 | 1.98E-02 |
| ENSG00000290105 | | 0.87 | 0.57 | 9.62E-03 | 2.44E-02 |
| ENSG00000232873 | RPL23AP93 | 0.87 | 0.69 | 9.89E-03 | 2.50E-02 |
| ENSG00000152672 | CLEC4F | 0.87 | 0.69 | 9.98E-03 | 2.52E-02 |
| ENSG00000011422 | PLAUR | 0.86 | 0.05 | 1.49E-69 | 5.08E-68 |
| ENSG00000187189 | TSPYL4 | 0.86 | 0.05 | 7.24E-57 | 1.90E-55 |
| ENSG00000116584 | ARHGEF2 | 0.86 | 0.06 | 3.66E-51 | 8.45E-50 |
| ENSG00000151553 | FHIP2A | 0.86 | 0.06 | 4.93E-51 | 1.13E-49 |
| ENSG00000070540 | WIPI1 | 0.86 | 0.07 | 5.91E-41 | 1.09E-39 |
| ENSG00000184903 | IMMP2L | 0.86 | 0.07 | 1.29E-34 | 1.99E-33 |
| ENSG00000101104 | PABPC1L | 0.86 | 0.07 | 1.30E-34 | 2.01E-33 |
| ENSG00000196323 | ZBTB44 | 0.86 | 0.07 | 1.77E-33 | 2.66E-32 |
| ENSG00000084444 | FAM234B | 0.86 | 0.07 | 9.03E-33 | 1.32E-31 |
| ENSG00000150527 | MIA2 | 0.86 | 0.08 | 6.46E-31 | 8.98E-30 |

| | | | | | |
|---|---|---|---|---|---|
| ENSG00000131669 | NINJ1 | 0.86 | 0.09 | 8.69E-24 | 9.38E-23 |
| ENSG00000142920 | AZIN2 | 0.86 | 0.1 | 3.82E-17 | 3.13E-16 |
| ENSG00000105613 | MAST1 | 0.86 | 0.12 | 3.02E-14 | 2.16E-13 |
| ENSG00000123989 | CHPF | 0.86 | 0.14 | 4.48E-11 | 2.65E-10 |
| ENSG00000232110 | | 0.86 | 0.14 | 1.04E-10 | 6.01E-10 |
| ENSG00000272872 | | 0.86 | 0.16 | 8.44E-09 | 4.25E-08 |
| ENSG00000225746 | MEG8 | 0.86 | 0.17 | 5.25E-08 | 2.49E-07 |
| ENSG00000260329 | TMEM263-DT | 0.86 | 0.18 | 1.21E-07 | 5.55E-07 |
| ENSG00000177854 | TMEM187 | 0.86 | 0.18 | 1.25E-07 | 5.77E-07 |
| ENSG00000260643 | | 0.86 | 0.22 | 1.28E-05 | 4.89E-05 |
| ENSG00000186976 | EFCAB6 | 0.86 | 0.23 | 2.14E-05 | 8.00E-05 |
| ENSG00000272983 | | 0.86 | 0.25 | 6.98E-05 | 2.47E-04 |
| ENSG00000177025 | C19orf18 | 0.86 | 0.26 | 8.64E-05 | 3.03E-04 |
| ENSG00000167861 | HID1 | 0.86 | 0.3 | 4.05E-04 | 1.31E-03 |
| ENSG00000290549 | | 0.86 | 0.32 | 8.42E-04 | 2.59E-03 |
| ENSG00000253354 | | 0.86 | 0.84 | 3.66E-03 | 1.01E-02 |
| ENSG00000182585 | EPGN | 0.86 | 0.83 | 3.99E-03 | 1.09E-02 |
| ENSG00000237399 | PITRM1-AS1 | 0.86 | 0.46 | 5.92E-03 | 1.57E-02 |
| ENSG00000253309 | SERPINE3 | 0.86 | 0.49 | 6.86E-03 | 1.80E-02 |
| ENSG00000244414 | CFHR1 | 0.86 | 0.76 | 7.82E-03 | 2.03E-02 |
| ENSG00000272432 | | 0.86 | 0.75 | 8.26E-03 | 2.13E-02 |
| ENSG00000262228 | | 0.86 | 0.58 | 9.71E-03 | 2.46E-02 |
| ENSG00000081051 | AFP | 0.86 | 0.69 | 1.07E-02 | 2.68E-02 |
| ENSG00000047056 | WDR37 | 0.85 | 0.04 | 5.62E-86 | 2.55E-84 |
| ENSG00000062716 | VMP1 | 0.85 | 0.04 | 9.78E-85 | 4.41E-83 |
| ENSG00000156273 | BACH1 | 0.85 | 0.05 | 1.42E-75 | 5.42E-74 |
| ENSG00000152332 | UHMK1 | 0.85 | 0.05 | 7.57E-70 | 2.59E-68 |
| ENSG00000172260 | NEGR1 | 0.85 | 0.05 | 9.81E-69 | 3.26E-67 |
| ENSG00000180488 | MIGA1 | 0.85 | 0.05 | 2.14E-63 | 6.39E-62 |
| ENSG00000108797 | CNTNAP1 | 0.85 | 0.07 | 2.55E-38 | 4.41E-37 |
| ENSG00000104497 | SNX16 | 0.85 | 0.07 | 5.29E-37 | 8.78E-36 |
| ENSG00000153558 | FBXL2 | 0.85 | 0.07 | 2.57E-32 | 3.71E-31 |
| ENSG00000120658 | ENOX1 | 0.85 | 0.07 | 3.48E-32 | 5.01E-31 |
| ENSG00000127080 | IPPK | 0.85 | 0.09 | 1.15E-24 | 1.28E-23 |
| ENSG00000130066 | SAT1 | 0.85 | 0.09 | 1.47E-21 | 1.46E-20 |
| ENSG00000104946 | TBC1D17 | 0.85 | 0.09 | 6.08E-20 | 5.63E-19 |
| ENSG00000116885 | OSCP1 | 0.85 | 0.1 | 2.56E-19 | 2.31E-18 |
| ENSG00000230333 | | 0.85 | 0.13 | 2.50E-12 | 1.60E-11 |
| ENSG00000188290 | HES4 | 0.85 | 0.13 | 8.89E-12 | 5.50E-11 |
| ENSG00000180425 | C11orf71 | 0.85 | 0.15 | 8.75E-10 | 4.74E-09 |
| ENSG00000154710 | RABGEF1 | 0.85 | 0.15 | 3.97E-09 | 2.05E-08 |
| ENSG00000164076 | CAMKV | 0.85 | 0.16 | 2.94E-08 | 1.42E-07 |
| ENSG00000203497 | PDCD4-AS1 | 0.85 | 0.19 | 1.01E-06 | 4.26E-06 |
| ENSG00000173928 | SWSAP1 | 0.85 | 0.2 | 2.34E-06 | 9.58E-06 |
| ENSG00000010610 | CD4 | 0.85 | 0.2 | 3.11E-06 | 1.26E-05 |
| ENSG00000269054 | ZNF497-AS1 | 0.85 | 0.3 | 4.76E-04 | 1.52E-03 |
| ENSG00000269600 | | 0.85 | 0.4 | 3.46E-03 | 9.59E-03 |
| ENSG00000227704 | | 0.85 | 0.56 | 1.01E-02 | 2.54E-02 |
| ENSG00000274317 | LINC02334 | 0.85 | 0.6 | 1.10E-02 | 2.76E-02 |
| ENSG00000127377 | CRYGN | 0.85 | 0.6 | 1.15E-02 | 2.86E-02 |

| | | | | | |
|---|---|---|---|---|---|
| ENSG00000253719 | ATXN7L3B | 0.84 | 0.04 | 1.42E-96 | 7.77E-95 |
| ENSG00000148730 | EIF4EBP2 | 0.84 | 0.04 | 2.07E-80 | 8.54E-79 |
| ENSG00000133657 | ATP13A3 | 0.84 | 0.05 | 4.22E-68 | 1.38E-66 |
| ENSG00000157796 | WDR19 | 0.84 | 0.05 | 2.47E-61 | 7.06E-60 |
| ENSG00000198046 | ZNF667 | 0.84 | 0.06 | 9.49E-45 | 1.90E-43 |
| ENSG00000196352 | CD55 | 0.84 | 0.06 | 1.66E-42 | 3.18E-41 |
| ENSG00000130150 | MOSPD2 | 0.84 | 0.07 | 2.14E-30 | 2.92E-29 |
| ENSG00000161010 | MRNIP | 0.84 | 0.08 | 1.56E-28 | 1.99E-27 |
| ENSG00000245694 | CRNDE | 0.84 | 0.08 | 4.19E-27 | 5.10E-26 |
| ENSG00000109736 | MFSD10 | 0.84 | 0.09 | 2.60E-22 | 2.64E-21 |
| ENSG00000154359 | LONRF1 | 0.84 | 0.09 | 3.35E-22 | 3.40E-21 |
| ENSG00000099992 | TBC1D10A | 0.84 | 0.11 | 3.65E-15 | 2.73E-14 |
| ENSG00000105696 | TMEM59L | 0.84 | 0.12 | 4.88E-14 | 3.45E-13 |
| ENSG00000087085 | ACHE | 0.84 | 0.12 | 8.48E-14 | 5.92E-13 |
| ENSG00000279088 | | 0.84 | 0.13 | 3.88E-12 | 2.46E-11 |
| ENSG00000289554 | | 0.84 | 0.13 | 4.98E-12 | 3.14E-11 |
| ENSG00000261455 | LINC01003 | 0.84 | 0.15 | 3.07E-09 | 1.60E-08 |
| ENSG00000268001 | CARD8-AS1 | 0.84 | 0.17 | 1.15E-07 | 5.29E-07 |
| ENSG00000268941 | LINC01711 | 0.84 | 0.18 | 2.61E-07 | 1.17E-06 |
| ENSG00000049249 | TNFRSF9 | 0.84 | 0.18 | 2.99E-07 | 1.34E-06 |
| ENSG00000204963 | PCDHA7 | 0.84 | 0.18 | 4.40E-07 | 1.94E-06 |
| ENSG00000154143 | PANX3 | 0.84 | 0.25 | 1.01E-04 | 3.52E-04 |
| ENSG00000212724 | KRTAP2-3 | 0.84 | 0.26 | 1.28E-04 | 4.40E-04 |
| ENSG00000285533 | RELA-DT | 0.84 | 0.26 | 1.66E-04 | 5.62E-04 |
| ENSG00000212124 | TAS2R19 | 0.84 | 0.31 | 8.09E-04 | 2.50E-03 |
| ENSG00000260228 | CDH13-AS2 | 0.84 | 0.32 | 1.06E-03 | 3.22E-03 |
| ENSG00000277959 | | 0.84 | 0.33 | 1.13E-03 | 3.42E-03 |
| ENSG00000240889 | NDUFB2-AS1 | 0.84 | 0.35 | 1.56E-03 | 4.60E-03 |
| ENSG00000142405 | NLRP12 | 0.84 | 0.4 | 3.44E-03 | 9.54E-03 |
| ENSG00000181997 | AQP7P2 | 0.84 | 0.84 | 3.80E-03 | 1.05E-02 |
| ENSG00000257580 | | 0.84 | 0.83 | 4.18E-03 | 1.14E-02 |
| ENSG00000137270 | GCM1 | 0.84 | 0.81 | 5.42E-03 | 1.45E-02 |
| ENSG00000272787 | | 0.84 | 0.45 | 6.14E-03 | 1.62E-02 |
| ENSG00000276103 | U4 | 0.84 | 0.79 | 6.89E-03 | 1.80E-02 |
| ENSG00000279452 | | 0.84 | 0.48 | 7.31E-03 | 1.90E-02 |
| ENSG00000232671 | ZNF687-AS1 | 0.84 | 0.55 | 1.07E-02 | 2.68E-02 |
| ENSG00000236047 | RPL13AP12 | 0.84 | 0.71 | 1.08E-02 | 2.72E-02 |
| ENSG00000142538 | PTH2 | 0.84 | 0.59 | 1.18E-02 | 2.93E-02 |
| ENSG00000144671 | SLC22A14 | 0.84 | 0.63 | 1.20E-02 | 2.98E-02 |
| ENSG00000159720 | ATP6V0D1 | 0.83 | 0.04 | 7.04E-91 | 3.51E-89 |
| ENSG00000117139 | KDM5B | 0.83 | 0.04 | 6.85E-88 | 3.22E-86 |
| ENSG00000123352 | SPATS2 | 0.83 | 0.05 | 3.24E-55 | 8.27E-54 |
| ENSG00000161091 | MFSD12 | 0.83 | 0.07 | 8.91E-36 | 1.44E-34 |
| ENSG00000189143 | CLDN4 | 0.83 | 0.08 | 4.48E-24 | 4.89E-23 |
| ENSG00000115137 | DNAJC27 | 0.83 | 0.09 | 7.86E-23 | 8.17E-22 |
| ENSG00000171105 | INSR | 0.83 | 0.09 | 8.94E-22 | 8.92E-21 |
| ENSG00000135540 | NHSL1 | 0.83 | 0.09 | 1.46E-20 | 1.38E-19 |
| ENSG00000244486 | SCARF2 | 0.83 | 0.09 | 3.42E-20 | 3.21E-19 |
| ENSG00000171223 | JUNB | 0.83 | 0.09 | 3.75E-20 | 3.51E-19 |
| ENSG00000100075 | SLC25A1 | 0.83 | 0.09 | 2.68E-19 | 2.42E-18 |

| | | | | | |
|---|---|---|---|---|---|
| ENSG00000198517 | MAFK | 0.83 | 0.11 | 1.52E-14 | 1.10E-13 |
| ENSG00000183793 | NPIPA5 | 0.83 | 0.13 | 6.71E-12 | 4.18E-11 |
| ENSG00000205138 | SDHAF1 | 0.83 | 0.13 | 3.19E-11 | 1.91E-10 |
| ENSG00000103269 | RHBDL1 | 0.83 | 0.16 | 2.03E-08 | 9.92E-08 |
| ENSG00000225975 | ZNF567-DT | 0.83 | 0.17 | 1.52E-07 | 6.93E-07 |
| ENSG00000105404 | RABAC1 | 0.83 | 0.18 | 5.73E-07 | 2.50E-06 |
| ENSG00000290928 | | 0.83 | 0.19 | 1.30E-06 | 5.44E-06 |
| ENSG00000237424 | FOXD2-AS1 | 0.83 | 0.2 | 4.27E-06 | 1.71E-05 |
| ENSG00000235919 | ASH1L-AS1 | 0.83 | 0.21 | 6.81E-06 | 2.68E-05 |
| ENSG00000291263 | SMG1P7 | 0.83 | 0.26 | 1.81E-04 | 6.11E-04 |
| ENSG00000273723 | SUGT1-DT | 0.83 | 0.28 | 3.84E-04 | 1.24E-03 |
| ENSG00000125266 | EFNB2 | 0.83 | 0.29 | 4.60E-04 | 1.47E-03 |
| ENSG00000283141 | LINC02666 | 0.83 | 0.29 | 5.25E-04 | 1.67E-03 |
| ENSG00000236516 | KLF2P4 | 0.83 | 0.3 | 6.75E-04 | 2.11E-03 |
| ENSG00000280007 | | 0.83 | 0.37 | 2.89E-03 | 8.14E-03 |
| ENSG00000260884 | | 0.83 | 0.39 | 3.41E-03 | 9.47E-03 |
| ENSG00000290073 | | 0.83 | 0.47 | 7.64E-03 | 1.98E-02 |
| ENSG00000289537 | | 0.83 | 0.48 | 8.18E-03 | 2.11E-02 |
| ENSG00000287760 | | 0.83 | 0.51 | 1.02E-02 | 2.57E-02 |
| ENSG00000261366 | MANEA-DT | 0.83 | 0.52 | 1.03E-02 | 2.59E-02 |
| ENSG00000277895 | | 0.83 | 0.67 | 1.23E-02 | 3.05E-02 |
| ENSG00000203987 | | 0.83 | 0.61 | 1.25E-02 | 3.08E-02 |
| ENSG00000250796 | CIDECP2 | 0.83 | 0.59 | 1.25E-02 | 3.09E-02 |
| ENSG00000270802 | | 0.83 | 0.67 | 1.26E-02 | 3.10E-02 |
| ENSG00000223878 | PPIAP53 | 0.83 | 0.67 | 1.28E-02 | 3.16E-02 |
| ENSG00000154277 | UCHL1 | 0.82 | 0.03 | 2.12E-172 | 2.91E-170 |
| ENSG00000111711 | GOLT1B | 0.82 | 0.05 | 1.94E-62 | 5.69E-61 |
| ENSG00000111276 | CDKN1B | 0.82 | 0.05 | 1.67E-59 | 4.66E-58 |
| ENSG00000276023 | DUSP14 | 0.82 | 0.05 | 2.44E-58 | 6.60E-57 |
| ENSG00000163257 | DCAF16 | 0.82 | 0.05 | 1.49E-56 | 3.89E-55 |
| ENSG00000255112 | CHMP1B | 0.82 | 0.05 | 2.54E-52 | 6.07E-51 |
| ENSG00000091879 | ANGPT2 | 0.82 | 0.06 | 4.02E-45 | 8.12E-44 |
| ENSG00000102287 | GABRE | 0.82 | 0.07 | 2.33E-34 | 3.57E-33 |
| ENSG00000110721 | CHKA | 0.82 | 0.07 | 9.41E-31 | 1.30E-29 |
| ENSG00000109220 | CHIC2 | 0.82 | 0.08 | 1.56E-27 | 1.93E-26 |
| ENSG00000181264 | TLCD5 | 0.82 | 0.08 | 2.29E-24 | 2.52E-23 |
| ENSG00000105321 | CCDC9 | 0.82 | 0.1 | 7.57E-17 | 6.13E-16 |
| ENSG00000198054 | DSCR8 | 0.82 | 0.12 | 3.71E-13 | 2.50E-12 |
| ENSG00000068137 | PLEKHH3 | 0.82 | 0.12 | 6.53E-12 | 4.06E-11 |
| ENSG00000229131 | | 0.82 | 0.18 | 1.20E-06 | 5.05E-06 |
| ENSG00000128692 | EIF2S2P4 | 0.82 | 0.19 | 1.76E-06 | 7.30E-06 |
| ENSG00000276952 | | 0.82 | 0.27 | 2.97E-04 | 9.74E-04 |
| ENSG00000183032 | SLC25A21 | 0.82 | 0.28 | 3.35E-04 | 1.09E-03 |
| ENSG00000177721 | ANXA2R | 0.82 | 0.28 | 4.61E-04 | 1.48E-03 |
| ENSG00000257557 | PPP1R12A-AS1 | 0.82 | 0.3 | 8.17E-04 | 2.52E-03 |
| ENSG00000156875 | MFSD14A | 0.82 | 0.31 | 9.97E-04 | 3.03E-03 |
| ENSG00000145491 | ROPN1L | 0.82 | 0.34 | 1.87E-03 | 5.44E-03 |
| ENSG00000135917 | SLC19A3 | 0.82 | 0.34 | 1.88E-03 | 5.49E-03 |
| ENSG00000235706 | DICER1-AS1 | 0.82 | 0.36 | 2.55E-03 | 7.27E-03 |
| ENSG00000288033 | | 0.82 | 0.42 | 5.23E-03 | 1.40E-02 |

| | | | | | |
|---|---|---|---|---|---|
| ENSG00000179826 | MRGPRX3 | 0.82 | 0.79 | 6.76E-03 | 1.77E-02 |
| ENSG00000100253 | MIOX | 0.82 | 0.79 | 7.36E-03 | 1.92E-02 |
| ENSG00000237505 | PKN2-AS1 | 0.82 | 0.78 | 7.57E-03 | 1.97E-02 |
| ENSG00000163121 | NEURL3 | 0.82 | 0.78 | 7.82E-03 | 2.03E-02 |
| ENSG00000234345 | ELF2P1 | 0.82 | 0.49 | 9.31E-03 | 2.37E-02 |
| ENSG00000100156 | SLC16A8 | 0.82 | 0.55 | 1.19E-02 | 2.95E-02 |
| ENSG00000279255 | | 0.82 | 0.55 | 1.22E-02 | 3.01E-02 |
| ENSG00000229598 | PRDX3P1 | 0.82 | 0.68 | 1.26E-02 | 3.11E-02 |
| ENSG00000291273 | | 0.82 | 0.6 | 1.33E-02 | 3.27E-02 |
| ENSG00000166483 | WEE1 | 0.81 | 0.04 | 1.15E-88 | 5.48E-87 |
| ENSG00000122641 | INHBA | 0.81 | 0.04 | 1.24E-76 | 4.88E-75 |
| ENSG00000150991 | UBC | 0.81 | 0.05 | 7.42E-64 | 2.23E-62 |
| ENSG00000141510 | TP53 | 0.81 | 0.05 | 7.76E-58 | 2.08E-56 |
| ENSG00000095564 | BTAF1 | 0.81 | 0.05 | 7.50E-56 | 1.94E-54 |
| ENSG00000159256 | MORC3 | 0.81 | 0.06 | 6.93E-48 | 1.48E-46 |
| ENSG00000100906 | NFKBIA | 0.81 | 0.06 | 7.72E-48 | 1.65E-46 |
| ENSG00000130703 | OSBPL2 | 0.81 | 0.06 | 1.17E-46 | 2.45E-45 |
| ENSG00000166598 | HSP90B1 | 0.81 | 0.06 | 5.33E-46 | 1.10E-44 |
| ENSG00000158480 | SPATA2 | 0.81 | 0.06 | 1.96E-39 | 3.46E-38 |
| ENSG00000256087 | ZNF432 | 0.81 | 0.07 | 1.58E-35 | 2.52E-34 |
| ENSG00000163001 | CFAP36 | 0.81 | 0.07 | 4.42E-29 | 5.77E-28 |
| ENSG00000225697 | SLC26A6 | 0.81 | 0.07 | 5.49E-29 | 7.13E-28 |
| ENSG00000083535 | PIBF1 | 0.81 | 0.09 | 3.82E-20 | 3.58E-19 |
| ENSG00000168237 | GLYCTK | 0.81 | 0.1 | 2.09E-17 | 1.73E-16 |
| ENSG00000267254 | ZNF790-AS1 | 0.81 | 0.11 | 1.69E-15 | 1.28E-14 |
| ENSG00000130758 | MAP3K10 | 0.81 | 0.12 | 4.06E-13 | 2.72E-12 |
| ENSG00000115507 | OTX1 | 0.81 | 0.15 | 1.88E-08 | 9.22E-08 |
| ENSG00000223764 | LINC02593 | 0.81 | 0.16 | 2.79E-08 | 1.35E-07 |
| ENSG00000213638 | ADAT3 | 0.81 | 0.18 | 7.21E-07 | 3.11E-06 |
| ENSG00000205209 | SCGB2B2 | 0.81 | 0.21 | 1.83E-05 | 6.88E-05 |
| ENSG00000284770 | TBCE | 0.81 | 0.23 | 6.85E-05 | 2.43E-04 |
| ENSG00000286667 | | 0.81 | 0.24 | 8.97E-05 | 3.13E-04 |
| ENSG00000291224 | | 0.81 | 0.25 | 1.16E-04 | 3.99E-04 |
| ENSG00000162997 | PRORSD1P | 0.81 | 0.26 | 2.67E-04 | 8.82E-04 |
| ENSG00000168454 | TXNDC2 | 0.81 | 0.29 | 5.93E-04 | 1.87E-03 |
| ENSG00000286342 | | 0.81 | 0.3 | 7.57E-04 | 2.34E-03 |
| ENSG00000281005 | LINC00921 | 0.81 | 0.38 | 3.89E-03 | 1.07E-02 |
| ENSG00000260425 | | 0.81 | 0.38 | 3.92E-03 | 1.08E-02 |
| ENSG00000222317 | RNA5SP118 | 0.81 | 0.48 | 9.36E-03 | 2.38E-02 |
| ENSG00000271869 | DCTN6-DT | 0.81 | 0.49 | 9.93E-03 | 2.51E-02 |
| ENSG00000271662 | PPP4R2P6 | 0.81 | 0.5 | 1.04E-02 | 2.63E-02 |
| ENSG00000229951 | FOSL2-AS1 | 0.81 | 0.74 | 1.07E-02 | 2.69E-02 |
| ENSG00000196611 | MMP1 | 0.81 | 0.53 | 1.21E-02 | 3.00E-02 |
| ENSG00000252759 | Y_RNA | 0.81 | 0.69 | 1.29E-02 | 3.17E-02 |
| ENSG00000259804 | CTCF-DT | 0.81 | 0.55 | 1.31E-02 | 3.21E-02 |
| ENSG00000099985 | OSM | 0.81 | 0.57 | 1.32E-02 | 3.23E-02 |
| ENSG00000279249 | | 0.81 | 0.69 | 1.32E-02 | 3.24E-02 |
| ENSG00000234134 | | 0.81 | 0.55 | 1.33E-02 | 3.26E-02 |
| ENSG00000103353 | UBFD1 | 0.8 | 0.03 | 2.60E-121 | 1.95E-119 |
| ENSG00000133318 | RTN3 | 0.8 | 0.04 | 3.32E-84 | 1.48E-82 |

| | | | | | |
|---|---|---|---|---|---|
| ENSG00000094975 | SUCO | 0.8 | 0.05 | 3.27E-57 | 8.64E-56 |
| ENSG00000019995 | ZRANB1 | 0.8 | 0.06 | 2.87E-40 | 5.18E-39 |
| ENSG00000188215 | DCUN1D3 | 0.8 | 0.06 | 1.64E-39 | 2.91E-38 |
| ENSG00000119878 | CRIPT | 0.8 | 0.07 | 2.74E-30 | 3.72E-29 |
| ENSG00000132471 | WBP2 | 0.8 | 0.07 | 5.45E-29 | 7.09E-28 |
| ENSG00000260920 | | 0.8 | 0.07 | 1.12E-27 | 1.39E-26 |
| ENSG00000164970 | FAM219A | 0.8 | 0.08 | 1.57E-26 | 1.87E-25 |
| ENSG00000170190 | SLC16A5 | 0.8 | 0.08 | 8.32E-25 | 9.34E-24 |
| ENSG00000006025 | OSBPL7 | 0.8 | 0.09 | 7.59E-21 | 7.30E-20 |
| ENSG00000160570 | DEDD2 | 0.8 | 0.09 | 8.67E-20 | 7.99E-19 |
| ENSG00000128228 | SDF2L1 | 0.8 | 0.1 | 3.10E-15 | 2.33E-14 |
| ENSG00000108590 | MED31 | 0.8 | 0.12 | 1.46E-11 | 8.94E-11 |
| ENSG00000261326 | LINC01355 | 0.8 | 0.17 | 2.33E-07 | 1.05E-06 |
| ENSG00000262655 | SPON1 | 0.8 | 0.18 | 1.69E-06 | 7.02E-06 |
| ENSG00000204922 | UQCC3 | 0.8 | 0.21 | 1.15E-05 | 4.43E-05 |
| ENSG00000259081 | | 0.8 | 0.25 | 2.08E-04 | 6.97E-04 |
| ENSG00000233030 | | 0.8 | 0.26 | 2.27E-04 | 7.55E-04 |
| ENSG00000273066 | | 0.8 | 0.26 | 2.87E-04 | 9.42E-04 |
| ENSG00000196696 | PDXDC2P-NPIPB14P | 0.8 | 0.32 | 1.37E-03 | 4.06E-03 |
| ENSG00000289207 | | 0.8 | 0.34 | 2.15E-03 | 6.19E-03 |
| ENSG00000004776 | HSPB6 | 0.8 | 0.35 | 2.72E-03 | 7.71E-03 |
| ENSG00000278952 | | 0.8 | 0.36 | 3.22E-03 | 9.00E-03 |
| ENSG00000289155 | | 0.8 | 0.38 | 4.18E-03 | 1.14E-02 |
| ENSG00000164185 | ZNF474 | 0.8 | 0.38 | 4.19E-03 | 1.14E-02 |
| ENSG00000204584 | | 0.8 | 0.45 | 8.42E-03 | 2.16E-02 |
| ENSG00000254285 | KRT8P3 | 0.8 | 0.45 | 8.92E-03 | 2.28E-02 |
| ENSG00000234692 | | 0.8 | 0.46 | 9.00E-03 | 2.30E-02 |
| ENSG00000248971 | KRT8P46 | 0.8 | 0.47 | 9.19E-03 | 2.34E-02 |
| ENSG00000270116 | | 0.8 | 0.51 | 1.20E-02 | 2.97E-02 |
| ENSG00000123096 | SSPN | 0.8 | 0.56 | 1.40E-02 | 3.42E-02 |
| ENSG00000287192 | | 0.8 | 0.58 | 1.43E-02 | 3.49E-02 |
| ENSG00000228323 | MYADM-AS1 | 0.8 | 0.56 | 1.43E-02 | 3.49E-02 |
| ENSG00000267033 | | 0.8 | 0.65 | 1.47E-02 | 3.58E-02 |
| ENSG00000229292 | RFPL4AL1 | 0.8 | 0.57 | 1.48E-02 | 3.59E-02 |
| ENSG00000257219 | LNCOG | 0.8 | 0.66 | 1.52E-02 | 3.68E-02 |
| ENSG00000260394 | STUB1-DT | 0.8 | 0.6 | 1.55E-02 | 3.74E-02 |
| ENSG00000140105 | WARS1 | 0.79 | 0.03 | 3.11E-119 | 2.26E-117 |
| ENSG00000041982 | TNC | 0.79 | 0.04 | 6.62E-79 | 2.69E-77 |
| ENSG00000128699 | ORMDL1 | 0.79 | 0.05 | 2.12E-53 | 5.19E-52 |
| ENSG00000133639 | BTG1 | 0.79 | 0.05 | 4.61E-52 | 1.09E-50 |
| ENSG00000162104 | ADCY9 | 0.79 | 0.06 | 3.22E-41 | 5.99E-40 |
| ENSG00000132376 | INPP5K | 0.79 | 0.06 | 8.18E-36 | 1.32E-34 |
| ENSG00000165997 | ARL5B | 0.79 | 0.06 | 4.55E-35 | 7.16E-34 |
| ENSG00000088179 | PTPN4 | 0.79 | 0.07 | 1.66E-30 | 2.28E-29 |
| ENSG00000130714 | POMT1 | 0.79 | 0.07 | 3.08E-27 | 3.77E-26 |
| ENSG00000158411 | MITD1 | 0.79 | 0.1 | 2.32E-17 | 1.92E-16 |
| ENSG00000102057 | KCND1 | 0.79 | 0.11 | 1.96E-14 | 1.41E-13 |
| ENSG00000125966 | MMP24 | 0.79 | 0.16 | 8.05E-08 | 3.76E-07 |
| ENSG00000256967 | | 0.79 | 0.17 | 2.61E-07 | 1.17E-06 |
| ENSG00000288838 | | 0.79 | 0.18 | 1.01E-06 | 4.30E-06 |

| | | | | | |
|---|---|---|---|---|---|
| ENSG00000102904 | TSNAXIP1 | 0.79 | 0.19 | 5.17E-06 | 2.06E-05 |
| ENSG00000259891 | | 0.79 | 0.2 | 1.05E-05 | 4.04E-05 |
| ENSG00000071073 | MGAT4A | 0.79 | 0.21 | 2.08E-05 | 7.78E-05 |
| ENSG00000143543 | JTB | 0.79 | 0.23 | 9.00E-05 | 3.15E-04 |
| ENSG00000214425 | LRRC37A4P | 0.79 | 0.28 | 7.24E-04 | 2.25E-03 |
| ENSG00000161955 | TNFSF13 | 0.79 | 0.33 | 2.02E-03 | 5.84E-03 |
| ENSG00000228350 | LINC02585 | 0.79 | 0.37 | 3.92E-03 | 1.08E-02 |
| ENSG00000224429 | LINC00539 | 0.79 | 0.37 | 4.31E-03 | 1.17E-02 |
| ENSG00000286989 | | 0.79 | 0.38 | 4.52E-03 | 1.23E-02 |
| ENSG00000227177 | AIMP1P2 | 0.79 | 0.81 | 5.81E-03 | 1.54E-02 |
| ENSG00000290127 | | 0.79 | 0.73 | 1.18E-02 | 2.93E-02 |
| ENSG00000106927 | AMBP | 0.79 | 0.51 | 1.28E-02 | 3.16E-02 |
| ENSG00000289241 | | 0.79 | 0.7 | 1.41E-02 | 3.44E-02 |
| ENSG00000285693 | | 0.79 | 0.53 | 1.41E-02 | 3.44E-02 |
| ENSG00000289005 | | 0.79 | 0.68 | 1.51E-02 | 3.66E-02 |
| ENSG00000260034 | LCMT1-AS2 | 0.79 | 0.6 | 1.57E-02 | 3.79E-02 |
| ENSG00000137880 | GCHFR | 0.79 | 0.62 | 1.60E-02 | 3.86E-02 |
| ENSG00000236595 | MED15P5 | 0.79 | 0.61 | 1.60E-02 | 3.86E-02 |
| ENSG00000285804 | | 0.79 | 0.6 | 1.62E-02 | 3.89E-02 |
| ENSG00000108406 | DHX40 | 0.78 | 0.05 | 2.44E-51 | 5.67E-50 |
| ENSG00000115963 | RND3 | 0.78 | 0.06 | 3.49E-43 | 6.80E-42 |
| ENSG00000011405 | PIK3C2A | 0.78 | 0.06 | 4.40E-42 | 8.38E-41 |
| ENSG00000137817 | PARP6 | 0.78 | 0.06 | 2.09E-39 | 3.70E-38 |
| ENSG00000154447 | SH3RF1 | 0.78 | 0.06 | 2.37E-34 | 3.63E-33 |
| ENSG00000066739 | ATG2B | 0.78 | 0.06 | 2.87E-34 | 4.38E-33 |
| ENSG00000080189 | SLC35C2 | 0.78 | 0.07 | 7.45E-32 | 1.06E-30 |
| ENSG00000183624 | HMCES | 0.78 | 0.07 | 2.79E-30 | 3.78E-29 |
| ENSG00000186866 | POFUT2 | 0.78 | 0.07 | 1.90E-28 | 2.41E-27 |
| ENSG00000132510 | KDM6B | 0.78 | 0.07 | 2.21E-27 | 2.72E-26 |
| ENSG00000280128 | | 0.78 | 0.09 | 2.42E-19 | 2.19E-18 |
| ENSG00000139190 | VAMP1 | 0.78 | 0.1 | 6.65E-17 | 5.41E-16 |
| ENSG00000197530 | MIB2 | 0.78 | 0.1 | 1.73E-15 | 1.31E-14 |
| ENSG00000177374 | HIC1 | 0.78 | 0.11 | 3.99E-13 | 2.68E-12 |
| ENSG00000227827 | PKD1P2 | 0.78 | 0.15 | 4.19E-08 | 2.00E-07 |
| ENSG00000290854 | | 0.78 | 0.2 | 1.29E-05 | 4.94E-05 |
| ENSG00000291120 | MRPL45P2 | 0.78 | 0.31 | 1.50E-03 | 4.42E-03 |
| ENSG00000121905 | HPCA | 0.78 | 0.31 | 1.61E-03 | 4.73E-03 |
| ENSG00000272438 | | 0.78 | 0.32 | 1.87E-03 | 5.45E-03 |
| ENSG00000271857 | | 0.78 | 0.36 | 3.83E-03 | 1.05E-02 |
| ENSG00000249628 | LINC00942 | 0.78 | 0.37 | 4.57E-03 | 1.24E-02 |
| ENSG00000241489 | | 0.78 | 0.41 | 6.53E-03 | 1.72E-02 |
| ENSG00000289269 | | 0.78 | 0.42 | 7.57E-03 | 1.97E-02 |
| ENSG00000248664 | | 0.78 | 0.45 | 9.91E-03 | 2.51E-02 |
| ENSG00000215895 | HSPA5P1 | 0.78 | 0.46 | 1.05E-02 | 2.65E-02 |
| ENSG00000254419 | | 0.78 | 0.7 | 1.49E-02 | 3.61E-02 |
| ENSG00000289592 | | 0.78 | 0.54 | 1.50E-02 | 3.63E-02 |
| ENSG00000277715 | | 0.78 | 0.54 | 1.50E-02 | 3.65E-02 |
| ENSG00000270061 | | 0.78 | 0.53 | 1.51E-02 | 3.66E-02 |
| ENSG00000099194 | SCD | 0.77 | 0.03 | 3.89E-111 | 2.58E-109 |
| ENSG00000023909 | GCLM | 0.77 | 0.04 | 7.85E-83 | 3.43E-81 |

| | | | | | |
|---|---|---|---|---|---|
| ENSG00000167004 | PDIA3 | 0.77 | 0.05 | 9.29E-56 | 2.40E-54 |
| ENSG00000164953 | TMEM67 | 0.77 | 0.06 | 2.22E-38 | 3.84E-37 |
| ENSG00000119899 | SLC17A5 | 0.77 | 0.07 | 4.30E-28 | 5.39E-27 |
| ENSG00000259330 | INAFM2 | 0.77 | 0.07 | 2.33E-27 | 2.87E-26 |
| ENSG00000104998 | IL27RA | 0.77 | 0.08 | 3.02E-24 | 3.31E-23 |
| ENSG00000129925 | PGAP6 | 0.77 | 0.09 | 2.69E-20 | 2.54E-19 |
| ENSG00000134590 | RTL8C | 0.77 | 0.09 | 1.62E-18 | 1.40E-17 |
| ENSG00000157379 | DHRS1 | 0.77 | 0.12 | 5.66E-11 | 3.33E-10 |
| ENSG00000254602 | MIR130AHG | 0.77 | 0.14 | 3.44E-09 | 1.78E-08 |
| ENSG00000213398 | LCAT | 0.77 | 0.14 | 6.99E-09 | 3.54E-08 |
| ENSG00000183891 | TTC32 | 0.77 | 0.15 | 1.53E-08 | 7.56E-08 |
| ENSG00000247516 | MIR4458HG | 0.77 | 0.15 | 3.34E-08 | 1.61E-07 |
| ENSG00000283236 | | 0.77 | 0.17 | 1.32E-06 | 5.57E-06 |
| ENSG00000269293 | ZSCAN16-AS1 | 0.77 | 0.18 | 1.98E-06 | 8.16E-06 |
| ENSG00000225218 | | 0.77 | 0.22 | 7.48E-05 | 2.64E-04 |
| ENSG00000183458 | PKD1P3 | 0.77 | 0.23 | 1.09E-04 | 3.78E-04 |
| ENSG00000196862 | RGPD4 | 0.77 | 0.24 | 1.99E-04 | 6.69E-04 |
| ENSG00000116678 | LEPR | 0.77 | 0.27 | 6.79E-04 | 2.12E-03 |
| ENSG00000290578 | | 0.77 | 0.28 | 8.75E-04 | 2.68E-03 |
| ENSG00000290956 | FAM21EP | 0.77 | 0.28 | 8.91E-04 | 2.73E-03 |
| ENSG00000212864 | RNF208 | 0.77 | 0.28 | 9.40E-04 | 2.87E-03 |
| ENSG00000274422 | | 0.77 | 0.3 | 1.39E-03 | 4.14E-03 |
| ENSG00000234498 | RPL13AP20 | 0.77 | 0.3 | 1.42E-03 | 4.22E-03 |
| ENSG00000198857 | HSD3BP5 | 0.77 | 0.33 | 2.65E-03 | 7.52E-03 |
| ENSG00000251081 | | 0.77 | 0.36 | 4.13E-03 | 1.13E-02 |
| ENSG00000183248 | PRR36 | 0.77 | 0.37 | 5.12E-03 | 1.38E-02 |
| ENSG00000289180 | | 0.77 | 0.79 | 7.87E-03 | 2.04E-02 |
| ENSG00000254739 | | 0.77 | 0.44 | 1.02E-02 | 2.57E-02 |
| ENSG00000254109 | RBPMS-AS1 | 0.77 | 0.45 | 1.10E-02 | 2.76E-02 |
| ENSG00000252244 | RNU7-3P | 0.77 | 0.46 | 1.12E-02 | 2.80E-02 |
| ENSG00000237159 | CNTFR-AS1 | 0.77 | 0.75 | 1.17E-02 | 2.92E-02 |
| ENSG00000259657 | PIGHP1 | 0.77 | 0.75 | 1.18E-02 | 2.93E-02 |
| ENSG00000286677 | | 0.77 | 0.74 | 1.21E-02 | 3.00E-02 |
| ENSG00000271855 | | 0.77 | 0.48 | 1.22E-02 | 3.02E-02 |
| ENSG00000240204 | SMKR1 | 0.77 | 0.51 | 1.44E-02 | 3.51E-02 |
| ENSG00000289110 | | 0.77 | 0.67 | 1.67E-02 | 3.99E-02 |
| ENSG00000285886 | | 0.77 | 0.66 | 1.71E-02 | 4.10E-02 |
| ENSG00000285130 | | 0.77 | 0.64 | 1.79E-02 | 4.25E-02 |
| ENSG00000230615 | | 0.77 | 0.64 | 1.80E-02 | 4.29E-02 |
| ENSG00000166224 | SGPL1 | 0.76 | 0.04 | 5.57E-87 | 2.54E-85 |
| ENSG00000119048 | UBE2B | 0.76 | 0.05 | 1.45E-62 | 4.24E-61 |
| ENSG00000152409 | JMY | 0.76 | 0.05 | 1.07E-50 | 2.45E-49 |
| ENSG00000204386 | NEU1 | 0.76 | 0.06 | 7.33E-41 | 1.35E-39 |
| ENSG00000178381 | ZFAND2A | 0.76 | 0.06 | 1.09E-37 | 1.86E-36 |
| ENSG00000137161 | CNPY3 | 0.76 | 0.06 | 5.52E-35 | 8.64E-34 |
| ENSG00000173915 | ATP5MK | 0.76 | 0.07 | 4.47E-28 | 5.60E-27 |
| ENSG00000185946 | RNPC3 | 0.76 | 0.08 | 1.78E-20 | 1.69E-19 |
| ENSG00000197774 | EME2 | 0.76 | 0.09 | 2.01E-17 | 1.67E-16 |
| ENSG00000146826 | TRAPPC14 | 0.76 | 0.09 | 8.17E-17 | 6.61E-16 |
| ENSG00000178449 | COX14 | 0.76 | 0.09 | 9.16E-17 | 7.39E-16 |

| | | | | | |
|---|---|---|---|---|---|
| ENSG00000225953 | SATB2-AS1 | 0.76 | 0.12 | 1.65E-11 | 1.00E-10 |
| ENSG00000258947 | TUBB3 | 0.76 | 0.14 | 8.89E-09 | 4.46E-08 |
| ENSG00000135976 | ANKRD36 | 0.76 | 0.14 | 8.90E-09 | 4.47E-08 |
| ENSG00000215030 | RPL13P12 | 0.76 | 0.15 | 4.82E-08 | 2.29E-07 |
| ENSG00000054148 | PHPT1 | 0.76 | 0.16 | 1.88E-07 | 8.53E-07 |
| ENSG00000270344 | POC1B-AS1 | 0.76 | 0.16 | 5.10E-07 | 2.23E-06 |
| ENSG00000261799 | | 0.76 | 0.16 | 6.06E-07 | 2.63E-06 |
| ENSG00000196741 | LINC01560 | 0.76 | 0.22 | 7.42E-05 | 2.62E-04 |
| ENSG00000268218 | | 0.76 | 0.25 | 3.34E-04 | 1.09E-03 |
| ENSG00000188227 | ZNF793 | 0.76 | 0.25 | 3.52E-04 | 1.14E-03 |
| ENSG00000196668 | LINC00173 | 0.76 | 0.25 | 3.95E-04 | 1.27E-03 |
| ENSG00000257815 | PRANCR | 0.76 | 0.25 | 3.95E-04 | 1.27E-03 |
| ENSG00000288766 | | 0.76 | 0.28 | 8.62E-04 | 2.64E-03 |
| ENSG00000226711 | FAM66C | 0.76 | 0.3 | 1.76E-03 | 5.13E-03 |
| ENSG00000272145 | NFYC-AS1 | 0.76 | 0.32 | 2.37E-03 | 6.79E-03 |
| ENSG00000086506 | HBQ1 | 0.76 | 0.37 | 5.85E-03 | 1.55E-02 |
| ENSG00000286909 | | 0.76 | 0.39 | 6.87E-03 | 1.80E-02 |
| ENSG00000289021 | | 0.76 | 0.41 | 8.14E-03 | 2.10E-02 |
| ENSG00000285748 | | 0.76 | 0.79 | 8.32E-03 | 2.14E-02 |
| ENSG00000153162 | BMP6 | 0.76 | 0.45 | 1.15E-02 | 2.87E-02 |
| ENSG00000258643 | BCL2L2-PABPN1 | 0.76 | 0.45 | 1.17E-02 | 2.90E-02 |
| ENSG00000272848 | | 0.76 | 0.45 | 1.19E-02 | 2.95E-02 |
| ENSG00000225339 | | 0.76 | 0.47 | 1.32E-02 | 3.25E-02 |
| ENSG00000266560 | | 0.76 | 0.48 | 1.34E-02 | 3.28E-02 |
| ENSG00000278075 | | 0.76 | 0.48 | 1.40E-02 | 3.43E-02 |
| ENSG00000207614 | MIR193A | 0.76 | 0.71 | 1.56E-02 | 3.77E-02 |
| ENSG00000279254 | | 0.76 | 0.69 | 1.66E-02 | 3.98E-02 |
| ENSG00000125895 | TMEM74B | 0.76 | 0.68 | 1.75E-02 | 4.17E-02 |
| ENSG00000267795 | SMIM22 | 0.76 | 0.67 | 1.79E-02 | 4.26E-02 |
| ENSG00000251138 | LINC02882 | 0.76 | 0.56 | 1.85E-02 | 4.38E-02 |
| ENSG00000279837 | | 0.76 | 0.62 | 1.87E-02 | 4.43E-02 |
| ENSG00000264520 | | 0.76 | 0.65 | 1.90E-02 | 4.50E-02 |
| ENSG00000179889 | PDXDC1 | 0.75 | 0.04 | 9.60E-93 | 5.03E-91 |
| ENSG00000183955 | KMT5A | 0.75 | 0.04 | 1.40E-87 | 6.50E-86 |
| ENSG00000115216 | NRBP1 | 0.75 | 0.04 | 8.70E-80 | 3.57E-78 |
| ENSG00000006125 | AP2B1 | 0.75 | 0.04 | 4.77E-79 | 1.94E-77 |
| ENSG00000198431 | TXNRD1 | 0.75 | 0.04 | 4.09E-73 | 1.51E-71 |
| ENSG00000142871 | CCN1 | 0.75 | 0.04 | 6.94E-70 | 2.37E-68 |
| ENSG00000135250 | SRPK2 | 0.75 | 0.05 | 5.29E-59 | 1.46E-57 |
| ENSG00000116793 | PHTF1 | 0.75 | 0.05 | 4.13E-50 | 9.27E-49 |
| ENSG00000157224 | CLDN12 | 0.75 | 0.05 | 6.92E-45 | 1.39E-43 |
| ENSG00000231074 | HCG18 | 0.75 | 0.05 | 3.55E-43 | 6.91E-42 |
| ENSG00000150457 | LATS2 | 0.75 | 0.06 | 2.41E-41 | 4.49E-40 |
| ENSG00000172795 | DCP2 | 0.75 | 0.06 | 4.28E-37 | 7.11E-36 |
| ENSG00000102100 | SLC35A2 | 0.75 | 0.07 | 9.18E-26 | 1.06E-24 |
| ENSG00000055211 | GINM1 | 0.75 | 0.07 | 1.00E-24 | 1.12E-23 |
| ENSG00000100836 | PABPN1 | 0.75 | 0.08 | 6.18E-20 | 5.73E-19 |
| ENSG00000197562 | RAB40C | 0.75 | 0.13 | 5.58E-10 | 3.07E-09 |
| ENSG00000267106 | ZNF561-AS1 | 0.75 | 0.16 | 2.71E-07 | 1.21E-06 |
| ENSG00000265298 | | 0.75 | 0.16 | 3.76E-07 | 1.67E-06 |

| | | | | | |
|---|---|---|---|---|---|
| ENSG00000166145 | SPINT1 | 0.75 | 0.17 | 8.47E-07 | 3.62E-06 |
| ENSG00000157680 | DGKI | 0.75 | 0.19 | 9.50E-06 | 3.68E-05 |
| ENSG00000205670 | SMIM11 | 0.75 | 0.24 | 2.89E-04 | 9.49E-04 |
| ENSG00000140993 | TIGD7 | 0.75 | 0.25 | 3.19E-04 | 1.04E-03 |
| ENSG00000280152 | | 0.75 | 0.26 | 5.26E-04 | 1.67E-03 |
| ENSG00000278948 | | 0.75 | 0.26 | 5.66E-04 | 1.79E-03 |
| ENSG00000229422 | | 0.75 | 0.28 | 9.30E-04 | 2.84E-03 |
| ENSG00000203279 | | 0.75 | 0.28 | 1.22E-03 | 3.65E-03 |
| ENSG00000272221 | | 0.75 | 0.3 | 1.79E-03 | 5.24E-03 |
| ENSG00000248752 | | 0.75 | 0.33 | 3.20E-03 | 8.94E-03 |
| ENSG00000025708 | TYMP | 0.75 | 0.35 | 4.64E-03 | 1.26E-02 |
| ENSG00000164707 | SLC13A4 | 0.75 | 0.37 | 5.87E-03 | 1.56E-02 |
| ENSG00000100290 | BIK | 0.75 | 0.38 | 6.45E-03 | 1.70E-02 |
| ENSG00000107201 | RIGI | 0.75 | 0.4 | 7.94E-03 | 2.05E-02 |
| ENSG00000285725 | | 0.75 | 0.4 | 8.53E-03 | 2.19E-02 |
| ENSG00000287569 | | 0.75 | 0.41 | 8.90E-03 | 2.28E-02 |
| ENSG00000286004 | | 0.75 | 0.43 | 1.11E-02 | 2.78E-02 |
| ENSG00000273084 | LINC03073 | 0.75 | 0.43 | 1.14E-02 | 2.85E-02 |
| ENSG00000223803 | RPS20P14 | 0.75 | 0.47 | 1.42E-02 | 3.46E-02 |
| ENSG00000278156 | TSC22D1-AS1 | 0.75 | 0.48 | 1.51E-02 | 3.66E-02 |
| ENSG00000289603 | | 0.75 | 0.62 | 1.96E-02 | 4.62E-02 |
| ENSG00000261202 | TNRC6B-DT | 0.75 | 0.64 | 2.01E-02 | 4.74E-02 |
| ENSG00000086232 | EIF2AK1 | 0.74 | 0.04 | 3.06E-91 | 1.54E-89 |
| ENSG00000128606 | LRRC17 | 0.74 | 0.04 | 2.85E-63 | 8.49E-62 |
| ENSG00000120742 | SERP1 | 0.74 | 0.05 | 6.07E-57 | 1.60E-55 |
| ENSG00000100221 | JOSD1 | 0.74 | 0.05 | 4.07E-50 | 9.15E-49 |
| ENSG00000113328 | CCNG1 | 0.74 | 0.06 | 3.00E-41 | 5.59E-40 |
| ENSG00000162695 | SLC30A7 | 0.74 | 0.06 | 3.99E-39 | 7.02E-38 |
| ENSG00000105281 | SLC1A5 | 0.74 | 0.06 | 4.82E-35 | 7.56E-34 |
| ENSG00000177606 | JUN | 0.74 | 0.06 | 1.11E-34 | 1.72E-33 |
| ENSG00000114933 | INO80D | 0.74 | 0.06 | 5.73E-32 | 8.18E-31 |
| ENSG00000143013 | LMO4 | 0.74 | 0.06 | 1.16E-30 | 1.59E-29 |
| ENSG00000197860 | SGTB | 0.74 | 0.07 | 3.83E-29 | 5.00E-28 |
| ENSG00000196365 | LONP1 | 0.74 | 0.07 | 7.31E-29 | 9.46E-28 |
| ENSG00000113811 | SELENOK | 0.74 | 0.07 | 3.24E-27 | 3.96E-26 |
| ENSG00000164603 | BMT2 | 0.74 | 0.07 | 6.83E-26 | 7.95E-25 |
| ENSG00000157764 | BRAF | 0.74 | 0.07 | 9.65E-26 | 1.12E-24 |
| ENSG00000188153 | COL4A5 | 0.74 | 0.07 | 1.53E-23 | 1.63E-22 |
| ENSG00000222009 | BTBD19 | 0.74 | 0.08 | 4.09E-21 | 3.99E-20 |
| ENSG00000260565 | ERVK13-1 | 0.74 | 0.08 | 3.50E-19 | 3.14E-18 |
| ENSG00000130203 | APOE | 0.74 | 0.08 | 3.79E-19 | 3.39E-18 |
| ENSG00000173933 | RBM4 | 0.74 | 0.09 | 1.54E-17 | 1.28E-16 |
| ENSG00000198832 | SELENOM | 0.74 | 0.09 | 2.48E-16 | 1.96E-15 |
| ENSG00000291087 | CRYBB2P1 | 0.74 | 0.1 | 1.86E-13 | 1.27E-12 |
| ENSG00000228594 | FNDC10 | 0.74 | 0.12 | 1.54E-10 | 8.82E-10 |
| ENSG00000133111 | RFXAP | 0.74 | 0.16 | 3.35E-07 | 1.49E-06 |
| ENSG00000225973 | PIGBOS1 | 0.74 | 0.21 | 6.56E-05 | 2.33E-04 |
| ENSG00000160282 | FTCD | 0.74 | 0.22 | 1.02E-04 | 3.55E-04 |
| ENSG00000099834 | CDHR5 | 0.74 | 0.24 | 3.47E-04 | 1.13E-03 |
| ENSG00000062582 | MRPS24 | 0.74 | 0.31 | 2.47E-03 | 7.05E-03 |

| | | | | | |
|---|---|---|---|---|---|
| ENSG00000249637 | ANKHD1-DT | 0.74 | 0.33 | 4.02E-03 | 1.10E-02 |
| ENSG00000102796 | DHRS12 | 0.74 | 0.37 | 7.28E-03 | 1.90E-02 |
| ENSG00000231966 | LINC02818 | 0.74 | 0.39 | 7.79E-03 | 2.02E-02 |
| ENSG00000288746 | | 0.74 | 0.4 | 8.86E-03 | 2.27E-02 |
| ENSG00000286408 | | 0.74 | 0.4 | 9.43E-03 | 2.40E-02 |
| ENSG00000261251 | EFCAB6-DT | 0.74 | 0.44 | 1.20E-02 | 2.97E-02 |
| ENSG00000257097 | CLIP1-AS1 | 0.74 | 0.44 | 1.28E-02 | 3.15E-02 |
| ENSG00000232645 | LINC01431 | 0.74 | 0.44 | 1.31E-02 | 3.21E-02 |
| ENSG00000223725 | | 0.74 | 0.73 | 1.49E-02 | 3.61E-02 |
| ENSG00000271009 | | 0.74 | 0.47 | 1.51E-02 | 3.66E-02 |
| ENSG00000267289 | PIN1-DT | 0.74 | 0.72 | 1.63E-02 | 3.91E-02 |
| ENSG00000225513 | RPL29P20 | 0.74 | 0.51 | 1.76E-02 | 4.20E-02 |
| ENSG00000250910 | MAP9-AS1 | 0.74 | 0.51 | 1.87E-02 | 4.43E-02 |
| ENSG00000124249 | KCNK15 | 0.74 | 0.53 | 1.91E-02 | 4.53E-02 |
| ENSG00000272097 | TMEM14B-DT | 0.74 | 0.59 | 2.07E-02 | 4.85E-02 |
| ENSG00000136197 | C7orf25 | 0.74 | 0.59 | 2.11E-02 | 4.95E-02 |
| ENSG00000131910 | NR0B2 | 0.74 | 0.62 | 2.11E-02 | 4.96E-02 |
| ENSG00000119537 | KDSR | 0.73 | 0.05 | 6.94E-49 | 1.52E-47 |
| ENSG00000178567 | EPM2AIP1 | 0.73 | 0.05 | 1.29E-43 | 2.54E-42 |
| ENSG00000134086 | VHL | 0.73 | 0.05 | 7.18E-43 | 1.39E-41 |
| ENSG00000111962 | UST | 0.73 | 0.06 | 3.38E-40 | 6.08E-39 |
| ENSG00000000419 | DPM1 | 0.73 | 0.06 | 3.71E-33 | 5.48E-32 |
| ENSG00000105426 | PTPRS | 0.73 | 0.07 | 5.30E-28 | 6.62E-27 |
| ENSG00000175137 | SH3BP5L | 0.73 | 0.07 | 1.08E-27 | 1.34E-26 |
| ENSG00000140450 | ARRDC4 | 0.73 | 0.07 | 1.44E-26 | 1.72E-25 |
| ENSG00000186395 | KRT10 | 0.73 | 0.07 | 6.96E-26 | 8.10E-25 |
| ENSG00000114383 | TUSC2 | 0.73 | 0.07 | 7.03E-26 | 8.17E-25 |
| ENSG00000283183 | | 0.73 | 0.07 | 1.62E-25 | 1.86E-24 |
| ENSG00000168591 | TMUB2 | 0.73 | 0.07 | 2.81E-25 | 3.20E-24 |
| ENSG00000147421 | HMBOX1 | 0.73 | 0.07 | 3.63E-25 | 4.12E-24 |
| ENSG00000237187 | NR2F1-AS1 | 0.73 | 0.07 | 1.23E-24 | 1.36E-23 |
| ENSG00000008283 | CYB561 | 0.73 | 0.07 | 5.03E-24 | 5.48E-23 |
| ENSG00000148384 | INPP5E | 0.73 | 0.08 | 9.46E-19 | 8.30E-18 |
| ENSG00000177666 | PNPLA2 | 0.73 | 0.09 | 5.49E-18 | 4.66E-17 |
| ENSG00000116983 | HPCAL4 | 0.73 | 0.09 | 1.74E-16 | 1.38E-15 |
| ENSG00000126062 | TMEM115 | 0.73 | 0.09 | 5.44E-16 | 4.23E-15 |
| ENSG00000229618 | | 0.73 | 0.1 | 2.94E-15 | 2.21E-14 |
| ENSG00000170271 | FAXDC2 | 0.73 | 0.11 | 9.72E-13 | 6.38E-12 |
| ENSG00000078814 | MYH7B | 0.73 | 0.12 | 7.11E-10 | 3.89E-09 |
| ENSG00000176182 | MYPOP | 0.73 | 0.12 | 7.78E-10 | 4.23E-09 |
| ENSG00000213085 | CFAP45 | 0.73 | 0.14 | 2.92E-08 | 1.41E-07 |
| ENSG00000260231 | KDM7A-DT | 0.73 | 0.15 | 4.29E-07 | 1.89E-06 |
| ENSG00000184451 | CCR10 | 0.73 | 0.16 | 7.04E-07 | 3.04E-06 |
| ENSG00000247373 | TMED2-DT | 0.73 | 0.19 | 2.75E-05 | 1.02E-04 |
| ENSG00000224687 | RASAL2-AS1 | 0.73 | 0.19 | 2.86E-05 | 1.05E-04 |
| ENSG00000227946 | INO80D-AS1 | 0.73 | 0.21 | 9.23E-05 | 3.22E-04 |
| ENSG00000118785 | SPP1 | 0.73 | 0.22 | 1.70E-04 | 5.75E-04 |
| ENSG00000198870 | STKLD1 | 0.73 | 0.23 | 2.57E-04 | 8.51E-04 |
| ENSG00000247157 | LINC01252 | 0.73 | 0.25 | 5.30E-04 | 1.68E-03 |
| ENSG00000291317 | TMEM276 | 0.73 | 0.29 | 1.96E-03 | 5.70E-03 |

| | | | | | |
|---|---|---|---|---|---|
| ENSG00000261126 | RBFADN | 0.73 | 0.29 | 2.00E-03 | 5.80E-03 |
| ENSG00000224546 | EIF4BP3 | 0.73 | 0.31 | 2.72E-03 | 7.69E-03 |
| ENSG00000276071 | | 0.73 | 0.32 | 3.14E-03 | 8.78E-03 |
| ENSG00000220685 | | 0.73 | 0.31 | 3.15E-03 | 8.83E-03 |
| ENSG00000164509 | IL31RA | 0.73 | 0.32 | 3.71E-03 | 1.02E-02 |
| ENSG00000291054 | | 0.73 | 0.33 | 3.88E-03 | 1.07E-02 |
| ENSG00000198075 | SULT1C4 | 0.73 | 0.33 | 4.00E-03 | 1.10E-02 |
| ENSG00000260911 | | 0.73 | 0.33 | 4.23E-03 | 1.15E-02 |
| ENSG00000269901 | | 0.73 | 0.34 | 4.98E-03 | 1.34E-02 |
| ENSG00000224903 | RNF32-AS1 | 0.73 | 0.37 | 6.96E-03 | 1.82E-02 |
| ENSG00000224046 | DMTF1-AS1 | 0.73 | 0.38 | 7.94E-03 | 2.05E-02 |
| ENSG00000248916 | NUP210P3 | 0.73 | 0.79 | 8.01E-03 | 2.07E-02 |
| ENSG00000259376 | | 0.73 | 0.4 | 1.04E-02 | 2.63E-02 |
| ENSG00000290357 | | 0.73 | 0.77 | 1.05E-02 | 2.65E-02 |
| ENSG00000273448 | | 0.73 | 0.41 | 1.08E-02 | 2.70E-02 |
| ENSG00000184303 | DRD5P1 | 0.73 | 0.76 | 1.24E-02 | 3.05E-02 |
| ENSG00000143365 | RORC | 0.73 | 0.45 | 1.52E-02 | 3.67E-02 |
| ENSG00000185527 | PDE6G | 0.73 | 0.45 | 1.53E-02 | 3.70E-02 |
| ENSG00000259744 | | 0.73 | 0.49 | 1.74E-02 | 4.16E-02 |
| ENSG00000289915 | | 0.73 | 0.7 | 1.87E-02 | 4.43E-02 |
| ENSG00000263142 | LRRC37A17P | 0.73 | 0.69 | 1.89E-02 | 4.47E-02 |
| ENSG00000290008 | | 0.73 | 0.53 | 2.04E-02 | 4.79E-02 |
| ENSG00000009335 | UBE3C | 0.72 | 0.04 | 8.80E-92 | 4.48E-90 |
| ENSG00000060982 | BCAT1 | 0.72 | 0.05 | 5.26E-55 | 1.34E-53 |
| ENSG00000134324 | LPIN1 | 0.72 | 0.05 | 3.77E-50 | 8.48E-49 |
| ENSG00000136436 | CALCOCO2 | 0.72 | 0.05 | 1.06E-47 | 2.26E-46 |
| ENSG00000213281 | NRAS | 0.72 | 0.05 | 3.45E-43 | 6.72E-42 |
| ENSG00000106799 | TGFBR1 | 0.72 | 0.06 | 3.60E-40 | 6.47E-39 |
| ENSG00000104312 | RIPK2 | 0.72 | 0.06 | 6.58E-35 | 1.03E-33 |
| ENSG00000187164 | SHTN1 | 0.72 | 0.06 | 3.78E-30 | 5.10E-29 |
| ENSG00000008300 | CELSR3 | 0.72 | 0.06 | 5.22E-30 | 7.01E-29 |
| ENSG00000165915 | SLC39A13 | 0.72 | 0.07 | 1.15E-24 | 1.28E-23 |
| ENSG00000108840 | HDAC5 | 0.72 | 0.07 | 3.68E-24 | 4.02E-23 |
| ENSG00000124067 | SLC12A4 | 0.72 | 0.08 | 1.27E-21 | 1.27E-20 |
| ENSG00000243335 | KCTD7 | 0.72 | 0.08 | 1.76E-20 | 1.67E-19 |
| ENSG00000105520 | PLPPR2 | 0.72 | 0.08 | 6.24E-20 | 5.78E-19 |
| ENSG00000242474 | LINC03015 | 0.72 | 0.08 | 7.54E-19 | 6.64E-18 |
| ENSG00000169710 | FASN | 0.72 | 0.08 | 1.27E-18 | 1.11E-17 |
| ENSG00000198453 | ZNF568 | 0.72 | 0.09 | 7.32E-17 | 5.94E-16 |
| ENSG00000174943 | KCTD13 | 0.72 | 0.09 | 9.72E-17 | 7.82E-16 |
| ENSG00000130176 | CNN1 | 0.72 | 0.09 | 4.21E-15 | 3.14E-14 |
| ENSG00000172375 | C2CD2L | 0.72 | 0.1 | 9.92E-15 | 7.27E-14 |
| ENSG00000148090 | AUH | 0.72 | 0.1 | 1.28E-14 | 9.31E-14 |
| ENSG00000141570 | CBX8 | 0.72 | 0.1 | 8.03E-14 | 5.61E-13 |
| ENSG00000137171 | KLC4 | 0.72 | 0.1 | 1.37E-13 | 9.45E-13 |
| ENSG00000104886 | PLEKHJ1 | 0.72 | 0.11 | 1.18E-11 | 7.25E-11 |
| ENSG00000147155 | EBP | 0.72 | 0.11 | 2.38E-11 | 1.44E-10 |
| ENSG00000155530 | LRGUK | 0.72 | 0.16 | 1.39E-06 | 5.83E-06 |
| ENSG00000184719 | RNLS | 0.72 | 0.16 | 1.56E-06 | 6.50E-06 |
| ENSG00000284669 | | 0.72 | 0.19 | 2.32E-05 | 8.66E-05 |

| | | | | | |
|---|---|---|---|---|---|
| ENSG00000250802 | ZBED3-AS1 | 0.72 | 0.19 | 3.22E-05 | 1.18E-04 |
| ENSG00000213889 | PPM1N | 0.72 | 0.2 | 4.65E-05 | 1.68E-04 |
| ENSG00000150281 | CTF1 | 0.72 | 0.21 | 1.32E-04 | 4.53E-04 |
| ENSG00000260708 | TBC1D22A-DT | 0.72 | 0.23 | 2.32E-04 | 7.70E-04 |
| ENSG00000232815 | DUX4L50 | 0.72 | 0.22 | 2.40E-04 | 7.95E-04 |
| ENSG00000291158 | LINC00869 | 0.72 | 0.23 | 3.08E-04 | 1.01E-03 |
| ENSG00000231952 | DPY19L1P2 | 0.72 | 0.24 | 4.36E-04 | 1.40E-03 |
| ENSG00000236700 | LINC01010 | 0.72 | 0.24 | 5.16E-04 | 1.64E-03 |
| ENSG00000260001 | TGFBR3L | 0.72 | 0.25 | 6.78E-04 | 2.12E-03 |
| ENSG00000289483 | | 0.72 | 0.25 | 7.23E-04 | 2.25E-03 |
| ENSG00000130307 | USHBP1 | 0.72 | 0.28 | 1.93E-03 | 5.61E-03 |
| ENSG00000237638 | LINC02245 | 0.72 | 0.3 | 2.83E-03 | 7.99E-03 |
| ENSG00000075073 | TACR2 | 0.72 | 0.3 | 2.96E-03 | 8.32E-03 |
| ENSG00000289019 | | 0.72 | 0.33 | 4.62E-03 | 1.25E-02 |
| ENSG00000271754 | | 0.72 | 0.77 | 1.09E-02 | 2.73E-02 |
| ENSG00000198626 | RYR2 | 0.72 | 0.4 | 1.09E-02 | 2.73E-02 |
| ENSG00000287817 | | 0.72 | 0.41 | 1.19E-02 | 2.95E-02 |
| ENSG00000078081 | LAMP3 | 0.72 | 0.42 | 1.36E-02 | 3.33E-02 |
| ENSG00000101489 | CELF4 | 0.72 | 0.73 | 1.56E-02 | 3.77E-02 |
| ENSG00000289890 | | 0.72 | 0.45 | 1.58E-02 | 3.82E-02 |
| ENSG00000256084 | | 0.72 | 0.47 | 1.74E-02 | 4.16E-02 |
| ENSG00000168374 | ARF4 | 0.71 | 0.04 | 1.55E-65 | 4.87E-64 |
| ENSG00000008083 | JARID2 | 0.71 | 0.04 | 6.32E-62 | 1.83E-60 |
| ENSG00000125977 | EIF2S2 | 0.71 | 0.04 | 8.14E-60 | 2.28E-58 |
| ENSG00000075420 | FNDC3B | 0.71 | 0.04 | 2.09E-56 | 5.42E-55 |
| ENSG00000109436 | TBC1D9 | 0.71 | 0.05 | 2.40E-45 | 4.87E-44 |
| ENSG00000155252 | PI4K2A | 0.71 | 0.06 | 2.19E-34 | 3.35E-33 |
| ENSG00000165516 | KLHDC2 | 0.71 | 0.06 | 2.32E-30 | 3.16E-29 |
| ENSG00000160752 | FDPS | 0.71 | 0.06 | 2.06E-28 | 2.61E-27 |
| ENSG00000119242 | CCDC92 | 0.71 | 0.07 | 7.91E-25 | 8.88E-24 |
| ENSG00000035115 | SH3YL1 | 0.71 | 0.08 | 1.88E-20 | 1.78E-19 |
| ENSG00000256537 | SMIM10L1 | 0.71 | 0.08 | 6.54E-19 | 5.78E-18 |
| ENSG00000152117 | SMPD4BP | 0.71 | 0.09 | 1.25E-15 | 9.51E-15 |
| ENSG00000171291 | ZNF439 | 0.71 | 0.09 | 5.86E-15 | 4.34E-14 |
| ENSG00000251322 | SHANK3 | 0.71 | 0.11 | 1.20E-11 | 7.40E-11 |
| ENSG00000224877 | NDUFAF8 | 0.71 | 0.11 | 1.34E-11 | 8.23E-11 |
| ENSG00000257698 | GIHCG | 0.71 | 0.11 | 2.72E-11 | 1.64E-10 |
| ENSG00000204344 | STK19 | 0.71 | 0.12 | 3.38E-10 | 1.89E-09 |
| ENSG00000236144 | TMEM147-AS1 | 0.71 | 0.13 | 2.07E-09 | 1.09E-08 |
| ENSG00000241749 | RPSAP52 | 0.71 | 0.14 | 4.10E-08 | 1.96E-07 |
| ENSG00000152669 | CCNO | 0.71 | 0.14 | 9.08E-08 | 4.23E-07 |
| ENSG00000269929 | MIRLET7A1HG | 0.71 | 0.14 | 1.17E-07 | 5.41E-07 |
| ENSG00000174004 | NRROS | 0.71 | 0.15 | 5.32E-07 | 2.32E-06 |
| ENSG00000291122 | CASTOR3P | 0.71 | 0.16 | 1.56E-06 | 6.49E-06 |
| ENSG00000100445 | SDR39U1 | 0.71 | 0.16 | 1.99E-06 | 8.21E-06 |
| ENSG00000277142 | LINC00235 | 0.71 | 0.2 | 5.94E-05 | 2.12E-04 |
| ENSG00000161509 | GRIN2C | 0.71 | 0.22 | 1.92E-04 | 6.45E-04 |
| ENSG00000206561 | COLQ | 0.71 | 0.23 | 3.40E-04 | 1.10E-03 |
| ENSG00000275457 | | 0.71 | 0.23 | 3.49E-04 | 1.13E-03 |
| ENSG00000233058 | ATP13A3-DT | 0.71 | 0.32 | 4.19E-03 | 1.14E-02 |

| | | | | | |
|---|---|---|---|---|---|
| ENSG00000278376 | | 0.71 | 0.34 | 5.96E-03 | 1.58E-02 |
| ENSG00000230798 | FOXD3-AS1 | 0.71 | 0.34 | 6.19E-03 | 1.64E-02 |
| ENSG00000255568 | BRWD1-AS2 | 0.71 | 0.37 | 8.94E-03 | 2.29E-02 |
| ENSG00000229930 | | 0.71 | 0.78 | 9.47E-03 | 2.41E-02 |
| ENSG00000150510 | FAM124A | 0.71 | 0.4 | 1.18E-02 | 2.93E-02 |
| ENSG00000223813 | PRR15-DT | 0.71 | 0.44 | 1.54E-02 | 3.72E-02 |
| ENSG00000279900 | | 0.71 | 0.44 | 1.61E-02 | 3.87E-02 |
| ENSG00000213413 | PVRIG | 0.71 | 0.46 | 1.87E-02 | 4.42E-02 |
| ENSG00000214212 | C19orf38 | 0.71 | 0.48 | 1.93E-02 | 4.55E-02 |
| ENSG00000185127 | C6orf120 | 0.7 | 0.04 | 5.63E-78 | 2.25E-76 |
| ENSG00000075391 | RASAL2 | 0.7 | 0.05 | 8.85E-43 | 1.71E-41 |
| ENSG00000145050 | MANF | 0.7 | 0.05 | 5.54E-42 | 1.05E-40 |
| ENSG00000092964 | DPYSL2 | 0.7 | 0.06 | 1.81E-37 | 3.05E-36 |
| ENSG00000196867 | ZFP28 | 0.7 | 0.06 | 2.55E-35 | 4.05E-34 |
| ENSG00000151746 | BICD1 | 0.7 | 0.06 | 2.97E-28 | 3.74E-27 |
| ENSG00000046604 | DSG2 | 0.7 | 0.07 | 2.96E-26 | 3.49E-25 |
| ENSG00000003249 | DBNDD1 | 0.7 | 0.07 | 3.58E-26 | 4.21E-25 |
| ENSG00000204310 | AGPAT1 | 0.7 | 0.07 | 3.74E-24 | 4.09E-23 |
| ENSG00000187801 | ZFP69B | 0.7 | 0.07 | 2.02E-22 | 2.06E-21 |
| ENSG00000178053 | MLF1 | 0.7 | 0.08 | 1.41E-18 | 1.23E-17 |
| ENSG00000132846 | ZBED3 | 0.7 | 0.09 | 3.98E-14 | 2.83E-13 |
| ENSG00000064547 | LPAR2 | 0.7 | 0.11 | 8.31E-12 | 5.15E-11 |
| ENSG00000258366 | RTEL1 | 0.7 | 0.11 | 1.06E-10 | 6.15E-10 |
| ENSG00000147894 | C9orf72 | 0.7 | 0.11 | 2.20E-10 | 1.25E-09 |
| ENSG00000223403 | MEG9 | 0.7 | 0.12 | 2.76E-09 | 1.44E-08 |
| ENSG00000260916 | CCPG1 | 0.7 | 0.13 | 3.93E-09 | 2.03E-08 |
| ENSG00000230844 | ZNF674-AS1 | 0.7 | 0.13 | 6.73E-09 | 3.41E-08 |
| ENSG00000197261 | C6orf141 | 0.7 | 0.13 | 8.94E-09 | 4.49E-08 |
| ENSG00000260942 | CAPN10-DT | 0.7 | 0.13 | 1.30E-08 | 6.44E-08 |
| ENSG00000186787 | SPIN2B | 0.7 | 0.15 | 6.10E-07 | 2.65E-06 |
| ENSG00000224934 | GOT1-DT | 0.7 | 0.19 | 3.29E-05 | 1.21E-04 |
| ENSG00000254595 | | 0.7 | 0.22 | 2.81E-04 | 9.24E-04 |
| ENSG00000155066 | PROM2 | 0.7 | 0.25 | 8.24E-04 | 2.54E-03 |
| ENSG00000250337 | PURPL | 0.7 | 0.25 | 9.73E-04 | 2.96E-03 |
| ENSG00000240370 | RPL13P5 | 0.7 | 0.26 | 1.37E-03 | 4.06E-03 |
| ENSG00000165171 | METTL27 | 0.7 | 0.35 | 7.48E-03 | 1.95E-02 |
| ENSG00000130045 | NXNL2 | 0.7 | 0.41 | 1.41E-02 | 3.44E-02 |
| ENSG00000284602 | | 0.7 | 0.41 | 1.48E-02 | 3.59E-02 |
| ENSG00000226747 | FSIP2-AS2 | 0.7 | 0.72 | 1.80E-02 | 4.28E-02 |
| ENSG00000207959 | MIR656 | 0.7 | 0.46 | 1.95E-02 | 4.61E-02 |
| ENSG00000289704 | | 0.7 | 0.71 | 1.97E-02 | 4.66E-02 |
| ENSG00000289689 | | 0.7 | 0.46 | 2.04E-02 | 4.80E-02 |
| ENSG00000123091 | RNF11 | 0.69 | 0.04 | 1.45E-75 | 5.55E-74 |
| ENSG00000089063 | TMEM230 | 0.69 | 0.04 | 7.90E-71 | 2.78E-69 |
| ENSG00000143575 | HAX1 | 0.69 | 0.04 | 2.52E-53 | 6.16E-52 |
| ENSG00000148154 | UGCG | 0.69 | 0.05 | 6.68E-52 | 1.57E-50 |
| ENSG00000179218 | CALR | 0.69 | 0.05 | 1.18E-50 | 2.69E-49 |
| ENSG00000184203 | PPP1R2 | 0.69 | 0.05 | 5.16E-44 | 1.02E-42 |
| ENSG00000123384 | LRP1 | 0.69 | 0.05 | 3.61E-40 | 6.48E-39 |
| ENSG00000204619 | PPP1R11 | 0.69 | 0.05 | 7.41E-39 | 1.30E-37 |

| | | | | | |
|---|---|---|---|---|---|
| ENSG00000034713 | GABARAPL2 | 0.69 | 0.05 | 2.15E-38 | 3.73E-37 |
| ENSG00000097007 | ABL1 | 0.69 | 0.05 | 8.62E-38 | 1.47E-36 |
| ENSG00000145362 | ANK2 | 0.69 | 0.06 | 4.83E-35 | 7.59E-34 |
| ENSG00000168894 | RNF181 | 0.69 | 0.06 | 1.49E-34 | 2.29E-33 |
| ENSG00000143067 | ZNF697 | 0.69 | 0.06 | 7.60E-33 | 1.11E-31 |
| ENSG00000157933 | SKI | 0.69 | 0.06 | 4.45E-31 | 6.21E-30 |
| ENSG00000127920 | GNG11 | 0.69 | 0.06 | 1.74E-30 | 2.38E-29 |
| ENSG00000171735 | CAMTA1 | 0.69 | 0.07 | 5.77E-26 | 6.73E-25 |
| ENSG00000189410 | SH2D5 | 0.69 | 0.07 | 1.93E-22 | 1.98E-21 |
| ENSG00000173875 | ZNF791 | 0.69 | 0.07 | 4.37E-21 | 4.26E-20 |
| ENSG00000090266 | NDUFB2 | 0.69 | 0.08 | 4.89E-19 | 4.35E-18 |
| ENSG00000107731 | UNC5B | 0.69 | 0.08 | 2.89E-18 | 2.48E-17 |
| ENSG00000181773 | GPR3 | 0.69 | 0.09 | 1.21E-15 | 9.23E-15 |
| ENSG00000180354 | MTURN | 0.69 | 0.1 | 3.78E-13 | 2.54E-12 |
| ENSG00000290873 | LINC03009 | 0.69 | 0.1 | 8.72E-13 | 5.74E-12 |
| ENSG00000170390 | DCLK2 | 0.69 | 0.1 | 2.48E-12 | 1.59E-11 |
| ENSG00000181588 | MEX3D | 0.69 | 0.11 | 1.63E-11 | 9.91E-11 |
| ENSG00000212694 | LINC01089 | 0.69 | 0.11 | 2.34E-11 | 1.41E-10 |
| ENSG00000154134 | ROBO3 | 0.69 | 0.11 | 1.24E-10 | 7.17E-10 |
| ENSG00000120693 | SMAD9 | 0.69 | 0.12 | 1.64E-09 | 8.73E-09 |
| ENSG00000231312 | MAP4K3-DT | 0.69 | 0.13 | 8.05E-09 | 4.06E-08 |
| ENSG00000239779 | WBP1 | 0.69 | 0.13 | 3.26E-08 | 1.57E-07 |
| ENSG00000276900 | | 0.69 | 0.18 | 1.61E-05 | 6.09E-05 |
| ENSG00000242622 | GSK3B-DT | 0.69 | 0.18 | 2.86E-05 | 1.05E-04 |
| ENSG00000196605 | ZNF846 | 0.69 | 0.19 | 6.29E-05 | 2.24E-04 |
| ENSG00000181652 | ATG9B | 0.69 | 0.2 | 8.34E-05 | 2.93E-04 |
| ENSG00000171189 | GRIK1 | 0.69 | 0.22 | 3.52E-04 | 1.14E-03 |
| ENSG00000091622 | PITPNM3 | 0.69 | 0.24 | 7.32E-04 | 2.27E-03 |
| ENSG00000197301 | HMGA2-AS1 | 0.69 | 0.27 | 1.94E-03 | 5.65E-03 |
| ENSG00000070831 | CDC42 | 0.69 | 0.3 | 3.63E-03 | 1.01E-02 |
| ENSG00000186409 | CCDC30 | 0.69 | 0.32 | 5.10E-03 | 1.37E-02 |
| ENSG00000075790 | BCAP29 | 0.69 | 0.32 | 5.40E-03 | 1.44E-02 |
| ENSG00000288891 | | 0.69 | 0.34 | 7.11E-03 | 1.86E-02 |
| ENSG00000289523 | | 0.69 | 0.34 | 7.54E-03 | 1.96E-02 |
| ENSG00000261430 | | 0.69 | 0.79 | 7.86E-03 | 2.03E-02 |
| ENSG00000253658 | LINC01592 | 0.69 | 0.4 | 1.34E-02 | 3.30E-02 |
| ENSG00000261441 | POLG-DT | 0.69 | 0.41 | 1.56E-02 | 3.76E-02 |
| ENSG00000199092 | MIR410 | 0.69 | 0.74 | 1.62E-02 | 3.90E-02 |
| ENSG00000249884 | RNF103-CHMP3 | 0.69 | 0.42 | 1.63E-02 | 3.91E-02 |
| ENSG00000240871 | KRTAP4-7 | 0.69 | 0.71 | 2.06E-02 | 4.84E-02 |
| ENSG00000131323 | TRAF3 | 0.68 | 0.04 | 1.17E-53 | 2.89E-52 |
| ENSG00000166741 | NNMT | 0.68 | 0.05 | 1.61E-52 | 3.86E-51 |
| ENSG00000139372 | TDG | 0.68 | 0.05 | 5.69E-41 | 1.05E-39 |
| ENSG00000115520 | COQ10B | 0.68 | 0.05 | 2.03E-38 | 3.52E-37 |
| ENSG00000100554 | ATP6V1D | 0.68 | 0.05 | 1.11E-35 | 1.79E-34 |
| ENSG00000152137 | HSPB8 | 0.68 | 0.06 | 1.87E-34 | 2.87E-33 |
| ENSG00000164031 | DNAJB14 | 0.68 | 0.06 | 6.61E-33 | 9.71E-32 |
| ENSG00000136560 | TANK | 0.68 | 0.06 | 2.51E-31 | 3.53E-30 |
| ENSG00000198856 | OSTC | 0.68 | 0.06 | 2.94E-29 | 3.86E-28 |
| ENSG00000125356 | NDUFA1 | 0.68 | 0.06 | 1.50E-27 | 1.86E-26 |

| | | | | | |
|---|---|---|---|---|---|
| ENSG00000163629 | PTPN13 | 0.68 | 0.07 | 2.16E-26 | 2.57E-25 |
| ENSG00000168944 | CEP120 | 0.68 | 0.07 | 1.74E-24 | 1.93E-23 |
| ENSG00000206418 | RAB12 | 0.68 | 0.08 | 6.28E-20 | 5.82E-19 |
| ENSG00000187531 | SIRT7 | 0.68 | 0.08 | 3.43E-19 | 3.08E-18 |
| ENSG00000287151 | C2orf27A | 0.68 | 0.08 | 3.12E-17 | 2.57E-16 |
| ENSG00000126351 | THRA | 0.68 | 0.09 | 3.12E-16 | 2.46E-15 |
| ENSG00000247828 | TMEM161B-DT | 0.68 | 0.09 | 3.99E-16 | 3.12E-15 |
| ENSG00000065833 | ME1 | 0.68 | 0.09 | 3.24E-15 | 2.43E-14 |
| ENSG00000174851 | YIF1A | 0.68 | 0.09 | 4.57E-15 | 3.40E-14 |
| ENSG00000185189 | NRBP2 | 0.68 | 0.09 | 7.50E-14 | 5.25E-13 |
| ENSG00000116761 | CTH | 0.68 | 0.09 | 1.95E-13 | 1.34E-12 |
| ENSG00000136842 | TMOD1 | 0.68 | 0.11 | 2.42E-11 | 1.46E-10 |
| ENSG00000182400 | TRAPPC6B | 0.68 | 0.11 | 2.46E-10 | 1.39E-09 |
| ENSG00000115226 | FNDC4 | 0.68 | 0.11 | 4.52E-10 | 2.50E-09 |
| ENSG00000197119 | SLC25A29 | 0.68 | 0.13 | 1.14E-08 | 5.69E-08 |
| ENSG00000160200 | CBS | 0.68 | 0.16 | 3.96E-06 | 1.59E-05 |
| ENSG00000137103 | TMEM8B | 0.68 | 0.17 | 1.70E-05 | 6.41E-05 |
| ENSG00000215067 | ALOX12-AS1 | 0.68 | 0.18 | 4.25E-05 | 1.54E-04 |
| ENSG00000291092 | PMS2P3 | 0.68 | 0.18 | 4.31E-05 | 1.56E-04 |
| ENSG00000053702 | NRIP2 | 0.68 | 0.19 | 6.74E-05 | 2.40E-04 |
| ENSG00000261512 | | 0.68 | 0.2 | 1.16E-04 | 4.01E-04 |
| ENSG00000103723 | AP3B2 | 0.68 | 0.21 | 1.73E-04 | 5.85E-04 |
| ENSG00000235961 | PNMA6A | 0.68 | 0.24 | 7.62E-04 | 2.36E-03 |
| ENSG00000158716 | DUSP23 | 0.68 | 0.26 | 1.51E-03 | 4.45E-03 |
| ENSG00000204052 | LRRC73 | 0.68 | 0.26 | 1.82E-03 | 5.30E-03 |
| ENSG00000279198 | | 0.68 | 0.28 | 3.06E-03 | 8.58E-03 |
| ENSG00000260136 | | 0.68 | 0.32 | 6.97E-03 | 1.82E-02 |
| ENSG00000188959 | C9orf152 | 0.68 | 0.34 | 7.76E-03 | 2.01E-02 |
| ENSG00000224358 | TMCO1-AS1 | 0.68 | 0.33 | 7.90E-03 | 2.04E-02 |
| ENSG00000167011 | NAT16 | 0.68 | 0.34 | 8.09E-03 | 2.09E-02 |
| ENSG00000142530 | GARIN5A | 0.68 | 0.34 | 8.43E-03 | 2.17E-02 |
| ENSG00000279568 | | 0.68 | 0.34 | 8.45E-03 | 2.17E-02 |
| ENSG00000281912 | LINC01144 | 0.68 | 0.38 | 1.28E-02 | 3.15E-02 |
| ENSG00000286379 | | 0.68 | 0.4 | 1.63E-02 | 3.91E-02 |
| ENSG00000256732 | | 0.68 | 0.74 | 1.65E-02 | 3.95E-02 |
| ENSG00000287710 | | 0.68 | 0.42 | 1.84E-02 | 4.36E-02 |
| ENSG00000183562 | | 0.68 | 0.43 | 1.89E-02 | 4.48E-02 |
| ENSG00000286147 | | 0.68 | 0.43 | 2.00E-02 | 4.72E-02 |
| ENSG00000152291 | TGOLN2 | 0.67 | 0.03 | 1.14E-106 | 7.18E-105 |
| ENSG00000204262 | COL5A2 | 0.67 | 0.04 | 1.17E-74 | 4.40E-73 |
| ENSG00000197081 | IGF2R | 0.67 | 0.04 | 6.66E-62 | 1.92E-60 |
| ENSG00000281649 | EBLN3P | 0.67 | 0.04 | 2.94E-57 | 7.78E-56 |
| ENSG00000154945 | ANKRD40 | 0.67 | 0.04 | 4.55E-52 | 1.08E-50 |
| ENSG00000124151 | NCOA3 | 0.67 | 0.04 | 7.79E-52 | 1.83E-50 |
| ENSG00000129083 | COPB1 | 0.67 | 0.05 | 2.43E-50 | 5.50E-49 |
| ENSG00000075223 | SEMA3C | 0.67 | 0.05 | 2.60E-50 | 5.88E-49 |
| ENSG00000054523 | KIF1B | 0.67 | 0.05 | 6.77E-46 | 1.39E-44 |
| ENSG00000134245 | WNT2B | 0.67 | 0.05 | 2.25E-45 | 4.56E-44 |
| ENSG00000203880 | PCMTD2 | 0.67 | 0.05 | 5.37E-41 | 9.97E-40 |
| ENSG00000114200 | BCHE | 0.67 | 0.05 | 2.60E-37 | 4.35E-36 |

| | | | | | |
|---|---|---|---|---|---|
| ENSG00000127314 | RAP1B | 0.67 | 0.06 | 1.85E-34 | 2.84E-33 |
| ENSG00000166398 | GARRE1 | 0.67 | 0.06 | 3.08E-34 | 4.71E-33 |
| ENSG00000088448 | ANKRD10 | 0.67 | 0.06 | 3.52E-32 | 5.07E-31 |
| ENSG00000204524 | ZNF805 | 0.67 | 0.06 | 4.42E-30 | 5.95E-29 |
| ENSG00000135631 | RAB11FIP5 | 0.67 | 0.07 | 1.49E-24 | 1.65E-23 |
| ENSG00000149308 | NPAT | 0.67 | 0.07 | 9.09E-24 | 9.81E-23 |
| ENSG00000116514 | RNF19B | 0.67 | 0.07 | 3.20E-22 | 3.25E-21 |
| ENSG00000146830 | GIGYF1 | 0.67 | 0.07 | 3.87E-21 | 3.78E-20 |
| ENSG00000151151 | IPMK | 0.67 | 0.07 | 2.13E-20 | 2.02E-19 |
| ENSG00000168505 | GBX2 | 0.67 | 0.1 | 3.71E-12 | 2.35E-11 |
| ENSG00000147509 | RGS20 | 0.67 | 0.1 | 9.18E-12 | 5.68E-11 |
| ENSG00000259366 | | 0.67 | 0.1 | 3.18E-11 | 1.91E-10 |
| ENSG00000160685 | ZBTB7B | 0.67 | 0.11 | 4.09E-10 | 2.27E-09 |
| ENSG00000286058 | | 0.67 | 0.11 | 5.77E-10 | 3.17E-09 |
| ENSG00000258744 | | 0.67 | 0.12 | 2.90E-09 | 1.51E-08 |
| ENSG00000170919 | TPT1-AS1 | 0.67 | 0.13 | 6.66E-08 | 3.13E-07 |
| ENSG00000125878 | TCF15 | 0.67 | 0.18 | 2.34E-05 | 8.72E-05 |
| ENSG00000230082 | PRRT3-AS1 | 0.67 | 0.2 | 1.28E-04 | 4.39E-04 |
| ENSG00000196912 | ANKRD36B | 0.67 | 0.2 | 1.37E-04 | 4.70E-04 |
| ENSG00000261888 | | 0.67 | 0.22 | 3.50E-04 | 1.14E-03 |
| ENSG00000104863 | LIN7B | 0.67 | 0.22 | 5.85E-04 | 1.85E-03 |
| ENSG00000133392 | MYH11 | 0.67 | 0.24 | 9.62E-04 | 2.93E-03 |
| ENSG00000176428 | VPS37D | 0.67 | 0.25 | 1.74E-03 | 5.09E-03 |
| ENSG00000260233 | ZNRD2-DT | 0.67 | 0.31 | 5.34E-03 | 1.43E-02 |
| ENSG00000266777 | SH3GL1P1 | 0.67 | 0.31 | 5.69E-03 | 1.51E-02 |
| ENSG00000224281 | SLC25A5-AS1 | 0.67 | 0.31 | 6.03E-03 | 1.60E-02 |
| ENSG00000223396 | RPS10P7 | 0.67 | 0.36 | 1.20E-02 | 2.97E-02 |
| ENSG00000224269 | | 0.67 | 0.38 | 1.48E-02 | 3.60E-02 |
| ENSG00000232810 | TNF | 0.67 | 0.39 | 1.51E-02 | 3.66E-02 |
| ENSG00000110025 | SNX15 | 0.67 | 0.41 | 1.83E-02 | 4.36E-02 |
| ENSG00000237118 | CYP2F2P | 0.67 | 0.43 | 2.06E-02 | 4.83E-02 |
| ENSG00000198074 | AKR1B10 | 0.67 | 0.71 | 2.12E-02 | 4.97E-02 |
| ENSG00000119820 | YIPF4 | 0.66 | 0.04 | 4.57E-57 | 1.21E-55 |
| ENSG00000171867 | PRNP | 0.66 | 0.04 | 5.48E-54 | 1.37E-52 |
| ENSG00000068697 | LAPTM4A | 0.66 | 0.05 | 1.28E-44 | 2.56E-43 |
| ENSG00000254337 | | 0.66 | 0.05 | 6.23E-41 | 1.15E-39 |
| ENSG00000064601 | CTSA | 0.66 | 0.05 | 2.11E-40 | 3.81E-39 |
| ENSG00000140948 | ZCCHC14 | 0.66 | 0.05 | 1.11E-38 | 1.93E-37 |
| ENSG00000167972 | ABCA3 | 0.66 | 0.05 | 1.25E-37 | 2.13E-36 |
| ENSG00000100934 | SEC23A | 0.66 | 0.05 | 2.29E-37 | 3.85E-36 |
| ENSG00000159461 | AMFR | 0.66 | 0.05 | 2.48E-37 | 4.15E-36 |
| ENSG00000138760 | SCARB2 | 0.66 | 0.05 | 1.58E-35 | 2.53E-34 |
| ENSG00000164631 | ZNF12 | 0.66 | 0.06 | 3.22E-33 | 4.78E-32 |
| ENSG00000116044 | NFE2L2 | 0.66 | 0.06 | 7.55E-32 | 1.07E-30 |
| ENSG00000115239 | ASB3 | 0.66 | 0.06 | 1.77E-31 | 2.50E-30 |
| ENSG00000117614 | SYF2 | 0.66 | 0.06 | 2.31E-30 | 3.15E-29 |
| ENSG00000136802 | LRRC8A | 0.66 | 0.06 | 6.81E-29 | 8.82E-28 |
| ENSG00000140931 | CMTM3 | 0.66 | 0.06 | 1.79E-26 | 2.14E-25 |
| ENSG00000155970 | MICU3 | 0.66 | 0.06 | 2.86E-26 | 3.37E-25 |
| ENSG00000178502 | KLHL11 | 0.66 | 0.06 | 7.36E-26 | 8.55E-25 |

| | | | | | |
|---|---|---|---|---|---|
| ENSG00000137500 | CCDC90B | 0.66 | 0.07 | 7.04E-24 | 7.63E-23 |
| ENSG00000125952 | MAX | 0.66 | 0.07 | 1.39E-22 | 1.43E-21 |
| ENSG00000181027 | FKRP | 0.66 | 0.07 | 5.77E-21 | 5.58E-20 |
| ENSG00000231721 | LINC-PINT | 0.66 | 0.07 | 2.53E-20 | 2.38E-19 |
| ENSG00000181192 | DHTKD1 | 0.66 | 0.08 | 9.88E-18 | 8.30E-17 |
| ENSG00000234072 | GTF3C2-AS2 | 0.66 | 0.08 | 1.64E-17 | 1.36E-16 |
| ENSG00000090581 | GNPTG | 0.66 | 0.08 | 6.53E-16 | 5.06E-15 |
| ENSG00000204406 | MBD5 | 0.66 | 0.08 | 6.74E-16 | 5.21E-15 |
| ENSG00000011478 | QPCTL | 0.66 | 0.09 | 5.79E-15 | 4.29E-14 |
| ENSG00000178764 | ZHX2 | 0.66 | 0.09 | 1.53E-14 | 1.11E-13 |
| ENSG00000183496 | MEX3B | 0.66 | 0.09 | 1.25E-13 | 8.65E-13 |
| ENSG00000105419 | MEIS3 | 0.66 | 0.1 | 1.04E-11 | 6.43E-11 |
| ENSG00000126458 | RRAS | 0.66 | 0.1 | 6.10E-11 | 3.58E-10 |
| ENSG00000196604 | POTEF | 0.66 | 0.11 | 8.68E-11 | 5.05E-10 |
| ENSG00000159753 | CARMIL2 | 0.66 | 0.12 | 6.07E-09 | 3.09E-08 |
| ENSG00000177301 | KCNA2 | 0.66 | 0.13 | 3.94E-08 | 1.88E-07 |
| ENSG00000165804 | ZNF219 | 0.66 | 0.13 | 5.36E-08 | 2.54E-07 |
| ENSG00000163463 | KRTCAP2 | 0.66 | 0.14 | 4.10E-07 | 1.81E-06 |
| ENSG00000002330 | BAD | 0.66 | 0.19 | 9.42E-05 | 3.28E-04 |
| ENSG00000179698 | WDR97 | 0.66 | 0.19 | 1.09E-04 | 3.77E-04 |
| ENSG00000233461 | | 0.66 | 0.19 | 1.35E-04 | 4.61E-04 |
| ENSG00000162300 | ZFPL1 | 0.66 | 0.22 | 5.91E-04 | 1.86E-03 |
| ENSG00000274471 | | 0.66 | 0.22 | 6.23E-04 | 1.96E-03 |
| ENSG00000205456 | TP53TG3D | 0.66 | 0.24 | 1.36E-03 | 4.05E-03 |
| ENSG00000251361 | | 0.66 | 0.27 | 3.01E-03 | 8.46E-03 |
| ENSG00000287168 | | 0.66 | 0.28 | 3.85E-03 | 1.06E-02 |
| ENSG00000226889 | ATF6-DT | 0.66 | 0.28 | 4.10E-03 | 1.12E-02 |
| ENSG00000183760 | ACP7 | 0.66 | 0.29 | 4.40E-03 | 1.19E-02 |
| ENSG00000234630 | | 0.66 | 0.34 | 1.04E-02 | 2.62E-02 |
| ENSG00000226450 | CYP2D8P | 0.66 | 0.34 | 1.09E-02 | 2.74E-02 |
| ENSG00000237529 | | 0.66 | 0.42 | 2.13E-02 | 4.98E-02 |
| ENSG00000111961 | SASH1 | 0.65 | 0.03 | 4.43E-84 | 1.97E-82 |
| ENSG00000137504 | CREBZF | 0.65 | 0.05 | 2.10E-46 | 4.36E-45 |
| ENSG00000071553 | ATP6AP1 | 0.65 | 0.05 | 1.55E-41 | 2.90E-40 |
| ENSG00000150712 | MTMR12 | 0.65 | 0.05 | 4.91E-35 | 7.70E-34 |
| ENSG00000173334 | TRIB1 | 0.65 | 0.06 | 7.55E-29 | 9.75E-28 |
| ENSG00000197780 | TAF13 | 0.65 | 0.06 | 9.50E-29 | 1.22E-27 |
| ENSG00000142867 | BCL10 | 0.65 | 0.06 | 1.06E-28 | 1.36E-27 |
| ENSG00000204388 | HSPA1B | 0.65 | 0.06 | 1.06E-24 | 1.18E-23 |
| ENSG00000168159 | RNF187 | 0.65 | 0.07 | 6.86E-24 | 7.44E-23 |
| ENSG00000178252 | WDR6 | 0.65 | 0.07 | 6.64E-21 | 6.41E-20 |
| ENSG00000159596 | TMEM69 | 0.65 | 0.07 | 3.58E-20 | 3.36E-19 |
| ENSG00000146066 | HIGD2A | 0.65 | 0.07 | 2.01E-19 | 1.82E-18 |
| ENSG00000100226 | GTPBP1 | 0.65 | 0.07 | 3.26E-19 | 2.93E-18 |
| ENSG00000143373 | ZNF687 | 0.65 | 0.08 | 2.69E-18 | 2.32E-17 |
| ENSG00000099814 | CEP170B | 0.65 | 0.09 | 6.45E-15 | 4.77E-14 |
| ENSG00000089639 | GMIP | 0.65 | 0.09 | 1.21E-13 | 8.41E-13 |
| ENSG00000100116 | GCAT | 0.65 | 0.09 | 2.53E-13 | 1.72E-12 |
| ENSG00000173258 | ZNF483 | 0.65 | 0.11 | 2.54E-10 | 1.43E-09 |
| ENSG00000115844 | DLX2 | 0.65 | 0.12 | 2.08E-08 | 1.02E-07 |

| | | | | | |
|---|---|---|---|---|---|
| ENSG00000131116 | ZNF428 | 0.65 | 0.12 | 2.62E-08 | 1.27E-07 |
| ENSG00000141295 | SCRN2 | 0.65 | 0.13 | 5.18E-08 | 2.46E-07 |
| ENSG00000181722 | ZBTB20 | 0.65 | 0.13 | 2.52E-07 | 1.13E-06 |
| ENSG00000235316 | DUSP8P5 | 0.65 | 0.15 | 4.14E-06 | 1.66E-05 |
| ENSG00000161328 | LRRC56 | 0.65 | 0.16 | 6.30E-06 | 2.49E-05 |
| ENSG00000259877 | ZNF778-DT | 0.65 | 0.18 | 7.07E-05 | 2.51E-04 |
| ENSG00000259802 | MARCHF6-DT | 0.65 | 0.21 | 3.82E-04 | 1.23E-03 |
| ENSG00000158769 | F11R | 0.65 | 0.23 | 9.66E-04 | 2.95E-03 |
| ENSG00000185291 | IL3RA | 0.65 | 0.25 | 2.26E-03 | 6.49E-03 |
| ENSG00000273295 | | 0.65 | 0.25 | 2.27E-03 | 6.51E-03 |
| ENSG00000085563 | ABCB1 | 0.65 | 0.31 | 7.05E-03 | 1.84E-02 |
| ENSG00000242902 | FLNC-AS1 | 0.65 | 0.32 | 7.87E-03 | 2.04E-02 |
| ENSG00000246898 | LINC00920 | 0.65 | 0.32 | 9.29E-03 | 2.37E-02 |
| ENSG00000248278 | SUMO2P17 | 0.65 | 0.33 | 9.58E-03 | 2.43E-02 |
| ENSG00000175773 | ZBTB44-DT | 0.65 | 0.78 | 1.06E-02 | 2.65E-02 |
| ENSG00000229325 | ACAP2-IT1 | 0.65 | 0.33 | 1.08E-02 | 2.71E-02 |
| ENSG00000109758 | HGFAC | 0.65 | 0.35 | 1.28E-02 | 3.16E-02 |
| ENSG00000253948 | VPS13B-DT | 0.65 | 0.36 | 1.47E-02 | 3.57E-02 |
| ENSG00000279491 | | 0.65 | 0.75 | 1.58E-02 | 3.82E-02 |
| ENSG00000058262 | SEC61A1 | 0.64 | 0.04 | 6.95E-73 | 2.52E-71 |
| ENSG00000058063 | ATP11B | 0.64 | 0.04 | 6.60E-55 | 1.68E-53 |
| ENSG00000113163 | CERT1 | 0.64 | 0.04 | 1.53E-48 | 3.31E-47 |
| ENSG00000169016 | E2F6 | 0.64 | 0.05 | 1.64E-40 | 2.98E-39 |
| ENSG00000103978 | TMEM87A | 0.64 | 0.05 | 3.15E-34 | 4.80E-33 |
| ENSG00000102580 | DNAJC3 | 0.64 | 0.05 | 8.75E-34 | 1.32E-32 |
| ENSG00000068400 | GRIPAP1 | 0.64 | 0.05 | 2.42E-32 | 3.49E-31 |
| ENSG00000092108 | SCFD1 | 0.64 | 0.06 | 5.76E-31 | 8.03E-30 |
| ENSG00000132485 | ZRANB2 | 0.64 | 0.06 | 6.87E-31 | 9.54E-30 |
| ENSG00000140995 | DEF8 | 0.64 | 0.06 | 4.54E-27 | 5.52E-26 |
| ENSG00000254999 | BRK1 | 0.64 | 0.06 | 5.57E-27 | 6.76E-26 |
| ENSG00000160285 | LSS | 0.64 | 0.07 | 1.48E-22 | 1.52E-21 |
| ENSG00000177125 | ZBTB34 | 0.64 | 0.07 | 1.95E-22 | 2.00E-21 |
| ENSG00000140598 | EFL1 | 0.64 | 0.07 | 1.85E-21 | 1.83E-20 |
| ENSG00000146083 | RNF44 | 0.64 | 0.07 | 4.38E-21 | 4.26E-20 |
| ENSG00000141480 | ARRB2 | 0.64 | 0.08 | 1.13E-17 | 9.47E-17 |
| ENSG00000244754 | N4BP2L2 | 0.64 | 0.08 | 8.57E-17 | 6.92E-16 |
| ENSG00000134049 | IER3IP1 | 0.64 | 0.08 | 1.33E-15 | 1.01E-14 |
| ENSG00000166987 | MBD6 | 0.64 | 0.09 | 2.82E-14 | 2.03E-13 |
| ENSG00000188522 | FAM83G | 0.64 | 0.09 | 2.13E-13 | 1.45E-12 |
| ENSG00000213614 | HEXA | 0.64 | 0.09 | 6.28E-13 | 4.17E-12 |
| ENSG00000105373 | NOP53 | 0.64 | 0.09 | 8.18E-13 | 5.40E-12 |
| ENSG00000165782 | PIP4P1 | 0.64 | 0.09 | 3.23E-12 | 2.06E-11 |
| ENSG00000130706 | ADRM1 | 0.64 | 0.1 | 2.33E-11 | 1.41E-10 |
| ENSG00000276570 | | 0.64 | 0.11 | 7.48E-10 | 4.08E-09 |
| ENSG00000189298 | ZKSCAN3 | 0.64 | 0.11 | 1.54E-09 | 8.20E-09 |
| ENSG00000174886 | NDUFA11 | 0.64 | 0.13 | 2.17E-07 | 9.80E-07 |
| ENSG00000180089 | TMEM86B | 0.64 | 0.15 | 4.80E-06 | 1.92E-05 |
| ENSG00000231240 | KLF2P1 | 0.64 | 0.16 | 1.56E-05 | 5.90E-05 |
| ENSG00000262903 | CTNS-AS1 | 0.64 | 0.19 | 1.36E-04 | 4.66E-04 |
| ENSG00000263266 | RPS7P1 | 0.64 | 0.19 | 2.00E-04 | 6.71E-04 |

| | | | | | |
|---|---|---|---|---|---|
| ENSG00000135722 | FBXL8 | 0.64 | 0.2 | 3.62E-04 | 1.17E-03 |
| ENSG00000259826 | CDK13-DT | 0.64 | 0.2 | 3.97E-04 | 1.28E-03 |
| ENSG00000205085 | GARIN1A | 0.64 | 0.24 | 1.78E-03 | 5.20E-03 |
| ENSG00000188672 | RHCE | 0.64 | 0.24 | 1.98E-03 | 5.74E-03 |
| ENSG00000231327 | LINC01816 | 0.64 | 0.27 | 4.20E-03 | 1.15E-02 |
| ENSG00000263823 | | 0.64 | 0.3 | 7.38E-03 | 1.92E-02 |
| ENSG00000219200 | RNASEK | 0.64 | 0.31 | 7.71E-03 | 2.00E-02 |
| ENSG00000261542 | | 0.64 | 0.33 | 1.05E-02 | 2.63E-02 |
| ENSG00000263412 | NFE2L1-DT | 0.64 | 0.32 | 1.06E-02 | 2.66E-02 |
| ENSG00000286575 | | 0.64 | 0.77 | 1.13E-02 | 2.83E-02 |
| ENSG00000188981 | MSANTD1 | 0.64 | 0.34 | 1.19E-02 | 2.96E-02 |
| ENSG00000270049 | ATP6V0D1-DT | 0.64 | 0.35 | 1.36E-02 | 3.32E-02 |
| ENSG00000206341 | HLA-H | 0.64 | 0.39 | 1.99E-02 | 4.69E-02 |
| ENSG00000144233 | AMMECR1L | 0.63 | 0.05 | 3.01E-45 | 6.09E-44 |
| ENSG00000160551 | TAOK1 | 0.63 | 0.05 | 6.76E-41 | 1.25E-39 |
| ENSG00000100650 | SRSF5 | 0.63 | 0.05 | 2.48E-35 | 3.94E-34 |
| ENSG00000130589 | HELZ2 | 0.63 | 0.06 | 8.08E-29 | 1.04E-27 |
| ENSG00000175745 | NR2F1 | 0.63 | 0.06 | 1.24E-28 | 1.59E-27 |
| ENSG00000135622 | SEMA4F | 0.63 | 0.06 | 4.47E-27 | 5.44E-26 |
| ENSG00000134107 | BHLHE40 | 0.63 | 0.06 | 6.72E-27 | 8.12E-26 |
| ENSG00000138463 | SLC49A4 | 0.63 | 0.06 | 9.51E-26 | 1.10E-24 |
| ENSG00000135452 | TSPAN31 | 0.63 | 0.06 | 3.23E-24 | 3.54E-23 |
| ENSG00000113851 | CRBN | 0.63 | 0.07 | 2.10E-21 | 2.07E-20 |
| ENSG00000165966 | PDZRN4 | 0.63 | 0.07 | 2.88E-20 | 2.71E-19 |
| ENSG00000214293 | APTR | 0.63 | 0.07 | 5.95E-20 | 5.52E-19 |
| ENSG00000175130 | MARCKSL1 | 0.63 | 0.07 | 1.16E-19 | 1.06E-18 |
| ENSG00000179988 | PSTK | 0.63 | 0.07 | 1.29E-19 | 1.18E-18 |
| ENSG00000164180 | TMEM161B | 0.63 | 0.07 | 3.09E-19 | 2.78E-18 |
| ENSG00000178860 | MSC | 0.63 | 0.07 | 6.88E-19 | 6.07E-18 |
| ENSG00000155256 | ZFYVE27 | 0.63 | 0.07 | 1.03E-18 | 9.01E-18 |
| ENSG00000100344 | PNPLA3 | 0.63 | 0.08 | 1.24E-16 | 9.96E-16 |
| ENSG00000169359 | SLC33A1 | 0.63 | 0.08 | 3.48E-16 | 2.73E-15 |
| ENSG00000167595 | PROSER3 | 0.63 | 0.08 | 3.69E-16 | 2.89E-15 |
| ENSG00000242732 | RTL5 | 0.63 | 0.09 | 2.72E-13 | 1.84E-12 |
| ENSG00000073350 | LLGL2 | 0.63 | 0.09 | 3.20E-12 | 2.04E-11 |
| ENSG00000277363 | SRCIN1 | 0.63 | 0.09 | 5.71E-12 | 3.57E-11 |
| ENSG00000183186 | C2CD4C | 0.63 | 0.12 | 4.27E-08 | 2.04E-07 |
| ENSG00000291153 | STAG3L4 | 0.63 | 0.12 | 4.96E-08 | 2.35E-07 |
| ENSG00000135469 | COQ10A | 0.63 | 0.13 | 2.74E-07 | 1.23E-06 |
| ENSG00000198816 | ZNF358 | 0.63 | 0.14 | 1.20E-06 | 5.06E-06 |
| ENSG00000173805 | HAP1 | 0.63 | 0.15 | 4.08E-06 | 1.64E-05 |
| ENSG00000165698 | SPACA9 | 0.63 | 0.16 | 1.65E-05 | 6.25E-05 |
| ENSG00000231770 | TMEM44-AS1 | 0.63 | 0.16 | 2.07E-05 | 7.77E-05 |
| ENSG00000245573 | BDNF-AS | 0.63 | 0.16 | 2.08E-05 | 7.79E-05 |
| ENSG00000273270 | LINC03072 | 0.63 | 0.17 | 4.44E-05 | 1.60E-04 |
| ENSG00000259583 | ALDH1A3-AS1 | 0.63 | 0.2 | 2.72E-04 | 8.98E-04 |
| ENSG00000173077 | DELEC1 | 0.63 | 0.2 | 4.15E-04 | 1.33E-03 |
| ENSG00000169020 | ATP5ME | 0.63 | 0.21 | 6.58E-04 | 2.06E-03 |
| ENSG00000279278 | | 0.63 | 0.21 | 7.23E-04 | 2.25E-03 |
| ENSG00000129757 | CDKN1C | 0.63 | 0.23 | 1.52E-03 | 4.48E-03 |

| | | | | | |
|---|---|---|---|---|---|
| ENSG00000204583 | LRCOL1 | 0.63 | 0.24 | 2.22E-03 | 6.39E-03 |
| ENSG00000185101 | ANO9 | 0.63 | 0.28 | 4.94E-03 | 1.33E-02 |
| ENSG00000149557 | FEZ1 | 0.63 | 0.28 | 5.17E-03 | 1.39E-02 |
| ENSG00000231331 | | 0.63 | 0.35 | 1.53E-02 | 3.69E-02 |
| ENSG00000262766 | | 0.63 | 0.37 | 1.91E-02 | 4.52E-02 |
| ENSG00000270299 | | 0.63 | 0.38 | 2.10E-02 | 4.93E-02 |
| ENSG00000057019 | DCBLD2 | 0.62 | 0.04 | 3.22E-59 | 8.92E-58 |
| ENSG00000134108 | ARL8B | 0.62 | 0.04 | 3.09E-50 | 6.98E-49 |
| ENSG00000198466 | ZNF587 | 0.62 | 0.04 | 4.46E-50 | 1.00E-48 |
| ENSG00000117481 | NSUN4 | 0.62 | 0.04 | 1.19E-49 | 2.65E-48 |
| ENSG00000086598 | TMED2 | 0.62 | 0.04 | 5.96E-48 | 1.28E-46 |
| ENSG00000140319 | SRP14 | 0.62 | 0.05 | 4.49E-43 | 8.72E-42 |
| ENSG00000108588 | CCDC47 | 0.62 | 0.05 | 2.80E-37 | 4.67E-36 |
| ENSG00000072042 | RDH11 | 0.62 | 0.05 | 8.74E-37 | 1.45E-35 |
| ENSG00000072041 | SLC6A15 | 0.62 | 0.05 | 3.81E-34 | 5.80E-33 |
| ENSG00000139651 | ZNF740 | 0.62 | 0.05 | 6.90E-32 | 9.84E-31 |
| ENSG00000160445 | ZER1 | 0.62 | 0.06 | 1.42E-28 | 1.81E-27 |
| ENSG00000217128 | FNIP1 | 0.62 | 0.06 | 3.15E-27 | 3.86E-26 |
| ENSG00000074935 | TUBE1 | 0.62 | 0.06 | 9.99E-27 | 1.20E-25 |
| ENSG00000121749 | TBC1D15 | 0.62 | 0.06 | 1.39E-24 | 1.55E-23 |
| ENSG00000185621 | LMLN | 0.62 | 0.06 | 3.29E-23 | 3.46E-22 |
| ENSG00000182504 | CEP97 | 0.62 | 0.07 | 1.94E-20 | 1.84E-19 |
| ENSG00000165802 | NSMF | 0.62 | 0.07 | 4.20E-20 | 3.92E-19 |
| ENSG00000162191 | UBXN1 | 0.62 | 0.07 | 6.53E-19 | 5.77E-18 |
| ENSG00000129245 | FXR2 | 0.62 | 0.07 | 2.72E-17 | 2.25E-16 |
| ENSG00000150401 | DCUN1D2 | 0.62 | 0.08 | 2.77E-15 | 2.08E-14 |
| ENSG00000106993 | CDC37L1 | 0.62 | 0.09 | 2.34E-13 | 1.59E-12 |
| ENSG00000124587 | PEX6 | 0.62 | 0.09 | 5.33E-12 | 3.34E-11 |
| ENSG00000182272 | B4GALNT4 | 0.62 | 0.1 | 3.93E-11 | 2.34E-10 |
| ENSG00000136040 | PLXNC1 | 0.62 | 0.1 | 8.27E-11 | 4.82E-10 |
| ENSG00000125657 | TNFSF9 | 0.62 | 0.1 | 9.03E-11 | 5.25E-10 |
| ENSG00000123395 | ATG101 | 0.62 | 0.1 | 1.53E-10 | 8.76E-10 |
| ENSG00000112667 | DNPH1 | 0.62 | 0.1 | 3.67E-10 | 2.04E-09 |
| ENSG00000198556 | ZNF789 | 0.62 | 0.11 | 4.20E-09 | 2.16E-08 |
| ENSG00000147804 | SLC39A4 | 0.62 | 0.12 | 5.37E-08 | 2.54E-07 |
| ENSG00000185338 | SOCS1 | 0.62 | 0.13 | 6.68E-07 | 2.89E-06 |
| ENSG00000285796 | | 0.62 | 0.14 | 3.91E-06 | 1.57E-05 |
| ENSG00000244733 | | 0.62 | 0.15 | 6.70E-06 | 2.64E-05 |
| ENSG00000214389 | RPS3AP26 | 0.62 | 0.16 | 1.69E-05 | 6.37E-05 |
| ENSG00000117425 | PTCH2 | 0.62 | 0.18 | 1.38E-04 | 4.73E-04 |
| ENSG00000185133 | INPP5J | 0.62 | 0.19 | 2.41E-04 | 7.98E-04 |
| ENSG00000265778 | ZNF516-AS1 | 0.62 | 0.19 | 2.73E-04 | 9.01E-04 |
| ENSG00000110723 | EXPH5 | 0.62 | 0.19 | 2.95E-04 | 9.67E-04 |
| ENSG00000239857 | GET4 | 0.62 | 0.19 | 3.18E-04 | 1.04E-03 |
| ENSG00000260051 | | 0.62 | 0.21 | 7.26E-04 | 2.26E-03 |
| ENSG00000009950 | MLXIPL | 0.62 | 0.23 | 1.71E-03 | 4.99E-03 |
| ENSG00000261659 | | 0.62 | 0.24 | 2.62E-03 | 7.44E-03 |
| ENSG00000186940 | CHCHD2P9 | 0.62 | 0.29 | 6.79E-03 | 1.78E-02 |
| ENSG00000260806 | PAPOLA-DT | 0.62 | 0.3 | 7.92E-03 | 2.05E-02 |
| ENSG00000218510 | LINC00339 | 0.62 | 0.3 | 8.71E-03 | 2.23E-02 |

| | | | | | |
|---|---|---|---|---|---|
| ENSG00000123965 | PMS2P5 | 0.62 | 0.3 | 9.62E-03 | 2.44E-02 |
| ENSG00000227540 | DNAJC9-AS1 | 0.62 | 0.33 | 1.34E-02 | 3.28E-02 |
| ENSG00000259218 | LINC00928 | 0.62 | 0.76 | 1.45E-02 | 3.53E-02 |
| ENSG00000140043 | PTGR2 | 0.62 | 0.35 | 1.57E-02 | 3.79E-02 |
| ENSG00000289018 | | 0.62 | 0.35 | 1.73E-02 | 4.13E-02 |
| ENSG00000288759 | | 0.62 | 0.36 | 1.82E-02 | 4.34E-02 |
| ENSG00000275339 | | 0.62 | 0.74 | 1.83E-02 | 4.34E-02 |
| ENSG00000274460 | | 0.62 | 0.36 | 1.85E-02 | 4.39E-02 |
| ENSG00000234230 | ZFX-AS1 | 0.62 | 0.73 | 2.04E-02 | 4.81E-02 |
| ENSG00000142657 | PGD | 0.61 | 0.04 | 3.82E-65 | 1.19E-63 |
| ENSG00000137494 | ANKRD42 | 0.61 | 0.05 | 4.43E-36 | 7.19E-35 |
| ENSG00000078804 | TP53INP2 | 0.61 | 0.05 | 4.14E-32 | 5.95E-31 |
| ENSG00000168795 | ZBTB5 | 0.61 | 0.06 | 1.16E-28 | 1.49E-27 |
| ENSG00000159167 | STC1 | 0.61 | 0.06 | 3.69E-28 | 4.64E-27 |
| ENSG00000179134 | SAMD4B | 0.61 | 0.06 | 2.08E-26 | 2.47E-25 |
| ENSG00000136048 | DRAM1 | 0.61 | 0.06 | 2.18E-24 | 2.40E-23 |
| ENSG00000198689 | SLC9A6 | 0.61 | 0.06 | 6.74E-22 | 6.76E-21 |
| ENSG00000173230 | GOLGB1 | 0.61 | 0.07 | 1.95E-20 | 1.84E-19 |
| ENSG00000161671 | EMC10 | 0.61 | 0.07 | 1.11E-19 | 1.02E-18 |
| ENSG00000165389 | SPTSSA | 0.61 | 0.07 | 3.72E-19 | 3.33E-18 |
| ENSG00000134825 | TMEM258 | 0.61 | 0.07 | 1.05E-17 | 8.78E-17 |
| ENSG00000221944 | TIGD1 | 0.61 | 0.08 | 1.02E-16 | 8.17E-16 |
| ENSG00000251003 | ZFPM2-AS1 | 0.61 | 0.09 | 1.11E-12 | 7.27E-12 |
| ENSG00000141519 | CCDC40 | 0.61 | 0.09 | 2.51E-12 | 1.61E-11 |
| ENSG00000130770 | ATP5IF1 | 0.61 | 0.09 | 1.12E-11 | 6.92E-11 |
| ENSG00000181135 | ZNF707 | 0.61 | 0.09 | 2.55E-11 | 1.54E-10 |
| ENSG00000176209 | SMIM19 | 0.61 | 0.1 | 7.11E-10 | 3.89E-09 |
| ENSG00000214413 | BBIP1 | 0.61 | 0.1 | 7.29E-10 | 3.98E-09 |
| ENSG00000015568 | RGPD5 | 0.61 | 0.1 | 8.32E-10 | 4.52E-09 |
| ENSG00000236104 | ZBTB22 | 0.61 | 0.11 | 1.83E-09 | 9.69E-09 |
| ENSG00000256060 | TRAPPC2B | 0.61 | 0.11 | 6.07E-09 | 3.08E-08 |
| ENSG00000248049 | UBA6-DT | 0.61 | 0.11 | 7.52E-09 | 3.80E-08 |
| ENSG00000125652 | ALKBH7 | 0.61 | 0.12 | 1.88E-07 | 8.54E-07 |
| ENSG00000165948 | IFI27L1 | 0.61 | 0.13 | 4.62E-07 | 2.03E-06 |
| ENSG00000235217 | TSPY26P | 0.61 | 0.13 | 8.35E-07 | 3.57E-06 |
| ENSG00000174292 | TNK1 | 0.61 | 0.14 | 5.01E-06 | 2.00E-05 |
| ENSG00000124074 | ENKD1 | 0.61 | 0.14 | 5.16E-06 | 2.06E-05 |
| ENSG00000197568 | HHLA3 | 0.61 | 0.14 | 5.20E-06 | 2.07E-05 |
| ENSG00000174007 | CEP19 | 0.61 | 0.14 | 5.45E-06 | 2.17E-05 |
| ENSG00000168143 | FAM83B | 0.61 | 0.15 | 7.14E-06 | 2.80E-05 |
| ENSG00000245146 | MALINC1 | 0.61 | 0.15 | 8.87E-06 | 3.44E-05 |
| ENSG00000268996 | MAN1B1-DT | 0.61 | 0.15 | 1.51E-05 | 5.75E-05 |
| ENSG00000166707 | ZCCHC18 | 0.61 | 0.17 | 9.21E-05 | 3.21E-04 |
| ENSG00000197558 | SSPOP | 0.61 | 0.19 | 3.94E-04 | 1.27E-03 |
| ENSG00000153982 | GDPD1 | 0.61 | 0.2 | 5.20E-04 | 1.65E-03 |
| ENSG00000249492 | | 0.61 | 0.2 | 6.52E-04 | 2.04E-03 |
| ENSG00000234028 | EIF2AK3-DT | 0.61 | 0.21 | 9.87E-04 | 3.00E-03 |
| ENSG00000271576 | PRKACB-DT | 0.61 | 0.21 | 9.88E-04 | 3.01E-03 |
| ENSG00000290415 | | 0.61 | 0.22 | 1.26E-03 | 3.76E-03 |
| ENSG00000234771 | SLC25A25-AS1 | 0.61 | 0.22 | 1.33E-03 | 3.95E-03 |

| | | | | | |
|---|---|---|---|---|---|
| ENSG00000270504 | | 0.61 | 0.22 | 1.34E-03 | 4.00E-03 |
| ENSG00000276931 | | 0.61 | 0.25 | 4.24E-03 | 1.16E-02 |
| ENSG00000249456 | | 0.61 | 0.28 | 7.19E-03 | 1.88E-02 |
| ENSG00000256546 | LINC02985 | 0.61 | 0.29 | 7.58E-03 | 1.97E-02 |
| ENSG00000206567 | | 0.61 | 0.29 | 7.70E-03 | 2.00E-02 |
| ENSG00000279611 | | 0.61 | 0.32 | 1.26E-02 | 3.10E-02 |
| ENSG00000289009 | | 0.61 | 0.32 | 1.36E-02 | 3.32E-02 |
| ENSG00000272023 | AFF4-DT | 0.61 | 0.75 | 1.65E-02 | 3.95E-02 |
| ENSG00000104140 | RHOV | 0.61 | 0.34 | 1.72E-02 | 4.12E-02 |
| ENSG00000231193 | | 0.61 | 0.74 | 1.98E-02 | 4.67E-02 |
| ENSG00000069812 | HES2 | 0.61 | 0.74 | 2.05E-02 | 4.81E-02 |
| ENSG00000286460 | | 0.61 | 0.73 | 2.09E-02 | 4.90E-02 |
| ENSG00000122218 | COPA | 0.6 | 0.03 | 4.97E-89 | 2.39E-87 |
| ENSG00000115806 | GORASP2 | 0.6 | 0.03 | 2.90E-68 | 9.53E-67 |
| ENSG00000113384 | GOLPH3 | 0.6 | 0.04 | 1.87E-46 | 3.89E-45 |
| ENSG00000101191 | DIDO1 | 0.6 | 0.04 | 1.10E-43 | 2.17E-42 |
| ENSG00000198911 | SREBF2 | 0.6 | 0.05 | 1.13E-33 | 1.71E-32 |
| ENSG00000159199 | ATP5MC1 | 0.6 | 0.05 | 4.31E-28 | 5.40E-27 |
| ENSG00000147364 | FBXO25 | 0.6 | 0.06 | 2.01E-25 | 2.31E-24 |
| ENSG00000164609 | SLU7 | 0.6 | 0.06 | 3.28E-25 | 3.73E-24 |
| ENSG00000154640 | BTG3 | 0.6 | 0.06 | 3.66E-24 | 4.00E-23 |
| ENSG00000185359 | HGS | 0.6 | 0.06 | 6.56E-22 | 6.59E-21 |
| ENSG00000198909 | MAP3K3 | 0.6 | 0.06 | 7.54E-22 | 7.54E-21 |
| ENSG00000108523 | RNF167 | 0.6 | 0.06 | 1.05E-21 | 1.05E-20 |
| ENSG00000082996 | RNF13 | 0.6 | 0.06 | 3.48E-21 | 3.41E-20 |
| ENSG00000106400 | ZNHIT1 | 0.6 | 0.07 | 3.87E-20 | 3.62E-19 |
| ENSG00000059728 | MXD1 | 0.6 | 0.07 | 9.70E-20 | 8.93E-19 |
| ENSG00000130827 | PLXNA3 | 0.6 | 0.07 | 1.23E-19 | 1.13E-18 |
| ENSG00000183691 | NOG | 0.6 | 0.07 | 1.81E-19 | 1.64E-18 |
| ENSG00000172497 | ACOT12 | 0.6 | 0.08 | 2.36E-15 | 1.78E-14 |
| ENSG00000116017 | ARID3A | 0.6 | 0.08 | 7.04E-15 | 5.19E-14 |
| ENSG00000135185 | TMEM243 | 0.6 | 0.08 | 1.84E-14 | 1.33E-13 |
| ENSG00000268362 | | 0.6 | 0.08 | 1.14E-13 | 7.91E-13 |
| ENSG00000133275 | CSNK1G2 | 0.6 | 0.08 | 3.21E-13 | 2.17E-12 |
| ENSG00000137821 | LRRC49 | 0.6 | 0.09 | 4.46E-13 | 2.99E-12 |
| ENSG00000128311 | TST | 0.6 | 0.09 | 6.67E-11 | 3.91E-10 |
| ENSG00000058453 | CROCC | 0.6 | 0.1 | 1.17E-10 | 6.76E-10 |
| ENSG00000182809 | CRIP2 | 0.6 | 0.1 | 2.04E-10 | 1.16E-09 |
| ENSG00000149823 | VPS51 | 0.6 | 0.1 | 6.51E-10 | 3.57E-09 |
| ENSG00000197070 | ARRDC1 | 0.6 | 0.11 | 4.48E-09 | 2.30E-08 |
| ENSG00000167554 | ZNF610 | 0.6 | 0.11 | 7.88E-09 | 3.97E-08 |
| ENSG00000142279 | WTIP | 0.6 | 0.11 | 1.45E-08 | 7.19E-08 |
| ENSG00000228474 | OST4 | 0.6 | 0.11 | 2.55E-08 | 1.24E-07 |
| ENSG00000224870 | MRPL20-AS1 | 0.6 | 0.12 | 8.60E-08 | 4.01E-07 |
| ENSG00000280355 | | 0.6 | 0.12 | 1.47E-07 | 6.73E-07 |
| ENSG00000114670 | NEK11 | 0.6 | 0.12 | 2.31E-07 | 1.04E-06 |
| ENSG00000182108 | DEXI | 0.6 | 0.14 | 4.39E-06 | 1.76E-05 |
| ENSG00000233611 | GBX2-AS1 | 0.6 | 0.15 | 2.16E-05 | 8.07E-05 |
| ENSG00000103599 | IQCH | 0.6 | 0.16 | 4.96E-05 | 1.78E-04 |
| ENSG00000288829 | | 0.6 | 0.17 | 1.43E-04 | 4.87E-04 |

| | | | | | |
|---|---|---|---|---|---|
| ENSG00000291175 | | 0.6 | 0.21 | 9.12E-04 | 2.79E-03 |
| ENSG00000275223 | | 0.6 | 0.22 | 2.03E-03 | 5.88E-03 |
| ENSG00000221184 | | 0.6 | 0.24 | 3.00E-03 | 8.43E-03 |
| ENSG00000250993 | | 0.6 | 0.24 | 3.35E-03 | 9.33E-03 |
| ENSG00000260924 | LINC01311 | 0.6 | 0.25 | 3.94E-03 | 1.08E-02 |
| ENSG00000225793 | TMEM30A-DT | 0.6 | 0.26 | 4.88E-03 | 1.32E-02 |
| ENSG00000254893 | RAP1BL | 0.6 | 0.26 | 4.93E-03 | 1.33E-02 |
| ENSG00000162482 | AKR7A3 | 0.6 | 0.28 | 7.73E-03 | 2.00E-02 |
| ENSG00000269318 | | 0.6 | 0.3 | 1.03E-02 | 2.60E-02 |
| ENSG00000170044 | ZPLD1 | 0.6 | 0.3 | 1.07E-02 | 2.69E-02 |
| ENSG00000273148 | LCDR | 0.6 | 0.3 | 1.14E-02 | 2.84E-02 |
| ENSG00000198022 | SAGE2P | 0.6 | 0.3 | 1.14E-02 | 2.85E-02 |
| ENSG00000285884 | | 0.6 | 0.31 | 1.21E-02 | 3.00E-02 |
| ENSG00000268883 | PNMA6B | 0.6 | 0.31 | 1.27E-02 | 3.12E-02 |
| ENSG00000236519 | LINC01424 | 0.6 | 0.31 | 1.32E-02 | 3.24E-02 |
| ENSG00000188869 | TMC3 | 0.6 | 0.76 | 1.52E-02 | 3.68E-02 |
| ENSG00000278341 | | 0.6 | 0.33 | 1.68E-02 | 4.02E-02 |
| ENSG00000290699 | CYP4F30P | 0.6 | 0.75 | 1.68E-02 | 4.02E-02 |
| ENSG00000201643 | SNORA14A | 0.6 | 0.74 | 1.84E-02 | 4.36E-02 |
| ENSG00000124467 | PSG8 | 0.6 | 0.74 | 2.00E-02 | 4.70E-02 |
| ENSG00000103056 | SMPD3 | 0.6 | 0.35 | 2.05E-02 | 4.82E-02 |
| ENSG00000236383 | CCDC200 | 0.6 | 0.35 | 2.07E-02 | 4.86E-02 |
| ENSG00000157954 | WIPI2 | 0.59 | 0.04 | 6.24E-52 | 1.47E-50 |
| ENSG00000160213 | CSTB | 0.59 | 0.05 | 4.11E-38 | 7.06E-37 |
| ENSG00000176142 | TMEM39A | 0.59 | 0.05 | 4.55E-38 | 7.80E-37 |
| ENSG00000187605 | TET3 | 0.59 | 0.05 | 1.78E-36 | 2.91E-35 |
| ENSG00000136888 | ATP6V1G1 | 0.59 | 0.05 | 6.29E-36 | 1.02E-34 |
| ENSG00000138413 | IDH1 | 0.59 | 0.05 | 3.80E-35 | 5.97E-34 |
| ENSG00000089248 | ERP29 | 0.59 | 0.05 | 2.27E-27 | 2.79E-26 |
| ENSG00000163516 | ANKZF1 | 0.59 | 0.06 | 1.47E-25 | 1.69E-24 |
| ENSG00000132424 | PNISR | 0.59 | 0.06 | 4.56E-21 | 4.43E-20 |
| ENSG00000164615 | CAMLG | 0.59 | 0.07 | 2.78E-20 | 2.62E-19 |
| ENSG00000022567 | SLC45A4 | 0.59 | 0.07 | 3.60E-17 | 2.96E-16 |
| ENSG00000172932 | ANKRD13D | 0.59 | 0.07 | 1.85E-16 | 1.47E-15 |
| ENSG00000165185 | KIAA1958 | 0.59 | 0.07 | 3.20E-16 | 2.52E-15 |
| ENSG00000173559 | NABP1 | 0.59 | 0.08 | 7.25E-16 | 5.59E-15 |
| ENSG00000069974 | RAB27A | 0.59 | 0.08 | 4.09E-14 | 2.90E-13 |
| ENSG00000164713 | BRI3 | 0.59 | 0.08 | 4.16E-14 | 2.95E-13 |
| ENSG00000063180 | CA11 | 0.59 | 0.08 | 8.33E-14 | 5.81E-13 |
| ENSG00000158286 | RNF207 | 0.59 | 0.1 | 5.93E-10 | 3.26E-09 |
| ENSG00000170537 | TMC7 | 0.59 | 0.1 | 3.84E-09 | 1.98E-08 |
| ENSG00000160284 | SPATC1L | 0.59 | 0.11 | 1.31E-08 | 6.49E-08 |
| ENSG00000172366 | MCRIP2 | 0.59 | 0.12 | 3.13E-07 | 1.40E-06 |
| ENSG00000185885 | IFITM1 | 0.59 | 0.13 | 2.94E-06 | 1.19E-05 |
| ENSG00000257261 | SLC38A4-AS1 | 0.59 | 0.14 | 6.48E-06 | 2.55E-05 |
| ENSG00000175938 | ORAI3 | 0.59 | 0.15 | 2.41E-05 | 8.95E-05 |
| ENSG00000167136 | ENDOG | 0.59 | 0.2 | 8.57E-04 | 2.63E-03 |
| ENSG00000291147 | | 0.59 | 0.21 | 1.12E-03 | 3.38E-03 |
| ENSG00000255224 | | 0.59 | 0.22 | 1.90E-03 | 5.54E-03 |
| ENSG00000066230 | SLC9A3 | 0.59 | 0.23 | 3.11E-03 | 8.73E-03 |

| | | | | | |
|---|---|---|---|---|---|
| ENSG00000168491 | CCDC110 | 0.59 | 0.25 | 4.69E-03 | 1.27E-02 |
| ENSG00000274372 | LINC02804 | 0.59 | 0.26 | 6.85E-03 | 1.79E-02 |
| ENSG00000185522 | LMNTD2 | 0.59 | 0.28 | 8.74E-03 | 2.24E-02 |
| ENSG00000289135 | | 0.59 | 0.28 | 9.13E-03 | 2.33E-02 |
| ENSG00000253320 | MAILR | 0.59 | 0.32 | 1.63E-02 | 3.92E-02 |
| ENSG00000272620 | AFAP1-AS1 | 0.59 | 0.34 | 2.06E-02 | 4.85E-02 |
| ENSG00000136240 | KDELR2 | 0.58 | 0.03 | 5.45E-69 | 1.83E-67 |
| ENSG00000129657 | SEC14L1 | 0.58 | 0.04 | 1.63E-50 | 3.72E-49 |
| ENSG00000184014 | DENND5A | 0.58 | 0.04 | 3.20E-40 | 5.76E-39 |
| ENSG00000080815 | PSEN1 | 0.58 | 0.05 | 5.15E-34 | 7.81E-33 |
| ENSG00000127481 | UBR4 | 0.58 | 0.05 | 6.49E-33 | 9.53E-32 |
| ENSG00000119328 | ABITRAM | 0.58 | 0.05 | 2.23E-32 | 3.23E-31 |
| ENSG00000059758 | CDK17 | 0.58 | 0.05 | 5.63E-32 | 8.04E-31 |
| ENSG00000137941 | TTLL7 | 0.58 | 0.06 | 6.36E-26 | 7.41E-25 |
| ENSG00000135968 | GCC2 | 0.58 | 0.06 | 3.71E-20 | 3.48E-19 |
| ENSG00000179941 | BBS10 | 0.58 | 0.06 | 1.09E-19 | 9.99E-19 |
| ENSG00000164114 | MAP9 | 0.58 | 0.07 | 2.00E-18 | 1.73E-17 |
| ENSG00000117410 | ATP6V0B | 0.58 | 0.07 | 7.91E-16 | 6.08E-15 |
| ENSG00000198087 | CD2AP | 0.58 | 0.07 | 8.30E-16 | 6.38E-15 |
| ENSG00000205133 | TRIQK | 0.58 | 0.08 | 1.77E-13 | 1.22E-12 |
| ENSG00000101460 | MAP1LC3A | 0.58 | 0.08 | 2.00E-13 | 1.37E-12 |
| ENSG00000088727 | KIF9 | 0.58 | 0.09 | 2.41E-11 | 1.45E-10 |
| ENSG00000121104 | FAM117A | 0.58 | 0.09 | 6.07E-11 | 3.57E-10 |
| ENSG00000243147 | MRPL33 | 0.58 | 0.11 | 9.68E-09 | 4.86E-08 |
| ENSG00000165810 | BTNL9 | 0.58 | 0.11 | 1.20E-08 | 5.98E-08 |
| ENSG00000170222 | ADPRM | 0.58 | 0.12 | 7.07E-07 | 3.05E-06 |
| ENSG00000118503 | TNFAIP3 | 0.58 | 0.13 | 3.83E-06 | 1.54E-05 |
| ENSG00000174939 | ASPHD1 | 0.58 | 0.14 | 4.24E-06 | 1.70E-05 |
| ENSG00000274828 | | 0.58 | 0.14 | 8.23E-06 | 3.21E-05 |
| ENSG00000060140 | STYK1 | 0.58 | 0.14 | 1.11E-05 | 4.27E-05 |
| ENSG00000279041 | | 0.58 | 0.14 | 1.52E-05 | 5.78E-05 |
| ENSG00000205464 | ATP6AP1L | 0.58 | 0.15 | 2.43E-05 | 9.01E-05 |
| ENSG00000290872 | | 0.58 | 0.16 | 1.14E-04 | 3.93E-04 |
| ENSG00000213199 | ASIC3 | 0.58 | 0.17 | 1.77E-04 | 5.98E-04 |
| ENSG00000163703 | CRELD1 | 0.58 | 0.19 | 5.76E-04 | 1.82E-03 |
| ENSG00000249936 | RAC1P2 | 0.58 | 0.19 | 6.09E-04 | 1.92E-03 |
| ENSG00000181513 | ACBD4 | 0.58 | 0.19 | 7.36E-04 | 2.28E-03 |
| ENSG00000015479 | MATR3 | 0.58 | 0.2 | 9.25E-04 | 2.83E-03 |
| ENSG00000175170 | FAM182B | 0.58 | 0.22 | 2.51E-03 | 7.15E-03 |
| ENSG00000239382 | ALKBH6 | 0.58 | 0.22 | 2.56E-03 | 7.29E-03 |
| ENSG00000287576 | | 0.58 | 0.25 | 6.03E-03 | 1.60E-02 |
| ENSG00000279539 | | 0.58 | 0.27 | 8.22E-03 | 2.12E-02 |
| ENSG00000167617 | CDC42EP5 | 0.58 | 0.27 | 8.73E-03 | 2.24E-02 |
| ENSG00000108950 | FAM20A | 0.58 | 0.3 | 1.52E-02 | 3.67E-02 |
| ENSG00000168000 | BSCL2 | 0.58 | 0.31 | 1.53E-02 | 3.71E-02 |
| ENSG00000272894 | MIOS-DT | 0.58 | 0.31 | 1.61E-02 | 3.86E-02 |
| ENSG00000111846 | GCNT2 | 0.58 | 0.32 | 1.67E-02 | 3.99E-02 |
| ENSG00000251450 | RASGRF2-AS1 | 0.58 | 0.31 | 1.67E-02 | 4.00E-02 |
| ENSG00000228022 | HCG20 | 0.58 | 0.75 | 1.67E-02 | 4.01E-02 |
| ENSG00000271918 | | 0.58 | 0.31 | 1.68E-02 | 4.02E-02 |

| | | | | | |
|---|---|---|---|---|---|
| ENSG00000236991 | EDRF1-AS1 | 0.58 | 0.32 | 1.90E-02 | 4.49E-02 |
| ENSG00000181378 | CFAP65 | 0.58 | 0.33 | 2.01E-02 | 4.73E-02 |
| ENSG00000170558 | CDH2 | 0.57 | 0.04 | 2.03E-57 | 5.42E-56 |
| ENSG00000197746 | PSAP | 0.57 | 0.04 | 4.23E-54 | 1.06E-52 |
| ENSG00000100201 | DDX17 | 0.57 | 0.04 | 1.64E-50 | 3.72E-49 |
| ENSG00000001629 | ANKIB1 | 0.57 | 0.04 | 6.62E-42 | 1.25E-40 |
| ENSG00000170385 | SLC30A1 | 0.57 | 0.04 | 1.33E-40 | 2.44E-39 |
| ENSG00000147535 | PLPP5 | 0.57 | 0.04 | 2.37E-40 | 4.27E-39 |
| ENSG00000077044 | DGKD | 0.57 | 0.05 | 5.35E-37 | 8.87E-36 |
| ENSG00000140650 | PMM2 | 0.57 | 0.05 | 4.34E-34 | 6.59E-33 |
| ENSG00000050165 | DKK3 | 0.57 | 0.05 | 3.16E-33 | 4.69E-32 |
| ENSG00000176171 | BNIP3 | 0.57 | 0.05 | 5.05E-33 | 7.44E-32 |
| ENSG00000203879 | GDI1 | 0.57 | 0.05 | 6.82E-32 | 9.72E-31 |
| ENSG00000167110 | GOLGA2 | 0.57 | 0.05 | 2.56E-27 | 3.14E-26 |
| ENSG00000266472 | MRPS21 | 0.57 | 0.05 | 2.93E-27 | 3.59E-26 |
| ENSG00000143198 | MGST3 | 0.57 | 0.05 | 1.20E-26 | 1.45E-25 |
| ENSG00000131871 | SELENOS | 0.57 | 0.06 | 5.36E-25 | 6.05E-24 |
| ENSG00000214655 | ZSWIM8 | 0.57 | 0.06 | 3.27E-20 | 3.07E-19 |
| ENSG00000163931 | TKT | 0.57 | 0.06 | 3.35E-20 | 3.14E-19 |
| ENSG00000205423 | CNEP1R1 | 0.57 | 0.07 | 6.63E-18 | 5.61E-17 |
| ENSG00000006652 | IFRD1 | 0.57 | 0.07 | 5.58E-16 | 4.33E-15 |
| ENSG00000116717 | GADD45A | 0.57 | 0.07 | 1.04E-15 | 7.95E-15 |
| ENSG00000156232 | WHAMM | 0.57 | 0.08 | 1.67E-14 | 1.21E-13 |
| ENSG00000075240 | GRAMD4 | 0.57 | 0.08 | 2.55E-13 | 1.73E-12 |
| ENSG00000180979 | LRRC57 | 0.57 | 0.08 | 3.45E-13 | 2.32E-12 |
| ENSG00000047634 | SCML1 | 0.57 | 0.08 | 1.79E-12 | 1.16E-11 |
| ENSG00000143774 | GUK1 | 0.57 | 0.09 | 7.38E-12 | 4.58E-11 |
| ENSG00000164117 | FBXO8 | 0.57 | 0.09 | 1.34E-11 | 8.23E-11 |
| ENSG00000196981 | WDR5B | 0.57 | 0.09 | 2.00E-11 | 1.21E-10 |
| ENSG00000130584 | ZBTB46 | 0.57 | 0.09 | 1.57E-10 | 8.98E-10 |
| ENSG00000185507 | IRF7 | 0.57 | 0.09 | 2.28E-10 | 1.29E-09 |
| ENSG00000169570 | DTWD2 | 0.57 | 0.1 | 8.90E-10 | 4.82E-09 |
| ENSG00000142330 | CAPN10 | 0.57 | 0.1 | 8.46E-09 | 4.26E-08 |
| ENSG00000203930 | LINC00632 | 0.57 | 0.11 | 5.42E-08 | 2.57E-07 |
| ENSG00000179240 | GVQW3 | 0.57 | 0.11 | 7.11E-08 | 3.34E-07 |
| ENSG00000260630 | SNAI3-AS1 | 0.57 | 0.12 | 7.26E-07 | 3.13E-06 |
| ENSG00000228630 | HOTAIR | 0.57 | 0.12 | 8.45E-07 | 3.61E-06 |
| ENSG00000111186 | WNT5B | 0.57 | 0.13 | 2.17E-06 | 8.93E-06 |
| ENSG00000196387 | ZNF140 | 0.57 | 0.14 | 1.55E-05 | 5.87E-05 |
| ENSG00000139537 | CCDC65 | 0.57 | 0.16 | 8.06E-05 | 2.84E-04 |
| ENSG00000268230 | | 0.57 | 0.16 | 9.36E-05 | 3.27E-04 |
| ENSG00000285053 | | 0.57 | 0.17 | 2.69E-04 | 8.87E-04 |
| ENSG00000261468 | | 0.57 | 0.17 | 2.78E-04 | 9.14E-04 |
| ENSG00000130958 | SLC35D2 | 0.57 | 0.18 | 3.75E-04 | 1.21E-03 |
| ENSG00000229931 | ATXN1-AS1 | 0.57 | 0.18 | 5.20E-04 | 1.65E-03 |
| ENSG00000160588 | MPZL3 | 0.57 | 0.19 | 7.68E-04 | 2.38E-03 |
| ENSG00000163888 | CAMK2N2 | 0.57 | 0.19 | 9.82E-04 | 2.99E-03 |
| ENSG00000286214 | COPG2IT1 | 0.57 | 0.2 | 1.10E-03 | 3.33E-03 |
| ENSG00000255182 | | 0.57 | 0.2 | 1.41E-03 | 4.17E-03 |
| ENSG00000089199 | CHGB | 0.57 | 0.21 | 1.63E-03 | 4.77E-03 |

| | | | | | |
|---|---|---|---|---|---|
| ENSG00000272240 | | 0.57 | 0.24 | 4.72E-03 | 1.28E-02 |
| ENSG00000196366 | C9orf163 | 0.57 | 0.24 | 5.06E-03 | 1.36E-02 |
| ENSG00000228784 | LINC00954 | 0.57 | 0.27 | 1.00E-02 | 2.53E-02 |
| ENSG00000242299 | RPS18P5 | 0.57 | 0.27 | 1.01E-02 | 2.55E-02 |
| ENSG00000104967 | NOVA2 | 0.57 | 0.28 | 1.06E-02 | 2.65E-02 |
| ENSG00000235863 | B3GALT4 | 0.57 | 0.29 | 1.37E-02 | 3.34E-02 |
| ENSG00000282995 | FRG1EP | 0.57 | 0.29 | 1.37E-02 | 3.36E-02 |
| ENSG00000149489 | ROM1 | 0.57 | 0.29 | 1.41E-02 | 3.43E-02 |
| ENSG00000185495 | SEPTIN7P13 | 0.57 | 0.74 | 2.04E-02 | 4.81E-02 |
| ENSG00000204842 | ATXN2 | 0.56 | 0.04 | 3.96E-49 | 8.72E-48 |
| ENSG00000175470 | PPP2R2D | 0.56 | 0.04 | 5.17E-40 | 9.24E-39 |
| ENSG00000109814 | UGDH | 0.56 | 0.05 | 1.24E-34 | 1.92E-33 |
| ENSG00000165434 | PGM2L1 | 0.56 | 0.05 | 1.27E-34 | 1.96E-33 |
| ENSG00000115524 | SF3B1 | 0.56 | 0.05 | 3.09E-30 | 4.19E-29 |
| ENSG00000100106 | TRIOBP | 0.56 | 0.05 | 5.15E-28 | 6.45E-27 |
| ENSG00000243317 | STMP1 | 0.56 | 0.05 | 9.96E-26 | 1.15E-24 |
| ENSG00000213923 | CSNK1E | 0.56 | 0.06 | 2.83E-22 | 2.88E-21 |
| ENSG00000030110 | BAK1 | 0.56 | 0.06 | 6.32E-21 | 6.10E-20 |
| ENSG00000164251 | F2RL1 | 0.56 | 0.06 | 3.67E-19 | 3.29E-18 |
| ENSG00000036054 | TBC1D23 | 0.56 | 0.07 | 3.74E-17 | 3.07E-16 |
| ENSG00000196263 | ZNF471 | 0.56 | 0.07 | 1.86E-16 | 1.47E-15 |
| ENSG00000105223 | PLD3 | 0.56 | 0.07 | 9.20E-15 | 6.76E-14 |
| ENSG00000007545 | CRAMP1 | 0.56 | 0.07 | 2.99E-14 | 2.14E-13 |
| ENSG00000138433 | CIR1 | 0.56 | 0.08 | 3.44E-14 | 2.45E-13 |
| ENSG00000143434 | SEMA6C | 0.56 | 0.08 | 3.95E-14 | 2.81E-13 |
| ENSG00000154930 | ACSS1 | 0.56 | 0.08 | 5.22E-14 | 3.69E-13 |
| ENSG00000196458 | ZNF605 | 0.56 | 0.08 | 1.70E-13 | 1.17E-12 |
| ENSG00000008710 | PKD1 | 0.56 | 0.08 | 2.57E-13 | 1.75E-12 |
| ENSG00000011028 | MRC2 | 0.56 | 0.08 | 4.57E-13 | 3.06E-12 |
| ENSG00000009780 | FAM76A | 0.56 | 0.08 | 1.45E-12 | 9.39E-12 |
| ENSG00000168672 | LRATD2 | 0.56 | 0.09 | 4.55E-11 | 2.70E-10 |
| ENSG00000263072 | ZNF213-AS1 | 0.56 | 0.09 | 5.06E-10 | 2.79E-09 |
| ENSG00000182087 | TMEM259 | 0.56 | 0.1 | 1.38E-09 | 7.35E-09 |
| ENSG00000103024 | NME3 | 0.56 | 0.1 | 1.32E-08 | 6.53E-08 |
| ENSG00000167302 | TEPSIN | 0.56 | 0.1 | 1.68E-08 | 8.26E-08 |
| ENSG00000204498 | NFKBIL1 | 0.56 | 0.1 | 1.68E-08 | 8.27E-08 |
| ENSG00000096872 | IFT74 | 0.56 | 0.11 | 1.51E-07 | 6.90E-07 |
| ENSG00000204434 | POTEKP | 0.56 | 0.12 | 1.61E-06 | 6.71E-06 |
| ENSG00000291046 | | 0.56 | 0.14 | 1.79E-05 | 6.75E-05 |
| ENSG00000067836 | ROGDI | 0.56 | 0.15 | 3.74E-05 | 1.36E-04 |
| ENSG00000119508 | NR4A3 | 0.56 | 0.15 | 4.34E-05 | 1.57E-04 |
| ENSG00000051523 | CYBA | 0.56 | 0.15 | 8.11E-05 | 2.85E-04 |
| ENSG00000287023 | | 0.56 | 0.15 | 8.69E-05 | 3.04E-04 |
| ENSG00000187583 | PLEKHN1 | 0.56 | 0.17 | 2.20E-04 | 7.35E-04 |
| ENSG00000262580 | | 0.56 | 0.18 | 5.11E-04 | 1.62E-03 |
| ENSG00000111863 | ADTRP | 0.56 | 0.19 | 7.85E-04 | 2.43E-03 |
| ENSG00000185065 | UFD1-AS1 | 0.56 | 0.19 | 8.66E-04 | 2.66E-03 |
| ENSG00000224559 | LINC01087 | 0.56 | 0.19 | 9.46E-04 | 2.89E-03 |
| ENSG00000228232 | GAPDHP1 | 0.56 | 0.21 | 2.15E-03 | 6.20E-03 |
| ENSG00000155749 | FLACC1 | 0.56 | 0.21 | 2.40E-03 | 6.87E-03 |

| | | | | | |
|---|---|---|---|---|---|
| ENSG00000248489 | CHD1-DT | 0.56 | 0.21 | 2.76E-03 | 7.80E-03 |
| ENSG00000123576 | ESX1 | 0.56 | 0.23 | 4.02E-03 | 1.10E-02 |
| ENSG00000143570 | SLC39A1 | 0.56 | 0.23 | 4.93E-03 | 1.33E-02 |
| ENSG00000279968 | CCDC28A-AS1 | 0.56 | 0.24 | 5.30E-03 | 1.42E-02 |
| ENSG00000233954 | UQCRHL | 0.56 | 0.24 | 5.40E-03 | 1.44E-02 |
| ENSG00000248206 | | 0.56 | 0.25 | 6.97E-03 | 1.82E-02 |
| ENSG00000279500 | | 0.56 | 0.26 | 9.53E-03 | 2.42E-02 |
| ENSG00000287737 | | 0.56 | 0.28 | 1.15E-02 | 2.87E-02 |
| ENSG00000226942 | IL9RP3 | 0.56 | 0.3 | 1.69E-02 | 4.04E-02 |
| ENSG00000144357 | UBR3 | 0.55 | 0.04 | 1.52E-44 | 3.03E-43 |
| ENSG00000129562 | DAD1 | 0.55 | 0.04 | 1.53E-36 | 2.51E-35 |
| ENSG00000082701 | GSK3B | 0.55 | 0.05 | 8.29E-35 | 1.29E-33 |
| ENSG00000135046 | ANXA1 | 0.55 | 0.05 | 2.03E-33 | 3.04E-32 |
| ENSG00000118454 | ANKRD13C | 0.55 | 0.05 | 1.87E-30 | 2.56E-29 |
| ENSG00000187957 | DNER | 0.55 | 0.05 | 1.86E-27 | 2.30E-26 |
| ENSG00000159479 | MED8 | 0.55 | 0.05 | 2.31E-26 | 2.74E-25 |
| ENSG00000039560 | RAI14 | 0.55 | 0.05 | 1.98E-25 | 2.27E-24 |
| ENSG00000174695 | TMEM167A | 0.55 | 0.05 | 2.08E-24 | 2.30E-23 |
| ENSG00000113595 | TRIM23 | 0.55 | 0.05 | 2.45E-24 | 2.69E-23 |
| ENSG00000167695 | TLCD3A | 0.55 | 0.06 | 3.16E-22 | 3.21E-21 |
| ENSG00000162616 | DNAJB4 | 0.55 | 0.06 | 3.78E-22 | 3.83E-21 |
| ENSG00000107036 | RIC1 | 0.55 | 0.06 | 1.44E-21 | 1.43E-20 |
| ENSG00000119541 | VPS4B | 0.55 | 0.06 | 4.00E-21 | 3.91E-20 |
| ENSG00000122482 | ZNF644 | 0.55 | 0.06 | 2.33E-20 | 2.20E-19 |
| ENSG00000116095 | PLEKHA3 | 0.55 | 0.06 | 2.33E-20 | 2.20E-19 |
| ENSG00000173114 | LRRN3 | 0.55 | 0.06 | 4.25E-19 | 3.79E-18 |
| ENSG00000143457 | GOLPH3L | 0.55 | 0.06 | 4.61E-19 | 4.10E-18 |
| ENSG00000172057 | ORMDL3 | 0.55 | 0.06 | 6.65E-19 | 5.87E-18 |
| ENSG00000151881 | TMEM267 | 0.55 | 0.06 | 1.18E-18 | 1.03E-17 |
| ENSG00000131067 | GGT7 | 0.55 | 0.06 | 3.83E-18 | 3.27E-17 |
| ENSG00000167113 | COQ4 | 0.55 | 0.07 | 3.14E-16 | 2.47E-15 |
| ENSG00000091009 | RBM27 | 0.55 | 0.08 | 2.70E-13 | 1.83E-12 |
| ENSG00000121005 | CRISPLD1 | 0.55 | 0.08 | 5.41E-12 | 3.39E-11 |
| ENSG00000105136 | ZNF419 | 0.55 | 0.08 | 1.03E-11 | 6.35E-11 |
| ENSG00000173581 | CCDC106 | 0.55 | 0.08 | 1.04E-11 | 6.44E-11 |
| ENSG00000179526 | SHARPIN | 0.55 | 0.09 | 2.72E-11 | 1.64E-10 |
| ENSG00000178149 | DALRD3 | 0.55 | 0.09 | 8.12E-11 | 4.74E-10 |
| ENSG00000169692 | AGPAT2 | 0.55 | 0.09 | 9.64E-11 | 5.60E-10 |
| ENSG00000156313 | RPGR | 0.55 | 0.09 | 1.42E-10 | 8.17E-10 |
| ENSG00000167700 | MFSD3 | 0.55 | 0.09 | 1.82E-10 | 1.04E-09 |
| ENSG00000110046 | ATG2A | 0.55 | 0.09 | 2.83E-10 | 1.59E-09 |
| ENSG00000142552 | RCN3 | 0.55 | 0.09 | 1.22E-09 | 6.53E-09 |
| ENSG00000203761 | MSTO2P | 0.55 | 0.09 | 1.41E-09 | 7.51E-09 |
| ENSG00000196421 | C20orf204 | 0.55 | 0.09 | 1.44E-09 | 7.69E-09 |
| ENSG00000158201 | ABHD3 | 0.55 | 0.1 | 3.94E-09 | 2.03E-08 |
| ENSG00000185950 | IRS2 | 0.55 | 0.1 | 1.09E-08 | 5.47E-08 |
| ENSG00000104177 | MYEF2 | 0.55 | 0.11 | 9.68E-08 | 4.50E-07 |
| ENSG00000223865 | HLA-DPB1 | 0.55 | 0.11 | 4.48E-07 | 1.97E-06 |
| ENSG00000258839 | MC1R | 0.55 | 0.11 | 5.40E-07 | 2.36E-06 |
| ENSG00000145506 | NKD2 | 0.55 | 0.14 | 1.36E-05 | 5.18E-05 |

| | | | | | |
|---|---|---|---|---|---|
| ENSG00000179455 | MKRN3 | 0.55 | 0.14 | 3.75E-05 | 1.37E-04 |
| ENSG00000231764 | DLX6-AS1 | 0.55 | 0.15 | 1.16E-04 | 4.00E-04 |
| ENSG00000196388 | INCA1 | 0.55 | 0.17 | 4.33E-04 | 1.39E-03 |
| ENSG00000182902 | SLC25A18 | 0.55 | 0.18 | 8.46E-04 | 2.60E-03 |
| ENSG00000115286 | NDUFS7 | 0.55 | 0.2 | 2.04E-03 | 5.90E-03 |
| ENSG00000268364 | SMC5-DT | 0.55 | 0.22 | 3.36E-03 | 9.34E-03 |
| ENSG00000291213 | DPY19L2P3 | 0.55 | 0.23 | 5.64E-03 | 1.50E-02 |
| ENSG00000072818 | ACAP1 | 0.55 | 0.24 | 6.38E-03 | 1.68E-02 |
| ENSG00000273456 | | 0.55 | 0.25 | 8.50E-03 | 2.18E-02 |
| ENSG00000249087 | ZNF436-AS1 | 0.55 | 0.26 | 1.19E-02 | 2.96E-02 |
| ENSG00000187260 | WDR86 | 0.55 | 0.27 | 1.32E-02 | 3.24E-02 |
| ENSG00000160539 | PLPP7 | 0.55 | 0.28 | 1.57E-02 | 3.80E-02 |
| ENSG00000138674 | SEC31A | 0.54 | 0.03 | 7.12E-60 | 2.00E-58 |
| ENSG00000163125 | RPRD2 | 0.54 | 0.04 | 4.39E-41 | 8.16E-40 |
| ENSG00000105778 | AVL9 | 0.54 | 0.04 | 7.57E-39 | 1.33E-37 |
| ENSG00000076685 | NT5C2 | 0.54 | 0.05 | 8.68E-33 | 1.27E-31 |
| ENSG00000166073 | GPR176 | 0.54 | 0.05 | 2.42E-30 | 3.30E-29 |
| ENSG00000184432 | COPB2 | 0.54 | 0.05 | 5.81E-30 | 7.79E-29 |
| ENSG00000119231 | SENP5 | 0.54 | 0.05 | 6.64E-28 | 8.28E-27 |
| ENSG00000171988 | JMJD1C | 0.54 | 0.05 | 5.60E-25 | 6.31E-24 |
| ENSG00000103363 | ELOB | 0.54 | 0.05 | 3.75E-24 | 4.10E-23 |
| ENSG00000127947 | PTPN12 | 0.54 | 0.06 | 6.30E-21 | 6.09E-20 |
| ENSG00000196233 | LCOR | 0.54 | 0.06 | 9.74E-21 | 9.32E-20 |
| ENSG00000109270 | LAMTOR3 | 0.54 | 0.06 | 5.64E-19 | 4.99E-18 |
| ENSG00000128789 | PSMG2 | 0.54 | 0.06 | 1.88E-18 | 1.63E-17 |
| ENSG00000140299 | BNIP2 | 0.54 | 0.06 | 4.63E-18 | 3.94E-17 |
| ENSG00000114742 | WDR48 | 0.54 | 0.07 | 6.42E-17 | 5.22E-16 |
| ENSG00000180772 | AGTR2 | 0.54 | 0.07 | 2.02E-16 | 1.60E-15 |
| ENSG00000049449 | RCN1 | 0.54 | 0.07 | 3.98E-16 | 3.11E-15 |
| ENSG00000080822 | CLDND1 | 0.54 | 0.07 | 1.24E-15 | 9.41E-15 |
| ENSG00000100711 | ZFYVE21 | 0.54 | 0.07 | 1.33E-15 | 1.01E-14 |
| ENSG00000113721 | PDGFRB | 0.54 | 0.08 | 4.40E-13 | 2.95E-12 |
| ENSG00000183010 | PYCR1 | 0.54 | 0.08 | 7.92E-12 | 4.91E-11 |
| ENSG00000144747 | TMF1 | 0.54 | 0.08 | 1.91E-11 | 1.16E-10 |
| ENSG00000132819 | RBM38 | 0.54 | 0.08 | 2.24E-11 | 1.35E-10 |
| ENSG00000176978 | DPP7 | 0.54 | 0.08 | 4.36E-11 | 2.59E-10 |
| ENSG00000167969 | ECI1 | 0.54 | 0.09 | 3.57E-10 | 1.99E-09 |
| ENSG00000204237 | OXLD1 | 0.54 | 0.09 | 9.87E-10 | 5.33E-09 |
| ENSG00000125347 | IRF1 | 0.54 | 0.1 | 3.36E-08 | 1.62E-07 |
| ENSG00000011132 | APBA3 | 0.54 | 0.11 | 2.04E-07 | 9.24E-07 |
| ENSG00000291272 | | 0.54 | 0.15 | 6.24E-05 | 2.23E-04 |
| ENSG00000099364 | FBXL19 | 0.54 | 0.15 | 1.18E-04 | 4.08E-04 |
| ENSG00000260025 | CRIM1-DT | 0.54 | 0.16 | 2.71E-04 | 8.95E-04 |
| ENSG00000272768 | | 0.54 | 0.18 | 8.35E-04 | 2.57E-03 |
| ENSG00000187134 | AKR1C1 | 0.54 | 0.2 | 2.01E-03 | 5.82E-03 |
| ENSG00000228486 | C2orf92 | 0.54 | 0.21 | 2.79E-03 | 7.89E-03 |
| ENSG00000205702 | CYP2D7 | 0.54 | 0.21 | 3.13E-03 | 8.77E-03 |
| ENSG00000231887 | PRH1 | 0.54 | 0.24 | 8.21E-03 | 2.11E-02 |
| ENSG00000217716 | RPS10P3 | 0.54 | 0.25 | 9.30E-03 | 2.37E-02 |
| ENSG00000178026 | LRRC75B | 0.54 | 0.26 | 1.14E-02 | 2.84E-02 |

| | | | | | |
|---|---|---|---|---|---|
| ENSG00000269313 | MAGIX | 0.54 | 0.27 | 1.38E-02 | 3.38E-02 |
| ENSG00000290842 | CROCCP3 | 0.54 | 0.28 | 1.67E-02 | 4.01E-02 |
| ENSG00000287104 | | 0.54 | 0.29 | 1.87E-02 | 4.43E-02 |
| ENSG00000139644 | TMBIM6 | 0.53 | 0.03 | 2.51E-57 | 6.68E-56 |
| ENSG00000115170 | ACVR1 | 0.53 | 0.04 | 1.74E-33 | 2.61E-32 |
| ENSG00000291237 | SOD2 | 0.53 | 0.04 | 4.62E-33 | 6.81E-32 |
| ENSG00000235106 | BRD3OS | 0.53 | 0.05 | 9.29E-32 | 1.32E-30 |
| ENSG00000115020 | PIKFYVE | 0.53 | 0.05 | 1.30E-30 | 1.79E-29 |
| ENSG00000181472 | ZBTB2 | 0.53 | 0.05 | 1.96E-30 | 2.68E-29 |
| ENSG00000143079 | CTTNBP2NL | 0.53 | 0.05 | 2.01E-29 | 2.65E-28 |
| ENSG00000157837 | SPPL3 | 0.53 | 0.05 | 7.13E-29 | 9.23E-28 |
| ENSG00000147324 | MFHAS1 | 0.53 | 0.05 | 1.44E-27 | 1.79E-26 |
| ENSG00000120709 | FAM53C | 0.53 | 0.05 | 5.02E-27 | 6.09E-26 |
| ENSG00000090520 | DNAJB11 | 0.53 | 0.06 | 1.03E-21 | 1.02E-20 |
| ENSG00000169118 | CSNK1G1 | 0.53 | 0.06 | 4.32E-21 | 4.21E-20 |
| ENSG00000147162 | OGT | 0.53 | 0.06 | 5.77E-20 | 5.35E-19 |
| ENSG00000198925 | ATG9A | 0.53 | 0.06 | 1.42E-18 | 1.24E-17 |
| ENSG00000170100 | ZNF778 | 0.53 | 0.06 | 1.18E-17 | 9.92E-17 |
| ENSG00000165502 | RPL36AL | 0.53 | 0.06 | 8.32E-17 | 6.72E-16 |
| ENSG00000169976 | SF3B5 | 0.53 | 0.06 | 1.20E-16 | 9.64E-16 |
| ENSG00000204272 | NBDY | 0.53 | 0.07 | 3.69E-16 | 2.89E-15 |
| ENSG00000087095 | NLK | 0.53 | 0.07 | 6.79E-16 | 5.25E-15 |
| ENSG00000137876 | RSL24D1 | 0.53 | 0.07 | 3.66E-14 | 2.60E-13 |
| ENSG00000181690 | PLAG1 | 0.53 | 0.07 | 7.07E-14 | 4.96E-13 |
| ENSG00000007541 | PIGQ | 0.53 | 0.07 | 1.17E-13 | 8.14E-13 |
| ENSG00000168434 | COG7 | 0.53 | 0.07 | 1.19E-13 | 8.23E-13 |
| ENSG00000160888 | IER2 | 0.53 | 0.07 | 3.47E-13 | 2.34E-12 |
| ENSG00000106004 | HOXA5 | 0.53 | 0.08 | 3.36E-12 | 2.13E-11 |
| ENSG00000100243 | CYB5R3 | 0.53 | 0.08 | 5.98E-12 | 3.74E-11 |
| ENSG00000197457 | STMN3 | 0.53 | 0.08 | 2.79E-11 | 1.68E-10 |
| ENSG00000268658 | LINC00664 | 0.53 | 0.08 | 5.22E-11 | 3.08E-10 |
| ENSG00000177700 | POLR2L | 0.53 | 0.09 | 2.32E-10 | 1.31E-09 |
| ENSG00000145335 | SNCA | 0.53 | 0.09 | 2.78E-10 | 1.56E-09 |
| ENSG00000118985 | ELL2 | 0.53 | 0.09 | 2.88E-10 | 1.62E-09 |
| ENSG00000203965 | EFCAB7 | 0.53 | 0.09 | 6.58E-10 | 3.61E-09 |
| ENSG00000253276 | CCDC71L | 0.53 | 0.09 | 9.38E-10 | 5.07E-09 |
| ENSG00000255310 | | 0.53 | 0.09 | 2.43E-09 | 1.27E-08 |
| ENSG00000204599 | TRIM39 | 0.53 | 0.09 | 3.33E-09 | 1.73E-08 |
| ENSG00000130731 | METTL26 | 0.53 | 0.09 | 3.46E-09 | 1.79E-08 |
| ENSG00000106404 | CLDN15 | 0.53 | 0.1 | 1.17E-08 | 5.84E-08 |
| ENSG00000187792 | ZNF70 | 0.53 | 0.1 | 2.11E-08 | 1.03E-07 |
| ENSG00000210196 | MT-TP | 0.53 | 0.11 | 2.86E-07 | 1.28E-06 |
| ENSG00000233276 | GPX1 | 0.53 | 0.14 | 6.78E-05 | 2.41E-04 |
| ENSG00000198208 | RPS6KL1 | 0.53 | 0.16 | 2.12E-04 | 7.09E-04 |
| ENSG00000246174 | KCTD21-AS1 | 0.53 | 0.16 | 2.33E-04 | 7.75E-04 |
| ENSG00000250251 | PKD1P6 | 0.53 | 0.16 | 3.66E-04 | 1.18E-03 |
| ENSG00000167619 | TMEM145 | 0.53 | 0.17 | 5.29E-04 | 1.68E-03 |
| ENSG00000178096 | BOLA1 | 0.53 | 0.18 | 1.14E-03 | 3.42E-03 |
| ENSG00000287562 | | 0.53 | 0.2 | 2.52E-03 | 7.18E-03 |
| ENSG00000112619 | PRPH2 | 0.53 | 0.2 | 3.03E-03 | 8.50E-03 |

| | | | | | |
|---|---|---|---|---|---|
| ENSG00000216895 | RBM33-DT | 0.53 | 0.21 | 3.26E-03 | 9.09E-03 |
| ENSG00000230979 | | 0.53 | 0.21 | 4.33E-03 | 1.18E-02 |
| ENSG00000277945 | | 0.53 | 0.26 | 1.21E-02 | 3.01E-02 |
| ENSG00000288560 | | 0.53 | 0.27 | 1.57E-02 | 3.78E-02 |
| ENSG00000285437 | POLR2J3 | 0.53 | 0.28 | 1.86E-02 | 4.42E-02 |
| ENSG00000253559 | OSGEPL1-AS1 | 0.53 | 0.28 | 2.05E-02 | 4.81E-02 |
| ENSG00000170471 | RALGAPB | 0.52 | 0.04 | 1.15E-40 | 2.10E-39 |
| ENSG00000177156 | TALDO1 | 0.52 | 0.04 | 2.01E-34 | 3.08E-33 |
| ENSG00000166471 | TMEM41B | 0.52 | 0.04 | 6.35E-33 | 9.34E-32 |
| ENSG00000160299 | PCNT | 0.52 | 0.04 | 7.49E-32 | 1.07E-30 |
| ENSG00000141424 | SLC39A6 | 0.52 | 0.05 | 5.65E-31 | 7.88E-30 |
| ENSG00000114439 | BBX | 0.52 | 0.05 | 4.37E-30 | 5.89E-29 |
| ENSG00000173575 | CHD2 | 0.52 | 0.05 | 3.38E-29 | 4.42E-28 |
| ENSG00000071537 | SEL1L | 0.52 | 0.05 | 1.59E-27 | 1.97E-26 |
| ENSG00000139697 | SBNO1 | 0.52 | 0.05 | 2.68E-25 | 3.05E-24 |
| ENSG00000104979 | C19orf53 | 0.52 | 0.05 | 7.07E-25 | 7.95E-24 |
| ENSG00000072958 | AP1M1 | 0.52 | 0.05 | 1.24E-22 | 1.28E-21 |
| ENSG00000182752 | PAPPA | 0.52 | 0.05 | 1.91E-22 | 1.96E-21 |
| ENSG00000155254 | MARVELD1 | 0.52 | 0.06 | 2.28E-21 | 2.24E-20 |
| ENSG00000145740 | SLC30A5 | 0.52 | 0.06 | 9.43E-21 | 9.03E-20 |
| ENSG00000106367 | AP1S1 | 0.52 | 0.06 | 4.68E-20 | 4.36E-19 |
| ENSG00000111203 | ITFG2 | 0.52 | 0.06 | 1.78E-19 | 1.62E-18 |
| ENSG00000117479 | SLC19A2 | 0.52 | 0.06 | 5.16E-19 | 4.58E-18 |
| ENSG00000204520 | MICA | 0.52 | 0.06 | 6.48E-19 | 5.73E-18 |
| ENSG00000105556 | MIER2 | 0.52 | 0.06 | 4.61E-17 | 3.77E-16 |
| ENSG00000101745 | ANKRD12 | 0.52 | 0.07 | 1.46E-15 | 1.11E-14 |
| ENSG00000204681 | GABBR1 | 0.52 | 0.07 | 2.47E-15 | 1.86E-14 |
| ENSG00000159899 | NPR2 | 0.52 | 0.07 | 4.71E-15 | 3.50E-14 |
| ENSG00000144401 | METTL21A | 0.52 | 0.07 | 9.42E-14 | 6.56E-13 |
| ENSG00000127837 | AAMP | 0.52 | 0.07 | 5.42E-13 | 3.61E-12 |
| ENSG00000183354 | KIAA2026 | 0.52 | 0.08 | 2.39E-12 | 1.53E-11 |
| ENSG00000169926 | KLF13 | 0.52 | 0.08 | 2.12E-11 | 1.28E-10 |
| ENSG00000131408 | NR1H2 | 0.52 | 0.08 | 2.31E-11 | 1.40E-10 |
| ENSG00000099999 | RNF215 | 0.52 | 0.08 | 2.40E-11 | 1.45E-10 |
| ENSG00000183648 | NDUFB1 | 0.52 | 0.08 | 3.57E-11 | 2.13E-10 |
| ENSG00000142102 | PGGHG | 0.52 | 0.08 | 1.56E-10 | 8.95E-10 |
| ENSG00000174473 | GALNTL6 | 0.52 | 0.08 | 1.77E-10 | 1.01E-09 |
| ENSG00000001461 | NIPAL3 | 0.52 | 0.08 | 2.65E-10 | 1.49E-09 |
| ENSG00000133243 | BTBD2 | 0.52 | 0.09 | 1.01E-09 | 5.45E-09 |
| ENSG00000154370 | TRIM11 | 0.52 | 0.09 | 2.16E-09 | 1.14E-08 |
| ENSG00000123349 | PFDN5 | 0.52 | 0.1 | 1.93E-08 | 9.45E-08 |
| ENSG00000114779 | ABHD14B | 0.52 | 0.1 | 2.22E-08 | 1.08E-07 |
| ENSG00000173482 | PTPRM | 0.52 | 0.12 | 3.61E-06 | 1.46E-05 |
| ENSG00000269867 | | 0.52 | 0.12 | 3.77E-06 | 1.52E-05 |
| ENSG00000197380 | DACT3 | 0.52 | 0.13 | 2.11E-05 | 7.89E-05 |
| ENSG00000281903 | LINC02246 | 0.52 | 0.15 | 1.34E-04 | 4.58E-04 |
| ENSG00000290948 | | 0.52 | 0.15 | 2.07E-04 | 6.91E-04 |
| ENSG00000142303 | ADAMTS10 | 0.52 | 0.15 | 2.52E-04 | 8.34E-04 |
| ENSG00000280287 | | 0.52 | 0.16 | 3.35E-04 | 1.09E-03 |
| ENSG00000175505 | CLCF1 | 0.52 | 0.17 | 8.77E-04 | 2.69E-03 |

| | | | | | |
|---|---|---|---|---|---|
| ENSG00000066923 | STAG3 | 0.52 | 0.18 | 9.33E-04 | 2.85E-03 |
| ENSG00000215068 | ANXA2R-AS1 | 0.52 | 0.18 | 1.41E-03 | 4.19E-03 |
| ENSG00000281468 | | 0.52 | 0.18 | 1.51E-03 | 4.45E-03 |
| ENSG00000253955 | LINC02995 | 0.52 | 0.25 | 1.24E-02 | 3.07E-02 |
| ENSG00000187017 | ESPN | 0.52 | 0.25 | 1.26E-02 | 3.11E-02 |
| ENSG00000290085 | | 0.52 | 0.27 | 1.82E-02 | 4.32E-02 |
| ENSG00000139725 | RHOF | 0.52 | 0.28 | 1.89E-02 | 4.49E-02 |
| ENSG00000108587 | GOSR1 | 0.51 | 0.04 | 3.53E-38 | 6.07E-37 |
| ENSG00000136986 | DERL1 | 0.51 | 0.04 | 1.38E-35 | 2.21E-34 |
| ENSG00000181852 | RNF41 | 0.51 | 0.04 | 1.34E-33 | 2.01E-32 |
| ENSG00000065911 | MTHFD2 | 0.51 | 0.05 | 5.81E-27 | 7.04E-26 |
| ENSG00000140743 | CDR2 | 0.51 | 0.05 | 6.88E-27 | 8.31E-26 |
| ENSG00000008294 | SPAG9 | 0.51 | 0.05 | 4.28E-24 | 4.66E-23 |
| ENSG00000133059 | DSTYK | 0.51 | 0.05 | 2.74E-21 | 2.69E-20 |
| ENSG00000140577 | CRTC3 | 0.51 | 0.06 | 2.19E-20 | 2.07E-19 |
| ENSG00000101052 | IFT52 | 0.51 | 0.06 | 5.72E-19 | 5.06E-18 |
| ENSG00000242125 | SNHG3 | 0.51 | 0.06 | 7.83E-18 | 6.61E-17 |
| ENSG00000151914 | DST | 0.51 | 0.06 | 1.16E-17 | 9.72E-17 |
| ENSG00000176783 | RUFY1 | 0.51 | 0.06 | 2.37E-17 | 1.96E-16 |
| ENSG00000180875 | GREM2 | 0.51 | 0.07 | 4.25E-14 | 3.01E-13 |
| ENSG00000118508 | RAB32 | 0.51 | 0.07 | 8.49E-14 | 5.92E-13 |
| ENSG00000100083 | GGA1 | 0.51 | 0.07 | 4.03E-13 | 2.71E-12 |
| ENSG00000164576 | SAP30L | 0.51 | 0.07 | 1.01E-12 | 6.63E-12 |
| ENSG00000180190 | TDRP | 0.51 | 0.07 | 1.46E-12 | 9.50E-12 |
| ENSG00000137055 | PLAA | 0.51 | 0.07 | 3.20E-12 | 2.04E-11 |
| ENSG00000175105 | ZNF654 | 0.51 | 0.08 | 7.30E-11 | 4.26E-10 |
| ENSG00000213903 | LTB4R | 0.51 | 0.08 | 2.35E-10 | 1.33E-09 |
| ENSG00000175573 | C11orf68 | 0.51 | 0.09 | 1.19E-08 | 5.94E-08 |
| ENSG00000161551 | ZNF577 | 0.51 | 0.09 | 1.63E-08 | 8.04E-08 |
| ENSG00000142444 | TIMM29 | 0.51 | 0.09 | 2.63E-08 | 1.28E-07 |
| ENSG00000228709 | LINC02575 | 0.51 | 0.1 | 2.73E-08 | 1.33E-07 |
| ENSG00000069020 | MAST4 | 0.51 | 0.1 | 3.62E-08 | 1.74E-07 |
| ENSG00000111775 | COX6A1 | 0.51 | 0.1 | 9.18E-08 | 4.27E-07 |
| ENSG00000169330 | MINAR1 | 0.51 | 0.1 | 2.45E-07 | 1.10E-06 |
| ENSG00000171823 | FBXL14 | 0.51 | 0.1 | 2.65E-07 | 1.19E-06 |
| ENSG00000198246 | SLC29A3 | 0.51 | 0.1 | 3.17E-07 | 1.41E-06 |
| ENSG00000221869 | CEBPD | 0.51 | 0.1 | 3.21E-07 | 1.43E-06 |
| ENSG00000134253 | TRIM45 | 0.51 | 0.11 | 3.55E-06 | 1.43E-05 |
| ENSG00000136149 | RPL13AP25 | 0.51 | 0.13 | 3.93E-05 | 1.43E-04 |
| ENSG00000088766 | CRLS1 | 0.51 | 0.13 | 4.16E-05 | 1.51E-04 |
| ENSG00000130244 | FAM98C | 0.51 | 0.13 | 4.87E-05 | 1.75E-04 |
| ENSG00000282936 | | 0.51 | 0.14 | 7.50E-05 | 2.65E-04 |
| ENSG00000162510 | MATN1 | 0.51 | 0.15 | 2.34E-04 | 7.78E-04 |
| ENSG00000007372 | PAX6 | 0.51 | 0.15 | 2.64E-04 | 8.72E-04 |
| ENSG00000232611 | | 0.51 | 0.17 | 7.98E-04 | 2.46E-03 |
| ENSG00000056487 | PHF21B | 0.51 | 0.17 | 8.59E-04 | 2.64E-03 |
| ENSG00000213326 | RPS7P11 | 0.51 | 0.18 | 1.52E-03 | 4.47E-03 |
| ENSG00000289138 | | 0.51 | 0.2 | 3.43E-03 | 9.51E-03 |
| ENSG00000219665 | ZNF433-AS1 | 0.51 | 0.22 | 7.03E-03 | 1.84E-02 |
| ENSG00000267048 | | 0.51 | 0.24 | 1.09E-02 | 2.72E-02 |

| | | | | | |
|---|---|---|---|---|---|
| ENSG00000273828 | | 0.51 | 0.24 | 1.12E-02 | 2.79E-02 |
| ENSG00000146677 | RPL32P18 | 0.51 | 0.24 | 1.13E-02 | 2.82E-02 |
| ENSG00000249378 | LINC01060 | 0.51 | 0.25 | 1.35E-02 | 3.30E-02 |
| ENSG00000181458 | TMEM45A | 0.51 | 0.26 | 1.72E-02 | 4.11E-02 |
| ENSG00000288826 | | 0.51 | 0.26 | 1.73E-02 | 4.12E-02 |
| ENSG00000162521 | RBBP4 | 0.5 | 0.03 | 1.62E-52 | 3.89E-51 |
| ENSG00000104695 | PPP2CB | 0.5 | 0.04 | 9.41E-38 | 1.60E-36 |
| ENSG00000170266 | GLB1 | 0.5 | 0.04 | 9.60E-37 | 1.59E-35 |
| ENSG00000159346 | ADIPOR1 | 0.5 | 0.04 | 1.61E-36 | 2.65E-35 |
| ENSG00000147416 | ATP6V1B2 | 0.5 | 0.04 | 1.77E-34 | 2.72E-33 |
| ENSG00000197622 | CDC42SE1 | 0.5 | 0.05 | 1.19E-28 | 1.53E-27 |
| ENSG00000221978 | CCNL2 | 0.5 | 0.05 | 1.22E-26 | 1.47E-25 |
| ENSG00000163960 | UBXN7 | 0.5 | 0.05 | 1.72E-26 | 2.05E-25 |
| ENSG00000151694 | ADAM17 | 0.5 | 0.05 | 6.52E-26 | 7.60E-25 |
| ENSG00000110696 | C11orf58 | 0.5 | 0.05 | 5.65E-25 | 6.37E-24 |
| ENSG00000175215 | CTDSP2 | 0.5 | 0.05 | 2.83E-24 | 3.11E-23 |
| ENSG00000167196 | FBXO22 | 0.5 | 0.05 | 7.48E-22 | 7.49E-21 |
| ENSG00000127914 | AKAP9 | 0.5 | 0.06 | 1.19E-19 | 1.09E-18 |
| ENSG00000175352 | NRIP3 | 0.5 | 0.06 | 1.70E-19 | 1.54E-18 |
| ENSG00000000003 | TSPAN6 | 0.5 | 0.06 | 8.80E-19 | 7.73E-18 |
| ENSG00000141452 | RMC1 | 0.5 | 0.06 | 9.62E-17 | 7.75E-16 |
| ENSG00000100439 | ABHD4 | 0.5 | 0.06 | 1.90E-16 | 1.51E-15 |
| ENSG00000221823 | PPP3R1 | 0.5 | 0.06 | 3.63E-16 | 2.85E-15 |
| ENSG00000169241 | SLC50A1 | 0.5 | 0.06 | 3.73E-16 | 2.92E-15 |
| ENSG00000116786 | PLEKHM2 | 0.5 | 0.07 | 1.47E-14 | 1.07E-13 |
| ENSG00000170791 | CHCHD7 | 0.5 | 0.07 | 5.67E-14 | 3.99E-13 |
| ENSG00000185730 | ZNF696 | 0.5 | 0.07 | 1.89E-13 | 1.30E-12 |
| ENSG00000196453 | ZNF777 | 0.5 | 0.07 | 1.92E-12 | 1.24E-11 |
| ENSG00000170619 | COMMD5 | 0.5 | 0.07 | 2.62E-12 | 1.68E-11 |
| ENSG00000123739 | PLA2G12A | 0.5 | 0.07 | 3.09E-12 | 1.97E-11 |
| ENSG00000139793 | MBNL2 | 0.5 | 0.08 | 1.27E-11 | 7.78E-11 |
| ENSG00000165195 | PIGA | 0.5 | 0.08 | 2.69E-11 | 1.62E-10 |
| ENSG00000204946 | ZNF783 | 0.5 | 0.08 | 8.37E-11 | 4.88E-10 |
| ENSG00000179818 | PCBP1-AS1 | 0.5 | 0.08 | 2.61E-10 | 1.47E-09 |
| ENSG00000167523 | SPATA33 | 0.5 | 0.08 | 2.99E-10 | 1.68E-09 |
| ENSG00000198455 | ZXDB | 0.5 | 0.08 | 3.02E-10 | 1.69E-09 |
| ENSG00000149115 | TNKS1BP1 | 0.5 | 0.08 | 1.52E-09 | 8.08E-09 |
| ENSG00000196110 | ZNF699 | 0.5 | 0.08 | 1.63E-09 | 8.68E-09 |
| ENSG00000173818 | ENDOV | 0.5 | 0.09 | 3.73E-09 | 1.93E-08 |
| ENSG00000043355 | ZIC2 | 0.5 | 0.09 | 3.74E-09 | 1.93E-08 |
| ENSG00000168118 | RAB4A | 0.5 | 0.09 | 3.95E-09 | 2.03E-08 |
| ENSG00000143224 | PPOX | 0.5 | 0.09 | 1.19E-08 | 5.93E-08 |
| ENSG00000080823 | MOK | 0.5 | 0.09 | 3.30E-08 | 1.59E-07 |
| ENSG00000103126 | AXIN1 | 0.5 | 0.1 | 1.08E-07 | 5.01E-07 |
| ENSG00000100417 | PMM1 | 0.5 | 0.1 | 1.12E-07 | 5.19E-07 |
| ENSG00000159388 | BTG2 | 0.5 | 0.11 | 7.65E-07 | 3.28E-06 |
| ENSG00000164967 | RPP25L | 0.5 | 0.11 | 8.57E-07 | 3.66E-06 |
| ENSG00000214226 | C17orf67 | 0.5 | 0.12 | 6.88E-06 | 2.70E-05 |
| ENSG00000213722 | DDAH2 | 0.5 | 0.12 | 8.01E-06 | 3.13E-05 |
| ENSG00000182759 | MAFA | 0.5 | 0.13 | 3.14E-05 | 1.15E-04 |

| | | | | | |
|---|---|---|---|---|---|
| ENSG00000123810 | B9D2 | 0.5 | 0.14 | 1.08E-04 | 3.74E-04 |
| ENSG00000095637 | SORBS1 | 0.5 | 0.16 | 5.26E-04 | 1.67E-03 |
| ENSG00000165312 | OTUD1 | 0.5 | 0.16 | 5.45E-04 | 1.73E-03 |
| ENSG00000278616 | BEND3P3 | 0.5 | 0.16 | 6.27E-04 | 1.97E-03 |
| ENSG00000197182 | MIRLET7BHG | 0.5 | 0.16 | 6.56E-04 | 2.05E-03 |
| ENSG00000124171 | PARD6B | 0.5 | 0.16 | 9.50E-04 | 2.90E-03 |
| ENSG00000125531 | FNDC11 | 0.5 | 0.2 | 4.38E-03 | 1.19E-02 |
| ENSG00000187824 | TMEM220 | 0.5 | 0.21 | 5.60E-03 | 1.49E-02 |
| ENSG00000276728 | | 0.5 | 0.22 | 7.98E-03 | 2.06E-02 |
| ENSG00000291110 | TRIM16L | 0.5 | 0.23 | 1.20E-02 | 2.99E-02 |
| ENSG00000133872 | SARAF | 0.49 | 0.04 | 5.25E-34 | 7.96E-33 |
| ENSG00000166780 | BMERB1 | 0.49 | 0.04 | 1.51E-28 | 1.93E-27 |
| ENSG00000184863 | RBM33 | 0.49 | 0.05 | 1.60E-27 | 1.97E-26 |
| ENSG00000135047 | CTSL | 0.49 | 0.05 | 1.82E-27 | 2.24E-26 |
| ENSG00000106803 | SEC61B | 0.49 | 0.05 | 2.37E-26 | 2.81E-25 |
| ENSG00000145391 | SETD7 | 0.49 | 0.05 | 4.93E-26 | 5.77E-25 |
| ENSG00000110395 | CBL | 0.49 | 0.05 | 5.20E-26 | 6.08E-25 |
| ENSG00000136810 | TXN | 0.49 | 0.05 | 9.29E-22 | 9.26E-21 |
| ENSG00000047249 | ATP6V1H | 0.49 | 0.05 | 4.82E-20 | 4.48E-19 |
| ENSG00000198862 | LTN1 | 0.49 | 0.06 | 1.66E-18 | 1.45E-17 |
| ENSG00000112290 | WASF1 | 0.49 | 0.06 | 1.37E-17 | 1.15E-16 |
| ENSG00000134248 | LAMTOR5 | 0.49 | 0.06 | 2.58E-17 | 2.13E-16 |
| ENSG00000106052 | TAX1BP1 | 0.49 | 0.06 | 7.00E-17 | 5.68E-16 |
| ENSG00000162551 | ALPL | 0.49 | 0.06 | 8.68E-16 | 6.65E-15 |
| ENSG00000176155 | CCDC57 | 0.49 | 0.06 | 6.61E-15 | 4.88E-14 |
| ENSG00000154429 | CCSAP | 0.49 | 0.07 | 1.25E-13 | 8.67E-13 |
| ENSG00000102103 | PQBP1 | 0.49 | 0.07 | 6.86E-13 | 4.54E-12 |
| ENSG00000184588 | PDE4B | 0.49 | 0.07 | 1.11E-12 | 7.24E-12 |
| ENSG00000111911 | HINT3 | 0.49 | 0.07 | 1.60E-12 | 1.04E-11 |
| ENSG00000197324 | LRP10 | 0.49 | 0.07 | 2.46E-12 | 1.57E-11 |
| ENSG00000197608 | ZNF841 | 0.49 | 0.08 | 6.18E-11 | 3.63E-10 |
| ENSG00000179361 | ARID3B | 0.49 | 0.08 | 9.61E-10 | 5.19E-09 |
| ENSG00000099804 | CDC34 | 0.49 | 0.08 | 1.86E-09 | 9.83E-09 |
| ENSG00000138495 | COX17 | 0.49 | 0.08 | 2.20E-09 | 1.16E-08 |
| ENSG00000126456 | IRF3 | 0.49 | 0.09 | 1.15E-08 | 5.72E-08 |
| ENSG00000291093 | SVIL-AS1 | 0.49 | 0.09 | 1.48E-08 | 7.30E-08 |
| ENSG00000111786 | SRSF9 | 0.49 | 0.09 | 1.49E-08 | 7.35E-08 |
| ENSG00000107140 | TESK1 | 0.49 | 0.09 | 2.49E-08 | 1.21E-07 |
| ENSG00000069667 | RORA | 0.49 | 0.09 | 2.57E-08 | 1.25E-07 |
| ENSG00000133101 | CCNA1 | 0.49 | 0.09 | 5.74E-08 | 2.71E-07 |
| ENSG00000170638 | TRABD | 0.49 | 0.1 | 1.33E-07 | 6.11E-07 |
| ENSG00000197363 | ZNF517 | 0.49 | 0.1 | 1.91E-07 | 8.69E-07 |
| ENSG00000168101 | NUDT16L1 | 0.49 | 0.1 | 2.51E-07 | 1.13E-06 |
| ENSG00000010404 | IDS | 0.49 | 0.1 | 2.52E-07 | 1.13E-06 |
| ENSG00000062524 | LTK | 0.49 | 0.1 | 3.53E-07 | 1.57E-06 |
| ENSG00000181085 | MAPK15 | 0.49 | 0.11 | 2.17E-06 | 8.93E-06 |
| ENSG00000204991 | SPIRE2 | 0.49 | 0.12 | 1.01E-05 | 3.91E-05 |
| ENSG00000175182 | FAM131A | 0.49 | 0.12 | 1.02E-05 | 3.94E-05 |
| ENSG00000115841 | RMDN2 | 0.49 | 0.12 | 1.86E-05 | 6.98E-05 |
| ENSG00000188848 | BEND4 | 0.49 | 0.12 | 2.34E-05 | 8.73E-05 |

| | | | | | |
|---|---|---|---|---|---|
| ENSG00000277156 | TOMM40P1 | 0.49 | 0.12 | 3.22E-05 | 1.18E-04 |
| ENSG00000233806 | LINC01237 | 0.49 | 0.13 | 5.63E-05 | 2.02E-04 |
| ENSG00000234684 | SDCBP2-AS1 | 0.49 | 0.14 | 1.13E-04 | 3.90E-04 |
| ENSG00000168061 | SAC3D1 | 0.49 | 0.14 | 1.27E-04 | 4.34E-04 |
| ENSG00000146232 | NFKBIE | 0.49 | 0.14 | 2.00E-04 | 6.72E-04 |
| ENSG00000179588 | ZFPM1 | 0.49 | 0.15 | 5.96E-04 | 1.88E-03 |
| ENSG00000166762 | CATSPER2 | 0.49 | 0.18 | 2.25E-03 | 6.47E-03 |
| ENSG00000235070 | | 0.49 | 0.18 | 2.73E-03 | 7.72E-03 |
| ENSG00000146021 | KLHL3 | 0.49 | 0.19 | 2.98E-03 | 8.38E-03 |
| ENSG00000203772 | SPRN | 0.49 | 0.19 | 3.74E-03 | 1.03E-02 |
| ENSG00000205220 | PSMB10 | 0.49 | 0.22 | 8.84E-03 | 2.26E-02 |
| ENSG00000274925 | ZKSCAN2-DT | 0.49 | 0.22 | 8.85E-03 | 2.27E-02 |
| ENSG00000135750 | KCNK1 | 0.49 | 0.22 | 9.23E-03 | 2.35E-02 |
| ENSG00000117480 | FAAH | 0.49 | 0.22 | 9.33E-03 | 2.37E-02 |
| ENSG00000188779 | SKOR1 | 0.49 | 0.22 | 1.04E-02 | 2.63E-02 |
| ENSG00000137393 | RNF144B | 0.49 | 0.23 | 1.29E-02 | 3.17E-02 |
| ENSG00000272138 | LINC01607 | 0.49 | 0.24 | 1.61E-02 | 3.87E-02 |
| ENSG00000099968 | BCL2L13 | 0.48 | 0.04 | 6.32E-32 | 9.02E-31 |
| ENSG00000136068 | FLNB | 0.48 | 0.04 | 8.13E-32 | 1.16E-30 |
| ENSG00000143702 | CEP170 | 0.48 | 0.04 | 1.09E-28 | 1.40E-27 |
| ENSG00000100296 | THOC5 | 0.48 | 0.04 | 1.41E-28 | 1.80E-27 |
| ENSG00000119280 | C1orf198 | 0.48 | 0.05 | 7.65E-24 | 8.28E-23 |
| ENSG00000113732 | ATP6V0E1 | 0.48 | 0.05 | 6.44E-23 | 6.72E-22 |
| ENSG00000118402 | ELOVL4 | 0.48 | 0.05 | 2.53E-22 | 2.58E-21 |
| ENSG00000143753 | DEGS1 | 0.48 | 0.05 | 3.65E-22 | 3.69E-21 |
| ENSG00000123983 | ACSL3 | 0.48 | 0.05 | 2.67E-21 | 2.62E-20 |
| ENSG00000124788 | ATXN1 | 0.48 | 0.05 | 3.13E-21 | 3.07E-20 |
| ENSG00000105438 | KDELR1 | 0.48 | 0.05 | 3.11E-20 | 2.92E-19 |
| ENSG00000156671 | SAMD8 | 0.48 | 0.05 | 2.76E-19 | 2.49E-18 |
| ENSG00000107669 | ATE1 | 0.48 | 0.05 | 3.68E-19 | 3.29E-18 |
| ENSG00000104691 | UBXN8 | 0.48 | 0.06 | 5.10E-18 | 4.34E-17 |
| ENSG00000135503 | ACVR1B | 0.48 | 0.06 | 5.09E-17 | 4.15E-16 |
| ENSG00000114120 | SLC25A36 | 0.48 | 0.06 | 1.08E-16 | 8.66E-16 |
| ENSG00000138081 | FBXO11 | 0.48 | 0.06 | 1.74E-16 | 1.38E-15 |
| ENSG00000155097 | ATP6V1C1 | 0.48 | 0.06 | 1.89E-16 | 1.50E-15 |
| ENSG00000140830 | TXNL4B | 0.48 | 0.06 | 3.10E-16 | 2.44E-15 |
| ENSG00000141682 | PMAIP1 | 0.48 | 0.06 | 3.83E-16 | 2.99E-15 |
| ENSG00000150779 | TIMM8B | 0.48 | 0.06 | 6.12E-16 | 4.75E-15 |
| ENSG00000086619 | ERO1B | 0.48 | 0.06 | 1.35E-14 | 9.80E-14 |
| ENSG00000131375 | CAPN7 | 0.48 | 0.06 | 2.16E-14 | 1.55E-13 |
| ENSG00000087206 | UIMC1 | 0.48 | 0.06 | 2.35E-14 | 1.69E-13 |
| ENSG00000131398 | KCNC3 | 0.48 | 0.07 | 5.30E-14 | 3.74E-13 |
| ENSG00000162433 | AK4 | 0.48 | 0.07 | 1.14E-13 | 7.90E-13 |
| ENSG00000174233 | ADCY6 | 0.48 | 0.07 | 3.72E-13 | 2.51E-12 |
| ENSG00000137522 | RNF121 | 0.48 | 0.07 | 4.02E-13 | 2.70E-12 |
| ENSG00000127952 | STYXL1 | 0.48 | 0.07 | 1.33E-12 | 8.67E-12 |
| ENSG00000105568 | PPP2R1A | 0.48 | 0.07 | 5.11E-12 | 3.21E-11 |
| ENSG00000184164 | CRELD2 | 0.48 | 0.07 | 5.38E-12 | 3.37E-11 |
| ENSG00000170892 | TSEN34 | 0.48 | 0.07 | 5.69E-12 | 3.56E-11 |
| ENSG00000030582 | GRN | 0.48 | 0.07 | 9.34E-12 | 5.78E-11 |

| | | | | | |
|---|---|---|---|---|---|
| ENSG00000101199 | ARFGAP1 | 0.48 | 0.07 | 2.84E-11 | 1.70E-10 |
| ENSG00000141873 | SLC39A3 | 0.48 | 0.07 | 3.83E-11 | 2.28E-10 |
| ENSG00000066651 | TRMT11 | 0.48 | 0.08 | 8.54E-11 | 4.97E-10 |
| ENSG00000258405 | ZNF578 | 0.48 | 0.08 | 1.76E-10 | 1.01E-09 |
| ENSG00000210082 | MT-RNR2 | 0.48 | 0.08 | 6.22E-10 | 3.41E-09 |
| ENSG00000167526 | RPL13 | 0.48 | 0.08 | 6.95E-10 | 3.80E-09 |
| ENSG00000166562 | SEC11C | 0.48 | 0.08 | 8.24E-09 | 4.15E-08 |
| ENSG00000006327 | TNFRSF12A | 0.48 | 0.09 | 2.23E-08 | 1.09E-07 |
| ENSG00000258890 | CEP95 | 0.48 | 0.09 | 3.58E-08 | 1.72E-07 |
| ENSG00000224189 | HAGLR | 0.48 | 0.09 | 6.23E-08 | 2.94E-07 |
| ENSG00000164088 | PPM1M | 0.48 | 0.09 | 1.53E-07 | 6.99E-07 |
| ENSG00000205885 | C1RL-AS1 | 0.48 | 0.1 | 2.62E-07 | 1.17E-06 |
| ENSG00000274265 | | 0.48 | 0.12 | 1.04E-05 | 4.00E-05 |
| ENSG00000164897 | TMUB1 | 0.48 | 0.12 | 2.06E-05 | 7.73E-05 |
| ENSG00000257727 | CNPY2 | 0.48 | 0.13 | 5.90E-05 | 2.11E-04 |
| ENSG00000174021 | GNG5 | 0.48 | 0.13 | 6.92E-05 | 2.46E-04 |
| ENSG00000147889 | CDKN2A | 0.48 | 0.15 | 5.64E-04 | 1.78E-03 |
| ENSG00000124140 | SLC12A5 | 0.48 | 0.16 | 8.79E-04 | 2.69E-03 |
| ENSG00000165115 | KIF27 | 0.48 | 0.16 | 1.26E-03 | 3.76E-03 |
| ENSG00000272849 | | 0.48 | 0.17 | 1.81E-03 | 5.28E-03 |
| ENSG00000286545 | | 0.48 | 0.18 | 2.59E-03 | 7.35E-03 |
| ENSG00000115257 | PCSK4 | 0.48 | 0.2 | 6.17E-03 | 1.63E-02 |
| ENSG00000253837 | LOXL2-AS1 | 0.48 | 0.21 | 7.79E-03 | 2.02E-02 |
| ENSG00000237036 | ZEB1-AS1 | 0.48 | 0.21 | 8.99E-03 | 2.30E-02 |
| ENSG00000225733 | FGD5-AS1 | 0.47 | 0.04 | 2.26E-40 | 4.08E-39 |
| ENSG00000136238 | RAC1 | 0.47 | 0.04 | 1.72E-39 | 3.05E-38 |
| ENSG00000100030 | MAPK1 | 0.47 | 0.04 | 5.71E-36 | 9.24E-35 |
| ENSG00000166986 | MARS1 | 0.47 | 0.04 | 1.15E-35 | 1.85E-34 |
| ENSG00000162434 | JAK1 | 0.47 | 0.04 | 2.87E-29 | 3.77E-28 |
| ENSG00000156502 | SUPV3L1 | 0.47 | 0.04 | 1.11E-28 | 1.42E-27 |
| ENSG00000181555 | SETD2 | 0.47 | 0.05 | 1.73E-24 | 1.91E-23 |
| ENSG00000138443 | ABI2 | 0.47 | 0.05 | 1.03E-23 | 1.11E-22 |
| ENSG00000074695 | LMAN1 | 0.47 | 0.05 | 3.98E-22 | 4.02E-21 |
| ENSG00000111897 | SERINC1 | 0.47 | 0.05 | 4.64E-22 | 4.69E-21 |
| ENSG00000116030 | SUMO1 | 0.47 | 0.05 | 6.43E-22 | 6.46E-21 |
| ENSG00000164307 | ERAP1 | 0.47 | 0.05 | 7.51E-22 | 7.51E-21 |
| ENSG00000165688 | PMPCA | 0.47 | 0.05 | 1.05E-21 | 1.05E-20 |
| ENSG00000114331 | ACAP2 | 0.47 | 0.05 | 3.89E-20 | 3.63E-19 |
| ENSG00000023318 | ERP44 | 0.47 | 0.05 | 4.76E-20 | 4.43E-19 |
| ENSG00000131943 | C19orf12 | 0.47 | 0.05 | 1.21E-19 | 1.10E-18 |
| ENSG00000162642 | C1orf52 | 0.47 | 0.05 | 2.25E-19 | 2.04E-18 |
| ENSG00000121289 | CEP89 | 0.47 | 0.05 | 2.83E-19 | 2.55E-18 |
| ENSG00000138764 | CCNG2 | 0.47 | 0.05 | 2.87E-19 | 2.59E-18 |
| ENSG00000109320 | NFKB1 | 0.47 | 0.05 | 5.20E-19 | 4.62E-18 |
| ENSG00000143321 | HDGF | 0.47 | 0.05 | 5.85E-19 | 5.18E-18 |
| ENSG00000169126 | ODAD2 | 0.47 | 0.06 | 8.84E-17 | 7.14E-16 |
| ENSG00000141867 | BRD4 | 0.47 | 0.06 | 6.01E-16 | 4.66E-15 |
| ENSG00000234745 | HLA-B | 0.47 | 0.06 | 1.05E-15 | 8.02E-15 |
| ENSG00000118217 | ATF6 | 0.47 | 0.06 | 2.03E-15 | 1.54E-14 |
| ENSG00000085365 | SCAMP1 | 0.47 | 0.06 | 9.36E-15 | 6.87E-14 |

| | | | | | |
|---|---|---|---|---|---|
| ENSG00000204291 | COL15A1 | 0.47 | 0.06 | 2.10E-14 | 1.51E-13 |
| ENSG00000152443 | ZNF776 | 0.47 | 0.06 | 2.22E-14 | 1.60E-13 |
| ENSG00000103549 | RNF40 | 0.47 | 0.06 | 3.96E-14 | 2.81E-13 |
| ENSG00000164038 | SLC9B2 | 0.47 | 0.07 | 2.19E-13 | 1.50E-12 |
| ENSG00000198498 | TMA16 | 0.47 | 0.07 | 1.63E-12 | 1.05E-11 |
| ENSG00000204954 | UQCC6 | 0.47 | 0.07 | 2.94E-12 | 1.88E-11 |
| ENSG00000111676 | ATN1 | 0.47 | 0.07 | 5.59E-12 | 3.50E-11 |
| ENSG00000053501 | USE1 | 0.47 | 0.07 | 3.38E-11 | 2.01E-10 |
| ENSG00000187713 | TMEM203 | 0.47 | 0.07 | 6.14E-11 | 3.61E-10 |
| ENSG00000243943 | ZNF512 | 0.47 | 0.07 | 6.43E-11 | 3.77E-10 |
| ENSG00000171206 | TRIM8 | 0.47 | 0.07 | 1.49E-10 | 8.55E-10 |
| ENSG00000214253 | FIS1 | 0.47 | 0.08 | 2.13E-10 | 1.21E-09 |
| ENSG00000075399 | VPS9D1 | 0.47 | 0.08 | 2.43E-10 | 1.38E-09 |
| ENSG00000105726 | ATP13A1 | 0.47 | 0.08 | 3.24E-10 | 1.81E-09 |
| ENSG00000101216 | GMEB2 | 0.47 | 0.08 | 4.84E-10 | 2.68E-09 |
| ENSG00000181788 | SIAH2 | 0.47 | 0.08 | 5.49E-10 | 3.02E-09 |
| ENSG00000102901 | CENPT | 0.47 | 0.08 | 1.27E-09 | 6.83E-09 |
| ENSG00000033100 | CHPF2 | 0.47 | 0.08 | 1.65E-09 | 8.75E-09 |
| ENSG00000138613 | APH1B | 0.47 | 0.08 | 1.89E-09 | 9.99E-09 |
| ENSG00000233927 | RPS28 | 0.47 | 0.08 | 2.13E-09 | 1.12E-08 |
| ENSG00000168234 | TTC39C | 0.47 | 0.08 | 4.26E-09 | 2.19E-08 |
| ENSG00000264247 | ZNF407-AS1 | 0.47 | 0.08 | 4.72E-09 | 2.41E-08 |
| ENSG00000186665 | C17orf58 | 0.47 | 0.08 | 1.43E-08 | 7.08E-08 |
| ENSG00000166313 | APBB1 | 0.47 | 0.09 | 1.79E-08 | 8.77E-08 |
| ENSG00000065665 | SEC61A2 | 0.47 | 0.09 | 4.09E-08 | 1.95E-07 |
| ENSG00000176170 | SPHK1 | 0.47 | 0.09 | 8.76E-08 | 4.08E-07 |
| ENSG00000161179 | YDJC | 0.47 | 0.09 | 3.48E-07 | 1.55E-06 |
| ENSG00000260196 | | 0.47 | 0.1 | 1.10E-06 | 4.66E-06 |
| ENSG00000100330 | MTMR3 | 0.47 | 0.1 | 1.11E-06 | 4.67E-06 |
| ENSG00000100300 | TSPO | 0.47 | 0.1 | 1.27E-06 | 5.33E-06 |
| ENSG00000144029 | MRPS5 | 0.47 | 0.11 | 5.66E-06 | 2.24E-05 |
| ENSG00000156958 | GALK2 | 0.47 | 0.12 | 2.24E-05 | 8.35E-05 |
| ENSG00000127415 | IDUA | 0.47 | 0.12 | 3.17E-05 | 1.17E-04 |
| ENSG00000291066 | | 0.47 | 0.12 | 3.64E-05 | 1.33E-04 |
| ENSG00000267534 | S1PR2 | 0.47 | 0.12 | 4.93E-05 | 1.77E-04 |
| ENSG00000273599 | | 0.47 | 0.13 | 8.76E-05 | 3.07E-04 |
| ENSG00000126005 | MMP24OS | 0.47 | 0.13 | 1.85E-04 | 6.22E-04 |
| ENSG00000177943 | MAMDC4 | 0.47 | 0.13 | 1.88E-04 | 6.32E-04 |
| ENSG00000279170 | TSTD3 | 0.47 | 0.14 | 2.72E-04 | 8.97E-04 |
| ENSG00000272994 | C2orf49-DT | 0.47 | 0.15 | 6.32E-04 | 1.98E-03 |
| ENSG00000009724 | MASP2 | 0.47 | 0.15 | 8.60E-04 | 2.64E-03 |
| ENSG00000131495 | NDUFA2 | 0.47 | 0.15 | 1.07E-03 | 3.23E-03 |
| ENSG00000174276 | ZNHIT2 | 0.47 | 0.16 | 1.08E-03 | 3.26E-03 |
| ENSG00000188483 | IER5L | 0.47 | 0.16 | 1.20E-03 | 3.59E-03 |
| ENSG00000124370 | MCEE | 0.47 | 0.16 | 1.43E-03 | 4.25E-03 |
| ENSG00000143674 | MAP3K21 | 0.47 | 0.17 | 2.79E-03 | 7.89E-03 |
| ENSG00000290554 | ADCY10P1 | 0.47 | 0.19 | 4.79E-03 | 1.29E-02 |
| ENSG00000168939 | SPRY3 | 0.47 | 0.2 | 7.08E-03 | 1.85E-02 |
| ENSG00000290535 | | 0.47 | 0.2 | 7.12E-03 | 1.86E-02 |
| ENSG00000221955 | SLC12A8 | 0.47 | 0.2 | 8.13E-03 | 2.10E-02 |

| | | | | | |
|---|---|---|---|---|---|
| ENSG00000158023 | CFAP251 | 0.47 | 0.21 | 9.46E-03 | 2.40E-02 |
| ENSG00000100902 | PSMA6 | 0.47 | 0.21 | 9.57E-03 | 2.43E-02 |
| ENSG00000162426 | SLC45A1 | 0.47 | 0.21 | 1.11E-02 | 2.78E-02 |
| ENSG00000167397 | VKORC1 | 0.47 | 0.21 | 1.13E-02 | 2.81E-02 |
| ENSG00000109846 | CRYAB | 0.47 | 0.22 | 1.39E-02 | 3.39E-02 |
| ENSG00000184986 | TMEM121 | 0.47 | 0.22 | 1.39E-02 | 3.40E-02 |
| ENSG00000228274 | | 0.47 | 0.23 | 1.50E-02 | 3.63E-02 |
| ENSG00000260751 | | 0.47 | 0.24 | 1.96E-02 | 4.64E-02 |
| ENSG00000113615 | SEC24A | 0.46 | 0.04 | 4.47E-31 | 6.24E-30 |
| ENSG00000164244 | PRRC1 | 0.46 | 0.04 | 4.29E-25 | 4.86E-24 |
| ENSG00000113194 | FAF2 | 0.46 | 0.05 | 2.29E-24 | 2.52E-23 |
| ENSG00000134440 | NARS1 | 0.46 | 0.05 | 6.44E-23 | 6.72E-22 |
| ENSG00000155660 | PDIA4 | 0.46 | 0.05 | 4.37E-21 | 4.26E-20 |
| ENSG00000177885 | GRB2 | 0.46 | 0.05 | 4.89E-21 | 4.75E-20 |
| ENSG00000136021 | SCYL2 | 0.46 | 0.05 | 8.23E-21 | 7.91E-20 |
| ENSG00000157827 | FMNL2 | 0.46 | 0.05 | 1.10E-20 | 1.05E-19 |
| ENSG00000091317 | CMTM6 | 0.46 | 0.05 | 1.48E-20 | 1.40E-19 |
| ENSG00000169375 | SIN3A | 0.46 | 0.05 | 3.87E-19 | 3.46E-18 |
| ENSG00000077232 | DNAJC10 | 0.46 | 0.05 | 4.67E-19 | 4.15E-18 |
| ENSG00000006451 | RALA | 0.46 | 0.05 | 2.86E-18 | 2.45E-17 |
| ENSG00000170855 | TRIAP1 | 0.46 | 0.05 | 8.22E-18 | 6.93E-17 |
| ENSG00000102401 | ARMCX3 | 0.46 | 0.06 | 1.48E-16 | 1.18E-15 |
| ENSG00000100239 | PPP6R2 | 0.46 | 0.06 | 1.90E-16 | 1.51E-15 |
| ENSG00000116604 | MEF2D | 0.46 | 0.06 | 4.89E-16 | 3.81E-15 |
| ENSG00000105819 | PMPCB | 0.46 | 0.06 | 6.06E-16 | 4.70E-15 |
| ENSG00000024048 | UBR2 | 0.46 | 0.06 | 8.53E-16 | 6.55E-15 |
| ENSG00000170242 | USP47 | 0.46 | 0.06 | 8.93E-16 | 6.84E-15 |
| ENSG00000141562 | NARF | 0.46 | 0.06 | 1.73E-15 | 1.31E-14 |
| ENSG00000126214 | KLC1 | 0.46 | 0.06 | 1.76E-14 | 1.27E-13 |
| ENSG00000221988 | PPT2 | 0.46 | 0.06 | 1.27E-13 | 8.77E-13 |
| ENSG00000013016 | EHD3 | 0.46 | 0.07 | 1.12E-12 | 7.31E-12 |
| ENSG00000106605 | BLVRA | 0.46 | 0.07 | 1.19E-12 | 7.74E-12 |
| ENSG00000087448 | KLHL42 | 0.46 | 0.07 | 1.88E-12 | 1.22E-11 |
| ENSG00000170906 | NDUFA3 | 0.46 | 0.07 | 3.27E-12 | 2.08E-11 |
| ENSG00000083750 | RRAGB | 0.46 | 0.07 | 3.68E-12 | 2.34E-11 |
| ENSG00000125814 | NAPB | 0.46 | 0.07 | 4.22E-12 | 2.67E-11 |
| ENSG00000176595 | KBTBD11 | 0.46 | 0.07 | 7.26E-12 | 4.51E-11 |
| ENSG00000117122 | MFAP2 | 0.46 | 0.07 | 1.27E-11 | 7.80E-11 |
| ENSG00000115947 | ORC4 | 0.46 | 0.07 | 1.69E-11 | 1.03E-10 |
| ENSG00000134285 | FKBP11 | 0.46 | 0.07 | 5.33E-11 | 3.15E-10 |
| ENSG00000103502 | CDIPT | 0.46 | 0.07 | 8.16E-11 | 4.76E-10 |
| ENSG00000118197 | DDX59 | 0.46 | 0.07 | 1.85E-10 | 1.06E-09 |
| ENSG00000034677 | RNF19A | 0.46 | 0.08 | 1.53E-09 | 8.13E-09 |
| ENSG00000168763 | CNNM3 | 0.46 | 0.08 | 1.65E-09 | 8.76E-09 |
| ENSG00000105402 | NAPA | 0.46 | 0.08 | 5.78E-09 | 2.94E-08 |
| ENSG00000117984 | CTSD | 0.46 | 0.09 | 7.52E-08 | 3.52E-07 |
| ENSG00000177674 | AGTRAP | 0.46 | 0.09 | 8.24E-08 | 3.85E-07 |
| ENSG00000147996 | ZNG1E | 0.46 | 0.09 | 1.22E-07 | 5.60E-07 |
| ENSG00000188185 | LINC00265 | 0.46 | 0.09 | 3.85E-07 | 1.70E-06 |
| ENSG00000123353 | ORMDL2 | 0.46 | 0.1 | 8.25E-07 | 3.53E-06 |

| | | | | | |
|---|---|---|---|---|---|
| ENSG00000063169 | BICRA | 0.46 | 0.1 | 3.66E-06 | 1.48E-05 |
| ENSG00000160050 | CCDC28B | 0.46 | 0.11 | 1.00E-05 | 3.88E-05 |
| ENSG00000137825 | ITPKA | 0.46 | 0.13 | 1.42E-04 | 4.84E-04 |
| ENSG00000167550 | RHEBL1 | 0.46 | 0.14 | 3.85E-04 | 1.24E-03 |
| ENSG00000160323 | ADAMTS13 | 0.46 | 0.14 | 3.88E-04 | 1.25E-03 |
| ENSG00000229119 | RPLP0P9 | 0.46 | 0.14 | 5.88E-04 | 1.85E-03 |
| ENSG00000180806 | HOXC9 | 0.46 | 0.15 | 9.15E-04 | 2.80E-03 |
| ENSG00000289843 | | 0.46 | 0.15 | 1.18E-03 | 3.56E-03 |
| ENSG00000270069 | MIR222HG | 0.46 | 0.16 | 2.17E-03 | 6.25E-03 |
| ENSG00000189409 | MMP23B | 0.46 | 0.18 | 3.71E-03 | 1.02E-02 |
| ENSG00000189227 | C15orf61 | 0.46 | 0.18 | 3.72E-03 | 1.03E-02 |
| ENSG00000154217 | PITPNC1 | 0.46 | 0.18 | 3.76E-03 | 1.04E-02 |
| ENSG00000224699 | LAMTOR5-AS1 | 0.46 | 0.18 | 4.86E-03 | 1.31E-02 |
| ENSG00000266969 | CEP192-DT | 0.46 | 0.19 | 5.32E-03 | 1.42E-02 |
| ENSG00000275734 | | 0.46 | 0.18 | 5.70E-03 | 1.52E-02 |
| ENSG00000264575 | LINC00526 | 0.46 | 0.2 | 1.05E-02 | 2.63E-02 |
| ENSG00000101180 | HRH3 | 0.46 | 0.21 | 1.13E-02 | 2.82E-02 |
| ENSG00000084628 | NKAIN1 | 0.46 | 0.21 | 1.29E-02 | 3.17E-02 |
| ENSG00000230409 | TCEA1P2 | 0.46 | 0.22 | 1.57E-02 | 3.78E-02 |
| ENSG00000291156 | | 0.46 | 0.22 | 1.62E-02 | 3.88E-02 |
| ENSG00000279407 | | 0.46 | 0.23 | 1.82E-02 | 4.32E-02 |
| ENSG00000213221 | DNLZ | 0.46 | 0.23 | 2.00E-02 | 4.71E-02 |
| ENSG00000116285 | ERRFI1 | 0.45 | 0.04 | 1.02E-28 | 1.32E-27 |
| ENSG00000125304 | TM9SF2 | 0.45 | 0.04 | 1.35E-26 | 1.62E-25 |
| ENSG00000113319 | RASGRF2 | 0.45 | 0.04 | 2.27E-26 | 2.69E-25 |
| ENSG00000072364 | AFF4 | 0.45 | 0.05 | 1.49E-23 | 1.60E-22 |
| ENSG00000182670 | TTC3 | 0.45 | 0.05 | 1.89E-23 | 2.01E-22 |
| ENSG00000067167 | TRAM1 | 0.45 | 0.05 | 3.51E-22 | 3.55E-21 |
| ENSG00000117335 | CD46 | 0.45 | 0.05 | 4.30E-22 | 4.35E-21 |
| ENSG00000122550 | KLHL7 | 0.45 | 0.05 | 6.36E-20 | 5.89E-19 |
| ENSG00000127526 | SLC35E1 | 0.45 | 0.05 | 1.09E-19 | 1.00E-18 |
| ENSG00000186594 | MIR22HG | 0.45 | 0.05 | 1.42E-18 | 1.23E-17 |
| ENSG00000169914 | OTUD3 | 0.45 | 0.05 | 1.85E-17 | 1.54E-16 |
| ENSG00000170456 | DENND5B | 0.45 | 0.05 | 2.62E-17 | 2.17E-16 |
| ENSG00000111647 | BLTP3B | 0.45 | 0.06 | 1.02E-16 | 8.20E-16 |
| ENSG00000169217 | CD2BP2 | 0.45 | 0.06 | 1.23E-16 | 9.87E-16 |
| ENSG00000168288 | MMADHC | 0.45 | 0.06 | 7.31E-16 | 5.63E-15 |
| ENSG00000155744 | HYCC2 | 0.45 | 0.06 | 8.66E-16 | 6.64E-15 |
| ENSG00000128524 | ATP6V1F | 0.45 | 0.06 | 4.65E-15 | 3.46E-14 |
| ENSG00000164949 | GEM | 0.45 | 0.06 | 1.01E-14 | 7.39E-14 |
| ENSG00000175115 | PACS1 | 0.45 | 0.06 | 2.98E-14 | 2.13E-13 |
| ENSG00000164054 | SHISA5 | 0.45 | 0.06 | 4.76E-14 | 3.36E-13 |
| ENSG00000101004 | NINL | 0.45 | 0.06 | 1.40E-13 | 9.66E-13 |
| ENSG00000083799 | CYLD | 0.45 | 0.06 | 1.65E-13 | 1.13E-12 |
| ENSG00000070081 | NUCB2 | 0.45 | 0.06 | 1.04E-12 | 6.81E-12 |
| ENSG00000122042 | UBL3 | 0.45 | 0.07 | 2.00E-12 | 1.29E-11 |
| ENSG00000142784 | WDTC1 | 0.45 | 0.07 | 7.82E-12 | 4.85E-11 |
| ENSG00000169136 | ATF5 | 0.45 | 0.07 | 1.03E-11 | 6.36E-11 |
| ENSG00000147905 | ZCCHC7 | 0.45 | 0.07 | 3.26E-11 | 1.95E-10 |
| ENSG00000140836 | ZFHX3 | 0.45 | 0.07 | 3.84E-11 | 2.28E-10 |

| | | | | | |
|---|---|---|---|---|---|
| ENSG00000100441 | KHNYN | 0.45 | 0.07 | 5.68E-11 | 3.34E-10 |
| ENSG00000178385 | PLEKHM3 | 0.45 | 0.07 | 1.19E-10 | 6.85E-10 |
| ENSG00000104980 | TIMM44 | 0.45 | 0.07 | 2.26E-10 | 1.28E-09 |
| ENSG00000149084 | HSD17B12 | 0.45 | 0.07 | 2.48E-10 | 1.40E-09 |
| ENSG00000163512 | AZI2 | 0.45 | 0.07 | 3.07E-10 | 1.72E-09 |
| ENSG00000197122 | SRC | 0.45 | 0.07 | 4.31E-10 | 2.39E-09 |
| ENSG00000172428 | COPS9 | 0.45 | 0.08 | 8.54E-10 | 4.63E-09 |
| ENSG00000084731 | KIF3C | 0.45 | 0.08 | 1.07E-09 | 5.77E-09 |
| ENSG00000177058 | SLC38A9 | 0.45 | 0.08 | 8.52E-09 | 4.28E-08 |
| ENSG00000141384 | TAF4B | 0.45 | 0.08 | 1.39E-08 | 6.86E-08 |
| ENSG00000291136 | | 0.45 | 0.08 | 2.91E-08 | 1.41E-07 |
| ENSG00000168701 | TMEM208 | 0.45 | 0.08 | 3.60E-08 | 1.72E-07 |
| ENSG00000168056 | LTBP3 | 0.45 | 0.08 | 3.89E-08 | 1.86E-07 |
| ENSG00000161996 | WDR90 | 0.45 | 0.09 | 9.13E-08 | 4.25E-07 |
| ENSG00000145979 | TBC1D7 | 0.45 | 0.09 | 9.97E-08 | 4.63E-07 |
| ENSG00000186104 | CYP2R1 | 0.45 | 0.09 | 1.84E-07 | 8.36E-07 |
| ENSG00000100084 | HIRA | 0.45 | 0.09 | 2.11E-07 | 9.53E-07 |
| ENSG00000145916 | RMND5B | 0.45 | 0.09 | 2.36E-07 | 1.06E-06 |
| ENSG00000133460 | SLC2A11 | 0.45 | 0.09 | 3.30E-07 | 1.47E-06 |
| ENSG00000258441 | LINC00641 | 0.45 | 0.09 | 3.91E-07 | 1.73E-06 |
| ENSG00000171045 | TSNARE1 | 0.45 | 0.09 | 5.38E-07 | 2.35E-06 |
| ENSG00000166450 | PRTG | 0.45 | 0.09 | 6.86E-07 | 2.96E-06 |
| ENSG00000100151 | PICK1 | 0.45 | 0.1 | 9.17E-07 | 3.91E-06 |
| ENSG00000138678 | GPAT3 | 0.45 | 0.1 | 2.45E-06 | 1.00E-05 |
| ENSG00000144134 | RABL2A | 0.45 | 0.1 | 3.93E-06 | 1.58E-05 |
| ENSG00000177272 | KCNA3 | 0.45 | 0.1 | 4.04E-06 | 1.62E-05 |
| ENSG00000198740 | ZNF652 | 0.45 | 0.1 | 6.82E-06 | 2.68E-05 |
| ENSG00000173681 | BCLAF3 | 0.45 | 0.11 | 2.38E-05 | 8.85E-05 |
| ENSG00000236552 | RPL13AP5 | 0.45 | 0.11 | 2.97E-05 | 1.09E-04 |
| ENSG00000163376 | KBTBD8 | 0.45 | 0.12 | 5.12E-05 | 1.84E-04 |
| ENSG00000214106 | PAXIP1-AS2 | 0.45 | 0.12 | 7.76E-05 | 2.74E-04 |
| ENSG00000149150 | SLC43A1 | 0.45 | 0.12 | 1.03E-04 | 3.57E-04 |
| ENSG00000128872 | TMOD2 | 0.45 | 0.13 | 1.92E-04 | 6.46E-04 |
| ENSG00000079313 | REXO1 | 0.45 | 0.14 | 7.44E-04 | 2.31E-03 |
| ENSG00000159214 | CCDC24 | 0.45 | 0.16 | 1.92E-03 | 5.59E-03 |
| ENSG00000162496 | DHRS3 | 0.45 | 0.16 | 2.01E-03 | 5.82E-03 |
| ENSG00000184047 | DIABLO | 0.45 | 0.16 | 2.60E-03 | 7.39E-03 |
| ENSG00000272341 | | 0.45 | 0.17 | 3.38E-03 | 9.40E-03 |
| ENSG00000130429 | ARPC1B | 0.45 | 0.17 | 4.16E-03 | 1.14E-02 |
| ENSG00000244687 | UBE2V1 | 0.45 | 0.18 | 4.38E-03 | 1.19E-02 |
| ENSG00000225031 | EIF4BP7 | 0.45 | 0.21 | 1.41E-02 | 3.43E-02 |
| ENSG00000270012 | | 0.45 | 0.22 | 1.83E-02 | 4.35E-02 |
| ENSG00000205572 | SERF1B | 0.45 | 0.23 | 2.00E-02 | 4.71E-02 |
| ENSG00000135931 | ARMC9 | 0.44 | 0.04 | 2.55E-35 | 4.05E-34 |
| ENSG00000136813 | ECPAS | 0.44 | 0.04 | 3.02E-26 | 3.55E-25 |
| ENSG00000008256 | CYTH3 | 0.44 | 0.05 | 4.96E-23 | 5.20E-22 |
| ENSG00000070018 | LRP6 | 0.44 | 0.05 | 1.27E-20 | 1.21E-19 |
| ENSG00000134851 | TMEM165 | 0.44 | 0.05 | 9.81E-20 | 9.02E-19 |
| ENSG00000065154 | OAT | 0.44 | 0.05 | 4.50E-19 | 4.01E-18 |
| ENSG00000153561 | RMND5A | 0.44 | 0.05 | 1.80E-18 | 1.56E-17 |

| | | | | | |
|---|---|---|---|---|---|
| ENSG00000160209 | PDXK | 0.44 | 0.05 | 6.11E-18 | 5.17E-17 |
| ENSG00000104765 | BNIP3L | 0.44 | 0.05 | 6.11E-18 | 5.18E-17 |
| ENSG00000078246 | TULP3 | 0.44 | 0.05 | 6.69E-18 | 5.66E-17 |
| ENSG00000124209 | RAB22A | 0.44 | 0.05 | 6.41E-17 | 5.22E-16 |
| ENSG00000108828 | VAT1 | 0.44 | 0.06 | 5.78E-15 | 4.28E-14 |
| ENSG00000115128 | SF3B6 | 0.44 | 0.06 | 2.64E-13 | 1.79E-12 |
| ENSG00000105254 | TBCB | 0.44 | 0.06 | 3.81E-13 | 2.56E-12 |
| ENSG00000213676 | ATF6B | 0.44 | 0.06 | 7.80E-13 | 5.16E-12 |
| ENSG00000131941 | RHPN2 | 0.44 | 0.06 | 6.35E-12 | 3.96E-11 |
| ENSG00000204116 | CHIC1 | 0.44 | 0.07 | 1.72E-11 | 1.05E-10 |
| ENSG00000156110 | ADK | 0.44 | 0.07 | 2.07E-11 | 1.26E-10 |
| ENSG00000065600 | PACC1 | 0.44 | 0.07 | 3.30E-11 | 1.97E-10 |
| ENSG00000105287 | PRKD2 | 0.44 | 0.07 | 8.70E-11 | 5.06E-10 |
| ENSG00000285967 | NIPBL-DT | 0.44 | 0.07 | 9.92E-11 | 5.76E-10 |
| ENSG00000173801 | JUP | 0.44 | 0.07 | 1.17E-10 | 6.76E-10 |
| ENSG00000127948 | POR | 0.44 | 0.07 | 1.54E-10 | 8.84E-10 |
| ENSG00000090263 | MRPS33 | 0.44 | 0.07 | 2.93E-10 | 1.64E-09 |
| ENSG00000167766 | ZNF83 | 0.44 | 0.07 | 4.77E-10 | 2.64E-09 |
| ENSG00000269556 | TMEM185A | 0.44 | 0.07 | 5.96E-10 | 3.27E-09 |
| ENSG00000152439 | ZNF773 | 0.44 | 0.07 | 1.37E-09 | 7.30E-09 |
| ENSG00000099246 | RAB18 | 0.44 | 0.07 | 1.93E-09 | 1.02E-08 |
| ENSG00000204220 | PFDN6 | 0.44 | 0.08 | 6.30E-09 | 3.20E-08 |
| ENSG00000146063 | TRIM41 | 0.44 | 0.08 | 7.93E-09 | 4.00E-08 |
| ENSG00000131373 | HACL1 | 0.44 | 0.08 | 1.21E-08 | 6.00E-08 |
| ENSG00000116871 | MAP7D1 | 0.44 | 0.08 | 1.54E-08 | 7.60E-08 |
| ENSG00000118473 | SGIP1 | 0.44 | 0.08 | 1.77E-08 | 8.68E-08 |
| ENSG00000112511 | PHF1 | 0.44 | 0.08 | 4.60E-08 | 2.19E-07 |
| ENSG00000182134 | TDRKH | 0.44 | 0.08 | 9.39E-08 | 4.36E-07 |
| ENSG00000113391 | FAM172A | 0.44 | 0.09 | 1.56E-07 | 7.11E-07 |
| ENSG00000169857 | AVEN | 0.44 | 0.09 | 5.11E-07 | 2.23E-06 |
| ENSG00000114859 | CLCN2 | 0.44 | 0.09 | 7.45E-07 | 3.20E-06 |
| ENSG00000147799 | ARHGAP39 | 0.44 | 0.09 | 9.98E-07 | 4.24E-06 |
| ENSG00000198171 | DDRGK1 | 0.44 | 0.1 | 1.84E-06 | 7.60E-06 |
| ENSG00000102780 | DGKH | 0.44 | 0.1 | 5.84E-06 | 2.31E-05 |
| ENSG00000061656 | SPAG4 | 0.44 | 0.1 | 7.18E-06 | 2.81E-05 |
| ENSG00000290865 | RRN3P3 | 0.44 | 0.1 | 1.14E-05 | 4.39E-05 |
| ENSG00000116815 | CD58 | 0.44 | 0.11 | 2.64E-05 | 9.77E-05 |
| ENSG00000237940 | LINC01238 | 0.44 | 0.12 | 1.10E-04 | 3.81E-04 |
| ENSG00000269918 | | 0.44 | 0.14 | 8.59E-04 | 2.64E-03 |
| ENSG00000151967 | SCHIP1 | 0.44 | 0.15 | 2.16E-03 | 6.23E-03 |
| ENSG00000271533 | | 0.44 | 0.16 | 3.41E-03 | 9.48E-03 |
| ENSG00000167371 | PRRT2 | 0.44 | 0.17 | 5.05E-03 | 1.36E-02 |
| ENSG00000269352 | PTOV1-AS2 | 0.44 | 0.18 | 5.65E-03 | 1.51E-02 |
| ENSG00000222036 | POTEM | 0.44 | 0.18 | 6.63E-03 | 1.74E-02 |
| ENSG00000187537 | POTEG | 0.44 | 0.18 | 8.01E-03 | 2.07E-02 |
| ENSG00000214595 | EML6 | 0.44 | 0.19 | 8.28E-03 | 2.13E-02 |
| ENSG00000260597 | | 0.44 | 0.2 | 1.27E-02 | 3.13E-02 |
| ENSG00000227615 | | 0.44 | 0.21 | 1.54E-02 | 3.72E-02 |
| ENSG00000143384 | MCL1 | 0.43 | 0.03 | 2.10E-40 | 3.81E-39 |
| ENSG00000175970 | UNC119B | 0.43 | 0.04 | 3.68E-30 | 4.97E-29 |

| | | | | | |
|---|---|---|---|---|---|
| ENSG00000141252 | VPS53 | 0.43 | 0.04 | 1.16E-22 | 1.20E-21 |
| ENSG00000110048 | OSBP | 0.43 | 0.05 | 3.36E-21 | 3.29E-20 |
| ENSG00000164294 | GPX8 | 0.43 | 0.05 | 6.93E-21 | 6.68E-20 |
| ENSG00000182985 | CADM1 | 0.43 | 0.05 | 1.29E-20 | 1.23E-19 |
| ENSG00000066117 | SMARCD1 | 0.43 | 0.05 | 1.63E-20 | 1.55E-19 |
| ENSG00000171862 | PTEN | 0.43 | 0.05 | 3.44E-20 | 3.23E-19 |
| ENSG00000164122 | ASB5 | 0.43 | 0.05 | 4.37E-20 | 4.08E-19 |
| ENSG00000187742 | SECISBP2 | 0.43 | 0.05 | 3.45E-19 | 3.10E-18 |
| ENSG00000028203 | VEZT | 0.43 | 0.05 | 5.19E-17 | 4.24E-16 |
| ENSG00000103187 | COTL1 | 0.43 | 0.05 | 7.03E-16 | 5.43E-15 |
| ENSG00000072840 | EVC | 0.43 | 0.05 | 8.34E-16 | 6.40E-15 |
| ENSG00000165476 | REEP3 | 0.43 | 0.05 | 2.73E-15 | 2.05E-14 |
| ENSG00000240230 | COX19 | 0.43 | 0.06 | 3.13E-15 | 2.35E-14 |
| ENSG00000143756 | FBXO28 | 0.43 | 0.06 | 6.53E-15 | 4.83E-14 |
| ENSG00000064961 | HMG20B | 0.43 | 0.06 | 8.11E-15 | 5.97E-14 |
| ENSG00000166783 | MARF1 | 0.43 | 0.06 | 8.73E-14 | 6.09E-13 |
| ENSG00000137642 | SORL1 | 0.43 | 0.06 | 1.31E-13 | 9.05E-13 |
| ENSG00000170291 | ELP5 | 0.43 | 0.06 | 5.18E-12 | 3.25E-11 |
| ENSG00000289234 | ZNF496-DT | 0.43 | 0.07 | 2.37E-11 | 1.43E-10 |
| ENSG00000134824 | FADS2 | 0.43 | 0.06 | 2.40E-11 | 1.45E-10 |
| ENSG00000144566 | RAB5A | 0.43 | 0.07 | 2.87E-11 | 1.72E-10 |
| ENSG00000204308 | RNF5 | 0.43 | 0.07 | 5.90E-11 | 3.47E-10 |
| ENSG00000183617 | MRPL54 | 0.43 | 0.07 | 1.07E-10 | 6.22E-10 |
| ENSG00000090061 | CCNK | 0.43 | 0.07 | 4.10E-10 | 2.27E-09 |
| ENSG00000160908 | ZNF394 | 0.43 | 0.07 | 1.17E-09 | 6.29E-09 |
| ENSG00000125995 | ROMO1 | 0.43 | 0.08 | 3.79E-09 | 1.96E-08 |
| ENSG00000268858 | | 0.43 | 0.07 | 5.29E-09 | 2.70E-08 |
| ENSG00000138600 | SPPL2A | 0.43 | 0.08 | 1.83E-08 | 8.96E-08 |
| ENSG00000101928 | MOSPD1 | 0.43 | 0.08 | 2.95E-08 | 1.42E-07 |
| ENSG00000140264 | SERF2 | 0.43 | 0.08 | 5.66E-08 | 2.68E-07 |
| ENSG00000121413 | ZSCAN18 | 0.43 | 0.08 | 1.01E-07 | 4.70E-07 |
| ENSG00000198270 | TMEM116 | 0.43 | 0.08 | 1.04E-07 | 4.82E-07 |
| ENSG00000172354 | GNB2 | 0.43 | 0.08 | 1.07E-07 | 4.95E-07 |
| ENSG00000185033 | SEMA4B | 0.43 | 0.08 | 1.55E-07 | 7.07E-07 |
| ENSG00000100124 | ANKRD54 | 0.43 | 0.08 | 1.74E-07 | 7.92E-07 |
| ENSG00000171044 | XKR6 | 0.43 | 0.09 | 4.69E-07 | 2.06E-06 |
| ENSG00000107864 | CPEB3 | 0.43 | 0.09 | 8.30E-07 | 3.55E-06 |
| ENSG00000126453 | BCL2L12 | 0.43 | 0.1 | 4.88E-06 | 1.95E-05 |
| ENSG00000161647 | MPP3 | 0.43 | 0.1 | 6.84E-06 | 2.69E-05 |
| ENSG00000174109 | UQCC4 | 0.43 | 0.1 | 1.08E-05 | 4.15E-05 |
| ENSG00000176101 | SSNA1 | 0.43 | 0.1 | 1.20E-05 | 4.61E-05 |
| ENSG00000145217 | SLC26A1 | 0.43 | 0.11 | 2.41E-05 | 8.96E-05 |
| ENSG00000167543 | TP53I13 | 0.43 | 0.11 | 3.29E-05 | 1.20E-04 |
| ENSG00000102977 | ACD | 0.43 | 0.12 | 1.13E-04 | 3.91E-04 |
| ENSG00000162959 | MEMO1 | 0.43 | 0.12 | 1.66E-04 | 5.62E-04 |
| ENSG00000197978 | GOLGA6L9 | 0.43 | 0.12 | 1.88E-04 | 6.31E-04 |
| ENSG00000286051 | MAGEA6-DT | 0.43 | 0.13 | 3.94E-04 | 1.27E-03 |
| ENSG00000130748 | TMEM160 | 0.43 | 0.13 | 5.55E-04 | 1.76E-03 |
| ENSG00000227372 | TP73-AS1 | 0.43 | 0.14 | 8.83E-04 | 2.71E-03 |
| ENSG00000021300 | PLEKHB1 | 0.43 | 0.14 | 1.04E-03 | 3.16E-03 |

| | | | | | |
|---|---|---|---|---|---|
| ENSG00000228192 | ZNF691-DT | 0.43 | 0.16 | 2.72E-03 | 7.70E-03 |
| ENSG00000188613 | NANOS1 | 0.43 | 0.16 | 3.65E-03 | 1.01E-02 |
| ENSG00000250397 | | 0.43 | 0.17 | 4.97E-03 | 1.34E-02 |
| ENSG00000267216 | ZNF8-ERVK3-1 | 0.43 | 0.17 | 5.31E-03 | 1.42E-02 |
| ENSG00000243477 | NAA80 | 0.43 | 0.17 | 6.03E-03 | 1.60E-02 |
| ENSG00000278594 | | 0.43 | 0.19 | 1.06E-02 | 2.65E-02 |
| ENSG00000138650 | PCDH10 | 0.43 | 0.19 | 1.26E-02 | 3.11E-02 |
| ENSG00000291154 | PMS2P4 | 0.43 | 0.2 | 1.44E-02 | 3.50E-02 |
| ENSG00000163794 | UCN | 0.43 | 0.2 | 1.50E-02 | 3.64E-02 |
| ENSG00000204256 | BRD2 | 0.42 | 0.04 | 1.47E-29 | 1.94E-28 |
| ENSG00000116478 | HDAC1 | 0.42 | 0.04 | 6.40E-27 | 7.74E-26 |
| ENSG00000162923 | WDR26 | 0.42 | 0.04 | 6.51E-27 | 7.87E-26 |
| ENSG00000130939 | UBE4B | 0.42 | 0.04 | 7.98E-27 | 9.62E-26 |
| ENSG00000113300 | CNOT6 | 0.42 | 0.04 | 1.81E-23 | 1.93E-22 |
| ENSG00000104687 | GSR | 0.42 | 0.04 | 3.33E-23 | 3.50E-22 |
| ENSG00000164543 | STK17A | 0.42 | 0.04 | 2.82E-22 | 2.87E-21 |
| ENSG00000145817 | YIPF5 | 0.42 | 0.05 | 4.18E-21 | 4.08E-20 |
| ENSG00000155011 | DKK2 | 0.42 | 0.05 | 1.09E-19 | 1.00E-18 |
| ENSG00000173011 | TADA2B | 0.42 | 0.05 | 5.16E-18 | 4.38E-17 |
| ENSG00000128595 | CALU | 0.42 | 0.05 | 9.72E-18 | 8.17E-17 |
| ENSG00000038219 | BOD1L1 | 0.42 | 0.05 | 6.04E-17 | 4.92E-16 |
| ENSG00000090054 | SPTLC1 | 0.42 | 0.05 | 5.24E-16 | 4.08E-15 |
| ENSG00000182952 | HMGN4 | 0.42 | 0.05 | 1.09E-15 | 8.30E-15 |
| ENSG00000114354 | TFG | 0.42 | 0.05 | 1.46E-15 | 1.11E-14 |
| ENSG00000101152 | DNAJC5 | 0.42 | 0.06 | 1.14E-14 | 8.34E-14 |
| ENSG00000135297 | MTO1 | 0.42 | 0.05 | 1.72E-14 | 1.24E-13 |
| ENSG00000063587 | ZNF275 | 0.42 | 0.06 | 4.24E-14 | 3.00E-13 |
| ENSG00000151748 | SAV1 | 0.42 | 0.06 | 5.18E-13 | 3.46E-12 |
| ENSG00000076554 | TPD52 | 0.42 | 0.06 | 8.70E-13 | 5.73E-12 |
| ENSG00000178104 | PDE4DIP | 0.42 | 0.06 | 8.72E-13 | 5.74E-12 |
| ENSG00000144674 | GOLGA4 | 0.42 | 0.06 | 9.86E-12 | 6.09E-11 |
| ENSG00000149179 | CSTPP1 | 0.42 | 0.06 | 2.62E-11 | 1.58E-10 |
| ENSG00000042753 | AP2S1 | 0.42 | 0.06 | 3.16E-11 | 1.89E-10 |
| ENSG00000188211 | NCR3LG1 | 0.42 | 0.07 | 5.62E-11 | 3.31E-10 |
| ENSG00000155366 | RHOC | 0.42 | 0.07 | 1.30E-10 | 7.46E-10 |
| ENSG00000082258 | CCNT2 | 0.42 | 0.07 | 1.35E-10 | 7.74E-10 |
| ENSG00000063177 | RPL18 | 0.42 | 0.07 | 2.14E-10 | 1.21E-09 |
| ENSG00000105723 | GSK3A | 0.42 | 0.07 | 2.22E-10 | 1.26E-09 |
| ENSG00000134058 | CDK7 | 0.42 | 0.07 | 2.96E-10 | 1.66E-09 |
| ENSG00000167770 | OTUB1 | 0.42 | 0.07 | 3.98E-10 | 2.21E-09 |
| ENSG00000196588 | MRTFA | 0.42 | 0.07 | 4.41E-10 | 2.44E-09 |
| ENSG00000167380 | ZNF226 | 0.42 | 0.07 | 4.89E-10 | 2.70E-09 |
| ENSG00000198408 | OGA | 0.42 | 0.07 | 5.76E-10 | 3.17E-09 |
| ENSG00000100139 | MICALL1 | 0.42 | 0.07 | 1.97E-09 | 1.04E-08 |
| ENSG00000117569 | PTBP2 | 0.42 | 0.07 | 2.07E-09 | 1.09E-08 |
| ENSG00000177888 | ZBTB41 | 0.42 | 0.07 | 2.35E-09 | 1.23E-08 |
| ENSG00000144426 | NBEAL1 | 0.42 | 0.07 | 3.65E-09 | 1.89E-08 |
| ENSG00000129518 | EAPP | 0.42 | 0.08 | 1.62E-08 | 8.00E-08 |
| ENSG00000088256 | GNA11 | 0.42 | 0.08 | 2.52E-08 | 1.22E-07 |
| ENSG00000087301 | TXNDC16 | 0.42 | 0.08 | 4.29E-08 | 2.04E-07 |

| | | | | | |
|---|---|---|---|---|---|
| ENSG00000231345 | BEND3P1 | 0.42 | 0.08 | 4.42E-08 | 2.10E-07 |
| ENSG00000161921 | CXCL16 | 0.42 | 0.08 | 6.12E-08 | 2.89E-07 |
| ENSG00000147854 | UHRF2 | 0.42 | 0.09 | 4.38E-07 | 1.93E-06 |
| ENSG00000099949 | LZTR1 | 0.42 | 0.09 | 8.72E-07 | 3.72E-06 |
| ENSG00000149089 | APIP | 0.42 | 0.09 | 2.11E-06 | 8.68E-06 |
| ENSG00000124721 | DNAH8 | 0.42 | 0.09 | 3.64E-06 | 1.47E-05 |
| ENSG00000085644 | ZNF213 | 0.42 | 0.11 | 2.71E-05 | 1.00E-04 |
| ENSG00000198355 | PIM3 | 0.42 | 0.11 | 8.53E-05 | 2.99E-04 |
| ENSG00000142396 | ERVK3-1 | 0.42 | 0.11 | 9.12E-05 | 3.19E-04 |
| ENSG00000236453 | | 0.42 | 0.11 | 9.28E-05 | 3.24E-04 |
| ENSG00000159885 | ZNF222 | 0.42 | 0.13 | 6.45E-04 | 2.02E-03 |
| ENSG00000243660 | ZNF487 | 0.42 | 0.13 | 7.47E-04 | 2.31E-03 |
| ENSG00000205336 | ADGRG1 | 0.42 | 0.13 | 7.91E-04 | 2.44E-03 |
| ENSG00000247627 | MTND4P12 | 0.42 | 0.14 | 1.21E-03 | 3.64E-03 |
| ENSG00000272933 | TRIM8-DT | 0.42 | 0.15 | 1.68E-03 | 4.94E-03 |
| ENSG00000240476 | LINC00973 | 0.42 | 0.15 | 3.44E-03 | 9.56E-03 |
| ENSG00000266885 | | 0.42 | 0.16 | 4.10E-03 | 1.12E-02 |
| ENSG00000269896 | | 0.42 | 0.17 | 6.13E-03 | 1.62E-02 |
| ENSG00000111886 | GABRR2 | 0.42 | 0.18 | 8.66E-03 | 2.22E-02 |
| ENSG00000287110 | ACAD9-DT | 0.42 | 0.18 | 9.88E-03 | 2.50E-02 |
| ENSG00000130054 | NALF2 | 0.42 | 0.19 | 1.24E-02 | 3.05E-02 |
| ENSG00000160460 | SPTBN4 | 0.42 | 0.19 | 1.51E-02 | 3.67E-02 |
| ENSG00000196724 | ZNF418 | 0.42 | 0.2 | 1.56E-02 | 3.76E-02 |
| ENSG00000272568 | CPVL-AS2 | 0.42 | 0.21 | 2.03E-02 | 4.77E-02 |
| ENSG00000143612 | C1orf43 | 0.41 | 0.03 | 2.30E-34 | 3.52E-33 |
| ENSG00000107651 | SEC23IP | 0.41 | 0.04 | 3.02E-31 | 4.24E-30 |
| ENSG00000115762 | PLEKHB2 | 0.41 | 0.04 | 4.87E-30 | 6.55E-29 |
| ENSG00000111371 | SLC38A1 | 0.41 | 0.04 | 4.12E-29 | 5.38E-28 |
| ENSG00000149485 | FADS1 | 0.41 | 0.04 | 3.28E-27 | 4.01E-26 |
| ENSG00000104368 | PLAT | 0.41 | 0.04 | 2.67E-26 | 3.16E-25 |
| ENSG00000185896 | LAMP1 | 0.41 | 0.04 | 8.43E-23 | 8.75E-22 |
| ENSG00000131143 | COX4I1 | 0.41 | 0.04 | 1.80E-21 | 1.78E-20 |
| ENSG00000090905 | TNRC6A | 0.41 | 0.04 | 2.07E-21 | 2.04E-20 |
| ENSG00000068878 | PSME4 | 0.41 | 0.05 | 4.16E-19 | 3.71E-18 |
| ENSG00000131051 | RBM39 | 0.41 | 0.05 | 6.41E-19 | 5.66E-18 |
| ENSG00000165819 | METTL3 | 0.41 | 0.05 | 3.19E-18 | 2.73E-17 |
| ENSG00000198169 | ZNF251 | 0.41 | 0.05 | 2.85E-17 | 2.36E-16 |
| ENSG00000141012 | GALNS | 0.41 | 0.05 | 2.77E-16 | 2.18E-15 |
| ENSG00000138138 | ATAD1 | 0.41 | 0.05 | 3.00E-16 | 2.36E-15 |
| ENSG00000159377 | PSMB4 | 0.41 | 0.05 | 4.31E-15 | 3.21E-14 |
| ENSG00000157191 | NECAP2 | 0.41 | 0.05 | 7.40E-15 | 5.46E-14 |
| ENSG00000186501 | TMEM222 | 0.41 | 0.05 | 4.98E-14 | 3.52E-13 |
| ENSG00000165417 | GTF2A1 | 0.41 | 0.06 | 9.88E-14 | 6.88E-13 |
| ENSG00000020256 | ZFP64 | 0.41 | 0.06 | 1.00E-13 | 6.99E-13 |
| ENSG00000152223 | EPG5 | 0.41 | 0.06 | 2.94E-13 | 1.99E-12 |
| ENSG00000157077 | ZFYVE9 | 0.41 | 0.06 | 4.14E-13 | 2.77E-12 |
| ENSG00000224531 | SMIM13 | 0.41 | 0.06 | 8.75E-13 | 5.75E-12 |
| ENSG00000181467 | RAP2B | 0.41 | 0.06 | 2.43E-12 | 1.56E-11 |
| ENSG00000107249 | GLIS3 | 0.41 | 0.06 | 2.96E-12 | 1.89E-11 |
| ENSG00000046651 | OFD1 | 0.41 | 0.06 | 1.33E-11 | 8.17E-11 |

| | | | | | |
|---|---|---|---|---|---|
| ENSG00000110911 | SLC11A2 | 0.41 | 0.06 | 2.09E-11 | 1.27E-10 |
| ENSG00000166750 | SLFN5 | 0.41 | 0.06 | 3.11E-11 | 1.86E-10 |
| ENSG00000102882 | MAPK3 | 0.41 | 0.06 | 4.79E-11 | 2.83E-10 |
| ENSG00000166275 | BORCS7 | 0.41 | 0.07 | 2.07E-10 | 1.18E-09 |
| ENSG00000176102 | CSTF3 | 0.41 | 0.07 | 3.66E-10 | 2.04E-09 |
| ENSG00000100744 | GSKIP | 0.41 | 0.07 | 5.49E-10 | 3.02E-09 |
| ENSG00000040487 | SLC66A1 | 0.41 | 0.07 | 1.66E-09 | 8.80E-09 |
| ENSG00000087152 | ATXN7L3 | 0.41 | 0.07 | 7.41E-09 | 3.74E-08 |
| ENSG00000172007 | RAB33B | 0.41 | 0.07 | 2.49E-08 | 1.21E-07 |
| ENSG00000287263 | | 0.41 | 0.08 | 3.56E-08 | 1.71E-07 |
| ENSG00000163935 | SFMBT1 | 0.41 | 0.08 | 7.14E-08 | 3.35E-07 |
| ENSG00000157107 | FCHO2 | 0.41 | 0.08 | 1.34E-07 | 6.17E-07 |
| ENSG00000065268 | WDR18 | 0.41 | 0.08 | 1.37E-07 | 6.27E-07 |
| ENSG00000131174 | COX7B | 0.41 | 0.08 | 2.04E-07 | 9.25E-07 |
| ENSG00000162585 | FAAP20 | 0.41 | 0.08 | 3.95E-07 | 1.75E-06 |
| ENSG00000173327 | MAP3K11 | 0.41 | 0.08 | 4.28E-07 | 1.89E-06 |
| ENSG00000185049 | NELFA | 0.41 | 0.08 | 4.85E-07 | 2.13E-06 |
| ENSG00000163378 | EOGT | 0.41 | 0.08 | 7.63E-07 | 3.28E-06 |
| ENSG00000167674 | HDGFL2 | 0.41 | 0.09 | 1.05E-06 | 4.43E-06 |
| ENSG00000104859 | CLASRP | 0.41 | 0.09 | 1.64E-06 | 6.82E-06 |
| ENSG00000173320 | STOX2 | 0.41 | 0.09 | 2.90E-06 | 1.18E-05 |
| ENSG00000196476 | C20orf96 | 0.41 | 0.09 | 4.84E-06 | 1.93E-05 |
| ENSG00000068354 | TBC1D25 | 0.41 | 0.09 | 8.75E-06 | 3.40E-05 |
| ENSG00000114923 | SLC4A3 | 0.41 | 0.1 | 1.39E-05 | 5.30E-05 |
| ENSG00000141542 | RAB40B | 0.41 | 0.1 | 3.41E-05 | 1.25E-04 |
| ENSG00000149328 | GLB1L2 | 0.41 | 0.11 | 8.12E-05 | 2.85E-04 |
| ENSG00000182154 | MRPL41 | 0.41 | 0.11 | 1.52E-04 | 5.18E-04 |
| ENSG00000037757 | MRI1 | 0.41 | 0.12 | 5.53E-04 | 1.75E-03 |
| ENSG00000184381 | PLA2G6 | 0.41 | 0.13 | 6.72E-04 | 2.10E-03 |
| ENSG00000168038 | ULK4 | 0.41 | 0.13 | 7.44E-04 | 2.31E-03 |
| ENSG00000174332 | GLIS1 | 0.41 | 0.13 | 1.11E-03 | 3.35E-03 |
| ENSG00000131188 | PRR7 | 0.41 | 0.14 | 1.56E-03 | 4.59E-03 |
| ENSG00000183426 | NPIPA1 | 0.41 | 0.14 | 1.63E-03 | 4.78E-03 |
| ENSG00000182700 | IGIP | 0.41 | 0.14 | 2.36E-03 | 6.75E-03 |
| ENSG00000291228 | | 0.41 | 0.16 | 4.07E-03 | 1.11E-02 |
| ENSG00000272106 | | 0.41 | 0.16 | 4.26E-03 | 1.16E-02 |
| ENSG00000237973 | MTCO1P12 | 0.41 | 0.17 | 8.39E-03 | 2.16E-02 |
| ENSG00000071994 | PDCD2 | 0.4 | 0.03 | 2.81E-30 | 3.81E-29 |
| ENSG00000142192 | APP | 0.4 | 0.04 | 1.06E-28 | 1.36E-27 |
| ENSG00000159592 | GPBP1L1 | 0.4 | 0.04 | 4.26E-24 | 4.65E-23 |
| ENSG00000163430 | FSTL1 | 0.4 | 0.04 | 4.77E-23 | 5.01E-22 |
| ENSG00000154734 | ADAMTS1 | 0.4 | 0.04 | 7.48E-22 | 7.49E-21 |
| ENSG00000253352 | TUG1 | 0.4 | 0.04 | 1.65E-21 | 1.63E-20 |
| ENSG00000134287 | ARF3 | 0.4 | 0.04 | 7.87E-21 | 7.57E-20 |
| ENSG00000131016 | AKAP12 | 0.4 | 0.04 | 5.07E-20 | 4.72E-19 |
| ENSG00000242372 | EIF6 | 0.4 | 0.04 | 9.22E-20 | 8.49E-19 |
| ENSG00000074054 | CLASP1 | 0.4 | 0.04 | 3.01E-19 | 2.71E-18 |
| ENSG00000176407 | KCMF1 | 0.4 | 0.05 | 9.50E-19 | 8.33E-18 |
| ENSG00000151135 | TMEM263 | 0.4 | 0.05 | 2.58E-18 | 2.22E-17 |
| ENSG00000108433 | GOSR2 | 0.4 | 0.05 | 3.26E-17 | 2.69E-16 |

| | | | | | |
|---|---|---|---|---|---|
| ENSG00000119523 | ALG2 | 0.4 | 0.05 | 3.36E-16 | 2.64E-15 |
| ENSG00000143420 | ENSA | 0.4 | 0.05 | 5.76E-16 | 4.47E-15 |
| ENSG00000134884 | ARGLU1 | 0.4 | 0.05 | 9.02E-16 | 6.91E-15 |
| ENSG00000148634 | HERC4 | 0.4 | 0.05 | 1.10E-15 | 8.39E-15 |
| ENSG00000178988 | MRFAP1L1 | 0.4 | 0.05 | 1.37E-15 | 1.04E-14 |
| ENSG00000151498 | ACAD8 | 0.4 | 0.05 | 3.99E-15 | 2.98E-14 |
| ENSG00000120265 | PCMT1 | 0.4 | 0.05 | 2.55E-14 | 1.83E-13 |
| ENSG00000071794 | HLTF | 0.4 | 0.05 | 5.43E-14 | 3.83E-13 |
| ENSG00000115268 | RPS15 | 0.4 | 0.05 | 7.82E-14 | 5.47E-13 |
| ENSG00000172262 | ZNF131 | 0.4 | 0.06 | 1.74E-13 | 1.20E-12 |
| ENSG00000167723 | TRPV3 | 0.4 | 0.06 | 2.77E-13 | 1.88E-12 |
| ENSG00000181904 | C5orf24 | 0.4 | 0.06 | 7.97E-13 | 5.26E-12 |
| ENSG00000162769 | FLVCR1 | 0.4 | 0.06 | 2.50E-12 | 1.60E-11 |
| ENSG00000162601 | MYSM1 | 0.4 | 0.06 | 3.02E-12 | 1.93E-11 |
| ENSG00000197928 | ZNF677 | 0.4 | 0.06 | 7.83E-12 | 4.86E-11 |
| ENSG00000144597 | EAF1 | 0.4 | 0.06 | 3.36E-11 | 2.01E-10 |
| ENSG00000175581 | MRPL48 | 0.4 | 0.06 | 4.61E-11 | 2.73E-10 |
| ENSG00000134759 | ELP2 | 0.4 | 0.06 | 1.20E-10 | 6.90E-10 |
| ENSG00000183077 | AFMID | 0.4 | 0.06 | 1.42E-10 | 8.14E-10 |
| ENSG00000172273 | HINFP | 0.4 | 0.07 | 4.35E-10 | 2.41E-09 |
| ENSG00000138031 | ADCY3 | 0.4 | 0.07 | 8.28E-10 | 4.49E-09 |
| ENSG00000163156 | SCNM1 | 0.4 | 0.07 | 1.23E-09 | 6.61E-09 |
| ENSG00000196954 | CASP4 | 0.4 | 0.07 | 1.43E-09 | 7.63E-09 |
| ENSG00000187240 | DYNC2H1 | 0.4 | 0.07 | 1.97E-09 | 1.04E-08 |
| ENSG00000111224 | PARP11 | 0.4 | 0.07 | 3.47E-09 | 1.80E-08 |
| ENSG00000066855 | MTFR1 | 0.4 | 0.07 | 1.05E-08 | 5.26E-08 |
| ENSG00000131931 | THAP1 | 0.4 | 0.07 | 1.76E-08 | 8.66E-08 |
| ENSG00000235531 | MSC-AS1 | 0.4 | 0.07 | 3.47E-08 | 1.67E-07 |
| ENSG00000120616 | EPC1 | 0.4 | 0.08 | 8.24E-08 | 3.85E-07 |
| ENSG00000163155 | LYSMD1 | 0.4 | 0.08 | 1.25E-07 | 5.74E-07 |
| ENSG00000168890 | TMEM150A | 0.4 | 0.08 | 1.99E-07 | 9.03E-07 |
| ENSG00000185019 | UBOX5 | 0.4 | 0.08 | 2.95E-07 | 1.32E-06 |
| ENSG00000109062 | NHERF1 | 0.4 | 0.08 | 2.97E-07 | 1.33E-06 |
| ENSG00000290931 | | 0.4 | 0.08 | 7.24E-07 | 3.12E-06 |
| ENSG00000167470 | MIDN | 0.4 | 0.08 | 1.01E-06 | 4.27E-06 |
| ENSG00000182362 | YBEY | 0.4 | 0.09 | 1.75E-06 | 7.27E-06 |
| ENSG00000085872 | CHERP | 0.4 | 0.09 | 3.48E-06 | 1.40E-05 |
| ENSG00000132432 | SEC61G | 0.4 | 0.09 | 7.48E-06 | 2.93E-05 |
| ENSG00000100211 | CBY1 | 0.4 | 0.09 | 9.60E-06 | 3.72E-05 |
| ENSG00000188385 | JAKMIP3 | 0.4 | 0.1 | 1.36E-05 | 5.20E-05 |
| ENSG00000273230 | | 0.4 | 0.1 | 1.88E-05 | 7.09E-05 |
| ENSG00000176049 | JAKMIP2 | 0.4 | 0.1 | 2.33E-05 | 8.67E-05 |
| ENSG00000197982 | C1orf122 | 0.4 | 0.1 | 6.78E-05 | 2.41E-04 |
| ENSG00000213185 | FAM24B | 0.4 | 0.11 | 8.41E-05 | 2.95E-04 |
| ENSG00000107077 | KDM4C | 0.4 | 0.11 | 1.07E-04 | 3.72E-04 |
| ENSG00000213563 | C8orf82 | 0.4 | 0.11 | 1.34E-04 | 4.58E-04 |
| ENSG00000176165 | FOXG1 | 0.4 | 0.11 | 1.79E-04 | 6.05E-04 |
| ENSG00000158856 | DMTN | 0.4 | 0.12 | 3.27E-04 | 1.06E-03 |
| ENSG00000232593 | KANTR | 0.4 | 0.12 | 4.15E-04 | 1.33E-03 |
| ENSG00000226328 | NUP50-DT | 0.4 | 0.14 | 2.57E-03 | 7.31E-03 |

| | | | | | |
|---|---|---|---|---|---|
| ENSG00000162576 | MXRA8 | 0.4 | 0.14 | 2.68E-03 | 7.60E-03 |
| ENSG00000239922 | | 0.4 | 0.14 | 3.03E-03 | 8.51E-03 |
| ENSG00000214021 | TTLL3 | 0.4 | 0.17 | 9.18E-03 | 2.34E-02 |
| ENSG00000159588 | CCDC17 | 0.4 | 0.17 | 1.01E-02 | 2.55E-02 |
| ENSG00000257315 | ZBED6 | 0.4 | 0.18 | 1.35E-02 | 3.31E-02 |
| ENSG00000230383 | RPL6P19 | 0.4 | 0.18 | 1.36E-02 | 3.34E-02 |
| ENSG00000251664 | PCDHA12 | 0.4 | 0.18 | 1.51E-02 | 3.65E-02 |
| ENSG00000245848 | CEBPA | 0.4 | 0.19 | 1.63E-02 | 3.91E-02 |
| ENSG00000290976 | | 0.4 | 0.19 | 2.00E-02 | 4.71E-02 |
| ENSG00000228878 | SEPTIN7-DT | 0.4 | 0.2 | 2.11E-02 | 4.95E-02 |
| ENSG00000084733 | RAB10 | 0.39 | 0.04 | 1.54E-26 | 1.84E-25 |
| ENSG00000151233 | GXYLT1 | 0.39 | 0.04 | 4.35E-23 | 4.57E-22 |
| ENSG00000118680 | MYL12B | 0.39 | 0.04 | 2.50E-22 | 2.55E-21 |
| ENSG00000113141 | IK | 0.39 | 0.04 | 3.60E-19 | 3.23E-18 |
| ENSG00000179010 | MRFAP1 | 0.39 | 0.04 | 1.30E-18 | 1.13E-17 |
| ENSG00000136854 | STXBP1 | 0.39 | 0.04 | 1.46E-18 | 1.27E-17 |
| ENSG00000188938 | FAM120AOS | 0.39 | 0.05 | 8.33E-18 | 7.02E-17 |
| ENSG00000175203 | DCTN2 | 0.39 | 0.05 | 9.85E-18 | 8.28E-17 |
| ENSG00000227097 | RPS28P7 | 0.39 | 0.05 | 1.13E-17 | 9.48E-17 |
| ENSG00000159873 | CCDC117 | 0.39 | 0.05 | 1.06E-15 | 8.06E-15 |
| ENSG00000170365 | SMAD1 | 0.39 | 0.05 | 6.99E-15 | 5.16E-14 |
| ENSG00000117036 | ETV3 | 0.39 | 0.05 | 7.72E-15 | 5.68E-14 |
| ENSG00000037749 | MFAP3 | 0.39 | 0.05 | 1.35E-14 | 9.84E-14 |
| ENSG00000159921 | GNE | 0.39 | 0.05 | 1.49E-13 | 1.03E-12 |
| ENSG00000100364 | KIAA0930 | 0.39 | 0.05 | 2.27E-13 | 1.55E-12 |
| ENSG00000131558 | EXOC4 | 0.39 | 0.06 | 5.43E-13 | 3.62E-12 |
| ENSG00000107672 | NSMCE4A | 0.39 | 0.06 | 6.39E-13 | 4.24E-12 |
| ENSG00000198315 | ZKSCAN8 | 0.39 | 0.06 | 4.29E-12 | 2.71E-11 |
| ENSG00000023041 | ZDHHC6 | 0.39 | 0.06 | 6.26E-12 | 3.91E-11 |
| ENSG00000147123 | NDUFB11 | 0.39 | 0.06 | 1.06E-11 | 6.52E-11 |
| ENSG00000090020 | SLC9A1 | 0.39 | 0.06 | 1.29E-11 | 7.94E-11 |
| ENSG00000166681 | BEX3 | 0.39 | 0.06 | 3.87E-11 | 2.30E-10 |
| ENSG00000126012 | KDM5C | 0.39 | 0.06 | 4.69E-11 | 2.77E-10 |
| ENSG00000230124 | ACBD6 | 0.39 | 0.06 | 5.60E-11 | 3.30E-10 |
| ENSG00000198198 | SZT2 | 0.39 | 0.06 | 7.78E-11 | 4.54E-10 |
| ENSG00000131023 | LATS1 | 0.39 | 0.06 | 8.52E-11 | 4.96E-10 |
| ENSG00000101439 | CST3 | 0.39 | 0.06 | 1.47E-10 | 8.46E-10 |
| ENSG00000003402 | CFLAR | 0.39 | 0.06 | 4.00E-10 | 2.22E-09 |
| ENSG00000160691 | SHC1 | 0.39 | 0.06 | 7.41E-10 | 4.04E-09 |
| ENSG00000115446 | UNC50 | 0.39 | 0.06 | 8.11E-10 | 4.40E-09 |
| ENSG00000141756 | FKBP10 | 0.39 | 0.06 | 8.20E-10 | 4.45E-09 |
| ENSG00000140326 | CDAN1 | 0.39 | 0.07 | 3.30E-09 | 1.71E-08 |
| ENSG00000101473 | ACOT8 | 0.39 | 0.07 | 4.05E-09 | 2.08E-08 |
| ENSG00000183935 | HTR7P1 | 0.39 | 0.07 | 4.40E-09 | 2.26E-08 |
| ENSG00000198858 | R3HDM4 | 0.39 | 0.07 | 4.55E-09 | 2.33E-08 |
| ENSG00000127663 | KDM4B | 0.39 | 0.07 | 9.55E-09 | 4.79E-08 |
| ENSG00000102897 | LYRM1 | 0.39 | 0.07 | 1.22E-08 | 6.06E-08 |
| ENSG00000136152 | COG3 | 0.39 | 0.07 | 1.62E-08 | 7.99E-08 |
| ENSG00000122970 | IFT81 | 0.39 | 0.07 | 1.81E-08 | 8.90E-08 |
| ENSG00000085433 | WDR47 | 0.39 | 0.07 | 2.04E-08 | 9.98E-08 |

| | | | | | |
|---|---|---|---|---|---|
| ENSG00000139998 | RAB15 | 0.39 | 0.08 | 1.50E-07 | 6.85E-07 |
| ENSG00000104852 | SNRNP70 | 0.39 | 0.08 | 3.25E-07 | 1.45E-06 |
| ENSG00000184481 | FOXO4 | 0.39 | 0.08 | 3.86E-07 | 1.71E-06 |
| ENSG00000100241 | SBF1 | 0.39 | 0.08 | 4.35E-07 | 1.92E-06 |
| ENSG00000182912 | TSPEAR-AS2 | 0.39 | 0.08 | 5.45E-07 | 2.38E-06 |
| ENSG00000105655 | ISYNA1 | 0.39 | 0.08 | 6.07E-07 | 2.64E-06 |
| ENSG00000148343 | MIGA2 | 0.39 | 0.09 | 2.51E-06 | 1.03E-05 |
| ENSG00000070444 | MNT | 0.39 | 0.09 | 4.41E-06 | 1.76E-05 |
| ENSG00000165714 | BORCS5 | 0.39 | 0.09 | 8.93E-06 | 3.47E-05 |
| ENSG00000000457 | SCYL3 | 0.39 | 0.09 | 1.68E-05 | 6.35E-05 |
| ENSG00000143951 | WDPCP | 0.39 | 0.1 | 2.35E-05 | 8.74E-05 |
| ENSG00000187775 | DNAH17 | 0.39 | 0.1 | 6.99E-05 | 2.48E-04 |
| ENSG00000184281 | TSSC4 | 0.39 | 0.1 | 7.25E-05 | 2.57E-04 |
| ENSG00000148344 | PTGES | 0.39 | 0.11 | 9.49E-05 | 3.31E-04 |
| ENSG00000215270 | TOMM40P2 | 0.39 | 0.11 | 3.12E-04 | 1.02E-03 |
| ENSG00000159314 | ARHGAP27 | 0.39 | 0.13 | 1.01E-03 | 3.05E-03 |
| ENSG00000237004 | ZNRF2P1 | 0.39 | 0.13 | 1.18E-03 | 3.56E-03 |
| ENSG00000269609 | C10orf95-AS1 | 0.39 | 0.13 | 1.26E-03 | 3.77E-03 |
| ENSG00000125740 | FOSB | 0.39 | 0.13 | 1.59E-03 | 4.68E-03 |
| ENSG00000215274 | GAGE10 | 0.39 | 0.13 | 1.95E-03 | 5.67E-03 |
| ENSG00000273203 | | 0.39 | 0.14 | 2.57E-03 | 7.30E-03 |
| ENSG00000137691 | CFAP300 | 0.39 | 0.14 | 3.30E-03 | 9.19E-03 |
| ENSG00000177283 | FZD8 | 0.39 | 0.14 | 3.32E-03 | 9.24E-03 |
| ENSG00000176058 | TPRN | 0.39 | 0.14 | 3.45E-03 | 9.58E-03 |
| ENSG00000099250 | NRP1 | 0.39 | 0.15 | 4.84E-03 | 1.31E-02 |
| ENSG00000185829 | ARL17A | 0.39 | 0.15 | 5.63E-03 | 1.50E-02 |
| ENSG00000281207 | SLFNL1-AS1 | 0.39 | 0.15 | 6.00E-03 | 1.59E-02 |
| ENSG00000245213 | GALNT7-DT | 0.39 | 0.16 | 7.33E-03 | 1.91E-02 |
| ENSG00000290921 | | 0.39 | 0.16 | 9.13E-03 | 2.33E-02 |
| ENSG00000117407 | ARTN | 0.39 | 0.17 | 9.26E-03 | 2.36E-02 |
| ENSG00000153165 | RGPD3 | 0.39 | 0.17 | 1.32E-02 | 3.25E-02 |
| ENSG00000105982 | RNF32 | 0.39 | 0.17 | 1.38E-02 | 3.38E-02 |
| ENSG00000238058 | CAMSAP1-DT | 0.39 | 0.19 | 1.77E-02 | 4.22E-02 |
| ENSG00000231305 | IFT122P3 | 0.39 | 0.19 | 1.85E-02 | 4.39E-02 |
| ENSG00000159140 | SON | 0.38 | 0.03 | 7.69E-32 | 1.09E-30 |
| ENSG00000025772 | TOMM34 | 0.38 | 0.04 | 7.79E-23 | 8.10E-22 |
| ENSG00000115310 | RTN4 | 0.38 | 0.04 | 1.43E-22 | 1.47E-21 |
| ENSG00000187446 | CHP1 | 0.38 | 0.04 | 7.50E-22 | 7.51E-21 |
| ENSG00000151718 | WWC2 | 0.38 | 0.04 | 9.14E-21 | 8.76E-20 |
| ENSG00000147403 | RPL10 | 0.38 | 0.04 | 2.10E-19 | 1.90E-18 |
| ENSG00000072803 | FBXW11 | 0.38 | 0.04 | 2.11E-18 | 1.82E-17 |
| ENSG00000132963 | POMP | 0.38 | 0.04 | 1.00E-17 | 8.42E-17 |
| ENSG00000135148 | TRAFD1 | 0.38 | 0.05 | 1.62E-16 | 1.29E-15 |
| ENSG00000109332 | UBE2D3 | 0.38 | 0.05 | 1.91E-16 | 1.51E-15 |
| ENSG00000111725 | PRKAB1 | 0.38 | 0.05 | 3.08E-16 | 2.42E-15 |
| ENSG00000131100 | ATP6V1E1 | 0.38 | 0.05 | 6.65E-16 | 5.14E-15 |
| ENSG00000011566 | MAP4K3 | 0.38 | 0.05 | 2.29E-15 | 1.73E-14 |
| ENSG00000136156 | ITM2B | 0.38 | 0.05 | 1.59E-14 | 1.15E-13 |
| ENSG00000176853 | FAM91A1 | 0.38 | 0.05 | 1.66E-14 | 1.20E-13 |
| ENSG00000118873 | RAB3GAP2 | 0.38 | 0.05 | 4.85E-14 | 3.43E-13 |

| | | | | | |
|---|---|---|---|---|---|
| ENSG00000101132 | PFDN4 | 0.38 | 0.05 | 6.61E-14 | 4.64E-13 |
| ENSG00000198356 | GET3 | 0.38 | 0.05 | 1.21E-13 | 8.38E-13 |
| ENSG00000090316 | MAEA | 0.38 | 0.05 | 9.52E-13 | 6.25E-12 |
| ENSG00000170348 | TMED10 | 0.38 | 0.05 | 9.70E-13 | 6.37E-12 |
| ENSG00000181924 | COA4 | 0.38 | 0.05 | 1.14E-12 | 7.41E-12 |
| ENSG00000104904 | OAZ1 | 0.38 | 0.05 | 1.47E-12 | 9.56E-12 |
| ENSG00000132254 | ARFIP2 | 0.38 | 0.05 | 1.64E-12 | 1.06E-11 |
| ENSG00000242588 | | 0.38 | 0.06 | 2.86E-12 | 1.83E-11 |
| ENSG00000157741 | UBN2 | 0.38 | 0.06 | 3.83E-12 | 2.43E-11 |
| ENSG00000116857 | TMEM9 | 0.38 | 0.06 | 1.16E-11 | 7.11E-11 |
| ENSG00000126524 | SBDS | 0.38 | 0.06 | 1.17E-11 | 7.17E-11 |
| ENSG00000110066 | KMT5B | 0.38 | 0.06 | 1.33E-10 | 7.68E-10 |
| ENSG00000173542 | MOB1B | 0.38 | 0.06 | 1.40E-10 | 8.03E-10 |
| ENSG00000082269 | FAM135A | 0.38 | 0.06 | 1.98E-10 | 1.13E-09 |
| ENSG00000147687 | TATDN1 | 0.38 | 0.06 | 2.55E-10 | 1.44E-09 |
| ENSG00000130255 | RPL36 | 0.38 | 0.06 | 3.10E-10 | 1.74E-09 |
| ENSG00000198563 | DDX39B | 0.38 | 0.06 | 1.06E-09 | 5.73E-09 |
| ENSG00000011258 | MBTD1 | 0.38 | 0.06 | 1.36E-09 | 7.26E-09 |
| ENSG00000135472 | FAIM2 | 0.38 | 0.06 | 3.53E-09 | 1.83E-08 |
| ENSG00000132024 | CC2D1A | 0.38 | 0.07 | 1.15E-08 | 5.75E-08 |
| ENSG00000123144 | TRIR | 0.38 | 0.07 | 1.40E-08 | 6.94E-08 |
| ENSG00000124172 | ATP5F1E | 0.38 | 0.07 | 2.36E-08 | 1.15E-07 |
| ENSG00000186111 | PIP5K1C | 0.38 | 0.07 | 3.89E-08 | 1.86E-07 |
| ENSG00000148341 | SH3GLB2 | 0.38 | 0.07 | 2.82E-07 | 1.26E-06 |
| ENSG00000129197 | RPAIN | 0.38 | 0.08 | 2.82E-07 | 1.26E-06 |
| ENSG00000105701 | FKBP8 | 0.38 | 0.08 | 4.96E-07 | 2.17E-06 |
| ENSG00000142546 | NOSIP | 0.38 | 0.08 | 1.23E-06 | 5.19E-06 |
| ENSG00000149131 | SERPING1 | 0.38 | 0.08 | 1.70E-06 | 7.05E-06 |
| ENSG00000163132 | MSX1 | 0.38 | 0.08 | 1.72E-06 | 7.15E-06 |
| ENSG00000131242 | RAB11FIP4 | 0.38 | 0.08 | 2.00E-06 | 8.25E-06 |
| ENSG00000140320 | BAHD1 | 0.38 | 0.08 | 3.41E-06 | 1.38E-05 |
| ENSG00000270885 | RASL10B | 0.38 | 0.08 | 3.47E-06 | 1.40E-05 |
| ENSG00000185800 | DMWD | 0.38 | 0.09 | 5.41E-06 | 2.15E-05 |
| ENSG00000118046 | STK11 | 0.38 | 0.09 | 2.00E-05 | 7.50E-05 |
| ENSG00000116668 | SWT1 | 0.38 | 0.09 | 2.31E-05 | 8.62E-05 |
| ENSG00000162817 | C1orf115 | 0.38 | 0.1 | 4.34E-05 | 1.57E-04 |
| ENSG00000196411 | EPHB4 | 0.38 | 0.1 | 5.22E-05 | 1.87E-04 |
| ENSG00000234912 | SNHG20 | 0.38 | 0.1 | 5.75E-05 | 2.06E-04 |
| ENSG00000174775 | HRAS | 0.38 | 0.1 | 6.82E-05 | 2.42E-04 |
| ENSG00000182208 | MOB2 | 0.38 | 0.11 | 2.54E-04 | 8.41E-04 |
| ENSG00000162729 | IGSF8 | 0.38 | 0.11 | 2.97E-04 | 9.72E-04 |
| ENSG00000024862 | CCDC28A | 0.38 | 0.11 | 3.68E-04 | 1.19E-03 |
| ENSG00000126603 | GLIS2 | 0.38 | 0.11 | 4.44E-04 | 1.42E-03 |
| ENSG00000197497 | ZNF665 | 0.38 | 0.12 | 5.00E-04 | 1.59E-03 |
| ENSG00000122756 | CNTFR | 0.38 | 0.13 | 2.34E-03 | 6.72E-03 |
| ENSG00000182310 | SPACA6 | 0.38 | 0.15 | 5.31E-03 | 1.42E-02 |
| ENSG00000167065 | DUSP18 | 0.38 | 0.15 | 6.59E-03 | 1.73E-02 |
| ENSG00000185499 | MUC1 | 0.38 | 0.16 | 9.64E-03 | 2.44E-02 |
| ENSG00000023734 | STRAP | 0.37 | 0.04 | 7.94E-24 | 8.59E-23 |
| ENSG00000061676 | NCKAP1 | 0.37 | 0.04 | 8.98E-22 | 8.96E-21 |

| | | | | | |
|---|---|---|---|---|---|
| ENSG00000122642 | FKBP9 | 0.37 | 0.04 | 3.38E-20 | 3.17E-19 |
| ENSG00000101474 | APMAP | 0.37 | 0.04 | 4.59E-20 | 4.28E-19 |
| ENSG00000136827 | TOR1A | 0.37 | 0.04 | 2.30E-18 | 1.98E-17 |
| ENSG00000143149 | ALDH9A1 | 0.37 | 0.04 | 2.67E-18 | 2.30E-17 |
| ENSG00000090615 | GOLGA3 | 0.37 | 0.04 | 4.36E-18 | 3.71E-17 |
| ENSG00000102007 | PLP2 | 0.37 | 0.04 | 5.59E-17 | 4.56E-16 |
| ENSG00000092841 | MYL6 | 0.37 | 0.05 | 8.13E-17 | 6.57E-16 |
| ENSG00000132670 | PTPRA | 0.37 | 0.05 | 1.05E-16 | 8.41E-16 |
| ENSG00000163291 | PAQR3 | 0.37 | 0.05 | 1.31E-15 | 9.98E-15 |
| ENSG00000077147 | TM9SF3 | 0.37 | 0.05 | 2.60E-15 | 1.96E-14 |
| ENSG00000101290 | CDS2 | 0.37 | 0.05 | 1.11E-14 | 8.07E-14 |
| ENSG00000164164 | OTUD4 | 0.37 | 0.05 | 1.58E-13 | 1.08E-12 |
| ENSG00000155363 | MOV10 | 0.37 | 0.05 | 8.02E-13 | 5.30E-12 |
| ENSG00000102606 | ARHGEF7 | 0.37 | 0.05 | 1.29E-12 | 8.39E-12 |
| ENSG00000070010 | UFD1 | 0.37 | 0.05 | 2.38E-12 | 1.53E-11 |
| ENSG00000013275 | PSMC4 | 0.37 | 0.05 | 3.36E-12 | 2.14E-11 |
| ENSG00000270231 | NBPF8 | 0.37 | 0.05 | 5.32E-12 | 3.34E-11 |
| ENSG00000148842 | CNNM2 | 0.37 | 0.05 | 5.52E-12 | 3.46E-11 |
| ENSG00000197217 | ENTPD4 | 0.37 | 0.05 | 6.54E-12 | 4.07E-11 |
| ENSG00000112473 | SLC39A7 | 0.37 | 0.05 | 9.39E-12 | 5.81E-11 |
| ENSG00000173276 | ZBTB21 | 0.37 | 0.05 | 1.06E-11 | 6.53E-11 |
| ENSG00000133103 | COG6 | 0.37 | 0.06 | 1.17E-11 | 7.17E-11 |
| ENSG00000173548 | SNX33 | 0.37 | 0.06 | 1.78E-11 | 1.08E-10 |
| ENSG00000105053 | VRK3 | 0.37 | 0.06 | 5.19E-11 | 3.07E-10 |
| ENSG00000072778 | ACADVL | 0.37 | 0.06 | 7.33E-10 | 4.00E-09 |
| ENSG00000111737 | RAB35 | 0.37 | 0.06 | 1.40E-09 | 7.47E-09 |
| ENSG00000119922 | IFIT2 | 0.37 | 0.06 | 1.42E-09 | 7.58E-09 |
| ENSG00000106245 | BUD31 | 0.37 | 0.06 | 2.21E-09 | 1.16E-08 |
| ENSG00000198718 | TOGARAM1 | 0.37 | 0.06 | 2.26E-09 | 1.19E-08 |
| ENSG00000197961 | ZNF121 | 0.37 | 0.06 | 2.52E-09 | 1.32E-08 |
| ENSG00000163605 | PPP4R2 | 0.37 | 0.07 | 4.97E-09 | 2.54E-08 |
| ENSG00000072864 | NDE1 | 0.37 | 0.06 | 6.25E-09 | 3.17E-08 |
| ENSG00000078399 | HOXA9 | 0.37 | 0.06 | 6.34E-09 | 3.22E-08 |
| ENSG00000100916 | BRMS1L | 0.37 | 0.07 | 1.74E-08 | 8.55E-08 |
| ENSG00000151320 | AKAP6 | 0.37 | 0.07 | 3.37E-08 | 1.62E-07 |
| ENSG00000121964 | GTDC1 | 0.37 | 0.07 | 3.39E-08 | 1.63E-07 |
| ENSG00000142186 | SCYL1 | 0.37 | 0.07 | 4.22E-08 | 2.01E-07 |
| ENSG00000147533 | GOLGA7 | 0.37 | 0.07 | 5.15E-08 | 2.44E-07 |
| ENSG00000249859 | PVT1 | 0.37 | 0.07 | 7.36E-08 | 3.45E-07 |
| ENSG00000166797 | CIAO2A | 0.37 | 0.07 | 1.60E-07 | 7.29E-07 |
| ENSG00000175213 | ZNF408 | 0.37 | 0.07 | 3.05E-07 | 1.36E-06 |
| ENSG00000005486 | RHBDD2 | 0.37 | 0.08 | 4.98E-07 | 2.18E-06 |
| ENSG00000148411 | NACC2 | 0.37 | 0.08 | 6.58E-07 | 2.85E-06 |
| ENSG00000115275 | MOGS | 0.37 | 0.08 | 1.43E-06 | 5.97E-06 |
| ENSG00000055732 | MCOLN3 | 0.37 | 0.08 | 1.61E-06 | 6.70E-06 |
| ENSG00000114735 | HEMK1 | 0.37 | 0.08 | 4.22E-06 | 1.69E-05 |
| ENSG00000149231 | CCDC82 | 0.37 | 0.08 | 4.80E-06 | 1.92E-05 |
| ENSG00000269190 | FBXO17 | 0.37 | 0.09 | 9.00E-06 | 3.49E-05 |
| ENSG00000120519 | SLC10A7 | 0.37 | 0.09 | 1.03E-05 | 3.99E-05 |
| ENSG00000123933 | MXD4 | 0.37 | 0.09 | 1.29E-05 | 4.95E-05 |

| | | | | | |
|---|---|---|---|---|---|
| ENSG00000174938 | SEZ6L2 | 0.37 | 0.09 | 1.30E-05 | 4.96E-05 |
| ENSG00000173264 | GPR137 | 0.37 | 0.09 | 1.38E-05 | 5.27E-05 |
| ENSG00000241685 | ARPC1A | 0.37 | 0.09 | 1.41E-05 | 5.38E-05 |
| ENSG00000105325 | FZR1 | 0.37 | 0.09 | 1.48E-05 | 5.64E-05 |
| ENSG00000140854 | KATNB1 | 0.37 | 0.09 | 2.11E-05 | 7.90E-05 |
| ENSG00000184076 | UQCR10 | 0.37 | 0.09 | 2.43E-05 | 9.04E-05 |
| ENSG00000145214 | DGKQ | 0.37 | 0.09 | 3.46E-05 | 1.27E-04 |
| ENSG00000179965 | ZNF771 | 0.37 | 0.09 | 3.70E-05 | 1.35E-04 |
| ENSG00000006015 | REX1BD | 0.37 | 0.09 | 4.29E-05 | 1.55E-04 |
| ENSG00000237172 | B3GNT9 | 0.37 | 0.09 | 4.77E-05 | 1.72E-04 |
| ENSG00000205903 | ZNF316 | 0.37 | 0.09 | 5.35E-05 | 1.92E-04 |
| ENSG00000198205 | ZXDA | 0.37 | 0.1 | 7.14E-05 | 2.53E-04 |
| ENSG00000174672 | BRSK2 | 0.37 | 0.1 | 1.25E-04 | 4.28E-04 |
| ENSG00000196372 | ASB13 | 0.37 | 0.1 | 1.59E-04 | 5.40E-04 |
| ENSG00000204564 | C6orf136 | 0.37 | 0.1 | 1.61E-04 | 5.46E-04 |
| ENSG00000141576 | RNF157 | 0.37 | 0.1 | 2.23E-04 | 7.44E-04 |
| ENSG00000196268 | ZNF493 | 0.37 | 0.1 | 2.24E-04 | 7.48E-04 |
| ENSG00000235552 | RPL6P27 | 0.37 | 0.1 | 2.38E-04 | 7.90E-04 |
| ENSG00000168569 | TMEM223 | 0.37 | 0.1 | 2.62E-04 | 8.66E-04 |
| ENSG00000039650 | PNKP | 0.37 | 0.11 | 3.64E-04 | 1.18E-03 |
| ENSG00000241837 | ATP5PO | 0.37 | 0.12 | 6.70E-04 | 2.10E-03 |
| ENSG00000172031 | EPHX4 | 0.37 | 0.12 | 7.47E-04 | 2.32E-03 |
| ENSG00000291075 | | 0.37 | 0.11 | 7.67E-04 | 2.37E-03 |
| ENSG00000182057 | OGFRP1 | 0.37 | 0.12 | 1.07E-03 | 3.25E-03 |
| ENSG00000218426 | | 0.37 | 0.12 | 1.46E-03 | 4.33E-03 |
| ENSG00000108932 | SLC16A6 | 0.37 | 0.13 | 2.71E-03 | 7.67E-03 |
| ENSG00000121716 | PILRB | 0.37 | 0.13 | 2.91E-03 | 8.20E-03 |
| ENSG00000226084 | RPL17P6 | 0.37 | 0.13 | 3.39E-03 | 9.43E-03 |
| ENSG00000157833 | GAREM2 | 0.37 | 0.14 | 3.57E-03 | 9.87E-03 |
| ENSG00000204381 | LAYN | 0.37 | 0.14 | 5.34E-03 | 1.43E-02 |
| ENSG00000118690 | ARMC2 | 0.37 | 0.15 | 6.73E-03 | 1.77E-02 |
| ENSG00000234797 | RPS3AP6 | 0.37 | 0.16 | 1.15E-02 | 2.87E-02 |
| ENSG00000123154 | WDR83 | 0.37 | 0.16 | 1.22E-02 | 3.02E-02 |
| ENSG00000173040 | EVC2 | 0.37 | 0.17 | 1.78E-02 | 4.25E-02 |
| ENSG00000088812 | ATRN | 0.36 | 0.04 | 5.61E-19 | 4.97E-18 |
| ENSG00000171863 | RPS7 | 0.36 | 0.04 | 3.34E-18 | 2.86E-17 |
| ENSG00000087263 | OGFOD1 | 0.36 | 0.04 | 3.45E-18 | 2.95E-17 |
| ENSG00000141580 | WDR45B | 0.36 | 0.04 | 5.68E-18 | 4.82E-17 |
| ENSG00000169855 | ROBO1 | 0.36 | 0.04 | 5.83E-18 | 4.95E-17 |
| ENSG00000128908 | INO80 | 0.36 | 0.04 | 9.67E-18 | 8.13E-17 |
| ENSG00000103222 | ABCC1 | 0.36 | 0.05 | 5.51E-14 | 3.89E-13 |
| ENSG00000086062 | B4GALT1 | 0.36 | 0.05 | 1.10E-13 | 7.64E-13 |
| ENSG00000244306 | DUXAP10 | 0.36 | 0.05 | 1.27E-13 | 8.81E-13 |
| ENSG00000110925 | CSRNP2 | 0.36 | 0.05 | 1.79E-13 | 1.22E-12 |
| ENSG00000198625 | MDM4 | 0.36 | 0.05 | 2.72E-13 | 1.84E-12 |
| ENSG00000106346 | USP42 | 0.36 | 0.05 | 6.11E-13 | 4.07E-12 |
| ENSG00000177731 | FLII | 0.36 | 0.05 | 4.32E-12 | 2.73E-11 |
| ENSG00000204590 | GNL1 | 0.36 | 0.05 | 5.03E-12 | 3.16E-11 |
| ENSG00000177383 | MAGEF1 | 0.36 | 0.05 | 5.24E-12 | 3.28E-11 |
| ENSG00000173517 | PEAK1 | 0.36 | 0.05 | 2.99E-11 | 1.79E-10 |

| | | | | | |
|---|---|---|---|---|---|
| ENSG00000116406 | EDEM3 | 0.36 | 0.06 | 6.08E-11 | 3.57E-10 |
| ENSG00000104219 | ZDHHC2 | 0.36 | 0.06 | 8.05E-11 | 4.70E-10 |
| ENSG00000182973 | CNOT10 | 0.36 | 0.06 | 1.25E-10 | 7.22E-10 |
| ENSG00000154803 | FLCN | 0.36 | 0.06 | 1.85E-10 | 1.05E-09 |
| ENSG00000155975 | VPS37A | 0.36 | 0.06 | 1.92E-10 | 1.10E-09 |
| ENSG00000153714 | LURAP1L | 0.36 | 0.06 | 3.38E-10 | 1.89E-09 |
| ENSG00000187535 | IFT140 | 0.36 | 0.06 | 5.60E-10 | 3.08E-09 |
| ENSG00000101966 | XIAP | 0.36 | 0.06 | 1.18E-09 | 6.32E-09 |
| ENSG00000116260 | QSOX1 | 0.36 | 0.06 | 1.53E-09 | 8.13E-09 |
| ENSG00000139117 | CPNE8 | 0.36 | 0.06 | 1.87E-09 | 9.90E-09 |
| ENSG00000215712 | TMEM242 | 0.36 | 0.06 | 6.59E-09 | 3.34E-08 |
| ENSG00000149806 | FAU | 0.36 | 0.06 | 8.20E-09 | 4.12E-08 |
| ENSG00000157593 | SLC35B2 | 0.36 | 0.06 | 9.76E-09 | 4.90E-08 |
| ENSG00000142556 | ZNF614 | 0.36 | 0.06 | 1.07E-08 | 5.37E-08 |
| ENSG00000167186 | COQ7 | 0.36 | 0.06 | 1.55E-08 | 7.65E-08 |
| ENSG00000177917 | ARL6IP6 | 0.36 | 0.06 | 2.12E-08 | 1.04E-07 |
| ENSG00000138744 | NAAA | 0.36 | 0.07 | 2.75E-08 | 1.33E-07 |
| ENSG00000198920 | KIAA0753 | 0.36 | 0.07 | 9.11E-08 | 4.24E-07 |
| ENSG00000119705 | SLIRP | 0.36 | 0.07 | 1.25E-07 | 5.77E-07 |
| ENSG00000141349 | G6PC3 | 0.36 | 0.07 | 1.73E-07 | 7.88E-07 |
| ENSG00000174996 | KLC2 | 0.36 | 0.07 | 1.90E-07 | 8.64E-07 |
| ENSG00000116918 | TSNAX | 0.36 | 0.07 | 2.15E-07 | 9.73E-07 |
| ENSG00000125505 | MBOAT7 | 0.36 | 0.07 | 2.93E-07 | 1.31E-06 |
| ENSG00000075426 | FOSL2 | 0.36 | 0.07 | 5.00E-07 | 2.19E-06 |
| ENSG00000107438 | PDLIM1 | 0.36 | 0.07 | 7.14E-07 | 3.08E-06 |
| ENSG00000148331 | ASB6 | 0.36 | 0.07 | 7.67E-07 | 3.29E-06 |
| ENSG00000172809 | RPL38 | 0.36 | 0.07 | 8.59E-07 | 3.67E-06 |
| ENSG00000164258 | NDUFS4 | 0.36 | 0.08 | 1.01E-06 | 4.30E-06 |
| ENSG00000150977 | RILPL2 | 0.36 | 0.08 | 1.42E-06 | 5.96E-06 |
| ENSG00000043093 | DCUN1D1 | 0.36 | 0.08 | 1.49E-06 | 6.24E-06 |
| ENSG00000175324 | LSM1 | 0.36 | 0.08 | 2.50E-06 | 1.02E-05 |
| ENSG00000090674 | MCOLN1 | 0.36 | 0.08 | 2.85E-06 | 1.16E-05 |
| ENSG00000213918 | DNASE1 | 0.36 | 0.08 | 4.42E-06 | 1.77E-05 |
| ENSG00000130811 | EIF3G | 0.36 | 0.08 | 5.10E-06 | 2.03E-05 |
| ENSG00000131725 | WDR44 | 0.36 | 0.08 | 5.50E-06 | 2.18E-05 |
| ENSG00000283633 | | 0.36 | 0.08 | 7.36E-06 | 2.89E-05 |
| ENSG00000132635 | PCED1A | 0.36 | 0.08 | 1.26E-05 | 4.84E-05 |
| ENSG00000175854 | SWI5 | 0.36 | 0.09 | 1.62E-05 | 6.13E-05 |
| ENSG00000196422 | PPP1R26 | 0.36 | 0.09 | 2.50E-05 | 9.27E-05 |
| ENSG00000101624 | CEP76 | 0.36 | 0.09 | 2.53E-05 | 9.37E-05 |
| ENSG00000114270 | COL7A1 | 0.36 | 0.09 | 5.55E-05 | 1.99E-04 |
| ENSG00000144935 | TRPC1 | 0.36 | 0.09 | 6.14E-05 | 2.19E-04 |
| ENSG00000185842 | DNAH14 | 0.36 | 0.09 | 8.27E-05 | 2.90E-04 |
| ENSG00000175087 | PDIK1L | 0.36 | 0.1 | 1.09E-04 | 3.77E-04 |
| ENSG00000171224 | FAM241B | 0.36 | 0.11 | 7.94E-04 | 2.45E-03 |
| ENSG00000186132 | C2orf76 | 0.36 | 0.12 | 2.38E-03 | 6.82E-03 |
| ENSG00000090932 | DLL3 | 0.36 | 0.13 | 2.58E-03 | 7.35E-03 |
| ENSG00000187837 | H1-2 | 0.36 | 0.13 | 3.28E-03 | 9.15E-03 |
| ENSG00000233476 | EEF1A1P6 | 0.36 | 0.13 | 3.55E-03 | 9.81E-03 |
| ENSG00000103326 | CAPN15 | 0.36 | 0.14 | 6.84E-03 | 1.79E-02 |

| | | | | | |
|---|---|---|---|---|---|
| ENSG00000272168 | CASC15 | 0.36 | 0.15 | 8.43E-03 | 2.17E-02 |
| ENSG00000115657 | ABCB6 | 0.36 | 0.16 | 1.33E-02 | 3.26E-02 |
| ENSG00000224152 | BAZ2B-AS1 | 0.36 | 0.16 | 1.61E-02 | 3.88E-02 |
| ENSG00000176731 | RBIS | 0.36 | 0.17 | 1.78E-02 | 4.24E-02 |
| ENSG00000227354 | RBM26-AS1 | 0.36 | 0.17 | 1.98E-02 | 4.68E-02 |
| ENSG00000174437 | ATP2A2 | 0.35 | 0.03 | 1.12E-36 | 1.85E-35 |
| ENSG00000162244 | RPL29 | 0.35 | 0.04 | 5.28E-21 | 5.12E-20 |
| ENSG00000196510 | ANAPC7 | 0.35 | 0.04 | 1.99E-18 | 1.72E-17 |
| ENSG00000106615 | RHEB | 0.35 | 0.04 | 4.51E-18 | 3.84E-17 |
| ENSG00000275066 | SYNRG | 0.35 | 0.04 | 7.82E-18 | 6.60E-17 |
| ENSG00000162368 | CMPK1 | 0.35 | 0.04 | 4.83E-17 | 3.95E-16 |
| ENSG00000131504 | DIAPH1 | 0.35 | 0.04 | 1.24E-16 | 9.93E-16 |
| ENSG00000103429 | BFAR | 0.35 | 0.04 | 1.36E-16 | 1.09E-15 |
| ENSG00000153989 | NUS1 | 0.35 | 0.04 | 1.36E-16 | 1.09E-15 |
| ENSG00000120254 | MTHFD1L | 0.35 | 0.04 | 5.00E-16 | 3.89E-15 |
| ENSG00000136758 | YME1L1 | 0.35 | 0.04 | 7.26E-16 | 5.60E-15 |
| ENSG00000168615 | ADAM9 | 0.35 | 0.04 | 1.12E-15 | 8.53E-15 |
| ENSG00000147684 | NDUFB9 | 0.35 | 0.05 | 1.21E-13 | 8.36E-13 |
| ENSG00000166454 | ATMIN | 0.35 | 0.05 | 8.84E-13 | 5.81E-12 |
| ENSG00000198961 | PJA2 | 0.35 | 0.05 | 5.59E-12 | 3.49E-11 |
| ENSG00000173660 | UQCRH | 0.35 | 0.05 | 1.11E-11 | 6.85E-11 |
| ENSG00000248333 | CDK11B | 0.35 | 0.05 | 1.28E-11 | 7.84E-11 |
| ENSG00000104228 | TRIM35 | 0.35 | 0.06 | 2.21E-10 | 1.26E-09 |
| ENSG00000086065 | CHMP5 | 0.35 | 0.06 | 5.51E-10 | 3.03E-09 |
| ENSG00000177879 | AP3S1 | 0.35 | 0.06 | 6.48E-10 | 3.55E-09 |
| ENSG00000121022 | COPS5 | 0.35 | 0.06 | 1.01E-09 | 5.42E-09 |
| ENSG00000111843 | TMEM14C | 0.35 | 0.06 | 1.11E-09 | 5.99E-09 |
| ENSG00000114446 | IFT57 | 0.35 | 0.06 | 1.22E-09 | 6.53E-09 |
| ENSG00000130517 | PGPEP1 | 0.35 | 0.06 | 1.74E-09 | 9.22E-09 |
| ENSG00000198393 | ZNF26 | 0.35 | 0.06 | 2.42E-09 | 1.27E-08 |
| ENSG00000132405 | TBC1D14 | 0.35 | 0.06 | 3.40E-09 | 1.76E-08 |
| ENSG00000234456 | MAGI2-AS3 | 0.35 | 0.06 | 3.82E-09 | 1.97E-08 |
| ENSG00000169255 | B3GALNT1 | 0.35 | 0.06 | 3.82E-09 | 1.97E-08 |
| ENSG00000187607 | ZNF286A | 0.35 | 0.06 | 4.42E-09 | 2.27E-08 |
| ENSG00000111275 | ALDH2 | 0.35 | 0.07 | 9.42E-08 | 4.38E-07 |
| ENSG00000121653 | MAPK8IP1 | 0.35 | 0.07 | 1.45E-07 | 6.64E-07 |
| ENSG00000119844 | AFTPH | 0.35 | 0.07 | 1.51E-07 | 6.92E-07 |
| ENSG00000054598 | FOXC1 | 0.35 | 0.07 | 5.65E-07 | 2.46E-06 |
| ENSG00000164291 | ARSK | 0.35 | 0.07 | 6.89E-07 | 2.97E-06 |
| ENSG00000176148 | TCP11L1 | 0.35 | 0.07 | 9.16E-07 | 3.90E-06 |
| ENSG00000126432 | PRDX5 | 0.35 | 0.07 | 1.66E-06 | 6.91E-06 |
| ENSG00000170946 | DNAJC24 | 0.35 | 0.08 | 2.06E-06 | 8.48E-06 |
| ENSG00000141504 | SAT2 | 0.35 | 0.08 | 3.51E-06 | 1.42E-05 |
| ENSG00000067057 | PFKP | 0.35 | 0.08 | 4.39E-06 | 1.76E-05 |
| ENSG00000172732 | MUS81 | 0.35 | 0.08 | 5.76E-06 | 2.28E-05 |
| ENSG00000256043 | CTSO | 0.35 | 0.08 | 1.68E-05 | 6.36E-05 |
| ENSG00000131697 | NPHP4 | 0.35 | 0.08 | 2.09E-05 | 7.83E-05 |
| ENSG00000175756 | AURKAIP1 | 0.35 | 0.08 | 2.42E-05 | 9.01E-05 |
| ENSG00000125375 | DMAC2L | 0.35 | 0.09 | 3.03E-05 | 1.12E-04 |
| ENSG00000063854 | HAGH | 0.35 | 0.09 | 5.01E-05 | 1.80E-04 |

| | | | | | |
|---|---|---|---|---|---|
| ENSG00000173113 | TRMT112 | 0.35 | 0.09 | 5.77E-05 | 2.06E-04 |
| ENSG00000198804 | MT-CO1 | 0.35 | 0.09 | 7.94E-05 | 2.80E-04 |
| ENSG00000164074 | ABHD18 | 0.35 | 0.1 | 2.15E-04 | 7.17E-04 |
| ENSG00000160233 | LRRC3 | 0.35 | 0.1 | 4.83E-04 | 1.54E-03 |
| ENSG00000119865 | CNRIP1 | 0.35 | 0.1 | 5.02E-04 | 1.60E-03 |
| ENSG00000204764 | RANBP17 | 0.35 | 0.11 | 7.22E-04 | 2.25E-03 |
| ENSG00000043143 | JADE2 | 0.35 | 0.11 | 8.23E-04 | 2.54E-03 |
| ENSG00000160401 | CFAP157 | 0.35 | 0.11 | 1.11E-03 | 3.35E-03 |
| ENSG00000280670 | CCDC163 | 0.35 | 0.13 | 2.94E-03 | 8.27E-03 |
| ENSG00000161682 | FAM171A2 | 0.35 | 0.13 | 3.42E-03 | 9.49E-03 |
| ENSG00000213888 | LINC01521 | 0.35 | 0.13 | 3.48E-03 | 9.64E-03 |
| ENSG00000038002 | AGA | 0.35 | 0.13 | 5.21E-03 | 1.40E-02 |
| ENSG00000215908 | CROCCP2 | 0.35 | 0.14 | 6.45E-03 | 1.70E-02 |
| ENSG00000291203 | | 0.35 | 0.14 | 7.52E-03 | 1.96E-02 |
| ENSG00000151743 | AMN1 | 0.35 | 0.14 | 9.97E-03 | 2.52E-02 |
| ENSG00000185614 | INKA1 | 0.35 | 0.15 | 1.05E-02 | 2.63E-02 |
| ENSG00000229152 | ANKRD10-IT1 | 0.35 | 0.15 | 1.19E-02 | 2.96E-02 |
| ENSG00000205913 | SRRM2-AS1 | 0.35 | 0.15 | 1.25E-02 | 3.09E-02 |
| ENSG00000105204 | DYRK1B | 0.35 | 0.16 | 1.50E-02 | 3.63E-02 |
| ENSG00000144362 | PHOSPHO2 | 0.35 | 0.16 | 1.59E-02 | 3.84E-02 |
| ENSG00000153485 | LYSET | 0.35 | 0.16 | 1.91E-02 | 4.53E-02 |
| ENSG00000047849 | MAP4 | 0.34 | 0.03 | 3.53E-26 | 4.15E-25 |
| ENSG00000124214 | STAU1 | 0.34 | 0.04 | 7.22E-21 | 6.95E-20 |
| ENSG00000136485 | DCAF7 | 0.34 | 0.04 | 2.40E-19 | 2.17E-18 |
| ENSG00000143870 | PDIA6 | 0.34 | 0.04 | 9.21E-19 | 8.09E-18 |
| ENSG00000088832 | FKBP1A | 0.34 | 0.04 | 3.61E-18 | 3.08E-17 |
| ENSG00000106771 | TMEM245 | 0.34 | 0.04 | 2.41E-17 | 2.00E-16 |
| ENSG00000007168 | PAFAH1B1 | 0.34 | 0.04 | 8.13E-17 | 6.57E-16 |
| ENSG00000123562 | MORF4L2 | 0.34 | 0.04 | 1.09E-16 | 8.78E-16 |
| ENSG00000255248 | MIR100HG | 0.34 | 0.04 | 2.54E-16 | 2.00E-15 |
| ENSG00000144635 | DYNC1LI1 | 0.34 | 0.05 | 4.33E-14 | 3.07E-13 |
| ENSG00000163444 | TMEM183A | 0.34 | 0.05 | 5.97E-14 | 4.20E-13 |
| ENSG00000159111 | MRPL10 | 0.34 | 0.05 | 1.38E-13 | 9.53E-13 |
| ENSG00000132323 | ILKAP | 0.34 | 0.05 | 1.91E-13 | 1.31E-12 |
| ENSG00000166855 | CLPX | 0.34 | 0.05 | 2.65E-13 | 1.80E-12 |
| ENSG00000075151 | EIF4G3 | 0.34 | 0.05 | 3.87E-13 | 2.61E-12 |
| ENSG00000100814 | CCNB1IP1 | 0.34 | 0.05 | 5.75E-13 | 3.83E-12 |
| ENSG00000104763 | ASAH1 | 0.34 | 0.05 | 8.88E-13 | 5.83E-12 |
| ENSG00000146966 | DENND2A | 0.34 | 0.05 | 4.07E-12 | 2.58E-11 |
| ENSG00000163249 | CCNYL1 | 0.34 | 0.05 | 9.29E-12 | 5.74E-11 |
| ENSG00000124767 | GLO1 | 0.34 | 0.05 | 1.74E-10 | 9.91E-10 |
| ENSG00000182247 | UBE2E2 | 0.34 | 0.06 | 1.66E-09 | 8.81E-09 |
| ENSG00000158711 | ELK4 | 0.34 | 0.06 | 2.10E-09 | 1.11E-08 |
| ENSG00000163714 | U2SURP | 0.34 | 0.06 | 2.89E-09 | 1.51E-08 |
| ENSG00000134569 | LRP4 | 0.34 | 0.06 | 5.97E-09 | 3.04E-08 |
| ENSG00000100401 | RANGAP1 | 0.34 | 0.06 | 1.13E-08 | 5.64E-08 |
| ENSG00000100284 | TOM1 | 0.34 | 0.06 | 1.59E-08 | 7.82E-08 |
| ENSG00000141447 | OSBPL1A | 0.34 | 0.06 | 2.24E-08 | 1.10E-07 |
| ENSG00000148798 | INA | 0.34 | 0.06 | 2.40E-08 | 1.17E-07 |
| ENSG00000103510 | KAT8 | 0.34 | 0.06 | 2.47E-08 | 1.20E-07 |

| | | | | | |
|---|---|---|---|---|---|
| ENSG00000109184 | DCUN1D4 | 0.34 | 0.06 | 5.83E-08 | 2.76E-07 |
| ENSG00000174343 | CHRNA9 | 0.34 | 0.06 | 6.68E-08 | 3.14E-07 |
| ENSG00000123179 | EBPL | 0.34 | 0.07 | 1.42E-07 | 6.52E-07 |
| ENSG00000114127 | XRN1 | 0.34 | 0.07 | 2.23E-07 | 1.01E-06 |
| ENSG00000168303 | MPLKIP | 0.34 | 0.07 | 9.85E-07 | 4.18E-06 |
| ENSG00000130177 | CDC16 | 0.34 | 0.07 | 2.19E-06 | 9.00E-06 |
| ENSG00000204856 | FAM216A | 0.34 | 0.07 | 2.75E-06 | 1.12E-05 |
| ENSG00000060069 | CTDP1 | 0.34 | 0.07 | 3.36E-06 | 1.36E-05 |
| ENSG00000130956 | HABP4 | 0.34 | 0.07 | 4.19E-06 | 1.68E-05 |
| ENSG00000130856 | ZNF236 | 0.34 | 0.08 | 6.29E-06 | 2.48E-05 |
| ENSG00000065491 | TBC1D22B | 0.34 | 0.08 | 8.52E-06 | 3.32E-05 |
| ENSG00000141198 | TOM1L1 | 0.34 | 0.08 | 1.21E-05 | 4.63E-05 |
| ENSG00000132589 | FLOT2 | 0.34 | 0.08 | 1.41E-05 | 5.38E-05 |
| ENSG00000141376 | BCAS3 | 0.34 | 0.09 | 3.56E-05 | 1.30E-04 |
| ENSG00000133265 | HSPBP1 | 0.34 | 0.09 | 9.72E-05 | 3.39E-04 |
| ENSG00000092020 | PPP2R3C | 0.34 | 0.09 | 1.01E-04 | 3.52E-04 |
| ENSG00000150787 | PTS | 0.34 | 0.09 | 1.22E-04 | 4.19E-04 |
| ENSG00000164331 | ANKRA2 | 0.34 | 0.1 | 2.24E-04 | 7.48E-04 |
| ENSG00000140104 | CLBA1 | 0.34 | 0.1 | 2.78E-04 | 9.16E-04 |
| ENSG00000132005 | RFX1 | 0.34 | 0.1 | 3.10E-04 | 1.01E-03 |
| ENSG00000116151 | MORN1 | 0.34 | 0.11 | 1.33E-03 | 3.96E-03 |
| ENSG00000125388 | GRK4 | 0.34 | 0.11 | 1.47E-03 | 4.34E-03 |
| ENSG00000050438 | SLC4A8 | 0.34 | 0.11 | 1.67E-03 | 4.91E-03 |
| ENSG00000228801 | PCMTD1-DT | 0.34 | 0.12 | 2.53E-03 | 7.22E-03 |
| ENSG00000198478 | SH3BGRL2 | 0.34 | 0.12 | 2.76E-03 | 7.80E-03 |
| ENSG00000108830 | RND2 | 0.34 | 0.12 | 3.60E-03 | 9.96E-03 |
| ENSG00000274605 | PCCA-DT | 0.34 | 0.12 | 3.83E-03 | 1.06E-02 |
| ENSG00000116254 | CHD5 | 0.34 | 0.12 | 4.02E-03 | 1.10E-02 |
| ENSG00000285410 | GABPB1-IT1 | 0.34 | 0.13 | 4.69E-03 | 1.27E-02 |
| ENSG00000205808 | PLPP6 | 0.34 | 0.13 | 5.88E-03 | 1.56E-02 |
| ENSG00000139194 | RBP5 | 0.34 | 0.15 | 1.26E-02 | 3.11E-02 |
| ENSG00000177675 | CD163L1 | 0.34 | 0.15 | 1.47E-02 | 3.56E-02 |
| ENSG00000048052 | HDAC9 | 0.34 | 0.15 | 1.78E-02 | 4.24E-02 |
| ENSG00000161920 | MED11 | 0.34 | 0.16 | 1.80E-02 | 4.27E-02 |
| ENSG00000102878 | HSF4 | 0.34 | 0.16 | 2.09E-02 | 4.90E-02 |
| ENSG00000229117 | RPL41 | 0.33 | 0.03 | 3.40E-24 | 3.72E-23 |
| ENSG00000022840 | RNF10 | 0.33 | 0.04 | 5.95E-18 | 5.05E-17 |
| ENSG00000063046 | EIF4B | 0.33 | 0.04 | 6.19E-18 | 5.24E-17 |
| ENSG00000175390 | EIF3F | 0.33 | 0.04 | 5.87E-16 | 4.55E-15 |
| ENSG00000170540 | ARL6IP1 | 0.33 | 0.04 | 5.21E-15 | 3.87E-14 |
| ENSG00000168092 | PAFAH1B2 | 0.33 | 0.04 | 1.09E-14 | 7.95E-14 |
| ENSG00000179295 | PTPN11 | 0.33 | 0.05 | 1.33E-13 | 9.17E-13 |
| ENSG00000112893 | MAN2A1 | 0.33 | 0.05 | 4.03E-13 | 2.71E-12 |
| ENSG00000146535 | GNA12 | 0.33 | 0.05 | 5.89E-13 | 3.92E-12 |
| ENSG00000064607 | SUGP2 | 0.33 | 0.05 | 8.27E-13 | 5.45E-12 |
| ENSG00000138430 | OLA1 | 0.33 | 0.05 | 9.60E-13 | 6.30E-12 |
| ENSG00000189266 | PNRC2 | 0.33 | 0.05 | 1.38E-12 | 8.95E-12 |
| ENSG00000109118 | PHF12 | 0.33 | 0.05 | 2.22E-12 | 1.43E-11 |
| ENSG00000042445 | RETSAT | 0.33 | 0.05 | 1.96E-11 | 1.19E-10 |
| ENSG00000147383 | NSDHL | 0.33 | 0.05 | 2.76E-11 | 1.66E-10 |

| | | | | | |
|---|---|---|---|---|---|
| ENSG00000205213 | LGR4 | 0.33 | 0.05 | 4.98E-11 | 2.94E-10 |
| ENSG00000052749 | RRP12 | 0.33 | 0.05 | 1.13E-10 | 6.53E-10 |
| ENSG00000112983 | BRD8 | 0.33 | 0.05 | 1.63E-10 | 9.29E-10 |
| ENSG00000114209 | PDCD10 | 0.33 | 0.05 | 2.00E-10 | 1.14E-09 |
| ENSG00000154305 | MIA3 | 0.33 | 0.05 | 8.23E-10 | 4.47E-09 |
| ENSG00000185825 | BCAP31 | 0.33 | 0.05 | 1.43E-09 | 7.65E-09 |
| ENSG00000144048 | DUSP11 | 0.33 | 0.06 | 1.73E-09 | 9.17E-09 |
| ENSG00000120784 | ZFP30 | 0.33 | 0.06 | 1.86E-09 | 9.83E-09 |
| ENSG00000156345 | CDK20 | 0.33 | 0.05 | 2.04E-09 | 1.08E-08 |
| ENSG00000112031 | MTRF1L | 0.33 | 0.06 | 3.18E-09 | 1.65E-08 |
| ENSG00000145246 | ATP10D | 0.33 | 0.06 | 3.62E-09 | 1.87E-08 |
| ENSG00000159228 | CBR1 | 0.33 | 0.06 | 3.65E-09 | 1.89E-08 |
| ENSG00000008394 | MGST1 | 0.33 | 0.06 | 7.76E-09 | 3.92E-08 |
| ENSG00000145833 | DDX46 | 0.33 | 0.06 | 1.81E-08 | 8.87E-08 |
| ENSG00000153066 | TXNDC11 | 0.33 | 0.06 | 2.93E-08 | 1.42E-07 |
| ENSG00000273841 | TAF9 | 0.33 | 0.06 | 3.62E-08 | 1.74E-07 |
| ENSG00000105640 | RPL18A | 0.33 | 0.06 | 4.14E-08 | 1.97E-07 |
| ENSG00000106080 | FKBP14 | 0.33 | 0.06 | 4.32E-08 | 2.06E-07 |
| ENSG00000179532 | DNHD1 | 0.33 | 0.06 | 1.27E-07 | 5.84E-07 |
| ENSG00000182117 | NOP10 | 0.33 | 0.06 | 1.30E-07 | 5.99E-07 |
| ENSG00000169609 | C15orf40 | 0.33 | 0.06 | 1.31E-07 | 6.01E-07 |
| ENSG00000136636 | KCTD3 | 0.33 | 0.06 | 1.89E-07 | 8.58E-07 |
| ENSG00000054267 | ARID4B | 0.33 | 0.06 | 2.14E-07 | 9.67E-07 |
| ENSG00000152484 | USP12 | 0.33 | 0.07 | 3.26E-07 | 1.45E-06 |
| ENSG00000107833 | NPM3 | 0.33 | 0.07 | 4.26E-07 | 1.88E-06 |
| ENSG00000005302 | MSL3 | 0.33 | 0.07 | 5.76E-07 | 2.51E-06 |
| ENSG00000226752 | CUTALP | 0.33 | 0.07 | 6.01E-07 | 2.61E-06 |
| ENSG00000268916 | CSAG3 | 0.33 | 0.07 | 7.08E-07 | 3.05E-06 |
| ENSG00000089351 | GRAMD1A | 0.33 | 0.07 | 9.06E-07 | 3.86E-06 |
| ENSG00000117016 | RIMS3 | 0.33 | 0.07 | 1.18E-06 | 4.96E-06 |
| ENSG00000175221 | MED16 | 0.33 | 0.07 | 1.43E-06 | 5.97E-06 |
| ENSG00000104969 | SGTA | 0.33 | 0.07 | 2.15E-06 | 8.84E-06 |
| ENSG00000119013 | NDUFB3 | 0.33 | 0.07 | 2.24E-06 | 9.18E-06 |
| ENSG00000077585 | GPR137B | 0.33 | 0.07 | 2.61E-06 | 1.07E-05 |
| ENSG00000140199 | SLC12A6 | 0.33 | 0.07 | 2.74E-06 | 1.12E-05 |
| ENSG00000145022 | TCTA | 0.33 | 0.07 | 2.87E-06 | 1.17E-05 |
| ENSG00000164309 | CMYA5 | 0.33 | 0.07 | 3.75E-06 | 1.51E-05 |
| ENSG00000081177 | EXD2 | 0.33 | 0.07 | 5.68E-06 | 2.25E-05 |
| ENSG00000278318 | ZNF229 | 0.33 | 0.08 | 6.68E-06 | 2.63E-05 |
| ENSG00000152359 | POC5 | 0.33 | 0.08 | 6.80E-06 | 2.67E-05 |
| ENSG00000008324 | SS18L2 | 0.33 | 0.08 | 7.73E-06 | 3.02E-05 |
| ENSG00000180998 | GPR137C | 0.33 | 0.08 | 7.91E-06 | 3.09E-05 |
| ENSG00000138439 | FAM117B | 0.33 | 0.08 | 1.10E-05 | 4.24E-05 |
| ENSG00000251369 | ZNF550 | 0.33 | 0.08 | 1.22E-05 | 4.67E-05 |
| ENSG00000167967 | E4F1 | 0.33 | 0.08 | 1.52E-05 | 5.78E-05 |
| ENSG00000171763 | SPATA5L1 | 0.33 | 0.08 | 1.71E-05 | 6.44E-05 |
| ENSG00000093010 | COMT | 0.33 | 0.08 | 1.91E-05 | 7.19E-05 |
| ENSG00000249915 | PDCD6 | 0.33 | 0.08 | 2.32E-05 | 8.65E-05 |
| ENSG00000133612 | AGAP3 | 0.33 | 0.08 | 2.71E-05 | 1.00E-04 |
| ENSG00000137210 | TMEM14B | 0.33 | 0.08 | 4.04E-05 | 1.47E-04 |

| | | | | | |
|---|---|---|---|---|---|
| ENSG00000131115 | ZNF227 | 0.33 | 0.08 | 4.78E-05 | 1.72E-04 |
| ENSG00000249673 | NOP14-AS1 | 0.33 | 0.09 | 8.30E-05 | 2.91E-04 |
| ENSG00000054282 | SDCCAG8 | 0.33 | 0.09 | 1.59E-04 | 5.40E-04 |
| ENSG00000221890 | NPTXR | 0.33 | 0.09 | 2.30E-04 | 7.67E-04 |
| ENSG00000236675 | MTX1P1 | 0.33 | 0.1 | 4.91E-04 | 1.57E-03 |
| ENSG00000184897 | H1-10 | 0.33 | 0.11 | 1.10E-03 | 3.33E-03 |
| ENSG00000123609 | NMI | 0.33 | 0.11 | 1.24E-03 | 3.71E-03 |
| ENSG00000274827 | LINC01297 | 0.33 | 0.11 | 1.96E-03 | 5.68E-03 |
| ENSG00000037965 | HOXC8 | 0.33 | 0.11 | 2.19E-03 | 6.30E-03 |
| ENSG00000159784 | FAM131B | 0.33 | 0.12 | 3.00E-03 | 8.43E-03 |
| ENSG00000174586 | ZNF497 | 0.33 | 0.12 | 3.34E-03 | 9.29E-03 |
| ENSG00000169972 | PUSL1 | 0.33 | 0.12 | 3.69E-03 | 1.02E-02 |
| ENSG00000290682 | STAG3L5P | 0.33 | 0.13 | 6.13E-03 | 1.62E-02 |
| ENSG00000167363 | FN3K | 0.33 | 0.13 | 6.25E-03 | 1.65E-02 |
| ENSG00000103266 | STUB1 | 0.33 | 0.13 | 6.78E-03 | 1.78E-02 |
| ENSG00000180479 | ZNF571 | 0.33 | 0.13 | 8.89E-03 | 2.27E-02 |
| ENSG00000233237 | LINC00472 | 0.33 | 0.14 | 1.06E-02 | 2.67E-02 |
| ENSG00000257194 | | 0.33 | 0.14 | 1.16E-02 | 2.89E-02 |
| ENSG00000171450 | CDK5R2 | 0.33 | 0.14 | 1.19E-02 | 2.96E-02 |
| ENSG00000213741 | RPS29 | 0.33 | 0.14 | 1.30E-02 | 3.19E-02 |
| ENSG00000105717 | PBX4 | 0.33 | 0.14 | 1.57E-02 | 3.79E-02 |
| ENSG00000102144 | PGK1 | 0.32 | 0.03 | 6.56E-20 | 6.07E-19 |
| ENSG00000197965 | MPZL1 | 0.32 | 0.04 | 1.08E-18 | 9.47E-18 |
| ENSG00000116209 | TMEM59 | 0.32 | 0.04 | 1.12E-17 | 9.39E-17 |
| ENSG00000114770 | ABCC5 | 0.32 | 0.04 | 2.15E-17 | 1.79E-16 |
| ENSG00000130227 | XPO7 | 0.32 | 0.04 | 3.25E-17 | 2.67E-16 |
| ENSG00000204217 | BMPR2 | 0.32 | 0.04 | 2.09E-14 | 1.50E-13 |
| ENSG00000164292 | RHOBTB3 | 0.32 | 0.04 | 4.07E-14 | 2.89E-13 |
| ENSG00000256235 | SMIM3 | 0.32 | 0.04 | 8.17E-14 | 5.71E-13 |
| ENSG00000136732 | GYPC | 0.32 | 0.04 | 2.51E-13 | 1.71E-12 |
| ENSG00000109046 | WSB1 | 0.32 | 0.04 | 3.28E-13 | 2.21E-12 |
| ENSG00000087460 | GNAS | 0.32 | 0.04 | 4.43E-13 | 2.97E-12 |
| ENSG00000108219 | TSPAN14 | 0.32 | 0.05 | 5.52E-12 | 3.45E-11 |
| ENSG00000127022 | CANX | 0.32 | 0.05 | 1.01E-11 | 6.24E-11 |
| ENSG00000119414 | PPP6C | 0.32 | 0.05 | 1.23E-11 | 7.58E-11 |
| ENSG00000086666 | ZFAND6 | 0.32 | 0.05 | 1.30E-11 | 7.96E-11 |
| ENSG00000106153 | CHCHD2 | 0.32 | 0.05 | 2.81E-11 | 1.69E-10 |
| ENSG00000118200 | CAMSAP2 | 0.32 | 0.05 | 1.58E-10 | 9.01E-10 |
| ENSG00000131507 | NDFIP1 | 0.32 | 0.05 | 3.09E-10 | 1.73E-09 |
| ENSG00000158109 | TPRG1L | 0.32 | 0.05 | 7.55E-10 | 4.11E-09 |
| ENSG00000213639 | PPP1CB | 0.32 | 0.05 | 1.52E-09 | 8.09E-09 |
| ENSG00000197045 | GMFB | 0.32 | 0.06 | 6.06E-09 | 3.08E-08 |
| ENSG00000143878 | RHOB | 0.32 | 0.06 | 7.83E-09 | 3.95E-08 |
| ENSG00000070501 | POLB | 0.32 | 0.06 | 1.01E-08 | 5.06E-08 |
| ENSG00000167930 | FAM234A | 0.32 | 0.06 | 1.21E-08 | 6.00E-08 |
| ENSG00000108510 | MED13 | 0.32 | 0.06 | 1.43E-08 | 7.08E-08 |
| ENSG00000084073 | ZMPSTE24 | 0.32 | 0.06 | 1.59E-08 | 7.82E-08 |
| ENSG00000175073 | VCPIP1 | 0.32 | 0.06 | 2.29E-08 | 1.12E-07 |
| ENSG00000130311 | DDA1 | 0.32 | 0.06 | 2.44E-08 | 1.19E-07 |
| ENSG00000105193 | RPS16 | 0.32 | 0.06 | 2.76E-08 | 1.34E-07 |

| | | | | | |
|---|---|---|---|---|---|
| ENSG00000108107 | RPL28 | 0.32 | 0.06 | 3.06E-08 | 1.48E-07 |
| ENSG00000101972 | STAG2 | 0.32 | 0.06 | 4.04E-08 | 1.93E-07 |
| ENSG00000197548 | ATG7 | 0.32 | 0.06 | 9.01E-08 | 4.20E-07 |
| ENSG00000008441 | NFIX | 0.32 | 0.06 | 1.23E-07 | 5.65E-07 |
| ENSG00000178074 | C2orf69 | 0.32 | 0.06 | 3.07E-07 | 1.37E-06 |
| ENSG00000048392 | RRM2B | 0.32 | 0.06 | 7.59E-07 | 3.26E-06 |
| ENSG00000163635 | ATXN7 | 0.32 | 0.07 | 1.72E-06 | 7.12E-06 |
| ENSG00000138592 | USP8 | 0.32 | 0.07 | 2.14E-06 | 8.80E-06 |
| ENSG00000135924 | DNAJB2 | 0.32 | 0.07 | 2.27E-06 | 9.32E-06 |
| ENSG00000166171 | DPCD | 0.32 | 0.07 | 2.34E-06 | 9.60E-06 |
| ENSG00000138777 | PPA2 | 0.32 | 0.07 | 5.54E-06 | 2.20E-05 |
| ENSG00000164930 | FZD6 | 0.32 | 0.07 | 8.33E-06 | 3.24E-05 |
| ENSG00000078902 | TOLLIP | 0.32 | 0.07 | 9.62E-06 | 3.73E-05 |
| ENSG00000107829 | FBXW4 | 0.32 | 0.08 | 1.19E-05 | 4.57E-05 |
| ENSG00000198886 | MT-ND4 | 0.32 | 0.08 | 1.56E-05 | 5.91E-05 |
| ENSG00000172270 | BSG | 0.32 | 0.08 | 2.24E-05 | 8.38E-05 |
| ENSG00000040531 | CTNS | 0.32 | 0.08 | 2.29E-05 | 8.53E-05 |
| ENSG00000110711 | AIP | 0.32 | 0.08 | 2.42E-05 | 9.01E-05 |
| ENSG00000231806 | PCAT7 | 0.32 | 0.08 | 3.96E-05 | 1.44E-04 |
| ENSG00000245937 | SLC12A2-DT | 0.32 | 0.08 | 4.63E-05 | 1.67E-04 |
| ENSG00000223960 | CHROMR | 0.32 | 0.08 | 6.46E-05 | 2.30E-04 |
| ENSG00000176422 | SPRYD4 | 0.32 | 0.08 | 7.73E-05 | 2.73E-04 |
| ENSG00000128891 | CCDC32 | 0.32 | 0.09 | 1.14E-04 | 3.92E-04 |
| ENSG00000153487 | ING1 | 0.32 | 0.09 | 1.46E-04 | 4.98E-04 |
| ENSG00000136826 | KLF4 | 0.32 | 0.09 | 1.53E-04 | 5.20E-04 |
| ENSG00000130193 | THEM6 | 0.32 | 0.09 | 1.91E-04 | 6.40E-04 |
| ENSG00000214063 | TSPAN4 | 0.32 | 0.09 | 2.40E-04 | 7.97E-04 |
| ENSG00000105229 | PIAS4 | 0.32 | 0.09 | 3.33E-04 | 1.08E-03 |
| ENSG00000092531 | SNAP23 | 0.32 | 0.09 | 3.96E-04 | 1.28E-03 |
| ENSG00000114841 | DNAH1 | 0.32 | 0.1 | 4.27E-04 | 1.37E-03 |
| ENSG00000143507 | DUSP10 | 0.32 | 0.1 | 4.91E-04 | 1.56E-03 |
| ENSG00000290720 | ZNF542P | 0.32 | 0.1 | 7.27E-04 | 2.26E-03 |
| ENSG00000167920 | KRT10-AS1 | 0.32 | 0.1 | 1.03E-03 | 3.13E-03 |
| ENSG00000188092 | GPR89B | 0.32 | 0.1 | 1.52E-03 | 4.50E-03 |
| ENSG00000088538 | DOCK3 | 0.32 | 0.12 | 3.93E-03 | 1.08E-02 |
| ENSG00000143919 | CAMKMT | 0.32 | 0.12 | 5.39E-03 | 1.44E-02 |
| ENSG00000055163 | CYFIP2 | 0.32 | 0.13 | 9.39E-03 | 2.39E-02 |
| ENSG00000242071 | RPL7AP6 | 0.32 | 0.13 | 1.16E-02 | 2.89E-02 |
| ENSG00000179431 | FJX1 | 0.32 | 0.14 | 1.27E-02 | 3.13E-02 |
| ENSG00000103254 | ANTKMT | 0.32 | 0.14 | 1.33E-02 | 3.27E-02 |
| ENSG00000122121 | XPNPEP2 | 0.32 | 0.14 | 1.34E-02 | 3.28E-02 |
| ENSG00000264112 | | 0.32 | 0.15 | 1.78E-02 | 4.24E-02 |
| ENSG00000135108 | FBXO21 | 0.31 | 0.04 | 2.35E-18 | 2.03E-17 |
| ENSG00000154736 | ADAMTS5 | 0.31 | 0.04 | 5.51E-18 | 4.68E-17 |
| ENSG00000168610 | STAT3 | 0.31 | 0.04 | 7.07E-16 | 5.45E-15 |
| ENSG00000143319 | ISG20L2 | 0.31 | 0.04 | 7.68E-16 | 5.91E-15 |
| ENSG00000142676 | RPL11 | 0.31 | 0.04 | 2.36E-14 | 1.70E-13 |
| ENSG00000138029 | HADHB | 0.31 | 0.04 | 4.40E-14 | 3.11E-13 |
| ENSG00000163681 | SLMAP | 0.31 | 0.04 | 2.67E-13 | 1.81E-12 |
| ENSG00000231500 | RPS18 | 0.31 | 0.04 | 4.47E-13 | 2.99E-12 |

| | | | | | |
|---|---|---|---|---|---|
| ENSG00000118515 | SGK1 | 0.31 | 0.04 | 5.95E-13 | 3.96E-12 |
| ENSG00000096717 | SIRT1 | 0.31 | 0.04 | 6.26E-13 | 4.16E-12 |
| ENSG00000172239 | PAIP1 | 0.31 | 0.04 | 1.20E-12 | 7.80E-12 |
| ENSG00000014641 | MDH1 | 0.31 | 0.04 | 1.90E-12 | 1.23E-11 |
| ENSG00000185963 | BICD2 | 0.31 | 0.05 | 4.39E-12 | 2.77E-11 |
| ENSG00000139726 | DENR | 0.31 | 0.05 | 9.27E-12 | 5.73E-11 |
| ENSG00000185532 | PRKG1 | 0.31 | 0.05 | 1.55E-10 | 8.86E-10 |
| ENSG00000263465 | SRSF8 | 0.31 | 0.05 | 2.68E-10 | 1.51E-09 |
| ENSG00000163428 | LRRC58 | 0.31 | 0.05 | 5.56E-10 | 3.06E-09 |
| ENSG00000114857 | NKTR | 0.31 | 0.05 | 2.56E-09 | 1.34E-08 |
| ENSG00000132549 | VPS13B | 0.31 | 0.05 | 4.06E-09 | 2.09E-08 |
| ENSG00000168438 | CDC40 | 0.31 | 0.05 | 4.38E-09 | 2.25E-08 |
| ENSG00000149357 | LAMTOR1 | 0.31 | 0.05 | 6.87E-09 | 3.48E-08 |
| ENSG00000167635 | ZNF146 | 0.31 | 0.05 | 1.24E-08 | 6.14E-08 |
| ENSG00000067208 | EVI5 | 0.31 | 0.06 | 1.74E-08 | 8.58E-08 |
| ENSG00000165006 | UBAP1 | 0.31 | 0.05 | 1.79E-08 | 8.80E-08 |
| ENSG00000103042 | SLC38A7 | 0.31 | 0.06 | 2.11E-08 | 1.03E-07 |
| ENSG00000151532 | VTI1A | 0.31 | 0.06 | 7.83E-08 | 3.66E-07 |
| ENSG00000151327 | FAM177A1 | 0.31 | 0.06 | 9.08E-08 | 4.23E-07 |
| ENSG00000106608 | URGCP | 0.31 | 0.06 | 2.27E-07 | 1.02E-06 |
| ENSG00000021574 | SPAST | 0.31 | 0.06 | 3.33E-07 | 1.48E-06 |
| ENSG00000112701 | SENP6 | 0.31 | 0.06 | 4.00E-07 | 1.77E-06 |
| ENSG00000204178 | MACO1 | 0.31 | 0.06 | 7.05E-07 | 3.04E-06 |
| ENSG00000149257 | SERPINH1 | 0.31 | 0.07 | 1.72E-06 | 7.12E-06 |
| ENSG00000128563 | PRKRIP1 | 0.31 | 0.07 | 2.08E-06 | 8.58E-06 |
| ENSG00000100142 | POLR2F | 0.31 | 0.07 | 3.53E-06 | 1.43E-05 |
| ENSG00000214087 | ARL16 | 0.31 | 0.07 | 5.20E-06 | 2.07E-05 |
| ENSG00000198746 | GPATCH3 | 0.31 | 0.07 | 5.71E-06 | 2.26E-05 |
| ENSG00000110717 | NDUFS8 | 0.31 | 0.07 | 1.61E-05 | 6.08E-05 |
| ENSG00000170043 | TRAPPC1 | 0.31 | 0.07 | 1.74E-05 | 6.56E-05 |
| ENSG00000088826 | SMOX | 0.31 | 0.08 | 2.15E-05 | 8.04E-05 |
| ENSG00000225439 | BOLA3-DT | 0.31 | 0.08 | 2.24E-05 | 8.36E-05 |
| ENSG00000172716 | SLFN11 | 0.31 | 0.08 | 3.13E-05 | 1.15E-04 |
| ENSG00000113638 | TTC33 | 0.31 | 0.08 | 1.03E-04 | 3.58E-04 |
| ENSG00000133114 | GPALPP1 | 0.31 | 0.08 | 1.22E-04 | 4.20E-04 |
| ENSG00000170260 | ZNF212 | 0.31 | 0.09 | 1.80E-04 | 6.07E-04 |
| ENSG00000158552 | ZFAND2B | 0.31 | 0.09 | 2.32E-04 | 7.71E-04 |
| ENSG00000100299 | ARSA | 0.31 | 0.09 | 2.70E-04 | 8.92E-04 |
| ENSG00000171703 | TCEA2 | 0.31 | 0.09 | 3.00E-04 | 9.83E-04 |
| ENSG00000052126 | PLEKHA5 | 0.31 | 0.09 | 3.14E-04 | 1.03E-03 |
| ENSG00000132563 | REEP2 | 0.31 | 0.09 | 3.32E-04 | 1.08E-03 |
| ENSG00000213983 | AP1G2 | 0.31 | 0.09 | 3.67E-04 | 1.19E-03 |
| ENSG00000197283 | SYNGAP1 | 0.31 | 0.09 | 5.00E-04 | 1.59E-03 |
| ENSG00000217236 | SP9 | 0.31 | 0.09 | 5.05E-04 | 1.61E-03 |
| ENSG00000166886 | NAB2 | 0.31 | 0.09 | 5.18E-04 | 1.65E-03 |
| ENSG00000059145 | UNKL | 0.31 | 0.09 | 6.33E-04 | 1.98E-03 |
| ENSG00000184194 | GPR173 | 0.31 | 0.09 | 6.50E-04 | 2.04E-03 |
| ENSG00000141337 | ARSG | 0.31 | 0.1 | 9.59E-04 | 2.92E-03 |
| ENSG00000168067 | MAP4K2 | 0.31 | 0.1 | 9.94E-04 | 3.02E-03 |
| ENSG00000185187 | SIGIRR | 0.31 | 0.1 | 1.11E-03 | 3.34E-03 |

| | | | | | |
|---|---|---|---|---|---|
| ENSG00000182093 | GET1 | 0.31 | 0.1 | 1.21E-03 | 3.62E-03 |
| ENSG00000143452 | HORMAD1 | 0.31 | 0.1 | 1.42E-03 | 4.23E-03 |
| ENSG00000173653 | RCE1 | 0.31 | 0.1 | 1.50E-03 | 4.43E-03 |
| ENSG00000177738 | ANXA2R-OT1 | 0.31 | 0.1 | 1.50E-03 | 4.44E-03 |
| ENSG00000182324 | KCNJ14 | 0.31 | 0.1 | 1.93E-03 | 5.62E-03 |
| ENSG00000102221 | JADE3 | 0.31 | 0.11 | 2.36E-03 | 6.76E-03 |
| ENSG00000167874 | TMEM88 | 0.31 | 0.11 | 3.16E-03 | 8.83E-03 |
| ENSG00000175471 | MCTP1 | 0.31 | 0.11 | 3.71E-03 | 1.02E-02 |
| ENSG00000243199 | RPL6P10 | 0.31 | 0.12 | 5.15E-03 | 1.38E-02 |
| ENSG00000129354 | AP1M2 | 0.31 | 0.12 | 6.07E-03 | 1.61E-02 |
| ENSG00000187987 | ZSCAN23 | 0.31 | 0.14 | 1.49E-02 | 3.61E-02 |
| ENSG00000205609 | EIF3CL | 0.31 | 0.14 | 1.59E-02 | 3.82E-02 |
| ENSG00000183291 | SELENOF | 0.3 | 0.03 | 2.39E-18 | 2.06E-17 |
| ENSG00000113648 | MACROH2A1 | 0.3 | 0.04 | 1.84E-15 | 1.39E-14 |
| ENSG00000148606 | POLR3A | 0.3 | 0.04 | 3.02E-15 | 2.27E-14 |
| ENSG00000166913 | YWHAB | 0.3 | 0.04 | 1.06E-14 | 7.73E-14 |
| ENSG00000183726 | TMEM50A | 0.3 | 0.04 | 2.64E-13 | 1.80E-12 |
| ENSG00000108298 | RPL19 | 0.3 | 0.04 | 3.04E-13 | 2.06E-12 |
| ENSG00000100813 | ACIN1 | 0.3 | 0.04 | 3.48E-13 | 2.35E-12 |
| ENSG00000110931 | CAMKK2 | 0.3 | 0.04 | 5.88E-13 | 3.92E-12 |
| ENSG00000143751 | SDE2 | 0.3 | 0.04 | 1.22E-12 | 7.98E-12 |
| ENSG00000100796 | PPP4R3A | 0.3 | 0.04 | 3.22E-12 | 2.05E-11 |
| ENSG00000008869 | HEATR5B | 0.3 | 0.04 | 7.13E-12 | 4.44E-11 |
| ENSG00000160218 | TRAPPC10 | 0.3 | 0.05 | 4.40E-11 | 2.61E-10 |
| ENSG00000170852 | KBTBD2 | 0.3 | 0.05 | 5.66E-11 | 3.33E-10 |
| ENSG00000093183 | SEC22C | 0.3 | 0.05 | 6.53E-11 | 3.83E-10 |
| ENSG00000122965 | RBM19 | 0.3 | 0.05 | 6.82E-11 | 3.99E-10 |
| ENSG00000103479 | RBL2 | 0.3 | 0.05 | 8.75E-11 | 5.09E-10 |
| ENSG00000102471 | NDFIP2 | 0.3 | 0.05 | 1.46E-10 | 8.35E-10 |
| ENSG00000107679 | PLEKHA1 | 0.3 | 0.05 | 1.59E-10 | 9.09E-10 |
| ENSG00000163682 | RPL9 | 0.3 | 0.05 | 6.10E-10 | 3.35E-09 |
| ENSG00000242247 | ARFGAP3 | 0.3 | 0.05 | 6.96E-10 | 3.81E-09 |
| ENSG00000185787 | MORF4L1 | 0.3 | 0.05 | 1.60E-09 | 8.51E-09 |
| ENSG00000167118 | URM1 | 0.3 | 0.05 | 8.05E-09 | 4.06E-08 |
| ENSG00000100614 | PPM1A | 0.3 | 0.05 | 8.13E-09 | 4.09E-08 |
| ENSG00000135837 | CEP350 | 0.3 | 0.05 | 1.68E-08 | 8.27E-08 |
| ENSG00000175376 | EIF1AD | 0.3 | 0.05 | 2.32E-08 | 1.13E-07 |
| ENSG00000146425 | DYNLT1 | 0.3 | 0.06 | 5.20E-08 | 2.46E-07 |
| ENSG00000114423 | CBLB | 0.3 | 0.06 | 9.69E-08 | 4.50E-07 |
| ENSG00000116521 | SCAMP3 | 0.3 | 0.06 | 1.16E-07 | 5.34E-07 |
| ENSG00000111726 | CMAS | 0.3 | 0.06 | 1.29E-07 | 5.94E-07 |
| ENSG00000092978 | GPATCH2 | 0.3 | 0.06 | 1.42E-07 | 6.52E-07 |
| ENSG00000112941 | TENT4A | 0.3 | 0.06 | 2.48E-07 | 1.11E-06 |
| ENSG00000146247 | PHIP | 0.3 | 0.06 | 2.90E-07 | 1.30E-06 |
| ENSG00000124357 | NAGK | 0.3 | 0.06 | 4.89E-07 | 2.14E-06 |
| ENSG00000197016 | ZNF470 | 0.3 | 0.06 | 5.39E-07 | 2.35E-06 |
| ENSG00000185043 | CIB1 | 0.3 | 0.06 | 6.29E-07 | 2.73E-06 |
| ENSG00000134046 | MBD2 | 0.3 | 0.07 | 4.56E-06 | 1.82E-05 |
| ENSG00000187051 | RPS19BP1 | 0.3 | 0.07 | 1.54E-05 | 5.84E-05 |
| ENSG00000104231 | ZFAND1 | 0.3 | 0.07 | 1.97E-05 | 7.39E-05 |

| | | | | | |
|---|---|---|---|---|---|
| ENSG00000213190 | MLLT11 | 0.3 | 0.07 | 2.14E-05 | 8.03E-05 |
| ENSG00000115317 | HTRA2 | 0.3 | 0.07 | 2.61E-05 | 9.67E-05 |
| ENSG00000181649 | PHLDA2 | 0.3 | 0.07 | 3.18E-05 | 1.17E-04 |
| ENSG00000257093 | DENND11 | 0.3 | 0.07 | 4.33E-05 | 1.57E-04 |
| ENSG00000174405 | LIG4 | 0.3 | 0.08 | 4.74E-05 | 1.71E-04 |
| ENSG00000090565 | RAB11FIP3 | 0.3 | 0.08 | 5.35E-05 | 1.92E-04 |
| ENSG00000224032 | EPB41L4A-AS1 | 0.3 | 0.08 | 6.46E-05 | 2.30E-04 |
| ENSG00000166582 | CENPV | 0.3 | 0.08 | 7.39E-05 | 2.61E-04 |
| ENSG00000034152 | MAP2K3 | 0.3 | 0.08 | 1.08E-04 | 3.73E-04 |
| ENSG00000166912 | MTMR10 | 0.3 | 0.08 | 1.43E-04 | 4.88E-04 |
| ENSG00000013306 | SLC25A39 | 0.3 | 0.08 | 1.47E-04 | 5.02E-04 |
| ENSG00000003509 | NDUFAF7 | 0.3 | 0.08 | 1.63E-04 | 5.53E-04 |
| ENSG00000196642 | RABL6 | 0.3 | 0.08 | 1.70E-04 | 5.76E-04 |
| ENSG00000147853 | AK3 | 0.3 | 0.08 | 1.73E-04 | 5.84E-04 |
| ENSG00000188818 | ZDHHC11 | 0.3 | 0.08 | 3.02E-04 | 9.89E-04 |
| ENSG00000106089 | STX1A | 0.3 | 0.09 | 3.28E-04 | 1.07E-03 |
| ENSG00000147027 | TMEM47 | 0.3 | 0.09 | 6.17E-04 | 1.94E-03 |
| ENSG00000170340 | B3GNT2 | 0.3 | 0.09 | 6.36E-04 | 1.99E-03 |
| ENSG00000167985 | SDHAF2 | 0.3 | 0.09 | 8.42E-04 | 2.59E-03 |
| ENSG00000041988 | THAP3 | 0.3 | 0.09 | 9.98E-04 | 3.03E-03 |
| ENSG00000180626 | ZNF594 | 0.3 | 0.1 | 1.23E-03 | 3.69E-03 |
| ENSG00000143545 | RAB13 | 0.3 | 0.1 | 1.56E-03 | 4.60E-03 |
| ENSG00000233016 | SNHG7 | 0.3 | 0.1 | 1.95E-03 | 5.67E-03 |
| ENSG00000127666 | TICAM1 | 0.3 | 0.1 | 1.96E-03 | 5.69E-03 |
| ENSG00000198933 | TBKBP1 | 0.3 | 0.1 | 2.44E-03 | 6.97E-03 |
| ENSG00000116977 | LGALS8 | 0.3 | 0.11 | 3.52E-03 | 9.75E-03 |
| ENSG00000174080 | CTSF | 0.3 | 0.11 | 4.10E-03 | 1.12E-02 |
| ENSG00000214654 | B3GALT9 | 0.3 | 0.12 | 6.97E-03 | 1.82E-02 |
| ENSG00000168040 | FADD | 0.3 | 0.12 | 7.09E-03 | 1.85E-02 |
| ENSG00000174977 | PAIP1P2 | 0.3 | 0.12 | 7.74E-03 | 2.01E-02 |
| ENSG00000182600 | SNORC | 0.3 | 0.13 | 1.16E-02 | 2.88E-02 |
| ENSG00000280206 | | 0.3 | 0.13 | 1.46E-02 | 3.54E-02 |
| ENSG00000197483 | ZNF628 | 0.3 | 0.13 | 1.75E-02 | 4.17E-02 |
| ENSG00000163466 | ARPC2 | 0.29 | 0.04 | 2.88E-17 | 2.38E-16 |
| ENSG00000166847 | DCTN5 | 0.29 | 0.04 | 1.76E-16 | 1.40E-15 |
| ENSG00000136816 | TOR1B | 0.29 | 0.04 | 1.69E-14 | 1.23E-13 |
| ENSG00000187678 | SPRY4 | 0.29 | 0.04 | 5.85E-14 | 4.12E-13 |
| ENSG00000167986 | DDB1 | 0.29 | 0.04 | 6.04E-14 | 4.25E-13 |
| ENSG00000157916 | RER1 | 0.29 | 0.04 | 1.84E-13 | 1.26E-12 |
| ENSG00000198369 | SPRED2 | 0.29 | 0.04 | 1.91E-12 | 1.23E-11 |
| ENSG00000078369 | GNB1 | 0.29 | 0.04 | 3.77E-12 | 2.39E-11 |
| ENSG00000101654 | RNMT | 0.29 | 0.04 | 3.99E-11 | 2.37E-10 |
| ENSG00000148248 | SURF4 | 0.29 | 0.05 | 6.06E-11 | 3.56E-10 |
| ENSG00000091844 | RGS17 | 0.29 | 0.05 | 1.25E-09 | 6.71E-09 |
| ENSG00000182220 | ATP6AP2 | 0.29 | 0.05 | 2.09E-09 | 1.10E-08 |
| ENSG00000125386 | FAM193A | 0.29 | 0.05 | 2.34E-09 | 1.23E-08 |
| ENSG00000070087 | PFN2 | 0.29 | 0.05 | 3.49E-09 | 1.81E-08 |
| ENSG00000164442 | CITED2 | 0.29 | 0.05 | 5.45E-09 | 2.78E-08 |
| ENSG00000077721 | UBE2A | 0.29 | 0.05 | 9.89E-09 | 4.96E-08 |
| ENSG00000115464 | USP34 | 0.29 | 0.05 | 1.51E-08 | 7.44E-08 |

| | | | | | |
|---|---|---|---|---|---|
| ENSG00000072849 | DERL2 | 0.29 | 0.05 | 5.30E-08 | 2.51E-07 |
| ENSG00000257103 | LSM14A | 0.29 | 0.05 | 8.69E-08 | 4.05E-07 |
| ENSG00000189339 | SLC35E2B | 0.29 | 0.05 | 1.00E-07 | 4.65E-07 |
| ENSG00000112378 | PERP | 0.29 | 0.06 | 1.24E-07 | 5.71E-07 |
| ENSG00000163719 | MTMR14 | 0.29 | 0.06 | 1.69E-07 | 7.71E-07 |
| ENSG00000198887 | SMC5 | 0.29 | 0.06 | 1.91E-07 | 8.69E-07 |
| ENSG00000188994 | ZNF292 | 0.29 | 0.06 | 2.14E-07 | 9.67E-07 |
| ENSG00000163138 | PACRGL | 0.29 | 0.06 | 2.14E-07 | 9.68E-07 |
| ENSG00000163531 | NFASC | 0.29 | 0.06 | 2.41E-07 | 1.09E-06 |
| ENSG00000162139 | NEU3 | 0.29 | 0.06 | 2.69E-07 | 1.21E-06 |
| ENSG00000122674 | CCZ1 | 0.29 | 0.06 | 3.25E-07 | 1.45E-06 |
| ENSG00000143776 | CDC42BPA | 0.29 | 0.06 | 3.39E-07 | 1.50E-06 |
| ENSG00000103091 | WDR59 | 0.29 | 0.06 | 3.79E-07 | 1.68E-06 |
| ENSG00000122643 | NT5C3A | 0.29 | 0.06 | 4.43E-07 | 1.95E-06 |
| ENSG00000085832 | EPS15 | 0.29 | 0.06 | 5.73E-07 | 2.49E-06 |
| ENSG00000027697 | IFNGR1 | 0.29 | 0.06 | 1.45E-06 | 6.09E-06 |
| ENSG00000135365 | PHF21A | 0.29 | 0.06 | 1.55E-06 | 6.46E-06 |
| ENSG00000129235 | TXNDC17 | 0.29 | 0.06 | 3.48E-06 | 1.40E-05 |
| ENSG00000138698 | RAP1GDS1 | 0.29 | 0.06 | 4.28E-06 | 1.72E-05 |
| ENSG00000138382 | METTL5 | 0.29 | 0.07 | 6.09E-06 | 2.41E-05 |
| ENSG00000103494 | RPGRIP1L | 0.29 | 0.07 | 8.38E-06 | 3.26E-05 |
| ENSG00000205659 | LIN52 | 0.29 | 0.07 | 1.56E-05 | 5.92E-05 |
| ENSG00000154114 | TBCEL | 0.29 | 0.07 | 1.69E-05 | 6.40E-05 |
| ENSG00000140400 | MAN2C1 | 0.29 | 0.07 | 1.90E-05 | 7.14E-05 |
| ENSG00000163602 | RYBP | 0.29 | 0.07 | 1.95E-05 | 7.33E-05 |
| ENSG00000113211 | PCDHB6 | 0.29 | 0.07 | 2.16E-05 | 8.09E-05 |
| ENSG00000205629 | LCMT1 | 0.29 | 0.07 | 3.09E-05 | 1.14E-04 |
| ENSG00000116685 | KIAA2013 | 0.29 | 0.07 | 6.78E-05 | 2.41E-04 |
| ENSG00000188186 | LAMTOR4 | 0.29 | 0.07 | 8.25E-05 | 2.90E-04 |
| ENSG00000136111 | TBC1D4 | 0.29 | 0.07 | 8.91E-05 | 3.12E-04 |
| ENSG00000120159 | CAAP1 | 0.29 | 0.08 | 9.03E-05 | 3.16E-04 |
| ENSG00000185813 | PCYT2 | 0.29 | 0.08 | 1.14E-04 | 3.93E-04 |
| ENSG00000121542 | SEC22A | 0.29 | 0.08 | 1.85E-04 | 6.24E-04 |
| ENSG00000026950 | BTN3A1 | 0.29 | 0.08 | 2.35E-04 | 7.79E-04 |
| ENSG00000182916 | TCEAL7 | 0.29 | 0.08 | 3.00E-04 | 9.81E-04 |
| ENSG00000197948 | FCHSD1 | 0.29 | 0.08 | 3.89E-04 | 1.26E-03 |
| ENSG00000182318 | ZSCAN22 | 0.29 | 0.08 | 4.73E-04 | 1.51E-03 |
| ENSG00000158050 | DUSP2 | 0.29 | 0.09 | 6.22E-04 | 1.95E-03 |
| ENSG00000136247 | ZDHHC4 | 0.29 | 0.09 | 6.71E-04 | 2.10E-03 |
| ENSG00000130734 | ATG4D | 0.29 | 0.09 | 9.60E-04 | 2.93E-03 |
| ENSG00000286067 | | 0.29 | 0.09 | 9.77E-04 | 2.98E-03 |
| ENSG00000007047 | MARK4 | 0.29 | 0.09 | 1.08E-03 | 3.26E-03 |
| ENSG00000256053 | COA8 | 0.29 | 0.09 | 1.29E-03 | 3.86E-03 |
| ENSG00000160961 | ZNF333 | 0.29 | 0.09 | 1.35E-03 | 4.01E-03 |
| ENSG00000123689 | G0S2 | 0.29 | 0.1 | 1.46E-03 | 4.32E-03 |
| ENSG00000151657 | KIN | 0.29 | 0.09 | 1.46E-03 | 4.33E-03 |
| ENSG00000185015 | CA13 | 0.29 | 0.1 | 2.28E-03 | 6.55E-03 |
| ENSG00000126890 | CTAG2 | 0.29 | 0.1 | 2.36E-03 | 6.75E-03 |
| ENSG00000181638 | ZFP41 | 0.29 | 0.1 | 2.61E-03 | 7.40E-03 |
| ENSG00000128266 | GNAZ | 0.29 | 0.1 | 3.16E-03 | 8.84E-03 |

| | | | | | |
|---|---|---|---|---|---|
| ENSG00000049192 | ADAMTS6 | 0.29 | 0.1 | 3.43E-03 | 9.53E-03 |
| ENSG00000178338 | ZNF354B | 0.29 | 0.1 | 3.45E-03 | 9.58E-03 |
| ENSG00000187867 | PALM3 | 0.29 | 0.11 | 3.66E-03 | 1.01E-02 |
| ENSG00000173621 | LRFN4 | 0.29 | 0.1 | 4.03E-03 | 1.10E-02 |
| ENSG00000291091 | SDHAP4 | 0.29 | 0.11 | 4.09E-03 | 1.12E-02 |
| ENSG00000171798 | KNDC1 | 0.29 | 0.1 | 4.30E-03 | 1.17E-02 |
| ENSG00000291072 | CROCCP2 | 0.29 | 0.1 | 4.47E-03 | 1.21E-02 |
| ENSG00000108691 | CCL2 | 0.29 | 0.11 | 4.54E-03 | 1.23E-02 |
| ENSG00000160404 | TOR2A | 0.29 | 0.11 | 5.94E-03 | 1.58E-02 |
| ENSG00000108187 | PBLD | 0.29 | 0.11 | 7.09E-03 | 1.85E-02 |
| ENSG00000131015 | ULBP2 | 0.29 | 0.11 | 7.54E-03 | 1.96E-02 |
| ENSG00000134709 | HOOK1 | 0.29 | 0.12 | 1.03E-02 | 2.59E-02 |
| ENSG00000092871 | RFFL | 0.29 | 0.12 | 1.08E-02 | 2.71E-02 |
| ENSG00000165716 | DIPK1B | 0.29 | 0.13 | 1.57E-02 | 3.79E-02 |
| ENSG00000204628 | RACK1 | 0.28 | 0.03 | 2.26E-19 | 2.04E-18 |
| ENSG00000026025 | VIM | 0.28 | 0.04 | 7.98E-13 | 5.27E-12 |
| ENSG00000108001 | EBF3 | 0.28 | 0.04 | 1.32E-11 | 8.08E-11 |
| ENSG00000142534 | RPS11 | 0.28 | 0.04 | 1.39E-11 | 8.48E-11 |
| ENSG00000132388 | UBE2G1 | 0.28 | 0.04 | 1.09E-10 | 6.28E-10 |
| ENSG00000111481 | COPZ1 | 0.28 | 0.04 | 1.53E-10 | 8.74E-10 |
| ENSG00000114902 | SPCS1 | 0.28 | 0.04 | 2.30E-10 | 1.30E-09 |
| ENSG00000113448 | PDE4D | 0.28 | 0.05 | 4.98E-10 | 2.75E-09 |
| ENSG00000156650 | KAT6B | 0.28 | 0.05 | 1.39E-09 | 7.43E-09 |
| ENSG00000197345 | MRPL21 | 0.28 | 0.05 | 5.43E-09 | 2.77E-08 |
| ENSG00000149639 | SOGA1 | 0.28 | 0.05 | 6.53E-09 | 3.31E-08 |
| ENSG00000237550 | | 0.28 | 0.05 | 6.56E-09 | 3.32E-08 |
| ENSG00000119314 | PTBP3 | 0.28 | 0.05 | 7.15E-09 | 3.61E-08 |
| ENSG00000122884 | P4HA1 | 0.28 | 0.05 | 9.78E-09 | 4.90E-08 |
| ENSG00000182899 | RPL35A | 0.28 | 0.05 | 1.16E-08 | 5.79E-08 |
| ENSG00000137177 | KIF13A | 0.28 | 0.05 | 1.23E-08 | 6.14E-08 |
| ENSG00000074319 | TSG101 | 0.28 | 0.05 | 6.10E-08 | 2.88E-07 |
| ENSG00000112282 | MED23 | 0.28 | 0.05 | 6.63E-08 | 3.12E-07 |
| ENSG00000143164 | DCAF6 | 0.28 | 0.05 | 8.05E-08 | 3.76E-07 |
| ENSG00000100697 | DICER1 | 0.28 | 0.05 | 9.08E-08 | 4.23E-07 |
| ENSG00000163660 | CCNL1 | 0.28 | 0.05 | 1.18E-07 | 5.45E-07 |
| ENSG00000226950 | DANCR | 0.28 | 0.05 | 1.57E-07 | 7.16E-07 |
| ENSG00000170632 | ARMC10 | 0.28 | 0.05 | 1.81E-07 | 8.24E-07 |
| ENSG00000121039 | RDH10 | 0.28 | 0.06 | 2.95E-07 | 1.32E-06 |
| ENSG00000111231 | GPN3 | 0.28 | 0.06 | 3.92E-07 | 1.73E-06 |
| ENSG00000151292 | CSNK1G3 | 0.28 | 0.06 | 4.40E-07 | 1.93E-06 |
| ENSG00000144278 | GALNT13 | 0.28 | 0.06 | 7.54E-07 | 3.24E-06 |
| ENSG00000137947 | GTF2B | 0.28 | 0.06 | 2.00E-06 | 8.25E-06 |
| ENSG00000121741 | ZMYM2 | 0.28 | 0.06 | 2.72E-06 | 1.11E-05 |
| ENSG00000065135 | GNAI3 | 0.28 | 0.06 | 3.05E-06 | 1.24E-05 |
| ENSG00000089159 | PXN | 0.28 | 0.06 | 3.61E-06 | 1.46E-05 |
| ENSG00000197756 | RPL37A | 0.28 | 0.06 | 4.78E-06 | 1.91E-05 |
| ENSG00000104131 | EIF3J | 0.28 | 0.06 | 7.73E-06 | 3.02E-05 |
| ENSG00000116273 | PHF13 | 0.28 | 0.06 | 1.12E-05 | 4.32E-05 |
| ENSG00000152942 | RAD17 | 0.28 | 0.07 | 1.93E-05 | 7.24E-05 |
| ENSG00000168488 | ATXN2L | 0.28 | 0.07 | 2.16E-05 | 8.07E-05 |

| | | | | | |
|---|---|---|---|---|---|
| ENSG00000087157 | PGS1 | 0.28 | 0.07 | 3.59E-05 | 1.31E-04 |
| ENSG00000151092 | NGLY1 | 0.28 | 0.07 | 4.61E-05 | 1.67E-04 |
| ENSG00000078808 | SDF4 | 0.28 | 0.07 | 6.52E-05 | 2.32E-04 |
| ENSG00000116741 | RGS2 | 0.28 | 0.07 | 8.72E-05 | 3.05E-04 |
| ENSG00000177971 | IMP3 | 0.28 | 0.07 | 8.73E-05 | 3.06E-04 |
| ENSG00000198783 | ZNF830 | 0.28 | 0.07 | 9.13E-05 | 3.19E-04 |
| ENSG00000105063 | PPP6R1 | 0.28 | 0.07 | 1.04E-04 | 3.61E-04 |
| ENSG00000089335 | ZNF302 | 0.28 | 0.07 | 1.19E-04 | 4.10E-04 |
| ENSG00000168216 | LMBRD1 | 0.28 | 0.07 | 1.69E-04 | 5.70E-04 |
| ENSG00000135916 | ITM2C | 0.28 | 0.08 | 2.41E-04 | 7.99E-04 |
| ENSG00000115289 | PCGF1 | 0.28 | 0.08 | 2.56E-04 | 8.46E-04 |
| ENSG00000186350 | RXRA | 0.28 | 0.08 | 3.09E-04 | 1.01E-03 |
| ENSG00000113575 | PPP2CA | 0.28 | 0.08 | 3.23E-04 | 1.05E-03 |
| ENSG00000170954 | ZNF415 | 0.28 | 0.08 | 4.09E-04 | 1.32E-03 |
| ENSG00000007866 | TEAD3 | 0.28 | 0.08 | 4.17E-04 | 1.34E-03 |
| ENSG00000170836 | PPM1D | 0.28 | 0.08 | 4.42E-04 | 1.42E-03 |
| ENSG00000161016 | RPL8 | 0.28 | 0.08 | 4.45E-04 | 1.42E-03 |
| ENSG00000160072 | ATAD3B | 0.28 | 0.08 | 4.59E-04 | 1.47E-03 |
| ENSG00000150630 | VEGFC | 0.28 | 0.08 | 4.76E-04 | 1.52E-03 |
| ENSG00000160161 | CILP2 | 0.28 | 0.09 | 1.05E-03 | 3.18E-03 |
| ENSG00000067191 | CACNB1 | 0.28 | 0.09 | 1.22E-03 | 3.66E-03 |
| ENSG00000130540 | SULT4A1 | 0.28 | 0.09 | 1.30E-03 | 3.87E-03 |
| ENSG00000290993 | | 0.28 | 0.09 | 1.32E-03 | 3.94E-03 |
| ENSG00000198841 | KTI12 | 0.28 | 0.09 | 1.99E-03 | 5.78E-03 |
| ENSG00000113108 | APBB3 | 0.28 | 0.09 | 2.29E-03 | 6.56E-03 |
| ENSG00000118292 | C1orf54 | 0.28 | 0.1 | 2.53E-03 | 7.21E-03 |
| ENSG00000077463 | SIRT6 | 0.28 | 0.1 | 4.79E-03 | 1.29E-02 |
| ENSG00000159079 | CFAP298 | 0.28 | 0.1 | 4.81E-03 | 1.30E-02 |
| ENSG00000291074 | CCDC144NL | 0.28 | 0.11 | 6.17E-03 | 1.63E-02 |
| ENSG00000169715 | MT1E | 0.28 | 0.11 | 6.22E-03 | 1.64E-02 |
| ENSG00000188542 | DUSP28 | 0.28 | 0.11 | 6.91E-03 | 1.81E-02 |
| ENSG00000197124 | ZNF682 | 0.28 | 0.11 | 8.68E-03 | 2.23E-02 |
| ENSG00000176845 | METRNL | 0.28 | 0.11 | 9.09E-03 | 2.32E-02 |
| ENSG00000261221 | ZNF865 | 0.28 | 0.12 | 1.25E-02 | 3.10E-02 |
| ENSG00000240771 | ARHGEF25 | 0.28 | 0.12 | 1.40E-02 | 3.43E-02 |
| ENSG00000105499 | PLA2G4C | 0.28 | 0.12 | 1.67E-02 | 4.00E-02 |
| ENSG00000266173 | STRADA | 0.28 | 0.13 | 1.75E-02 | 4.17E-02 |
| ENSG00000236859 | NIFK-AS1 | 0.28 | 0.13 | 2.05E-02 | 4.82E-02 |
| ENSG00000163513 | TGFBR2 | 0.27 | 0.04 | 1.05E-14 | 7.67E-14 |
| ENSG00000029364 | SLC39A9 | 0.27 | 0.04 | 5.32E-13 | 3.55E-12 |
| ENSG00000148358 | GPR107 | 0.27 | 0.04 | 1.51E-11 | 9.20E-11 |
| ENSG00000167978 | SRRM2 | 0.27 | 0.04 | 1.38E-10 | 7.95E-10 |
| ENSG00000126267 | COX6B1 | 0.27 | 0.04 | 1.52E-10 | 8.70E-10 |
| ENSG00000277791 | PSMB3 | 0.27 | 0.04 | 1.84E-10 | 1.05E-09 |
| ENSG00000124226 | RNF114 | 0.27 | 0.04 | 4.00E-10 | 2.22E-09 |
| ENSG00000020577 | SAMD4A | 0.27 | 0.04 | 5.32E-10 | 2.93E-09 |
| ENSG00000100528 | CNIH1 | 0.27 | 0.04 | 1.34E-09 | 7.16E-09 |
| ENSG00000177119 | ANO6 | 0.27 | 0.04 | 1.46E-09 | 7.81E-09 |
| ENSG00000105372 | RPS19 | 0.27 | 0.05 | 2.99E-09 | 1.56E-08 |
| ENSG00000063322 | MED29 | 0.27 | 0.05 | 6.02E-09 | 3.06E-08 |

| | | | | | |
|---|---|---|---|---|---|
| ENSG00000143727 | ACP1 | 0.27 | 0.05 | 1.47E-08 | 7.25E-08 |
| ENSG00000139977 | NAA30 | 0.27 | 0.05 | 3.47E-08 | 1.66E-07 |
| ENSG00000185624 | P4HB | 0.27 | 0.05 | 5.78E-08 | 2.73E-07 |
| ENSG00000135334 | AKIRIN2 | 0.27 | 0.05 | 1.01E-07 | 4.67E-07 |
| ENSG00000144468 | RHBDD1 | 0.27 | 0.05 | 1.87E-07 | 8.51E-07 |
| ENSG00000163608 | NEPRO | 0.27 | 0.05 | 2.43E-07 | 1.09E-06 |
| ENSG00000175354 | PTPN2 | 0.27 | 0.05 | 4.33E-07 | 1.91E-06 |
| ENSG00000234616 | JRK | 0.27 | 0.06 | 6.23E-07 | 2.70E-06 |
| ENSG00000139163 | ETNK1 | 0.27 | 0.05 | 6.39E-07 | 2.77E-06 |
| ENSG00000129473 | BCL2L2 | 0.27 | 0.06 | 6.69E-07 | 2.89E-06 |
| ENSG00000168653 | NDUFS5 | 0.27 | 0.05 | 7.21E-07 | 3.11E-06 |
| ENSG00000067445 | TRO | 0.27 | 0.06 | 7.61E-07 | 3.27E-06 |
| ENSG00000161021 | MAML1 | 0.27 | 0.06 | 8.31E-07 | 3.56E-06 |
| ENSG00000116251 | RPL22 | 0.27 | 0.06 | 1.35E-06 | 5.69E-06 |
| ENSG00000185420 | SMYD3 | 0.27 | 0.06 | 1.46E-06 | 6.11E-06 |
| ENSG00000148158 | SNX30 | 0.27 | 0.06 | 1.47E-06 | 6.16E-06 |
| ENSG00000131966 | ACTR10 | 0.27 | 0.06 | 2.15E-06 | 8.86E-06 |
| ENSG00000100567 | PSMA3 | 0.27 | 0.06 | 2.39E-06 | 9.78E-06 |
| ENSG00000175061 | SNHG29 | 0.27 | 0.06 | 2.73E-06 | 1.11E-05 |
| ENSG00000173039 | RELA | 0.27 | 0.06 | 1.15E-05 | 4.43E-05 |
| ENSG00000107938 | EDRF1 | 0.27 | 0.06 | 1.23E-05 | 4.73E-05 |
| ENSG00000117155 | SSX2IP | 0.27 | 0.07 | 1.95E-05 | 7.31E-05 |
| ENSG00000087087 | SRRT | 0.27 | 0.06 | 2.30E-05 | 8.58E-05 |
| ENSG00000075975 | MKRN2 | 0.27 | 0.07 | 2.39E-05 | 8.87E-05 |
| ENSG00000066455 | GOLGA5 | 0.27 | 0.07 | 3.51E-05 | 1.28E-04 |
| ENSG00000114956 | DGUOK | 0.27 | 0.07 | 3.81E-05 | 1.39E-04 |
| ENSG00000081307 | UBA5 | 0.27 | 0.07 | 4.94E-05 | 1.78E-04 |
| ENSG00000145354 | CISD2 | 0.27 | 0.07 | 5.13E-05 | 1.84E-04 |
| ENSG00000152944 | MED21 | 0.27 | 0.07 | 6.31E-05 | 2.25E-04 |
| ENSG00000079805 | DNM2 | 0.27 | 0.07 | 7.24E-05 | 2.56E-04 |
| ENSG00000196961 | AP2A1 | 0.27 | 0.07 | 8.21E-05 | 2.88E-04 |
| ENSG00000112200 | ZNF451 | 0.27 | 0.07 | 2.10E-04 | 7.03E-04 |
| ENSG00000168807 | SNTB2 | 0.27 | 0.07 | 2.75E-04 | 9.06E-04 |
| ENSG00000100979 | PLTP | 0.27 | 0.08 | 3.56E-04 | 1.15E-03 |
| ENSG00000148814 | LRRC27 | 0.27 | 0.08 | 5.89E-04 | 1.86E-03 |
| ENSG00000128594 | LRRC4 | 0.27 | 0.08 | 8.25E-04 | 2.54E-03 |
| ENSG00000074181 | NOTCH3 | 0.27 | 0.09 | 1.25E-03 | 3.73E-03 |
| ENSG00000244398 | RPL36AP37 | 0.27 | 0.09 | 1.53E-03 | 4.50E-03 |
| ENSG00000132740 | IGHMBP2 | 0.27 | 0.09 | 1.62E-03 | 4.76E-03 |
| ENSG00000064932 | SBNO2 | 0.27 | 0.09 | 1.91E-03 | 5.55E-03 |
| ENSG00000224914 | LINC00863 | 0.27 | 0.09 | 2.02E-03 | 5.86E-03 |
| ENSG00000213462 | ERV3-1 | 0.27 | 0.09 | 2.64E-03 | 7.51E-03 |
| ENSG00000108352 | RAPGEFL1 | 0.27 | 0.1 | 3.40E-03 | 9.43E-03 |
| ENSG00000100036 | SLC35E4 | 0.27 | 0.09 | 3.40E-03 | 9.44E-03 |
| ENSG00000168994 | PXDC1 | 0.27 | 0.1 | 4.71E-03 | 1.27E-02 |
| ENSG00000177548 | RABEP2 | 0.27 | 0.1 | 4.84E-03 | 1.31E-02 |
| ENSG00000147119 | CHST7 | 0.27 | 0.1 | 4.86E-03 | 1.31E-02 |
| ENSG00000124444 | ZNF576 | 0.27 | 0.1 | 6.24E-03 | 1.65E-02 |
| ENSG00000071889 | FAM3A | 0.27 | 0.1 | 8.34E-03 | 2.15E-02 |
| ENSG00000168297 | PXK | 0.27 | 0.11 | 9.55E-03 | 2.42E-02 |

| | | | | | |
|---|---|---|---|---|---|
| ENSG00000172977 | KAT5 | 0.27 | 0.11 | 1.07E-02 | 2.69E-02 |
| ENSG00000204650 | LINC02210 | 0.27 | 0.11 | 1.41E-02 | 3.43E-02 |
| ENSG00000135587 | SMPD2 | 0.27 | 0.12 | 1.75E-02 | 4.17E-02 |
| ENSG00000136448 | NMT1 | 0.26 | 0.04 | 1.95E-13 | 1.34E-12 |
| ENSG00000076641 | PAG1 | 0.26 | 0.04 | 2.49E-13 | 1.69E-12 |
| ENSG00000197694 | SPTAN1 | 0.26 | 0.04 | 2.63E-13 | 1.79E-12 |
| ENSG00000100138 | SNU13 | 0.26 | 0.04 | 6.74E-12 | 4.19E-11 |
| ENSG00000196367 | TRRAP | 0.26 | 0.04 | 3.58E-11 | 2.13E-10 |
| ENSG00000162736 | NCSTN | 0.26 | 0.04 | 2.41E-10 | 1.36E-09 |
| ENSG00000112697 | TMEM30A | 0.26 | 0.04 | 2.83E-10 | 1.59E-09 |
| ENSG00000084754 | HADHA | 0.26 | 0.04 | 4.45E-10 | 2.46E-09 |
| ENSG00000008018 | PSMB1 | 0.26 | 0.04 | 1.45E-09 | 7.72E-09 |
| ENSG00000179051 | RCC2 | 0.26 | 0.05 | 7.46E-09 | 3.77E-08 |
| ENSG00000187239 | FNBP1 | 0.26 | 0.05 | 1.90E-08 | 9.31E-08 |
| ENSG00000126261 | UBA2 | 0.26 | 0.05 | 6.88E-08 | 3.23E-07 |
| ENSG00000089234 | BRAP | 0.26 | 0.05 | 1.07E-07 | 4.96E-07 |
| ENSG00000125868 | DSTN | 0.26 | 0.05 | 1.35E-07 | 6.20E-07 |
| ENSG00000145901 | TNIP1 | 0.26 | 0.05 | 1.38E-07 | 6.33E-07 |
| ENSG00000108100 | CCNY | 0.26 | 0.05 | 1.66E-07 | 7.54E-07 |
| ENSG00000124177 | CHD6 | 0.26 | 0.05 | 3.53E-07 | 1.57E-06 |
| ENSG00000117362 | APH1A | 0.26 | 0.05 | 4.26E-07 | 1.88E-06 |
| ENSG00000121067 | SPOP | 0.26 | 0.05 | 4.90E-07 | 2.15E-06 |
| ENSG00000163558 | PRKCI | 0.26 | 0.05 | 5.52E-07 | 2.41E-06 |
| ENSG00000157800 | SLC37A3 | 0.26 | 0.05 | 5.65E-07 | 2.46E-06 |
| ENSG00000172939 | OXSR1 | 0.26 | 0.05 | 5.95E-07 | 2.59E-06 |
| ENSG00000149262 | INTS4 | 0.26 | 0.05 | 6.49E-07 | 2.81E-06 |
| ENSG00000129566 | TEP1 | 0.26 | 0.05 | 7.26E-07 | 3.13E-06 |
| ENSG00000078747 | ITCH | 0.26 | 0.05 | 1.41E-06 | 5.90E-06 |
| ENSG00000129625 | REEP5 | 0.26 | 0.06 | 1.66E-06 | 6.92E-06 |
| ENSG00000088854 | DNAAF9 | 0.26 | 0.06 | 1.76E-06 | 7.27E-06 |
| ENSG00000005075 | POLR2J | 0.26 | 0.06 | 2.15E-06 | 8.83E-06 |
| ENSG00000136950 | ARPC5L | 0.26 | 0.06 | 2.46E-06 | 1.01E-05 |
| ENSG00000129636 | ITFG1 | 0.26 | 0.06 | 4.02E-06 | 1.62E-05 |
| ENSG00000112851 | ERBIN | 0.26 | 0.06 | 7.31E-06 | 2.86E-05 |
| ENSG00000004779 | NDUFAB1 | 0.26 | 0.06 | 8.12E-06 | 3.17E-05 |
| ENSG00000105669 | COPE | 0.26 | 0.06 | 1.48E-05 | 5.62E-05 |
| ENSG00000206503 | HLA-A | 0.26 | 0.06 | 1.89E-05 | 7.10E-05 |
| ENSG00000132581 | SDF2 | 0.26 | 0.06 | 2.56E-05 | 9.47E-05 |
| ENSG00000198756 | COLGALT2 | 0.26 | 0.06 | 2.70E-05 | 9.97E-05 |
| ENSG00000126602 | TRAP1 | 0.26 | 0.06 | 2.94E-05 | 1.08E-04 |
| ENSG00000087338 | GMCL1 | 0.26 | 0.06 | 3.43E-05 | 1.25E-04 |
| ENSG00000136942 | RPL35 | 0.26 | 0.06 | 3.52E-05 | 1.29E-04 |
| ENSG00000108239 | TBC1D12 | 0.26 | 0.07 | 6.27E-05 | 2.24E-04 |
| ENSG00000186020 | ZNF529 | 0.26 | 0.07 | 6.63E-05 | 2.36E-04 |
| ENSG00000168872 | DDX19A | 0.26 | 0.07 | 7.78E-05 | 2.74E-04 |
| ENSG00000075945 | KIFAP3 | 0.26 | 0.07 | 1.05E-04 | 3.63E-04 |
| ENSG00000104529 | EEF1D | 0.26 | 0.07 | 1.11E-04 | 3.85E-04 |
| ENSG00000105185 | PDCD5 | 0.26 | 0.07 | 1.22E-04 | 4.18E-04 |
| ENSG00000134758 | RNF138 | 0.26 | 0.07 | 1.41E-04 | 4.81E-04 |
| ENSG00000167114 | SLC27A4 | 0.26 | 0.07 | 1.84E-04 | 6.21E-04 |

| | | | | | |
|---|---|---|---|---|---|
| ENSG00000158805 | ZNF276 | 0.26 | 0.07 | 2.85E-04 | 9.36E-04 |
| ENSG00000130382 | MLLT1 | 0.26 | 0.07 | 2.95E-04 | 9.67E-04 |
| ENSG00000263002 | ZNF234 | 0.26 | 0.07 | 3.18E-04 | 1.04E-03 |
| ENSG00000166169 | POLL | 0.26 | 0.07 | 3.32E-04 | 1.08E-03 |
| ENSG00000127922 | SEM1 | 0.26 | 0.07 | 3.55E-04 | 1.15E-03 |
| ENSG00000105245 | NUMBL | 0.26 | 0.08 | 3.55E-04 | 1.15E-03 |
| ENSG00000159263 | SIM2 | 0.26 | 0.08 | 3.85E-04 | 1.24E-03 |
| ENSG00000101246 | ARFRP1 | 0.26 | 0.08 | 4.35E-04 | 1.40E-03 |
| ENSG00000122507 | BBS9 | 0.26 | 0.08 | 5.50E-04 | 1.74E-03 |
| ENSG00000291080 | ZNF252P | 0.26 | 0.08 | 9.09E-04 | 2.78E-03 |
| ENSG00000159069 | FBXW5 | 0.26 | 0.08 | 9.69E-04 | 2.95E-03 |
| ENSG00000105072 | C19orf44 | 0.26 | 0.08 | 1.06E-03 | 3.20E-03 |
| ENSG00000166813 | KIF7 | 0.26 | 0.08 | 1.07E-03 | 3.23E-03 |
| ENSG00000105767 | CADM4 | 0.26 | 0.08 | 1.19E-03 | 3.58E-03 |
| ENSG00000136932 | TRMO | 0.26 | 0.08 | 1.21E-03 | 3.63E-03 |
| ENSG00000137806 | NDUFAF1 | 0.26 | 0.08 | 1.39E-03 | 4.14E-03 |
| ENSG00000137764 | MAP2K5 | 0.26 | 0.08 | 1.40E-03 | 4.17E-03 |
| ENSG00000156482 | RPL30 | 0.26 | 0.08 | 1.58E-03 | 4.66E-03 |
| ENSG00000141552 | ANAPC11 | 0.26 | 0.09 | 1.99E-03 | 5.75E-03 |
| ENSG00000291006 | | 0.26 | 0.09 | 2.43E-03 | 6.93E-03 |
| ENSG00000152904 | GGPS1 | 0.26 | 0.09 | 3.09E-03 | 8.66E-03 |
| ENSG00000233184 | DPH5-DT | 0.26 | 0.09 | 3.12E-03 | 8.75E-03 |
| ENSG00000144445 | KANSL1L | 0.26 | 0.09 | 3.32E-03 | 9.24E-03 |
| ENSG00000099889 | ARVCF | 0.26 | 0.09 | 3.88E-03 | 1.07E-02 |
| ENSG00000115468 | EFHD1 | 0.26 | 0.09 | 4.03E-03 | 1.10E-02 |
| ENSG00000257621 | PSMA3-AS1 | 0.26 | 0.1 | 5.00E-03 | 1.34E-02 |
| ENSG00000126768 | TIMM17B | 0.26 | 0.1 | 5.20E-03 | 1.39E-02 |
| ENSG00000198089 | SFI1 | 0.26 | 0.1 | 6.14E-03 | 1.62E-02 |
| ENSG00000062370 | ZNF112 | 0.26 | 0.1 | 7.04E-03 | 1.84E-02 |
| ENSG00000168778 | TCTN2 | 0.26 | 0.1 | 7.65E-03 | 1.99E-02 |
| ENSG00000181038 | METTL23 | 0.26 | 0.1 | 8.64E-03 | 2.22E-02 |
| ENSG00000104960 | PTOV1 | 0.26 | 0.11 | 1.11E-02 | 2.77E-02 |
| ENSG00000132950 | ZMYM5 | 0.26 | 0.11 | 1.21E-02 | 3.00E-02 |
| ENSG00000148362 | PAXX | 0.26 | 0.11 | 1.31E-02 | 3.21E-02 |
| ENSG00000088970 | KIZ | 0.26 | 0.11 | 1.44E-02 | 3.50E-02 |
| ENSG00000188707 | ZBED10P | 0.26 | 0.12 | 2.05E-02 | 4.82E-02 |
| ENSG00000104341 | LAPTM4B | 0.25 | 0.03 | 2.82E-13 | 1.91E-12 |
| ENSG00000143337 | TOR1AIP1 | 0.25 | 0.04 | 6.33E-12 | 3.95E-11 |
| ENSG00000124120 | TTPAL | 0.25 | 0.04 | 1.02E-10 | 5.92E-10 |
| ENSG00000137076 | TLN1 | 0.25 | 0.04 | 9.03E-10 | 4.88E-09 |
| ENSG00000145495 | MARCHF6 | 0.25 | 0.04 | 9.35E-10 | 5.05E-09 |
| ENSG00000054965 | FAM168A | 0.25 | 0.04 | 2.52E-09 | 1.32E-08 |
| ENSG00000197930 | ERO1A | 0.25 | 0.04 | 3.05E-09 | 1.59E-08 |
| ENSG00000108010 | GLRX3 | 0.25 | 0.04 | 3.63E-09 | 1.88E-08 |
| ENSG00000153914 | SREK1 | 0.25 | 0.04 | 6.43E-09 | 3.26E-08 |
| ENSG00000163297 | ANTXR2 | 0.25 | 0.04 | 8.13E-09 | 4.09E-08 |
| ENSG00000134508 | CABLES1 | 0.25 | 0.05 | 2.34E-08 | 1.14E-07 |
| ENSG00000101161 | PRPF6 | 0.25 | 0.05 | 3.35E-08 | 1.61E-07 |
| ENSG00000166860 | ZBTB39 | 0.25 | 0.05 | 3.50E-08 | 1.68E-07 |
| ENSG00000063978 | RNF4 | 0.25 | 0.05 | 1.14E-07 | 5.26E-07 |

| | | | | | |
|---|---|---|---|---|---|
| ENSG00000089597 | GANAB | 0.25 | 0.05 | 1.34E-07 | 6.15E-07 |
| ENSG00000136937 | NCBP1 | 0.25 | 0.05 | 1.45E-07 | 6.64E-07 |
| ENSG00000121644 | DESI2 | 0.25 | 0.05 | 2.81E-07 | 1.26E-06 |
| ENSG00000175582 | RAB6A | 0.25 | 0.05 | 4.55E-07 | 2.00E-06 |
| ENSG00000176444 | CLK2 | 0.25 | 0.05 | 6.86E-07 | 2.96E-06 |
| ENSG00000101365 | IDH3B | 0.25 | 0.05 | 1.03E-06 | 4.37E-06 |
| ENSG00000170525 | PFKFB3 | 0.25 | 0.05 | 1.89E-06 | 7.81E-06 |
| ENSG00000164151 | ICE1 | 0.25 | 0.05 | 2.23E-06 | 9.16E-06 |
| ENSG00000181222 | POLR2A | 0.25 | 0.05 | 2.29E-06 | 9.39E-06 |
| ENSG00000120925 | RNF170 | 0.25 | 0.05 | 2.76E-06 | 1.13E-05 |
| ENSG00000184402 | SS18L1 | 0.25 | 0.05 | 2.82E-06 | 1.15E-05 |
| ENSG00000034510 | TMSB10 | 0.25 | 0.06 | 4.68E-06 | 1.87E-05 |
| ENSG00000179195 | ZNF664 | 0.25 | 0.06 | 5.73E-06 | 2.27E-05 |
| ENSG00000148925 | BTBD10 | 0.25 | 0.06 | 6.95E-06 | 2.73E-05 |
| ENSG00000169902 | TPST1 | 0.25 | 0.06 | 8.48E-06 | 3.30E-05 |
| ENSG00000141627 | DYM | 0.25 | 0.06 | 1.22E-05 | 4.68E-05 |
| ENSG00000117000 | RLF | 0.25 | 0.06 | 1.42E-05 | 5.40E-05 |
| ENSG00000150403 | TMCO3 | 0.25 | 0.06 | 1.59E-05 | 6.01E-05 |
| ENSG00000040199 | PHLPP2 | 0.25 | 0.06 | 1.90E-05 | 7.15E-05 |
| ENSG00000100592 | DAAM1 | 0.25 | 0.06 | 1.95E-05 | 7.33E-05 |
| ENSG00000183688 | RFLNB | 0.25 | 0.06 | 4.34E-05 | 1.57E-04 |
| ENSG00000173209 | AHSA2P | 0.25 | 0.06 | 7.28E-05 | 2.58E-04 |
| ENSG00000104064 | GABPB1 | 0.25 | 0.07 | 1.19E-04 | 4.10E-04 |
| ENSG00000082212 | ME2 | 0.25 | 0.07 | 1.24E-04 | 4.26E-04 |
| ENSG00000139168 | ZCRB1 | 0.25 | 0.07 | 1.45E-04 | 4.96E-04 |
| ENSG00000113441 | LNPEP | 0.25 | 0.07 | 1.51E-04 | 5.13E-04 |
| ENSG00000168566 | SNRNP48 | 0.25 | 0.07 | 1.92E-04 | 6.46E-04 |
| ENSG00000143156 | NME7 | 0.25 | 0.07 | 2.27E-04 | 7.55E-04 |
| ENSG00000102858 | MGRN1 | 0.25 | 0.07 | 2.37E-04 | 7.85E-04 |
| ENSG00000128581 | IFT22 | 0.25 | 0.07 | 2.64E-04 | 8.72E-04 |
| ENSG00000111058 | ACSS3 | 0.25 | 0.07 | 3.29E-04 | 1.07E-03 |
| ENSG00000173141 | MRPL57 | 0.25 | 0.07 | 3.74E-04 | 1.21E-03 |
| ENSG00000103496 | STX4 | 0.25 | 0.07 | 4.33E-04 | 1.39E-03 |
| ENSG00000186310 | NAP1L3 | 0.25 | 0.08 | 5.97E-04 | 1.88E-03 |
| ENSG00000100632 | ERH | 0.25 | 0.08 | 6.35E-04 | 1.99E-03 |
| ENSG00000131368 | MRPS25 | 0.25 | 0.07 | 6.70E-04 | 2.10E-03 |
| ENSG00000179943 | FIZ1 | 0.25 | 0.08 | 8.08E-04 | 2.49E-03 |
| ENSG00000084092 | NOA1 | 0.25 | 0.08 | 8.48E-04 | 2.61E-03 |
| ENSG00000143158 | MPC2 | 0.25 | 0.08 | 9.09E-04 | 2.78E-03 |
| ENSG00000023171 | GRAMD1B | 0.25 | 0.08 | 9.73E-04 | 2.96E-03 |
| ENSG00000105656 | ELL | 0.25 | 0.08 | 1.11E-03 | 3.35E-03 |
| ENSG00000005812 | FBXL3 | 0.25 | 0.08 | 1.24E-03 | 3.72E-03 |
| ENSG00000137343 | ATAT1 | 0.25 | 0.08 | 1.39E-03 | 4.14E-03 |
| ENSG00000183011 | NAA38 | 0.25 | 0.08 | 1.39E-03 | 4.14E-03 |
| ENSG00000083937 | CHMP2B | 0.25 | 0.08 | 1.84E-03 | 5.35E-03 |
| ENSG00000104324 | CPQ | 0.25 | 0.08 | 1.84E-03 | 5.37E-03 |
| ENSG00000053254 | FOXN3 | 0.25 | 0.08 | 2.21E-03 | 6.36E-03 |
| ENSG00000114388 | NPRL2 | 0.25 | 0.09 | 2.58E-03 | 7.34E-03 |
| ENSG00000066697 | MSANTD3 | 0.25 | 0.09 | 2.70E-03 | 7.65E-03 |
| ENSG00000131127 | ZNF141 | 0.25 | 0.09 | 2.97E-03 | 8.35E-03 |

| | | | | | |
|---|---|---|---|---|---|
| ENSG00000137038 | DMAC1 | 0.25 | 0.09 | 3.07E-03 | 8.61E-03 |
| ENSG00000196878 | LAMB3 | 0.25 | 0.09 | 3.27E-03 | 9.12E-03 |
| ENSG00000005238 | ATOSB | 0.25 | 0.09 | 3.40E-03 | 9.44E-03 |
| ENSG00000181315 | ZNF322 | 0.25 | 0.09 | 3.51E-03 | 9.73E-03 |
| ENSG00000203778 | FAM229B | 0.25 | 0.09 | 4.08E-03 | 1.12E-02 |
| ENSG00000169750 | RAC3 | 0.25 | 0.09 | 5.89E-03 | 1.56E-02 |
| ENSG00000151338 | MIPOL1 | 0.25 | 0.09 | 6.14E-03 | 1.62E-02 |
| ENSG00000143303 | METTL25B | 0.25 | 0.1 | 8.58E-03 | 2.20E-02 |
| ENSG00000163521 | GLB1L | 0.25 | 0.1 | 9.33E-03 | 2.37E-02 |
| ENSG00000014914 | MTMR11 | 0.25 | 0.1 | 1.01E-02 | 2.54E-02 |
| ENSG00000105792 | CFAP69 | 0.25 | 0.1 | 1.17E-02 | 2.91E-02 |
| ENSG00000221843 | C2orf16 | 0.25 | 0.11 | 1.23E-02 | 3.05E-02 |
| ENSG00000157353 | FCSK | 0.25 | 0.11 | 1.28E-02 | 3.15E-02 |
| ENSG00000106701 | FSD1L | 0.25 | 0.1 | 1.30E-02 | 3.21E-02 |
| ENSG00000124243 | BCAS4 | 0.25 | 0.11 | 1.35E-02 | 3.31E-02 |
| ENSG00000110011 | DNAJC4 | 0.25 | 0.11 | 1.41E-02 | 3.45E-02 |
| ENSG00000177463 | NR2C2 | 0.25 | 0.11 | 1.79E-02 | 4.25E-02 |
| ENSG00000117151 | CTBS | 0.25 | 0.11 | 2.03E-02 | 4.78E-02 |
| ENSG00000155506 | LARP1 | 0.24 | 0.03 | 2.18E-12 | 1.41E-11 |
| ENSG00000181789 | COPG1 | 0.24 | 0.04 | 5.06E-12 | 3.18E-11 |
| ENSG00000145860 | RNF145 | 0.24 | 0.04 | 2.03E-11 | 1.23E-10 |
| ENSG00000054118 | THRAP3 | 0.24 | 0.04 | 3.58E-11 | 2.13E-10 |
| ENSG00000123066 | MED13L | 0.24 | 0.04 | 1.56E-10 | 8.94E-10 |
| ENSG00000166135 | HIF1AN | 0.24 | 0.04 | 2.99E-10 | 1.68E-09 |
| ENSG00000135269 | TES | 0.24 | 0.04 | 4.50E-09 | 2.30E-08 |
| ENSG00000140157 | NIPA2 | 0.24 | 0.04 | 5.90E-08 | 2.78E-07 |
| ENSG00000083845 | RPS5 | 0.24 | 0.05 | 1.27E-07 | 5.82E-07 |
| ENSG00000189067 | LITAF | 0.24 | 0.05 | 1.57E-07 | 7.17E-07 |
| ENSG00000146872 | TLK2 | 0.24 | 0.05 | 2.52E-07 | 1.13E-06 |
| ENSG00000116539 | ASH1L | 0.24 | 0.05 | 4.00E-07 | 1.77E-06 |
| ENSG00000187498 | COL4A1 | 0.24 | 0.05 | 5.98E-07 | 2.60E-06 |
| ENSG00000196663 | TECPR2 | 0.24 | 0.05 | 6.04E-07 | 2.62E-06 |
| ENSG00000135390 | ATP5MC2 | 0.24 | 0.05 | 7.36E-07 | 3.17E-06 |
| ENSG00000130338 | TULP4 | 0.24 | 0.05 | 8.03E-07 | 3.44E-06 |
| ENSG00000122406 | RPL5 | 0.24 | 0.05 | 8.80E-07 | 3.75E-06 |
| ENSG00000149100 | EIF3M | 0.24 | 0.05 | 9.60E-07 | 4.08E-06 |
| ENSG00000174227 | PIGG | 0.24 | 0.05 | 1.10E-06 | 4.65E-06 |
| ENSG00000074201 | CLNS1A | 0.24 | 0.05 | 1.40E-06 | 5.86E-06 |
| ENSG00000104660 | LEPROTL1 | 0.24 | 0.05 | 1.83E-06 | 7.56E-06 |
| ENSG00000170145 | SIK2 | 0.24 | 0.05 | 1.91E-06 | 7.88E-06 |
| ENSG00000070961 | ATP2B1 | 0.24 | 0.05 | 2.00E-06 | 8.26E-06 |
| ENSG00000198742 | SMURF1 | 0.24 | 0.05 | 2.11E-06 | 8.70E-06 |
| ENSG00000177663 | IL17RA | 0.24 | 0.05 | 2.56E-06 | 1.05E-05 |
| ENSG00000124486 | USP9X | 0.24 | 0.05 | 2.85E-06 | 1.16E-05 |
| ENSG00000162613 | FUBP1 | 0.24 | 0.05 | 3.33E-06 | 1.35E-05 |
| ENSG00000196937 | FAM3C | 0.24 | 0.05 | 6.13E-06 | 2.42E-05 |
| ENSG00000163346 | PBXIP1 | 0.24 | 0.05 | 6.19E-06 | 2.44E-05 |
| ENSG00000184465 | WDR27 | 0.24 | 0.05 | 6.95E-06 | 2.73E-05 |
| ENSG00000135999 | EPC2 | 0.24 | 0.06 | 7.96E-06 | 3.11E-05 |
| ENSG00000081154 | PCNP | 0.24 | 0.05 | 8.72E-06 | 3.39E-05 |

| | | | | | |
|---|---|---|---|---|---|
| ENSG00000171603 | CLSTN1 | 0.24 | 0.06 | 1.49E-05 | 5.67E-05 |
| ENSG00000176095 | IP6K1 | 0.24 | 0.06 | 1.62E-05 | 6.13E-05 |
| ENSG00000116489 | CAPZA1 | 0.24 | 0.06 | 1.95E-05 | 7.31E-05 |
| ENSG00000141646 | SMAD4 | 0.24 | 0.06 | 2.05E-05 | 7.68E-05 |
| ENSG00000011243 | AKAP8L | 0.24 | 0.06 | 2.52E-05 | 9.36E-05 |
| ENSG00000160991 | ORAI2 | 0.24 | 0.06 | 3.32E-05 | 1.21E-04 |
| ENSG00000129315 | CCNT1 | 0.24 | 0.06 | 3.41E-05 | 1.25E-04 |
| ENSG00000205189 | ZBTB10 | 0.24 | 0.06 | 4.14E-05 | 1.50E-04 |
| ENSG00000155906 | RMND1 | 0.24 | 0.06 | 4.16E-05 | 1.51E-04 |
| ENSG00000104142 | VPS18 | 0.24 | 0.06 | 4.21E-05 | 1.53E-04 |
| ENSG00000069956 | MAPK6 | 0.24 | 0.06 | 4.30E-05 | 1.56E-04 |
| ENSG00000169905 | TOR1AIP2 | 0.24 | 0.06 | 4.41E-05 | 1.60E-04 |
| ENSG00000126226 | PCID2 | 0.24 | 0.06 | 4.44E-05 | 1.60E-04 |
| ENSG00000141378 | PTRH2 | 0.24 | 0.06 | 4.54E-05 | 1.64E-04 |
| ENSG00000020129 | NCDN | 0.24 | 0.06 | 4.65E-05 | 1.68E-04 |
| ENSG00000092330 | TINF2 | 0.24 | 0.06 | 5.15E-05 | 1.85E-04 |
| ENSG00000182636 | NDN | 0.24 | 0.06 | 5.22E-05 | 1.87E-04 |
| ENSG00000176105 | YES1 | 0.24 | 0.06 | 6.03E-05 | 2.15E-04 |
| ENSG00000106028 | SSBP1 | 0.24 | 0.06 | 6.56E-05 | 2.33E-04 |
| ENSG00000171858 | RPS21 | 0.24 | 0.06 | 7.35E-05 | 2.60E-04 |
| ENSG00000135049 | AGTPBP1 | 0.24 | 0.06 | 7.82E-05 | 2.75E-04 |
| ENSG00000156265 | MAP3K7CL | 0.24 | 0.06 | 1.06E-04 | 3.69E-04 |
| ENSG00000115318 | LOXL3 | 0.24 | 0.06 | 1.51E-04 | 5.13E-04 |
| ENSG00000137269 | LRRC1 | 0.24 | 0.07 | 1.53E-04 | 5.20E-04 |
| ENSG00000142197 | DOP1B | 0.24 | 0.06 | 1.80E-04 | 6.08E-04 |
| ENSG00000168884 | TNIP2 | 0.24 | 0.07 | 1.90E-04 | 6.39E-04 |
| ENSG00000104671 | DCTN6 | 0.24 | 0.07 | 2.02E-04 | 6.78E-04 |
| ENSG00000076053 | RBM7 | 0.24 | 0.07 | 2.81E-04 | 9.23E-04 |
| ENSG00000115514 | TXNDC9 | 0.24 | 0.07 | 3.54E-04 | 1.15E-03 |
| ENSG00000164050 | PLXNB1 | 0.24 | 0.07 | 5.28E-04 | 1.68E-03 |
| ENSG00000162413 | KLHL21 | 0.24 | 0.07 | 5.93E-04 | 1.87E-03 |
| ENSG00000142655 | PEX14 | 0.24 | 0.07 | 6.20E-04 | 1.95E-03 |
| ENSG00000185716 | MOSMO | 0.24 | 0.07 | 7.02E-04 | 2.19E-03 |
| ENSG00000177105 | RHOG | 0.24 | 0.07 | 7.19E-04 | 2.24E-03 |
| ENSG00000130254 | SAFB2 | 0.24 | 0.07 | 7.27E-04 | 2.26E-03 |
| ENSG00000105677 | TMEM147 | 0.24 | 0.07 | 7.91E-04 | 2.44E-03 |
| ENSG00000141098 | GFOD2 | 0.24 | 0.07 | 8.43E-04 | 2.59E-03 |
| ENSG00000103174 | NAGPA | 0.24 | 0.07 | 9.22E-04 | 2.82E-03 |
| ENSG00000144579 | CTDSP1 | 0.24 | 0.07 | 1.10E-03 | 3.32E-03 |
| ENSG00000107223 | EDF1 | 0.24 | 0.07 | 1.17E-03 | 3.53E-03 |
| ENSG00000133112 | TPT1 | 0.24 | 0.08 | 1.33E-03 | 3.97E-03 |
| ENSG00000182903 | ZNF721 | 0.24 | 0.08 | 1.72E-03 | 5.02E-03 |
| ENSG00000167657 | DAPK3 | 0.24 | 0.08 | 2.03E-03 | 5.88E-03 |
| ENSG00000204673 | AKT1S1 | 0.24 | 0.08 | 2.43E-03 | 6.94E-03 |
| ENSG00000184110 | EIF3C | 0.24 | 0.08 | 2.56E-03 | 7.29E-03 |
| ENSG00000068971 | PPP2R5B | 0.24 | 0.08 | 2.59E-03 | 7.36E-03 |
| ENSG00000126934 | MAP2K2 | 0.24 | 0.08 | 2.67E-03 | 7.58E-03 |
| ENSG00000145476 | CYP4V2 | 0.24 | 0.08 | 2.71E-03 | 7.69E-03 |
| ENSG00000155903 | RASA2 | 0.24 | 0.08 | 2.87E-03 | 8.08E-03 |
| ENSG00000153208 | MERTK | 0.24 | 0.09 | 5.91E-03 | 1.57E-02 |

| | | | | | |
|---|---|---|---|---|---|
| ENSG00000054967 | RELT | 0.24 | 0.09 | 8.07E-03 | 2.08E-02 |
| ENSG00000120832 | MTERF2 | 0.24 | 0.09 | 8.35E-03 | 2.15E-02 |
| ENSG00000068028 | RASSF1 | 0.24 | 0.1 | 9.56E-03 | 2.43E-02 |
| ENSG00000105617 | LENG1 | 0.24 | 0.1 | 1.20E-02 | 2.99E-02 |
| ENSG00000182325 | FBXL6 | 0.24 | 0.1 | 1.27E-02 | 3.14E-02 |
| ENSG00000198185 | ZNF334 | 0.24 | 0.1 | 1.38E-02 | 3.36E-02 |
| ENSG00000164616 | FBXL21P | 0.24 | 0.1 | 1.41E-02 | 3.44E-02 |
| ENSG00000167232 | ZNF91 | 0.24 | 0.11 | 1.69E-02 | 4.05E-02 |
| ENSG00000156381 | ANKRD9 | 0.24 | 0.11 | 2.00E-02 | 4.70E-02 |
| ENSG00000025800 | KPNA6 | 0.23 | 0.03 | 1.02E-12 | 6.68E-12 |
| ENSG00000089220 | PEBP1 | 0.23 | 0.03 | 1.50E-12 | 9.69E-12 |
| ENSG00000107929 | LARP4B | 0.23 | 0.04 | 2.04E-10 | 1.16E-09 |
| ENSG00000101367 | MAPRE1 | 0.23 | 0.04 | 1.44E-08 | 7.12E-08 |
| ENSG00000221983 | UBA52 | 0.23 | 0.04 | 1.66E-08 | 8.18E-08 |
| ENSG00000132475 | H3-3B | 0.23 | 0.04 | 5.90E-08 | 2.79E-07 |
| ENSG00000115839 | RAB3GAP1 | 0.23 | 0.05 | 4.53E-07 | 1.99E-06 |
| ENSG00000176986 | SEC24C | 0.23 | 0.05 | 5.71E-07 | 2.49E-06 |
| ENSG00000162694 | EXTL2 | 0.23 | 0.05 | 5.72E-07 | 2.49E-06 |
| ENSG00000131238 | PPT1 | 0.23 | 0.05 | 1.19E-06 | 5.00E-06 |
| ENSG00000272391 | POM121C | 0.23 | 0.05 | 1.46E-06 | 6.10E-06 |
| ENSG00000165629 | ATP5F1C | 0.23 | 0.05 | 1.63E-06 | 6.80E-06 |
| ENSG00000105854 | PON2 | 0.23 | 0.05 | 1.70E-06 | 7.05E-06 |
| ENSG00000052723 | SIKE1 | 0.23 | 0.05 | 1.74E-06 | 7.21E-06 |
| ENSG00000137574 | TGS1 | 0.23 | 0.05 | 2.97E-06 | 1.21E-05 |
| ENSG00000010810 | FYN | 0.23 | 0.05 | 4.12E-06 | 1.65E-05 |
| ENSG00000062598 | ELMO2 | 0.23 | 0.05 | 4.28E-06 | 1.71E-05 |
| ENSG00000196950 | SLC39A10 | 0.23 | 0.05 | 5.47E-06 | 2.17E-05 |
| ENSG00000114353 | GNAI2 | 0.23 | 0.05 | 1.02E-05 | 3.93E-05 |
| ENSG00000166889 | PATL1 | 0.23 | 0.05 | 1.20E-05 | 4.59E-05 |
| ENSG00000196821 | ILRUN | 0.23 | 0.05 | 1.33E-05 | 5.09E-05 |
| ENSG00000144895 | EIF2A | 0.23 | 0.05 | 1.60E-05 | 6.07E-05 |
| ENSG00000078043 | PIAS2 | 0.23 | 0.05 | 2.07E-05 | 7.75E-05 |
| ENSG00000197265 | GTF2E2 | 0.23 | 0.06 | 6.29E-05 | 2.24E-04 |
| ENSG00000136908 | DPM2 | 0.23 | 0.06 | 6.47E-05 | 2.30E-04 |
| ENSG00000138767 | CNOT6L | 0.23 | 0.06 | 7.26E-05 | 2.57E-04 |
| ENSG00000136280 | CCM2 | 0.23 | 0.06 | 1.04E-04 | 3.61E-04 |
| ENSG00000165355 | FBXO33 | 0.23 | 0.06 | 1.06E-04 | 3.67E-04 |
| ENSG00000131148 | EMC8 | 0.23 | 0.06 | 1.07E-04 | 3.70E-04 |
| ENSG00000025796 | SEC63 | 0.23 | 0.06 | 1.42E-04 | 4.85E-04 |
| ENSG00000135749 | PCNX2 | 0.23 | 0.06 | 1.61E-04 | 5.45E-04 |
| ENSG00000133812 | SBF2 | 0.23 | 0.06 | 1.70E-04 | 5.73E-04 |
| ENSG00000177600 | RPLP2 | 0.23 | 0.06 | 2.20E-04 | 7.34E-04 |
| ENSG00000066422 | ZBTB11 | 0.23 | 0.06 | 2.54E-04 | 8.39E-04 |
| ENSG00000116649 | SRM | 0.23 | 0.06 | 2.86E-04 | 9.38E-04 |
| ENSG00000138660 | AP1AR | 0.23 | 0.06 | 3.24E-04 | 1.06E-03 |
| ENSG00000188070 | ZFTA | 0.23 | 0.07 | 5.15E-04 | 1.64E-03 |
| ENSG00000198586 | TLK1 | 0.23 | 0.07 | 5.27E-04 | 1.67E-03 |
| ENSG00000115694 | STK25 | 0.23 | 0.07 | 5.61E-04 | 1.78E-03 |
| ENSG00000099381 | SETD1A | 0.23 | 0.07 | 7.08E-04 | 2.20E-03 |
| ENSG00000112624 | BICRAL | 0.23 | 0.07 | 7.16E-04 | 2.23E-03 |

| | | | | | |
|---|---|---|---|---|---|
| ENSG00000127452 | FBXL12 | 0.23 | 0.07 | 7.56E-04 | 2.34E-03 |
| ENSG00000196655 | TRAPPC4 | 0.23 | 0.07 | 1.08E-03 | 3.27E-03 |
| ENSG00000156973 | PDE6D | 0.23 | 0.07 | 1.08E-03 | 3.27E-03 |
| ENSG00000175198 | PCCA | 0.23 | 0.07 | 1.19E-03 | 3.58E-03 |
| ENSG00000116213 | WRAP73 | 0.23 | 0.07 | 1.25E-03 | 3.75E-03 |
| ENSG00000169100 | SLC25A6 | 0.23 | 0.07 | 1.34E-03 | 3.99E-03 |
| ENSG00000198840 | MT-ND3 | 0.23 | 0.07 | 1.45E-03 | 4.31E-03 |
| ENSG00000100354 | TNRC6B | 0.23 | 0.07 | 1.50E-03 | 4.44E-03 |
| ENSG00000120458 | MSANTD2 | 0.23 | 0.08 | 2.09E-03 | 6.03E-03 |
| ENSG00000224271 | EPIC1 | 0.23 | 0.08 | 4.07E-03 | 1.11E-02 |
| ENSG00000182141 | ZNF708 | 0.23 | 0.08 | 4.18E-03 | 1.14E-02 |
| ENSG00000167925 | GHDC | 0.23 | 0.08 | 4.27E-03 | 1.16E-02 |
| ENSG00000141741 | MIEN1 | 0.23 | 0.08 | 4.56E-03 | 1.24E-02 |
| ENSG00000140406 | TLNRD1 | 0.23 | 0.08 | 4.61E-03 | 1.25E-02 |
| ENSG00000105737 | GRIK5 | 0.23 | 0.09 | 6.91E-03 | 1.81E-02 |
| ENSG00000140983 | RHOT2 | 0.23 | 0.09 | 7.02E-03 | 1.84E-02 |
| ENSG00000163795 | ZNF513 | 0.23 | 0.09 | 8.00E-03 | 2.07E-02 |
| ENSG00000250510 | GPR162 | 0.23 | 0.09 | 8.07E-03 | 2.08E-02 |
| ENSG00000241468 | ATP5MF | 0.23 | 0.09 | 9.15E-03 | 2.33E-02 |
| ENSG00000242516 | LINC00960 | 0.23 | 0.09 | 1.12E-02 | 2.81E-02 |
| ENSG00000064545 | TMEM161A | 0.23 | 0.09 | 1.16E-02 | 2.90E-02 |
| ENSG00000103061 | SLC7A6OS | 0.23 | 0.1 | 1.25E-02 | 3.08E-02 |
| ENSG00000026036 | RTEL1-TNFRSF6B | 0.23 | 0.1 | 1.41E-02 | 3.43E-02 |
| ENSG00000135317 | SNX14 | 0.23 | 0.1 | 1.50E-02 | 3.63E-02 |
| ENSG00000170775 | GPR37 | 0.23 | 0.1 | 1.76E-02 | 4.21E-02 |
| ENSG00000175701 | MTLN | 0.23 | 0.1 | 1.98E-02 | 4.68E-02 |
| ENSG00000158156 | XKR8 | 0.23 | 0.1 | 2.00E-02 | 4.72E-02 |
| ENSG00000070756 | PABPC1 | 0.22 | 0.03 | 1.45E-12 | 9.40E-12 |
| ENSG00000108561 | C1QBP | 0.22 | 0.03 | 3.25E-10 | 1.82E-09 |
| ENSG00000173473 | SMARCC1 | 0.22 | 0.04 | 4.68E-09 | 2.39E-08 |
| ENSG00000084234 | APLP2 | 0.22 | 0.04 | 3.42E-08 | 1.64E-07 |
| ENSG00000103194 | USP10 | 0.22 | 0.04 | 6.31E-08 | 2.97E-07 |
| ENSG00000166908 | PIP4K2C | 0.22 | 0.04 | 2.13E-07 | 9.65E-07 |
| ENSG00000130803 | ZNF317 | 0.22 | 0.04 | 2.26E-07 | 1.02E-06 |
| ENSG00000129422 | MTUS1 | 0.22 | 0.05 | 6.28E-07 | 2.72E-06 |
| ENSG00000111328 | CDK2AP1 | 0.22 | 0.05 | 8.90E-07 | 3.79E-06 |
| ENSG00000136930 | PSMB7 | 0.22 | 0.05 | 1.13E-06 | 4.78E-06 |
| ENSG00000164587 | RPS14 | 0.22 | 0.04 | 1.15E-06 | 4.86E-06 |
| ENSG00000132676 | DAP3 | 0.22 | 0.05 | 1.49E-06 | 6.21E-06 |
| ENSG00000070610 | GBA2 | 0.22 | 0.05 | 1.70E-06 | 7.05E-06 |
| ENSG00000174106 | LEMD3 | 0.22 | 0.05 | 1.86E-06 | 7.68E-06 |
| ENSG00000125818 | PSMF1 | 0.22 | 0.05 | 2.18E-06 | 8.96E-06 |
| ENSG00000124562 | SNRPC | 0.22 | 0.05 | 3.67E-06 | 1.48E-05 |
| ENSG00000005893 | LAMP2 | 0.22 | 0.05 | 7.76E-06 | 3.03E-05 |
| ENSG00000188895 | MSL1 | 0.22 | 0.05 | 9.50E-06 | 3.68E-05 |
| ENSG00000105401 | CDC37 | 0.22 | 0.05 | 1.31E-05 | 5.00E-05 |
| ENSG00000144659 | SLC25A38 | 0.22 | 0.05 | 1.61E-05 | 6.11E-05 |
| ENSG00000152952 | PLOD2 | 0.22 | 0.05 | 2.22E-05 | 8.31E-05 |
| ENSG00000091640 | SPAG7 | 0.22 | 0.05 | 2.45E-05 | 9.11E-05 |
| ENSG00000196199 | MPHOSPH8 | 0.22 | 0.05 | 2.76E-05 | 1.02E-04 |

| | | | | | |
|---|---|---|---|---|---|
| ENSG00000198642 | KLHL9 | 0.22 | 0.05 | 2.87E-05 | 1.06E-04 |
| ENSG00000130024 | PHF10 | 0.22 | 0.05 | 3.43E-05 | 1.25E-04 |
| ENSG00000142541 | RPL13A | 0.22 | 0.05 | 3.70E-05 | 1.35E-04 |
| ENSG00000125991 | ERGIC3 | 0.22 | 0.06 | 4.85E-05 | 1.75E-04 |
| ENSG00000104067 | TJP1 | 0.22 | 0.06 | 6.56E-05 | 2.34E-04 |
| ENSG00000213523 | SRA1 | 0.22 | 0.06 | 6.84E-05 | 2.43E-04 |
| ENSG00000170185 | USP38 | 0.22 | 0.06 | 6.85E-05 | 2.43E-04 |
| ENSG00000078304 | PPP2R5C | 0.22 | 0.06 | 9.87E-05 | 3.44E-04 |
| ENSG00000162961 | DPY30 | 0.22 | 0.06 | 1.20E-04 | 4.13E-04 |
| ENSG00000103404 | USP31 | 0.22 | 0.06 | 1.44E-04 | 4.91E-04 |
| ENSG00000119522 | DENND1A | 0.22 | 0.06 | 1.66E-04 | 5.63E-04 |
| ENSG00000108469 | RECQL5 | 0.22 | 0.06 | 2.14E-04 | 7.14E-04 |
| ENSG00000140497 | SCAMP2 | 0.22 | 0.06 | 2.31E-04 | 7.67E-04 |
| ENSG00000100023 | PPIL2 | 0.22 | 0.06 | 2.48E-04 | 8.21E-04 |
| ENSG00000129696 | TTI2 | 0.22 | 0.06 | 2.90E-04 | 9.51E-04 |
| ENSG00000125691 | RPL23 | 0.22 | 0.06 | 2.94E-04 | 9.62E-04 |
| ENSG00000284024 | MSANTD7 | 0.22 | 0.06 | 3.82E-04 | 1.23E-03 |
| ENSG00000096093 | EFHC1 | 0.22 | 0.06 | 4.79E-04 | 1.53E-03 |
| ENSG00000126767 | ELK1 | 0.22 | 0.07 | 5.28E-04 | 1.68E-03 |
| ENSG00000124659 | TBCC | 0.22 | 0.07 | 5.72E-04 | 1.81E-03 |
| ENSG00000105700 | KXD1 | 0.22 | 0.07 | 8.10E-04 | 2.50E-03 |
| ENSG00000167257 | RNF214 | 0.22 | 0.07 | 8.19E-04 | 2.52E-03 |
| ENSG00000140474 | ULK3 | 0.22 | 0.07 | 1.09E-03 | 3.31E-03 |
| ENSG00000165775 | FUNDC2 | 0.22 | 0.07 | 1.25E-03 | 3.73E-03 |
| ENSG00000138834 | MAPK8IP3 | 0.22 | 0.07 | 2.27E-03 | 6.51E-03 |
| ENSG00000029639 | TFB1M | 0.22 | 0.07 | 2.45E-03 | 7.00E-03 |
| ENSG00000197136 | PCNX3 | 0.22 | 0.07 | 3.25E-03 | 9.06E-03 |
| ENSG00000122696 | SLC25A51 | 0.22 | 0.08 | 3.74E-03 | 1.03E-02 |
| ENSG00000198055 | GRK6 | 0.22 | 0.08 | 3.81E-03 | 1.05E-02 |
| ENSG00000133316 | WDR74 | 0.22 | 0.08 | 4.33E-03 | 1.18E-02 |
| ENSG00000137193 | PIM1 | 0.22 | 0.08 | 4.55E-03 | 1.23E-02 |
| ENSG00000163131 | CTSS | 0.22 | 0.08 | 6.52E-03 | 1.72E-02 |
| ENSG00000269994 | LINC02893 | 0.22 | 0.09 | 8.15E-03 | 2.10E-02 |
| ENSG00000145912 | NHP2 | 0.22 | 0.09 | 9.21E-03 | 2.35E-02 |
| ENSG00000242086 | MUC20-OT1 | 0.22 | 0.09 | 9.93E-03 | 2.51E-02 |
| ENSG00000184990 | SIVA1 | 0.22 | 0.09 | 1.06E-02 | 2.65E-02 |
| ENSG00000139910 | NOVA1 | 0.22 | 0.09 | 1.11E-02 | 2.78E-02 |
| ENSG00000141858 | SAMD1 | 0.22 | 0.09 | 1.28E-02 | 3.15E-02 |
| ENSG00000152465 | NMT2 | 0.22 | 0.09 | 1.38E-02 | 3.38E-02 |
| ENSG00000175309 | PHYKPL | 0.22 | 0.09 | 1.44E-02 | 3.51E-02 |
| ENSG00000160799 | CCDC12 | 0.22 | 0.09 | 1.49E-02 | 3.62E-02 |
| ENSG00000100258 | LMF2 | 0.22 | 0.1 | 1.86E-02 | 4.41E-02 |
| ENSG00000170876 | TMEM43 | 0.21 | 0.03 | 2.37E-09 | 1.24E-08 |
| ENSG00000116688 | MFN2 | 0.21 | 0.04 | 3.73E-08 | 1.78E-07 |
| ENSG00000156508 | EEF1A1 | 0.21 | 0.04 | 6.96E-08 | 3.27E-07 |
| ENSG00000116580 | GON4L | 0.21 | 0.04 | 1.81E-07 | 8.25E-07 |
| ENSG00000092847 | AGO1 | 0.21 | 0.04 | 2.74E-07 | 1.23E-06 |
| ENSG00000107341 | UBE2R2 | 0.21 | 0.04 | 1.31E-06 | 5.50E-06 |
| ENSG00000099341 | PSMD8 | 0.21 | 0.04 | 2.29E-06 | 9.40E-06 |
| ENSG00000117222 | RBBP5 | 0.21 | 0.05 | 2.62E-06 | 1.07E-05 |

| | | | | | |
|---|---|---|---|---|---|
| ENSG00000065978 | YBX1 | 0.21 | 0.05 | 2.74E-06 | 1.12E-05 |
| ENSG00000086589 | RBM22 | 0.21 | 0.05 | 2.90E-06 | 1.18E-05 |
| ENSG00000213401 | MAGEA12 | 0.21 | 0.05 | 3.54E-06 | 1.43E-05 |
| ENSG00000121390 | PSPC1 | 0.21 | 0.04 | 3.61E-06 | 1.46E-05 |
| ENSG00000064651 | SLC12A2 | 0.21 | 0.05 | 4.20E-06 | 1.69E-05 |
| ENSG00000186432 | KPNA4 | 0.21 | 0.05 | 4.36E-06 | 1.75E-05 |
| ENSG00000140992 | PDPK1 | 0.21 | 0.05 | 5.48E-06 | 2.18E-05 |
| ENSG00000116754 | SRSF11 | 0.21 | 0.05 | 5.94E-06 | 2.35E-05 |
| ENSG00000143418 | CERS2 | 0.21 | 0.05 | 6.80E-06 | 2.67E-05 |
| ENSG00000211460 | TSN | 0.21 | 0.05 | 7.20E-06 | 2.82E-05 |
| ENSG00000140443 | IGF1R | 0.21 | 0.05 | 8.94E-06 | 3.47E-05 |
| ENSG00000029363 | BCLAF1 | 0.21 | 0.05 | 2.48E-05 | 9.20E-05 |
| ENSG00000108848 | LUC7L3 | 0.21 | 0.05 | 3.03E-05 | 1.11E-04 |
| ENSG00000103423 | DNAJA3 | 0.21 | 0.05 | 3.98E-05 | 1.44E-04 |
| ENSG00000198373 | WWP2 | 0.21 | 0.05 | 4.46E-05 | 1.61E-04 |
| ENSG00000184007 | PTP4A2 | 0.21 | 0.05 | 5.49E-05 | 1.97E-04 |
| ENSG00000111596 | CNOT2 | 0.21 | 0.05 | 6.23E-05 | 2.22E-04 |
| ENSG00000151465 | CDC123 | 0.21 | 0.05 | 6.31E-05 | 2.25E-04 |
| ENSG00000163162 | RNF149 | 0.21 | 0.05 | 6.95E-05 | 2.47E-04 |
| ENSG00000117906 | RCN2 | 0.21 | 0.05 | 7.22E-05 | 2.56E-04 |
| ENSG00000131508 | UBE2D2 | 0.21 | 0.05 | 8.40E-05 | 2.94E-04 |
| ENSG00000119953 | SMNDC1 | 0.21 | 0.05 | 8.41E-05 | 2.95E-04 |
| ENSG00000163913 | IFT122 | 0.21 | 0.06 | 1.26E-04 | 4.34E-04 |
| ENSG00000048649 | RSF1 | 0.21 | 0.05 | 1.32E-04 | 4.52E-04 |
| ENSG00000101182 | PSMA7 | 0.21 | 0.06 | 1.35E-04 | 4.62E-04 |
| ENSG00000132478 | UNK | 0.21 | 0.05 | 1.60E-04 | 5.42E-04 |
| ENSG00000114904 | NEK4 | 0.21 | 0.06 | 1.96E-04 | 6.56E-04 |
| ENSG00000103160 | HSDL1 | 0.21 | 0.06 | 2.18E-04 | 7.26E-04 |
| ENSG00000062194 | GPBP1 | 0.21 | 0.06 | 2.23E-04 | 7.43E-04 |
| ENSG00000185344 | ATP6V0A2 | 0.21 | 0.06 | 2.24E-04 | 7.48E-04 |
| ENSG00000183735 | TBK1 | 0.21 | 0.06 | 2.78E-04 | 9.15E-04 |
| ENSG00000168397 | ATG4B | 0.21 | 0.06 | 3.22E-04 | 1.05E-03 |
| ENSG00000196504 | PRPF40A | 0.21 | 0.06 | 3.86E-04 | 1.25E-03 |
| ENSG00000126945 | HNRNPH2 | 0.21 | 0.06 | 4.23E-04 | 1.36E-03 |
| ENSG00000132970 | WASF3 | 0.21 | 0.06 | 4.72E-04 | 1.51E-03 |
| ENSG00000206195 | DUXAP8 | 0.21 | 0.06 | 4.80E-04 | 1.53E-03 |
| ENSG00000065485 | PDIA5 | 0.21 | 0.06 | 5.14E-04 | 1.64E-03 |
| ENSG00000174485 | DENND4A | 0.21 | 0.06 | 6.20E-04 | 1.95E-03 |
| ENSG00000104823 | ECH1 | 0.21 | 0.06 | 6.75E-04 | 2.11E-03 |
| ENSG00000153130 | SCOC | 0.21 | 0.06 | 7.47E-04 | 2.32E-03 |
| ENSG00000115966 | ATF2 | 0.21 | 0.06 | 7.68E-04 | 2.38E-03 |
| ENSG00000153071 | DAB2 | 0.21 | 0.06 | 8.86E-04 | 2.72E-03 |
| ENSG00000214160 | ALG3 | 0.21 | 0.06 | 8.92E-04 | 2.73E-03 |
| ENSG00000153094 | BCL2L11 | 0.21 | 0.07 | 9.85E-04 | 3.00E-03 |
| ENSG00000109762 | SNX25 | 0.21 | 0.07 | 1.08E-03 | 3.27E-03 |
| ENSG00000267575 | LINC02987 | 0.21 | 0.07 | 1.19E-03 | 3.58E-03 |
| ENSG00000107331 | ABCA2 | 0.21 | 0.07 | 1.27E-03 | 3.81E-03 |
| ENSG00000140153 | WDR20 | 0.21 | 0.07 | 1.33E-03 | 3.97E-03 |
| ENSG00000156256 | USP16 | 0.21 | 0.07 | 1.42E-03 | 4.22E-03 |
| ENSG00000011201 | ANOS1 | 0.21 | 0.07 | 1.71E-03 | 5.02E-03 |

| | | | | | |
|---|---|---|---|---|---|
| ENSG00000099282 | TSPAN15 | 0.21 | 0.07 | 2.52E-03 | 7.19E-03 |
| ENSG00000178719 | GRINA | 0.21 | 0.07 | 3.53E-03 | 9.77E-03 |
| ENSG00000164405 | UQCRQ | 0.21 | 0.07 | 3.64E-03 | 1.01E-02 |
| ENSG00000106460 | TMEM106B | 0.21 | 0.08 | 3.81E-03 | 1.05E-02 |
| ENSG00000138032 | PPM1B | 0.21 | 0.07 | 4.46E-03 | 1.21E-02 |
| ENSG00000174132 | FAM174A | 0.21 | 0.08 | 5.67E-03 | 1.51E-02 |
| ENSG00000186625 | KATNA1 | 0.21 | 0.08 | 5.89E-03 | 1.56E-02 |
| ENSG00000115145 | STAM2 | 0.21 | 0.08 | 5.91E-03 | 1.57E-02 |
| ENSG00000128923 | MINDY2 | 0.21 | 0.08 | 6.89E-03 | 1.80E-02 |
| ENSG00000198712 | MT-CO2 | 0.21 | 0.08 | 8.86E-03 | 2.27E-02 |
| ENSG00000171100 | MTM1 | 0.21 | 0.08 | 9.31E-03 | 2.37E-02 |
| ENSG00000251022 | THAP9-AS1 | 0.21 | 0.08 | 9.57E-03 | 2.43E-02 |
| ENSG00000074842 | MYDGF | 0.21 | 0.08 | 1.11E-02 | 2.77E-02 |
| ENSG00000136521 | NDUFB5 | 0.21 | 0.08 | 1.15E-02 | 2.85E-02 |
| ENSG00000125656 | CLPP | 0.21 | 0.09 | 1.22E-02 | 3.03E-02 |
| ENSG00000090006 | LTBP4 | 0.21 | 0.09 | 1.25E-02 | 3.08E-02 |
| ENSG00000130725 | UBE2M | 0.21 | 0.09 | 1.35E-02 | 3.30E-02 |
| ENSG00000225210 | DUXAP9 | 0.21 | 0.09 | 1.44E-02 | 3.50E-02 |
| ENSG00000240857 | RDH14 | 0.21 | 0.09 | 1.44E-02 | 3.51E-02 |
| ENSG00000173451 | THAP2 | 0.21 | 0.09 | 1.60E-02 | 3.85E-02 |
| ENSG00000204172 | AGAP9 | 0.21 | 0.09 | 1.62E-02 | 3.89E-02 |
| ENSG00000188026 | RILPL1 | 0.21 | 0.09 | 1.80E-02 | 4.28E-02 |
| ENSG00000256223 | ZNF10 | 0.21 | 0.09 | 1.82E-02 | 4.33E-02 |
| ENSG00000166441 | RPL27A | 0.2 | 0.03 | 7.01E-10 | 3.83E-09 |
| ENSG00000182934 | SRPRA | 0.2 | 0.03 | 1.59E-09 | 8.45E-09 |
| ENSG00000139641 | ESYT1 | 0.2 | 0.04 | 8.90E-08 | 4.15E-07 |
| ENSG00000105993 | DNAJB6 | 0.2 | 0.04 | 3.03E-07 | 1.35E-06 |
| ENSG00000085733 | CTTN | 0.2 | 0.04 | 3.96E-07 | 1.75E-06 |
| ENSG00000148303 | RPL7A | 0.2 | 0.04 | 3.97E-07 | 1.76E-06 |
| ENSG00000120063 | GNA13 | 0.2 | 0.04 | 1.39E-06 | 5.83E-06 |
| ENSG00000117758 | STX12 | 0.2 | 0.04 | 1.67E-06 | 6.93E-06 |
| ENSG00000131626 | PPFIA1 | 0.2 | 0.04 | 2.95E-06 | 1.20E-05 |
| ENSG00000107164 | FUBP3 | 0.2 | 0.04 | 3.09E-06 | 1.25E-05 |
| ENSG00000143614 | GATAD2B | 0.2 | 0.04 | 4.59E-06 | 1.83E-05 |
| ENSG00000177628 | GBA1 | 0.2 | 0.05 | 6.35E-06 | 2.51E-05 |
| ENSG00000052841 | TTC17 | 0.2 | 0.05 | 1.63E-05 | 6.17E-05 |
| ENSG00000144524 | COPS7B | 0.2 | 0.05 | 2.58E-05 | 9.55E-05 |
| ENSG00000166900 | STX3 | 0.2 | 0.05 | 3.09E-05 | 1.14E-04 |
| ENSG00000047315 | POLR2B | 0.2 | 0.05 | 3.10E-05 | 1.14E-04 |
| ENSG00000175387 | SMAD2 | 0.2 | 0.05 | 3.29E-05 | 1.21E-04 |
| ENSG00000140750 | ARHGAP17 | 0.2 | 0.05 | 3.34E-05 | 1.22E-04 |
| ENSG00000198952 | SMG5 | 0.2 | 0.05 | 3.88E-05 | 1.41E-04 |
| ENSG00000136051 | WASHC4 | 0.2 | 0.05 | 8.75E-05 | 3.06E-04 |
| ENSG00000174099 | MSRB3 | 0.2 | 0.05 | 8.80E-05 | 3.08E-04 |
| ENSG00000073712 | FERMT2 | 0.2 | 0.05 | 1.14E-04 | 3.93E-04 |
| ENSG00000128708 | HAT1 | 0.2 | 0.05 | 1.25E-04 | 4.31E-04 |
| ENSG00000102226 | USP11 | 0.2 | 0.05 | 1.78E-04 | 5.99E-04 |
| ENSG00000114316 | USP4 | 0.2 | 0.05 | 2.03E-04 | 6.80E-04 |
| ENSG00000064763 | FAR2 | 0.2 | 0.06 | 3.08E-04 | 1.01E-03 |
| ENSG00000204351 | SKIC2 | 0.2 | 0.06 | 3.13E-04 | 1.02E-03 |

| | | | | | |
|---|---|---|---|---|---|
| ENSG00000101935 | AMMECR1 | 0.2 | 0.06 | 4.01E-04 | 1.29E-03 |
| ENSG00000117625 | RCOR3 | 0.2 | 0.06 | 4.63E-04 | 1.48E-03 |
| ENSG00000116266 | STXBP3 | 0.2 | 0.06 | 5.45E-04 | 1.73E-03 |
| ENSG00000179933 | C14orf119 | 0.2 | 0.06 | 6.24E-04 | 1.96E-03 |
| ENSG00000055609 | KMT2C | 0.2 | 0.06 | 6.73E-04 | 2.10E-03 |
| ENSG00000135164 | DMTF1 | 0.2 | 0.06 | 6.94E-04 | 2.16E-03 |
| ENSG00000166997 | CNPY4 | 0.2 | 0.06 | 8.06E-04 | 2.49E-03 |
| ENSG00000115421 | PAPOLG | 0.2 | 0.06 | 1.20E-03 | 3.60E-03 |
| ENSG00000033178 | UBA6 | 0.2 | 0.06 | 1.20E-03 | 3.60E-03 |
| ENSG00000100938 | GMPR2 | 0.2 | 0.06 | 1.20E-03 | 3.61E-03 |
| ENSG00000106636 | YKT6 | 0.2 | 0.06 | 1.43E-03 | 4.23E-03 |
| ENSG00000174684 | B4GAT1 | 0.2 | 0.06 | 1.46E-03 | 4.33E-03 |
| ENSG00000136738 | STAM | 0.2 | 0.06 | 1.47E-03 | 4.34E-03 |
| ENSG00000151445 | VIPAS39 | 0.2 | 0.06 | 1.54E-03 | 4.54E-03 |
| ENSG00000197969 | VPS13A | 0.2 | 0.07 | 1.63E-03 | 4.80E-03 |
| ENSG00000129003 | VPS13C | 0.2 | 0.07 | 1.74E-03 | 5.08E-03 |
| ENSG00000189319 | FAM53B | 0.2 | 0.06 | 2.00E-03 | 5.78E-03 |
| ENSG00000013441 | CLK1 | 0.2 | 0.07 | 2.15E-03 | 6.19E-03 |
| ENSG00000213064 | SFT2D2 | 0.2 | 0.07 | 2.35E-03 | 6.73E-03 |
| ENSG00000164144 | ARFIP1 | 0.2 | 0.07 | 2.66E-03 | 7.55E-03 |
| ENSG00000104805 | NUCB1 | 0.2 | 0.07 | 3.21E-03 | 8.97E-03 |
| ENSG00000140987 | ZSCAN32 | 0.2 | 0.07 | 3.42E-03 | 9.51E-03 |
| ENSG00000127418 | FGFRL1 | 0.2 | 0.07 | 3.80E-03 | 1.05E-02 |
| ENSG00000135637 | CCDC142 | 0.2 | 0.07 | 4.07E-03 | 1.11E-02 |
| ENSG00000125835 | SNRPB | 0.2 | 0.07 | 4.29E-03 | 1.17E-02 |
| ENSG00000106397 | PLOD3 | 0.2 | 0.07 | 4.46E-03 | 1.21E-02 |
| ENSG00000099204 | ABLIM1 | 0.2 | 0.07 | 4.73E-03 | 1.28E-02 |
| ENSG00000153339 | TRAPPC8 | 0.2 | 0.07 | 4.87E-03 | 1.31E-02 |
| ENSG00000128159 | TUBGCP6 | 0.2 | 0.07 | 5.70E-03 | 1.52E-02 |
| ENSG00000168528 | SERINC2 | 0.2 | 0.07 | 5.75E-03 | 1.53E-02 |
| ENSG00000196295 | GARS1-DT | 0.2 | 0.07 | 5.81E-03 | 1.54E-02 |
| ENSG00000290418 | TPTEP1 | 0.2 | 0.07 | 5.84E-03 | 1.55E-02 |
| ENSG00000185324 | CDK10 | 0.2 | 0.08 | 6.39E-03 | 1.68E-02 |
| ENSG00000196683 | TOMM7 | 0.2 | 0.07 | 6.50E-03 | 1.71E-02 |
| ENSG00000131165 | CHMP1A | 0.2 | 0.08 | 6.54E-03 | 1.72E-02 |
| ENSG00000125734 | GPR108 | 0.2 | 0.08 | 6.67E-03 | 1.75E-02 |
| ENSG00000291207 | | 0.2 | 0.08 | 7.16E-03 | 1.87E-02 |
| ENSG00000198727 | MT-CYB | 0.2 | 0.08 | 1.43E-02 | 3.49E-02 |
| ENSG00000113916 | BCL6 | 0.2 | 0.08 | 1.54E-02 | 3.72E-02 |
| ENSG00000169592 | INO80E | 0.2 | 0.08 | 1.68E-02 | 4.02E-02 |
| ENSG00000088451 | TGDS | 0.2 | 0.09 | 1.82E-02 | 4.33E-02 |
| ENSG00000162929 | SANBR | 0.2 | 0.09 | 1.85E-02 | 4.39E-02 |
| ENSG00000170270 | GON7 | 0.2 | 0.09 | 1.87E-02 | 4.44E-02 |
| ENSG00000198793 | MTOR | 0.19 | 0.03 | 6.93E-09 | 3.50E-08 |
| ENSG00000089009 | RPL6 | 0.19 | 0.04 | 7.79E-07 | 3.34E-06 |
| ENSG00000171314 | PGAM1 | 0.19 | 0.04 | 1.20E-06 | 5.04E-06 |
| ENSG00000074696 | HACD3 | 0.19 | 0.04 | 1.56E-06 | 6.51E-06 |
| ENSG00000139842 | CUL4A | 0.19 | 0.04 | 2.82E-06 | 1.15E-05 |
| ENSG00000106638 | TBL2 | 0.19 | 0.04 | 1.17E-05 | 4.51E-05 |
| ENSG00000111361 | EIF2B1 | 0.19 | 0.04 | 1.43E-05 | 5.44E-05 |

| | | | | | |
|---|---|---|---|---|---|
| ENSG00000224470 | ATXN1L | 0.19 | 0.05 | 2.45E-05 | 9.09E-05 |
| ENSG00000185219 | ZNF445 | 0.19 | 0.05 | 5.05E-05 | 1.81E-04 |
| ENSG00000137692 | DCUN1D5 | 0.19 | 0.05 | 5.67E-05 | 2.03E-04 |
| ENSG00000069329 | VPS35 | 0.19 | 0.05 | 7.97E-05 | 2.81E-04 |
| ENSG00000120705 | ETF1 | 0.19 | 0.05 | 9.16E-05 | 3.20E-04 |
| ENSG00000168395 | ING5 | 0.19 | 0.05 | 9.50E-05 | 3.31E-04 |
| ENSG00000163904 | SENP2 | 0.19 | 0.05 | 1.07E-04 | 3.70E-04 |
| ENSG00000172661 | WASHC2C | 0.19 | 0.05 | 1.10E-04 | 3.81E-04 |
| ENSG00000255302 | EID1 | 0.19 | 0.05 | 1.22E-04 | 4.20E-04 |
| ENSG00000012174 | MBTPS2 | 0.19 | 0.05 | 1.25E-04 | 4.31E-04 |
| ENSG00000114062 | UBE3A | 0.19 | 0.05 | 1.48E-04 | 5.04E-04 |
| ENSG00000115233 | PSMD14 | 0.19 | 0.05 | 1.55E-04 | 5.26E-04 |
| ENSG00000138614 | INTS14 | 0.19 | 0.05 | 1.66E-04 | 5.60E-04 |
| ENSG00000183020 | AP2A2 | 0.19 | 0.05 | 1.84E-04 | 6.19E-04 |
| ENSG00000182551 | ADI1 | 0.19 | 0.05 | 2.27E-04 | 7.57E-04 |
| ENSG00000147050 | KDM6A | 0.19 | 0.05 | 2.47E-04 | 8.19E-04 |
| ENSG00000197956 | S100A6 | 0.19 | 0.05 | 2.74E-04 | 9.04E-04 |
| ENSG00000100997 | ABHD12 | 0.19 | 0.05 | 3.21E-04 | 1.05E-03 |
| ENSG00000085719 | CPNE3 | 0.19 | 0.05 | 3.61E-04 | 1.17E-03 |
| ENSG00000162231 | NXF1 | 0.19 | 0.05 | 3.79E-04 | 1.22E-03 |
| ENSG00000204634 | TBC1D8 | 0.19 | 0.06 | 4.03E-04 | 1.30E-03 |
| ENSG00000100731 | PCNX1 | 0.19 | 0.05 | 4.22E-04 | 1.36E-03 |
| ENSG00000126003 | PLAGL2 | 0.19 | 0.06 | 6.36E-04 | 1.99E-03 |
| ENSG00000144677 | CTDSPL | 0.19 | 0.06 | 6.93E-04 | 2.16E-03 |
| ENSG00000081791 | DELE1 | 0.19 | 0.06 | 7.47E-04 | 2.32E-03 |
| ENSG00000136279 | DBNL | 0.19 | 0.06 | 8.95E-04 | 2.74E-03 |
| ENSG00000205765 | RIMOC1 | 0.19 | 0.06 | 9.11E-04 | 2.79E-03 |
| ENSG00000136933 | RABEPK | 0.19 | 0.06 | 9.37E-04 | 2.86E-03 |
| ENSG00000118482 | PHF3 | 0.19 | 0.06 | 1.08E-03 | 3.27E-03 |
| ENSG00000140265 | ZSCAN29 | 0.19 | 0.06 | 1.11E-03 | 3.35E-03 |
| ENSG00000183207 | RUVBL2 | 0.19 | 0.06 | 1.51E-03 | 4.46E-03 |
| ENSG00000188641 | DPYD | 0.19 | 0.06 | 1.61E-03 | 4.72E-03 |
| ENSG00000134909 | ARHGAP32 | 0.19 | 0.06 | 1.61E-03 | 4.73E-03 |
| ENSG00000183741 | CBX6 | 0.19 | 0.06 | 2.66E-03 | 7.55E-03 |
| ENSG00000105255 | FSD1 | 0.19 | 0.07 | 2.76E-03 | 7.79E-03 |
| ENSG00000168743 | NPNT | 0.19 | 0.06 | 3.00E-03 | 8.43E-03 |
| ENSG00000058600 | POLR3E | 0.19 | 0.07 | 3.10E-03 | 8.69E-03 |
| ENSG00000196911 | KPNA5 | 0.19 | 0.07 | 3.12E-03 | 8.74E-03 |
| ENSG00000169992 | NLGN2 | 0.19 | 0.06 | 3.15E-03 | 8.80E-03 |
| ENSG00000032444 | PNPLA6 | 0.19 | 0.07 | 3.80E-03 | 1.05E-02 |
| ENSG00000135541 | AHI1 | 0.19 | 0.07 | 3.86E-03 | 1.06E-02 |
| ENSG00000104218 | CSPP1 | 0.19 | 0.07 | 3.88E-03 | 1.07E-02 |
| ENSG00000177556 | ATOX1 | 0.19 | 0.07 | 3.89E-03 | 1.07E-02 |
| ENSG00000253738 | OTUD6B-AS1 | 0.19 | 0.07 | 3.90E-03 | 1.07E-02 |
| ENSG00000198707 | CEP290 | 0.19 | 0.07 | 4.10E-03 | 1.12E-02 |
| ENSG00000129071 | MBD4 | 0.19 | 0.07 | 5.51E-03 | 1.47E-02 |
| ENSG00000124733 | MEA1 | 0.19 | 0.07 | 7.87E-03 | 2.04E-02 |
| ENSG00000159335 | PTMS | 0.19 | 0.07 | 8.16E-03 | 2.10E-02 |
| ENSG00000168904 | LRRC28 | 0.19 | 0.07 | 8.58E-03 | 2.20E-02 |
| ENSG00000105290 | APLP1 | 0.19 | 0.07 | 8.81E-03 | 2.26E-02 |

| | | | | | |
|---|---|---|---|---|---|
| ENSG00000152782 | PANK1 | 0.19 | 0.08 | 9.06E-03 | 2.31E-02 |
| ENSG00000067365 | METTL22 | 0.19 | 0.08 | 1.04E-02 | 2.63E-02 |
| ENSG00000115998 | C2orf42 | 0.19 | 0.08 | 1.27E-02 | 3.14E-02 |
| ENSG00000159433 | STARD9 | 0.19 | 0.08 | 1.43E-02 | 3.48E-02 |
| ENSG00000028839 | TBPL1 | 0.19 | 0.08 | 1.46E-02 | 3.54E-02 |
| ENSG00000177697 | CD151 | 0.19 | 0.08 | 1.50E-02 | 3.64E-02 |
| ENSG00000117318 | ID3 | 0.19 | 0.08 | 1.65E-02 | 3.95E-02 |
| ENSG00000108591 | DRG2 | 0.19 | 0.08 | 1.68E-02 | 4.03E-02 |
| ENSG00000109103 | UNC119 | 0.19 | 0.08 | 1.83E-02 | 4.35E-02 |
| ENSG00000153391 | INO80C | 0.19 | 0.09 | 2.12E-02 | 4.97E-02 |
| ENSG00000135862 | LAMC1 | 0.18 | 0.03 | 3.04E-08 | 1.47E-07 |
| ENSG00000173120 | KDM2A | 0.18 | 0.03 | 4.26E-08 | 2.03E-07 |
| ENSG00000160208 | RRP1B | 0.18 | 0.03 | 9.82E-08 | 4.56E-07 |
| ENSG00000205531 | NAP1L4 | 0.18 | 0.04 | 4.33E-07 | 1.91E-06 |
| ENSG00000166619 | BLCAP | 0.18 | 0.04 | 4.39E-07 | 1.93E-06 |
| ENSG00000176871 | WSB2 | 0.18 | 0.04 | 1.71E-06 | 7.11E-06 |
| ENSG00000168036 | CTNNB1 | 0.18 | 0.04 | 3.80E-06 | 1.53E-05 |
| ENSG00000185728 | YTHDF3 | 0.18 | 0.04 | 5.15E-06 | 2.05E-05 |
| ENSG00000166147 | FBN1 | 0.18 | 0.04 | 1.21E-05 | 4.64E-05 |
| ENSG00000180543 | TSPYL5 | 0.18 | 0.04 | 1.25E-05 | 4.80E-05 |
| ENSG00000134371 | CDC73 | 0.18 | 0.04 | 3.26E-05 | 1.19E-04 |
| ENSG00000065802 | ASB1 | 0.18 | 0.05 | 4.77E-05 | 1.72E-04 |
| ENSG00000099995 | SF3A1 | 0.18 | 0.05 | 4.85E-05 | 1.75E-04 |
| ENSG00000138802 | SEC24B | 0.18 | 0.05 | 6.25E-05 | 2.23E-04 |
| ENSG00000170162 | VGLL2 | 0.18 | 0.05 | 6.31E-05 | 2.25E-04 |
| ENSG00000134444 | RELCH | 0.18 | 0.05 | 7.07E-05 | 2.50E-04 |
| ENSG00000139970 | RTN1 | 0.18 | 0.05 | 7.16E-05 | 2.53E-04 |
| ENSG00000106635 | BCL7B | 0.18 | 0.04 | 7.52E-05 | 2.66E-04 |
| ENSG00000134779 | TPGS2 | 0.18 | 0.04 | 8.86E-05 | 3.10E-04 |
| ENSG00000213619 | NDUFS3 | 0.18 | 0.05 | 1.22E-04 | 4.20E-04 |
| ENSG00000144713 | RPL32 | 0.18 | 0.05 | 1.36E-04 | 4.65E-04 |
| ENSG00000162889 | MAPKAPK2 | 0.18 | 0.05 | 1.54E-04 | 5.24E-04 |
| ENSG00000090686 | USP48 | 0.18 | 0.05 | 1.97E-04 | 6.62E-04 |
| ENSG00000159792 | PSKH1 | 0.18 | 0.05 | 2.04E-04 | 6.83E-04 |
| ENSG00000175224 | ATG13 | 0.18 | 0.05 | 2.13E-04 | 7.11E-04 |
| ENSG00000169756 | LIMS1 | 0.18 | 0.05 | 2.16E-04 | 7.22E-04 |
| ENSG00000117450 | PRDX1 | 0.18 | 0.05 | 2.91E-04 | 9.56E-04 |
| ENSG00000125633 | CCDC93 | 0.18 | 0.05 | 5.30E-04 | 1.68E-03 |
| ENSG00000120820 | GLT8D2 | 0.18 | 0.05 | 6.27E-04 | 1.97E-03 |
| ENSG00000130724 | CHMP2A | 0.18 | 0.05 | 6.77E-04 | 2.11E-03 |
| ENSG00000107984 | DKK1 | 0.18 | 0.05 | 7.03E-04 | 2.19E-03 |
| ENSG00000133884 | DPF2 | 0.18 | 0.05 | 7.06E-04 | 2.20E-03 |
| ENSG00000042286 | AIFM2 | 0.18 | 0.05 | 7.29E-04 | 2.26E-03 |
| ENSG00000139112 | GABARAPL1 | 0.18 | 0.06 | 8.41E-04 | 2.59E-03 |
| ENSG00000065548 | ZC3H15 | 0.18 | 0.05 | 9.89E-04 | 3.01E-03 |
| ENSG00000183475 | ASB7 | 0.18 | 0.06 | 1.14E-03 | 3.42E-03 |
| ENSG00000136243 | NUP42 | 0.18 | 0.05 | 1.14E-03 | 3.43E-03 |
| ENSG00000137207 | YIPF3 | 0.18 | 0.06 | 1.15E-03 | 3.47E-03 |
| ENSG00000164220 | F2RL2 | 0.18 | 0.06 | 1.38E-03 | 4.11E-03 |
| ENSG00000090372 | STRN4 | 0.18 | 0.06 | 1.39E-03 | 4.14E-03 |

| | | | | | |
|---|---|---|---|---|---|
| ENSG00000169018 | FEM1B | 0.18 | 0.06 | 1.45E-03 | 4.29E-03 |
| ENSG00000083099 | LYRM2 | 0.18 | 0.06 | 1.57E-03 | 4.63E-03 |
| ENSG00000163389 | POGLUT1 | 0.18 | 0.06 | 2.08E-03 | 6.01E-03 |
| ENSG00000169967 | MAP3K2 | 0.18 | 0.06 | 2.33E-03 | 6.67E-03 |
| ENSG00000180879 | SSR4 | 0.18 | 0.06 | 3.06E-03 | 8.57E-03 |
| ENSG00000113742 | CPEB4 | 0.18 | 0.06 | 3.11E-03 | 8.72E-03 |
| ENSG00000159164 | SV2A | 0.18 | 0.06 | 3.26E-03 | 9.08E-03 |
| ENSG00000165392 | WRN | 0.18 | 0.06 | 3.94E-03 | 1.08E-02 |
| ENSG00000159692 | CTBP1 | 0.18 | 0.06 | 5.07E-03 | 1.36E-02 |
| ENSG00000168291 | PDHB | 0.18 | 0.07 | 5.14E-03 | 1.38E-02 |
| ENSG00000155016 | CYP2U1 | 0.18 | 0.07 | 5.20E-03 | 1.39E-02 |
| ENSG00000164332 | UBLCP1 | 0.18 | 0.07 | 5.22E-03 | 1.40E-02 |
| ENSG00000108604 | SMARCD2 | 0.18 | 0.07 | 5.26E-03 | 1.41E-02 |
| ENSG00000183155 | RABIF | 0.18 | 0.07 | 5.87E-03 | 1.56E-02 |
| ENSG00000078124 | ACER3 | 0.18 | 0.07 | 6.17E-03 | 1.63E-02 |
| ENSG00000136783 | NIPSNAP3A | 0.18 | 0.07 | 6.58E-03 | 1.73E-02 |
| ENSG00000117707 | PROX1 | 0.18 | 0.07 | 6.72E-03 | 1.76E-02 |
| ENSG00000108669 | CYTH1 | 0.18 | 0.07 | 7.28E-03 | 1.90E-02 |
| ENSG00000158019 | BABAM2 | 0.18 | 0.07 | 7.62E-03 | 1.98E-02 |
| ENSG00000139116 | KIF21A | 0.18 | 0.07 | 7.69E-03 | 1.99E-02 |
| ENSG00000076924 | XAB2 | 0.18 | 0.07 | 7.95E-03 | 2.06E-02 |
| ENSG00000163093 | BBS5 | 0.18 | 0.07 | 8.69E-03 | 2.23E-02 |
| ENSG00000132406 | TMEM128 | 0.18 | 0.07 | 8.94E-03 | 2.28E-02 |
| ENSG00000136935 | GOLGA1 | 0.18 | 0.07 | 9.86E-03 | 2.50E-02 |
| ENSG00000184277 | TM2D3 | 0.18 | 0.07 | 1.02E-02 | 2.56E-02 |
| ENSG00000070269 | TMEM260 | 0.18 | 0.07 | 1.03E-02 | 2.59E-02 |
| ENSG00000096092 | TMEM14A | 0.18 | 0.07 | 1.14E-02 | 2.85E-02 |
| ENSG00000105698 | USF2 | 0.18 | 0.07 | 1.29E-02 | 3.18E-02 |
| ENSG00000139505 | MTMR6 | 0.18 | 0.08 | 1.32E-02 | 3.25E-02 |
| ENSG00000117533 | VAMP4 | 0.18 | 0.07 | 1.35E-02 | 3.30E-02 |
| ENSG00000176022 | B3GALT6 | 0.18 | 0.08 | 1.40E-02 | 3.41E-02 |
| ENSG00000166478 | ZNF143 | 0.18 | 0.08 | 1.51E-02 | 3.66E-02 |
| ENSG00000131748 | STARD3 | 0.18 | 0.08 | 1.52E-02 | 3.69E-02 |
| ENSG00000132801 | ZSWIM3 | 0.18 | 0.08 | 2.13E-02 | 4.99E-02 |
| ENSG00000146072 | TNFRSF21 | 0.17 | 0.04 | 6.30E-06 | 2.49E-05 |
| ENSG00000175029 | CTBP2 | 0.17 | 0.04 | 1.11E-05 | 4.27E-05 |
| ENSG00000144043 | TEX261 | 0.17 | 0.04 | 1.33E-05 | 5.10E-05 |
| ENSG00000139289 | PHLDA1 | 0.17 | 0.04 | 1.36E-05 | 5.18E-05 |
| ENSG00000102158 | MAGT1 | 0.17 | 0.04 | 2.12E-05 | 7.95E-05 |
| ENSG00000101856 | PGRMC1 | 0.17 | 0.04 | 2.35E-05 | 8.74E-05 |
| ENSG00000101608 | MYL12A | 0.17 | 0.04 | 8.26E-05 | 2.90E-04 |
| ENSG00000186566 | GPATCH8 | 0.17 | 0.04 | 8.54E-05 | 2.99E-04 |
| ENSG00000136715 | SAP130 | 0.17 | 0.04 | 9.18E-05 | 3.21E-04 |
| ENSG00000221914 | PPP2R2A | 0.17 | 0.04 | 9.30E-05 | 3.24E-04 |
| ENSG00000206053 | JPT2 | 0.17 | 0.04 | 1.09E-04 | 3.77E-04 |
| ENSG00000158195 | WASF2 | 0.17 | 0.04 | 1.21E-04 | 4.16E-04 |
| ENSG00000090060 | PAPOLA | 0.17 | 0.05 | 1.23E-04 | 4.23E-04 |
| ENSG00000138757 | G3BP2 | 0.17 | 0.05 | 2.96E-04 | 9.70E-04 |
| ENSG00000113013 | HSPA9 | 0.17 | 0.05 | 3.11E-04 | 1.02E-03 |
| ENSG00000105186 | ANKRD27 | 0.17 | 0.05 | 3.32E-04 | 1.08E-03 |

| | | | | | |
|---|---|---|---|---|---|
| ENSG00000264522 | OTUD7B | 0.17 | 0.05 | 3.70E-04 | 1.20E-03 |
| ENSG00000121064 | SCPEP1 | 0.17 | 0.05 | 4.23E-04 | 1.36E-03 |
| ENSG00000110435 | PDHX | 0.17 | 0.05 | 4.90E-04 | 1.56E-03 |
| ENSG00000121073 | SLC35B1 | 0.17 | 0.05 | 4.98E-04 | 1.59E-03 |
| ENSG00000085063 | CD59 | 0.17 | 0.05 | 5.37E-04 | 1.70E-03 |
| ENSG00000140575 | IQGAP1 | 0.17 | 0.05 | 5.57E-04 | 1.76E-03 |
| ENSG00000106144 | CASP2 | 0.17 | 0.05 | 6.05E-04 | 1.90E-03 |
| ENSG00000178741 | COX5A | 0.17 | 0.05 | 7.17E-04 | 2.23E-03 |
| ENSG00000112276 | BVES | 0.17 | 0.05 | 1.21E-03 | 3.62E-03 |
| ENSG00000120509 | PDZD11 | 0.17 | 0.05 | 1.25E-03 | 3.73E-03 |
| ENSG00000112695 | COX7A2 | 0.17 | 0.05 | 1.43E-03 | 4.24E-03 |
| ENSG00000174282 | ZBTB4 | 0.17 | 0.05 | 1.49E-03 | 4.39E-03 |
| ENSG00000113269 | RNF130 | 0.17 | 0.05 | 1.54E-03 | 4.54E-03 |
| ENSG00000179562 | GCC1 | 0.17 | 0.06 | 1.65E-03 | 4.84E-03 |
| ENSG00000182150 | ERCC6L2 | 0.17 | 0.06 | 1.88E-03 | 5.47E-03 |
| ENSG00000136754 | ABI1 | 0.17 | 0.06 | 1.97E-03 | 5.72E-03 |
| ENSG00000103064 | SLC7A6 | 0.17 | 0.06 | 2.05E-03 | 5.92E-03 |
| ENSG00000172046 | USP19 | 0.17 | 0.06 | 2.23E-03 | 6.40E-03 |
| ENSG00000198722 | UNC13B | 0.17 | 0.06 | 2.74E-03 | 7.74E-03 |
| ENSG00000204227 | RING1 | 0.17 | 0.06 | 2.80E-03 | 7.91E-03 |
| ENSG00000119900 | OGFRL1 | 0.17 | 0.06 | 2.85E-03 | 8.04E-03 |
| ENSG00000109065 | NAT9 | 0.17 | 0.06 | 2.90E-03 | 8.15E-03 |
| ENSG00000105671 | DDX49 | 0.17 | 0.06 | 3.18E-03 | 8.88E-03 |
| ENSG00000136631 | VPS45 | 0.17 | 0.06 | 3.24E-03 | 9.04E-03 |
| ENSG00000221867 | MAGEA3 | 0.17 | 0.06 | 3.86E-03 | 1.06E-02 |
| ENSG00000120333 | MRPS14 | 0.17 | 0.06 | 4.38E-03 | 1.19E-02 |
| ENSG00000236287 | ZBED5 | 0.17 | 0.06 | 4.41E-03 | 1.20E-02 |
| ENSG00000168394 | TAP1 | 0.17 | 0.06 | 4.53E-03 | 1.23E-02 |
| ENSG00000135845 | PIGC | 0.17 | 0.06 | 4.53E-03 | 1.23E-02 |
| ENSG00000075275 | CELSR1 | 0.17 | 0.06 | 4.70E-03 | 1.27E-02 |
| ENSG00000103037 | SETD6 | 0.17 | 0.06 | 4.99E-03 | 1.34E-02 |
| ENSG00000104812 | GYS1 | 0.17 | 0.06 | 5.03E-03 | 1.35E-02 |
| ENSG00000168522 | FNTA | 0.17 | 0.06 | 5.85E-03 | 1.55E-02 |
| ENSG00000232119 | MCTS1 | 0.17 | 0.06 | 6.00E-03 | 1.59E-02 |
| ENSG00000166881 | NEMP1 | 0.17 | 0.06 | 6.44E-03 | 1.69E-02 |
| ENSG00000114023 | FAM162A | 0.17 | 0.06 | 7.10E-03 | 1.85E-02 |
| ENSG00000164073 | MFSD8 | 0.17 | 0.06 | 7.18E-03 | 1.87E-02 |
| ENSG00000269893 | SNHG8 | 0.17 | 0.06 | 7.49E-03 | 1.95E-02 |
| ENSG00000099331 | MYO9B | 0.17 | 0.07 | 7.50E-03 | 1.95E-02 |
| ENSG00000140307 | GTF2A2 | 0.17 | 0.07 | 7.94E-03 | 2.05E-02 |
| ENSG00000008952 | SEC62 | 0.17 | 0.07 | 9.15E-03 | 2.33E-02 |
| ENSG00000171262 | FAM98B | 0.17 | 0.07 | 9.19E-03 | 2.34E-02 |
| ENSG00000131171 | SH3BGRL | 0.17 | 0.07 | 9.83E-03 | 2.49E-02 |
| ENSG00000140092 | FBLN5 | 0.17 | 0.07 | 1.11E-02 | 2.78E-02 |
| ENSG00000006704 | GTF2IRD1 | 0.17 | 0.07 | 1.14E-02 | 2.84E-02 |
| ENSG00000176386 | CDC26 | 0.17 | 0.07 | 1.14E-02 | 2.84E-02 |
| ENSG00000197150 | ABCB8 | 0.17 | 0.07 | 1.15E-02 | 2.87E-02 |
| ENSG00000139625 | MAP3K12 | 0.17 | 0.07 | 1.20E-02 | 2.98E-02 |
| ENSG00000057935 | MTA3 | 0.17 | 0.07 | 1.21E-02 | 3.00E-02 |
| ENSG00000139546 | TARBP2 | 0.17 | 0.07 | 1.22E-02 | 3.02E-02 |

| | | | | | |
|---|---|---|---|---|---|
| ENSG00000177030 | DEAF1 | 0.17 | 0.07 | 1.27E-02 | 3.14E-02 |
| ENSG00000171861 | MRM3 | 0.17 | 0.07 | 1.33E-02 | 3.27E-02 |
| ENSG00000153885 | KCTD15 | 0.17 | 0.07 | 1.34E-02 | 3.28E-02 |
| ENSG00000213753 | CENPBD2P | 0.17 | 0.07 | 1.47E-02 | 3.57E-02 |
| ENSG00000197603 | CPLANE1 | 0.17 | 0.07 | 1.54E-02 | 3.72E-02 |
| ENSG00000011009 | LYPLA2 | 0.17 | 0.07 | 1.56E-02 | 3.76E-02 |
| ENSG00000131446 | MGAT1 | 0.17 | 0.07 | 1.63E-02 | 3.91E-02 |
| ENSG00000007384 | RHBDF1 | 0.17 | 0.07 | 1.66E-02 | 3.97E-02 |
| ENSG00000118276 | B4GALT6 | 0.17 | 0.07 | 1.69E-02 | 4.04E-02 |
| ENSG00000179271 | GADD45GIP1 | 0.17 | 0.07 | 1.76E-02 | 4.20E-02 |
| ENSG00000176473 | WDR25 | 0.17 | 0.07 | 1.77E-02 | 4.21E-02 |
| ENSG00000285756 | | 0.17 | 0.07 | 1.91E-02 | 4.53E-02 |
| ENSG00000156873 | PHKG2 | 0.17 | 0.08 | 2.04E-02 | 4.80E-02 |
| ENSG00000110917 | MLEC | 0.16 | 0.03 | 1.21E-07 | 5.59E-07 |
| ENSG00000137154 | RPS6 | 0.16 | 0.03 | 1.67E-07 | 7.62E-07 |
| ENSG00000140943 | MBTPS1 | 0.16 | 0.03 | 3.48E-07 | 1.54E-06 |
| ENSG00000165637 | VDAC2 | 0.16 | 0.03 | 4.26E-07 | 1.88E-06 |
| ENSG00000148672 | GLUD1 | 0.16 | 0.04 | 6.65E-05 | 2.37E-04 |
| ENSG00000132305 | IMMT | 0.16 | 0.04 | 7.78E-05 | 2.74E-04 |
| ENSG00000011007 | ELOA | 0.16 | 0.04 | 1.00E-04 | 3.49E-04 |
| ENSG00000160075 | SSU72 | 0.16 | 0.04 | 1.57E-04 | 5.32E-04 |
| ENSG00000178913 | TAF7 | 0.16 | 0.04 | 1.66E-04 | 5.62E-04 |
| ENSG00000111640 | GAPDH | 0.16 | 0.04 | 2.06E-04 | 6.88E-04 |
| ENSG00000118855 | MFSD1 | 0.16 | 0.04 | 2.21E-04 | 7.36E-04 |
| ENSG00000118181 | RPS25 | 0.16 | 0.04 | 3.17E-04 | 1.03E-03 |
| ENSG00000162851 | TFB2M | 0.16 | 0.05 | 5.95E-04 | 1.87E-03 |
| ENSG00000162852 | CNST | 0.16 | 0.05 | 6.45E-04 | 2.02E-03 |
| ENSG00000167112 | TRUB2 | 0.16 | 0.05 | 6.88E-04 | 2.15E-03 |
| ENSG00000114416 | FXR1 | 0.16 | 0.05 | 7.05E-04 | 2.20E-03 |
| ENSG00000142751 | GPN2 | 0.16 | 0.05 | 7.05E-04 | 2.20E-03 |
| ENSG00000149480 | MTA2 | 0.16 | 0.05 | 7.72E-04 | 2.39E-03 |
| ENSG00000157985 | AGAP1 | 0.16 | 0.05 | 8.16E-04 | 2.52E-03 |
| ENSG00000278311 | GGNBP2 | 0.16 | 0.05 | 1.38E-03 | 4.10E-03 |
| ENSG00000083520 | DIS3 | 0.16 | 0.05 | 1.44E-03 | 4.25E-03 |
| ENSG00000125875 | TBC1D20 | 0.16 | 0.05 | 1.55E-03 | 4.57E-03 |
| ENSG00000204160 | ZDHHC18 | 0.16 | 0.05 | 1.55E-03 | 4.57E-03 |
| ENSG00000141279 | NPEPPS | 0.16 | 0.05 | 1.67E-03 | 4.91E-03 |
| ENSG00000166340 | TPP1 | 0.16 | 0.05 | 1.73E-03 | 5.07E-03 |
| ENSG00000173064 | HECTD4 | 0.16 | 0.05 | 1.76E-03 | 5.16E-03 |
| ENSG00000143771 | CNIH4 | 0.16 | 0.05 | 2.04E-03 | 5.90E-03 |
| ENSG00000123684 | LPGAT1 | 0.16 | 0.05 | 2.46E-03 | 7.02E-03 |
| ENSG00000163625 | WDFY3 | 0.16 | 0.05 | 2.79E-03 | 7.89E-03 |
| ENSG00000107185 | RGP1 | 0.16 | 0.05 | 2.86E-03 | 8.05E-03 |
| ENSG00000166068 | SPRED1 | 0.16 | 0.05 | 3.07E-03 | 8.62E-03 |
| ENSG00000099290 | WASHC2A | 0.16 | 0.05 | 3.27E-03 | 9.11E-03 |
| ENSG00000170871 | KIAA0232 | 0.16 | 0.05 | 3.46E-03 | 9.60E-03 |
| ENSG00000156374 | PCGF6 | 0.16 | 0.06 | 4.05E-03 | 1.11E-02 |
| ENSG00000124459 | ZNF45 | 0.16 | 0.06 | 4.16E-03 | 1.14E-02 |
| ENSG00000100883 | SRP54 | 0.16 | 0.06 | 4.85E-03 | 1.31E-02 |
| ENSG00000100897 | DCAF11 | 0.16 | 0.06 | 5.69E-03 | 1.51E-02 |

| | | | | | |
|---|---|---|---|---|---|
| ENSG00000163848 | ZNF148 | 0.16 | 0.06 | 6.11E-03 | 1.62E-02 |
| ENSG00000176953 | NFATC2IP | 0.16 | 0.06 | 7.79E-03 | 2.02E-02 |
| ENSG00000106290 | TAF6 | 0.16 | 0.06 | 9.46E-03 | 2.40E-02 |
| ENSG00000173960 | UBXN2A | 0.16 | 0.07 | 1.30E-02 | 3.20E-02 |
| ENSG00000204525 | HLA-C | 0.16 | 0.07 | 1.36E-02 | 3.34E-02 |
| ENSG00000277586 | NEFL | 0.16 | 0.06 | 1.40E-02 | 3.41E-02 |
| ENSG00000105429 | MEGF8 | 0.16 | 0.06 | 1.44E-02 | 3.49E-02 |
| ENSG00000089289 | IGBP1 | 0.16 | 0.07 | 1.56E-02 | 3.77E-02 |
| ENSG00000127993 | RBM48 | 0.16 | 0.07 | 1.61E-02 | 3.88E-02 |
| ENSG00000117419 | ERI3 | 0.16 | 0.07 | 1.64E-02 | 3.92E-02 |
| ENSG00000120370 | GORAB | 0.16 | 0.07 | 1.89E-02 | 4.49E-02 |
| ENSG00000241553 | ARPC4 | 0.16 | 0.07 | 2.03E-02 | 4.78E-02 |
| ENSG00000142002 | DPP9 | 0.16 | 0.07 | 2.07E-02 | 4.87E-02 |
| ENSG00000290770 | MFSD14CP | 0.16 | 0.07 | 2.13E-02 | 4.99E-02 |
| ENSG00000038382 | TRIO | 0.15 | 0.03 | 3.64E-06 | 1.47E-05 |
| ENSG00000100811 | YY1 | 0.15 | 0.03 | 2.01E-05 | 7.54E-05 |
| ENSG00000109111 | SUPT6H | 0.15 | 0.04 | 5.44E-05 | 1.95E-04 |
| ENSG00000104723 | TUSC3 | 0.15 | 0.04 | 5.55E-05 | 1.99E-04 |
| ENSG00000165280 | VCP | 0.15 | 0.04 | 2.19E-04 | 7.31E-04 |
| ENSG00000164329 | TENT2 | 0.15 | 0.04 | 2.73E-04 | 8.99E-04 |
| ENSG00000102898 | NUTF2 | 0.15 | 0.04 | 3.55E-04 | 1.15E-03 |
| ENSG00000113716 | HMGXB3 | 0.15 | 0.04 | 3.57E-04 | 1.16E-03 |
| ENSG00000182473 | EXOC7 | 0.15 | 0.04 | 3.87E-04 | 1.25E-03 |
| ENSG00000108953 | YWHAE | 0.15 | 0.05 | 7.78E-04 | 2.41E-03 |
| ENSG00000090530 | P3H2 | 0.15 | 0.05 | 7.82E-04 | 2.42E-03 |
| ENSG00000068745 | IP6K2 | 0.15 | 0.05 | 8.11E-04 | 2.50E-03 |
| ENSG00000153201 | RANBP2 | 0.15 | 0.04 | 8.45E-04 | 2.60E-03 |
| ENSG00000089157 | RPLP0 | 0.15 | 0.05 | 1.05E-03 | 3.18E-03 |
| ENSG00000163110 | PDLIM5 | 0.15 | 0.05 | 1.35E-03 | 4.03E-03 |
| ENSG00000198818 | SFT2D1 | 0.15 | 0.05 | 1.51E-03 | 4.45E-03 |
| ENSG00000085788 | DDHD2 | 0.15 | 0.05 | 1.66E-03 | 4.86E-03 |
| ENSG00000146648 | EGFR | 0.15 | 0.05 | 3.07E-03 | 8.62E-03 |
| ENSG00000117280 | RAB29 | 0.15 | 0.05 | 3.33E-03 | 9.28E-03 |
| ENSG00000011260 | UTP18 | 0.15 | 0.05 | 3.38E-03 | 9.39E-03 |
| ENSG00000149273 | RPS3 | 0.15 | 0.05 | 3.70E-03 | 1.02E-02 |
| ENSG00000128989 | ARPP19 | 0.15 | 0.05 | 3.72E-03 | 1.03E-02 |
| ENSG00000198612 | COPS8 | 0.15 | 0.05 | 4.57E-03 | 1.24E-02 |
| ENSG00000143748 | NVL | 0.15 | 0.05 | 5.06E-03 | 1.36E-02 |
| ENSG00000198860 | TSEN15 | 0.15 | 0.05 | 5.13E-03 | 1.38E-02 |
| ENSG00000072736 | NFATC3 | 0.15 | 0.05 | 5.43E-03 | 1.45E-02 |
| ENSG00000143815 | LBR | 0.15 | 0.06 | 5.90E-03 | 1.57E-02 |
| ENSG00000029725 | RABEP1 | 0.15 | 0.05 | 6.01E-03 | 1.59E-02 |
| ENSG00000240849 | PEDS1 | 0.15 | 0.06 | 6.92E-03 | 1.81E-02 |
| ENSG00000197535 | MYO5A | 0.15 | 0.05 | 7.05E-03 | 1.84E-02 |
| ENSG00000167468 | GPX4 | 0.15 | 0.06 | 7.27E-03 | 1.90E-02 |
| ENSG00000168246 | UBTD2 | 0.15 | 0.06 | 7.36E-03 | 1.92E-02 |
| ENSG00000164919 | COX6C | 0.15 | 0.06 | 8.12E-03 | 2.09E-02 |
| ENSG00000144036 | EXOC6B | 0.15 | 0.06 | 9.55E-03 | 2.42E-02 |
| ENSG00000249464 | LINC01091 | 0.15 | 0.06 | 1.01E-02 | 2.55E-02 |
| ENSG00000166012 | TAF1D | 0.15 | 0.06 | 1.04E-02 | 2.62E-02 |

| | | | | | |
|---|---|---|---|---|---|
| ENSG00000158417 | EIF5B | 0.15 | 0.06 | 1.31E-02 | 3.22E-02 |
| ENSG00000133398 | MED10 | 0.15 | 0.06 | 1.32E-02 | 3.25E-02 |
| ENSG00000102804 | TSC22D1 | 0.15 | 0.06 | 1.33E-02 | 3.26E-02 |
| ENSG00000100196 | KDELR3 | 0.15 | 0.06 | 1.52E-02 | 3.67E-02 |
| ENSG00000159082 | SYNJ1 | 0.15 | 0.06 | 1.58E-02 | 3.81E-02 |
| ENSG00000148300 | REXO4 | 0.15 | 0.06 | 1.58E-02 | 3.82E-02 |
| ENSG00000165175 | MID1IP1 | 0.15 | 0.06 | 1.60E-02 | 3.86E-02 |
| ENSG00000078142 | PIK3C3 | 0.15 | 0.07 | 1.81E-02 | 4.29E-02 |
| ENSG00000183255 | PTTG1IP | 0.14 | 0.04 | 8.23E-05 | 2.89E-04 |
| ENSG00000065883 | CDK13 | 0.14 | 0.04 | 2.09E-04 | 7.00E-04 |
| ENSG00000100335 | MIEF1 | 0.14 | 0.04 | 3.36E-04 | 1.09E-03 |
| ENSG00000135720 | DYNC1LI2 | 0.14 | 0.04 | 5.50E-04 | 1.74E-03 |
| ENSG00000185904 | LINC00839 | 0.14 | 0.04 | 7.97E-04 | 2.46E-03 |
| ENSG00000005100 | DHX33 | 0.14 | 0.04 | 8.25E-04 | 2.54E-03 |
| ENSG00000197102 | DYNC1H1 | 0.14 | 0.04 | 9.05E-04 | 2.77E-03 |
| ENSG00000189043 | NDUFA4 | 0.14 | 0.04 | 9.53E-04 | 2.91E-03 |
| ENSG00000142168 | SOD1 | 0.14 | 0.04 | 1.11E-03 | 3.34E-03 |
| ENSG00000106571 | GLI3 | 0.14 | 0.04 | 1.15E-03 | 3.46E-03 |
| ENSG00000197157 | SND1 | 0.14 | 0.04 | 1.29E-03 | 3.84E-03 |
| ENSG00000171067 | C11orf24 | 0.14 | 0.04 | 1.39E-03 | 4.14E-03 |
| ENSG00000166887 | VPS39 | 0.14 | 0.04 | 1.44E-03 | 4.27E-03 |
| ENSG00000122203 | KIAA1191 | 0.14 | 0.04 | 1.51E-03 | 4.45E-03 |
| ENSG00000101266 | CSNK2A1 | 0.14 | 0.05 | 1.70E-03 | 4.98E-03 |
| ENSG00000110958 | PTGES3 | 0.14 | 0.05 | 2.22E-03 | 6.38E-03 |
| ENSG00000133835 | HSD17B4 | 0.14 | 0.05 | 2.47E-03 | 7.06E-03 |
| ENSG00000136938 | ANP32B | 0.14 | 0.05 | 2.53E-03 | 7.21E-03 |
| ENSG00000121892 | PDS5A | 0.14 | 0.05 | 2.54E-03 | 7.25E-03 |
| ENSG00000112335 | SNX3 | 0.14 | 0.05 | 2.98E-03 | 8.38E-03 |
| ENSG00000156113 | KCNMA1 | 0.14 | 0.05 | 3.16E-03 | 8.84E-03 |
| ENSG00000083896 | YTHDC1 | 0.14 | 0.05 | 3.91E-03 | 1.07E-02 |
| ENSG00000152270 | PDE3B | 0.14 | 0.05 | 4.67E-03 | 1.26E-02 |
| ENSG00000115825 | PRKD3 | 0.14 | 0.05 | 6.63E-03 | 1.74E-02 |
| ENSG00000071082 | RPL31 | 0.14 | 0.05 | 7.65E-03 | 1.99E-02 |
| ENSG00000155508 | CNOT8 | 0.14 | 0.05 | 8.99E-03 | 2.30E-02 |
| ENSG00000155876 | RRAGA | 0.14 | 0.06 | 9.41E-03 | 2.39E-02 |
| ENSG00000179119 | SPTY2D1 | 0.14 | 0.05 | 9.43E-03 | 2.40E-02 |
| ENSG00000131263 | RLIM | 0.14 | 0.05 | 9.51E-03 | 2.41E-02 |
| ENSG00000116350 | SRSF4 | 0.14 | 0.05 | 9.59E-03 | 2.43E-02 |
| ENSG00000006715 | VPS41 | 0.14 | 0.05 | 1.12E-02 | 2.80E-02 |
| ENSG00000141644 | MBD1 | 0.14 | 0.06 | 1.13E-02 | 2.83E-02 |
| ENSG00000162702 | ZNF281 | 0.14 | 0.06 | 1.33E-02 | 3.26E-02 |
| ENSG00000150995 | ITPR1 | 0.14 | 0.06 | 1.35E-02 | 3.31E-02 |
| ENSG00000185404 | SP140L | 0.14 | 0.06 | 1.40E-02 | 3.41E-02 |
| ENSG00000162738 | VANGL2 | 0.14 | 0.06 | 1.43E-02 | 3.49E-02 |
| ENSG00000101871 | MID1 | 0.14 | 0.06 | 1.53E-02 | 3.69E-02 |
| ENSG00000150776 | NKAPD1 | 0.14 | 0.06 | 1.57E-02 | 3.78E-02 |
| ENSG00000160932 | LY6E | 0.14 | 0.06 | 2.10E-02 | 4.92E-02 |
| ENSG00000158669 | GPAT4 | 0.13 | 0.04 | 3.46E-04 | 1.12E-03 |
| ENSG00000180398 | MCFD2 | 0.13 | 0.04 | 5.61E-04 | 1.77E-03 |
| ENSG00000164466 | SFXN1 | 0.13 | 0.04 | 9.19E-04 | 2.81E-03 |

| | | | | | |
|---|---|---|---|---|---|
| ENSG00000069248 | NUP133 | 0.13 | 0.04 | 1.02E-03 | 3.11E-03 |
| ENSG00000156515 | HK1 | 0.13 | 0.04 | 1.35E-03 | 4.03E-03 |
| ENSG00000084623 | EIF3I | 0.13 | 0.04 | 1.60E-03 | 4.70E-03 |
| ENSG00000071127 | WDR1 | 0.13 | 0.04 | 1.79E-03 | 5.22E-03 |
| ENSG00000101310 | SEC23B | 0.13 | 0.04 | 1.98E-03 | 5.74E-03 |
| ENSG00000198876 | DCAF12 | 0.13 | 0.04 | 2.35E-03 | 6.73E-03 |
| ENSG00000166526 | ZNF3 | 0.13 | 0.04 | 4.09E-03 | 1.12E-02 |
| ENSG00000006468 | ETV1 | 0.13 | 0.05 | 6.06E-03 | 1.60E-02 |
| ENSG00000135870 | RC3H1 | 0.13 | 0.05 | 6.10E-03 | 1.61E-02 |
| ENSG00000164674 | SYTL3 | 0.13 | 0.05 | 6.76E-03 | 1.77E-02 |
| ENSG00000154144 | TBRG1 | 0.13 | 0.05 | 7.20E-03 | 1.88E-02 |
| ENSG00000004897 | CDC27 | 0.13 | 0.05 | 7.68E-03 | 1.99E-02 |
| ENSG00000143514 | TP53BP2 | 0.13 | 0.05 | 1.14E-02 | 2.85E-02 |
| ENSG00000136273 | HUS1 | 0.13 | 0.05 | 1.33E-02 | 3.26E-02 |
| ENSG00000181666 | ZNF875 | 0.13 | 0.05 | 1.35E-02 | 3.31E-02 |
| ENSG00000135249 | RINT1 | 0.13 | 0.06 | 1.66E-02 | 3.99E-02 |
| ENSG00000120533 | ENY2 | 0.13 | 0.05 | 1.90E-02 | 4.51E-02 |
| ENSG00000110075 | PPP6R3 | 0.13 | 0.06 | 2.00E-02 | 4.71E-02 |
| ENSG00000187109 | NAP1L1 | 0.13 | 0.06 | 2.03E-02 | 4.78E-02 |
| ENSG00000130764 | LRRC47 | 0.13 | 0.06 | 2.06E-02 | 4.84E-02 |
| ENSG00000149948 | HMGA2 | 0.12 | 0.03 | 5.66E-04 | 1.79E-03 |
| ENSG00000090863 | GLG1 | 0.12 | 0.03 | 5.96E-04 | 1.88E-03 |
| ENSG00000198752 | CDC42BPB | 0.12 | 0.04 | 6.76E-04 | 2.11E-03 |
| ENSG00000143398 | PIP5K1A | 0.12 | 0.04 | 7.61E-04 | 2.36E-03 |
| ENSG00000176915 | ANKLE2 | 0.12 | 0.04 | 1.44E-03 | 4.26E-03 |
| ENSG00000005339 | CREBBP | 0.12 | 0.04 | 1.85E-03 | 5.39E-03 |
| ENSG00000117592 | PRDX6 | 0.12 | 0.04 | 1.94E-03 | 5.64E-03 |
| ENSG00000124198 | ARFGEF2 | 0.12 | 0.04 | 2.30E-03 | 6.60E-03 |
| ENSG00000139292 | LGR5 | 0.12 | 0.04 | 3.63E-03 | 1.00E-02 |
| ENSG00000273749 | CYFIP1 | 0.12 | 0.04 | 4.04E-03 | 1.11E-02 |
| ENSG00000112977 | DAP | 0.12 | 0.04 | 5.13E-03 | 1.38E-02 |
| ENSG00000197713 | RPE | 0.12 | 0.05 | 5.86E-03 | 1.56E-02 |
| ENSG00000145715 | RASA1 | 0.12 | 0.04 | 6.16E-03 | 1.63E-02 |
| ENSG00000168385 | SEPTIN2 | 0.12 | 0.05 | 6.74E-03 | 1.77E-02 |
| ENSG00000214078 | CPNE1 | 0.12 | 0.05 | 8.52E-03 | 2.19E-02 |
| ENSG00000111639 | MRPL51 | 0.12 | 0.05 | 8.62E-03 | 2.21E-02 |
| ENSG00000122085 | MTERF4 | 0.12 | 0.05 | 8.93E-03 | 2.28E-02 |
| ENSG00000091129 | NRCAM | 0.12 | 0.05 | 1.01E-02 | 2.55E-02 |
| ENSG00000140612 | SEC11A | 0.12 | 0.05 | 1.02E-02 | 2.57E-02 |
| ENSG00000147065 | MSN | 0.12 | 0.05 | 1.09E-02 | 2.73E-02 |
| ENSG00000169398 | PTK2 | 0.12 | 0.05 | 1.10E-02 | 2.76E-02 |
| ENSG00000134717 | BTF3L4 | 0.12 | 0.05 | 1.18E-02 | 2.93E-02 |
| ENSG00000146147 | MLIP | 0.12 | 0.05 | 1.29E-02 | 3.19E-02 |
| ENSG00000102921 | N4BP1 | 0.12 | 0.05 | 1.39E-02 | 3.39E-02 |
| ENSG00000225921 | NOL7 | 0.12 | 0.05 | 1.49E-02 | 3.62E-02 |
| ENSG00000124155 | PIGT | 0.12 | 0.05 | 1.62E-02 | 3.89E-02 |
| ENSG00000118263 | KLF7 | 0.12 | 0.05 | 1.64E-02 | 3.92E-02 |
| ENSG00000146701 | MDH2 | 0.12 | 0.05 | 1.66E-02 | 3.99E-02 |
| ENSG00000099954 | CECR2 | 0.12 | 0.05 | 1.68E-02 | 4.02E-02 |
| ENSG00000101782 | RIOK3 | 0.12 | 0.05 | 1.75E-02 | 4.17E-02 |

| | | | | | |
|---|---|---|---|---|---|
| ENSG00000174748 | RPL15 | 0.11 | 0.04 | 1.60E-03 | 4.71E-03 |
| ENSG00000147457 | CHMP7 | 0.11 | 0.04 | 2.75E-03 | 7.77E-03 |
| ENSG00000185164 | NOMO2 | 0.11 | 0.04 | 3.98E-03 | 1.09E-02 |
| ENSG00000164032 | H2AZ1 | 0.11 | 0.04 | 5.99E-03 | 1.59E-02 |
| ENSG00000162734 | PEA15 | 0.11 | 0.04 | 6.30E-03 | 1.66E-02 |
| ENSG00000213719 | CLIC1 | 0.11 | 0.04 | 6.52E-03 | 1.71E-02 |
| ENSG00000055483 | USP36 | 0.11 | 0.04 | 8.26E-03 | 2.13E-02 |
| ENSG00000139146 | SINHCAF | 0.11 | 0.04 | 1.29E-02 | 3.17E-02 |
| ENSG00000077235 | GTF3C1 | 0.11 | 0.05 | 1.62E-02 | 3.89E-02 |
| ENSG00000136003 | ISCU | 0.11 | 0.05 | 1.67E-02 | 4.01E-02 |
| ENSG00000170776 | AKAP13 | 0.11 | 0.05 | 1.72E-02 | 4.11E-02 |
| ENSG00000092148 | HECTD1 | 0.11 | 0.05 | 1.74E-02 | 4.16E-02 |
| ENSG00000103018 | CYB5B | 0.11 | 0.05 | 1.86E-02 | 4.42E-02 |
| ENSG00000140391 | TSPAN3 | 0.1 | 0.03 | 3.07E-03 | 8.62E-03 |
| ENSG00000169439 | SDC2 | 0.1 | 0.04 | 4.16E-03 | 1.14E-02 |
| ENSG00000059573 | ALDH18A1 | 0.1 | 0.04 | 6.88E-03 | 1.80E-02 |
| ENSG00000125743 | SNRPD2 | 0.1 | 0.04 | 7.30E-03 | 1.90E-02 |
| ENSG00000102265 | TIMP1 | 0.1 | 0.04 | 7.66E-03 | 1.99E-02 |
| ENSG00000276293 | PIP4K2B | 0.1 | 0.04 | 8.20E-03 | 2.11E-02 |
| ENSG00000108091 | CCDC6 | 0.1 | 0.04 | 1.21E-02 | 3.00E-02 |
| ENSG00000134247 | PTGFRN | 0.1 | 0.04 | 1.94E-02 | 4.58E-02 |
| ENSG00000173726 | TOMM20 | 0.1 | 0.04 | 1.94E-02 | 4.60E-02 |
| ENSG00000101193 | GID8 | 0.09 | 0.03 | 8.19E-03 | 2.11E-02 |
| ENSG00000111530 | CAND1 | 0.09 | 0.04 | 1.35E-02 | 3.32E-02 |
| ENSG00000075415 | SLC25A3 | 0.09 | 0.04 | 2.08E-02 | 4.89E-02 |
| ENSG00000134644 | PUM1 | 0.08 | 0.03 | 1.98E-02 | 4.67E-02 |

**Table S3. Human primers used for RT-qPCR analysis.**

| | **PRIMER (5'-3')** |
|---|---|
| DHCR7 forward | TCCACAGCCATGTGACCAATGC |
| DHCR7 reverse | CGAAGTGGTCATGGCAGATGTC |
| HMGCR forward | ACTTCGTGTTCATGACTTTC |
| HMGCR reverse | GACATAATCATCTTGACCCTC |
| PERK forward | GTCCCAAGGCTTTGGAATCTGTC |
| PERK reverse | CCTACCAAGACAGGAGTTCTGG |
| ATF4 forward | TTCTCCAGCGACAAGGCTAAGG |
| ATF4 reverse | CTCCAACATCCAATCTGTCCCG |
| CHOP forward | GGTATGAGGACCTGCAAGAGGT |
| CHOP reverse | CTTGTGACCTCTGCTGGTTCTG |
| GADD34 forward | TCCGACTGCAAAGGCGGCTCA |
| GADD34 reverse | CAGCCAGGAAATGGACAGTGAC |
| PROX1 forward | CTGAAGACCTACTTCTCCGACG |
| PROX1 reverse | GATGGCTTGACGTGCGTACTTC |
| HPRT forward | CTCATGGACTGATTATGGACAGGAC |
| HPRT reverse | GCAGGTCAGCAAAGAACTTATAGCC |
